# Genetically stable kill-switch using “demon and angel” expression construct of essential genes

**DOI:** 10.1101/2023.11.12.566782

**Authors:** Yusuke Kato, Hirotada Mori

## Abstract

Genetic instability of synthetic genetic devices is a key obstacle for practical use. This problem is particularly critical in kill-switches for conditional host killing. Here, we propose a genetically stable kill-switch based on a “demon and angel” expression construct of a toxic essential gene. The kill-switch conditionally overexpresses the toxic essential gene. Additionally, the identical essential gene is deleted in the genome. The essential gene is expressed at a low level to maintain host survival in the OFF state and kills the host by the overexpression in the ON state. The single expression construct is responsible for both killing the hosts and maintaining viability, reducing the emergence of loss-of-function mutants. We constructed the kill-switch using the toxic essential gene encoding tyrosyl-tRNA synthetase, *tyrS*, in *Escherichia coli*. The bacteria harboring the kill-switch were conditionally suicidal over 300 generations. Toxic overexpression of essential genes has also been found in other organisms, suggesting that the “demon and angel” kill switch is scalable to various organisms.

**GRAPHICAL ABSTRACT:** 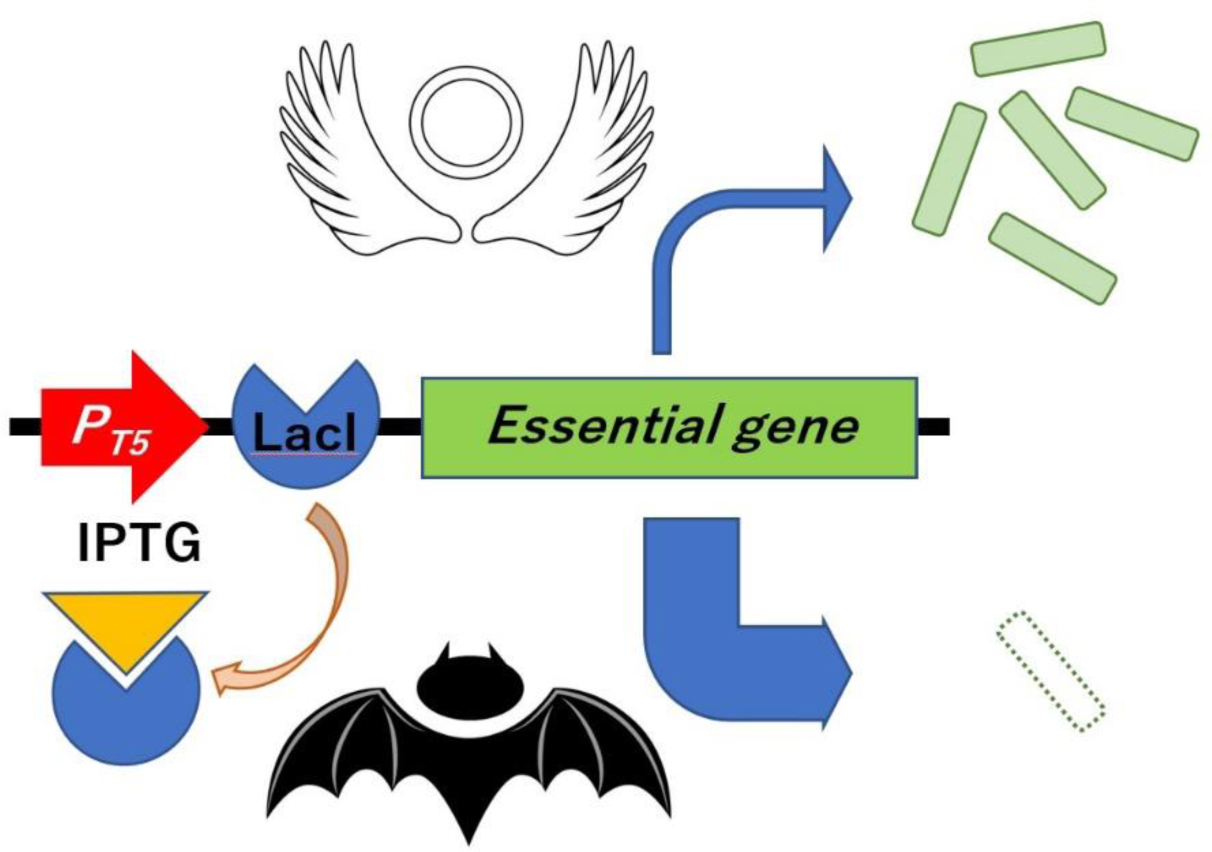

## INTRODUCTION

Synthetic gene circuits for bioengineering purposes are usually not essential for host survival, but rather a burden [1–3]. Therefore, the synthetic gene circuits are genetically unstable since loss-of-function mutants have a higher fitness. This problem is a critical obstacle to the practical application of synthetic biology. Kill-switch is a genetic device that conditionally kills host cells [4]. The kill-switch is used as a core component in biological containment and density control systems [5–7]. The application of genetically engineered microorganisms with biological containment in the open environments is considered revolutionary in the bioindustry [8–10]. In addition, highly efficient microbial production with precise growth control, particularly in multi-cell systems, is also a useful application [11,12]. Because the kill-switches contain toxic genes, they harm the cells even in the OFF-state by leakage toxic expression and are therefore particularly prone to genetic instability, allowing loss-of-function mutants to emerge and predominate through passaging [13]. To construct a useful kill-switch, it is essential to resolve the genetic instability.

To date, several methods have been proposed to reduce the genetic instability of kill-switches. Strategies fall into two major categories. One is to reduce the frequency of mutations that cause loss-of-function. Methods include removal of transposable elements and IS elements [14–16], deletion of mutagenic genes [17], exclusion of repetitive sequences [18] and hosting of reduced genome cells [6]. Maintenance using toxin-antitoxin systems [19], mobile plasmids [20] and integration into the genome [21] have been effective in preventing the loss of kill-switches encoded on plasmids. Removal of mutants by periodic renewal of the cell population is also a promising solution [22]. The other is eliminating the fitness advantage of loss-of-function mutants. Minimizing unintended toxicity in the OFF state mitigates the reduced fitness of cells harboring the kill-switch [13,23]. Linking the functional expression of essential genes to kill-switches can reduce the fitness of mutants [24,25].

In the bacterium *Escherichia coli*, we cloned the all ORFs of protein-coding genes in the ASKA plasmid library [26]. The cloned ORF can be overexpressed in the presence of IPTG by the coliphage T5 promoter with *lacO* operators [27]. Interestingly, overexpression of the ORFs was often toxic to the host [26]. These toxic ORFs include many essential genes.

Here, we propose a novel principle to construct a genetically stable kill-switch using a “demon and angel” essential gene expression construct (Figure 1). To construct the kill-switch, a plasmid conditionally overexpressing the toxic essential gene is transformed into the host. The ASKA plasmids meet this requirement. Meanwhile, the identical essential gene is disrupted in the genome. Therefore, the resulting bacteria must supply the essential gene product from the ASKA plasmid. The overexpression is suppressed in the absence of IPTG (OFF state), but a small amount of product is produced due to the leakage expression [28], maintaining the bacteria in survival and growth. However, in the presence of IPTG (ON state), the essential gene is overexpressed and kills the host bacteria. In this kill-switch, a single expression construct provides not only the host-killing toxin but also the essential gene product for survival, predicting that loss-of-function mutations are largely prevented. In addition, leakage expression of toxins, which is a cause of genetic instability in conventional kill switches, is not a problem because such low-level expression is essential for survival, not poisonous. In this study, we demonstrate the principle of the “demon and angel” kill-switch using the ASKA plasmids in *E. coli*.

**Figure 1.**
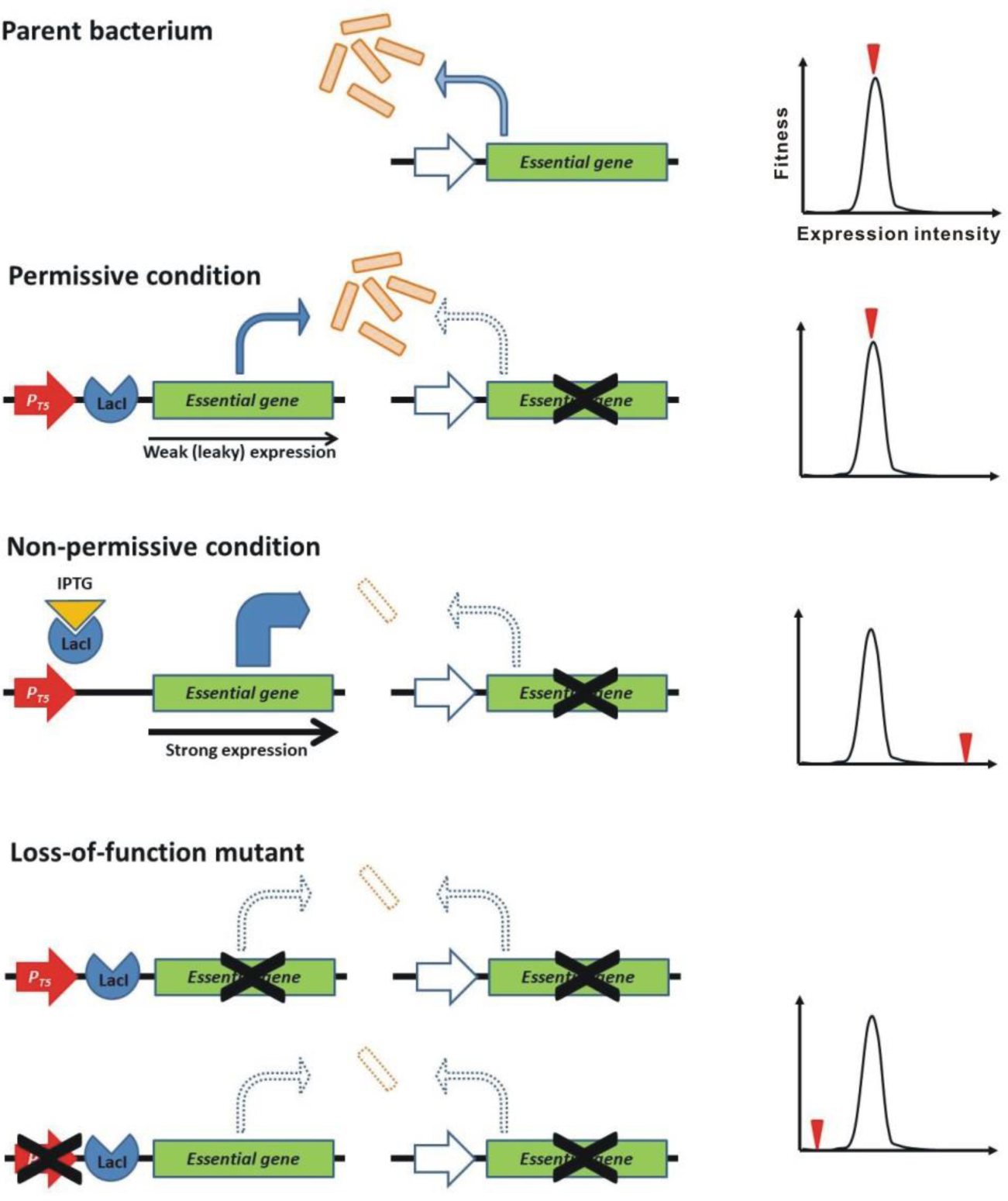
Kill-switch using the “demon and angel” expression construct of an essential gene. A schematic diagram is shown. Yellow squares represent bacterial cells. *P_T5_*, the coliphage T5 promoter. The genes shown on the right and left are the original essential gene in the genome and the synthetic overexpression construct in the plasmid, respectively.

## MATERIAL AND METHODS

### Bacterial strains

*E. coli* K-12, strain AG1 [*recA1 endA1 gyrA96 thi-1 hsdR17(rKmKþ) supE44 relA1*] harboring ASKA+ and ASKA-plasmid libraries, W3110, and W3110 ts 29-13 were obtained from the National BioResource Project (NIG, Japan): *E. coli* (https://shigen.nig.ac.jp/ecoli/strain/). BL21-AI [F^-^*omp*T *hsd*S_B_(r ^-^ m ^-^) *gal dcm ara*B::T7RNAP-*tet*A] was purchased commercially (ThermoFisher Scientific, MA, USA).

### Bacterial culture and transformation

LB medium (1% BactoTriptone, 0.5% Bacto Yeast Extract, 1% NaCl) was used to culture all *E. coli* strains. Antibiotics (chloramphenicol 50 mg/L, kanamycin 25 mg/L, and tetracycline 1.25 mg/L) were added as needed. Liquid culture was performed in an air incubator with shaking (180 rpm). To prepare solid media, 2% agar was added. Bacteria were incubated at 37°C unless otherwise noted. Bacterial transformation was performed by electroporation using a Gene Pulser II^TM^ electroporator (Bio-Rad Laboratories, CA, USA).

### Evaluation for toxic overexpression

The procedure for preliminary toxicity evaluation for all ORFs using ASKA+ plasmid library has been described previously [26]. For quantitative assays, overnight cultures of *E. coli* harboring the ASKA-plasmid encoding the tested ORFs were used. The cultures diluted 10 to 10^3^-fold were inoculated 250 μl onto solid medium containing 1 mM IPTG and chloramphenicol, a selection marker for ASKA-, and surviving colonies were counted. The total number of bacteria was determined by inoculating 10^6^-fold diluted cultures onto a solid medium without IPTG. The toxicity due to overexpression was assessed by the survivor frequency (survivors/total CFU).

### Preparation of BL21-AI [*tyrS^-^*; ASKA-(*tyrS*)]

The ASKA-clone encoding *tyrS* was isolated from AG1 carrying this plasmid using the Qiagen Plasmid Midi (Qiagen Inc., CA, USA) and transformed into BL21-AI. The *tyrS* in the genome of the resulting BL21-AI [*tyrS^+^*; ASKA-(*tyrS*)] strain was substituted with a knock-out cassette containing the neomycin resistance gene using λRed recombinase [29]. Among individual colonies resistant to both chloramphenicol and kanamycin, clones with successful substitution were identified by PCR using the genomic DNA as a template. Finally, the deletion of *tyrS* was directly confirmed by sequencing the *tyrS* locus using the Illumina sequencing, as detailed below.

### Passage test

Plasmid loss was tested for BL21-AI [*tyrS^-^*; ASKA-(*tyrS*)] and the control parent strain, BL21-AI [*tyrS^+^*; ASKA-(*tyrS*)]. Every 24 hours, the bacteria were diluted 10^3^-fold (1 μL/1 mL fresh LB medium without selection marker antibiotics) and incubated again at 25°C. This dilution and reculturing was repeated 12 times, approximately 120 generations. On inoculation into a new medium, an aliquot of the culture was withdrawn, diluted 10^6^-fold, and inoculated (250 μL) onto solid medium. Sixty-four colonies were selected and inoculated on solid medium containing chloramphenicol to determine plasmid maintenance.

To verify genetic stability, bacteria were cultured in the presence of chloramphenicol and kanamycin. Dilution and reculturing procedures were identical to those used in the plasmid loss test. On inoculation for reculturing, an aliquot of the cultures was withdrawn, diluted 10-fold, and inoculated (250 μL) onto solid medium containing 5 mM IPTG. Surviving colonies were counted after overnight incubation. The total number of bacteria was also measured by inoculating 10^6^-fold diluted cultures on IPTG-free solid medium.

### Genome sequencing

The procedure for genome sequencing has been described previously [30]. Briefly, total DNA, including genomic and plasmid DNAs, was extracted using the DNeasy Blood & Tissue Kit (Qiagen). After checking the integrity of the extracted DNA, sequencing libraries were prepared using a TruSeq DNA PCR-free Sample Preparation Kit (Illumina, Inc., San Diego, CA, USA). The purified libraries were subjected to paired-end (2 × 150 bp) sequencing by Macrogen using the NovaSeq6000 platform. Raw reads were trimmed to remove adapter sequences, mapped to the *E. coli* BL21-AI (NZ_CP047231.1) and ASKA-(*tyrS*) reference sequence using BWA-MEM v0.7.17. Variants (SNPs and small INDELs) were identified using Samtools. Variant functional annotation was performed with SnpEff v.5.0e (https://pcingola.github.io/SnpEff/).

### RNA-seq

Bacterial samples were homogenized using a Multi-beads Shocker^®^ (Yasuikikai, Osaka, Japan) at 1,500 rpm for 2 minutes. Total RNA was extracted using the RNeasy Mini Kit and the QIAcube Connect Extraction System (Qiagen). The concentration of the total RNA samples was determined using the Synergy LX multimode reader (BioTek, VT, USA) and the QuantiFluor RNA System (Promega, WI, USA). The RNA quality was evaluated using the 5200 Fragment Analyzer System and the Agilent HS RNA Kit (Agilent Technologies, Inc., CA, USA). After the ribosomal RNA depletion with the RiboPOOL kit (siTOOLs Biotech GmbH, Planegg, Germany), the cDNA libraries were prepared using the MGIEasy RNA Directional Library Prep Kit (MGI-Tech, Shenzhen, China). The concentration of libraries was determined as described above. A library quality check was performed using the dsDNA 915 reagent kit (Agilent Technologies) and the Fragment Analyzer. Circularized DNA and DNA nanoballs were prepared using the MGIEasy circularization kit and the DNBSEQ-G400RS high-throughput sequencing kit (MGI-Tech). The DNBs were sequenced using the DNBSEQ-G400 (MGI-Tech). The cutadapt (ver. 4.0) and sickle tools (ver. 1.33) were used to remove adapter sequences and low-quality sequence reads. The resulting reads were mapped using Bowtie 2 (ver. 2.5.0) to the *E. coli* BL21-AI genome (CP047231.1) and the plasmid ASKA-(*tyrS*). The mapped reads were quantified with FeatureCounts (ver.2.0.3). The row counts were normalized using the transcripts-per-million (TPM) method. The expression level was evaluated relative to *gltA*, a constitutively expressed gene [31].

## RESULTS

### Screening of “toxic” essential genes

The ASKA plasmid is a multi-copy plasmid containing the colE1 origin [26]. There are two types of ASKA plasmid: ASKA+ in which *gfp* is fused at the 3’ terminal end of the ORF, and ASKA-which excludes the *gfp*. We examined growth and GFP expression of *E. coli* AG1 harboring ASKA+ plasmids in overexpression induction with 1 mM IPTG (Table S1). Total 4269 ORFs were tested. Growth inhibition was observed in 3301 ORFs (77%), of which 2149 (50%) were severely inhibited with almost no growth, as previously reported [26] (Figure 2). Of the ORFs that severely inhibited growth, 156 were essential genes. This represents 52% of all 300 essential genes. In addition, 112 of the highly growth inhibitory essential genes expressed little or no GFP. Thus, there are more than 10^2^ essential genes that are strongly harmful upon overexpression.

**Figure 2.**
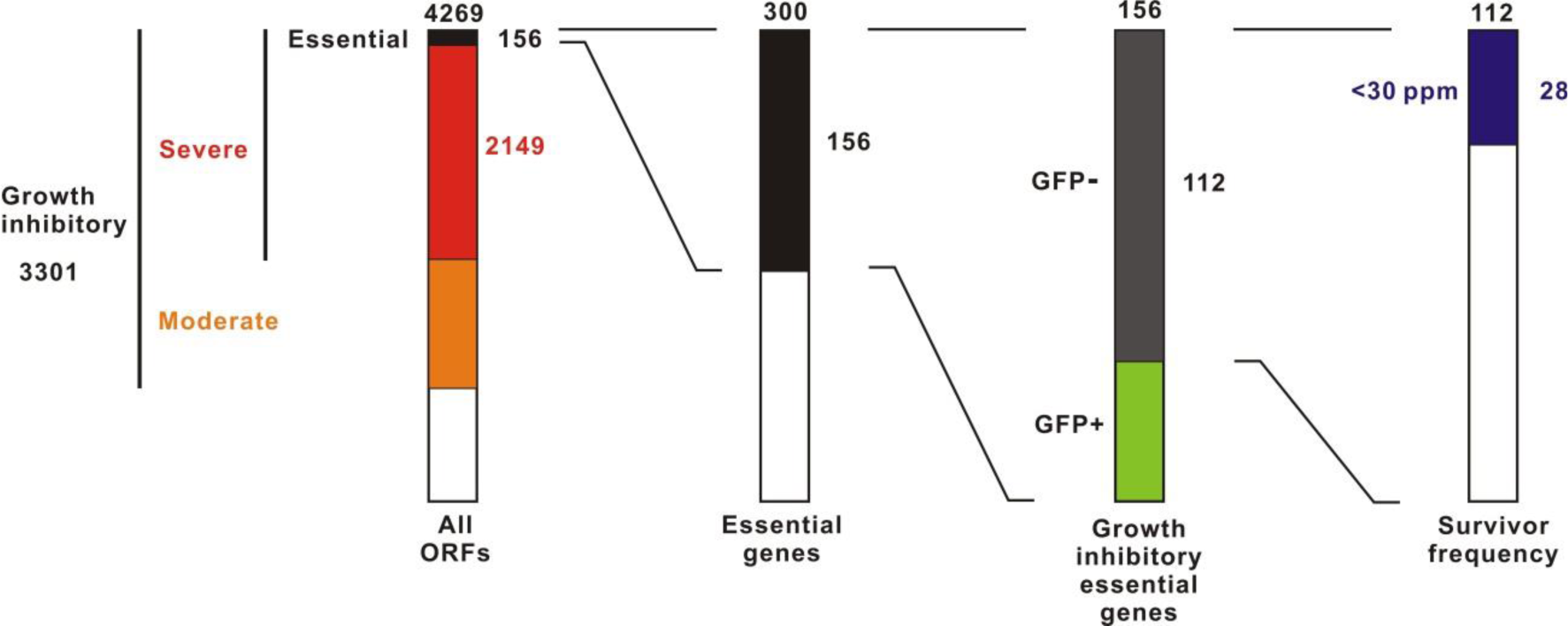
Screening for toxic essential genes in the overexpression. The ASKA+ plasmid library containing 4269 *E. coli* W3110 ORFs was screened. Left column is cited from ref.27 with some modifications. Survivor frequencies were evaluated using the ASKA-plasmid library to avoid the effect of GFP-tag. The value indicated at the top of each column is the total number.

For the 112 essential genes that exhibited both strong growth inhibition and suppression of GFP expression, we evaluated the survivor frequency upon overexpression (Table S2). To avoid the effects of *gfp*-tag, the ASKA-plasmids were used. The survivor frequency showed significant variability. For example, with *dnaX* and *tilS*, no colonies were detected in the presence of 1 mM IPTG after inoculating around 10^7^ CFU (Table S2, Figure S2). In contrast, some survivors were detected in other ORFs such as *murG* and *ftsE*. There were 28 (25%) ORFs that showed a survival frequency of <30 ppm. Of these, 16 were cytoplasmic proteins, 9 were integral membrane proteins, and 3 were membrane anchored proteins (Table S2).

### Realization of the “demon and angel” kill-switch

To realize the “demon and angel” kill-switch, we examined which essential gene would be suitable as a suitable model. The kill-switch must provide a small amount of functional product by leakage expression in the OFF state to maintain host viability. Membrane proteins are often difficult to express functionally and injure the host [32]. Therefore, we decided to select a candidate essential gene among genes encoding cytoplasmic proteins. Aminoacyl-tRNA synthetase (aaRS) genes were focussed. Total 26 aaRS genes, including 22 essential genes, were found in the ASKA+ library. Of the 24 genes tested, 23 showed toxicity upon overexpression (Table S3). Nine genes, *pheT, leuS, tyrS, ileS, tilS, proS, argS, glyS* and *poxA*, were highly growth inhibitory with no GFP expression. *tyrS* is the essential gene encoding tyrosyl-tRNA synthetase [33]. The survivor frequency of AG1[ASKA-(*tyrS*)] upon overexpression with 1 mM IPTG was only 22.5 ppm, indicating potent toxicity (Table S2). After the exposure to 5 mM IPTG for 1 h or less, AG1[ASKA-(*tyrS*)] rapidly and irreversibly lost the ability to form colonies, suggesting that the effect of *tyrS* overexpression is bactericidal and suitable for host-killing (Figure S3). The availability of temperature-sensitive mutants and the presumed low impact of genomic gene deletion, as described below, are advantages for the application of *tyrS* to the kill-switch.

*E. coli* W3110 ts 29-13 strain is a temperature-sensitive mutant of *tyrS* [34]. This strain shows normal growth at 30°C but fails to form colonies at 42°C (Fig 3A). We transfected the plasmid ASKA-(*tyrS*) into this strain. The resulting strain, W3110 ts 29-13 [ASKA-(*tyrS*)], recovered colony formation at 42°C in the absence of IPTG, suggesting that ASKA-(*tyrS*) can supply TyrS at levels required for host survival and proliferation by leakage expression (Fig 3A). No colonies were observed in the presence of 3 mM IPTG at both 30°C and 42°C, suggesting that *tyrS* overexpression kills the host (Fig 3B). These results suggest that the “demon and angel” kill-switch using *tyrS* is feasible to construct, as we confirmed the rescue of host survival and growth by leakage expression in the OFF state and the host-killing by overexpression in the ON state.

**Figure 3.**
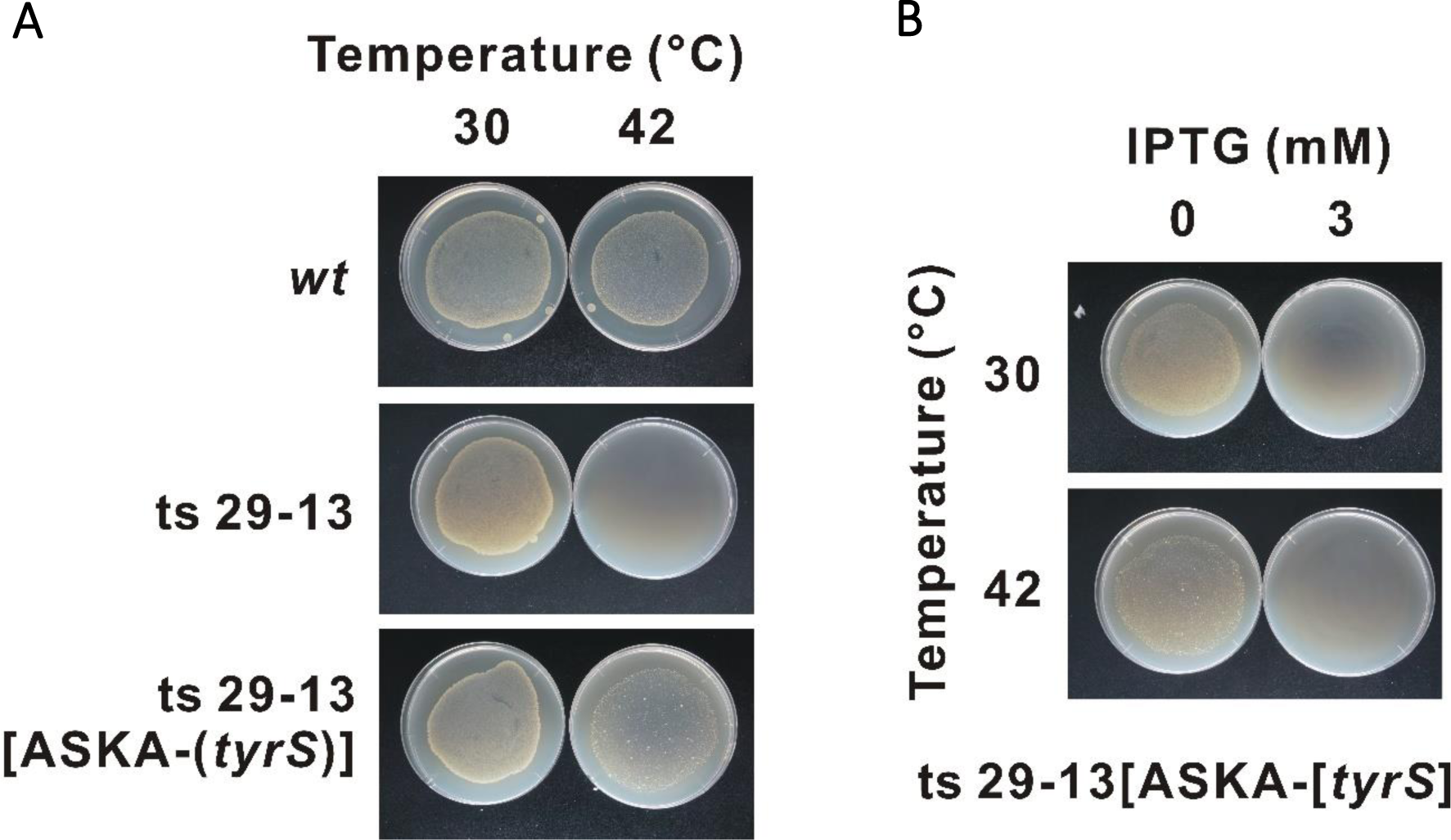
Evaluation of the “demon and angel” *tyrS* expression construct in W3110 ts 29-13. W3110 and its derivative strains were evaluated. (A) Maintenance of host survival and growth in the OFF state. W3110 ts 29-13 can grow at 30°C but not at 42°C, due to a temperature-sensitive mutation in *tyrS*. The ASKA-plasmid containing a conditional *tyrS* overexpression construct is carried in W3110 ts 29-13[ASKA-(tyrS)]. Since this experiment was performed in the absence of the inducer, IPTG, the overexpression of *tyrS* was not induced. Meanwhile, leakage expression of *tyrS* was expected. (B) Conditional host killing. Overexpression of *tyrS* encoded on the ASKA-plasmid is induced by 3 mM IPTG.

Next, we constructed the kill-switch in a *tyrS* deletion mutant strain. *tyrS* is part of the *pdxH-tyrS-pdxY* operon [35,36]. *pdxH* and *pdxY* are non-essential genes, suggesting that disruption of *tyrS* is unlikely to impair the function of other essential genes [37]. First, we transfected ASKA-(*tyrS*) into the unmodified BL21-AI to obtain BL21-AI [*tyrS^+^*; ASKA-(*tyrS*)] (Fig. 4A) [38]. The ORF of *tyrS* in the genome was subsequently deleted by substituting the neomycin resistance gene. The deletion was confirmed by genome sequencing. The resulting BL21-AI [*tyrS^-^*; ASKA-(*tyrS*)] strain survived and grew in the absence of IPTG, presumably due to leakage expression of the plasmid *tyrS* (Fig. 4B, upper panel). Meanwhile, BL21-AI [*tyrS^-^*; ASKA-(*tyrS*)] was killed in the presence of IPTG, as well as BL21-AI [*tyrS^+^*; ASKA-(*tyrS*)] (Figure S4).

**Figure 4.**
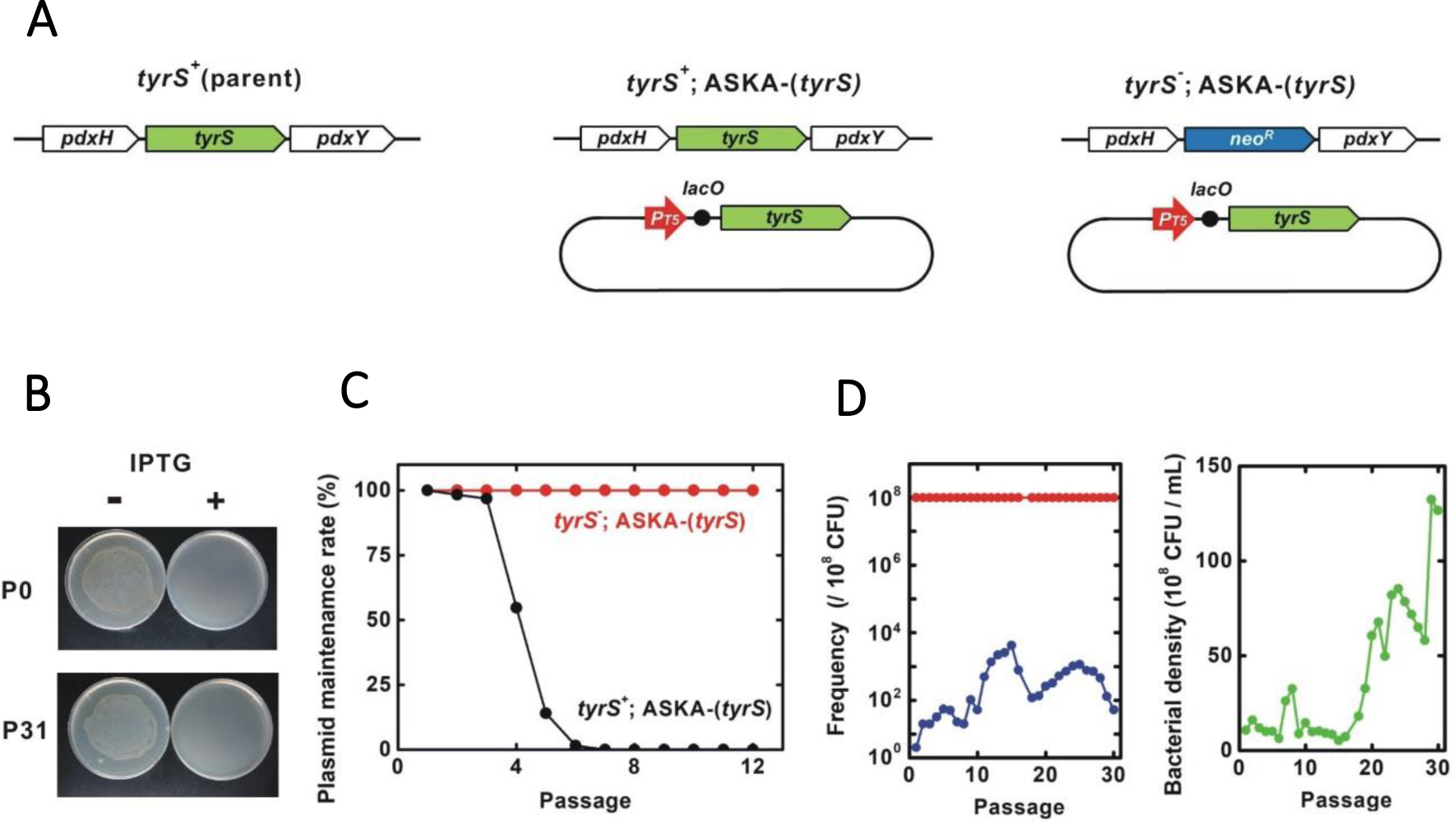
Genetic stability of the “demon and angel” kill-switch. (A) Genotype of the tested strains. BL21-AI was used as the parent strain. *neo^R^*, neomycin resistance gene. ASKA plasmids contain several other genes including *lacI^q^*and *cat.* (B) IPTG-induced host killing. Bacterial viability was tested in the presence and absence of 3 mM IPTG. P0 and P31, the BL21-AI [*tyrS^-^*; ASKA-(*tyrS*)] population after 0 and 31 passages, respectively. (C) Passage test to evaluate the maintenance of the ASKA-plasmid encoding *tyrS*. Bacterial cultures were diluted 10^3^-fold for each passage. Bacteria harboring the plasmid were identified by their resistance to chloramphenicol, the selection marker for ASKA-. ASKA-(*tyrS*) plasmid was stably maintained in BL21-AI [*tyrS^-^*; ASKA-(*tyrS*)] for >12 passages whereas BL21-AI [*tyrS^+^*; ASKA-(*tyrS*)] lost the plasmid after 6-7 passages. (D) Passage test to evaluate changes in the population frequency and bacterial density. Red and blue, IPTG-sensitive and -resistant bacteria, respectively. Green, bacterial density. Most of the bacteria remained IPTG-sensitive throughout the 30 passages. Survivor frequency showed complex changes. Bacterial density increased around the passage 18.

### Genetic stability of *tyrS* kill-switch

Passage tests showed that BL21-AI [*tyrS^+^*; ASKA-(*tyrS*)] strain, which retains *tyrS* in the genome, lost the ASKA plasmid after 6-7 passages (60-70 generations) in the absence of plasmid selection marker chloramphenicol (Fig.4C). In contrast, the ASKA plasmid was stably maintained for at least 120 generations in BL21-AI [*tyrS^-^*; ASKA-(*tyrS*)] in which the genomic *tyrS* was deleted, suggesting that the plasmid *tyrS* is essential for survival in this strain.

BL21-AI [*tyrS^-^*; ASKA-(*tyrS*)] was killed in the presence of IPTG (Fig. 4B, S4). In a passage test, the IPTG sensitivity was preserved well through more than 30 passages, equivalent to about 300 generations (Fig. 4B and 4D, left panel). Interestingly, the survivor frequency showed complex changes over the generations, with a range of less than 3 × 10^-5^, peaking at the passage 14-15 and 24-25. Survivors isolated from passages 14 and 30 (P14R and P30R, respectively) were IPTG-resistant, indicating the emergence of loss-of-function mutants (Fig. 5A). Notably, most bacteria (>99.997% at all time points of the 30 passages) maintained the kill-switch function and no rapid replacement with the loss-of-function mutants was observed (Fig. 4D, left panel).

**Figure 5.**
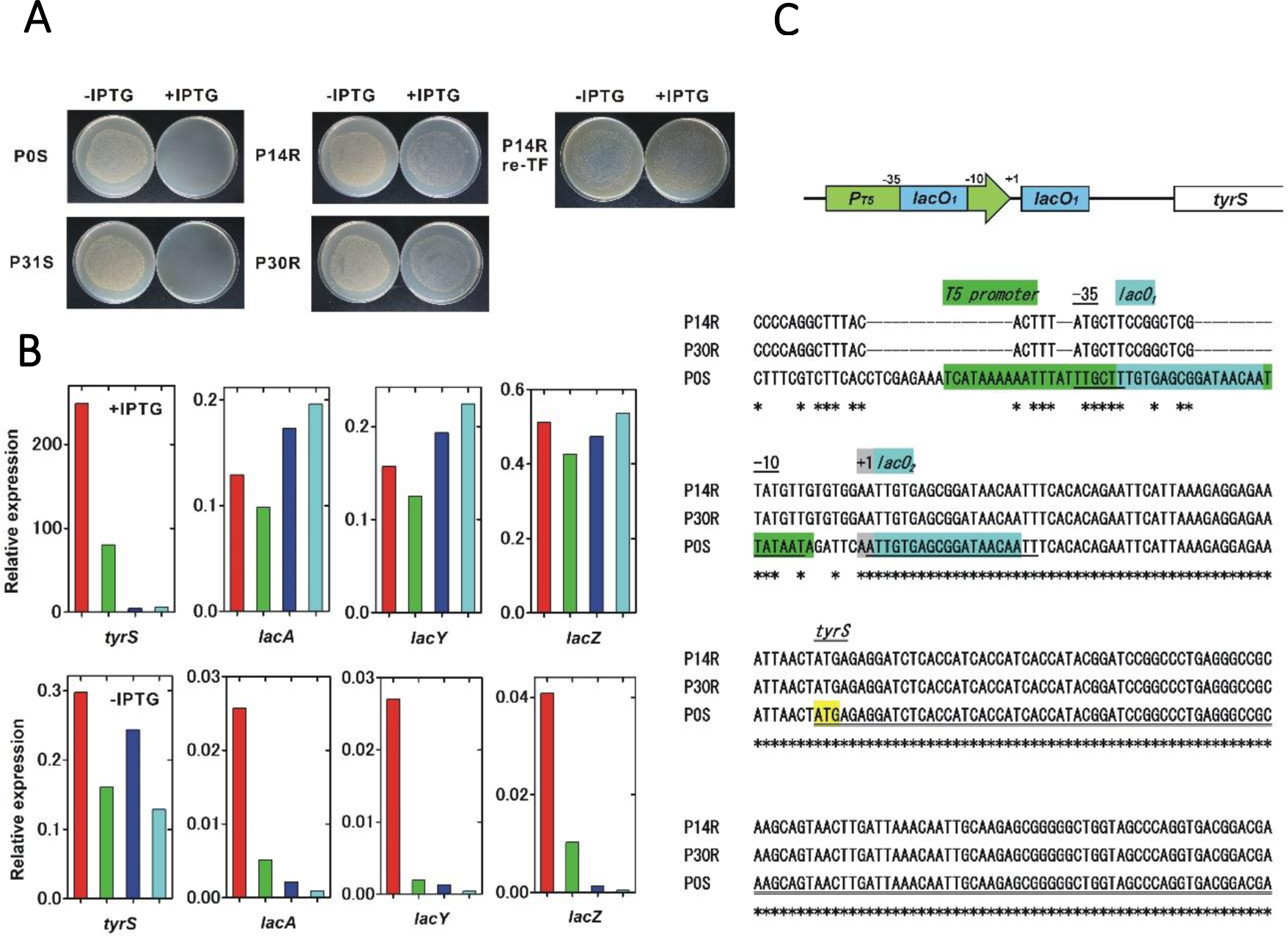
Characterization of the IPTG-resistant mutants. We characterized two IPTG-sensitive strains, P0S and P31S, and two IPTG-resistant mutants, P14R and P30R, which were isolated in the passage test shown in Figure 4. (A) IPTG sensitivity of the tested strains. P14R re-TF is the unmodified BL21-AI strain carrying the ASKA-plasmid isolated from P14R. (B) Relative expression of IPTG-inducible genes. A comprehensive analysis of gene expression was performed using RNA-seq. Red, green, dark blue and light blue columns indicate P0S, P31S, P14R and P30R, respectively. The expression level was normalized to that of a constitutively expressed gene, *gltA*, citrate synthase. (C) Mutations found in the plasmid *tyrS* regulatory region. A schematic diagram of the *tyrS* regulatory region in the ASKA-plasmid is shown in the upper panel. The nucleotide sequences of the ASKA-plasmids isolated from P14R, P30R and P0S are aligned and shown in the lower panel. An asterisk indicates a conserved base.

To identify the IPTG-resistant mechanisms, the transcriptomes in the IPTG-sensitive strains isolated from passage 0 and 31, P0S and P31S, and the IPTG-resistant strains, P14R and P30R, were analyzed using RNA-seq. We found that the induction of the IPTG-regulated *tyrS* on the ASKA plasmid remarkably decreased in the IPTG-resistant strains although the IPTG-inducibility still remained partial (Figure 5B). The *tyrS* induction ratio was 836.3, 499.1, 18.1 and 42.8, for P0S, P31S, P14R and P30R, respectively. Meanwhile, the basal expression of the plasmid *tyrS* was similar in both the IPTG-sensitive and resistant strains. Interestingly, the IPTG-induced expression level of the native IPTG-inducible genes in the genome, *lacA*, *lacY* and *lacZ*, was comparable between IPTG-sensitive and resistant strains. The basal expression of those genomic genes remarkably decreased in P14R and P30R although the expression of *lacI*, that negatively regulates the expression of the IPTG-inducible genes, was not clearly affected (Figure S5).

Next, we sequenced the total DNAs containing both genomic and plasmid DNA isolated from P0S, P14R and P30R, using Illumina sequencing. Severe mutations were found in the regulatory region of *tyrS* in the ASKA plasmid in P14R and P30R whose sequences were completely identical (Figure 5C). The coliphage T5 promoter region was largely mutated, but the -10 and -35 regions were relatively conserved. Although *lacO_1_* sequence embedded in the promoter was almost destroyed, *lacO_2_* located between the promoter and the translation initiation site was completely conserved. The plasmid *lacI* was intact. Some additional genomic mutations were also detected in P14R and P30R (Table S4). Finally, the ASKA plasmid was extracted from P14R and transformed into unmodified BL21-AI. The resulting strain was IPTG-resistant, suggesting that the mutations on the plasmid are sufficient to cause loss of the kill-switch function (Figure 5A).

Notably, the bacterial density clearly started to increase from around the passage 18 and grew to the passage 29-30 (Figure 4D, right panel). The growth rate of P31S (doubling time = 30.3 min) was greater than those of P0S (41.5 min) and P30R (33.3 min).

## DISCUSSION

In this paper, we propose a novel principle for constructing kill-switches with high genetic stability. The kill-switch is based on a “demon and angel” essential gene expression construct on which both viability maintenance and host-killing depend. We constructed the kill-switch in *E. coli* using *tyrS* as a model. The viability maintenance essentiality in the OFF state and the host-killing in the ON state were maintained at least for 120 and 300 generations, respectively. However, the *tyrS* kill-switch was not an ideal case because we detected some loss-of-function mutants. The following is an analysis and discussion of the underlying causes.

In the loss-of-function mutants, transcriptome analyses showed a significant decrease in the *tyrS* expression level upon IPTG induction although the native IPTG-inducible genes were normally upregulated. Re-transformation of the ASKA-(*tyrS*) plasmid isolated from the mutant reproduced the IPTG-resistant phenotype, indicating that the mutations in the plasmid were the main cause. NGS analyses detected substantial mutations in the regulatory region of the plasmidal *tyrS*, supporting this observation. In the mutated plasmids, the coliphage T5 promoter region was severely impaired, suggesting that the promoter activity substantially decreased. Additionally, the disruption of *lacO_1_* embedded in the promoter sequence may mitigate the reduction in *tyrS* leakage expression. The sequence of the plasmidal *tyrS* regulatory region was completely identical between P14R and P30R, suggesting that P14R or its sister strain acquired further mutations in the genome and evolved into P30R. Based on the above observation, we propose a possible mechanism for the complex change of survivor frequency observed in the passage test. (1) Mutations in the ASKA plasmid reduced the activity of the promoter that controls the expression of *tyrS*, resulting in the severe decrease of *tyrS* expression in the ON state. The reduced level of TyrS failed to kill the host, causing the IPTG-resistance in P14R and P30R. (2) The weaker promoter activity and the *lacO_1_* disruption also affected the *tyrS* leakage expression level. Overall, a similar level of tyrS expression in the OFF state was observed in both IPTG-sensitive and resistant strains, but the loss-of-function mutants had slightly higher fitness. Consequently, the proportion of the mutants gradually increased as the generation progressed. (3) Subsequently, the fraction of IPTG-resistant mutants decreased with the emergence of other mutants, including P31S that was IPTG-sensitive and more rapidly grew to higher density.

The “demon and angel” kill switch was designed to be highly resistant to point mutations and short insertion/deletion mutations, as described in the introduction. Surprisingly, the isolated loss-of-function mutants, P14R and P30R, acquired heavily deleted and substituted sequences in the *tyrS* regulatory region, suggesting that the large rearrangement of regulatory region is a serious obstacle. The loss-of-function mutants maintained *tyrS* expression levels in the OFF state, contributing to the higher fitness acquisition. To eliminate the fitness-advantage in the mutants, the expression level of the toxic essential gene should be adjusted to an optimal level in the bacteria harboring the functional kill-switch. This condition can be achieved by selecting and adjusting optimal genetic elements such as promoters, induction systems, RBSs, plasmid replication origins and toxic essential genes. P31S acquired a higher fitness, suggesting that evolutionary methods are also effective.

Of the 300 essential genes, 112 genes were associated with severely damaged phenotypes of massive growth inhibition and no GPF expression, suggesting that the essential genes with toxic overexpression phenotype are abundant. These toxic essential genes are candidate genetic parts to construct the “demon and angel” kill-switches.

Focusing on 24 aaRS genes, 23 genes were toxic, of which 9 genes exhibited the severely damaged phenotypes. The proper balance of the aaRS and the cognate tRNAs is critical in maintaining the fidelity of protein biosynthesis [39]. The production of many aaRSs is regulated via various negative autoregulatory mechanisms [40]. The toxic-overexpression of some aaRS genes, including *tyrS*, have been previously reported [41–44]. In *tyrS*, the toxicity is directly correlated to the enzymatic activity [44]. This correlation is an advantage for use in the “demon and angel” kill-switch because mutations reducing toxicity always reduce essential enzymatic activity. The effect of *tyrS* overexpression was bactericidal, indicating its suitability for use in kill switches. Interestingly, most natural amino acid misincorporations by misacylated aminoacyl-tRNAs are tolerated [45,46]. The informational disturbance caused by some aaRS overexpression could be more toxic than the misincorporation at the specific codons.

Genes that are toxic upon overexpression have been reported not only in bacteria but also in various unicellular and multicellular eukaryotes [47–50]. The application of the “demon and angel” kill switch should be scalable to various organisms.

## DATA AVAILABILITY

The data underlying this article are available in the article and in its online supplementary material.

## SUPPLEMENTARY DATA

Supplementary Data are available at NAR online.

