## Supplementary information for "Genetically stable kill-switch using “demon and angel” expression construct of essential genes"

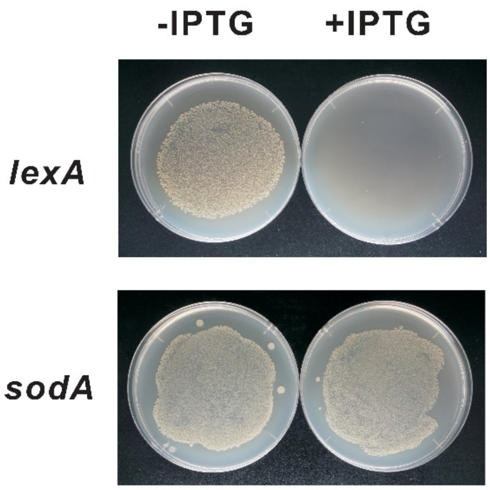


Figure S1. Toxic overexpression of endogenous genes. We tested the *E. coli* BL21-AI strain with an ASKA-plasmid containing an IPTG-inducible overexpression construct of the indicated gene. Overnight cultures were diluted $10^3$-fold and inoculated onto LB-agar plates with or without 5 mM IPTG. *lexA* and *sodA* showed the toxic and non-toxic overexpression phenotype, respectively.

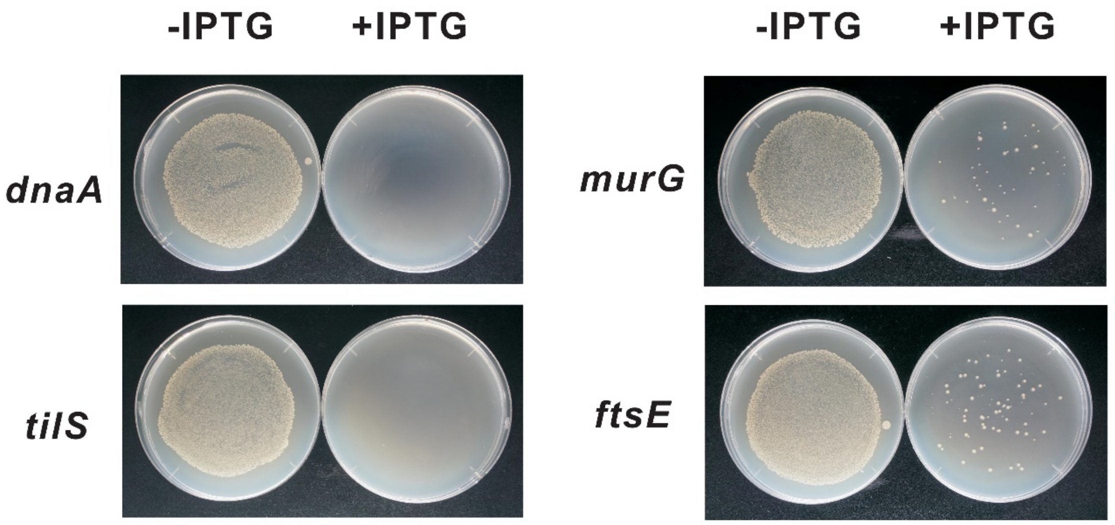


Figure S2. Various toxicity severity of essential genes. Overnight cultures were diluted $10^3$-fold and inoculated onto LB-agar plates with or without 5 mM IPTG.

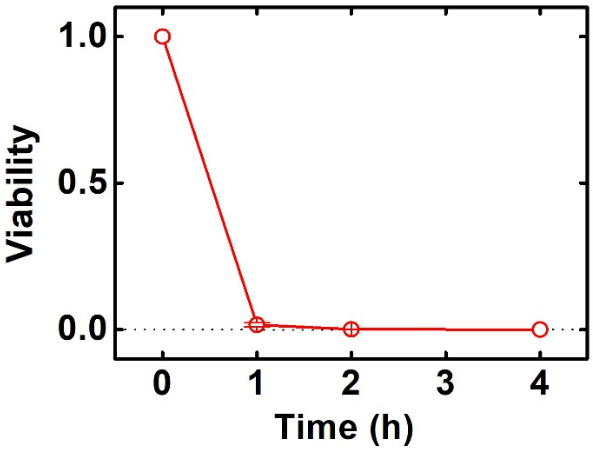


Figure S3. Mode of action for *tyrS* toxic-overexpression. Viability of BL21-AI [tyrS-; ASKA-(tyrS)] after addition of IPTG (final 5 mM) was determined. Data points represent the mean ± s.d. of three biological replicates.

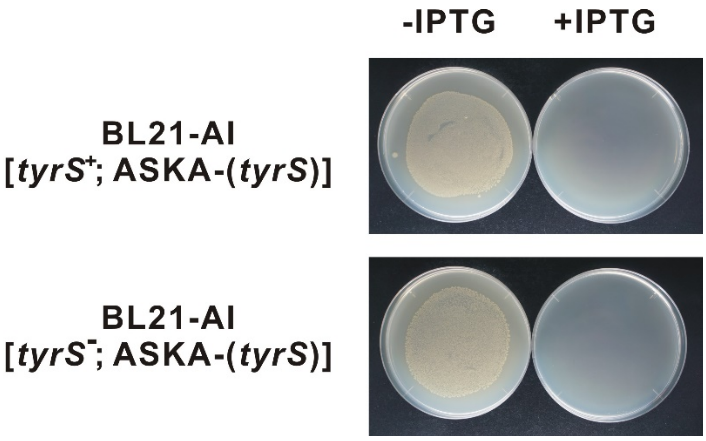


Figure S4. IPTG-sensitivity of BL21-AI carrying ASKA-(*tyrS*). Overnight cultures were diluted $10^3$-fold and inoculated onto LB-agar plates with or without 5 mM IPTG.

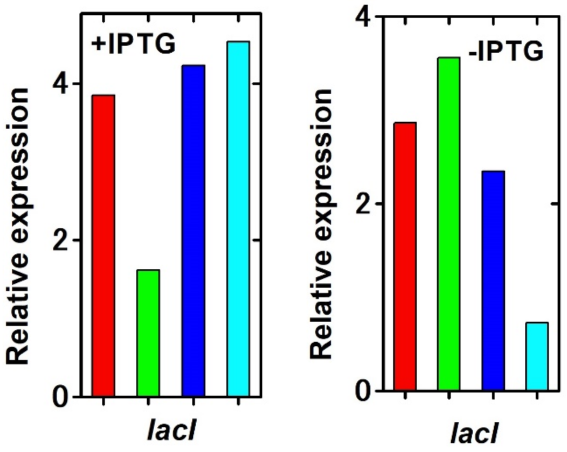


Figure S5. Relative expression of *lacI*. Comprehensive analysis of gene expression was performed using RNA-seq. Red, green, dark blue and light blue column indicate P0S, P31S, P14R and P30R, respectively. The expression level was normalized by that of a constitutively expressed gene, *gltA*, citrate synthase.

Table S1. Growth inhibition and GFP expression in overexpression of ASKA+ plasmids. Total 4269 clones were tested. Results are listed in alphabetical order by gene name. The intensity of GFP expression was evaluated on the following three levels： gg, strong; g, weak; o, not detected. Growth inhibition was evaluated on the following three levels: tt, no growth; t, slower growth; o, normal growth.

| Gene | Feature | Product | Location | Function | Process | GFP fluorescence | IPTG growth |
|---|---|---|---|---|---|---|---|
| *aaeA* | CDS | p-hydroxybenzoic acid efflux system component | Periplasmic | COG1566;Multidrug resistance efflux pump | | | |
| *aaeB* | CDS | p-hydroxybenzoic acid efflux system component | Integral Membrane Protein | COG1289;Predicted membrane protein | | o | tt |
| *aaeR* | CDS | predicted DNA-binding transcriptional regulator, efflux system | Cytoplasmic | COG0583;Transcriptional regulator | GO:0006350 transcription | o | tt |
| *aaeX* | CDS | membrane protein of efflux system | Integral Membrane Protein | | | | |
| *aas* | CDS | fused 2-acylglycerophospho-ethanolamine acyl transferase -!- acyl-acyl carrier protein synthetase | Integral Membrane Protein | COG0204;1-acyl-sn-glycerol-3-phosphate acyltransferase -!- COG0318;Acyl-CoA synthetases (AMP-forming)/AMP-acid ligases II | GO:0006464 protein modification | o | tt |
| *aat* | CDS | leucyl/phenylalanyl-tRNA-protein transferase | Cytoplasmic | COG2360;Leu/Phe-tRNA-protein transferase | GO:0006508 proteolysis and peptidolysis | gg | tt |
| *abgA* | CDS | predicted peptidase, aminobenzoyl-glutamate utilization protein | Cytoplasmic | | GO:0016052 carbohydrate catabolism | | |
| *abgB* | CDS | predicted peptidase, aminobenzoyl-glutamate utilization protein | Cytoplasmic | COG1473;Metal-dependent amidase/aminoacylase/carboxypeptidase | GO:0016052 carbohydrate catabolism | gg | t |
| *abgR* | CDS | predicted DNA-binding transcriptional regulator | Cytoplasmic | COG0583;Transcriptional regulator | GO:0016052 carbohydrate catabolism -!- GO:0006350 | gg | tt |
| *abgT* | CDS | predicted cryptic aminobenzoyl-glutamate transporter | Integral Membrane Protein | COG2978;Putative p-aminobenzoyl-glutamate transporter | | | |
| *abrB* | CDS | predicted regulator | Integral Membrane Protein | | | | |
| *accA* | CDS | acetylCoA carboxylase, carboxytransferase, alpha subunit | Cytoplasmic | COG0825;Acetyl-CoA carboxylase alpha subunit | | gg | o |
| *accB* | CDS | acetyl CoA carboxylase, BCCP subunit | Cytoplasmic | COG0511;Biotin carboxyl carrier protein | GO:0042966 biotin carboxyl carrier protein biosynthesis | o | tt |
| *accC* | CDS | acetyl-CoA carboxylase, biotin carboxylase subunit | Cytoplasmic | COG0439;Biotin carboxylase | | o | tt |
| *accD* | CDS | acetylCoA carboxylase, beta (carboxyltranferase) subunit | Cytoplasmic | COG0777;Acetyl-CoA carboxylase beta subunit | | gg | t |
| *aceA* | CDS | isocitrate lyase | Cytoplasmic | COG2224;Isocitrate lyase | GO:0006097 glyoxylate cycle | o | tt |
| *aceB* | CDS | malate synthase A | Cytoplasmic | COG2225;Malate synthase | GO:0006097 glyoxylate cycle | gg | t |
| *aceE* | CDS | pyruvate dehydrogenase, decarboxylase component E1, thiamin-binding | Cytoplasmic | COG2609;Pyruvate dehydrogenase complex, dehydrogenase (E1) component | GO:0016052 carbohydrate catabolism -!- GO:0006086 acetyl-CoA biosynthesis from pyruvate -!- GO:0006096 glycolysis -!- GO:0009436 glyoxylate catabolism -!- GO:0009061 anaerobic respiration | gg | tt |
| *aceF* | CDS | pyruvate dehydrogenase, dihydrolipoyltransacetylase component E2 | Cytoplasmic | COG0508;Pyruvate/2-oxoglutarate dehydrogenase complex, dihydrolipoamide acyltransferase (E2) component, and related enzymes | GO:0016052 carbohydrate catabolism -!- GO:0006086 acetyl-CoA biosynthesis from pyruvate -!- GO:0006096 glycolysis -!- GO:0009436 glyoxylate catabolism -!- GO:0009061 anaerobic respiration | o | tt |
| *aceK* | CDS | isocitrate dehydrogenase kinase/phosphatase | Cytoplasmic | COG4579;Isocitrate dehydrogenase kinase/phosphatase | GO:0006097 glyoxylate cycle -!- GO:0006464 protein modification | o | t |
| *ackA* | CDS | acetate kinase A and propionate kinase 2 | Cytoplasmic | COG0282;Acetate kinase | GO:0016052 carbohydrate catabolism -!- GO:0009063 amino acid catabolism -!- GO:0042867 pyruvate catabolism -!- GO:0045733 acetate catabolism | gg | t |
| *acnA* | CDS | aconitate hydratase 1 | Cytoplasmic | COG1048;Aconitase A | GO:0006099 tricarboxylic acid cycle -!- GO:0009061 anaerobic respiration | g | tt |
| *acnB* | CDS | bifunctional aconitate hydratase 2 -!- 2-methylisocitrate dehydratase | Cytoplasmic | COG1049;Aconitase B | GO:0006099 tricarboxylic acid cycle -!- GO:0009060 aerobic respiration -!- GO:0009061 anaerobic respiration | gg | o |
| *acpP* | CDS | acyl carrier protein (ACP) | Cytoplasmic | COG0236;Acyl carrier protein | GO:0042967 acyl-carrier protein biosynthesis | gg | t |
| *acpS* | CDS | holo-[acyl-carrier-protein] synthase 1 | Cytoplasmic | COG0736;Phosphopantetheinyl transferase (holo-ACP synthase) | | gg | t |
| *acpT* | CDS | holo-(acyl carrier protein) synthase 2 | Cytoplasmic | COG2091;Phosphopantetheinyl transferase | | o | t |
| *acrA* | CDS | multidrug efflux system | Inner Membrane Lipoprotein | COG0845;Membrane-fusion protein | GO:0042493 response to drug | o | tt |
| *acrB* | CDS | multidrug efflux system protein | Integral Membrane Protein | COG0841;Cation/multidrug efflux pump | GO:0042493 response to drug | o | tt |
| *acrD* | CDS | aminoglycoside/multidrug efflux system | Integral Membrane Protein | COG0841;Cation/multidrug efflux pump | GO:0042493 response to drug | o | tt |
| *acrE* | CDS | cytoplasmic membrane lipoprotein | Inner Membrane Lipoprotein | COG0845;Membrane-fusion protein | | o | tt |
| *acrF* | CDS | multidrug efflux system protein | Integral Membrane Protein | COG0841;Cation/multidrug efflux pump | GO:0042493 response to drug | o | o |
| *acrR* | CDS | DNA-binding transcriptional regulator | Cytoplasmic | COG1309;Transcriptional regulator | GO:0006350 transcription -!- GO:0042493 response to drug | gg | tt |
| *acs* | CDS | bifunctional acetyl-CoA synthetase -!- propionyl-CoA synthetase | Cytoplasmic | COG0365;Acyl-coenzyme A synthetases/AMP-(fatty) acid ligases | GO:0016052 carbohydrate catabolism -!- GO:0045733 acetate | gg | t |
| *actP* | CDS | acetate transporter | Integral Membrane Protein | COG4147;Predicted symporter | | o | tt |
| *ada* | CDS | fused DNA-binding transcriptional dual regulator -!- O6-methylguanine-DNA methyltransferase | Cytoplasmic | COG2169;Adenosine deaminase -!- COG0350;Methylated DNA-protein cysteine methyltransferase | GO:0006281 DNA repair -!- GO:0006350 transcription | o | tt |
| *add* | CDS | adenosine deaminase | Cytoplasmic | COG1816;Adenosine deaminase | GO:0015949 nucleobase, nucleoside and nucleotide interconversion | gg | o |
| *ade* | CDS | cryptic adenine deaminase | Cytoplasmic | COG1001;Adenine deaminase | GO:0008152 metabolism | g | o |
| *adhE* | CDS | fused acetaldehyde-CoA dehydrogenase -!- iron-dependent alcohol dehydrogenase -!- pyruvate-formate lyase deactivase | Cytoplasmic | COG1012;NAD-dependent aldehyde dehydrogenases -!- COG1454;Alcohol dehydrogenase, class IV | GO:0006113 fermentation | gg | t |
| *adhP* | CDS | alcohol dehydrogenase, 1-propanol preferring | Cytoplasmic | | GO:0009061 anaerobic respiration | gg | t |
| *adiA* | CDS | biodegradative arginine decarboxylase | Cytoplasmic | COG1982;Arginine/lysine/ornithine decarboxylases | GO:0009063 amino acid catabolism | | |
| *adiC* | CDS | arginine:agmatin | Integral Membrane Protein | COG0531;Amino acid transporters | | o | tt |
| *adiY* | CDS | DNA-binding transcriptional activator | Cytoplasmic | COG2207;AraC-type DNA-binding domain-containing proteins | GO:0006350 transcription | gg | t |
| *adk* | CDS | adenylate kinase | Cytoplasmic | COG0563;Adenylate kinase and related kinases | GO:0015949 nucleobase, nucleoside and nucleotide interconversion -!- GO:0009152 purine ribonucleotide biosynthesis | gg | o |
| *aegA* | CDS | fused predicted FeS binding subunit -!- predicted NAD/FAD-binding subunit of oxidoreductase | Cytoplasmic | COG1142;Fe-S-cluster-containing hydrogenase components 2 -!- COG0493;NADPH-dependent glutamate synthase beta chain and related oxidoreductases | | gg | t |
| *aer* | CDS | fused signal transducer for aerotaxis sensory component -!- methyl accepting chemotaxis component | Membrane Anchored | COG2202;FOG: PAS/PAC domain -!- COG0840;Methyl-accepting chemotaxis protein | GO:0016052 carbohydrate catabolism -!- GO:0042330 taxis | o | tt |
| *aes* | CDS | acetyl esterase | Cytoplasmic | COG0657;Esterase/lipase | GO:0016052 carbohydrate | o | t |
| *afuB* | CDS | CP4-6 prophage; predicted ferric trasnporter subunit -!- membrane component of ABC superfamily | Integral Membrane Protein | COG1178;ABC-type Fe3+ transport system, permease component | | o | tt |
| *afuC* | CDS | CP4-6 prophage; predicted ferric transporter subunit -!- ATP-binding component of ABC superfamily | Cytoplasmic | COG3842;ABC-type spermidine/putrescine transport systems, ATPase components | | gg | tt |
| *agaA* | CDS | predicted truncated N-acetylgalactosamine-6-phosphate deacetylase | Cytoplasmic | | | | |
| *agaB* | CDS | N-acetylgalactosamine-specific enzyme IIB component of PTS | Cytoplasmic | COG3444;Phosphotransferase system, mannose/fructose/N-acetylgalactosamine-specific component IIB | | gg | t |
| *agaC* | CDS | N-acetylgalactosamine-specific enzyme IIC component of PTS | Integral Membrane Protein | COG3715;Phosphotransferase system, mannose/fructose/N-acetylgalactosamine-specific component IIC | | o | tt |
| *agaD* | CDS | N-acetylgalactosamine-specific enzyme IID component of PTS | Integral Membrane Protein | COG3716;Phosphotransferase system, mannose/fructose/N-acetylgalactosamine-specific component IID | GO:0046349 amino sugar biosynthesis | o | tt |
| *agaI* | CDS | galactosamine-6-phosphate isomerase | Cytoplasmic | COG0363;6-phosphogluconolactonase/Glucosamine-6-ph osphate isomerase/deaminase | | gg | t |
| *agaR* | CDS | DNA-binding transcriptional dual regulator | Cytoplasmic | COG1349;Transcriptional regulators of sugar metabolism | GO:0046349 amino sugar biosynthesis -!- GO:0006350 transcription | g | tt |
| *agaS* | CDS | tagatose-6-phosphate ketose/aldose isomerase | Cytoplasmic | COG2222;Predicted phosphosugar isomerases | | gg | o |
| *agaV* | CDS | N-acetylgalactosamine-specific enzyme IIB component of PTS | Cytoplasmic | | GO:0046349 amino sugar biosynthesis | gg | o |
| *agaW* | CDS | N-acetylgalactosamine-specific enzyme IIC component of PTS, fragment (pseudogene) | Integral Membrane Protein | COG3715;Phosphotransferase system, mannose/fructose/N-acetylgalactosamine-specific component IIC | GO:0046349 amino sugar biosynthesis | o | tt |
| *agp* | CDS | glucose-1-phosphatase/inositol phosphatase | Periplasmic | | GO:0006006 glucose metabolism | o | tt |
| *ahpC* | CDS | alkyl hydroperoxide reductase, C22 subunit | Cytoplasmic | COG0450;Peroxiredoxin | GO:0006805 xenobiotic metabolism | gg | t |
| *ahpF* | CDS | alkyl hydroperoxide reductase, F52a subunit, FAD/NAD(P)-binding | Cytoplasmic | COG3634;Alkyl hydroperoxide reductase, large subunit | GO:0006805 xenobiotic metabolism | gg | tt |
| *aidB* | CDS | isovaleryl CoA dehydrogenase | Cytoplasmic | | GO:0006950 response to stress | | |
| *ais* | CDS | conserved protein | Membrane Anchored | | | g | tt |
| *alaS* | CDS | alanyl-tRNA synthetase | Cytoplasmic | COG0013;Alanyl-tRNA synthetase | GO:0006418 amino acid activation | gg | tt |
| *alaT* | tRNA | tRNA-Ala(UGC) (Alanine tRNA 1B) | | | | | |
| *alaU* | tRNA | tRNA-Ala(UGC) (Alanine tRNA 1B) | | | | | |
| *alaV* | tRNA | tRNA-Ala(UGC) (Alanine tRNA 1B) | | | | | |
| *alaW* | tRNA | tRNA-Ala(GGC), (Alanine tRNA 2) | | | | | |
| *alaX* | tRNA | tRNA-Ala(GGC) (Alanine tRNA 2) | | | | | |
| *aldA* | CDS | aldehyde dehydrogenase A, NAD-linked | Cytoplasmic | COG1012;NAD-dependent aldehyde dehydrogenases | GO:0016052 carbohydrate catabolism -!- GO:0009438 methylglyoxal metabolism -!- GO:0006113 fermentation -!- GO:0009441 glycolate metabolism | o | t |
| *aldB* | CDS | aldehyde dehydrogenase B | Cytoplasmic | | GO:0016052 carbohydrate catabolism -!- GO:0009438 methylglyoxal metabolism | g | o |
| *alkA* | CDS | 3-methyl-adenine DNA glycosylase II | Cytoplasmic | COG0122;3-methyladenine DNA glycosylase/8-oxoguanine DNA | GO:0006281 DNA repair | g | o |
| *alkB* | CDS | oxidative demethylase of N1-methyladenine or N3-methylcytosine DNA lesions | Cytoplasmic | COG3145;Alkylated DNA repair protein | GO:0006281 DNA repair | gg | o |
| *allA* | CDS | ureidoglycolate hydrolase | Cytoplasmic | COG3194;Ureidoglycolate hydrolase | GO:0006807 nitrogen metabolism -!- GO:0009442 allantoin assimilation | gg | t |
| *allB* | CDS | allantoinase | Cytoplasmic | COG0044;Dihydroorotase and related cyclic amidohydrolases | GO:0006807 nitrogen metabolism -!- GO:0009442 allantoin assimilation | gg | t |

| | | | | | | | |
|---|---|---|---|---|---|---|---|
| *allC* | CDS | allantoate amidohydrolase | Cytoplasmic | COG0624;Acetylornithine deacetylase/Succinyl-diaminopimelate desuccinylase and related deacylases | GO:0006807 nitrogen metabolism -!- GO:0009442 allantoin assimilation | o | t |
| *allD* | CDS | ureidoglycolate dehydrogenase | Cytoplasmic | COG2055;Malate/L-lactate dehydrogenases | GO:0006807 nitrogen metabolism -!- GO:0009442 allantoin assimilation | gg | t |
| *allR* | CDS | DNA-binding transcriptional regulator | Cytoplasmic | COG1414;Transcriptional regulator | GO:0006350 transcription | gg | tt |
| *alpA* | CDS | CP4-57 prophage; DNA-binding transcriptional activator | Cytoplasmic | COG3311;Predicted transcriptional regulator | GO:0006350 transcription | o | tt |
| *alr* | CDS | alanine racemase 1, PLP-binding, biosynthetic | Cytoplasmic | COG0787;Alanine racemase | GO:0006523 alanine biosynthesis | gg | o |
| *alsA* | CDS | fused D-allose transporter subunits -!- ATP-binding components of ABC superfamily | Cytoplasmic | COG1129;ABC-type sugar transport system, ATPase component | GO:0016052 carbohydrate catabolism | g | o |
| *alsB* | CDS | D-allose transporter subunit -!- periplasmic-binding component of ABC superfamily | Periplasmic | COG1879;ABC-type sugar transport system, periplasmic component | GO:0016052 carbohydrate catabolism | o | tt |
| *alsC* | CDS | D-allose transporter subunit -!- membrane component of ABC superfamily | Integral Membrane Protein | COG1172;Ribose/xylose/arabinose/galactoside ABC-type transport systems, permease components | GO:0016052 carbohydrate catabolism | o | tt |
| *alsE* | CDS | allulose-6-phosphate 3-epimerase | Cytoplasmic | COG0036;Pentose-5-phosphate-3-epimerase | | gg | o |
| *alsK* | CDS | D-allose kinase | Cytoplasmic | COG1940;Transcriptional regulator/sugar kinase | GO:0006355 regulation of transcription, DNA-dependent | | |
| *alx* | CDS | predicted inner membrane protein, part of terminus | Integral Membrane Protein | | GO:0006950 response to stress | | |
| *amiA* | CDS | N-acetylmuramoyl-l-alanine amidase I | Periplasmic | COG0860;N-acetylmuramoyl-L-alanine amidase | GO:0009252 peptidoglycan biosynthesis | gg | t |
| *amiB* | CDS | N-acetylmuramoyl-l-alanine amidase II | Periplasmic | COG0860;N-acetylmuramoyl-L-alanine amidase | GO:0009252 peptidoglycan biosynthesis | o | tt |
| *amiC* | CDS | N-acetylmuramoyl-L-alanine amidase | Periplasmic | | GO:0009252 peptidoglycan biosynthesis | | |
| *amn* | CDS | AMP nucleosidase | Cytoplasmic | COG0775;Nucleoside phosphorylase | GO:0015949 nucleobase, nucleoside and nucleotide interconversion | gg | o |
| *ampC* | CDS | beta-lactamase/D-alanine carboxypeptidase | Periplasmic | COG1680;Beta-lactamase class C and other penicillin binding proteins | GO:0009252 peptidoglycan biosynthesis -!- GO:0042493 response to drug | o | tt |
| *ampD* | CDS | N-acetyl-anhydromuranmyl-L-alanine amidase | Cytoplasmic | COG3023;Negative regulator of beta-lactamase expression | GO:0000270 peptidoglycan metabolism | gg | o |
| *ampE* | CDS | predicted inner membrane protein | Integral Membrane Protein | COG3725;Membrane protein required for beta-lactamase induction | GO:0042493 response to drug | o | tt |
| *ampG* | CDS | muropeptide transporter | Integral Membrane Protein | COG0477;Permeases of the major facilitator superfamily | GO:0000270 peptidoglycan metabolism | o | tt |
| *ampH* | CDS | beta-lactamase/D-alanine carboxypeptidase | Periplasmic | COG1680;Beta-lactamase class C and other penicillin binding proteins | GO:0042493 response to drug | | |
| *amtB* | CDS | ammonium transporter | Integral Membrane Protein | COG0004;Ammonia permease | | o | tt |
| *amyA* | CDS | cytoplasmic alpha-amylase | Cytoplasmic | COG0366;Glycosidases | GO:0016052 carbohydrate catabolism -!- GO:0000272 polysaccharide catabolism | o | tt |
| *ansA* | CDS | cytoplasmic L-asparaginase I | Cytoplasmic | COG0252;L-asparaginase/archaeal Glu-tRNAGln amidotransferase subunit D | GO:0009063 amino acid catabolism | gg | t |
| *ansB* | CDS | periplasmic L-asparaginase II | Periplasmic | COG0252;L-asparaginase/archaeal Glu-tRNAGln amidotransferase subunit D | GO:0009063 amino acid catabolism | g | tt |
| *ansP* | CDS | L-asparagine transporter | Integral Membrane Protein | | GO:0006529 asparagine | | |
| *apaG* | CDS | protein associated with Co2+ and Mg2+ efflux | Cytoplasmic | COG2967;Uncharacterized protein affecting Mg2+/Co2+ transport | | gg | tt |
| *apaH* | CDS | diadenosine tetraphosphatase | Cytoplasmic | COG0639;Diadenosine tetraphosphatase and related serine/threonine protein phosphatases | GO:0015949 nucleobase, nucleoside and nucleotide interconversion | gg | tt |
| *aphA* | CDS | acid phosphatase/phosphotransferase, class B, non-specific | Periplasmic | COG3700;Acid phosphatase (class B) | GO:0015949 nucleobase, nucleoside and nucleotide interconversion | o | tt |
| *appA* | CDS | phosphoanhydride phosphorylase | Periplasmic | | GO:0006793 phosphorus metabolism -!- GO:0042594 response to | o | tt |
| *appB* | CDS | cytochrome bd-II oxidase, subunit II | Integral Membrane Protein | COG1294;Cytochrome bd-type quinol oxidase, subunit 2 | | o | tt |
| *appC* | CDS | cytochrome bd-II oxidase, subunit I | Integral Membrane Protein | COG1271;Cytochrome bd-type quinol oxidase, subunit 1 | | o | tt |
| *appY* | CDS | DLP12 prophage; DNA-binding transcriptional activator | Cytoplasmic | COG2207;AraC-type DNA-binding domain-containing proteins | GO:0006350 transcription -!- GO:0042594 response to starvation | o | tt |
| *apt* | CDS | adenine phosphoribosyltransferase | Cytoplasmic | COG0503;Adenine/guanine phosphoribosyltransferases and related PRPP-binding proteins | GO:0015949 nucleobase, nucleoside and nucleotide interconversion | gg | o |
| *aqpZ* | CDS | aquaporin | Integral Membrane Protein | COG0580;Glycerol uptake facilitator and related permeases (Major Intrinsic Protein Family) | GO:0006970 response to osmotic stress | o | tt |
| *araA* | CDS | L-arabinose isomerase | Cytoplasmic | COG2160;L-arabinose isomerase | GO:0016052 carbohydrate | o | tt |
| *araB* | CDS | L-ribulokinase | Cytoplasmic | COG1069;Ribulose kinase | GO:0016052 carbohydrate | gg | o |
| *araC* | CDS | DNA-binding transcriptional dual regulator | Cytoplasmic | COG2207;AraC-type DNA-binding domain-containing proteins | GO:0016052 carbohydrate catabolism -!- GO:0006350 | gg | t |
| *araD* | CDS | L-ribulose-5-phosphate 4-epimerase | Cytoplasmic | COG0235;Ribulose-5-phosphate 4-epimerase and related epimerases and aldolases | GO:0016052 carbohydrate catabolism | gg | o |
| *araE* | CDS | arabinose transporter | Integral Membrane Protein | COG0477;Permeases of the major facilitator superfamily | GO:0016052 carbohydrate | o | tt |
| *araF* | CDS | L-arabinose transporter subunit -!- periplasmic-binding component of ABC superfamily | Periplasmic | COG1879;ABC-type sugar transport system, periplasmic component | GO:0016052 carbohydrate catabolism | o | tt |
| *araG* | CDS | fused L-arabinose transporter subunits -!- ATP-binding components of ABC superfamily | Cytoplasmic | COG1129;ABC-type sugar transport system, ATPase component | GO:0016052 carbohydrate catabolism | gg | o |
| *araH* | CDS | fused L-arabinose transporter subunits -!- membrane components of ABC superfamily | Integral Membrane Protein | | GO:0016052 carbohydrate catabolism | o | tt |
| *araJ* | CDS | predicted transporter | Integral Membrane Protein | | GO:0016052 carbohydrate | o | tt |
| *arcA* | CDS | DNA-binding response regulator in two-component regulatory system with ArcB or CpxA | Cytoplasmic | COG0745;Response regulators consisting of a CheY-like receiver domain and a winged-helix DNA-binding domain | GO:0006350 transcription | o | tt |
| *arcB* | CDS | hybrid sensory histidine kinase in two-component regulatory system with ArcA | Integral Membrane Protein | COG0642;Signal transduction histidine kinase -!- COG0784;FOG: CheY-like receiver | GO:0009060 aerobic respiration -!- GO:0006464 protein modification | | |
| *argA* | CDS | fused acetylglutamate kinase homolog (inactive) -!- amino acid N-acetyltransferase | Cytoplasmic | COG0548;Acetylglutamate kinase -!- COG1246;N-acetylglutamate synthase and related acetyltransferases | GO:0006526 arginine biosynthesis | gg | tt |
| *argB* | CDS | acetylglutamate kinase | Cytoplasmic | COG0548;Acetylglutamate kinase | GO:0006526 arginine biosynthesis | | |
| *argC* | CDS | N-acetyl-gamma-glutamylphosphate reductase, NAD(P)-binding | Cytoplasmic | COG0002;Acetylglutamate semialdehyde dehydrogenase | GO:0006526 arginine biosynthesis | gg | t |
| *argD* | CDS | bifunctional acetylornithine aminotransferase -!- succinyldiaminopimelate aminotransferase | Cytoplasmic | COG4992;Ornithine/acetylornithine aminotransferase | GO:0006526 arginine biosynthesis -!- GO:0009089 lysine biosynthesis via diaminopimelate | gg | tt |
| *argE* | CDS | acetylornithine deacetylase | Cytoplasmic | COG0624;Acetylornithine deacetylase/Succinyl-diaminopimelate desuccinylase and related deacylases | GO:0006526 arginine biosynthesis | o | tt |
| *argF* | CDS | CP4-6 prophage; ornithine carbamoyltransferase 2, chain F | Cytoplasmic | COG0078;Ornithine carbamoyltransferase | GO:0006526 arginine biosynthesis | gg | tt |
| *argG* | CDS | argininosuccinate synthetase | Cytoplasmic | COG0137;Argininosuccinate synthase | GO:0006526 arginine biosynthesis | gg | t |
| *argH* | CDS | argininosuccinate lyase | Cytoplasmic | COG0165;Argininosuccinate lyase | GO:0006526 arginine biosynthesis | o | tt |
| *argI* | CDS | ornithine carbamoyltransferase 1 | Cytoplasmic | COG0078;Ornithine carbamoyltransferase | GO:0006526 arginine biosynthesis | gg | o |
| *argK* | CDS | membrane ATPase/protein kinase | Cytoplasmic | COG1703;Putative periplasmic protein kinase ArgK and related GTPases of G3E family | | gg | t |
| *argO* | CDS | arginine transporter | Integral Membrane Protein | COG1279;Lysine efflux permease | | gg | tt |
| *argP* | CDS | DNA-binding transcriptional activator, replication initiation inhibitor | Cytoplasmic | COG0583;Transcriptional regulator | GO:0006261 DNA dependent DNA replication -!- GO:0006350 transcription | gg | t |
| *argQ* | tRNA | tRNA-Arg(ACG) (Arginine tRNA2) | | | | | |
| *argR* | CDS | DNA-binding transcriptional dual regulator, L-arginine-binding | Cytoplasmic | COG1438;Arginine repressor | GO:0006526 arginine biosynthesis -!- GO:0006350 transcription | o | tt |
| *argS* | CDS | arginyl-tRNA synthetase | Cytoplasmic | COG0018;Arginyl-tRNA synthetase | GO:0006418 amino acid activation | o | tt |
| *argT* | CDS | lysine/arginine/ornithine transporter subunit -!- periplasmic-binding component of ABC superfamily | Periplasmic | COG0834;ABC-type amino acid transport/signal transduction systems, periplasmic component/domain | GO:0009089 lysine biosynthesis via diaminopimelate -!- GO:0006526 arginine biosynthesis | o | tt |
| *argU* | tRNA | tRNA-Arg(UCU) (Arginine tRNA4) | | | | | |
| *argV* | tRNA | tRNA-Arg(ACG) (Arginine tRNA2) | | | | | |
| *argW* | tRNA | tRNA-Arg(CCU) (Arginine tRNA5) | | | | | |
| *argX* | tRNA | tRNA-Arg(CCG) (Arginine tRNA3) | | | | | |
| *argY* | tRNA | tRNA-Arg(ACG) (Arginine tRNA2) | | | | | |
| *argZ* | tRNA | tRNA-Arg(ACG) (Arginine tRNA2) | | | | | |
| *arnT* | CDS | 4-amino-4-deoxy-L-arabinose transferase | Integral Membrane Protein | COG1807;4-amino-4-deoxy-L-arabinose transferase and related glycosyltransferases of PMT family | GO:0006950 response to stress | g | tt |
| *aroA* | CDS | 5-enolpyruvylshikimate-3-phosphate synthetase | Cytoplasmic | COG0128;5-enolpyruvylshikimate-3-phosphate synthase | GO:0009423 chorismate biosynthesis | gg | o |
| *aroB* | CDS | 3-dehydroquinate synthase | Cytoplasmic | COG0337;3-dehydroquinate synthetase | GO:0009423 chorismate biosynthesis | gg | t |
| *aroC* | CDS | chorismate synthase | Cytoplasmic | COG0082;Chorismate synthase | GO:0009423 chorismate biosynthesis | gg | tt |
| *aroD* | CDS | 3-dehydroquinate dehydratase | Cytoplasmic | COG0710;3-dehydroquinate dehydratase | GO:0009423 chorismate biosynthesis | gg | o |
| *aroE* | CDS | dehydroshikimate reductase, NAD(P)-binding | Cytoplasmic | COG0169;Shikimate 5-dehydrogenase | GO:0009423 chorismate biosynthesis | gg | t |
| *aroF* | CDS | 3-deoxy-D-arabinoheptulosonate-7-phosphate synthase, tyrosine-repressible | Cytoplasmic | COG0722;3-deoxy-D-arabino-heptulosonate 7-phosphate (DAHP) synthase | GO:0009423 chorismate biosynthesis | gg | t |
| *aroG* | CDS | 3-deoxy-D-arabino-heptulosonate-7-phosphate synthase, phenylalanine repressible | Cytoplasmic | COG0722;3-deoxy-D-arabino-heptulosonate 7-phosphate (DAHP) synthase | GO:0009094 L-phenylalanine biosynthesis | gg | t |
| *aroH* | CDS | 3-deoxy-D-arabino-heptulosonate-7-phosphate synthase, tryptophan repressible | Cytoplasmic | COG0722;3-deoxy-D-arabino-heptulosonate 7-phosphate (DAHP) synthase | GO:0000162 tryptophan biosynthesis | gg | t |
| *aroK* | CDS | shikimate kinase I | Cytoplasmic | | GO:0009423 chorismate biosynthesis | | |
| *aroL* | CDS | shikimate kinase II | Cytoplasmic | COG0703;Shikimate kinase | GO:0009423 chorismate biosynthesis | gg | o |
| *aroM* | CDS | conserved protein | Cytoplasmic | | | o | tt |
| *aroP* | CDS | aromatic amino acid transporter | Integral Membrane Protein | COG1113;Gamma-aminobutyrate permease and related permeases | GO:0009094 L-phenylalanine biosynthesis -!- GO:0006571 tyrosine biosynthesis | gg | tt |
| *arpA* | CDS | regulator of acetyl CoA synthetase | Cytoplasmic | COG0666;FOG: Ankyrin repeat | | gg | o |
| *arpB* | CDS | predicted protein, N-ter fragment (pseudogene) | Cytoplasmic | | | | |
| *arpB* | ancestral | predicted protein (pseudogene) | Cytoplasmic | | | | |
| *arpB* | CDS | predicted protein, C-ter fragment (pseudogene) | Cytoplasmic | | | g | o |
| *arsB* | CDS | arsenite/antimonite transporter | Integral Membrane Protein | | GO:0006805 xenobiotic metabolism | o | tt |
| *arsC* | CDS | arsenate reductase | Cytoplasmic | COG1393;Arsenate reductase and related proteins, glutaredoxin family | GO:0042493 response to drug | gg | o |
| *arsR* | CDS | DNA-binding transcriptional regulator | Cytoplasmic | COG0640;Predicted transcriptional regulators | GO:0006350 transcription -!- GO:0042493 response to drug | o | tt |
| *artI* | CDS | arginine transporter subunit -!- periplasmic-binding component of ABC superfamily | Periplasmic | COG0834;ABC-type amino acid transport/signal transduction systems, periplasmic component/domain | GO:0006542 glutamine biosynthesis | o | tt |

| | | | | | | | |
|---|---|---|---|---|---|---|---|
| *artJ* | CDS | arginine transporter subunit -!- periplasmic-binding component of ABC superfamily | Periplasmic | COG0834;ABC-type amino acid transport/signal transduction systems, periplasmic component/domain | GO:0006526 arginine biosynthesis | o | tt |
| *artM* | CDS | arginine transporter subunit -!- membrane component of ABC superfamily | Integral Membrane Protein | COG4160;ABC-type arginine/histidine transport system, permease component | GO:0006526 arginine biosynthesis -!- GO:0009063 amino acid catabolism | o | tt |
| *artP* | CDS | arginine transporter subunit -!- ATP-binding component of ABC superfamily | Cytoplasmic | COG4161;ABC-type arginine transport system, ATPase component | GO:0006526 arginine biosynthesis | o | t |
| *artQ* | CDS | arginine transporter subunit -!-membrane component of ABC superfamily | Integral Membrane Protein | COG4215;ABC-type arginine transport system, permease component | GO:0006526 arginine biosynthesis | o | tt |
| *ascB* | CDS | cryptic 6-phospho-beta-glucosidase | Cytoplasmic | COG2723;Beta-glucosidase/6-phospho-beta-glucosidas e/beta-galactosidase | GO:0016052 carbohydrate catabolism | gg | t |
| *ascF* | CDS | fused cellobiose/arbutin/salicin-specific enzyme IIBC component of PTS | Integral Membrane Protein | COG1264;Phosphotransferase system IIB components -!- COG1263;Phosphotransferase system IIC components, glucose/maltose/N-acetylglucosamine-specific | GO:0016052 carbohydrate catabolism | | |
| *ascG* | CDS | DNA-binding transcriptional regulator | Cytoplasmic | COG1609;Transcriptional regulators | GO:0016052 carbohydrate catabolism -!- GO:0006350 | | |
| *asd* | CDS | aspartate-semialdehyde dehydrogenase, NAD(P)-binding | Cytoplasmic | COG0136;Aspartate-semialdehyde dehydrogenase | GO:0009089 lysine biosynthesis via diaminopimelate -!- GO:0009088 threonine biosynthesis -!- GO:0009086 methionine biosynthesis -!- GO:0009090 homoserine biosynthesis | g | t |
| *aslA* | CDS | acrylsulfatase-like enzyme | Periplasmic | COG3119;Arylsulfatase A and related enzymes | GO:0006790 sulfur metabolism | o | tt |
| *aslB* | CDS | predicted regulator of arylsulfatase activity | Cytoplasmic | | GO:0006790 sulfur metabolism -!- GO:0006457 protein folding | | |
| *asmA* | CDS | predicted assembly protein | Membrane Anchored | COG2982;Uncharacterized protein involved in outer membrane biogenesis | GO:0009103 lipopolysaccharide biosynthesis | o | tt |
| *asnA* | CDS | asparagine synthetase A | Cytoplasmic | COG2502;Asparagine synthetase A | GO:0009063 amino acid catabolism -!- GO:0006529 asparagine biosynthesis | gg | o |
| *asnB* | CDS | asparagine synthetase B | Cytoplasmic | COG0367;Asparagine synthase (glutamine-hydrolyzing) | GO:0009063 amino acid catabolism -!- GO:0006529 asparagine biosynthesis | gg | t |
| *asnC* | CDS | DNA-binding transcriptional dual regulator | Cytoplasmic | COG1522;Transcriptional regulators | GO:0006529 asparagine biosynthesis -!- GO:0006350 | gg | t |
| *asnS* | CDS | asparaginyl tRNA synthetase | Cytoplasmic | COG0017;Aspartyl/asparaginyl-tRNA synthetases | GO:0006418 amino acid activation | gg | t |
| *asnT* | tRNA | tRNA-Asn(GUU) (Asparagine tRNA) | | | | | |
| *asnU* | tRNA | tRNA-Asn(GUU) (Asparagine tRNA) | | | | | |
| *asnV* | tRNA | tRNA-Asn(GUU) (Asparagine tRNA) | | | | | |
| *asnW* | tRNA | tRNA-Asn(GUU) (Asparagine tRNA) | | | | | |
| *aspA* | CDS | aspartate ammonia-lyase | Cytoplasmic | | GO:0006520 amino acid metabolism -!- GO:0006529 asparagine biosynthesis -!- GO:0009089 lysine biosynthesis via diaminopimelate | o | tt |
| *aspC* | CDS | aspartate aminotransferase, PLP-dependent | Cytoplasmic | COG1448;Aspartate/tyrosine/aromatic aminotransferase | GO:0009063 amino acid catabolism -!- GO:0006532 aspartate | gg | o |
| *aspS* | CDS | aspartyl-tRNA synthetase | Cytoplasmic | COG0173;Aspartyl-tRNA synthetase | GO:0006418 amino acid activation | gg | t |
| *aspT* | tRNA | tRNA-Asp(GUC) (Aspartate tRNA1) | | | | | |
| *aspU* | tRNA | tRNA-Asp(GUC) (Aspartate tRNA1) | | | | | |
| *aspV* | tRNA | tRNA-Asp(GUC) (Aspartate tRNA1) | | | | | |
| *asr* | CDS | acid shock-inducible periplasmic protein | Periplasmic | | GO:0009268 response to pH | | |
| *astA* | CDS | arginine succinyltransferase | Cytoplasmic | COG3138;Arginine/ornithine N-succinyltransferase beta subunit | GO:0009063 amino acid catabolism | gg | t |
| *astB* | CDS | succinylarginine dihydrolase | Cytoplasmic | COG3724;Succinylarginine dihydrolase | GO:0009063 amino acid catabolism | o | t |
| *astC* | CDS | succinylornithine transaminase, PLP-dependent | Cytoplasmic | COG4992;Ornithine/acetylornithine aminotransferase | GO:0009063 amino acid catabolism | gg | o |
| *astD* | CDS | succinylglutamic semialdehyde dehydrogenase | Cytoplasmic | COG1012;NAD-dependent aldehyde dehydrogenases | GO:0009063 amino acid catabolism | | |
| *astE* | CDS | succinylglutamate desuccinylase | Cytoplasmic | COG2988;Succinylglutamate desuccinylase | GO:0009063 amino acid catabolism | gg | o |
| *atoA* | CDS | acetyl-CoA:acetoacetyl-CoA transferase, beta | Cytoplasmic | COG2057;Acyl CoA:acetate/3-ketoacid CoA transferase, beta subunit | GO:0019395 fatty acid oxidation | gg | o |
| *atoB* | CDS | acetyl-CoA acetyltransferase | Cytoplasmic | COG0183;Acetyl-CoA acetyltransferase | GO:0019395 fatty acid oxidation | o | tt |
| *atoC* | CDS | fused response regulator of ato opeon, in two-component system with AtoS | Cytoplasmic | COG2204;Response regulator containing CheY-like receiver, AAA-type ATPase, and DNA-binding domains | GO:0019395 fatty acid oxidation -!- GO:0006596 polyamine biosynthesis -!- GO:0006350 transcription | gg | t |
| *atoD* | CDS | acetyl-CoA:acetoacetyl-CoA transferase, alpha | Cytoplasmic | COG1788;Acyl CoA:acetate/3-ketoacid CoA transferase, alpha subunit | GO:0019395 fatty acid oxidation | gg | t |
| *atoE* | CDS | short chain fatty acid transporter | Integral Membrane Protein | COG2031;Short chain fatty acids transporter | GO:0019395 fatty acid oxidation | o | tt |
| *atoS* | CDS | sensory histidine kinase in two-component regulatory system with AtoC | Integral Membrane Protein | COG1012;NAD-dependent aldehyde dehydrogenases -!- COG0642;Signal transduction histidine kinase | GO:0019395 fatty acid oxidation -!- GO:0006464 protein modification | o | tt |
| *atpA* | CDS | F1 sector of membrane-bound ATP synthase, alpha subunit | Cytoplasmic | COG0056;F0F1-type ATP synthase, alpha subunit | GO:0015986 ATP synthesis coupled proton transport | gg | o |
| *atpB* | CDS | F0 sector of membrane-bound ATP synthase, subunit a | Integral Membrane Protein | COG0356;F0F1-type ATP synthase, subunit a | GO:0015986 ATP synthesis coupled proton transport -!- GO:0006457 protein folding | o | tt |
| *atpC* | CDS | F1 sector of membrane-bound ATP synthase, epsilon subunit | Cytoplasmic | COG0355;F0F1-type ATP synthase, epsilon subunit (mitochondrial delta subunit) | GO:0015986 ATP synthesis coupled proton transport | gg | t |
| *atpD* | CDS | F1 sector of membrane-bound ATP synthase, beta subunit | Cytoplasmic | COG0055;F0F1-type ATP synthase, beta subunit | GO:0015986 ATP synthesis coupled proton transport | g | t |
| *atpE* | CDS | F0 sector of membrane-bound ATP synthase, subunit c | Integral Membrane Protein | COG0636;F0F1-type ATP synthase, subunit c/Archaeal/vacuolar-type H+-ATPase, subunit K | GO:0015986 ATP synthesis coupled proton transport | o | tt |
| *atpF* | CDS | F0 sector of membrane-bound ATP synthase, subunit b | Membrane Anchored | COG0711;F0F1-type ATP synthase, subunit b | GO:0015986 ATP synthesis coupled proton transport | o | tt |
| *atpG* | CDS | F1 sector of membrane-bound ATP synthase, gamma subunit | Cytoplasmic | COG0224;F0F1-type ATP synthase, gamma subunit | GO:0015986 ATP synthesis coupled proton transport | gg | tt |
| *atpH* | CDS | F1 sector of membrane-bound ATP synthase, delta subunit | Cytoplasmic | COG0712;F0F1-type ATP synthase, delta subunit (mitochondrial oligomycin sensitivity protein) | GO:0015986 ATP synthesis coupled proton transport | o | o |
| *atpI* | CDS | ATP synthase, membrane-bound accesory subunit | Integral Membrane Protein | COG3312;F0F1-type ATP synthase, subunit I | GO:0015986 ATP synthesis coupled proton transport | | |
| *avtA* | CDS | valine-pyruvate aminotransferase | Cytoplasmic | | GO:0006523 alanine biosynthesis | | |
| *azoR* | CDS | NADH-azoreductase, FMN-dependent | Cytoplasmic | COG1182;Acyl carrier protein phosphodiesterase | | gg | o |
| *bacA* | CDS | undecaprenyl pyrophosphate phosphatase | Integral Membrane Protein | COG1968;Uncharacterized bacitracin resistance protein | GO:0042493 response to drug | o | tt |
| *baeR* | CDS | DNA-binding response regulator in two-component regulatory system with BaeS | Cytoplasmic | COG0745;Response regulators consisting of a CheY-like receiver domain and a winged-helix DNA-binding domain | GO:0006350 transcription | gg | o |
| *baeS* | CDS | sensory histidine kinase in two-component regulatory system with BaeR | Integral Membrane Protein | COG0642;Signal transduction histidine kinase | GO:0006464 protein modification | o | tt |
| *barA* | CDS | hybrid sensory histidine kinase, in two-component regulatory system with UvrY | Integral Membrane Protein | COG0642;Signal transduction histidine kinase -!- COG4999;Uncharacterized domain of BarA-like signal transduction histidine kinases -!- COG2201;Chemotaxis response regulator containing a CheY-like receiver domain and a methylesterase domain -!- | GO:0006464 protein modification | o | tt |
| *basR* | CDS | DNA-binding response regulator in two-component regulatory system with BasS | Cytoplasmic | COG0745;Response regulators consisting of a CheY-like receiver domain and a winged-helix DNA-binding domain | GO:0006350 transcription | o | tt |
| *basS* | CDS | sensory histidine kinase in two-component regulatory system with BasR | Integral Membrane Protein | COG0642;Signal transduction histidine kinase | GO:0006464 protein modification | o | tt |
| *bax* | CDS | conserved protein | Periplasmic | | | | |
| *bcp* | CDS | thiol peroxidase, thioredoxin-dependent | Cytoplasmic | COG1225;Peroxiredoxin | GO:0006805 xenobiotic metabolism | gg | t |
| *bcr* | CDS | bicyclomycin/multidrug efflux system | Integral Membrane Protein | COG0477;Permeases of the major facilitator superfamily | GO:0042493 response to drug | | |
| *bcsA* | CDS | cellulose synthase, catalytic subunit | Integral Membrane Protein | | | | |
| *bcsB* | CDS | regulator of cellulose synthase, cyclic di-GMP binding | Integral Membrane Protein | | | o | tt |
| *bcsC* | CDS | cellulose synthase subunit | Periplasmic | | | | |
| *bcsE* | CDS | conserved protein | Cytoplasmic | | | gg | t |
| *bcsF* | CDS | predicted protein | Membrane Anchored | | | | |
| *bcsG* | CDS | predicted inner membrane protein | Integral Membrane Protein | | | o | tt |
| *bcsZ* | CDS | endo-1,4-D-glucanase | Outer Membrane Lipoprotein | COG3405;Endoglucanase Y | GO:0016052 carbohydrate catabolism -!- GO:0000272 polysaccharide catabolism | o | tt |
| *bdm* | CDS | biofilm-dependent modulation protein | Cytoplasmic | | | | |
| *betA* | CDS | choline dehydrogenase, a flavoprotein | Cytoplasmic | COG2303;Choline dehydrogenase and related flavoproteins | GO:0006578 betaine biosynthesis -!- GO:0006970 response to osmotic stress | gg | t |
| *betB* | CDS | betaine aldehyde dehydrogenase, NAD-dependent | Cytoplasmic | COG1012;NAD-dependent aldehyde dehydrogenases | GO:0006578 betaine biosynthesis -!- GO:0006970 response to osmotic stress | gg | t |
| *betI* | CDS | DNA-binding transcriptional repressor | Cytoplasmic | | GO:0006350 transcription -!- GO:0006970 response to osmotic stress | o | o |
| *betT* | CDS | choline transporter of high affinity | Integral Membrane Protein | COG1292;Choline-glycine betaine transporter | GO:0006578 betaine biosynthesis | o | tt |
| *bfd* | CDS | bacterioferritin-associated ferredoxin | Cytoplasmic | COG2906;Bacterioferritin-associated ferredoxin | GO:0006826 iron ion transport | gg | t |
| *bfr* | CDS | bacterioferritin, iron storage and detoxification protein | Cytoplasmic | COG2193;Bacterioferritin (cytochrome b1) | GO:0006826 iron ion transport | gg | tt |
| *bglA* | CDS | 6-phospho-beta-glucosidase A | Cytoplasmic | COG2723;Beta-glucosidase/6-phospho-beta-glucosidas e/beta-galactosidase | GO:0016052 carbohydrate catabolism | gg | o |
| *bglB* | CDS | cryptic phospho-beta-glucosidase B | Cytoplasmic | COG2723;Beta-glucosidase/6-phospho-beta-glucosidas e/beta-galactosidase | GO:0016052 carbohydrate catabolism | gg | t |
| *bglF* | CDS | fused beta-glucoside-specific PTS enzyme IIABC components | Integral Membrane Protein | COG1264;Phosphotransferase system IIB components -!- COG1263;Phosphotransferase system IIC components, glucose/maltose/N-acetylglucosamine-specific -!- COG2190;Phosphotransferase system IIA components | GO:0016052 carbohydrate catabolism -!- GO:0006464 protein modification | o | tt |
| *bglG* | CDS | transcriptional antiterminator of the bgl operon | Cytoplasmic | COG3711;Transcriptional antiterminator | GO:0016052 carbohydrate catabolism -!- GO:0006350 | gg | t |
| *bglH* | CDS | carbohydrate-specific outer membrane porin, cryptic | Outer Membrane B-barrel protein | COG4580;Maltoporin (phage lambda and maltose receptor) | | o | tt |
| *bglJ* | CDS | DNA-binding transcriptional regulator | Cytoplasmic | COG2197;Response regulator containing a CheY-like receiver domain and an HTH DNA-binding domain | GO:0016052 carbohydrate catabolism -!- GO:0006350 | | |
| *bglX* | CDS | beta-D-glucoside glucohydrolase, periplasmic | Periplasmic | COG1472;Beta-glucosidase-related glycosidases | GO:0016052 carbohydrate | g | tt |
| *bioA* | CDS | 7,8-diaminopelargonic acid synthase, PLP- | Cytoplasmic | COG0161;Adenosylmethionine-8-amino-7-oxononanoate aminotransferase | GO:0009102 biotin biosynthesis | gg | t |
| *bioB* | CDS | biotin synthase | Cytoplasmic | COG0502;Biotin synthase and related enzymes | GO:0009102 biotin biosynthesis | gg | t |
| *bioC* | CDS | predicted methltransferase, enzyme of biotin | Cytoplasmic | COG0500;SAM-dependent methyltransferases | GO:0009102 biotin biosynthesis | g | t |

| | | | | | | | |
|---|---|---|---|---|---|---|---|
| *bioD* | CDS | dethiobiotin synthetase | Cytoplasmic | COG0132;Dethiobiotin synthetase | GO:0009102 biotin biosynthesis | g | t |
| *bioF* | CDS | 8-amino-7-oxononanoate synthase | Cytoplasmic | COG0156;7-keto-8-aminopelargonate synthetase and related enzymes | GO:0009102 biotin biosynthesis | g | t |
| *bioH* | CDS | carboxylesterase of pimeloyl-CoA synthesis | Cytoplasmic | COG0596;Predicted hydrolases or acyltransferases (alpha/beta hydrolase superfamily) | GO:0009102 biotin biosynthesis | gg | tt |
| *bipA* | CDS | GTP-binding protein | Cytoplasmic | | | | |
| *birA* | CDS | bifunctional biotin-[acetylCoA carboxylase] holoenzyme synthetase -!- DNA-binding transcriptional repressor, bio-5'-AMP-binding | Cytoplasmic | COG1654;Biotin operon repressor -!- COG0340;Biotin-(acetyl-CoA carboxylase) ligase | GO:0009102 biotin biosynthesis -!- GO:0006350 transcription | gg | t |
| *bisC* | CDS | biotin sulfoxide reductase | Cytoplasmic | COG0243;Anaerobic dehydrogenases, typically selenocysteine-containing | GO:0009102 biotin biosynthesis | | |
| *blc* | CDS | outer membrane lipoprotein (lipocalin) | Outer Membrane Lipoprotein | COG3040;Bacterial lipocalin | | o | tt |
| *blr* | CDS | beta-lactam resistance membrane protein | Integral Membrane Protein | | | | |
| *bolA* | CDS | regulator of penicillin binding proteins and beta lactamase transcription (morphogene) | Cytoplasmic | | GO:0006970 response to osmotic stress | | |
| *borD* | CDS | DLP12 prophage; predicted lipoprotein | Outer Membrane Lipoprotein | | | o | tt |
| *brnQ* | CDS | predicted branched chain amino acid transporter (LIV-II) | Integral Membrane Protein | COG1114;Branched-chain amino acid permeases | GO:0009098 leucine biosynthesis -!- GO:0009063 amino acid catabolism | o | tt |
| *btuB* | CDS | vitamin B12/cobalamin outer membrane transporter | Outer Membrane B-barrel protein | COG4206;Outer membrane cobalamin receptor protein | | | |
| *btuC* | CDS | vitamin B12 transporter subunit -!- membrane component of ABC superfamily | Integral Membrane Protein | COG4139;ABC-type cobalamin transport system, permease component | GO:0009236 vitamin B12 biosynthesis | o | tt |
| *btuD* | CDS | vitamin B12 transporter subunit -!- ATP-binding component of ABC superfamily | Cytoplasmic | COG4138;ABC-type cobalamin transport system, ATPase component | GO:0009236 vitamin B12 biosynthesis | g | t |
| *btuE* | CDS | predicted glutathione peroxidase | Cytoplasmic | COG0386;Glutathione peroxidase | GO:0009236 vitamin B12 | gg | t |
| *btuF* | CDS | vitamin B12 transporter subunit -!- periplasmic-binding component of ABC superfamily | Periplasmic | COG0614;ABC-type Fe3+-hydroxamate transport system, periplasmic component | | o | tt |
| *btuR* | CDS | cob(I)alamin adenolsyltransferase/cobinamide ATP-dependent adenolsyltransferase | Cytoplasmic | COG2109;ATP:corrinoid adenosyltransferase | GO:0009236 vitamin B12 biosynthesis | gg | t |
| *cadA* | CDS | lysine decarboxylase 1 | Cytoplasmic | COG1982;Arginine/lysine/ornithine decarboxylases | GO:0009063 amino acid catabolism -!- GO:0009089 lysine biosynthesis via diaminopimelate | g | t |
| *cadB* | CDS | predicted lysine/cadaverine transporter | Integral Membrane Protein | COG0531;Amino acid transporters | GO:0009089 lysine biosynthesis via diaminopimelate | o | tt |
| *cadC* | CDS | DNA-binding transcriptional activator | Membrane Anchored | COG3710;DNA-binding winged-HTH domains | GO:0009063 amino acid catabolism -!- GO:0006350 transcription | o | tt |
| *caiA* | CDS | crotonobetaine reductase subunit II, FAD-binding | Cytoplasmic | COG1960;Acyl-CoA dehydrogenases | GO:0006097 glyoxylate cycle -!- GO:0009310 amine catabolism -!- GO:0042413 carnitine catabolism | gg | t |
| *caiB* | CDS | crotonobetainyl CoA:carnitine CoA transferase | Cytoplasmic | COG1804;Predicted acyl-CoA transferases/carnitine dehydratase | GO:0042413 carnitine catabolism -!- GO:0009310 amine catabolism | gg | o |
| *caiC* | CDS | predicted crotonobetaine CoA ligase:carnitine CoA ligase | Cytoplasmic | | GO:0009310 amine catabolism | gg | o |
| *caiD* | CDS | crotonobetainyl CoA hydratase | Cytoplasmic | COG1024;Enoyl-CoA hydratase/carnithine racemase | GO:0009310 amine catabolism -!- GO:0042413 carnitine catabolism | o | tt |
| *caiE* | CDS | predicted acyl transferase | Cytoplasmic | COG0663;Carbonic anhydrases/acetyltransferases, isoleucine patch superfamily | GO:0009310 amine catabolism -!- GO:0042413 carnitine catabolism | | |
| *caiF* | CDS | DNA-binding transcriptional activator | Cytoplasmic | | GO:0009310 amine catabolism -!- GO:0006350 transcription | gg | o |
| *caiT* | CDS | predicted transporter | Integral Membrane Protein | COG1292;Choline-glycine betaine transporter | GO:0016052 carbohydrate | o | tt |
| *can* | CDS | carbonic anhydrase | Cytoplasmic | COG0288;Carbonic anhydrase | | o | t |
| *carA* | CDS | carbamoyl phosphate synthetase small subunit, glutamine amidotransferase | Cytoplasmic | COG0505;Carbamoylphosphate synthase small subunit | GO:0006526 arginine biosynthesis -!- GO:0006221 pyrimidine nucleotide biosynthesis | gg | o |
| *carB* | CDS | carbamoyl-phosphate synthase large subunit | Cytoplasmic | COG0458;Carbamoylphosphate synthase large subunit (split gene in MJ) | GO:0006526 arginine biosynthesis -!- GO:0006221 pyrimidine nucleotide biosynthesis | o | tt |
| *cbl* | CDS | DNA-binding transcriptional activator | Cytoplasmic | COG0583;Transcriptional regulator | GO:0019344 cysteine biosynthesis -!- GO:0006790 sulfur metabolism -!- GO:0006350 transcription | gg | t |
| *cbpA* | CDS | curved DNA-binding protein, DnaJ homologue that functions as a co-chaperone of DnaK | Cytoplasmic | COG2214;DnaJ-class molecular chaperone | GO:0006457 protein folding | gg | t |
| *cbpM* | CDS | modulator of CbpA co-chaperone | Cytoplasmic | | | gg | o |
| *cbrA* | CDS | predicted oxidoreductase, FAD/NAD(P)-binding domain | Cytoplasmic | COG0644;Dehydrogenases (flavoproteins) | | | |
| *cbrC* | CDS | conserved protein | Cytoplasmic | COG3196;Uncharacterized protein conserved in bacteria | | gg | tt |
| *cca* | CDS | fused tRNA nucleotidyl transferase -!- 2'3'-cyclic phosphodiesterase/2'nucleotidase/phosphatase | Cytoplasmic | COG0617;tRNA nucleotidyltransferase/poly(A) polymerase | GO:0009451 RNA modification | g | t |
| *cchA* | CDS | predicted carboxysome structural protein, ethanolamine utilization protein | Cytoplasmic | COG4577;Carbon dioxide concentrating mechanism/carboxysome shell protein | GO:0006805 xenobiotic metabolism | gg | o |
| *cchB* | CDS | predicted carboxysome structural protein, ethanolamine utilization protein | Cytoplasmic | COG4576;Carbon dioxide concentrating mechanism/carboxysome shell protein | GO:0006805 xenobiotic metabolism | gg | o |
| *ccmA* | CDS | heme exporter subunit -!- ATP-binding component of ABC superfamily | Cytoplasmic | COG4133;ABC-type transport system involved in cytochrome c biogenesis, ATPase component | GO:0017004 cytochrome biogenesis -!- GO:0006457 protein folding | | |
| *ccmB* | CDS | heme exporter subunit -!- membrane component of ABC superfamily | Integral Membrane Protein | COG2386;ABC-type transport system involved in cytochrome c biogenesis, permease component | GO:0017004 cytochrome biogenesis -!- GO:0006457 protein folding | o | tt |
| *ccmC* | CDS | heme exporter subunit -!- membrane component of ABC superfamily | Integral Membrane Protein | COG0755;ABC-type transport system involved in cytochrome c biogenesis, permease component | GO:0017004 cytochrome biogenesis -!- GO:0006457 protein folding | o | tt |
| *ccmD* | CDS | cytochrome c biogenesis protein | Membrane Anchored | COG3114;Heme exporter protein D | GO:0017004 cytochrome biogenesis -!- GO:0006457 protein folding | o | tt |
| *ccmE* | CDS | periplasmic heme chaperone | Periplasmic | COG2332;Cytochrome c-type biogenesis protein CcmE | GO:0017004 cytochrome biogenesis -!- GO:0006457 protein folding | o | tt |
| *ccmF* | CDS | heme lyase, CcmF subunit | Integral Membrane Protein | COG1138;Cytochrome c biogenesis factor | GO:0017004 cytochrome biogenesis | o | tt |
| *ccmG* | CDS | periplasmic thioredoxin of cytochrome c-type biogenesis | Membrane Anchored | COG0526;Thiol-disulfide isomerase and thioredoxins | GO:0017004 cytochrome biogenesis -!- GO:0006457 protein folding | o | tt |
| *ccmH* | CDS | heme lyase, CcmH subunit | Integral Membrane Protein | COG3088;Uncharacterized protein involved in biosynthesis of c-type cytochromes -!- COG4235;Cytochrome c biogenesis factor | GO:0017004 cytochrome biogenesis -!- GO:0006457 protein folding | o | tt |
| *cdaR* | CDS | DNA-binding transcriptional regulator | Cytoplasmic | | | | |
| *cdd* | CDS | cytidine/deoxycytidine deaminase | Cytoplasmic | COG0295;Cytidine deaminase | GO:0015949 nucleobase, nucleoside and nucleotide interconversion | g | t |
| *cdh* | CDS | CDP-diacylglycerol phosphotidylhydrolase | Membrane Anchored | COG2134;CDP-diacylglycerol pyrophosphatase | | o | tt |
| *cdsA* | CDS | CDP-diglyceride synthase | Integral Membrane Protein | COG0575;CDP-diglyceride synthetase | GO:0008654 phospholipid biosynthesis | | |
| *cedA* | CDS | cell division modulator | Cytoplasmic | | | gg | o |
| *cfa* | CDS | cyclopropane fatty acyl phospholipid synthase (unsaturated-phospholipid methyltransferase) | Cytoplasmic | COG2230;Cyclopropane fatty acid synthase and related methyltransferases | | gg | t |
| *chaA* | CDS | calcium/sodium:proton antiporter | Integral Membrane Protein | COG0387;Ca2+/H+ antiporter | | o | tt |
| *chaB* | CDS | predicted cation regulator | Cytoplasmic | COG4572;Putative cation transport regulator | GO:0006350 transcription | gg | o |
| *chaC* | CDS | regulatory protein for cation transport | Cytoplasmic | COG3703;Uncharacterized protein involved in cation transport | GO:0006350 transcription | gg | o |
| *chbA* | CDS | N,N'-diacetylchitobiose-specific enzyme IIA component of PTS | Cytoplasmic | COG1447;Phosphotransferase system cellobiose-specific component IIA | GO:0016052 carbohydrate catabolism | o | o |
| *chbB* | CDS | N,N'-diacetylchitobiose-specific enzyme IIB component of PTS | Periplasmic | COG1440;Phosphotransferase system cellobiose-specific component IIB | GO:0016052 carbohydrate catabolism | gg | o |
| *chbC* | CDS | N,N'-diacetylchitobiose-specific enzyme IIC component of PTS | Integral Membrane Protein | COG1455;Phosphotransferase system cellobiose-specific component IIC | GO:0016052 carbohydrate catabolism | gg | tt |
| *chbF* | CDS | cryptic phospho-beta-glucosidase, NAD(P)-binding | Cytoplasmic | COG1486;Alpha-galactosidases/6-phospho-beta-glucos idases, family 4 of glycosyl hydrolases | GO:0016052 carbohydrate catabolism -!- GO:0009435 nicotinamide adenine dinucleotide | gg | t |
| *chbG* | CDS | conserved protein | Cytoplasmic | COG3394;Uncharacterized protein conserved in bacteria | | o | tt |
| *chbR* | CDS | DNA-binding transcriptional dual regulator | Cytoplasmic | COG2207;AraC-type DNA-binding domain-containing proteins | GO:0016052 carbohydrate catabolism -!- GO:0006350 | o | tt |
| *cheA* | CDS | fused chemotactic sensory histidine kinase (soluble) in two-component regulatory system with CheB and CheY | Cytoplasmic | COG0643;Chemotaxis protein histidine kinase and related kinases | GO:0006464 protein modification -!- GO:0042330 taxis | gg | tt |
| *cheB* | CDS | fused chemotaxis regulator -!- protein-glutamate methylesterase, in two-component regulatory system with CheA | Cytoplasmic | COG2201;Chemotaxis response regulator containing a CheY-like receiver domain and a methylesterase domain | GO:0006464 protein modification -!- GO:0042330 taxis | gg | t |
| *cheR* | CDS | chemotaxis regulator | Cytoplasmic | COG1352;Methylase of chemotaxis methyl-accepting proteins | GO:0006464 protein modification -!- GO:0042330 taxis | g | t |
| *cheW* | CDS | purine-binding chemotaxis protein | Cytoplasmic | COG0835;Chemotaxis signal transduction protein | GO:0042330 taxis | gg | t |
| *cheY* | CDS | chemotaxis regulator transmitting signal to flagellar motor component | Cytoplasmic | COG0784;FOG: CheY-like receiver | GO:0042330 taxis | gg | o |
| *cheZ* | CDS | chemotaxis regulator | Cytoplasmic | COG3143;Chemotaxis protein | GO:0006464 protein modification -!- GO:0042330 taxis | gg | tt |
| *chiA* | CDS | periplasmic endochitinase | Periplasmic | | GO:0009057 macromolecule catabolism | g | tt |
| *chpA* | CDS | toxin of the ChpA-ChpR toxin-antitoxin system, endoribonuclease | Cytoplasmic | COG2337;Growth inhibitor | | o | tt |
| *chpB* | CDS | toxin of the ChpB-ChpS toxin-antitoxin system | Cytoplasmic | COG2337;Growth inhibitor | | o | tt |
| *chpR* | CDS | antitoxin of the ChpA-ChpR toxin-antitoxin system | Cytoplasmic | COG2336;Growth regulator | | gg | tt |
| *chpS* | CDS | antitoxin of the ChpB-ChpS toxin-antitoxin system | Cytoplasmic | COG2336;Growth regulator | | | |
| *cirA* | CDS | ferric iron-catecholate outer membrane transporter | Outer Membrane B-barrel protein | COG4771;Outer membrane receptor for ferrienterochelin and colicins | | g | t |
| *citA* | CDS | sensory histidine kinase in two-component regulatory system with citB | Integral Membrane Protein | COG3290;Signal transduction histidine kinase regulating citrate/malate metabolism | GO:0006464 protein modification | o | tt |
| *citB* | CDS | DNA-binding response regulator in two-component regulatory system with citA | Cytoplasmic | COG4565;Response regulator of citrate/malate metabolism | GO:0006350 transcription | o | tt |
| *citC* | CDS | citrate lyase synthetase | Cytoplasmic | | GO:0006464 protein modification | gg | t |
| *citD* | CDS | citrate lyase, acyl carrier (gamma) subunit | Cytoplasmic | COG3052;Citrate lyase, gamma subunit | GO:0042967 acyl-carrier protein biosynthesis | gg | t |
| *citE* | CDS | citrate lyase, citryl-ACP lyase (beta) subunit | Cytoplasmic | | | g | t |
| *citF* | CDS | citrate lyase, citrate-ACP transferase (alpha) subunit | Cytoplasmic | COG3051;Citrate lyase, alpha subunit | | | |

| | | | | | | | |
|---|---|---|---|---|---|---|---|
| *citG* | CDS | triphosphoribosyl-dephospho-CoA transferase | Cytoplasmic | COG1767;Triphosphoribosyl-dephospho-CoA synthetase | GO:0006412 protein biosynthesis | g | t |
| *citT* | CDS | citrate:succinate antiporter | Integral Membrane Protein | COG0471;Di- and tricarboxylate transporters | GO:0016052 carbohydrate | o | tt |
| *citX* | CDS | apo-citrate lyase phosphoribosyl-dephospho-CoA transferase | Cytoplasmic | COG3697;Phosphoribosyl-dephospho-CoA transferase (holo-ACP synthetase) | GO:0006464 protein modification | gg | t |
| *clcA* | CDS | chloride channel, voltage-gated | Integral Membrane Protein | COG0038;Chloride channel protein EriC | | | |
| *clcB* | CDS | predicted voltage-gated chloride channel | Integral Membrane Protein | COG0038;Chloride channel protein EriC | | | |
| *cld* | CDS | regulator of length of O-antigen component of lipopolysaccharide chains | Integral Membrane Protein | | GO:0009243 O antigen biosynthesis | | |
| *clpA* | CDS | ATPase and specificity subunit of ClpA-ClpP ATP-dependent serine protease, chaperone activity | Cytoplasmic | COG0542;ATPases with chaperone activity, ATP-binding subunit | | gg | tt |
| *clpB* | CDS | protein disaggregation chaperone | Cytoplasmic | COG0542;ATPases with chaperone activity, ATP-binding subunit | GO:0006457 protein folding | gg | tt |
| *clpP* | CDS | proteolytic subunit of ClpA-ClpP and ClpX-ClpP ATP-dependent serine proteases | Cytoplasmic | COG0740;Protease subunit of ATP-dependent Clp proteases | GO:0009266 response to temperature | gg | o |
| *clpS* | CDS | regulatory protein for ClpA substrate specificity | Cytoplasmic | COG2127;Uncharacterized conserved protein | | gg | o |
| *clpX* | CDS | ATPase and specificity subunit of ClpX-ClpP ATP-dependent serine protease | Cytoplasmic | COG1219;ATP-dependent protease Clp, ATPase subunit | GO:0006457 protein folding | o | tt |
| *cls* | CDS | cardiolipin synthase 1 | Integral Membrane Protein | COG1502;Phosphatidylserine/phosphatidylglycerophos phate/cardiolipin synthases and related enzymes | GO:0008654 phospholipid biosynthesis -!- GO:0042493 response to drug | o | tt |
| *cmk* | CDS | cytidylate kinase | Cytoplasmic | COG0283;Cytidylate kinase | GO:0015949 nucleobase, nucleoside and nucleotide interconversion | gg | o |
| *cmr* | CDS | multidrug efflux system protein | Integral Membrane Protein | COG0477;Permeases of the major facilitator superfamily | GO:0042493 response to drug | o | tt |
| *cmtA* | CDS | predicted fused mannitol-specific enzyme IIBC component of PTS | Integral Membrane Protein | COG2213;Phosphotransferase system, mannitol-specific IIBC component | GO:0016052 carbohydrate catabolism | o | tt |
| *cmtB* | CDS | predicted mannitol-specific enzyme IIA component of PTS | Cytoplasmic | COG1762;Phosphotransferase system mannitol/fructose-specific IIA domain (Ntr-type) | GO:0016052 carbohydrate catabolism | gg | o |
| *coaA* | CDS | pantothenate kinase | Cytoplasmic | COG1072;Panthothenate kinase | GO:0015937 coenzyme A biosynthesis | gg | tt |
| *coaD* | CDS | pantetheine-phosphate adenylyltransferase | Cytoplasmic | COG0669;Phosphopantetheine adenylyltransferase | | gg | o |
| *coaE* | CDS | dephospho-CoA kinase | Periplasmic | COG0237;Dephospho-CoA kinase | | gg | t |
| *cobB* | CDS | deacetylase of acetyl-CoA synthetase, NAD-dependent | Cytoplasmic | COG0846;NAD-dependent protein deacetylases, SIR2 family | GO:0006355 regulation of transcription, DNA-dependent | o | tt |
| *cobC* | CDS | predicted alpha-ribazole-5'-P phosphatase | Cytoplasmic | COG0406;Fructose-2,6-bisphosphatase | GO:0009236 vitamin B12 | gg | t |
| *cobS* | CDS | cobalamin 5'-phosphate synthase | Integral Membrane Protein | COG0368;Cobalamin-5-phosphate synthase | GO:0009236 vitamin B12 | o | tt |
| *cobT* | CDS | nicotinate-nucleotide dimethylbenzimidazole-P phophoribosyl transferase | Cytoplasmic | COG2038;NaMN:DMB phosphoribosyltransferase | GO:0009236 vitamin B12 biosynthesis | gg | o |
| *cobU* | CDS | bifunctional cobinamide kinase -!- cobinamide phosphate guanylyltransferase | Cytoplasmic | COG2087;Adenosyl cobinamide kinase/adenosyl cobinamide phosphate guanylyltransferase | GO:0009236 vitamin B12 biosynthesis | g | t |
| *codA* | CDS | cytosine deaminase | Cytoplasmic | COG0402;Cytosine deaminase and related metal-dependent hydrolases | GO:0015949 nucleobase, nucleoside and nucleotide interconversion | o | tt |
| *codB* | CDS | cytosine transporter | Integral Membrane Protein | COG1457;Purine-cytosine permease and related proteins | GO:0015949 nucleobase, nucleoside and nucleotide interconversion | o | tt |
| *cof* | CDS | thiamin pyrimidine pyrophosphate hydrolase | Cytoplasmic | COG0561;Predicted hydrolases of the HAD superfamily | | gg | o |
| *copA* | CDS | copper transporter | Integral Membrane Protein | COG2217;Cation transport ATPase | | o | tt |
| *corA* | CDS | magnesium/nickel/cobalt transporter | Integral Membrane Protein | COG0598;Mg2+ and Co2+ transporters | | o | tt |
| *cpdA* | CDS | cyclic 3',5'-adenosine monophosphate phosphodiesterase | Cytoplasmic | COG1409;Predicted phosphohydrolases | GO:0016052 carbohydrate catabolism | gg | t |
| *cpdB* | CDS | 2':3'-cyclic-nucleotide 2'-phosphodiesterase | Periplasmic | COG0737;5'-nucleotidase/2',3'-cyclic phosphodiesterase and related esterases | GO:0015949 nucleobase, nucleoside and nucleotide interconversion | o | tt |
| *cpsB* | CDS | mannose-1-phosphate guanyltransferase | Cytoplasmic | COG0836;Mannose-1-phosphate guanylyltransferase -!- COG0662;Mannose-6-phosphate isomerase | GO:0009242 colanic acid biosynthesis -!- GO:0016052 carbohydrate catabolism -!- GO:0009226 nucleotide-sugar biosynthesis -!- GO:0009269 | gg | t |
| *cpsG* | CDS | phosphomannomutase | Cytoplasmic | COG1109;Phosphomannomutase | GO:0009226 nucleotide-sugar biosynthesis -!- GO:0009242 colanic acid biosynthesis -!- GO:0009269 response to dessication | gg | t |
| *cpxA* | CDS | sensory histidine kinase in two-component regulatory system with CpxR | Integral Membrane Protein | COG0642;Signal transduction histidine kinase | GO:0006464 protein modification -!- GO:0006805 xenobiotic metabolism | o | tt |
| *cpxP* | CDS | periplasmic protein combats stress | Periplasmic | | GO:0006350 transcription -!- GO:0006805 xenobiotic metabolism | | |
| *cpxR* | CDS | DNA-binding response regulator in two-component regulatory system with CpxA | Cytoplasmic | COG0745;Response regulators consisting of a CheY-like receiver domain and a winged-helix DNA-binding domain | GO:0006350 transcription -!- GO:0006805 xenobiotic metabolism | o | tt |
| *crcA* | CDS | palmitoyl transferase for Lipid A | Outer Membrane B-barrel protein | | | o | t |
| *crcB* | CDS | predicted inner membrane protein associated with chromosome condensation | Integral Membrane Protein | COG0239;Integral membrane protein possibly involved in chromosome condensation | | o | tt |
| *creA* | CDS | conserved protein | Periplasmic | COG3045;Uncharacterized protein conserved in bacteria | | o | tt |
| *creB* | CDS | DNA-binding response regulator in two-component regulatory system with CreC | Cytoplasmic | COG0745;Response regulators consisting of a CheY-like receiver domain and a winged-helix DNA-binding domain | GO:0006350 transcription | o | t |
| *creC* | CDS | sensory histidine kinase in two-component regulatory system with CreB or PhoB, regulator of the CreBC regulon | Integral Membrane Protein | COG0642;Signal transduction histidine kinase | GO:0006464 protein modification | o | tt |
| *creD* | CDS | inner membrane protein | Integral Membrane Protein | COG4452;Inner membrane protein involved in colicin E2 resistance | | o | tt |
| *crl* | CDS | DNA-binding transcriptional regulator | Cytoplasmic | | GO:0006350 transcription | gg | o |
| *crp* | CDS | DNA-binding transcriptional dual regulator | Cytoplasmic | COG0664;cAMP-binding proteins - catabolite gene activator and regulatory subunit of cAMP-dependent protein kinases | GO:0006350 transcription | | |
| *crr* | CDS | glucose-specific enzyme IIA component of PTS | Cytoplasmic | COG2190;Phosphotransferase system IIA components | GO:0016052 carbohydrate | gg | o |
| *csdA* | CDS | cysteine sulfinate desulfinase | Cytoplasmic | COG0520;Selenocysteine lyase | GO:0006790 sulfur metabolism -!- GO:0006520 amino acid metabolism | gg | tt |
| *csgA* | CDS | cryptic curlin major subunit | Extracellular | | GO:0009101 glycoprotein biosynthesis | o | tt |
| *csgB* | CDS | curlin nucleator protein, minor subunit in curli complex | Extracellular | | GO:0009101 glycoprotein biosynthesis | o | tt |
| *csgC* | CDS | predicted curli production protein | Periplasmic | | | o | tt |
| *csgD* | CDS | DNA-binding transcriptional activator in two-component regulatory system | Cytoplasmic | COG2771;DNA-binding HTH domain-containing proteins | GO:0006350 transcription | o | tt |
| *csgE* | CDS | predicted transport protein | Periplasmic |  |  | o | tt |
| *csgF* | CDS | predicted transport protein | Periplasmic | | | o | tt |
| *csgG* | CDS | outer membrane lipoprotein | Outer Membrane Lipoprotein | COG1462;Uncharacterized protein involved in formation of curli polymers | | o | tt |
| *csiE* | CDS | stationary phase inducible protein | Cytoplasmic | COG3711;Transcriptional antiterminator | | | |
| *csiR* | CDS | DNA-binding transcriptional dual regulator | Cytoplasmic | | GO:0006350 transcription | gg | t |
| *cspA* | CDS | major cold shock protein | Cytoplasmic | COG1278;Cold shock proteins | GO:0006350 transcription -!- GO:0009266 response to | gg | tt |
| *cspB* | CDS | Qin prophage; cold shock protein | Cytoplasmic | COG1278;Cold shock proteins | | gg | t |
| *cspC* | CDS | stress protein, member of the CspA-family | Cytoplasmic | COG1278;Cold shock proteins | GO:0006350 transcription -!- GO:0009266 response to | gg | t |
| *cspD* | CDS | cold shock protein homolog | Cytoplasmic | COG1278;Cold shock proteins | | gg | tt |
| *cspE* | CDS | DNA-binding transcriptional repressor | Cytoplasmic | COG1278;Cold shock proteins | GO:0006350 transcription -!- GO:0009266 response to | gg | tt |
| *cspF* | CDS | Qin prophage; cold shock protein | Cytoplasmic | COG1278;Cold shock proteins | | o | t |
| *cspG* | CDS | DNA-binding transcriptional regulator | Cytoplasmic | COG1278;Cold shock proteins | GO:0009266 response to | o | tt |
| *cspH* | CDS | stress protein, member of the CspA-family | Cytoplasmic | COG1278;Cold shock proteins | GO:0009266 response to | | |
| *cspI* | CDS | Qin prophage; cold shock protein | Cytoplasmic | COG1278;Cold shock proteins | | gg | tt |
| *csrA* | CDS | pleiotropic regulatory protein for carbon source metabolism | Cytoplasmic | COG1551;Carbon storage regulator (could also regulate swarming and quorum sensing) | GO:0016052 carbohydrate catabolism -!- GO:0006096 glycolysis -!- GO:0006094 gluconeogenesis -!- GO:0006401 RNA catabolism -!- GO:0009386 translational attenuation -!- GO:0042330 taxis | gg | o |
| *csrB* | misc_RNA | regulatory sRNA | | | | | |
| *csrC* | misc_RNA | regulatory sRNA | | | | | |
| *cstA* | CDS | carbon starvation protein | Integral Membrane Protein | COG1966;Carbon starvation protein, predicted membrane protein | GO:0042594 response to starvation | o | tt |
| *cueO* | CDS | multicopper oxidase (laccase) | Periplasmic | COG2132;Putative multicopper oxidases | | gg | tt |
| *cueR* | CDS | DNA-binding transcriptional activator | Cytoplasmic | COG0789;Predicted transcriptional regulators | GO:0006350 transcription | gg | t |
| *cusA* | CDS | copper/silver efflux system, membrane component | Integral Membrane Protein | COG3696;Putative silver efflux pump | | o | tt |
| *cusB* | CDS | copper/silver efflux system, membrane fusion protein | Membrane Anchored | COG0845;Membrane-fusion protein | GO:0042493 response to drug | o | tt |
| *cusC* | CDS | copper/silver efflux system, outer membrane component | Outer Membrane Lipoprotein | COG1538;Outer membrane protein | | o | tt |
| *cusF* | CDS | periplasmic copper-binding protein | Periplasmic | | | o | tt |
| *cusR* | CDS | DNA-binding response regulator in two-component regulatory system with CusS | Cytoplasmic | COG0745;Response regulators consisting of a CheY-like receiver domain and a winged-helix DNA-binding domain | GO:0006350 transcription | gg | tt |
| *cusS* | CDS | sensory histidine kinase in two-component regulatory system with CusR, senses copper ions | Integral Membrane Protein | COG0642;Signal transduction histidine kinase | GO:0006464 protein modification | | |
| *cutA* | CDS | copper binding protein, copper sensitivity | Cytoplasmic | COG1324;Uncharacterized protein involved in tolerance to divalent cations | GO:0017004 cytochrome biogenesis | gg | o |
| *cutC* | CDS | copper homeostasis protein | Cytoplasmic | | GO:0006805 xenobiotic metabolism | gg | t |
| *cvpA* | CDS | membrane protein required for colicin V production | Integral Membrane Protein | COG1286;Uncharacterized membrane protein, required for colicin V production | | o | tt |
| *cvrA* | CDS | predicted cation/proton antiporter | Integral Membrane Protein | COG3263;NhaP-type Na+/H+ and K+/H+ antiporters with a unique C-terminal domain | | | |
| *cyaA* | CDS | adenylate cyclase | Cytoplasmic | COG3072;Adenylate cyclase | | o | tt |
| *cyaY* | CDS | frataxin, iron-binding and oxidizing protein | Cytoplasmic | COG1965;Protein implicated in iron transport, frataxin homolog | | gg | o |
| *cybB* | CDS | cytochrome b561 | Integral Membrane Protein | COG3038;Cytochrome B561 | GO:0017004 cytochrome biogenesis | | |
| *cybC* | CDS | cytochrome b562, truncated (pseudogene) | Cytoplasmic | | GO:0017004 cytochrome biogenesis | gg | o |

| | | | | | | | |
|---|---|---|---|---|---|---|---|
| *cycA* | CDS | D-alanine/D-serine/glycine transporter | Integral Membrane Protein | COG1113;Gamma-aminobutyrate permease and related permeases | GO:0009063 amino acid catabolism -!- GO:0009252 peptidoglycan biosynthesis -!- GO:0006545 glycine biosynthesis -!- GO:0006523 alanine biosynthesis -!- GO:0006164 purine nucleotide biosynthesis -!- GO:0009257 10-formyltetrahydrofolate biosynthesis -!- | o | tt |
| *cydA* | CDS | cytochrome d terminal oxidase, subunit I | Integral Membrane Protein | COG1271;Cytochrome bd-type quinol oxidase, subunit 1 | | o | tt |
| *cydB* | CDS | cytochrome d terminal oxidase, subunit II | Integral Membrane Protein | COG1294;Cytochrome bd-type quinol oxidase, subunit 2 | | o | tt |
| *cydC* | CDS | fused cysteine transporter subunits -!- membrane component and ATP-binding component of ABC superfamily | Integral Membrane Protein | COG4987;ABC-type transport system involved in cytochrome bd biosynthesis, fused ATPase and permease components | | o | tt |
| *cydD* | CDS | fused cysteine transporter subunits -!- membrane component and ATP-binding component of ABC superfamily | Integral Membrane Protein | COG4988;ABC-type transport system involved in cytochrome bd biosynthesis, ATPase and permease components | | o | tt |
| *cynR* | CDS | DNA-binding transcriptional dual regulator | Cytoplasmic | COG0583;Transcriptional regulator | GO:0006350 transcription -!- GO:0006805 xenobiotic metabolism | | |
| *cynS* | CDS | cyanate aminohydrolase | Cytoplasmic | COG1513;Cyanate lyase | GO:0009440 cyanate catabolism -!- GO:0006807 nitrogen metabolism -!- GO:0006805 xenobiotic metabolism | gg | t |
| *cynT* | CDS | carbonic anhydrase | Cytoplasmic | COG0288;Carbonic anhydrase | GO:0009440 cyanate catabolism -!- GO:0006805 xenobiotic metabolism | gg | t |
| *cynX* | CDS | predicted cyanate transporter | Integral Membrane Protein | COG2807;Cyanate permease | GO:0009440 cyanate catabolism -!- GO:0006805 xenobiotic metabolism | o | tt |
| *cyoA* | CDS | cytochrome o ubiquinol oxidase subunit II | Integral Membrane Protein | COG1622;Heme/copper-type cytochrome/quinol oxidases, subunit 2 | GO:0009060 aerobic respiration | o | tt |
| *cyoB* | CDS | cytochrome o ubiquinol oxidase subunit I | Integral Membrane Protein | COG0843;Heme/copper-type cytochrome/quinol oxidases, subunit 1 | GO:0009060 aerobic respiration | o | tt |
| *cyoC* | CDS | cytochrome o ubiquinol oxidase subunit III | Integral Membrane Protein | COG1845;Heme/copper-type cytochrome/quinol oxidase, subunit 3 | GO:0009060 aerobic respiration | o | tt |
| *cyoD* | CDS | cytochrome o ubiquinol oxidase subunit IV | Integral Membrane Protein | COG3125;Heme/copper-type cytochrome/quinol oxidase, subunit 4 | GO:0009060 aerobic respiration | o | tt |
| *cyoE* | CDS | protoheme IX farnesyltransferase | Integral Membrane Protein | COG0109;Polyprenyltransferase (cytochrome oxidase assembly factor) | GO:0009060 aerobic respiration | o | tt |
| *cysA* | CDS | sulfate/thiosulfate transporter subunit -!- ATP-binding component of ABC superfamily | Cytoplasmic | COG1118;ABC-type sulfate/molybdate transport systems, ATPase component | GO:0006790 sulfur metabolism -!- GO:0042493 response to drug | gg | t |
| *cysB* | CDS | DNA-binding transcriptional dual regulator, O-acetyl-L-serine-binding | Cytoplasmic | COG0583;Transcriptional regulator | GO:0019344 cysteine biosynthesis -!- GO:0006350 transcription | gg | tt |
| *cysC* | CDS | adenosine 5'-phosphosulfate kinase | Cytoplasmic | COG0529;Adenylylsulfate kinase and related kinases | GO:0006790 sulfur metabolism | gg | t |
| *cysD* | CDS | sulfate adenylyltransferase, subunit 2 | Cytoplasmic | COG0175;3'-phosphoadenosine 5'-phosphosulfate sulfotransferase (PAPS reductase)/FAD synthetase and related enzymes | GO:0006790 sulfur metabolism | gg | t |
| *cysE* | CDS | serine acetyltransferase | Cytoplasmic | COG1045;Serine acetyltransferase | GO:0019344 cysteine biosynthesis | gg | tt |
| *cysG* | CDS | fused siroheme synthase 1,3-dimethyluroporphyriongen III dehydrogenase/siroheme ferrochelatase -!- | Cytoplasmic | COG1648;Siroheme synthase (precorrin-2 oxidase/ferrochelatase domain) -!- COG0007;Uroporphyrinogen-III methylase | | o | tt |
| *cysH* | CDS | 3'-phosphoadenosine 5'-phosphosulfate reductase | Cytoplasmic | COG0175;3'-phosphoadenosine 5'-phosphosulfate sulfotransferase (PAPS reductase)/FAD synthetase and related enzymes | GO:0006790 sulfur metabolism | gg | t |
| *cysI* | CDS | sulfite reductase, beta subunit, NAD(P)-binding | Cytoplasmic | COG0155;Sulfite reductase, beta subunit (hemoprotein) | GO:0006790 sulfur metabolism | o | tt |
| *cysJ* | CDS | sulfite reductase, alpha subunit, flavoprotein | Cytoplasmic | COG0369;Sulfite reductase, alpha subunit (flavoprotein) | GO:0006790 sulfur metabolism | o | tt |
| *cysK* | CDS | cysteine synthase A, O-acetylserine sulfhydrolase A subunit | Cytoplasmic | COG0031;Cysteine synthase | GO:0019344 cysteine biosynthesis | gg | t |
| *cysM* | CDS | cysteine synthase B (O-acetylserine sulfhydrolase B) | Cytoplasmic | COG0031;Cysteine synthase | GO:0019344 cysteine biosynthesis | gg | t |
| *cysN* | CDS | sulfate adenylyltransferase, subunit 1 | Cytoplasmic | COG2895;GTPases - Sulfate adenylate transferase subunit 1 | GO:0006790 sulfur metabolism | gg | tt |
| *cysP* | CDS | thiosulfate transporter subunit -!- periplasmic-binding component of ABC superfamily | Periplasmic | COG4150;ABC-type sulfate transport system, periplasmic component | GO:0006790 sulfur metabolism | o | tt |
| *cysQ* | CDS | PAPS (adenosine 3'-phosphate 5'-phosphosulfate) 3'(2'),5'-bisphosphate nucleotidase | Cytoplasmic | COG1218;3'-Phosphoadenosine 5'-phosphosulfate (PAPS) 3'-phosphatase | GO:0006790 sulfur metabolism | gg | tt |
| *cysS* | CDS | cysteinyl-tRNA synthetase | Cytoplasmic | COG0215;Cysteinyl-tRNA synthetase | GO:0006418 amino acid activation | gg | tt |
| *cysT* | tRNA | tRNA-Cys(GCA) (Cysteine tRNA) | | | | | |
| *cysU* | CDS | sulfate/thiosulfate transporter subunit -!- membrane component of ABC superfamily | Integral Membrane Protein | COG0555;ABC-type sulfate transport system, permease component | GO:0006790 sulfur metabolism | o | tt |
| *cysW* | CDS | sulfate/thiosulfate transporter subunit -!- membrane component of ABC superfamily | Integral Membrane Protein | | GO:0006790 sulfur metabolism | gg | t |
| *cysZ* | CDS | predicted inner membrane protein | Integral Membrane Protein | COG2981;Uncharacterized protein involved in cysteine biosynthesis | GO:0006790 sulfur metabolism | g | tt |
| *cytR* | CDS | DNA-binding transcriptional dual regulator | Cytoplasmic | COG1609;Transcriptional regulators | GO:0015949 nucleobase, nucleoside and nucleotide interconversion -!- GO:0006350 transcription | o | tt |
| *dacA* | CDS | D-alanyl-D-alanine carboxypeptidase (penicillin-binding protein 5) | Periplasmic | COG1686;D-alanyl-D-alanine carboxypeptidase | GO:0042493 response to drug | o | tt |
| *dacB* | CDS | D-alanyl-D-alanine carboxypeptidase | Periplasmic | COG2027;D-alanyl-D-alanine carboxypeptidase (penicillin-binding protein 4) | GO:0009252 peptidoglycan biosynthesis -!- GO:0042493 response to drug | o | tt |
| *dacC* | CDS | D-alanyl-D-alanine carboxypeptidase (penicillin-binding protein 6a) | Periplasmic | COG1686;D-alanyl-D-alanine carboxypeptidase | GO:0042493 response to drug | o | tt |
| *dacD* | CDS | D-alanyl-D-alanine carboxypeptidase (penicillin-binding protein 6b) | Periplasmic | COG1686;D-alanyl-D-alanine carboxypeptidase | | | |
| *dadA* | CDS | D-amino acid dehydrogenase | Cytoplasmic | COG0665;Glycine/D-amino acid oxidases (deaminating) | GO:0009063 amino acid catabolism | o | tt |
| *dadX* | CDS | alanine racemase 2, PLP-binding | Cytoplasmic | COG0787;Alanine racemase | GO:0009063 amino acid catabolism -!- GO:0006523 alanine biosynthesis | o | tt |
| *dam* | CDS | DNA adenine methylase | Cytoplasmic | COG0338;Site-specific DNA methylase | GO:0006306 DNA methylation | o | tt |
| *damX* | CDS | predicted protein | Membrane Anchored | COG3266;Uncharacterized protein conserved in bacteria | | o | tt |
| *dapA* | CDS | dihydrodipicolinate synthase | Cytoplasmic | COG0329;Dihydrodipicolinate synthase/N-acetylneuraminate lyase | GO:0009089 lysine biosynthesis via diaminopimelate | gg | o |
| *dapB* | CDS | dihydrodipicolinate reductase | Cytoplasmic | COG0289;Dihydrodipicolinate reductase | GO:0009089 lysine biosynthesis via diaminopimelate | gg | o |
| *dapD* | CDS | 2,3,4,5-tetrahydropyridine-2-carboxylate N-succinyltransferase | Cytoplasmic | COG2171;Tetrahydrodipicolinate N-succinyltransferase | GO:0009089 lysine biosynthesis via diaminopimelate | gg | tt |
| *dapE* | CDS | N-succinyl-diaminopimelate deacylase | Cytoplasmic | COG0624;Acetylornithine deacetylase/Succinyl-diaminopimelate desuccinylase and related deacylases | GO:0009089 lysine biosynthesis via diaminopimelate | g | tt |
| *dapF* | CDS | diaminopimelate epimerase | Cytoplasmic | COG0253;Diaminopimelate epimerase | GO:0009089 lysine biosynthesis via diaminopimelate | | |
| *dbpA* | CDS | ATP-dependent RNA helicase specific for 23S rRNA | Cytoplasmic | COG0513;Superfamily II DNA and RNA helicases | GO:0009451 RNA modification | gg | t |
| *dcd* | CDS | 2'-deoxycytidine 5'-triphosphate deaminase | Cytoplasmic | COG0717;Deoxycytidine deaminase | GO:0015949 nucleobase, nucleoside and nucleotide interconversion | gg | o |
| *dcm* | CDS | DNA cytosine methylase | Cytoplasmic | COG0270;Site-specific DNA methylase | GO:0006306 DNA methylation | gg | tt |
| *dcp* | CDS | dipeptidyl carboxypeptidase II | Cytoplasmic | COG0339;Zn-dependent oligopeptidases | | gg | t |
| *dcrB* | CDS | periplasmic protein | Inner Membrane Lipoprotein | | | | |
| *dctA* | CDS | C4-dicarboxylic acid, orotate and citrate transporter | Integral Membrane Protein | COG1301;Na+/H+-dicarboxylate symporters | GO:0016052 carbohydrate | o | tt |
| *dcuA* | CDS | C4-dicarboxylate antiporter | Integral Membrane Protein | COG2704;Anaerobic C4-dicarboxylate transporter | GO:0016052 carbohydrate | | |
| *dcuB* | CDS | C4-dicarboxylate antiporter | Integral Membrane Protein | COG2704;Anaerobic C4-dicarboxylate transporter | GO:0016052 carbohydrate | o | tt |
| *dcuC* | CDS | anaerobic C4-dicarboxylate transport | Integral Membrane Protein | COG3069;C4-dicarboxylate transporter | GO:0016052 carbohydrate | | |
| *dcuC* | CDS | anaerobic C4-dicarboxylate transport | Integral Membrane Protein | | | o | tt |
| *dcuC* | CDS | anaerobic C4-dicarboxylate transport | Integral Membrane Protein | | | o | tt |
| *dcuD* | CDS | predicted transporter | Integral Membrane Protein | COG3069;C4-dicarboxylate transporter | | o | tt |
| *dcuR* | CDS | DNA-binding response regulator in two-component regulatory system with DcuS | Cytoplasmic | COG4565;Response regulator of citrate/malate metabolism | GO:0009061 anaerobic respiration -!- GO:0006350 transcription | o | o |
| *dcuS* | CDS | sensory histidine kinase in two-component regulatory system with DcuR, regulator of anaerobic fumarate respiration | Integral Membrane Protein | COG3290;Signal transduction histidine kinase regulating citrate/malate metabolism | GO:0009061 anaerobic respiration -!- GO:0006464 protein modification | o | tt |
| *ddg* | CDS | palmitoleoyl-acyl carrier protein (ACP)-dependent acyltransferase | Membrane Anchored | | GO:0009245 lipid A biosynthesis -!- GO:0009266 response to | gg | tt |
| *ddlA* | CDS | D-alanine-D-alanine ligase A | Cytoplasmic | COG1181;D-alanine-D-alanine ligase and related ATP-grasp enzymes | GO:0009252 peptidoglycan biosynthesis | gg | tt |
| *ddlB* | CDS | D-Alanine:D-alanine ligase | Cytoplasmic | COG1181;D-alanine-D-alanine ligase and related ATP-grasp enzymes | GO:0009252 peptidoglycan biosynthesis | gg | o |
| *ddpA* | CDS | D-Ala-D-Ala transporter subunit -!- periplasmic-binding component of ABC superfamily | Periplasmic | COG0747;ABC-type dipeptide transport system, periplasmic component | | | |
| *ddpB* | CDS | D-Ala-D-Ala transporter subunit -!- membrane component of ABC superfamily | Integral Membrane Protein | COG0601;ABC-type dipeptide/oligopeptide/nickel transport systems, permease components | | o | tt |
| *ddpC* | CDS | D-Ala-D-Ala transporter subunit -!- membrane component of ABC superfamily | Integral Membrane Protein | COG1173;ABC-type dipeptide/oligopeptide/nickel transport systems, permease components | | o | tt |
| *ddpD* | CDS | D-Ala-D-Ala transporter subunit -!- ATP-binding component of ABC superfamily | Cytoplasmic | COG0444;ABC-type dipeptide/oligopeptide/nickel transport system, ATPase component | | gg | o |
| *ddpF* | CDS | D-Ala-D-Ala transporter subunit -!- ATP-binding component of ABC superfamily | Cytoplasmic | COG1124;ABC-type dipeptide/oligopeptide/nickel transport system, ATPase component | | g | t |
| *ddpX* | CDS | D-Ala-D-Ala dipeptidase, Zn-dependent | Cytoplasmic | COG2173;D-alanyl-D-alanine dipeptidase | GO:0006970 response to osmotic stress | gg | t |
| *deaD* | CDS | ATP-dependent RNA helicase | Cytoplasmic | COG0513;Superfamily II DNA and RNA helicases | GO:0009451 RNA modification | | |
| *dedA* | CDS | conserved inner membrane protein | Integral Membrane Protein | COG0586;Uncharacterized membrane-associated protein | | o | tt |
| *dedD* | CDS | conserved protein | Membrane Anchored | COG3147;Uncharacterized protein conserved in bacteria | | | |
| *def* | CDS | peptide deformylase | Cytoplasmic | COG0242;N-formylmethionyl-tRNA deformylase | GO:0006464 protein modification | gg | t |
| *degP* | CDS | serine endoprotease (protease Do), membrane-associated | Periplasmic | COG0265;Trypsin-like serine proteases, typically periplasmic, contain C-terminal PDZ domain | GO:0009266 response to temperature | o | tt |
| *degQ* | CDS | serine endoprotease, periplasmic | Membrane Anchored | COG0265;Trypsin-like serine proteases, typically periplasmic, contain C-terminal PDZ domain | | o | tt |
| *degS* | CDS | serine endoprotease, periplasmic | Membrane Anchored | COG0265;Trypsin-like serine proteases, typically periplasmic, contain C-terminal PDZ domain | GO:0006508 proteolysis and peptidolysis | o | tt |
| *deoA* | CDS | thymidine phosphorylase | Cytoplasmic | COG0213;Thymidine phosphorylase | GO:0015949 nucleobase, nucleoside and nucleotide interconversion | gg | t |
| *deoB* | CDS | phosphopentomutase | Cytoplasmic | COG1015;Phosphopentomutase | GO:0015949 nucleobase, nucleoside and nucleotide interconversion | gg | t |
| *deoC* | CDS | 2-deoxyribose-5-phosphate aldolase, NAD(P)-linked | Cytoplasmic | COG0274;Deoxyribose-phosphate aldolase | GO:0015949 nucleobase, nucleoside and nucleotide interconversion | gg | o |

| | | | | | | | |
|---|---|---|---|---|---|---|---|
| *deoD* | CDS | purine-nucleoside phosphorylase | Cytoplasmic | COG0813;Purine-nucleoside phosphorylase | GO:0015949 nucleobase, nucleoside and nucleotide interconversion | g | o |
| *deoR* | CDS | DNA-binding transcriptional repressor | Cytoplasmic | COG1349;Transcriptional regulators of sugar metabolism | GO:0015949 nucleobase, nucleoside and nucleotide interconversion -!- GO:0006350 transcription | gg | t |
| *der* | CDS | predicted GTP-binding protein | Cytoplasmic | COG1160;Predicted GTPases | | | |
| *dfp* | CDS | fused 4'-phosphopantothenoylcysteine decarboxylase -!- phosphopantothenoylcysteine synthetase, FMN-binding | Cytoplasmic | | GO:0015937 coenzyme A biosynthesis -!- GO:0006261 DNA dependent DNA replication | | |
| *dgkA* | CDS | diacylglycerol kinase | Integral Membrane Protein | COG0818;Diacylglycerol kinase | GO:0008654 phospholipid biosynthesis | o | tt |
| *dgoA* | CDS | 2-oxo-3-deoxygalactonate 6-phosphate aldolase | Cytoplasmic | | GO:0016052 carbohydrate | | |
| *dgoD* | CDS | galactonate dehydratase | Cytoplasmic | | GO:0016052 carbohydrate | | |
| *dgoK* | CDS | 2-oxo-3-deoxygalactonate kinase | Cytoplasmic | COG3734;2-keto-3-deoxy-galactonokinase | GO:0016052 carbohydrate | g | t |
| *dgoR* | CDS | predicted DNA-binding transcriptional regulator | Cytoplasmic | | GO:0016052 carbohydrate catabolism -!- GO:0006350 | | |
| *dgoT* | CDS | D-galactonate transporter | Integral Membrane Protein | | GO:0016052 carbohydrate | | |
| *dgsA* | CDS | DNA-binding transcriptional repressor | Cytoplasmic | COG1940;Transcriptional regulator/sugar kinase | GO:0000271 polysaccharide biosynthesis -!- GO:0006350 transcription | o | tt |
| *dgt* | CDS | deoxyguanosine triphosphate triphosphohydrolase | Cytoplasmic | COG0232;dGTP triphosphohydrolase | GO:0015949 nucleobase, nucleoside and nucleotide interconversion | gg | t |
| *dhaH* | CDS | fused predicted dihydroxyacetone-specific PTS enzyme HPr component -!- EI component | Cytoplasmic | COG3412;Uncharacterized protein conserved in bacteria -!- COG1080;Phosphoenolpyruvate-protein kinase (PTS system EI component in bacteria) | | | |
| *dhaK* | CDS | dihydroxyacetone kinase, N-terminal domain | Cytoplasmic | | | | |
| *dhaL* | CDS | dihydroxyacetone kinase, C-terminal domain | Cytoplasmic | COG2376;Dihydroxyacetone kinase | | | |
| *dhaR* | CDS | predicted DNA-binding transcriptional regulator, dihydroxyacetone | Cytoplasmic | COG3284;Transcriptional activator of acetoin/glycerol metabolism | GO:0006350 transcription | | |
| *dicA* | CDS | Qin prophage; predicted regulator for DicB | Cytoplasmic | COG1396;Predicted transcriptional regulators | | o | tt |
| *dicB* | CDS | Qin prophage; cell division inhibition protein | Cytoplasmic | | | o | tt |
| *dicC* | CDS | Qin prophage; DNA-binding transcriptional regulator | Cytoplasmic | | | gg | o |
| *dicF* | misc_RNA | Qin prophage; DicF antisense RNA | | | | | |
| *dinB* | CDS | DNA polymerase IV | Cytoplasmic | COG0389;Nucleotidyltransferase/DNA polymerase involved in DNA repair | GO:0006261 DNA dependent DNA replication -!- GO:0009432 SOS response | o | tt |
| *dinD* | CDS | DNA-damage-inducible protein | Cytoplasmic | | GO:0006281 DNA repair | gg | o |
| *dinF* | CDS | DNA-damage-inducible SOS response protein | Integral Membrane Protein | COG0534;Na+-driven multidrug efflux pump | GO:0006281 DNA repair -!- GO:0009314 response to radiation -!- GO:0009432 SOS response | o | tt |
| *dinG* | CDS | ATP-dependent DNA helicase | Cytoplasmic | COG1199;Rad3-related DNA helicases | GO:0006281 DNA repair -!- GO:0009432 SOS response | o | tt |
| *dinI* | CDS | DNA damage-inducible protein I | Cytoplasmic | | GO:0006281 DNA repair -!- GO:0009432 SOS response | g | o |
| *dinJ* | CDS | predicted antitoxin of YafQ-DinJ toxin-antitoxin | Cytoplasmic | COG3077;DNA-damage-inducible protein J | GO:0006281 DNA repair | gg | t |
| *dipZ* | CDS | fused thiol:disulfide interchange protein | Integral Membrane Protein | COG4232;Thiol:disulfide interchange protein | GO:0006790 sulfur metabolism -!- GO:0017004 cytochrome biogenesis | | |
| *djlA* | CDS | DnaJ-like protein, membrane anchored | Membrane Anchored | COG1076;DnaJ-domain-containing proteins 1 | GO:0006457 protein folding | o | tt |
| *djlB* | CDS | predicted chaperone | Membrane Anchored | | GO:0006457 protein folding | o | tt |
| *djlC* | CDS | Hsc56 co-chaperone of HscC | Cytoplasmic | | | o | tt |
| *dkgA* | CDS | 2,5-diketo-D-gluconate reductase A | Cytoplasmic | | GO:0016052 carbohydrate | | |
| *dkgB* | CDS | 2,5-diketo-D-gluconate reductase B | Cytoplasmic | COG0656;Aldo/keto reductases, related to diketogulonate reductase | GO:0016052 carbohydrate | gg | o |
| *dksA* | CDS | transcriptional regulator of rRNA transcription, DnaK suppressor protein | Cytoplasmic | COG1734;DnaK suppressor protein | GO:0006457 protein folding | gg | o |
| *dld* | CDS | D-lactate dehydrogenase, FAD-binding, NADH independent | Cytoplasmic | COG0277;FAD/FMN-containing dehydrogenases | GO:0009060 aerobic respiration | gg | tt |
| *dmsA* | CDS | dimethyl sulfoxide reductase, anaerobic, subunit A | Periplasmic | | GO:0009061 anaerobic respiration | | |
| *dmsB* | CDS | dimethyl sulfoxide reductase, anaerobic, subunit B | Cytoplasmic | COG0437;Fe-S-cluster-containing hydrogenase components 1 | GO:0009061 anaerobic respiration | gg | t |
| *dmsC* | CDS | dimethyl sulfoxide reductase, anaerobic, subunit C | Integral Membrane Protein | COG3302;DMSO reductase anchor subunit | GO:0009061 anaerobic respiration | o | tt |
| *dmsD* | CDS | twin-argninine leader-binding protein for DmsA and TorA | Cytoplasmic | COG3381;Uncharacterized component of anaerobic dehydrogenases | | | |
| *dnaA* | CDS | chromosomal replication initiator protein DnaA, DNA-binding transcriptional dual regulator | Cytoplasmic | COG0593;ATPase involved in DNA replication initiation | GO:0006261 DNA dependent DNA replication | o | tt |
| *dnaB* | CDS | replicative DNA helicase | Cytoplasmic | COG0305;Replicative DNA helicase | GO:0006261 DNA dependent DNA replication | o | tt |
| *dnaC* | CDS | DNA biosynthesis protein | Cytoplasmic | COG1484;DNA replication protein | GO:0006261 DNA dependent DNA replication | o | tt |
| *dnaE* | CDS | DNA polymerase III alpha subunit | Cytoplasmic | COG0587;DNA polymerase III, alpha subunit | GO:0006261 DNA dependent DNA replication | o | tt |
| *dnaG* | CDS | DNA primase | Cytoplasmic | COG0358;DNA primase (bacterial type) | GO:0006261 DNA dependent DNA replication | gg | t |
| *dnaJ* | CDS | chaperone Hsp40, co-chaperone with DnaK | Cytoplasmic | COG0484;DnaJ-class molecular chaperone with C-terminal Zn finger domain | GO:0006457 protein folding | o | tt |
| *dnaK* | CDS | chaperone Hsp70, co-chaperone with DnaJ | Cytoplasmic | COG0443;Molecular chaperone | GO:0006457 protein folding -!- GO:0006970 response to osmotic stress | gg | t |
| *dnaN* | CDS | DNA polymerase III, beta subunit | Cytoplasmic | COG0592;DNA polymerase sliding clamp subunit (PCNA homolog) | GO:0006261 DNA dependent DNA replication | o | t |
| *dnaQ* | CDS | DNA polymerase III epsilon subunit | Cytoplasmic | COG0847;DNA polymerase III, epsilon subunit and related 3'-5' exonucleases | GO:0006261 DNA dependent DNA replication -!- GO:0006457 protein folding | gg | t |
| *dnaT* | CDS | DNA biosynthesis protein (primosomal protein I) | Cytoplasmic | | GO:0006261 DNA dependent DNA replication | o | t |
| *dnaX* | CDS | DNA polymerase III/DNA elongation factor III, tau and gamma subunits | Cytoplasmic | COG2812;DNA polymerase III, gamma/tau subunits | GO:0006261 DNA dependent DNA replication | o | tt |
| *dos* | CDS | cAMP phosphodiesterase, heme-regulated | Cytoplasmic | COG2202;FOG: PAS/PAC domain -!- COG2199;FOG: GGDEF domain -!- COG2200;FOG: EAL domain | | o | tt |
| *dppA* | CDS | dipeptide transporter -!- periplasmic-binding component of ABC superfamily | Periplasmic | COG0747;ABC-type dipeptide transport system, periplasmic component | GO:0009063 amino acid catabolism | o | tt |
| *dppB* | CDS | dipeptide transporter-!- membrane component of ABC superfamily | Integral Membrane Protein | COG0601;ABC-type dipeptide/oligopeptide/nickel transport systems, permease components | | o | o |
| *dppC* | CDS | dipeptide transporter -!- membrane component of ABC superfamily | Integral Membrane Protein | COG1173;ABC-type dipeptide/oligopeptide/nickel transport systems, permease components | | o | tt |
| *dppD* | CDS | dipeptide transporter -!- ATP-binding component of ABC superfamily | Cytoplasmic | COG0444;ABC-type dipeptide/oligopeptide/nickel transport system, ATPase component | GO:0009063 amino acid catabolism | g | t |
| *dppF* | CDS | dipeptide transporter -!- ATP-binding component of ABC superfamily | Cytoplasmic | COG4608;ABC-type oligopeptide transport system, ATPase component | GO:0009063 amino acid catabolism | o | tt |
| *dps* | CDS | Fe-binding and storage protein | Cytoplasmic | COG0783;DNA-binding ferritin-like protein (oxidative damage protectant) | GO:0042594 response to starvation | gg | t |
| *dsbA* | CDS | periplasmic protein disulfide isomerase I | Periplasmic | COG0526;Thiol-disulfide isomerase and thioredoxins | GO:0006457 protein folding -!- GO:0006950 response to stress | o | tt |
| *dsbB* | CDS | oxidoreductase that catalyzes reoxidation of DsbA protein disulfide isomerase I | Integral Membrane Protein | | GO:0009296 flagella biogenesis -!- GO:0006457 protein folding -!- GO:0042330 taxis | | |
| *dsbC* | CDS | protein disulfide isomerase II | Periplasmic | COG1651;Protein-disulfide isomerase | GO:0006457 protein folding | gg | tt |
| *dsbG* | CDS | periplasmic disulfide isomerase/thiol-disulphide oxidase | Periplasmic | COG1651;Protein-disulfide isomerase | GO:0006457 protein folding | o | tt |
| *dsdA* | CDS | D-serine ammonia-lyase | Cytoplasmic | COG3048;D-serine dehydratase | GO:0009063 amino acid catabolism | gg | t |
| *dsdC* | CDS | DNA-binding transcriptional dual regulator | Cytoplasmic | COG0583;Transcriptional regulator | GO:0009063 amino acid catabolism -!- GO:0006350 transcription | gg | t |
| *dsdX* | CDS | predicted transporter | Integral Membrane Protein | COG2610;H+/gluconate symporter and related permeases | | g | tt |
| *dsrA* | misc_RNA | regulatory, anti-sense RNA | | | | | |
| *dsrB* | CDS | predicted protein | Cytoplasmic | | | gg | o |
| *dtd* | CDS | D-Tyr-tRNA(Tyr) deacylase | Cytoplasmic | COG1490;D-Tyr-tRNAtyr deacylase | GO:0009451 RNA modification -!- GO:0006805 xenobiotic metabolism | o | t |
| *dusB* | CDS | tRNA-dihydrouridine synthase B | Cytoplasmic | COG0042;tRNA-dihydrouridine synthase | | g | t |
| *dusC* | CDS | tRNA-dihydrouridine synthase C | Cytoplasmic | COG0042;tRNA-dihydrouridine synthase | | gg | t |
| *dut* | CDS | deoxyuridinetriphosphatase | Cytoplasmic | COG0756;dUTPase | GO:0015949 nucleobase, nucleoside and nucleotide interconversion | gg | o |
| *dxr* | CDS | 1-deoxy-D-xylulose 5-phosphate reductoisomerase | Cytoplasmic | COG0743;1-deoxy-D-xylulose 5-phosphate reductoisomerase | | gg | t |
| *dxs* | CDS | 1-deoxyxylulose-5-phosphate synthase, thiamine-requiring, FAD-requiring | Cytoplasmic | COG1154;Deoxyxylulose-5-phosphate synthase | GO:0008615 pyridoxine biosynthesis -!- GO:0009228 thiamin biosynthesis | o | tt |
| *eaeH* | CDS | attaching and effacing protein, pathogenesis factor | Membrane Anchored | | | o | tt |
| *eamA* | CDS | cysteine and O-acetyl-L-serine efflux system | Integral Membrane Protein | | | | |
| *ebgA* | CDS | cryptic beta-D-galactosidase, alpha subunit | Cytoplasmic | | GO:0016052 carbohydrate | | |
| *ebgC* | CDS | cryptic beta-D-galactosidase, beta subunit | Cytoplasmic | COG2731;Beta-galactosidase, beta subunit | GO:0016052 carbohydrate | o | t |
| *ebgR* | CDS | DNA-binding transcriptional repressor | Cytoplasmic | COG1609;Transcriptional regulators | GO:0016052 carbohydrate catabolism -!- GO:0006350 | gg | tt |
| *ecnA* | CDS | entericidin A membrane lipoprotein, antidote entericidin B | Outer Membrane Lipoprotein | | | | |
| *ecnB* | CDS | entericidin B membrane lipoprotein | Outer Membrane Lipoprotein | | | o | o |
| *eco* | CDS | ecotin, a serine protease inhibitor | Periplasmic | COG4574;Serine protease inhibitor ecotin | | o | tt |
| *ecpD* | CDS | predicted periplasmic pilin chaperone | Periplasmic | COG3121;P pilus assembly protein, chaperone PapD | GO:0006457 protein folding | o | tt |
| *eda* | CDS | multifunctional 2-keto-3-deoxygluconate 6-phosphate aldolase -!- 2-keto-4-hydroxyglutarate aldolase -!- oxaloacetate decarboxylase | Cytoplasmic | COG0800;2-keto-3-deoxy-6-phosphogluconate aldolase | GO:0016052 carbohydrate catabolism -!- GO:0009255 Entner-Doudoroff pathway -!- GO:0009436 glyoxylate catabolism | gg | t |
| *edd* | CDS | 6-phosphogluconate dehydratase | Cytoplasmic | COG0129;Dihydroxyacid dehydratase/phosphogluconate dehydratase | GO:0009255 Entner-Doudoroff pathway | gg | o |

| | | | | | | | |
|---|---|---|---|---|---|---|---|
| *efp* | CDS | Elongation factor EF-P | Cytoplasmic | COG0231;Translation elongation factor P (EF-P)/translation initiation factor 5A (eIF-5A) | GO:0006412 protein biosynthesis | gg | o |
| *ego* | CDS | fused AI2 transporter subunits -!- ATP-binding components of ABC superfamily | Cytoplasmic | COG1129;ABC-type sugar transport system, ATPase component | | gg | t |
| *elaA* | CDS | predicted acyltransferase with acyl-CoA N-acyltransferase domain | Cytoplasmic | COG2153;Predicted acyltransferase | | gg | tt |
| *elaB* | CDS | conserved protein | Membrane Anchored | COG4575;Uncharacterized conserved protein | | g | o |
| *elaC* | CDS | binuclear zinc phosphodiesterase | Cytoplasmic | | | gg | t |
| *elaD* | CDS | predicted enzyme | Cytoplasmic | | | | |
| *elbA* | CDS | predicted protein | Cytoplasmic | | | o | o |
| *elbB* | CDS | isoprenoid biosynthesis protein with amidotransferase-like domain | Cytoplasmic | COG3155;Uncharacterized protein involved in an early stage of isoprenoid biosynthesis | | gg | t |
| *emrA* | CDS | multidrug efflux system | Membrane Anchored | COG1566;Multidrug resistance efflux pump | GO:0042493 response to drug | o | tt |
| *emrB* | CDS | multidrug efflux system protein | Integral Membrane Protein | COG0477;Permeases of the major facilitator superfamily | GO:0042493 response to drug | g | tt |
| *emrD* | CDS | multidrug efflux system protein | Integral Membrane Protein | COG0477;Permeases of the major facilitator superfamily | GO:0042493 response to drug | | |
| *emrE* | CDS | DLP12 prophage; multidrug resistance protein | Integral Membrane Protein | COG2076;Membrane transporters of cations and cationic drugs | GO:0042493 response to drug | o | tt |
| *emrK* | CDS | EmrKY-TolC multidrug resistance efflux pump, membrane fusion protein component | Periplasmic | COG1566;Multidrug resistance efflux pump | GO:0042493 response to drug | gg | tt |
| *emrY* | CDS | predicted multidrug efflux system | Integral Membrane Protein | COG0477;Permeases of the major facilitator superfamily | GO:0042493 response to drug | o | tt |
| *emtA* | CDS | lytic murein endotransglycosylase E | Outer Membrane Lipoprotein | COG0741;Soluble lytic murein transglycosylase and related regulatory proteins (some contain LysM/invasin domains) | GO:0009252 peptidoglycan biosynthesis | | |
| *endA* | CDS | DNA-specific endonuclease I | Periplasmic | COG2356;Endonuclease I | GO:0006308 DNA catabolism -!- GO:0006308 DNA catabolism | gg | tt |
| *eno* | CDS | enolase | Cytoplasmic | COG0148;Enolase | GO:0006096 glycolysis -!- GO:0009061 anaerobic respiration -!- GO:0006094 gluconeogenesis | gg | o |
| *entA* | CDS | 2,3-dihydro-2,3-dihydroxybenzoate dehydrogenase | Cytoplasmic | COG1028;Dehydrogenases with different specificities (related to short-chain alcohol dehydrogenases) | GO:0009239 enterobactin biosynthesis | gg | t |
| *entB* | CDS | isochorismatase | Cytoplasmic | COG1535;Isochorismate hydrolase -!- COG3433;Aryl carrier domain | GO:0009239 enterobactin biosynthesis -!- GO:0019184 nonribosomal peptide biosynthesis | gg | tt |
| *entC* | CDS | isochorismate synthase 1 | Cytoplasmic | COG1169;Isochorismate synthase | GO:0009239 enterobactin biosynthesis | gg | t |
| *entD* | CDS | phosphopantetheinyltransferase component of enterobactin synthase multienzyme complex | Membrane Anchored | COG2977;Phosphopantetheinyl transferase component of siderophore synthetase | GO:0009239 enterobactin biosynthesis | | |
| *entE* | CDS | 2,3-dihydroxybenzoate-AMP ligase component of enterobactin synthase multienzyme complex | Cytoplasmic | COG1021;Peptide arylation enzymes | GO:0009239 enterobactin biosynthesis -!- GO:0019184 nonribosomal peptide biosynthesis | gg | t |
| *entF* | CDS | enterobactin synthase multienzyme complex component, ATP-dependent | Cytoplasmic | COG1020;Non-ribosomal peptide synthetase modules and related proteins -!- COG3319;Thioesterase domains of type I polyketide synthases or non-ribosomal peptide synthetases | GO:0009239 enterobactin biosynthesis -!- GO:0015937 coenzyme A biosynthesis -!- GO:0019184 nonribosomal peptide biosynthesis -!- GO:0006826 iron ion transport | o | tt |
| *envC* | CDS | protease with a role in cell division | Periplasmic | | | | |
| *envR* | CDS | DNA-binding transcriptional regulator | Cytoplasmic | COG1309;Transcriptional regulator | GO:0006350 transcription -!- GO:0042493 response to drug | gg | t |
| *envY* | CDS | DNA-binding transcriptional activator | Cytoplasmic | COG2207;AraC-type DNA-binding domain-containing proteins | GO:0006350 transcription | gg | t |
| *envZ* | CDS | sensory histidine kinase in two-component regulatory system with OmpR | Integral Membrane Protein | COG0642;Signal transduction histidine kinase | GO:0006464 protein modification | o | tt |
| *epd* | CDS | D-erythrose 4-phosphate dehydrogenase | Cytoplasmic | COG0057;Glyceraldehyde-3-phosphate dehydrogenase/erythrose-4-phosphate dehydrogenase | GO:0008615 pyridoxine biosynthesis -!- GO:0006094 gluconeogenesis | g | o |
| *eptA* | CDS | predicted metal dependent hydrolase | Integral Membrane Protein | COG2194;Predicted membrane-associated, metal-dependent hydrolase | | | |
| *eptB* | CDS | predicted metal dependent hydrolase | Integral Membrane Protein | COG2194;Predicted membrane-associated, metal-dependent hydrolase | | | |
| *era* | CDS | membrane-associated, 16S rRNA-binding GTPase | Cytoplasmic | COG1159;GTPase | GO:0007049 cell cycle | gg | t |
| *erfK* | CDS | conserved protein with NAD(P)-binding Rossmann-fold domain | Periplasmic | COG1376;Uncharacterized protein conserved in bacteria | | o | tt |
| *essD* | CDS | DLP12 prophage; predicted phage lysis protein | Integral Membrane Protein | | | o | tt |
| *essQ* | CDS | Qin prophage; predicted S lysis protein | Integral Membrane Protein | | | | |
| *etp* | CDS | phosphotyrosine-protein phosphatase | Cytoplasmic | | GO:0006464 protein modification | | |
| *eutA* | CDS | reactivating factor for ethanolamine ammonia lyase | Cytoplasmic | COG4819;Ethanolamine utilization protein, possible chaperonin protecting lyase from inhibition | | gg | tt |
| *eutB* | CDS | ethanolamine ammonia-lyase, large subunit, heavy chain | Cytoplasmic | | GO:0009310 amine catabolism | g | tt |
| *eutC* | CDS | ethanolamine ammonia-lyase, small subunit (light chain) | Cytoplasmic | COG4302;Ethanolamine ammonia-lyase, small subunit | GO:0009310 amine catabolism | gg | t |
| *eutE* | CDS | predicted aldehyde dehydrogenase, ethanolamine utilization protein | Cytoplasmic | COG1012;NAD-dependent aldehyde dehydrogenases | GO:0009310 amine catabolism | g | t |
| *eutG* | CDS | predicted alcohol dehydrogenase in ethanolamine utilization | Cytoplasmic | | GO:0009310 amine catabolism | gg | tt |
| *eutH* | CDS | predicted inner membrane protein | Integral Membrane Protein | COG3192;Ethanolamine utilization protein | GO:0009310 amine catabolism | o | tt |
| *eutI* | CDS | predicted phosphotransacetylase subunit | Cytoplasmic | COG0280;Phosphotransacetylase | GO:0009310 amine catabolism | gg | t |
| *eutJ* | CDS | predicted chaperonin, ethanolamine utilization protein | Cytoplasmic | COG4820;Ethanolamine utilization protein, possible chaperonin | GO:0009310 amine catabolism -!- GO:0006457 protein folding | gg | t |
| *eutL* | CDS | predicted carboxysome structural protein with predicted role in ethanolamine utilization | Cytoplasmic | COG4816;Ethanolamine utilization protein | GO:0009310 amine catabolism | gg | t |
| *eutP* | CDS | conserved protein with nucleoside triphosphate hydrolase domain | Cytoplasmic | COG4917;Ethanolamine utilization protein | GO:0009310 amine catabolism | gg | t |
| *eutQ* | CDS | conserved protein | Cytoplasmic | COG4766;Ethanolamine utilization protein | GO:0009310 amine catabolism | gg | t |
| *eutT* | CDS | predicted cobalamin adenosyltransferase in ethanolamine utilization | Cytoplasmic | COG4812;Ethanolamine utilization cobalamin adenosyltransferase | GO:0009310 amine catabolism | g | tt |
| *evgA* | CDS | DNA-binding response regulator in two-component regulatory system with EvgS | Cytoplasmic | COG2197;Response regulator containing a CheY-like receiver domain and an HTH DNA-binding domain | GO:0006350 transcription | gg | o |
| *evgS* | CDS | hybrid sensory histidine kinase in two-component regulatory system with EvgA | Integral Membrane Protein | COG0834;ABC-type amino acid transport/signal transduction systems, periplasmic component/domain -!- COG0642;Signal transduction histidine kinase -!- COG0784;FOG: CheY-like receiver | GO:0006464 protein modification | o | tt |
| *exbB* | CDS | membrane spanning protein in TonB-ExbB-ExbD complex | Integral Membrane Protein | COG0811;Biopolymer transport proteins | GO:0006826 iron ion transport | o | tt |
| *exbD* | CDS | membrane spanning protein in TonB-ExbB-ExbD complex | Membrane Anchored | COG0848;Biopolymer transport protein | GO:0006826 iron ion transport | o | tt |
| *exo* | CDS | exonuclease IX (5'-3' exonuclease) | Cytoplasmic | | GO:0006308 DNA catabolism -!- GO:0006308 DNA catabolism | | |
| *exoX* | CDS | DNA exonuclease X | Cytoplasmic | COG0847;DNA polymerase III, epsilon subunit and related 3'-5' exonucleases | GO:0006308 DNA catabolism -!- GO:0006281 DNA repair -!- GO:0006308 DNA catabolism | g | t |
| *exuR* | CDS | DNA-binding transcriptional repressor | Cytoplasmic | COG2186;Transcriptional regulators | GO:0016052 carbohydrate catabolism -!- GO:0006350 | o | tt |
| *exuT* | CDS | hexuronate transporter | Integral Membrane Protein | COG0477;Permeases of the major facilitator superfamily | GO:0016052 carbohydrate | o | tt |
| *fabA* | CDS | beta-hydroxydecanoyl thioester dehydrase | Cytoplasmic | COG0764;3-hydroxymyristoyl/3-hydroxydecanoyl-(acyl carrier protein) dehydratases | | gg | t |
| *fabB* | CDS | 3-oxoacyl-[acyl-carrier-protein] synthase I | Cytoplasmic | COG0304;3-oxoacyl-(acyl-carrier-protein) synthase | | gg | tt |
| *fabD* | CDS | malonyl-CoA-[acyl-carrier-protein] transacylase | Cytoplasmic | COG0331;(acyl-carrier-protein) S-malonyltransferase | | gg | o |
| *fabF* | CDS | 3-oxoacyl-[acyl-carrier-protein] synthase II | Cytoplasmic | COG0304;3-oxoacyl-(acyl-carrier-protein) synthase | | g | t |
| *fabG* | CDS | 3-oxoacyl-[acyl-carrier-protein] reductase | Cytoplasmic | COG1028;Dehydrogenases with different specificities (related to short-chain alcohol dehydrogenases) | | gg | o |
| *fabH* | CDS | 3-oxoacyl-[acyl-carrier-protein] synthase III | Cytoplasmic | COG0332;3-oxoacyl-[acyl-carrier-protein] synthase III | | o | tt |
| *fabI* | CDS | enoyl-[acyl-carrier-protein] reductase, NADH-dependent | Cytoplasmic | COG0623;Enoyl-[acyl-carrier-protein] reductase (NADH) | | gg | o |
| *fabR* | CDS | DNA-binding transcriptional repressor | Cytoplasmic | COG1309;Transcriptional regulator | | gg | t |
| *fabZ* | CDS | (3R)-hydroxymyristol acyl carrier protein dehydratase | Cytoplasmic | COG0764;3-hydroxymyristoyl/3-hydroxydecanoyl-(acyl carrier protein) dehydratases | | gg | tt |
| *fadA* | CDS | 3-ketoacyl-CoA thiolase (thiolase I) | Cytoplasmic | | GO:0019395 fatty acid oxidation | | |
| *fadB* | CDS | fused 3-hydroxybutyryl-CoA epimerase/delta(3)-cis-delta(2)-trans-enoyl-CoA isomerase/enoyl-CoA hydratase -!- 3-hydroxyacyl-CoA dehydrogenase | Cytoplasmic | COG1024;Enoyl-CoA hydratase/carnithine racemase -!- COG1250;3-hydroxyacyl-CoA dehydrogenase | GO:0019395 fatty acid oxidation | o | tt |
| *fadD* | CDS | acyl-CoA synthetase (long-chain-fatty-acid--CoA ligase) | Cytoplasmic | COG0318;Acyl-CoA synthetases (AMP-forming)/AMP-acid ligases II | GO:0019395 fatty acid oxidation -!- GO:0008654 phospholipid biosynthesis | o | o |
| *fadE* | CDS | acyl coenzyme A dehydrogenase | Integral Membrane Protein | COG1960;Acyl-CoA dehydrogenases | GO:0019395 fatty acid oxidation | | |
| *fadH* | CDS | 2,4-dienoyl-CoA reductase, NADH and FMN-linked | Cytoplasmic | COG1902;NADH:flavin oxidoreductases, Old Yellow Enzyme family -!- COG0446;Uncharacterized NAD(FAD)-dependent dehydrogenases | GO:0019395 fatty acid oxidation | o | t |
| *fadL* | CDS | long-chain fatty acid outer membrane transporter | Outer Membrane B-barrel protein | COG2067;Long-chain fatty acid transport protein | GO:0019395 fatty acid oxidation | o | tt |
| *fadR* | CDS | DNA-binding transcriptional dual regulator | Cytoplasmic | COG2186;Transcriptional regulators | GO:0019395 fatty acid oxidation -!- GO:0006350 transcription | gg | t |
| *fbaA* | CDS | fructose-bisphosphate aldolase, class II | Cytoplasmic | COG0191;Fructose/tagatose bisphosphate aldolase | GO:0016052 carbohydrate | gg | o |
| *fbaB* | CDS | fructose-bisphosphate aldolase class I | Cytoplasmic | | GO:0006096 glycolysis | | |
| *fbp* | CDS | fructose-1,6-bisphosphatase I | Cytoplasmic | COG0158;Fructose-1,6-bisphosphatase | GO:0006094 gluconeogenesis | gg | t |
| *fcl* | CDS | bifunctional GDP-fucose synthetase: GDP-4-dehydro-6-deoxy-D-mannose epimerase and GDP-4-dehydro-6-L-deoxygalactose reductase | Cytoplasmic | COG0451;Nucleoside-diphosphate-sugar epimerases | GO:0009242 colanic acid biosynthesis -!- GO:0009269 response to dessication | gg | t |
| *fdhD* | CDS | formate dehydrogenase formation protein | Cytoplasmic | COG1526;Uncharacterized protein required for formate dehydrogenase activity | GO:0009061 anaerobic respiration -!- GO:0006457 protein folding | g | t |
| *fdhE* | CDS | formate dehydrogenase formation protein | Cytoplasmic | COG3058;Uncharacterized protein involved in formate dehydrogenase formation | GO:0009061 anaerobic respiration -!- GO:0006457 protein folding | gg | o |
| *fdhF* | CDS | formate dehydrogenase-H, selenopolypeptide | Cytoplasmic | COG0243;Anaerobic dehydrogenases, typically selenocysteine-containing | GO:0006113 fermentation | o | t |
| *fdnG* | CDS | formate dehydrogenase-N, alpha subunit, nitrate-inducible | Periplasmic | COG0243;Anaerobic dehydrogenases, typically selenocysteine-containing | GO:0009061 anaerobic respiration | o | o |
| *fdnH* | CDS | formate dehydrogenase-N, Fe-S (beta) subunit, nitrate-inducible | Membrane Anchored | COG0437;Fe-S-cluster-containing hydrogenase components 1 | GO:0009061 anaerobic respiration | o | tt |
| *fdnI* | CDS | formate dehydrogenase-N, cytochrome B556 (gamma) subunit, nitrate-inducible | Integral Membrane Protein | COG2864;Cytochrome b subunit of formate dehydrogenase | GO:0009061 anaerobic respiration -!- GO:0017004 cytochrome biogenesis | o | tt |

| | | | | | | | |
|---|---|---|---|---|---|---|---|
| *fdoG* | CDS | formate dehydrogenase-O, large subunit | Periplasmic | COG0243;Anaerobic dehydrogenases, typically selenocysteine-containing | GO:0009061 anaerobic respiration | o | o |
| *fdoH* | CDS | formate dehydrogenase-O, Fe-S subunit | Membrane Anchored | COG0437;Fe-S-cluster-containing hydrogenase components 1 | GO:0009061 anaerobic respiration | o | tt |
| *fdoI* | CDS | formate dehydrogenase-O, cytochrome b556 subunit | Integral Membrane Protein | COG2864;Cytochrome b subunit of formate dehydrogenase | GO:0009061 anaerobic respiration -!- GO:0017004 cytochrome biogenesis | o | tt |
| *fdrA* | CDS | predicted acyl-CoA synthetase with NAD(P)-binding Rossmann-fold domain | Integral Membrane Protein | COG0074;Succinyl-CoA synthetase, alpha subunit | | gg | t |
| *fdx* | CDS | [2Fe-2S] ferredoxin | Cytoplasmic | COG0633;Ferredoxin | | gg | tt |
| *feaB* | CDS | phenylacetaldehyde dehydrogenase | Cytoplasmic | | GO:0009310 amine catabolism -!- GO:0009435 nicotinamide adenine dinucleotide biosynthesis | o | tt |
| *feaR* | CDS | DNA-binding transcriptional regulator | Cytoplasmic | COG2207;AraC-type DNA-binding domain-containing proteins | GO:0009063 amino acid catabolism -!- GO:0009310 amine catabolism -!- GO:0006350 transcription | gg | t |
| *fecA* | CDS | KpLE2 phage-like element; ferric citrate outer membrane transporter | Outer Membrane B-barrel protein | COG4772;Outer membrane receptor for Fe3+-dicitrate | GO:0006826 iron ion transport | o | tt |
| *fecB* | CDS | KpLE2 phage-like element; iron-dicitrate transporter subunit -!- periplasmic-binding component of ABC superfamily | Periplasmic | COG4594;ABC-type Fe3+-citrate transport system, periplasmic component | GO:0006826 iron ion transport | o | tt |
| *fecC* | CDS | KpLE2 phage-like element; iron-dicitrate transporter subunit -!- membrane component of ABC superfamily | Integral Membrane Protein | COG0609;ABC-type Fe3+-siderophore transport system, permease component | GO:0006826 iron ion transport | o | tt |
| *fecD* | CDS | KpLE2 phage-like element; iron-dicitrate transporter subunit -!- membrane component of ABC superfamily | Integral Membrane Protein | COG0609;ABC-type Fe3+-siderophore transport system, permease component | | o | tt |
| *fecE* | CDS | KpLE2 phage-like element; iron-dicitrate transporter subunit -!- ATP-binding component of ABC superfamily | Cytoplasmic | COG1120;ABC-type cobalamin/Fe3+-siderophores transport systems, ATPase components | GO:0006826 iron ion transport | g | t |
| *fecI* | CDS | KpLE2 phage-like element; RNA polymerase, sigma 19 factor | Cytoplasmic | COG1595;DNA-directed RNA polymerase specialized sigma subunit, sigma24 homolog | GO:0006350 transcription | gg | t |
| *fecR* | CDS | KpLE2 phage-like element; transmembrane signal transducer for ferric citrate transport | Integral Membrane Protein | COG3712;Fe2+-dicitrate sensor, membrane component | GO:0006350 transcription -!- GO:0006826 iron ion transport | o | t |
| *feoA* | CDS | ferrous iron transporter, protein A | Cytoplasmic | COG1918;Fe2+ transport system protein A | GO:0006826 iron ion transport | gg | o |
| *feoB* | CDS | fused ferrous iron transporter, protein B | Integral Membrane Protein | COG0370;Fe2+ transport system protein B | GO:0006826 iron ion transport | o | tt |
| *fepA* | CDS | iron-enterobactin outer membrane transporter | Outer Membrane B-barrel protein | COG4771;Outer membrane receptor for ferrienterochelin and colicins | GO:0006826 iron ion transport | | |
| *fepB* | CDS | iron-enterobactin transporter subunit -!- periplasmic-binding component of ABC superfamily | Periplasmic | COG4592;ABC-type Fe2+-enterobactin transport system, periplasmic component | GO:0006826 iron ion transport | o | tt |
| *fepC* | CDS | iron-enterobactin transporter subunit -!- ATP-binding component of ABC superfamily | Cytoplasmic | COG1120;ABC-type cobalamin/Fe3+-siderophores transport systems, ATPase components | GO:0006826 iron ion transport | gg | t |
| *fepD* | CDS | iron-enterobactin transporter subunit -!- membrane component of ABC superfamily | Integral Membrane Protein | COG0609;ABC-type Fe3+-siderophore transport system, permease component | GO:0006826 iron ion transport | o | tt |
| *fepE* | CDS | regulator of length of O-antigen component of lipopolysaccharide chains | Integral Membrane Protein | COG3765;Chain length determinant protein | GO:0006826 iron ion transport | o | tt |
| *fepG* | CDS | iron-enterobactin transporter subunit -!- membrane component of ABC superfamily | Integral Membrane Protein | COG4779;ABC-type enterobactin transport system, permease component | GO:0006826 iron ion transport | o | tt |
| *fes* | CDS | enterobactin/ferric enterobactin esterase | Cytoplasmic | COG2382;Enterochelin esterase and related enzymes | GO:0006826 iron ion transport | gg | t |
| *ffh* | CDS | Signal Recognition Particle (SRP) component with 4.5S RNA (ffs) | Cytoplasmic | COG0541;Signal recognition particle GTPase | | | |
| *ffs* | misc_RNA | 4.5S RNA | | | | | |
| *fhiA* | CDS | flagellar system protein, promoterless fragment (pseudogene) | Integral Membrane Protein | COG1298;Flagellar biosynthesis pathway, component FlhA | | | |
| *fhlA* | CDS | DNA-binding transcriptional activator | Cytoplasmic | COG3604;Transcriptional regulator containing GAF, AAA-type ATPase, and DNA binding domains | GO:0006113 fermentation -!- GO:0006350 transcription | gg | t |
| *fhuA* | CDS | ferrichrome outer membrane transporter | Outer Membrane B-barrel protein | COG1629;Outer membrane receptor proteins, mostly Fe transport | | o | tt |
| *fhuB* | CDS | fused subunits of iron-hydroxamate transporter -!- membrane components of ABC superfamily transporter | Integral Membrane Protein | COG0609;ABC-type Fe3+-siderophore transport system, permease component | | o | tt |
| *fhuC* | CDS | iron-hydroxamate transporter subunit -!- ATP-binding component of ABC superfamily | Cytoplasmic | COG1120;ABC-type cobalamin/Fe3+-siderophores transport systems, ATPase components | | gg | o |
| *fhuD* | CDS | iron-hydroxamate transporter subunit -!- periplasmic-binding component of ABC superfamily | Periplasmic | COG0614;ABC-type Fe3+-hydroxamate transport system, periplasmic component | GO:0006826 iron ion transport | gg | t |
| *fhuE* | CDS | ferric-rhodotorulic acid outer membrane transporter | Outer Membrane B-barrel protein | COG4773;Outer membrane receptor for ferric coprogen and ferric-rhodotorulic acid | GO:0006826 iron ion transport | o | tt |
| *fhuF* | CDS | ferric iron reductase involved in ferric hydroximate transport | Cytoplasmic | COG4114;Uncharacterized Fe-S protein | GO:0006457 protein folding | o | tt |
| *fic* | CDS | stationary-phase protein, cell division | Cytoplasmic | COG2184;Protein involved in cell division | | gg | tt |
| *fieF* | CDS | zinc transporter | Integral Membrane Protein | COG0053;Predicted Co/Zn/Cd cation transporters | | o | tt |
| *fimA* | CDS | major type 1 subunit fimbrin (pilin) | Periplasmic | COG3539;P pilus assembly protein, pilin FimA | GO:0009101 glycoprotein biosynthesis | o | tt |
| *fimB* | CDS | tyrosine recombinase/inversion of on/off regulator of fimA | Cytoplasmic | COG0582;Integrase | GO:0009297 fimbrial biogenesis -!- GO:0006310 DNA recombination | o | tt |
| *fimC* | CDS | chaperone, periplasmic | Periplasmic | | GO:0006457 protein folding | o | o |
| *fimD* | CDS | outer membrane usher protein, type 1 fimbrial synthesis | Outer Membrane B-barrel protein | COG3188;P pilus assembly protein, porin PapC | | | |
| *fimE* | CDS | tyrosine recombinase/inversion of on/off regulator of fimA | Cytoplasmic | COG0582;Integrase | GO:0009297 fimbrial biogenesis -!- GO:0006310 DNA recombination | o | t |
| *fimF* | CDS | minor component of type 1 fimbriae | Periplasmic | COG3539;P pilus assembly protein, pilin FimA | GO:0009101 glycoprotein biosynthesis | o | o |
| *fimG* | CDS | minor component of type 1 fimbriae | Periplasmic | COG3539;P pilus assembly protein, pilin FimA | GO:0009101 glycoprotein biosynthesis | o | t |
| *fimH* | CDS | minor component of type 1 fimbriae | Membrane Anchored | | GO:0009101 glycoprotein biosynthesis | o | tt |
| *fimI* | CDS | fimbrial protein involved in type 1 pilus biosynthesis | Periplasmic | COG3539;P pilus assembly protein, pilin FimA | GO:0009101 glycoprotein biosynthesis | | |
| *fimZ* | CDS | predicted DNA-binding transcriptional regulator | Cytoplasmic | COG2197;Response regulator containing a CheY-like receiver domain and an HTH DNA-binding domain | GO:0006350 transcription | | |
| *fis* | CDS | global DNA-binding transcriptional dual regulator | Cytoplasmic | COG2901;Factor for inversion stimulation Fis, transcriptional activator | GO:0006310 DNA recombination | o | tt |
| *fiu* | CDS | predicted iron outer membrane transporter | Outer Membrane B-barrel protein | COG4774;Outer membrane receptor for monomeric catechols | GO:0006826 iron ion transport | o | tt |
| *fixA* | CDS | predicted electron transfer flavoprotein subunit, ETFP adenine nucleotide-binding domain | Cytoplasmic | COG2086;Electron transfer flavoprotein, beta subunit | GO:0016052 carbohydrate catabolism | gg | tt |
| *fixB* | CDS | predicted electron transfer flavoprotein, NAD/FAD-binding domain and ETFP adenine nucleotide-binding domain-like | Cytoplasmic | COG2025;Electron transfer flavoprotein, alpha subunit | GO:0016052 carbohydrate catabolism | gg | o |
| *fixC* | CDS | predicted oxidoreductase, FAD/NAD(P)-binding domain | Periplasmic | COG0644;Dehydrogenases (flavoproteins) | GO:0042413 carnitine catabolism | gg | tt |
| *fixX* | CDS | predicted 4Fe-4S ferredoxin-type protein | Cytoplasmic | COG2440;Ferredoxin-like protein | GO:0042413 carnitine catabolism | g | o |
| *fklB* | CDS | FKBP-type peptidyl-prolyl cis-trans isomerase (rotamase) | Cytoplasmic | | GO:0006457 protein folding | | |
| *fkpA* | CDS | FKBP-type peptidyl-prolyl cis-trans isomerase (rotamase) | Periplasmic | COG0545;FKBP-type peptidyl-prolyl cis-trans isomerases 1 | GO:0006457 protein folding | o | tt |
| *fkpB* | CDS | FKBP-type peptidyl-prolyl cis-trans isomerase (rotamase) | Cytoplasmic | COG1047;FKBP-type peptidyl-prolyl cis-trans isomerases 2 | GO:0006457 protein folding | gg | o |
| *fldA* | CDS | flavodoxin 1 | Cytoplasmic | COG0716;Flavodoxins | | gg | tt |
| *fldB* | CDS | flavodoxin 2 | Cytoplasmic | COG0716;Flavodoxins | GO:0009060 aerobic respiration | gg | tt |
| *flgA* | CDS | assembly protein for flagellar basal-body periplasmic P ring | Periplasmic | COG1261;Flagellar basal body P-ring biosynthesis protein | GO:0009296 flagella biogenesis -!- GO:0042330 taxis | g | tt |
| *flgB* | CDS | flagellar component of cell-proximal portion of basal-body rod | Periplasmic | COG1815;Flagellar basal body protein | GO:0009296 flagella biogenesis -!- GO:0042330 taxis | gg | t |
| *flgC* | CDS | flagellar component of cell-proximal portion of basal-body rod | Periplasmic | COG1558;Flagellar basal body rod protein | GO:0009296 flagella biogenesis -!- GO:0042330 taxis | gg | o |
| *flgD* | CDS | flagellar hook assembly protein | Periplasmic | COG1843;Flagellar hook capping protein | GO:0009296 flagella biogenesis -!- GO:0042330 taxis | gg | o |
| *flgE* | CDS | flagellar hook protein | Cytoplasmic | COG1749;Flagellar hook protein FlgE | GO:0009296 flagella biogenesis -!- GO:0042330 taxis | gg | t |
| *flgF* | CDS | flagellar component of cell-proximal portion of basal-body rod | Periplasmic | COG4787;Flagellar basal body rod protein | GO:0009296 flagella biogenesis -!- GO:0042330 taxis | gg | o |
| *flgG* | CDS | flagellar component of cell-distal portion of basal-body rod | Periplasmic | COG4786;Flagellar basal body rod protein | GO:0009296 flagella biogenesis -!- GO:0042330 taxis | gg | o |
| *flgH* | CDS | flagellar protein of basal-body outer-membrane L ring | Outer Membrane Lipoprotein | COG2063;Flagellar basal body L-ring protein | GO:0009296 flagella biogenesis -!- GO:0042330 taxis | | |
| *flgI* | CDS | predicted flagellar basal body protein | Periplasmic | | GO:0009296 flagella biogenesis -!- GO:0042330 taxis | o | tt |
| *flgJ* | CDS | muramidase | Cytoplasmic | COG3951;Rod binding protein -!- COG1705;Muramidase (flagellum-specific) | GO:0009296 flagella biogenesis -!- GO:0042330 taxis | gg | t |
| *flgK* | CDS | flagellar hook-filament junction protein 1 | Cytoplasmic | COG1256;Flagellar hook-associated protein | GO:0009296 flagella biogenesis -!- GO:0006457 protein folding -!- GO:0042330 taxis | gg | tt |
| *flgL* | CDS | flagellar hook-filament junction protein | Cytoplasmic | COG1344;Flagellin and related hook-associated proteins | GO:0009296 flagella biogenesis -!- GO:0006457 protein folding -!- GO:0042330 taxis | gg | t |
| *flgM* | CDS | anti-sigma factor for FliA (sigma 28) | Cytoplasmic | COG2747;Negative regulator of flagellin synthesis (anti-sigma28 factor) | GO:0006350 transcription -!- GO:0042330 taxis | gg | o |
| *flgN* | CDS | export chaperone for FlgK and FlgL | Cytoplasmic | COG3418;Flagellar biosynthesis/type III secretory pathway chaperone | GO:0009296 flagella biogenesis -!- GO:0006457 protein folding -!- GO:0042330 taxis | gg | o |
| *flhA* | CDS | predicted flagellar export pore protein | Integral Membrane Protein | COG1298;Flagellar biosynthesis pathway, component FlhA | GO:0009296 flagella biogenesis -!- GO:0042330 taxis | o | tt |
| *flhB* | CDS | predicted flagellar export pore protein | Integral Membrane Protein | COG1377;Flagellar biosynthesis pathway, component FlhB | | o | tt |
| *flhC* | CDS | DNA-binding transcriptional regulator with FlhD | Cytoplasmic | | GO:0009296 flagella biogenesis -!- GO:0006350 transcription -!- GO:0042330 taxis | gg | t |

| | | | | | | | |
|---|---|---|---|---|---|---|---|
| *flhD* | CDS | DNA-binding transcriptional dual regulator with FlhC | Cytoplasmic | | GO:0009296 flagella biogenesis -!- GO:0006350 transcription -!- GO:0042330 taxis | gg | o |
| *flhE* | CDS | flagellar protein | Periplasmic | | GO:0009296 flagella biogenesis -!- GO:0042330 taxis | o | tt |
| *fliA* | CDS | RNA polymerase, sigma 28 (sigma F) factor | Cytoplasmic | | GO:0006350 transcription -!- GO:0042330 taxis | gg | tt |
| *fliC* | CDS | flagellar filament structural protein (flagellin) | Cytoplasmic | COG1344;Flagellin and related hook-associated proteins | GO:0009296 flagella biogenesis -!- GO:0042330 taxis | gg | tt |
| *fliD* | CDS | flagellar filament capping protein | Cytoplasmic | COG1345;Flagellar capping protein | GO:0009296 flagella biogenesis -!- GO:0042330 taxis | gg | t |
| *fliE* | CDS | flagellar basal-body component | Cytoplasmic | COG1677;Flagellar hook-basal body protein | GO:0009296 flagella biogenesis -!- GO:0042330 taxis | gg | o |
| *fliF* | CDS | flagellar basal-body MS-ring and collar protein | Integral Membrane Protein | COG1766;Flagellar biosynthesis/type III secretory pathway lipoprotein | GO:0009296 flagella biogenesis -!- GO:0042330 taxis | o | tt |
| *fliG* | CDS | flagellar motor switching and energizing component | Cytoplasmic | COG1536;Flagellar motor switch protein | GO:0009296 flagella biogenesis -!- GO:0042330 taxis | gg | t |
| *fliH* | CDS | flagellar biosynthesis protein | Cytoplasmic | COG1317;Flagellar biosynthesis/type III secretory pathway protein | GO:0009296 flagella biogenesis -!- GO:0042330 taxis | gg | tt |
| *fliI* | CDS | flagellum-specific ATP synthase | Cytoplasmic | COG1157;Flagellar biosynthesis/type III secretory pathway ATPase | GO:0015986 ATP synthesis coupled proton transport | gg | t |
| *fliJ* | CDS | flagellar protein | Cytoplasmic | COG2882;Flagellar biosynthesis chaperone | GO:0009296 flagella biogenesis -!- GO:0042330 taxis | gg | o |
| *fliK* | CDS | flagellar hook-length control protein | Cytoplasmic | COG3144;Flagellar hook-length control protein | GO:0009296 flagella biogenesis -!- GO:0042330 taxis | gg | o |
| *fliL* | CDS | flagellar biosynthesis protein | Outer Membrane Lipoprotein | COG1580;Flagellar basal body-associated protein | GO:0009296 flagella biogenesis -!- GO:0042330 taxis | o | tt |
| *fliM* | CDS | flagellar motor switching and energizing component | Cytoplasmic | COG1868;Flagellar motor switch protein | GO:0009296 flagella biogenesis -!- GO:0042330 taxis | gg | o |
| *fliN* | CDS | flagellar motor switching and energizing component | Cytoplasmic | COG1886;Flagellar motor switch/type III secretory pathway protein | GO:0009296 flagella biogenesis -!- GO:0042330 taxis | gg | o |
| *fliO* | CDS | flagellar biosynthesis protein | Membrane Anchored | | GO:0009296 flagella biogenesis -!- GO:0042330 taxis | | |
| *fliP* | CDS | flagellar biosynthesis protein | Integral Membrane Protein | COG1338;Flagellar biosynthesis pathway, component FliP | GO:0009296 flagella biogenesis -!- GO:0042330 taxis | o | tt |
| *fliQ* | CDS | flagellar biosynthesis protein | Integral Membrane Protein | COG1987;Flagellar biosynthesis pathway, component FliQ | GO:0009101 glycoprotein biosynthesis -!- GO:0009296 flagella biogenesis | o | tt |
| *fliR* | CDS | flagellar export pore protein | Integral Membrane Protein | COG1684;Flagellar biosynthesis pathway, component FliR | | o | tt |
| *fliS* | CDS | flagellar protein potentiates polymerization | Cytoplasmic | COG1516;Flagellin-specific chaperone FliS | GO:0009296 flagella biogenesis -!- GO:0042330 taxis | gg | o |
| *fliT* | CDS | predicted chaperone | Cytoplasmic | | GO:0009296 flagella biogenesis -!- GO:0006457 protein folding -!- GO:0042330 taxis | gg | o |
| *fliY* | CDS | cystine transporter subunit -!- periplasmic-binding component of ABC superfamily | Periplasmic | COG0834;ABC-type amino acid transport/signal transduction systems, periplasmic component/domain | GO:0019344 cysteine biosynthesis -!- GO:0042883 L-cysteine transport | o | tt |
| *fliZ* | CDS | predicted regulator of FliA activity | Cytoplasmic | | | gg | tt |
| *flk* | CDS | predicted flagella assembly protein | Membrane Anchored | | | o | tt |
| *flu* | CDS | CP4-44 prophage; antigen 43 (Ag43) phase-variable biofilm formation autotransporter | Outer Membrane B-barrel protein | | | o | tt |
| *flxA* | CDS | Qin prophage; predicted protein | Cytoplasmic | | | gg | o |
| *fmt* | CDS | 10-formyltetrahydrofolate:L-methionyl-tRNA(fMet) N-formyltransferase | Cytoplasmic | COG0223;Methionyl-tRNA formyltransferase | GO:0009451 RNA modification | o | tt |
| *fnr* | CDS | DNA-binding transcriptional dual regulator, global regulator of anaerobic growth | Cytoplasmic | COG0664;cAMP-binding proteins - catabolite gene activator and regulatory subunit of cAMP-dependent protein kinases | GO:0009061 anaerobic respiration -!- GO:0006350 transcription | gg | t |
| *focA* | CDS | formate transporter | Integral Membrane Protein | COG2116;Formate/nitrite family of transporters | GO:0016052 carbohydrate | o | t |
| *focB* | CDS | predicted formate transporter | Integral Membrane Protein | COG2116;Formate/nitrite family of transporters | | o | tt |
| *folA* | CDS | dihydrofolate reductase | Cytoplasmic | COG0262;Dihydrofolate reductase | GO:0046656 folic acid biosynthesis -!- GO:0009257 10-formyltetrahydrofolate biosynthesis | gg | tt |
| *folB* | CDS | bifunctioal dihydroneopterin aldolase -!- dihydroneopterin triphosphate 2'-epimerase | Cytoplasmic | COG1539;Dihydroneopterin aldolase | GO:0046656 folic acid biosynthesis | g | t |
| *folC* | CDS | bifunctional folylpolyglutamate synthase -!- dihydrofolate synthase | Cytoplasmic | COG0285;Folylpolyglutamate synthase | GO:0046656 folic acid biosynthesis -!- GO:0009257 10-formyltetrahydrofolate biosynthesis | gg | o |
| *folD* | CDS | bifunctional 5,10-methylene-tetrahydrofolate dehydrogenase -!- 5,10-methylene-tetrahydrofolate cyclohydrolase | Cytoplasmic | COG0190;5,10-methylene-tetrahydrofolate dehydrogenase/Methenyl tetrahydrofolate cyclohydrolase | GO:0046656 folic acid biosynthesis -!- GO:0009257 10-formyltetrahydrofolate biosynthesis | gg | t |
| *folE* | CDS | GTP cyclohydrolase I | Cytoplasmic | COG0302;GTP cyclohydrolase I | GO:0046656 folic acid biosynthesis | gg | o |
| *folK* | CDS | 2-amino-4-hydroxy-6-hydroxymethyldihyropteridine pyrophosphokinase | Cytoplasmic | COG0801;7,8-dihydro-6-hydroxymethylpterin-pyrophos phokinase | GO:0046656 folic acid biosynthesis | gg | o |
| *folM* | CDS | dihydrofolate reductase isozyme | Cytoplasmic | COG1028;Dehydrogenases with different specificities (related to short-chain alcohol dehydrogenases) | | gg | t |
| *folP* | CDS | 7,8-dihydropteroate synthase | Cytoplasmic | | GO:0046656 folic acid biosynthesis | gg | t |
| *folX* | CDS | D-erythro-7,8-dihydroneopterin triphosphate 2'-epimerase and dihydroneopterin aldolase | Cytoplasmic | COG1539;Dihydroneopterin aldolase | | g | t |
| *fpr* | CDS | ferredoxin-NADP reductase | Cytoplasmic | COG1018;Flavodoxin reductases (ferredoxin-NADPH reductases) family 1 | GO:0006805 xenobiotic metabolism | gg | o |
| *frc* | CDS | formyl-CoA transferase, NAD(P)-binding | Cytoplasmic | COG1804;Predicted acyl-CoA transferases/carnitine dehydratase | | gg | t |
| *frdA* | CDS | fumarate reductase (anaerobic) catalytic and NAD/flavoprotein subunit | Cytoplasmic | COG1053;Succinate dehydrogenase/fumarate reductase, flavoprotein subunit | GO:0006113 fermentation -!- GO:0009061 anaerobic respiration | g | t |
| *frdB* | CDS | fumarate reductase (anaerobic), Fe-S subunit | Cytoplasmic | COG0479;Succinate dehydrogenase/fumarate reductase, Fe-S protein subunit | GO:0006113 fermentation -!- GO:0009061 anaerobic respiration | g | tt |
| *frdC* | CDS | fumarate reductase (anaerobic), membrane anchor subunit | Integral Membrane Protein | COG3029;Fumarate reductase subunit C | GO:0006113 fermentation -!- GO:0009061 anaerobic respiration | o | tt |
| *frdD* | CDS | fumarate reductase (anaerobic), membrane anchor subunit | Integral Membrane Protein | COG3080;Fumarate reductase subunit D | GO:0006113 fermentation -!- GO:0009061 anaerobic respiration | o | tt |
| *fre* | CDS | flavin reductase | Cytoplasmic | COG0543;2-polyprenylphenol hydroxylase and related flavodoxin oxidoreductases | | gg | o |
| *frlA* | CDS | predicted fructoselysine transporter | Integral Membrane Protein | COG0531;Amino acid transporters | | o | tt |
| *frlB* | CDS | fructoselysine-6-P-deglycase | Cytoplasmic | COG2222;Predicted phosphosugar isomerases | | | |
| *frlC* | CDS | predicted isomerase | Cytoplasmic | | | | |
| *frlD* | CDS | fructoselysine 6-kinase | Cytoplasmic | COG0524;Sugar kinases, ribokinase family | | o | tt |
| *frlR* | CDS | predicted DNA-binding transcriptional regulator | Cytoplasmic | | GO:0006350 transcription | | |
| *frmA* | CDS | alcohol dehydrogenase class III/glutathione-dependent formaldehyde dehydrogenase | Cytoplasmic | COG1062;Zn-dependent alcohol dehydrogenases, class III | GO:0006113 fermentation | gg | t |
| *frmB* | CDS | predicted esterase | Cytoplasmic | COG0627;Predicted esterase | | gg | t |
| *frmR* | CDS | regulator protein that represses frmRAB operon | Cytoplasmic | COG1937;Uncharacterized protein conserved in bacteria | | gg | t |
| *frr* | CDS | ribosome recycling factor | Cytoplasmic | COG0233;Ribosome recycling factor | GO:0006412 protein biosynthesis -!- GO:0009386 translational attenuation | o | tt |
| *frsA* | CDS | hydrolase, binds to enzyme IIA(Glc) | Cytoplasmic | COG1073;Hydrolases of the alpha/beta superfamily | | gg | tt |
| *fruA* | CDS | fused fructose-specific PTS enzyme IIB'BC components | Integral Membrane Protein | COG1445;Phosphotransferase system fructose-specific component IIB -!- COG1299;Phosphotransferase system, fructose-specific IIC component | GO:0016052 carbohydrate catabolism | gg | tt |
| *fruB* | CDS | fused fructose-specific PTS enzyme IIA component -!- HPr component | Cytoplasmic | COG1925;Phosphotransferase system, HPr-related proteins | GO:0016052 carbohydrate catabolism | gg | t |
| *fruK* | CDS | fructose-1-phosphate kinase | Cytoplasmic | COG1105;Fructose-1-phosphate kinase and related fructose-6-phosphate kinase (PfkB) | GO:0006096 glycolysis | gg | t |
| *fruR* | CDS | DNA-binding transcriptional dual regulator | Cytoplasmic | COG1609;Transcriptional regulators | GO:0006096 glycolysis -!- GO:0006350 transcription | gg | tt |
| *frvA* | CDS | predicted enzyme IIA component of PTS | Cytoplasmic | COG1762;Phosphotransferase system mannitol/fructose-specific IIA domain (Ntr-type) | GO:0016052 carbohydrate catabolism | g | t |
| *frvB* | CDS | fused predicted PTS enzyme IIBC components | Integral Membrane Protein | COG1445;Phosphotransferase system fructose-specific component IIB -!- COG1299;Phosphotransferase system, fructose-specific IIC component | | | |
| *frvR* | CDS | predicted regulator | Cytoplasmic | COG3711;Transcriptional antiterminator -!- COG1762;Phosphotransferase system mannitol/fructose-specific IIA domain (Ntr-type) | GO:0006350 transcription | o | t |
| *frvX* | CDS | predicted endo-1,4-beta-glucanase | Cytoplasmic | COG1363;Cellulase M and related proteins | GO:0000272 polysaccharide catabolism | o | tt |
| *frwB* | CDS | predicted enzyme IIB component of PTS | Periplasmic | COG1445;Phosphotransferase system fructose-specific component IIB | GO:0016052 carbohydrate | gg | t |
| *frwC* | CDS | predicted enzyme IIC component of PTS | Integral Membrane Protein | COG1299;Phosphotransferase system, fructose-specific IIC component | | o | tt |
| *frwD* | CDS | predicted enzyme IIB component of PTS | Periplasmic | COG1445;Phosphotransferase system fructose-specific component IIB | GO:0016052 carbohydrate | o | tt |
| *fsaA* | CDS | fructose-6-phosphate aldolase 1 | Cytoplasmic | | | | |
| *fsaB* | CDS | fructose-6-phosphate aldolase 2 | Cytoplasmic | COG0176;Transaldolase | | gg | o |
| *fsr* | CDS | predicted fosmidomycin efflux system | Integral Membrane Protein | COG0477;Permeases of the major facilitator superfamily | GO:0042493 response to drug | o | tt |
| *ftn* | CDS | ferritin iron storage protein (cytoplasmic) | Cytoplasmic | COG1528;Ferritin-like protein | GO:0006826 iron ion transport | o | o |
| *ftsA* | CDS | ATP-binding cell division protein involved in recruitment of FtsK to Z ring | Cytoplasmic | COG0849;Actin-like ATPase involved in cell division | | o | tt |
| *ftsB* | CDS | cell division protein | Membrane Anchored | COG2919;Septum formation initiator | | o | tt |
| *ftsE* | CDS | predicted transporter subunit -!- ATP-binding component of ABC superfamily | Cytoplasmic | COG2884;Predicted ATPase involved in cell division | | o | tt |
| *ftsH* | CDS | protease, ATP-dependent zinc-metallo | Integral Membrane Protein | COG0465;ATP-dependent Zn proteases | | o | tt |
| *ftsI* | CDS | transpeptidase involved in septal peptidoglycan synthesis (penicillin-binding protein 3) | Membrane Anchored | COG0768;Cell division protein FtsI/penicillin-binding protein 2 | GO:0009252 peptidoglycan biosynthesis -!- GO:0042493 response to drug | o | tt |
| *ftsK* | CDS | DNA-binding membrane protein required for chromosome resolution and partitioning | Integral Membrane Protein | COG1674;DNA segregation ATPase FtsK/SpoIIIE and related proteins | GO:0009432 SOS response | o | tt |
| *ftsL* | CDS | membrane bound cell division protein at septum containing leucine zipper motif | Membrane Anchored | COG3116;Cell division protein | | o | tt |
| *ftsN* | CDS | essential cell division protein | Membrane Anchored | COG3087;Cell division protein | | o | tt |
| *ftsQ* | CDS | membrane anchored protein involved in growth of wall at septum | Membrane Anchored | COG1589;Cell division septal protein | | o | tt |
| *ftsW* | CDS | integral membrane protein involved in stabilising FstZ ring during cell division | Integral Membrane Protein | COG0772;Bacterial cell division membrane protein | | o | tt |

| | | | | | | | |
|---|---|---|---|---|---|---|---|
| *ftsX* | CDS | predicted transporter subunit -!- membrane component of ABC superfamily | Integral Membrane Protein | COG2177;Cell division protein | | o | tt |
| *ftsY* | CDS | fused Signal Recognition Particle (SRP) receptor | Cytoplasmic | COG0552;Signal recognition particle GTPase | | o | tt |
| *ftsZ* | CDS | GTP-binding tubulin-like cell division protein | Cytoplasmic | COG0206;Cell division GTPase | | gg | tt |
| *fucA* | CDS | L-fuculose-1-phosphate aldolase | Cytoplasmic | COG0235;Ribulose-5-phosphate 4-epimerase and related epimerases and aldolases | GO:0016052 carbohydrate catabolism | gg | t |
| *fucI* | CDS | L-fucose isomerase | Cytoplasmic | COG2407;L-fucose isomerase and related proteins | GO:0016052 carbohydrate | gg | o |
| *fucK* | CDS | L-fuculokinase | Cytoplasmic | COG1070;Sugar (pentulose and hexulose) kinases | GO:0016052 carbohydrate | g | tt |
| *fucO* | CDS | L-1,2-propanediol oxidoreductase | Cytoplasmic | COG1454;Alcohol dehydrogenase, class IV | GO:0016052 carbohydrate catabolism -!- GO:0009441 glycolate | gg | tt |
| *fucP* | CDS | L-fucose transporter | Integral Membrane Protein | COG0738;Fucose permease | GO:0016052 carbohydrate | g | tt |
| *fucR* | CDS | DNA-binding transcriptional activator | Cytoplasmic | COG1349;Transcriptional regulators of sugar metabolism | GO:0016052 carbohydrate catabolism -!- GO:0006350 | gg | t |
| *fucU* | CDS | L-fucose mutarotase | Cytoplasmic | COG4154;Fucose dissimilation pathway protein FucU | | gg | o |
| *fumA* | CDS | aerobic Class I fumarate hydratase (fumarase A) | Cytoplasmic | COG1951;Tartrate dehydratase alpha subunit/Fumarate hydratase class I, N-terminal domain -!- COG1838;Tartrate dehydratase beta subunit/Fumarate hydratase class I, C-terminal domain | GO:0006099 tricarboxylic acid cycle | gg | tt |
| *fumB* | CDS | anaerobic class I fumarate hydratase (fumarase B) | Cytoplasmic | COG1951;Tartrate dehydratase alpha subunit/Fumarate hydratase class I, N-terminal domain -!- COG1838;Tartrate dehydratase beta subunit/Fumarate hydratase class I, C-terminal domain | GO:0006099 tricarboxylic acid cycle -!- GO:0006113 fermentation -!- GO:0009061 anaerobic respiration | gg | o |
| *fumC* | CDS | fumarate hydratase (fumarase C),aerobic Class II | Cytoplasmic | COG0114;Fumarase | GO:0006099 tricarboxylic acid cycle -!- GO:0006113 fermentation -!- GO:0009061 anaerobic respiration | gg | t |
| *fur* | CDS | DNA-binding transcriptional dual regulator | Cytoplasmic | COG0735;Fe2+/Zn2+ uptake regulation proteins | GO:0006350 transcription | o | tt |
| *fusA* | CDS | protein chain elongation factor EF-G | Cytoplasmic | COG0480;Translation elongation factors (GTPases) | GO:0006412 protein biosynthesis | gg | tt |
| *fxsA* | CDS | inner membrane protein | Integral Membrane Protein | COG3030;Protein affecting phage T7 exclusion by the F plasmid | | o | tt |
| *gabD* | CDS | succinate-semialdehyde dehydrogenase I, NADP-dependent | Cytoplasmic | COG1012;NAD-dependent aldehyde dehydrogenases | GO:0009450 aminobutyrate catabolism -!- GO:0009447 putrescine catabolism | gg | t |
| *gabP* | CDS | gamma-aminobutyrate transporter | Integral Membrane Protein | COG1113;Gamma-aminobutyrate permease and related permeases | GO:0016052 carbohydrate | o | tt |
| *gabT* | CDS | 4-aminobutyrate aminotransferase, PLP-dependent | Cytoplasmic | COG0160;4-aminobutyrate aminotransferase and related aminotransferases | GO:0009450 aminobutyrate catabolism -!- GO:0009447 putrescine catabolism | gg | t |
| *gadA* | CDS | glutamate decarboxylase A, PLP-dependent | Cytoplasmic | COG0076;Glutamate decarboxylase and related PLP-dependent proteins | GO:0009063 amino acid catabolism -!- GO:0006520 amino acid metabolism | gg | o |
| *gadB* | CDS | glutamate decarboxylase B, PLP-dependent | Cytoplasmic | COG0076;Glutamate decarboxylase and related PLP-dependent proteins | GO:0009063 amino acid catabolism -!- GO:0006520 amino acid metabolism | gg | t |
| *gadC* | CDS | predicted glutamate:gamma-aminobutyric acid antiporter | Integral Membrane Protein | COG0531;Amino acid transporters | | gg | tt |
| *gadE* | CDS | DNA-binding transcriptional activator | Cytoplasmic | COG2771;DNA-binding HTH domain-containing proteins | | g | t |
| *gadW* | CDS | DNA-binding transcriptional activator | Cytoplasmic | COG2207;AraC-type DNA-binding domain-containing proteins | GO:0006350 transcription | gg | t |
| *gadX* | CDS | DNA-binding transcriptional dual regulator | Cytoplasmic | COG2207;AraC-type DNA-binding domain-containing proteins | GO:0006350 transcription | o | tt |
| *gadY* | misc_RNA | small RNA | | | | | |
| *galE* | CDS | UDP-galactose-4-epimerase | Cytoplasmic | | GO:0006012 galactose metabolism -!- GO:0009242 colanic acid biosynthesis -!- GO:0016052 carbohydrate catabolism -!- GO:0009435 nicotinamide adenine dinucleotide biosynthesis -!- GO:0009269 response to dessication | gg | o |
| *galF* | CDS | predicted subunit with GalU | Cytoplasmic | COG1210;UDP-glucose pyrophosphorylase | GO:0009226 nucleotide-sugar biosynthesis | gg | t |
| *galK* | CDS | galactokinase | Cytoplasmic | COG0153;Galactokinase | GO:0006012 galactose metabolism -!- GO:0009242 colanic acid biosynthesis -!- GO:0016052 carbohydrate catabolism -!- GO:0009269 response to dessication | gg | t |
| *galM* | CDS | galactose-1-epimerase (mutarotase) | Cytoplasmic | COG2017;Galactose mutarotase and related enzymes | GO:0016052 carbohydrate | gg | t |
| *galP* | CDS | D-galactose transporter | Integral Membrane Protein | COG0477;Permeases of the major facilitator superfamily | GO:0016052 carbohydrate | o | tt |
| *galR* | CDS | DNA-binding transcriptional repressor | Cytoplasmic | COG1609;Transcriptional regulators | GO:0016052 carbohydrate catabolism -!- GO:0006350 | gg | tt |
| *galS* | CDS | DNA-binding transcriptional repressor | Cytoplasmic | COG1609;Transcriptional regulators | GO:0016052 carbohydrate catabolism -!- GO:0006350 | gg | t |
| *galT* | CDS | galactose-1-phosphate uridylyltransferase | Cytoplasmic | COG1085;Galactose-1-phosphate uridylyltransferase | GO:0006012 galactose metabolism -!- GO:0009242 colanic acid biosynthesis -!- GO:0016052 carbohydrate catabolism -!- GO:0009269 response to dessication | gg | t |
| *galU* | CDS | glucose-1-phosphate uridylyltransferase | Cytoplasmic | COG1210;UDP-glucose pyrophosphorylase | GO:0006012 galactose metabolism -!- GO:0006006 glucose metabolism -!- GO:0009242 colanic acid biosynthesis -!- GO:0016052 carbohydrate catabolism -!- GO:0009269 response to dessication | gg | t |
| *gapA* | CDS | glyceraldehyde-3-phosphate dehydrogenase A | Cytoplasmic | COG0057;Glyceraldehyde-3-phosphate dehydrogenase/erythrose-4-phosphate dehydrogenase | GO:0006096 glycolysis -!- GO:0008615 pyridoxine biosynthesis -!- GO:0006094 gluconeogenesis | gg | t |
| *gapC* | CDS | glyceraldehyde-3-phosphate dehydrogenase C, N-ter fragment (pseudogene) | Cytoplasmic | | | gg | t |
| *gapC* | ancestral | glyceraldehyde-3-phosphate dehydrogenase C (pseudogene) | Cytoplasmic | | | | |
| *gapC* | CDS | glyceraldehyde-3-phosphate dehydrogenase C, C-ter fragment (pseudogene) | Cytoplasmic | | | | |
| *garD* | CDS | (D)-galactarate dehydrogenase | Cytoplasmic | COG2721;Altronate dehydratase | GO:0016052 carbohydrate | gg | t |
| *garK* | CDS | glycerate kinase I | Cytoplasmic | | GO:0016052 carbohydrate | gg | tt |
| *garL* | CDS | alpha-dehydro-beta-deoxy-D-glucarate aldolase | Cytoplasmic | COG3836;2,4-dihydroxyhept-2-ene-1,7-dioic acid aldolase | GO:0016052 carbohydrate | gg | t |
| *garP* | CDS | predicted (D)-galactarate transporter | Integral Membrane Protein | COG0477;Permeases of the major facilitator superfamily | GO:0016052 carbohydrate | g | tt |
| *garR* | CDS | tartronate semialdehyde reductase | Cytoplasmic | COG2084;3-hydroxyisobutyrate dehydrogenase and related beta-hydroxyacid dehydrogenases | GO:0016052 carbohydrate catabolism -!- GO:0009441 glycolate | | |
| *gatA* | CDS | galactitol-specific enzyme IIA component of PTS | Cytoplasmic | COG1762;Phosphotransferase system mannitol/fructose-specific IIA domain (Ntr-type) | GO:0016052 carbohydrate catabolism | | |
| *gatA* | CDS | galactitol-specific enzyme IIA component of PTS | Cytoplasmic | | | o | o |
| *gatA* | CDS | galactitol-specific enzyme IIA component of PTS | Cytoplasmic | | | gg | t |
| *gatB* | CDS | galactitol-specific enzyme IIB component of PTS | Cytoplasmic | COG3414;Phosphotransferase system, galactitol-specific IIB component | GO:0016052 carbohydrate | gg | o |
| *gatC* | CDS | galactitol-specific enzyme IIC component of PTS | Integral Membrane Protein | COG3775;Phosphotransferase system, galactitol-specific IIC component | GO:0016052 carbohydrate | o | tt |
| *gatD* | CDS | galactitol-1-phosphate dehydrogenase, Zn-dependent and NAD(P)-binding | Cytoplasmic | COG1063;Threonine dehydrogenase and related Zn-dependent dehydrogenases | GO:0016052 carbohydrate catabolism | gg | t |
| *gatR* | CDS | DNA-binding transcriptional regulator, N-ter fragment (pseudogene) | Cytoplasmic | | | | |
| *gatR* | ancestral | DNA-binding transcriptional regulator (pseudogene) | Cytoplasmic | | | | |
| *gatR* | CDS | DNA-binding transcriptional regulator, C-ter fragment (pseudogene) | Cytoplasmic | | | o | tt |
| *gatY* | CDS | D-tagatose 1,6-bisphosphate aldolase 2, catalytic subunit | Cytoplasmic | COG0191;Fructose/tagatose bisphosphate aldolase | GO:0016052 carbohydrate catabolism | | |
| *gatZ* | CDS | D-tagatose 1,6-bisphosphate aldolase 2, subunit | Cytoplasmic | COG4573;Predicted tagatose 6-phosphate kinase | GO:0016052 carbohydrate | g | tt |
| *gcd* | CDS | glucose dehydrogenase | Integral Membrane Protein | COG4993;Glucose dehydrogenase | GO:0006006 glucose metabolism -!- GO:0016052 carbohydrate | o | tt |
| *gcl* | CDS | glyoxylate carboligase | Cytoplasmic | COG3960;Glyoxylate carboligase | GO:0009441 glycolate metabolism -!- GO:0009442 allantoin assimilation | o | o |
| *gcvA* | CDS | DNA-binding transcriptional dual regulator | Cytoplasmic | COG0583;Transcriptional regulator | GO:0009063 amino acid catabolism -!- GO:0009257 10-formyltetrahydrofolate biosynthesis -!- GO:0006350 transcription | o | tt |
| *gcvB* | misc_RNA | regulatory sRNA | | | | | |
| *gcvH* | CDS | glycine cleavage complex lipoylprotein | Cytoplasmic | COG0509;Glycine cleavage system H protein (lipoate-binding) | GO:0009063 amino acid catabolism -!- GO:0009257 10-formyltetrahydrofolate biosynthesis | gg | t |
| *gcvP* | CDS | glycine decarboxylase, PLP-dependent, subunit (protein P) of glycine cleavage complex | Cytoplasmic | COG0403;Glycine cleavage system protein P (pyridoxal-binding), N-terminal domain -!- COG1003;Glycine cleavage system protein P (pyridoxal-binding), C-terminal domain | GO:0009063 amino acid catabolism -!- GO:0009257 10-formyltetrahydrofolate biosynthesis | gg | tt |
| *gcvR* | CDS | DNA-binding transcriptional repressor, regulatory protein accessory to GcvA | Cytoplasmic | | GO:0009063 amino acid catabolism -!- GO:0009257 10-formyltetrahydrofolate biosynthesis -!- GO:0006350 transcription | o | t |
| *gcvT* | CDS | aminomethyltransferase, tetrahydrofolate-dependent, subunit (T protein) of glycine cleavage complex | Cytoplasmic | COG0404;Glycine cleavage system T protein (aminomethyltransferase) | GO:0009063 amino acid catabolism -!- GO:0009257 10-formyltetrahydrofolate biosynthesis | gg | o |
| *gdhA* | CDS | glutamate dehydrogenase, NADP-specific | Cytoplasmic | COG0334;Glutamate dehydrogenase/leucine dehydrogenase | GO:0006537 glutamate biosynthesis | gg | t |
| *ggt* | CDS | gamma-glutamyltranspeptidase | Periplasmic | COG0405;Gamma-glutamyltransferase | | o | tt |
| *gidA* | CDS | glucose-inhibited cell-division protein | Cytoplasmic | COG0445;NAD/FAD-utilizing enzyme apparently involved in cell division | | gg | t |
| *gidB* | CDS | methyltransferase, glucose-inhibited cell-division protein | Cytoplasmic | COG0357;Predicted S-adenosylmethionine-dependent methyltransferase involved in bacterial cell division | | gg | o |
| *glcB* | CDS | malate synthase G | Cytoplasmic | COG2225;Malate synthase | GO:0016052 carbohydrate catabolism -!- GO:0009436 glyoxylate | gg | t |
| *glcC* | CDS | DNA-binding transcriptional dual regulator, glycolate-binding | Cytoplasmic | COG2186;Transcriptional regulators | GO:0016052 carbohydrate catabolism -!- GO:0006350 | gg | t |
| *glcD* | CDS | glycolate oxidase subunit, FAD-linked | Cytoplasmic | COG0277;FAD/FMN-containing dehydrogenases | GO:0016052 carbohydrate | g | tt |
| *glcE* | CDS | glycolate oxidase FAD binding subunit | Cytoplasmic | | | | |

| | | | | | | | |
|---|---|---|---|---|---|---|---|
| *glcF* | CDS | glycolate oxidase iron-sulfur subunit | Cytoplasmic | | | | |
| *glcG* | CDS | conserved protein | Cytoplasmic | COG3193;Uncharacterized protein, possibly involved in utilization of glycolate and propanediol | | gg | o |
| *gldA* | CDS | glycerol dehydrogenase, NAD | Cytoplasmic | | GO:0006071 glycerol metabolism | | |
| *glf* | CDS | UDP-galactopyranose mutase, FAD/NAD(P)-binding | Cytoplasmic | COG0562;UDP-galactopyranose mutase | GO:0009243 O antigen biosynthesis | gg | o |
| *glgA* | CDS | glycogen synthase | Cytoplasmic | COG0297;Glycogen synthase | GO:0000271 polysaccharide biosynthesis | g | t |
| *glgB* | CDS | 1,4-alpha-glucan branching enzyme | Cytoplasmic | COG0296;1,4-alpha-glucan branching enzyme | GO:0000271 polysaccharide biosynthesis | o | tt |
| *glgC* | CDS | glucose-1-phosphate adenylyltransferase | Cytoplasmic | COG0448;ADP-glucose pyrophosphorylase | GO:0000271 polysaccharide biosynthesis | o | tt |
| *glgP* | CDS | glycogen phosphorylase | Cytoplasmic | COG0058;Glucan phosphorylase | GO:0016052 carbohydrate catabolism -!- GO:0000271 polysaccharide biosynthesis | o | tt |
| *glgS* | CDS | predicted glycogen synthesis protein | Cytoplasmic | | GO:0000271 polysaccharide biosynthesis | gg | o |
| *glgX* | CDS | glycogen debranching enzyme | Cytoplasmic | COG1523;Type II secretory pathway, pullulanase PulA and related glycosidases | GO:0016052 carbohydrate catabolism -!- GO:0000271 polysaccharide biosynthesis | o | tt |
| *glk* | CDS | glucokinase | Cytoplasmic | COG0837;Glucokinase | GO:0016052 carbohydrate | gg | o |
| *glmM* | CDS | phosphoglucosamine mutase | Cytoplasmic | COG1109;Phosphomannomutase | GO:0009243 O antigen biosynthesis -!- GO:0009246 enterobacterial common antigen biosynthesis -!- GO:0006006 glucose metabolism -!- GO:0009252 peptidoglycan biosynthesis | gg | t |
| *glmS* | CDS | L-glutamine:D-fructose-6-phosphate aminotransferase | Cytoplasmic | COG0449;Glucosamine 6-phosphate synthetase, contains amidotransferase and phosphosugar isomerase domains | GO:0009243 O antigen biosynthesis -!- GO:0009252 peptidoglycan biosynthesis -!- GO:0046349 amino sugar biosynthesis | gg | o |
| *glmU* | CDS | fused N-acetyl glucosamine-1-phosphate uridyltransferase -!- glucosamine-1-phosphate acetyl transferase | Cytoplasmic | COG1207;N-acetylglucosamine-1-phosphate uridyltransferase (contains nucleotidyltransferase and I-patch acetyltransferase domains) | GO:0009243 O antigen biosynthesis -!- GO:0009252 peptidoglycan biosynthesis -!- GO:0009226 nucleotide-sugar biosynthesis -!- GO:0009246 enterobacterial common antigen biosynthesis -!- GO:0009310 amine catabolism | gg | tt |
| *glnA* | CDS | glutamine synthetase | Cytoplasmic | COG0174;Glutamine synthetase | GO:0006542 glutamine biosynthesis -!- GO:0006807 nitrogen metabolism | g | t |
| *glnB* | CDS | regulatory protein P-II for glutamine synthetase | Cytoplasmic | COG0347;Nitrogen regulatory protein PII | GO:0006542 glutamine biosynthesis -!- GO:0006350 transcription | o | tt |
| *glnD* | CDS | uridylyltransferase | Cytoplasmic | COG2844;UTP:GlnB (protein PII) uridylyltransferase | GO:0006807 nitrogen metabolism -!- GO:0006542 glutamine biosynthesis -!- GO:0006464 protein modification | gg | tt |
| *glnE* | CDS | fused deadenylyltransferase -!- adenylyltransferase for glutamine synthetase | Cytoplasmic | COG1391;Glutamine synthetase adenylyltransferase | GO:0006807 nitrogen metabolism -!- GO:0006464 protein modification | o | t |
| *glnG* | CDS | fused DNA-binding response regulator in two-component regulatory system with GlnL, nitrogen regulator I (NRI) | Cytoplasmic | COG2204;Response regulator containing CheY-like receiver, AAA-type ATPase, and DNA-binding domains | GO:0006542 glutamine biosynthesis -!- GO:0006350 transcription | o | t |
| *glnH* | CDS | glutamine transporter subunit -!- periplasmic binding component of ABC superfamily | Periplasmic | COG0834;ABC-type amino acid transport/signal transduction systems, periplasmic component/domain | GO:0006542 glutamine biosynthesis | o | tt |
| *glnK* | CDS | nitrogen assimilation regulatory protein for GlnL, GlnE, and AmtB | Cytoplasmic | COG0347;Nitrogen regulatory protein PII | GO:0006807 nitrogen metabolism | gg | tt |
| *glnL* | CDS | sensory kinase in two-component regulatory system with GlnG | Cytoplasmic | COG3852;Signal transduction histidine kinase, nitrogen specific | GO:0006542 glutamine biosynthesis -!- GO:0006464 protein modification | o | o |
| *glnP* | CDS | glutamine transporter subunit -!- membrane component of ABC superfamily | Integral Membrane Protein | COG0765;ABC-type amino acid transport system, permease component | GO:0006542 glutamine biosynthesis | o | tt |
| *glnQ* | CDS | glutamine transporter subunit -!- ATP-binding component of ABC superfamily | Cytoplasmic | COG1126;ABC-type polar amino acid transport system, ATPase component | GO:0006542 glutamine biosynthesis | g | t |
| *glnS* | CDS | glutamyl-tRNA synthetase | Cytoplasmic | COG0008;Glutamyl- and glutaminyl-tRNA synthetases | GO:0006418 amino acid activation | gg | tt |
| *glnU* | tRNA | tRNA-Gln(UUG) (Glutamine tRNA1) | | | | | |
| *glnV* | tRNA | tRNA-Gln(CUG) (Glutamine tRNA2) | | | | | |
| *glnW* | tRNA | tRNA-Gln(UUG) (Glutamine tRNA1) | | | | | |
| *glnX* | tRNA | tRNA-Gln(CUG) (Glutamine tRNA2) | | | | | |
| *gloA* | CDS | glyoxalase I, Ni-dependent | Cytoplasmic | COG0346;Lactoylglutathione lyase and related lyases | GO:0009438 methylglyoxal metabolism | gg | o |
| *gloB* | CDS | predicted hydroxyacylglutathione hydrolase | Cytoplasmic | COG0491;Zn-dependent hydrolases, including glyoxylases | GO:0009438 methylglyoxal metabolism | gg | o |
| *glpA* | CDS | sn-glycerol-3-phosphate dehydrogenase (anaerobic), large subunit, FAD/NAD(P)-binding | Cytoplasmic | COG0578;Glycerol-3-phosphate dehydrogenase | GO:0009061 anaerobic respiration -!- GO:0006071 glycerol metabolism | gg | t |
| *glpB* | CDS | sn-glycerol-3-phosphate dehydrogenase (anaerobic), membrane anchor subunit | Membrane Associated | COG3075;Anaerobic glycerol-3-phosphate dehydrogenase | GO:0009061 anaerobic respiration -!- GO:0006071 glycerol metabolism | gg | t |
| *glpC* | CDS | sn-glycerol-3-phosphate dehydrogenase (anaerobic), small subunit | Cytoplasmic | COG0247;Fe-S oxidoreductase | GO:0009061 anaerobic respiration -!- GO:0006071 glycerol metabolism | g | t |
| *glpD* | CDS | sn-glycerol-3-phosphate dehydrogenase, aerobic, FAD/NAD(P)-binding | Cytoplasmic | COG0578;Glycerol-3-phosphate dehydrogenase | GO:0009060 aerobic respiration -!- GO:0006071 glycerol metabolism | o | tt |
| *glpE* | CDS | thiosulfate:cyanide sulfurtransferase (rhodanese) | Cytoplasmic | COG0607;Rhodanese-related sulfurtransferase | GO:0009061 anaerobic respiration | gg | tt |
| *glpF* | CDS | glycerol facilitator | Integral Membrane Protein | COG0580;Glycerol uptake facilitator and related permeases (Major Intrinsic Protein Family) | GO:0006071 glycerol metabolism | o | tt |
| *glpG* | CDS | predicted intramembrane serine protease | Integral Membrane Protein | | | | |
| *glpK* | CDS | glycerol kinase | Cytoplasmic | COG0554;Glycerol kinase | GO:0006071 glycerol metabolism | o | tt |
| *glpQ* | CDS | periplasmic glycerophosphodiester | Periplasmic | COG0584;Glycerophosphoryl diester phosphodiesterase | GO:0006071 glycerol metabolism | o | tt |
| *glpR* | CDS | DNA-binding transcriptional repressor | Cytoplasmic | COG1349;Transcriptional regulators of sugar metabolism | GO:0009061 anaerobic respiration -!- GO:0006350 transcription | o | tt |
| *glpT* | CDS | sn-glycerol-3-phosphate transporter | Integral Membrane Protein | COG2271;Sugar phosphate permease | GO:0009060 aerobic respiration -!- GO:0009061 anaerobic respiration -!- GO:0006071 glycerol metabolism | o | tt |
| *glpX* | CDS | fructose 1,6-bisphosphatase II | Cytoplasmic | COG1494;Fructose-1,6-bisphosphatase/sedoheptulose 1,7-bisphosphatase and related proteins | GO:0008652 amino acid biosynthesis | gg | t |
| *gltA* | CDS | citrate synthase | Cytoplasmic | COG0372;Citrate synthase | GO:0006099 tricarboxylic acid cycle -!- GO:0009061 anaerobic respiration | gg | t |
| *gltB* | CDS | glutamate synthase, large subunit | Cytoplasmic | COG0067;Glutamate synthase domain 1 -!- COG0069;Glutamate synthase domain 2 -!- COG0070;Glutamate synthase domain 3 | GO:0006537 glutamate biosynthesis -!- GO:0006807 nitrogen metabolism | g | o |
| *gltD* | CDS | glutamate synthase, 4Fe-4S protein, small subunit | Cytoplasmic | COG0493;NADPH-dependent glutamate synthase beta chain and related oxidoreductases | GO:0006537 glutamate biosynthesis -!- GO:0006807 nitrogen metabolism | g | t |
| *gltF* | CDS | periplasmic protein | Periplasmic | | GO:0006537 glutamate biosynthesis -!- GO:0006807 nitrogen metabolism -!- GO:0006355 regulation of transcription, DNA-dependent | o | tt |
| *gltI* | CDS | glutamate and aspartate transporter subunit -!- periplasmic-binding component of ABC superfamily | Periplasmic | COG0834;ABC-type amino acid transport/signal transduction systems, periplasmic component/domain | GO:0006537 glutamate biosynthesis | | |
| *gltJ* | CDS | glutamate and aspartate transporter subunit -!- membrane component of ABC superfamily | Integral Membrane Protein | COG0765;ABC-type amino acid transport system, permease component | GO:0006537 glutamate biosynthesis -!- GO:0006532 aspartate | o | tt |
| *gltK* | CDS | glutamate and aspartate transporter subunit -!- membrane component of ABC superfamily | Integral Membrane Protein | COG0765;ABC-type amino acid transport system, permease component | GO:0006537 glutamate biosynthesis -!- GO:0006532 aspartate | o | tt |
| *gltL* | CDS | glutamate and aspartate transporter subunit -!- ATP-binding component of ABC superfamily | Cytoplasmic | COG1126;ABC-type polar amino acid transport system, ATPase component | GO:0006537 glutamate biosynthesis -!- GO:0006532 aspartate | g | t |
| *gltP* | CDS | glutamate/aspartate:proton symporter | Integral Membrane Protein | COG1301;Na+/H+-dicarboxylate symporters | GO:0006537 glutamate biosynthesis -!- GO:0006532 aspartate | o | tt |
| *gltS* | CDS | glutamate transporter | Integral Membrane Protein | COG0786;Na+/glutamate symporter | GO:0006537 glutamate biosynthesis -!- GO:0006532 aspartate | o | tt |
| *gltT* | tRNA | tRNA-Glu(UUC) (Glutamate tRNA2) | | | | | |
| *gltU* | tRNA | tRNA-Glu(UUC) (Glutamate tRNA2) | | | | | |
| *gltV* | tRNA | tRNA-Glu(UUC) (Glutamate tRNA2) | | | | | |
| *gltW* | tRNA | tRNA-Glu(UUC) (Glutamate tRNA2) | | | | | |
| *gltX* | CDS | glutamyl-tRNA synthetase | Cytoplasmic | COG0008;Glutamyl- and glutaminyl-tRNA synthetases | GO:0006418 amino acid activation | gg | tt |
| *glvB* | CDS | arbutin specific enzyme IIB component of PTS | Membrane Anchored | | | o | tt |
| *glvC* | CDS | arbutin sp ecific enzyme IIC component of PTS | Integral Membrane Protein | | | o | tt |
| *glvG* | CDS | predicted 6-phospho-beta-glucosidase (pseudogene) | Membrane Anchored | COG1486;Alpha-galactosidases/6-phospho-beta-glucos idases, family 4 of glycosyl hydrolases | GO:0016052 carbohydrate catabolism -!- GO:0009435 nicotinamide adenine dinucleotide | gg | t |
| *glxK* | CDS | glycerate kinase II | Cytoplasmic | COG1929;Glycerate kinase | GO:0016052 carbohydrate catabolism -!- GO:0009441 glycolate metabolism -!- GO:0009442 allantoin | g | tt |
| *glxR* | CDS | tartronate semialdehyde reductase, NADH-dependent | Cytoplasmic | COG2084;3-hydroxyisobutyrate dehydrogenase and related beta-hydroxyacid dehydrogenases | GO:0016052 carbohydrate catabolism -!- GO:0009441 glycolate metabolism -!- GO:0009442 allantoin | gg | t |
| *glyA* | CDS | serine hydroxymethyltransferase | Cytoplasmic | COG0112;Glycine/serine hydroxymethyltransferase | GO:0009063 amino acid catabolism -!- GO:0006545 glycine biosynthesis -!- GO:0046656 folic acid biosynthesis -!- GO:0009257 10-formyltetrahydrofolate biosynthesis | gg | t |
| *glyQ* | CDS | glycine tRNA synthetase, alpha subunit | Cytoplasmic | COG0752;Glycyl-tRNA synthetase, alpha subunit | GO:0006418 amino acid activation | g | t |
| *glyS* | CDS | glycine tRNA synthetase, beta subunit | Cytoplasmic | COG0751;Glycyl-tRNA synthetase, beta subunit | GO:0006418 amino acid activation | o | tt |
| *glyT* | tRNA | tRNA-Gly(UCC) (Glycine tRNA2) | | | | | |
| *glyU* | tRNA | tRNA-Gly(CCC) (Glycine tRNA1) | | | | | |
| *glyV* | tRNA | tRNA-Gly(GCC) (Glycine tRNA3) | | | | | |
| *glyW* | tRNA | tRNA-Gly(GCC) (Glycine tRNA3) | | | | | |
| *glyX* | tRNA | tRNA-Gly(GCC) (Glycine tRNA3) | | | | | |
| *glyY* | tRNA | tRNA-Gly(GCC) (Glycine tRNA3) | | | | | |

| | | | | | | | |
|---|---|---|---|---|---|---|---|
| *gmd* | CDS | GDP-D-mannose dehydratase, NAD(P)-binding | Cytoplasmic | COG1089;GDP-D-mannose dehydratase | GO:0009242 colanic acid biosynthesis -!- GO:0009435 nicotinamide adenine dinucleotide biosynthesis -!- GO:0009269 | gg | t |
| *gmhB* | CDS | D,D-heptose 1,7-bisphosphate phosphatase | Cytoplasmic | COG0241;Histidinol phosphatase and related phosphatases | | gg | tt |
| *gmk* | CDS | guanylate kinase | Cytoplasmic | COG0194;Guanylate kinase | GO:0015949 nucleobase, nucleoside and nucleotide interconversion -!- GO:0009152 purine ribonucleotide biosynthesis | gg | t |
| *gmr* | CDS | modulator of Rnase II stability | Cytoplasmic | COG2202;FOG: PAS/PAC domain -!- COG2200;FOG: EAL domain | | o | t |
| *gnd* | CDS | gluconate-6-phosphate dehydrogenase, decarboxylating | Cytoplasmic | COG0362;6-phosphogluconate dehydrogenase | GO:0016052 carbohydrate catabolism -!- GO:0009051 pentose-phosphate shunt, oxidative branch -!- GO:0009052 pentose-phosphate shunt, non-oxidative branch | gg | o |
| *gnsA* | CDS | predicted regulator of phosphatidylethanolamine synthesis | Cytoplasmic | | | gg | t |
| *gnsB* | CDS | Qin prophage; predicted protein | Cytoplasmic | | | | |
| *gntK* | CDS | gluconate kinase 2 -!- gluconate transport, GNT I system | Cytoplasmic | COG3265;Gluconate kinase | GO:0006006 glucose metabolism | gg | o |
| *gntP* | CDS | fructuronate transporter | Integral Membrane Protein | COG2610;H+/gluconate symporter and related permeases | | o | tt |
| *gntR* | CDS | DNA-binding transcriptional repressor | Cytoplasmic | | GO:0009255 Entner-Doudoroff pathway -!- GO:0006350 transcription | | |
| *gntT* | CDS | gluconate transporter, high-affinity GNT I system | Integral Membrane Protein | | | | |
| *gntU* | CDS | gluconate transporter, low affinity GNT 1 system | Integral Membrane Protein | | | | |
| *gntX* | CDS | gluconate periplasmic binding protein with phosphoribosyltransferase domain, GNT I system | Cytoplasmic | COG1040;Predicted amidophosphoribosyltransferases | | | |
| *gntY* | CDS | predicted gluconate transport associated protein | Cytoplasmic | COG0316;Uncharacterized conserved protein -!- COG0694;Thioredoxin-like proteins and domains | | gg | t |
| *gor* | CDS | glutathione oxidoreductase | Cytoplasmic | COG1249;Pyruvate/2-oxoglutarate dehydrogenase complex, dihydrolipoamide dehydrogenase (E3) component, and related enzymes | | gg | t |
| *gph* | CDS | phosphoglycolate phosphatase | Cytoplasmic | COG0546;Predicted phosphatases | GO:0016052 carbohydrate | gg | tt |
| *gpmA* | CDS | phosphoglyceromutase 1 | Cytoplasmic | COG0588;Phosphoglycerate mutase 1 | GO:0006096 glycolysis -!- GO:0006094 gluconeogenesis | gg | t |
| *gpmI* | CDS | phosphoglycero mutase III, cofactor-independent | Cytoplasmic | COG0696;Phosphoglyceromutase | GO:0016052 carbohydrate | gg | tt |
| *gpp* | CDS | guanosine pentaphosphatase/exopolyphosphatase | Cytoplasmic | | GO:0015949 nucleobase, nucleoside and nucleotide interconversion -!- GO:0006793 phosphorus metabolism -!- GO:0042594 response to | | |
| *gpsA* | CDS | glycerol-3-phosphate dehydrogenase (NAD+) | Cytoplasmic | COG0240;Glycerol-3-phosphate dehydrogenase | GO:0006071 glycerol metabolism -!- GO:0006793 phosphorus metabolism | gg | t |
| *gpt* | CDS | guanine-hypoxanthine phosphoribosyltransferase | Cytoplasmic | COG0503;Adenine/guanine phosphoribosyltransferases and related PRPP-binding proteins | GO:0015949 nucleobase, nucleoside and nucleotide interconversion | gg | tt |
| *greA* | CDS | transcription elongation factor | Cytoplasmic | | GO:0006350 transcription -!- GO:0006355 regulation of transcription, DNA-dependent | gg | tt |
| *greB* | CDS | transcription elongation factor | Cytoplasmic | | GO:0006350 transcription -!- GO:0006355 regulation of transcription, DNA-dependent | gg | tt |
| *groL* | CDS | Cpn60 chaperonin GroEL, large subunit of GroESL | Cytoplasmic | COG0459;Chaperonin GroEL (HSP60 family) | GO:0006457 protein folding | o | tt |
| *groS* | CDS | Cpn10 chaperonin GroES, small subunit of GroESL | Cytoplasmic | COG0234;Co-chaperonin GroES (HSP10) | GO:0006457 protein folding | g | t |
| *grpE* | CDS | heat shock protein | Cytoplasmic | COG0576;Molecular chaperone GrpE (heat shock protein) | GO:0006457 protein folding | o | tt |
| *grxA* | CDS | glutaredoxin 1, redox coenzyme for ribonucleotide reductase (RNR1a) | Cytoplasmic | COG0695;Glutaredoxin and related proteins | | g | o |
| *grxB* | CDS | glutaredoxin 2 (Grx2) | Cytoplasmic | COG2999;Glutaredoxin 2 | | gg | o |
| *grxC* | CDS | glutaredoxin 3 | Cytoplasmic | COG0695;Glutaredoxin and related proteins | | gg | o |
| *gshA* | CDS | gamma-glutamate-cysteine ligase | Cytoplasmic | COG2918;Gamma-glutamylcysteine synthetase | GO:0006750 glutathione biosynthesis | gg | tt |
| *gshB* | CDS | glutathione synthetase | Cytoplasmic | COG0189;Glutathione synthase/Ribosomal protein S6 modification enzyme (glutaminyl transferase) | GO:0006750 glutathione biosynthesis | gg | o |
| *gsk* | CDS | inosine/guanosine kinase | Cytoplasmic | COG0524;Sugar kinases, ribokinase family | GO:0015949 nucleobase, nucleoside and nucleotide interconversion | o | tt |
| *gspA* | CDS | general secretory pathway component, cryptic | Membrane Anchored | | | o | tt |
| *gspC* | CDS | general secretory pathway component, cryptic | Membrane Anchored | COG3031;Type II secretory pathway, component PulC | | o | tt |
| *gspD* | CDS | general secretory pathway component, cryptic | Periplasmic | | | | |
| *gspE* | CDS | general secretory pathway component, cryptic | Cytoplasmic | COG2804;Type II secretory pathway, ATPase PulE/Tfp pilus assembly pathway, ATPase PilB | | o | t |
| *gspF* | CDS | general secretory pathway component, cryptic | Integral Membrane Protein | COG1459;Type II secretory pathway, component PulF | | o | tt |
| *gspG* | CDS | pseudopilin, cryptic, general secretion pathway | Membrane Anchored | COG2165;Type II secretory pathway, pseudopilin PulG | | o | tt |
| *gspH* | CDS | predicted general secretory pathway component, cryptic | Membrane Anchored | COG2165;Type II secretory pathway, pseudopilin PulG | | o | tt |
| *gspI* | CDS | general secretory pathway component, cryptic | Periplasmic | | | | |
| *gspJ* | CDS | predicted general secretory pathway component, cryptic | Periplasmic | COG4795;Type II secretory pathway, component PulJ | | o | tt |
| *gspK* | CDS | general secretory pathway component, cryptic | Periplasmic | COG3156;Type II secretory pathway, component PulK | | o | tt |
| *gspL* | CDS | general secretory pathway component, cryptic | Cytoplasmic | COG3297;Type II secretory pathway, component PulL | | | |
| *gspM* | CDS | general secretory pathway component, cryptic | Membrane Anchored | | | | |
| *gspO* | CDS | bifunctional prepilin leader peptidase -!- methylase | Integral Membrane Protein | COG1989;Type II secretory pathway, prepilin signal peptidase PulO and related peptidases | | o | tt |
| *gss* | CDS | fused glutathionylspermidine amidase -!- glutathionylspermidine synthetase | Membrane Anchored | COG0754;Glutathionylspermidine synthase | GO:0006596 polyamine biosynthesis | gg | t |
| *gst* | CDS | glutathionine S-transferase | Cytoplasmic | COG0625;Glutathione S-transferase | | gg | t |
| *guaA* | CDS | GMP synthetase (glutamine aminotransferase) | Cytoplasmic | COG0518;GMP synthase - Glutamine amidotransferase domain -!- COG0519;GMP synthase, PP-ATPase domain/subunit | GO:0006164 purine nucleotide biosynthesis -!- GO:0015949 nucleobase, nucleoside and nucleotide interconversion | gg | t |
| *guaB* | CDS | IMP dehydrogenase | Cytoplasmic | COG0516;IMP dehydrogenase/GMP reductase -!- COG2524;Predicted transcriptional regulator, contains C-terminal CBS domains -!- COG0516;IMP dehydrogenase/GMP reductase | GO:0006164 purine nucleotide biosynthesis -!- GO:0015949 nucleobase, nucleoside and nucleotide interconversion | | |
| *guaC* | CDS | GMP reductase | Cytoplasmic | COG0516;IMP dehydrogenase/GMP reductase | GO:0009152 purine ribonucleotide biosynthesis -!- GO:0015949 nucleobase, nucleoside and nucleotide interconversion | gg | o |
| *guaD* | CDS | guanine deaminase | Cytoplasmic | COG0402;Cytosine deaminase and related metal-dependent hydrolases | | | |
| *gudD* | CDS | (D)-glucarate dehydratase 1 | Cytoplasmic | COG4948;L-alanine-DL-glutamate epimerase and related enzymes of enolase superfamily | GO:0016052 carbohydrate catabolism | gg | t |
| *gudP* | CDS | predicted D-glucarate transporter | Integral Membrane Protein | COG0477;Permeases of the major facilitator superfamily | GO:0016052 carbohydrate | g | tt |
| *gudX* | CDS | predicted glucarate dehydratase | Cytoplasmic | COG4948;L-alanine-DL-glutamate epimerase and related enzymes of enolase superfamily | GO:0016052 carbohydrate catabolism | gg | o |
| *gutM* | CDS | DNA-binding transcriptional activator | Periplasmic | COG4578;Glucitol operon activator | GO:0016052 carbohydrate | o | tt |
| *gutQ* | CDS | predicted phosphosugar-binding protein | Cytoplasmic | COG0794;Predicted sugar phosphate isomerase involved in capsule formation -!- COG0517;FOG: CBS domain | | | |
| *gyrA* | CDS | DNA gyrase (type II topoisomerase), subunit A | Cytoplasmic | COG0188;Type IIA topoisomerase (DNA gyrase/topo II, topoisomerase IV), A subunit | GO:0006261 DNA dependent DNA replication -!- GO:0006350 transcription | gg | tt |
| *gyrB* | CDS | DNA gyrase, subunit B | Cytoplasmic | | GO:0006261 DNA dependent DNA replication -!- GO:0006350 transcription | | |
| *hcaB* | CDS | 2,3-dihydroxy-2,3-dihydrophenylpropionate dehydrogenase | Cytoplasmic | COG1028;Dehydrogenases with different specificities (related to short-chain alcohol dehydrogenases) | GO:0016052 carbohydrate catabolism | gg | t |
| *hcaC* | CDS | 3-phenylpropionate dioxygenase, predicted ferredoxin subunit | Cytoplasmic | COG2146;Ferredoxin subunits of nitrite reductase and ring-hydroxylating dioxygenases | GO:0016052 carbohydrate catabolism | gg | t |
| *hcaD* | CDS | phenylpropionate dioxygenase, ferredoxin reductase subunit | Periplasmic | COG0446;Uncharacterized NAD(FAD)-dependent dehydrogenases | GO:0016052 carbohydrate catabolism | gg | t |
| *hcaE* | CDS | 3-phenylpropionate dioxygenase, large (alpha) subunit | Cytoplasmic | COG4638;Phenylpropionate dioxygenase and related ring-hydroxylating dioxygenases, large terminal subunit | GO:0016052 carbohydrate catabolism | gg | t |
| *hcaF* | CDS | 3-phenylpropionate dioxygenase, small (beta) subunit | Cytoplasmic | | GO:0016052 carbohydrate | o | t |
| *hcaR* | CDS | DNA-binding transcriptional regulator | Cytoplasmic | COG0583;Transcriptional regulator | GO:0016052 carbohydrate catabolism -!- GO:0006350 | g | t |
| *hcaT* | CDS | predicted 3-phenylpropionic transporter | Integral Membrane Protein | COG0477;Permeases of the major facilitator superfamily | GO:0042889 3-phenylpropionic acid transport | o | tt |
| *hchA* | CDS | Hsp31 molecular chaperone | Cytoplasmic | COG0693;Putative intracellular protease/amidase | | gg | o |
| *hcp* | CDS | hybrid-cluster [4Fe-2S-2O] protein in anaerobic terminal reductases | Cytoplasmic | | | gg | t |
| *hcr* | CDS | HCP oxidoreductase, NADH-dependent | Cytoplasmic | COG1018;Flavodoxin reductases (ferredoxin-NADPH reductases) family 1 | | | |
| *hda* | CDS | ATPase regulatory factor involved in DnaA inactivation | Cytoplasmic | COG0593;ATPase involved in DNA replication initiation | GO:0006261 DNA dependent DNA replication | | |
| *hdeA* | CDS | stress response protein acid-resistance protein | Periplasmic | | | o | tt |
| *hdeB* | CDS | acid-resistance protein | Periplasmic | | | | |
| *hdeD* | CDS | acid-resistance membrane protein | Integral Membrane Protein | COG3247;Uncharacterized conserved protein | | o | tt |
| *hdfR* | CDS | DNA-binding transcriptional regulator | Cytoplasmic | | GO:0006350 transcription | | |
| *hdhA* | CDS | 7alpha-hydroxysteroid dehydrogenase, NAD-dependent | Cytoplasmic | COG1028;Dehydrogenases with different specificities (related to short-chain alcohol dehydrogenases) | | gg | t |
| *helD* | CDS | DNA helicase IV | Cytoplasmic | COG0210;Superfamily I DNA and RNA helicases | GO:0006261 DNA dependent DNA replication | gg | tt |
| *hemA* | CDS | glutamyl tRNA reductase | Cytoplasmic | COG0373;Glutamyl-tRNA reductase | GO:0042493 response to drug | o | tt |
| *hemB* | CDS | porphobilinogen synthase | Cytoplasmic | | | gg | t |
| *hemC* | CDS | hydroxymethylbilane synthase | Cytoplasmic | | | | |
| *hemD* | CDS | uroporphyrinogen III synthase | Cytoplasmic | COG1587;Uroporphyrinogen-III synthase | | o | t |
| *hemE* | CDS | uroporphyrinogen decarboxylase | Cytoplasmic | COG0407;Uroporphyrinogen-III decarboxylase | | gg | t |
| *hemF* | CDS | coproporphyrinogen III oxidase | Cytoplasmic | COG0408;Coproporphyrinogen III oxidase | | gg | o |

| | | | | | | | |
|---|---|---|---|---|---|---|---|
| *hemG* | CDS | protoporphyrin oxidase, flavoprotein | Cytoplasmic | COG4635;Flavodoxin | | o | tt |
| *hemH* | CDS | ferrochelatase | Cytoplasmic | COG0276;Protoheme ferro-lyase (ferrochelatase) | | o | tt |
| *hemL* | CDS | glutamate-1-semialdehyde aminotransferase (aminomutase) | Cytoplasmic | COG0001;Glutamate-1-semialdehyde aminotransferase | | gg | t |
| *hemN* | CDS | coproporphyrinogen III oxidase, SAM and NAD(P)H dependent, oxygen-independent | Cytoplasmic | COG0635;Coproporphyrinogen III oxidase and related Fe-S oxidoreductases | | gg | tt |
| *hemX* | CDS | uroporphyrinogen III methylase | Membrane Anchored | COG2959;Uncharacterized enzyme of heme biosynthesis | | o | tt |
| *hemY* | CDS | predicted protoheme IX synthesis protein | Integral Membrane Protein | COG3071;Uncharacterized enzyme of heme biosynthesis | | o | tt |
| *hepA* | CDS | RNA polymerase-associated helicase protein (ATPase and RNA polymerase recycling factor) | Cytoplasmic | COG0553;Superfamily II DNA/RNA helicases, SNF2 family | GO:0009451 RNA modification | o | tt |
| *hflC* | CDS | modulator for HflB protease specific for phage lambda cII repressor | Membrane Anchored | COG0330;Membrane protease subunits, stomatin/prohibitin homologs | | o | tt |
| *hflD* | CDS | predicted lysogenization regulator | Cytoplasmic | COG2915;Uncharacterized protein involved in purine metabolism | | | |
| *hflK* | CDS | modulator for HflB protease specific for phage lambda cII repressor | Membrane Anchored | COG0330;Membrane protease subunits, stomatin/prohibitin homologs | | o | tt |
| *hflX* | CDS | predicted GTPase | Cytoplasmic | COG2262;GTPases | | g | o |
| *hfq* | CDS | HF-I, host factor for RNA phage Q beta replication | Cytoplasmic | COG1923;Uncharacterized host factor I protein | GO:0009386 translational attenuation | g | t |
| *hha* | CDS | modulator of gene expression, with H-NS | Cytoplasmic | | GO:0006350 transcription | o | tt |
| *hinT* | CDS | purine nucleoside phosphoramidase | Cytoplasmic | | GO:0006412 protein biosynthesis | gg | o |
| *hipA* | CDS | regulator with hipB | Cytoplasmic | COG3550;Uncharacterized protein related to capsule biosynthesis enzymes | GO:0009252 peptidoglycan biosynthesis -!- GO:0006259 DNA metabolism | | |
| *hipB* | CDS | DNA-binding transcriptional regulator | Cytoplasmic | COG1396;Predicted transcriptional regulators | GO:0009252 peptidoglycan biosynthesis -!- GO:0006259 DNA metabolism | gg | tt |
| *hisA* | CDS | N-(5'-phospho-L-ribosyl-formimino)-5-amino-1-(5'-phosphoribosyl)-4-imidazolecarboxamide isomerase | Cytoplasmic | COG0106;Phosphoribosylformimino-5-aminoimidazole carboxamide ribonucleotide (ProFAR) isomerase | GO:0000105 histidine biosynthesis | gg | t |
| *hisB* | CDS | fused histidinol-phosphatase -!- imidazoleglycerol-phosphate dehydratase | Cytoplasmic | COG0241;Histidinol phosphatase and related phosphatases -!- COG0131;Imidazoleglycerol-phosphate dehydratase | GO:0000105 histidine biosynthesis | gg | t |
| *hisC* | CDS | histidinol-phosphate aminotransferase | Cytoplasmic | COG0079;Histidinol-phosphate/aromatic aminotransferase and cobyric acid decarboxylase | GO:0000105 histidine biosynthesis | o | o |
| *hisD* | CDS | bifunctional histidinal dehydrogenase -!- histidinol dehydrogenase | Cytoplasmic | COG0141;Histidinol dehydrogenase | GO:0000105 histidine biosynthesis | gg | t |
| *hisF* | CDS | imidazole glycerol phosphate synthase, catalytic subunit with HisF | Cytoplasmic | COG0107;Imidazoleglycerol-phosphate synthase | GO:0000105 histidine biosynthesis | gg | o |
| *hisG* | CDS | ATP phosphoribosyltransferase | Cytoplasmic | COG0040;ATP phosphoribosyltransferase | GO:0000105 histidine biosynthesis | gg | o |
| *hisH* | CDS | imidazole glycerol phosphate synthase, glutamine amidotransferase subunit | Cytoplasmic | COG0118;Glutamine amidotransferase | GO:0000105 histidine biosynthesis | gg | t |
| *hisI* | CDS | fused phosphoribosyl-AMP cyclohydrolase -!- phosphoribosyl-ATP pyrophosphatase | Cytoplasmic | COG0139;Phosphoribosyl-AMP cyclohydrolase -!- COG0140;Phosphoribosyl-ATP pyrophosphohydrolase | GO:0000105 histidine biosynthesis | gg | t |
| *hisJ* | CDS | histidine/lysine/arginine/ornithine transporter subunit -!- periplasmic-binding component of ABC superfamily | Periplasmic | COG0834;ABC-type amino acid transport/signal transduction systems, periplasmic component/domain | GO:0000105 histidine biosynthesis | o | tt |
| *hisL* | CDS | his operon leader peptide | Cytoplasmic | | GO:0000105 histidine biosynthesis | gg | o |
| *hisM* | CDS | histidine/lysine/arginine/ornithine transporter subunit -!- membrane component of ABC superfamily | Integral Membrane Protein | COG4160;ABC-type arginine/histidine transport system, permease component | GO:0009089 lysine biosynthesis via diaminopimelate -!- GO:0006526 arginine biosynthesis | o | tt |
| *hisP* | CDS | histidine/lysine/arginine/ornithine transporter subunit -!- ATP-binding component of ABC superfamily | Cytoplasmic | COG4598;ABC-type histidine transport system, ATPase component | GO:0009089 lysine biosynthesis via diaminopimelate -!- GO:0006526 arginine biosynthesis -!- GO:0000105 histidine biosynthesis -!- GO:0009063 amino acid catabolism | gg | t |
| *hisQ* | CDS | histidine/lysine/arginine/ornithine transporter subunit -!- membrane component of ABC superfamily | Integral Membrane Protein | COG4215;ABC-type arginine transport system, permease component | GO:0009089 lysine biosynthesis via diaminopimelate -!- GO:0006526 arginine biosynthesis | o | tt |
| *hisR* | tRNA | tRNA-His(GUG) (Histidine tRNA) | | | | | |
| *hisS* | CDS | histidyl tRNA synthetase | Cytoplasmic | COG0124;Histidyl-tRNA synthetase | GO:0006418 amino acid activation | gg | o |
| *hlpA* | CDS | periplasmic chaperone | Periplasmic | COG2825;Outer membrane protein | GO:0006457 protein folding | o | tt |
| *hlyE* | CDS | hemolysin E | Extracellular | | | | |
| *hmp* | CDS | fused nitric oxide dioxygenase -!- dihydropteridine reductase 2 | Cytoplasmic | COG1017;Hemoglobin-like flavoprotein -!- COG1018;Flavodoxin reductases (ferredoxin-NADPH reductases) family 1 | | gg | o |
| *hns* | CDS | global DNA-binding transcriptional dual regulator H-NS | Cytoplasmic | COG2916;DNA-binding protein H-NS | GO:0006350 transcription -!- GO:0006355 regulation of transcription, DNA-dependent -!- GO:0042330 taxis | o | tt |
| *hofB* | CDS | conserved protein with nucleoside triphosphate hydrolase domain | Cytoplasmic | COG2804;Type II secretory pathway, ATPase PulE/Tfp pilus assembly pathway, ATPase PilB | | gg | t |
| *hofC* | CDS | assembly protein in type IV pilin biogenesis, transmembrane protein | Integral Membrane Protein | COG1459;Type II secretory pathway, component PulF | | o | tt |
| *hofQ* | CDS | predicted fimbrial transporter | Periplasmic | COG4796;Type II secretory pathway, component HofQ | | o | tt |
| *hokA* | CDS | toxic polypeptide, small | Membrane Anchored | | | gg | t |
| *hokB* | CDS | toxic polypeptide, small | Membrane Anchored | | | | |
| *hokC* | CDS | toxic membrane protein, small | Periplasmic | | | | |
| *hokD* | CDS | Qin prophage; small toxic polypeptide | Periplasmic | | | o | tt |
| *hokE* | CDS | toxic polypeptide, small | Membrane Anchored | | | | |
| *holA* | CDS | DNA polymerase III, delta subunit | Cytoplasmic | COG1466;DNA polymerase III, delta subunit | GO:0006261 DNA dependent DNA replication | gg | tt |
| *holB* | CDS | DNA polymerase III, delta prime subunit | Cytoplasmic | COG0470;ATPase involved in DNA replication | GO:0006261 DNA dependent DNA replication | o | tt |
| *holC* | CDS | DNA polymerase III, chi subunit | Cytoplasmic | COG2927;DNA polymerase III, chi subunit | GO:0006261 DNA dependent DNA replication | o | tt |
| *holD* | CDS | DNA polymerase III, psi subunit | Cytoplasmic | COG3050;DNA polymerase III, psi subunit | GO:0006261 DNA dependent DNA replication | gg | o |
| *holE* | CDS | DNA polymerase III, theta subunit | Cytoplasmic | | GO:0006261 DNA dependent DNA replication | gg | o |
| *hpt* | CDS | hypoxanthine phosphoribosyltransferase | Cytoplasmic | | GO:0015949 nucleobase, nucleoside and nucleotide interconversion | | |
| *hrpA* | CDS | ATP-dependent helicase | Cytoplasmic | | GO:0006261 DNA dependent DNA replication | | |
| *hrpB* | CDS | predicted ATP-dependent helicase | Cytoplasmic | COG1643;HrpA-like helicases | GO:0006261 DNA dependent DNA replication | gg | t |
| *hscA* | CDS | DnaK-like molecular chaperone specific for IscU | Cytoplasmic | COG0443;Molecular chaperone | GO:0006457 protein folding | gg | tt |
| *hscB* | CDS | DnaJ-like molecular chaperone specific for IscU | Cytoplasmic | COG1076;DnaJ-domain-containing proteins 1 | GO:0006457 protein folding | gg | tt |
| *hscC* | CDS | Hsp70 family chaperone Hsc62, binds to RpoD and inhibits transcription | Cytoplasmic | COG0443;Molecular chaperone | GO:0006457 protein folding | o | tt |
| *hsdM* | CDS | DNA methylase M | Cytoplasmic | COG0286;Type I restriction-modification system methyltransferase subunit | GO:0006306 DNA methylation | gg | t |
| *hsdR* | CDS | endonuclease R | Cytoplasmic | COG4096;Type I site-specific restriction-modification system, R (restriction) subunit and related helicases | GO:0006308 DNA catabolism -!- GO:0006308 DNA catabolism | o | tt |
| *hsdS* | CDS | specificity determinant for hsdM and hsdR | Cytoplasmic | COG0732;Restriction endonuclease S subunits | GO:0006306 DNA methylation | gg | o |
| *hslJ* | CDS | heat-inducible protein | Outer Membrane Lipoprotein | COG3187;Heat shock protein | GO:0009266 response to temperature | o | tt |
| *hslO* | CDS | heat shock protein Hsp33 | Cytoplasmic | COG1281;Disulfide bond chaperones of the HSP33 family | GO:0006457 protein folding | | |
| *hslR* | CDS | ribosome-associated heat shock protein Hsp15 | Cytoplasmic | COG1188;Ribosome-associated heat shock protein implicated in the recycling of the 50S subunit (S4 paralog) | GO:0009266 response to temperature | gg | tt |
| *hslU* | CDS | molecular chaperone and ATPase component of HslUV protease | Cytoplasmic | COG1220;ATP-dependent protease HslVU (ClpYQ), ATPase subunit | GO:0006508 proteolysis and peptidolysis -!- GO:0006457 protein folding -!- GO:0009266 response to temperature | g | tt |
| *hslV* | CDS | peptidase component of the HslUV protease | Cytoplasmic | COG5405;ATP-dependent protease HslVU (ClpYQ), peptidase subunit | GO:0006508 proteolysis and peptidolysis -!- GO:0006457 protein folding | g | t |
| *hsrA* | CDS | predicted multidrug or homocysteine efflux system | Integral Membrane Protein | COG0477;Permeases of the major facilitator superfamily | | o | tt |
| *htgA* | CDS | predicted protein | Cytoplasmic | | GO:0006457 protein folding -!- GO:0006350 transcription -!- GO:0009266 response to | | |
| *htpG* | CDS | molecular chaperone HSP90 family | Cytoplasmic | COG0326;Molecular chaperone, HSP90 family | GO:0006457 protein folding | gg | t |
| *htpX* | CDS | predicted endopeptidase | Integral Membrane Protein | COG0501;Zn-dependent protease with chaperone function | GO:0009266 response to | g | tt |
| *htrC* | CDS | heat shock protein | Cytoplasmic | | GO:0009266 response to | o | tt |
| *htrE* | CDS | predicted outer membrane usher protein | Outer Membrane B-barrel protein | COG3188;P pilus assembly protein, porin PapC | | o | tt |
| *htrG* | CDS | predicted signal transduction protein (SH3 domain) | Membrane Anchored | COG3103;SH3 domain protein | | o | tt |
| *htrL* | CDS | predicted protein | Cytoplasmic | | GO:0009103 lipopolysaccharide biosynthesis | | |
| *hupA* | CDS | HU, DNA-binding transcriptional regulator, alpha subunit | Cytoplasmic | COG0776;Bacterial nucleoid DNA-binding protein | | gg | o |
| *hupB* | CDS | HU, DNA-binding transcriptional regulator, beta subunit | Cytoplasmic | COG0776;Bacterial nucleoid DNA-binding protein | | gg | t |
| *hyaA* | CDS | hydrogenase 1, small subunit | Integral Membrane Protein | COG1740;Ni,Fe-hydrogenase I small subunit | GO:0009060 aerobic respiration -!- GO:0009061 anaerobic respiration | gg | o |
| *hyaB* | CDS | hydrogenase 1, large subunit | Periplasmic | COG0374;Ni,Fe-hydrogenase I large subunit | GO:0009060 aerobic respiration -!- GO:0009061 anaerobic respiration | gg | t |
| *hyaC* | CDS | hydrogenase 1, b-type cytochrome subunit | Integral Membrane Protein | COG1969;Ni,Fe-hydrogenase I cytochrome b subunit | GO:0009060 aerobic respiration -!- GO:0009061 anaerobic respiration -!- GO:0017004 cytochrome biogenesis | o | tt |
| *hyaD* | CDS | protein involved in processing of HyaA and HyaB proteins | Cytoplasmic | COG0680;Ni,Fe-hydrogenase maturation factor | GO:0009060 aerobic respiration -!- GO:0009061 anaerobic respiration -!- GO:0006457 protein folding | gg | o |
| *hyaE* | CDS | protein involved in processing of HyaA and HyaB proteins | Cytoplasmic | | GO:0009060 aerobic respiration -!- GO:0009061 anaerobic respiration -!- GO:0006457 protein folding | gg | o |
| *hyaF* | CDS | protein involved in nickel incorporation into hydrogenase-1 proteins | Cytoplasmic | | GO:0009060 aerobic respiration -!- GO:0009061 anaerobic respiration -!- GO:0006457 protein folding | gg | tt |

| | | | | | | | |
|---|---|---|---|---|---|---|---|
| *hybA* | CDS | hydrogenase 2 4Fe-4S ferredoxin-type component | Periplasmic | COG0437;Fe-S-cluster-containing hydrogenase components 1 | | gg | t |
| *hybB* | CDS | predicted hydrogenase 2 cytochrome b type component | Integral Membrane Protein | | GO:0009061 anaerobic respiration -!- GO:0017004 cytochrome biogenesis | | |
| *hybC* | CDS | hydrogenase 2, large subunit | Periplasmic | COG0374;Ni,Fe-hydrogenase I large subunit | GO:0009061 anaerobic respiration | gg | t |
| *hybD* | CDS | predicted maturation element for hydrogenase 2 | Cytoplasmic | COG0680;Ni,Fe-hydrogenase maturation factor | GO:0009061 anaerobic respiration | gg | tt |
| *hybE* | CDS | hydrogenase 2-specific chaperone | Cytoplasmic | | | gg | t |
| *hybF* | CDS | protein involved with the maturation of hydrogenases 1 and 2 | Cytoplasmic | COG0375;Zn finger protein HypA/HybF (possibly regulating hydrogenase expression) | GO:0009061 anaerobic respiration -!- GO:0006457 protein folding | | |
| *hybG* | CDS | hydrogenase 2 accessory protein | Cytoplasmic | COG0298;Hydrogenase maturation factor | GO:0009061 anaerobic respiration -!- GO:0006457 protein folding | gg | t |
| *hybO* | CDS | hydrogenase 2, small subunit | Membrane Anchored | COG1740;Ni,Fe-hydrogenase I small subunit | GO:0009061 anaerobic respiration -!- GO:0006457 protein folding | gg | t |
| *hycA* | CDS | regulator of the transcriptional regulator FhlA | Cytoplasmic | | GO:0006113 fermentation -!- GO:0006350 transcription | gg | o |
| *hycB* | CDS | hydrogenase 3, Fe-S subunit | Cytoplasmic | COG1142;Fe-S-cluster-containing hydrogenase components 2 | GO:0006113 fermentation -!- GO:0009061 anaerobic respiration | gg | t |
| *hycC* | CDS | hydrogenase 3, membrane subunit | Integral Membrane Protein | COG0651;Formate hydrogenlyase subunit 3/Multisubunit Na+/H+ antiporter, MnhD subunit | GO:0006113 fermentation | o | tt |
| *hycD* | CDS | hydrogenase 3, membrane subunit | Integral Membrane Protein | COG0650;Formate hydrogenlyase subunit 4 | GO:0006113 fermentation | o | tt |
| *hycE* | CDS | hydrogenase 3, large subunit | Cytoplasmic | COG3262;Ni,Fe-hydrogenase III component G -!- COG3261;Ni,Fe-hydrogenase III large subunit | GO:0006113 fermentation -!- GO:0009061 anaerobic respiration | gg | t |
| *hycF* | CDS | formate hydrogenlyase complex iron-sulfur protein | Cytoplasmic | COG1143;Formate hydrogenlyase subunit 6/NADH:ubiquinone oxidoreductase 23 kD subunit (chain I) | GO:0006113 fermentation -!- GO:0009061 anaerobic respiration | gg | tt |
| *hycG* | CDS | hydrogenase 3 and formate hydrogenase complex, HycG subunit | Membrane Anchored | COG3260;Ni,Fe-hydrogenase III small subunit | GO:0006113 fermentation -!- GO:0009061 anaerobic respiration | o | t |
| *hycH* | CDS | protein required for maturation of hydrogenase 3 | Cytoplasmic | | GO:0006113 fermentation | gg | t |
| *hycI* | CDS | protease involved in processing C-terminal end of HycE | Cytoplasmic | COG0680;Ni,Fe-hydrogenase maturation factor | GO:0006457 protein folding | gg | t |
| *hydN* | CDS | formate dehydrogenase-H, [4Fe-4S] ferredoxin | Cytoplasmic | COG1142;Fe-S-cluster-containing hydrogenase components 2 | GO:0009061 anaerobic respiration | g | t |
| *hyfA* | CDS | hydrogenase 4, 4Fe-4S subunit | Cytoplasmic | | GO:0009061 anaerobic respiration | gg | t |
| *hyfB* | CDS | hydrogenase 4, membrane subunit | Integral Membrane Protein | COG0651;Formate hydrogenlyase subunit 3/Multisubunit Na+/H+ antiporter, MnhD subunit | GO:0009061 anaerobic respiration | o | tt |
| *hyfC* | CDS | hydrogenase 4, membrane subunit | Integral Membrane Protein | | GO:0009061 anaerobic respiration | gg | tt |
| *hyfD* | CDS | hydrogenase 4, membrane subunit | Integral Membrane Protein | COG1009;NADH:ubiquinone oxidoreductase subunit 5 (chain L)/Multisubunit Na+/H+ antiporter, MnhA subunit | GO:0009061 anaerobic respiration | o | tt |
| *hyfE* | CDS | hydrogenase 4, membrane subunit | Integral Membrane Protein | COG4237;Hydrogenase 4 membrane component (E) | GO:0009061 anaerobic respiration | o | tt |
| *hyfF* | CDS | hydrogenase 4, membrane subunit | Integral Membrane Protein | COG0651;Formate hydrogenlyase subunit 3/Multisubunit Na+/H+ antiporter, MnhD subunit | GO:0009061 anaerobic respiration | g | tt |
| *hyfG* | CDS | hydrogenase 4, subunit | Cytoplasmic | COG3262;Ni,Fe-hydrogenase III component G -!- COG3261;Ni,Fe-hydrogenase III large subunit | GO:0009061 anaerobic respiration | g | t |
| *hyfH* | CDS | hydrogenase 4, Fe-S subunit | Cytoplasmic | COG1143;Formate hydrogenlyase subunit 6/NADH:ubiquinone oxidoreductase 23 kD subunit (chain I) | GO:0009061 anaerobic respiration | gg | t |
| *hyfI* | CDS | hydrogenase 4, Fe-S subunit | Cytoplasmic | COG3260;Ni,Fe-hydrogenase III small subunit | GO:0009061 anaerobic respiration | | |
| *hyfJ* | CDS | predicted processing element hydrogenase 4 | Cytoplasmic | | GO:0006412 protein biosynthesis | g | t |
| *hyfR* | CDS | DNA-binding transcriptional activator, formate sensing | Cytoplasmic | | GO:0009061 anaerobic respiration -!- GO:0006350 transcription | gg | t |
| *hyi* | CDS | hydroxypyruvate isomerase | Cytoplasmic | COG3622;Hydroxypyruvate isomerase | | gg | o |
| *hypA* | CDS | protein involved in nickel insertion into hydrogenases 3 | Cytoplasmic | COG0375;Zn finger protein HypA/HybF (possibly regulating hydrogenase expression) | GO:0009061 anaerobic respiration | g | o |
| *hypB* | CDS | GTP hydrolase involved in nickel liganding into hydrogenases | Cytoplasmic | COG0378;Ni2+-binding GTPase involved in regulation of expression and maturation of urease and hydrogenase | GO:0006113 fermentation -!- GO:0006457 protein folding | gg | t |
| *hypC* | CDS | protein required for maturation of hydrogenases 1 and 3 | Cytoplasmic | COG0298;Hydrogenase maturation factor | GO:0009061 anaerobic respiration -!- GO:0006464 protein modification | gg | tt |
| *hypD* | CDS | protein required for maturation of hydrogenases | Cytoplasmic | COG0409;Hydrogenase maturation factor | GO:0009061 anaerobic respiration -!- GO:0006464 protein modification | gg | t |
| *hypE* | CDS | carbamoyl phosphate phosphatase, hydrogenase 3 maturation protein | Cytoplasmic | COG0309;Hydrogenase maturation factor | GO:0009061 anaerobic respiration -!- GO:0006457 protein folding | gg | t |
| *hypF* | CDS | carbamoyl phosphate phosphatase and maturation protein for [NiFe] hydrogenases | Cytoplasmic | COG0068;Hydrogenase maturation factor | GO:0009061 anaerobic respiration -!- GO:0006457 protein folding | | |
| *hyuA* | CDS | D-stereospecific phenylhydantoinase | Cytoplasmic | | | gg | t |
| *iaaA* | CDS | L-asparaginase | Cytoplasmic | COG1446;Asparaginase | GO:0009063 amino acid catabolism | gg | t |
| *iadA* | CDS | isoaspartyl dipeptidase | Cytoplasmic | | | g | t |
| *iap* | CDS | aminopeptidase in alkaline phosphatase isozyme conversion | Periplasmic | COG2234;Predicted aminopeptidases | GO:0006464 protein modification | o | tt |
| *ibpA* | CDS | heat shock chaperone | Cytoplasmic | COG0071;Molecular chaperone (small heat shock protein) | GO:0009266 response to | g | o |
| *ibpB* | CDS | heat shock chaperone | Cytoplasmic | COG0071;Molecular chaperone (small heat shock protein) | GO:0009266 response to | g | o |
| *icd* | CDS | e14 prophage; isocitrate dehydrogenase, specific for NADP+ | Cytoplasmic | COG0538;Isocitrate dehydrogenases | GO:0006099 tricarboxylic acid cycle -!- GO:0009061 anaerobic respiration | gg | o |
| *icdC* | CDS | conserved protein (pseudogene) | Cytoplasmic | | | | |
| *iclR* | CDS | DNA-binding transcriptional repressor | Cytoplasmic | COG1414;Transcriptional regulator | GO:0006097 glyoxylate cycle -!- GO:0006350 transcription | o | tt |
| *idi* | CDS | isopentenyl diphosphate isomerase | Cytoplasmic | COG1443;Isopentenyldiphosphate isomerase | | gg | o |
| *idnD* | CDS | L-idonate 5-dehydrogenase, NAD-binding | Cytoplasmic | COG1063;Threonine dehydrogenase and related Zn-dependent dehydrogenases | GO:0016052 carbohydrate catabolism | o | tt |
| *idnK* | CDS | D-gluconate kinase, thermosensitive | Cytoplasmic | COG3265;Gluconate kinase | GO:0016052 carbohydrate | gg | o |
| *idnO* | CDS | 5-keto-D-gluconate-5-reductase | Cytoplasmic | COG1028;Dehydrogenases with different specificities (related to short-chain alcohol dehydrogenases) | GO:0016052 carbohydrate catabolism | o | o |
| *idnR* | CDS | DNA-binding transcriptional repressor, 5-gluconate-binding | Cytoplasmic | COG1609;Transcriptional regulators | GO:0016052 carbohydrate catabolism -!- GO:0006350 | o | o |
| *idnT* | CDS | L-idonate and D-gluconate transporter | Integral Membrane Protein | COG2610;H+/gluconate symporter and related permeases | | o | tt |
| *ihfA* | CDS | integration host factor (IHF), DNA-binding protein, alpha subunit | Cytoplasmic | COG0776;Bacterial nucleoid DNA-binding protein | GO:0006310 DNA recombination | gg | o |
| *ihfB* | CDS | integration host factor (IHF), DNA-binding protein, beta subunit | Cytoplasmic | COG0776;Bacterial nucleoid DNA-binding protein | GO:0006310 DNA recombination | gg | o |
| *ileS* | CDS | isoleucyl-tRNA synthetase | Cytoplasmic | COG0060;Isoleucyl-tRNA synthetase | GO:0006418 amino acid activation | o | tt |
| *ileT* | tRNA | tRNA-Ile(GAU) (Isoleucine tRNA1) | | | | | |
| *ileU* | tRNA | tRNA-Ile(GAU) (Isoleucine tRNA1) | | | | | |
| *ileV* | tRNA | tRNA-Ile(GAU) (Isoleucine tRNA1) | | | | | |
| *ileX* | tRNA | tRNA-Ile(CAU) (Isoleucine tRNA2) | | | | | |
| *ileY* | tRNA | tRNA-Ile(CAU) (Isoleucine tRNA2 variant) | | | | | |
| *ilvA* | CDS | threonine deaminase | Cytoplasmic | COG1171;Threonine dehydratase | | g | t |
| *ilvB* | CDS | acetolactate synthase I, large subunit | Cytoplasmic | COG0028;Thiamine pyrophosphate-requiring enzymes [acetolactate synthase, pyruvate dehydrogenase (cytochrome), glyoxylate carboligase, phosphonopyruvate decarboxylase] | | gg | t |
| *ilvC* | CDS | ketol-acid reductoisomerase, NAD(P)-binding | Cytoplasmic | COG0059;Ketol-acid reductoisomerase | | o | tt |
| *ilvD* | CDS | dihydroxyacid dehydratase | Cytoplasmic | | | | |
| *ilvE* | CDS | branched-chain amino-acid aminotransferase | Cytoplasmic | | GO:0006523 alanine biosynthesis -!- GO:0009098 leucine biosynthesis | | |
| *ilvG* | CDS | acetolactate synthase II, large subunit, N-ter fragment (pseudogene) | Cytoplasmic | | | g | t |
| *ilvG* | ancestral | large subunit of acetolactate synthase II | Cytoplasmic | | | | |
| *ilvG* | CDS | acetolactate synthase II, large subunit, C-ter fragment (pseudogene) | Cytoplasmic | | | g | o |
| *ilvH* | CDS | acetolactate synthase III, thiamin-dependent, small subunit | Cytoplasmic | COG0440;Acetolactate synthase, small (regulatory) subunit | | gg | o |
| *ilvI* | CDS | acetolactate synthase III, large subunit | Cytoplasmic | | | gg | t |
| *ilvL* | CDS | ilv operon leader peptide | Cytoplasmic | | | o | t |
| *ilvM* | CDS | acetolactate synthase II, small subunit | Cytoplasmic | COG3978;Acetolactate synthase (isozyme II), small (regulatory) subunit | | gg | t |
| *ilvN* | CDS | acetolactate synthase I, small subunit | Cytoplasmic | COG0440;Acetolactate synthase, small (regulatory) subunit | | g | o |
| *ilvY* | CDS | DNA-binding transcriptional dual regulator | Cytoplasmic | COG0583;Transcriptional regulator | GO:0006350 transcription | o | tt |
| *imp* | CDS | exported protein required for envelope biosynthesis and integrity | Outer Membrane B-barrel protein | COG1452;Organic solvent tolerance protein OstA | | o | tt |
| *inaA* | CDS | conserved protein | Cytoplasmic | | | gg | o |
| *infA* | CDS | translation initiation factor IF-1 | Cytoplasmic | COG0361;Translation initiation factor 1 (IF-1) | GO:0006412 protein biosynthesis | gg | o |
| *infB* | CDS | fused protein chain initiation factor 2, IF2 | Cytoplasmic | COG0532;Translation initiation factor 2 (IF-2; GTPase) | GO:0006412 protein biosynthesis | o | tt |
| *infC* | CDS | protein chain initiation factor IF-3 | Cytoplasmic | COG0290;Translation initiation factor 3 (IF-3) | GO:0006412 protein biosynthesis | | |
| *insA* | CDS | IS1 element protein | Cytoplasmic | | | | |
| *insA* | CDS | IS1 repressor protein InsA | Cytoplasmic | COG3677;Transposase and inactivated derivatives | | | |
| *insA* | CDS | CP4-6 prophage; IS1 repressor protein InsA | Cytoplasmic | COG3677;Transposase and inactivated derivatives | | | |
| *insA* | CDS | CP4-6 prophage; IS1 repressor protein InsA | Cytoplasmic | COG3677;Transposase and inactivated derivatives | | | |
| *insA* | CDS | IS1 repressor protein InsA | Cytoplasmic | | | | |
| *insA* | CDS | IS1 repressor protein InsA | Cytoplasmic | COG3677;Transposase and inactivated derivatives | | | |
| *insA* | CDS | IS1 repressor protein InsA | Cytoplasmic | COG3677;Transposase and inactivated derivatives | | | |
| *insA* | CDS | KpLE2 phage-like element; IS1 repressor protein | Cytoplasmic | | | | |
| *insB* | CDS | IS1 element protein | Cytoplasmic | | | | |
| *insB* | CDS | IS1 transposase InsAB' | Cytoplasmic | COG1662;Transposase and inactivated derivatives, IS1 family | | | |
| *insB* | CDS | CP4-6 prophage; IS1 transposase InsAB' | Cytoplasmic | COG1662;Transposase and inactivated derivatives, IS1 family | | | |
| *insB* | CDS | CP4-6 prophage; IS1 transposase InsAB' | Cytoplasmic | COG1662;Transposase and inactivated derivatives, IS1 family | | | |
| *insB* | CDS | IS1 transposase InsAB' | Cytoplasmic | COG1662;Transposase and inactivated derivatives, IS1 family | | | |
| *insB* | CDS | IS1 transposase InsAB' | Cytoplasmic | COG1662;Transposase and inactivated derivatives, IS1 family | | | |
| *insB* | CDS | IS1 transposase InsAB' | Cytoplasmic | | | | |
| *insB* | CDS | IS1 transposase InsAB', N-ter fragment | Cytoplasmic | | | | |
| *insB* | ancestral | IS1 transposase InsAB' (pseudogene) | Cytoplasmic | | | | |
| *insB* | CDS | IS1 transposase InsAB', C-ter fragment | Cytoplasmic | | | | |
| *insC* | CDS | IS2 element protein | Cytoplasmic | | | | |
| *insC* | CDS | IS2 element protein | Cytoplasmic | | | | |
| *insC* | CDS | IS2 element protein | Cytoplasmic | | | | |
| *insC* | CDS | IS2 element protein | Cytoplasmic | | | | |
| *insC* | CDS | IS2 insertion element repressor InsA | Cytoplasmic | COG2963;Transposase and inactivated derivatives | | | |
| *insC* | CDS | IS2 insertion element repressor InsA | Cytoplasmic | COG2963;Transposase and inactivated derivatives | | | |

| | | | | | | | |
|---|---|---|---|---|---|---|---|
| *insC* | CDS | CP4-44 prophage; IS2 insertion element repressor InsA | Cytoplasmic | | | | |
| *insC* | CDS | IS2 insertion element repressor InsA | Cytoplasmic | COG2963;Transposase and inactivated derivatives | | | |
| *insC* | CDS | IS2 insertion element repressor InsA | Cytoplasmic | COG2963;Transposase and inactivated derivatives | | | |
| *insC* | CDS | KpLE2 phage-like element; IS2 insertion element repressor InsA | Cytoplasmic | COG2963;Transposase and inactivated derivatives | | | |
| *insD* | CDS | IS2 element protein | Cytoplasmic | | | | |
| *insD* | CDS | IS2 element protein | Cytoplasmic | | | | |
| *insD* | CDS | IS2 element protein | Cytoplasmic | | | | |
| *insD* | CDS | IS2 element protein | Cytoplasmic | | | | |
| *insD* | CDS | IS2 insertion element transposase InsAB' | Cytoplasmic | COG2801;Transposase and inactivated derivatives | | | |
| *insD* | CDS | IS2 insertion element transposase InsAB' | Cytoplasmic | COG2801;Transposase and inactivated derivatives | | | |
| *insD* | CDS | Qin prophage; predicted transposase | Cytoplasmic | COG2801;Transposase and inactivated derivatives | | | |
| *insD* | CDS | CP4-44 prophage; IS2 insertion element transposase InsAB' | Cytoplasmic | COG2801;Transposase and inactivated derivatives | | | |
| *insD* | CDS | IS2 insertion element transposase InsAB' | Cytoplasmic | COG2801;Transposase and inactivated derivatives | | | |
| *insD* | CDS | IS2 insertion element transposase InsAB' | Cytoplasmic | COG2801;Transposase and inactivated derivatives | | | |
| *insD* | CDS | KpLE2 phage-like element; IS2 insertion element transposase InsAB' | Cytoplasmic | COG2801;Transposase and inactivated derivatives | | | |
| *insE* | CDS | IS3 element protein | Cytoplasmic | COG2963;Transposase and inactivated derivatives | | | |
| *insE* | CDS | IS3 element protein | Cytoplasmic | COG2963;Transposase and inactivated derivatives | | | |
| *insE* | CDS | DLP12 prophage; IS3 element protein InsE | Cytoplasmic | COG2963;Transposase and inactivated derivatives | | | |
| *insE* | CDS | IS3 element protein | Cytoplasmic | COG2963;Transposase and inactivated derivatives | | | |
| *insE* | CDS | IS3 element protein InsE | Cytoplasmic | COG2963;Transposase and inactivated derivatives | | | |
| *insF* | CDS | IS3 element protein InsF | Cytoplasmic | COG2801;Transposase and inactivated derivatives | | | |
| *insF* | CDS | IS3 element protein InsF | Cytoplasmic | COG2801;Transposase and inactivated derivatives | | | |
| *insF* | CDS | DLP12 prophage; IS3 element protein InsF | Cytoplasmic | COG2801;Transposase and inactivated derivatives | | | |
| *insF* | CDS | IS3 element protein InsF | Cytoplasmic | COG2801;Transposase and inactivated derivatives | | | |
| *insF* | CDS | IS3 element protein InsF | Cytoplasmic | COG2801;Transposase and inactivated derivatives | | | |
| *insG* | CDS | KpLE2 phage-like element; IS4 predicted | Cytoplasmic | COG3385;FOG: Transposase and inactivated derivatives | | | |
| *insH* | CDS | IS5 element protein | Cytoplasmic | | | | |
| *insH* | CDS | IS5 element protein | Cytoplasmic | | | | |
| *insH* | CDS | IS5 element protein | Cytoplasmic | | | | |
| *insH* | CDS | IS5 element protein | Cytoplasmic | | | | |
| *insH* | CDS | IS5 element protein | Cytoplasmic | | | | |
| *insH* | CDS | IS5 element protein | Cytoplasmic | | | | |
| *insH* | CDS | IS5 element protein | Cytoplasmic | | | | |
| *insH* | CDS | CP4-6 prophage; IS5 transposase and trans- | Cytoplasmic | COG3039;Transposase and inactivated derivatives, IS5 family | | | |
| *insH* | CDS | DLP12 prophage; IS5 transposase and trans- | Cytoplasmic | COG3039;Transposase and inactivated derivatives, IS5 family | | | |
| *insH* | CDS | IS5 transposase and trans-activator | Cytoplasmic | COG3039;Transposase and inactivated derivatives, IS5 family | | | |
| *insH* | CDS | IS5 transposase and trans-activator | Cytoplasmic | COG3039;Transposase and inactivated derivatives, IS5 family | | | |
| *insH* | CDS | Rac prophage; IS5 transposase and trans-activator | Cytoplasmic | COG3039;Transposase and inactivated derivatives, IS5 family | | | |
| *insH* | CDS | CP4-44 prophage; IS5 transposase and trans- | Cytoplasmic | COG3039;Transposase and inactivated derivatives, IS5 family | | | |
| *insH* | CDS | IS5 transposase and trans-activator | Cytoplasmic | COG3039;Transposase and inactivated derivatives, IS5 family | | | |
| *insH* | CDS | IS5 transposase and trans-activator | Cytoplasmic | COG3039;Transposase and inactivated derivatives, IS5 family | | | |
| *insH* | CDS | IS5 transposase and trans-activator | Cytoplasmic | COG3039;Transposase and inactivated derivatives, IS5 family | | | |
| *insH* | CDS | IS5 transposase and trans-activator | Cytoplasmic | COG3039;Transposase and inactivated derivatives, IS5 family | | | |
| *insH* | CDS | IS5 transposase and trans-activator | Cytoplasmic | COG3039;Transposase and inactivated derivatives, IS5 family | | | |
| *insI* | CDS | CP4-6 prophage; IS30 transposase | Cytoplasmic | COG2826;Transposase and inactivated derivatives, IS30 family | | | |
| *insI* | CDS | IS30 transposase | Cytoplasmic | COG2826;Transposase and inactivated derivatives, IS30 family | | | |
| *insI* | CDS | KpLE2 phage-like element; IS30 transposase | Cytoplasmic | COG2826;Transposase and inactivated derivatives, IS30 family | | | |
| *insJ* | CDS | IS150 protein InsA. Couples formation of linear intermediate to transposition. | Cytoplasmic | COG2963;Transposase and inactivated derivatives | | | |
| *insK* | CDS | IS150 conserved protein InsB | Cytoplasmic | COG2801;Transposase and inactivated derivatives | | | |
| *insL* | CDS | IS186/IS421 transposase | Cytoplasmic | COG3385;FOG: Transposase and inactivated derivatives | | | |
| *insL* | CDS | IS186/IS421 transposase | Cytoplasmic | COG3385;FOG: Transposase and inactivated derivatives | | | |
| *insL* | CDS | predicted transposase | Cytoplasmic | COG3385;FOG: Transposase and inactivated derivatives | | | |
| *insM* | CDS | KpLE2 phage-like element; predicted transposase fragment (pseudogene) | Cytoplasmic | | | | |
| *insN* | CDS | CP4-6 prophage; partial regulator of insertion element IS911A | Cytoplasmic | COG2963;Transposase and inactivated derivatives | | | |
| *insN* | CDS | KpLE2 phage-like element; partial regulator of insertion element IS911B. Interrupted by insertion of IS30 element. | Cytoplasmic | | | | |
| *insO* | CDS | CP4-6 prophage; partial transposase of insertion element IS911A, interrupted by insertion of IS30 element. | Cytoplasmic | COG2801;Transposase and inactivated derivatives | | o | tt |
| *insO* | CDS | KpLE2 phage-like element; partial transposase (pseudogene) | Cytoplasmic | COG2801;Transposase and inactivated derivatives | | | |
| *intA* | CDS | CP4-57 prophage; integrase | Cytoplasmic | COG0582;Integrase | | o | tt |
| *intB* | CDS | KpLE2 phage-like element; predicted integrase | Cytoplasmic | COG0582;Integrase | | o | tt |
| *intD* | CDS | DLP12 prophage; predicted integrase | Cytoplasmic | COG0582;Integrase | | o | tt |
| *intE* | CDS | e14 prophage; predicted integrase | Cytoplasmic | COG0582;Integrase | | o | tt |
| *intF* | CDS | CP4-6 prophage; predicted phage integrase | Cytoplasmic | COG0582;Integrase | | o | tt |
| *intG* | CDS | predicted defective phage integrase (pseudogene) | Cytoplasmic | COG0582;Integrase | | | |
| *intQ* | CDS | Qin prophage; predicted defective integrase | Cytoplasmic | COG0582;Integrase | | gg | o |
| *intR* | CDS | Rac prophage; integrase | Cytoplasmic | COG0582;Integrase | | o | tt |
| *intS* | CDS | CPS-53 (KpLE1) prophage; predicted prophage CPS-53 integrase | Cytoplasmic | COG0582;Integrase | | g | tt |
| *intZ* | CDS | CPZ-55 prophage; predicted integrase | Cytoplasmic | COG0582;Integrase | | | |
| *iscA* | CDS | FeS cluster assembly protein | Cytoplasmic | COG0316;Uncharacterized conserved protein | GO:0006457 protein folding | gg | o |
| *iscR* | CDS | DNA-binding transcriptional activator | Cytoplasmic | COG1959;Predicted transcriptional regulator | GO:0006412 protein biosynthesis | gg | o |
| *iscS* | CDS | cysteine desulfurase (tRNA sulfurtransferase), PLP-dependent | Cytoplasmic | | GO:0009451 RNA modification | gg | t |
| *iscU* | CDS | scaffold protein | Cytoplasmic | COG0822;NifU homolog involved in Fe-S cluster formation | GO:0006412 protein biosynthesis | gg | t |
| *ispA* | CDS | geranyltranstransferase | Cytoplasmic | COG0142;Geranylgeranyl pyrophosphate synthase | | gg | t |
| *ispB* | CDS | octaprenyl diphosphate synthase | Cytoplasmic | COG0142;Geranylgeranyl pyrophosphate synthase | | gg | tt |
| *ispD* | CDS | 4-diphosphocytidyl-2C-methyl-D-erythritol synthase | Cytoplasmic | COG1211;4-diphosphocytidyl-2-methyl-D-erithritol synthase | | gg | t |
| *ispE* | CDS | 4-diphosphocytidyl-2-C-methylerythritol kinase | Cytoplasmic | COG1947;4-diphosphocytidyl-2C-methyl-D-erythritol 2-phosphate synthase | | gg | o |
| *ispF* | CDS | 2C-methyl-D-erythritol 2,4-cyclodiphosphate synthase | Cytoplasmic | COG0245;2C-methyl-D-erythritol 2,4-cyclodiphosphate synthase | | gg | tt |
| *ispG* | CDS | 1-hydroxy-2-methyl-2-(E)-butenyl 4-diphosphate synthase | Cytoplasmic | COG0821;Enzyme involved in the deoxyxylulose pathway of isoprenoid biosynthesis | | gg | t |
| *ispH* | CDS | 1-hydroxy-2-methyl-2-(E)-butenyl 4-diphosphate reductase, 4Fe-4S protein | Cytoplasmic | COG0761;Penicillin tolerance protein | GO:0042493 response to drug | g | tt |
| *ispU* | CDS | undecaprenyl pyrophosphate synthase | Cytoplasmic | COG0020;Undecaprenyl pyrophosphate synthase | | gg | tt |
| *isrA* | misc_RNA | small RNA | | | | | |
| *isrB* | misc_RNA | small RNA | | | | | |
| *isrC* | misc_RNA | small RNA | | | | | |
| *ivbL* | CDS | ilvB operon leader peptide | Cytoplasmic | | | gg | o |
| *ivy* | CDS | inhibitor of vertebrate C-lysozyme | Periplasmic | | | gg | tt |
| *katE* | CDS | hydroperoxidase HPII(III) (catalase) | Cytoplasmic | | GO:0006805 xenobiotic metabolism | gg | o |
| *katG* | CDS | catalase/hydroperoxidase HPI(I) | Cytoplasmic | COG0376;Catalase (peroxidase I) | GO:0006805 xenobiotic metabolism | g | tt |
| *kbaY* | CDS | tagatose 6-phosphate aldolase 1, kbaY subunit | Cytoplasmic | COG0191;Fructose/tagatose bisphosphate aldolase | GO:0016052 carbohydrate | gg | t |
| *kbaZ* | CDS | tagatose 6-phosphate aldolase 1, kbaZ subunit | Cytoplasmic | COG4573;Predicted tagatose 6-phosphate kinase | GO:0016052 carbohydrate | gg | t |
| *kbl* | CDS | glycine C-acetyltransferase | Cytoplasmic | COG0156;7-keto-8-aminopelargonate synthetase and related enzymes | GO:0009063 amino acid catabolism | gg | t |
| *kch* | CDS | voltage-gated potassium channel | Integral Membrane Protein | COG1226;Kef-type K+ transport systems, predicted NAD-binding | | gg | tt |
| *kdgK* | CDS | ketodeoxygluconokinase | Cytoplasmic | COG0524;Sugar kinases, ribokinase family | GO:0016052 carbohydrate | | |
| *kdgR* | CDS | predicted DNA-binding transcriptional regulator | Cytoplasmic | COG1414;Transcriptional regulator | GO:0006350 transcription | g | tt |
| *kdgT* | CDS | 2-keto-3-deoxy-D-gluconate transporter | Integral Membrane Protein | | GO:0016052 carbohydrate | | |
| *kdpA* | CDS | potassium translocating ATPase, subunit A | Integral Membrane Protein | COG2060;K+-transporting ATPase, A chain | | o | tt |
| *kdpB* | CDS | potassium translocating ATPase, subunit B | Integral Membrane Protein | COG2216;High-affinity K+ transport system, ATPase chain B | | o | tt |
| *kdpC* | CDS | potassium translocating ATPase, subunit C | Membrane Anchored | COG2156;K+-transporting ATPase, c chain | | o | tt |
| *kdpD* | CDS | fused sensory histidine kinase in two-component regulatory system with KdpE | Integral Membrane Protein | COG2205;Osmosensitive K+ channel histidine kinase | GO:0006464 protein modification | o | tt |
| *kdpE* | CDS | DNA-binding response regulator in two-component regulatory system with KdpD | Cytoplasmic | COG0745;Response regulators consisting of a CheY-like receiver domain and a winged-helix DNA-binding domain | GO:0006350 transcription | | |
| *kdpF* | CDS | potassium ion accessory transporter subunit | Membrane Anchored | | | o | tt |
| *kdsA* | CDS | 3-deoxy-D-manno-octulosonate 8-phosphate synthase | Cytoplasmic | COG2877;3-deoxy-D-manno-octulosonic acid (KDO) 8-phosphate synthase | GO:0009244 lipopolysaccharide core region biosynthesis -!- GO:0009248 K antigen biosynthesis | gg | t |
| *kdsB* | CDS | 3-deoxy-manno-octulosonate cytidylyltransferase | Cytoplasmic | COG1212;CMP-2-keto-3-deoxyoctulosonic acid synthetase | GO:0009244 lipopolysaccharide core region biosynthesis -!- GO:0009245 lipid A biosynthesis -!- GO:0009248 K antigen biosynthesis | gg | t |
| *kdsC* | CDS | 3-deoxy-D-manno-octulosonate 8-phosphate phosphatase | Periplasmic | COG1778;Low specificity phosphatase (HAD superfamily) | | o | tt |
| *kdsD* | CDS | D-arabinose 5-phosphate isomerase | Cytoplasmic | COG0794;Predicted sugar phosphate isomerase involved in capsule formation -!- COG0517;FOG: CBS domain | | gg | o |
| *kdtA* | CDS | 3-deoxy-D-manno-octulosonic-acid transferase (KDO transferase) | Cytoplasmic | COG1519;3-deoxy-D-manno-octulosonic-acid transferase | GO:0009245 lipid A biosynthesis -!- GO:0009244 lipopolysaccharide core region biosynthesis -!- GO:0009248 K antigen biosynthesis | o | tt |
| *kduD* | CDS | 2-deoxy-D-gluconate 3-dehydrogenase | Cytoplasmic | COG1028;Dehydrogenases with different specificities (related to short-chain alcohol dehydrogenases) | | gg | t |
| *kduI* | CDS | predicted 5-keto 4-deoxyuronate isomerase | Cytoplasmic | COG3717;5-keto 4-deoxyuronate isomerase | GO:0016052 carbohydrate | gg | t |
| *kefA* | CDS | fused mechanosensitive channel proteins | Integral Membrane Protein | COG3264;Small-conductance mechanosensitive channel | | o | tt |

| | | | | | | | |
|---|---|---|---|---|---|---|---|
| *kefB* | CDS | potassium:proton antiporter | Integral Membrane Protein | COG0475;Kef-type K+ transport systems, membrane components -!- COG1226;Kef-type K+ transport systems, predicted NAD-binding | | o | tt |
| *kefC* | CDS | potassium:proton antiporter | Integral Membrane Protein | COG0475;Kef-type K+ transport systems, membrane components -!- COG1226;Kef-type K+ transport systems, predicted NAD-binding | | o | tt |
| *kefF* | CDS | flavoprotein subunit for the KefC potassium efflux system | Cytoplasmic | COG2249;Putative NADPH-quinone reductase (modulator of drug activity B) | | gg | o |
| *kefG* | CDS | component of potassium efflux complex with KefB | Cytoplasmic | COG2249;Putative NADPH-quinone reductase (modulator of drug activity | | gg | o |
| *kgtP* | CDS | alpha-ketoglutarate transporter | Integral Membrane Protein | COG0477;Permeases of the major facilitator superfamily | GO:0016052 carbohydrate | o | tt |
| *kil* | CDS | Rac prophage; inhibitor of ftsZ, killing protein | Cytoplasmic | | | gg | tt |
| *kptA* | CDS | 2'-phosphotransferase | Cytoplasmic | | | | |
| *ksgA* | CDS | S-adenosylmethionine-6-N',N'-adenosyl (rRNA) dimethyltransferase | Cytoplasmic | COG0030;Dimethyladenosine transferase (rRNA methylation) | GO:0009451 RNA modification -!- GO:0042493 response to drug | gg | t |
| *lacA* | CDS | thiogalactoside acetyltransferase | Cytoplasmic | COG0110;Acetyltransferase (isoleucine patch superfamily) | GO:0016052 carbohydrate | gg | o |
| *lacI* | CDS | DNA-binding transcriptional repressor | Cytoplasmic | | GO:0016052 carbohydrate catabolism -!- GO:0006350 | o | tt |
| *lacY* | CDS | lactose/galactose transporter | Integral Membrane Protein | COG0477;Permeases of the major facilitator superfamily | GO:0016052 carbohydrate | o | tt |
| *lacZ* | CDS | beta-D-galactosidase | Cytoplasmic | COG3250;Beta-galactosidase/beta-glucuronidase | GO:0016052 carbohydrate | gg | t |
| *lamB* | CDS | maltose outer membrane porin (maltoporin) | Outer Membrane B-barrel protein | COG4580;Maltoporin (phage lambda and maltose receptor) | | o | tt |
| *lar* | CDS | Rac prophage; restriction alleviation protein | Cytoplasmic | | | | |
| *ldcA* | CDS | L,D-carboxypeptidase A | Cytoplasmic | COG1619;Uncharacterized proteins, homologs of microcin C7 resistance protein MccF | GO:0000270 peptidoglycan metabolism | gg | t |
| *ldcC* | CDS | lysine decarboxylase 2, constitutive | Cytoplasmic | COG1982;Arginine/lysine/ornithine decarboxylases | GO:0006096 glycolysis -!- GO:0016052 carbohydrate | o | tt |
| *ldhA* | CDS | fermentative D-lactate dehydrogenase, NAD-dependent | Cytoplasmic | COG1052;Lactate dehydrogenase and related dehydrogenases | GO:0006113 fermentation | gg | t |
| *ldrA* | CDS | toxic polypeptide, small | Cytoplasmic | | | | |
| *ldrB* | CDS | toxic polypeptide, small | Cytoplasmic | | | | |
| *ldrC* | CDS | toxic polypeptide, small | Cytoplasmic | | | | |
| *ldrD* | CDS | toxic polypeptide, small | Cytoplasmic | | | | |
| *lepA* | CDS | GTP binding membrane protein | Cytoplasmic | COG0481;Membrane GTPase LepA | GO:0006412 protein biosynthesis | o | tt |
| *lepB* | CDS | leader peptidase (signal peptidase I) | Integral Membrane Protein | COG0681;Signal peptidase I | | o | tt |
| *leuA* | CDS | 2-isopropylmalate synthase | Cytoplasmic | COG0119;Isopropylmalate/homocitrate/citramalate synthases | GO:0009098 leucine biosynthesis | gg | t |
| *leuB* | CDS | 3-isopropylmalate dehydrogenase | Cytoplasmic | | GO:0009098 leucine biosynthesis | | |
| *leuC* | CDS | 3-isopropylmalate isomerase subunit, dehydratase component | Cytoplasmic | COG0065;3-isopropylmalate dehydratase large subunit | GO:0009098 leucine biosynthesis | gg | t |
| *leuD* | CDS | 3-isopropylmalate isomerase subunit | Cytoplasmic | COG0066;3-isopropylmalate dehydratase small subunit | GO:0009098 leucine biosynthesis | gg | o |
| *leuL* | CDS | leu operon leader peptide | Cytoplasmic | | GO:0009098 leucine biosynthesis | gg | o |
| *leuO* | CDS | DNA-binding transcriptional activator | Cytoplasmic | | GO:0009098 leucine biosynthesis -!- GO:0006350 transcription | gg | t |
| *leuP* | tRNA | tRNA-Leu(CAG) (Leucine tRNA1) | | | | | |
| *leuQ* | tRNA | tRNA-Leu(CAG) (Leucine tRNA1) | | | | | |
| *leuS* | CDS | leucyl-tRNA synthetase | Cytoplasmic | COG0495;Leucyl-tRNA synthetase | GO:0006418 amino acid activation | o | tt |
| *leuT* | tRNA | tRNA-Leu(CAG) (Leucine tRNA10 | | | | | |
| *leuU* | tRNA | tRNA-Leu(GAG) (Leucine tRNA2) | | | | | |
| *leuV* | tRNA | tRNA-Leu(CAG) (Leucine tRNA1) | | | | | |
| *leuW* | tRNA | tRNA-Leu(UAG) (Leucine tRNA3) | | | | | |
| *leuX* | tRNA | tRNA-Leu(CAA) (Leucine tRNA5) | | | | | |
| *leuZ* | tRNA | tRNA-Leu(UAA) (Leucine tRNA4) | | | | | |
| *lexA* | CDS | DNA-binding transcriptional repressor | Cytoplasmic | COG1974;SOS-response transcriptional repressors (RecA-mediated autopeptidases) | GO:0006281 DNA repair -!- GO:0006350 transcription -!- GO:0009432 SOS response | o | tt |
| *lgt* | CDS | phosphatidylglycerol-prolipoprotein diacylglyceryl transferase | Integral Membrane Protein | COG0682;Prolipoprotein diacylglyceryltransferase | GO:0008654 phospholipid biosynthesis -!- GO:0006464 protein modification | o | tt |
| *lhr* | CDS | predicted ATP-dependent helicase | Cytoplasmic | COG1201;Lhr-like helicases | GO:0006261 DNA dependent DNA replication | o | t |
| *ligA* | CDS | DNA ligase, NAD(+)-dependent | Cytoplasmic | COG0272;NAD-dependent DNA ligase (contains BRCT domain type II) | GO:0006281 DNA repair | gg | tt |
| *ligB* | CDS | DNA ligase, NAD(+)-dependent | Cytoplasmic | COG0272;NAD-dependent DNA ligase (contains BRCT domain type II) | GO:0006259 DNA metabolism | o | tt |
| *ligT* | CDS | 2'-5' RNA ligase | Cytoplasmic | | GO:0009451 RNA modification | | |
| *lipA* | CDS | lipoate synthase | Cytoplasmic | COG0320;Lipoate synthase | GO:0009107 lipoate biosynthesis | gg | t |
| *lipB* | CDS | lipoyl-protein ligase | Cytoplasmic | COG0321;Lipoate-protein ligase B | GO:0009107 lipoate biosynthesis | | |
| *lit* | CDS | e14 prophage; cell death peptidase, inhibitor of T4 late gene expression | Cytoplasmic | | | o | o |
| *livF* | CDS | leucine/isoleucine/valine transporter subunit -!- ATP-binding component of ABC superfamily | Cytoplasmic | | GO:0009063 amino acid catabolism -!- GO:0009098 leucine biosynthesis | gg | t |
| *livG* | CDS | leucine/isoleucine/valine transporter subunit -!- ATP-binding component of ABC superfamily | Cytoplasmic | COG0411;ABC-type branched-chain amino acid transport systems, ATPase component | GO:0009063 amino acid catabolism -!- GO:0009098 leucine biosynthesis | g | tt |
| *livH* | CDS | leucine/isoleucine/valine transporter subunit -!- membrane component of ABC superfamily | Integral Membrane Protein | COG0559;Branched-chain amino acid ABC-type transport system, permease components | GO:0009098 leucine biosynthesis | o | tt |
| *livJ* | CDS | leucine/isoleucine/valine transporter subunit -!- periplasmic-binding component of ABC superfamily | Periplasmic | | GO:0009098 leucine biosynthesis | o | tt |
| *livK* | CDS | leucine transporter subunit -!- periplasmic-binding component of ABC superfamily | Periplasmic | COG0683;ABC-type branched-chain amino acid transport systems, periplasmic component | GO:0009098 leucine biosynthesis | o | tt |
| *livM* | CDS | leucine/isoleucine/valine transporter subunit -!- membrane component of ABC superfamily | Integral Membrane Protein | COG4177;ABC-type branched-chain amino acid transport system, permease component | GO:0009098 leucine biosynthesis | o | tt |
| *lldD* | CDS | L-lactate dehydrogenase, FMN-linked | Cytoplasmic | COG1304;L-lactate dehydrogenase (FMN-dependent) and related alpha-hydroxy acid dehydrogenases | GO:0009060 aerobic respiration -!- GO:0009061 anaerobic respiration | gg | t |
| *lldP* | CDS | L-lactate permease | Integral Membrane Protein | COG1620;L-lactate permease | GO:0016052 carbohydrate | o | tt |
| *lldR* | CDS | DNA-binding transcriptional repressor | Cytoplasmic | COG2186;Transcriptional regulators | GO:0009060 aerobic respiration -!- GO:0006350 transcription | o | tt |
| *lnt* | CDS | apolipoprotein N-acyltransferase | Integral Membrane Protein | COG0815;Apolipoprotein N-acyltransferase | GO:0042158 lipoprotein biosynthesis | o | tt |
| *lolA* | CDS | chaperone for lipoproteins | Periplasmic | COG2834;Outer membrane lipoprotein-sorting protein | | o | tt |
| *lolB* | CDS | chaperone for lipoproteins | Outer Membrane Lipoprotein | COG3017;Outer membrane lipoprotein involved in outer membrane biogenesis | | gg | tt |
| *lolC* | CDS | outer membrane-specific lipoprotein transporter subunit -!- membrane component of ABC superfamily | Integral Membrane Protein | COG4591;ABC-type transport system, involved in lipoprotein release, permease component | | | |
| *lolD* | CDS | outer membrane-specific lipoprotein transporter subunit -!- ATP-binding component of ABC superfamily | Cytoplasmic | | | | |
| *lolE* | CDS | outer membrane-specific lipoprotein transporter subunit -!- membrane component of ABC superfamily | Integral Membrane Protein | COG4591;ABC-type transport system, involved in lipoprotein release, permease component | | o | tt |
| *lomR* | CDS | Rac prophage; predicted protein, N-ter fragment (pseudogene) | Cytoplasmic | COG3637;Opacity protein and related surface antigens | | | |
| *lomR* | ancestral | Rac prophage; predicted protein (pseudogene) | Cytoplasmic | COG3637;Opacity protein and related surface antigens | | | |
| *lomR* | CDS | Rac prophage; predicted protein, C-ter fragment (pseudogene) | Cytoplasmic | COG3637;Opacity protein and related surface antigens | | | |
| *lon* | CDS | DNA-binding ATP-dependent protease La | Cytoplasmic | COG0466;ATP-dependent Lon protease, bacterial type | | o | tt |
| *lpcA* | CDS | D-sedoheptulose 7-phosphate isomerase | Cytoplasmic | COG0279;Phosphoheptose isomerase | GO:0009244 lipopolysaccharide core region biosynthesis | g | t |
| *lpd* | CDS | lipoamide dehydrogenase, E3 component is part of three enzyme complexes | Cytoplasmic | COG1249;Pyruvate/2-oxoglutarate dehydrogenase complex, dihydrolipoamide dehydrogenase (E3) component, and related enzymes | GO:0016052 carbohydrate catabolism -!- GO:0009063 amino acid catabolism -!- GO:0006096 glycolysis -!- GO:0006086 acetyl-CoA biosynthesis from pyruvate -!- GO:0006099 tricarboxylic acid cycle -!- GO:0009060 aerobic respiration -!- GO:0009061 anaerobic res | gg | o |
| *lplA* | CDS | lipoate-protein ligase A | Cytoplasmic | COG0095;Lipoate-protein ligase A | GO:0042158 lipoprotein biosynthesis -!- GO:0006418 amino acid activation | gg | t |
| *lpp* | CDS | murein lipoprotein | Outer Membrane Lipoprotein | COG4238;Murein lipoprotein | GO:0009252 peptidoglycan biosynthesis | o | tt |
| *lpxA* | CDS | UDP-N-acetylglucosamine acetyltransferase | Cytoplasmic | COG1043;Acyl-[acyl carrier protein]--UDP-N-acetylglucosamine O-acyltransferase | GO:0009245 lipid A biosynthesis | gg | o |
| *lpxB* | CDS | tetraacyldisaccharide-1-P synthase | Cytoplasmic | COG0763;Lipid A disaccharide synthetase | GO:0009245 lipid A biosynthesis | o | tt |
| *lpxC* | CDS | UDP-3-O-acyl N-acetylglucosamine deacetylase | Cytoplasmic | COG0774;UDP-3-O-acyl-N-acetylglucosamine deacetylase | GO:0009245 lipid A biosynthesis | o | tt |
| *lpxD* | CDS | UDP-3-O-(3-hydroxymyristoyl)-glucosamine N-acyltransferase | Cytoplasmic | COG1044;UDP-3-O-[3-hydroxymyristoyl] glucosamine N-acyltransferase | GO:0009245 lipid A biosynthesis | gg | tt |
| *lpxH* | CDS | UDP-2,3-diacylglucosamine pyrophosphatase | Cytoplasmic | COG2908;Uncharacterized protein conserved in bacteria | GO:0009245 lipid A biosynthesis | o | tt |
| *lpxK* | CDS | lipid A 4'kinase | Integral Membrane Protein | COG1663;Tetraacyldisaccharide-1-P 4'-kinase | GO:0009244 lipopolysaccharide core region biosynthesis -!- GO:0009245 lipid A biosynthesis -!- GO:0009248 K antigen biosynthesis | o | t |
| *lpxL* | CDS | lauryl-acyl carrier protein (ACP)-dependent acyltransferase | Membrane Anchored | COG1560;Lauroyl/myristoyl acyltransferase | GO:0009245 lipid A biosynthesis | o | tt |
| *lpxM* | CDS | myristoyl-acyl carrier protein (ACP)-dependent acyltransferase | Membrane Anchored | COG1560;Lauroyl/myristoyl acyltransferase | GO:0009245 lipid A biosynthesis | gg | tt |
| *lrhA* | CDS | DNA-binding transcriptional repressor | Cytoplasmic | COG0583;Transcriptional regulator | GO:0009060 aerobic respiration -!- GO:0006350 transcription | gg | tt |
| *lrp* | CDS | DNA-binding transcriptional dual regulator, leucine-binding | Cytoplasmic | COG1522;Transcriptional regulators | GO:0009098 leucine biosynthesis -!- GO:0006350 transcription | gg | tt |
| *lspA* | CDS | prolipoprotein signal peptidase (signal peptidase II) | Integral Membrane Protein | COG0597;Lipoprotein signal peptidase | | o | tt |
| *lsrB* | CDS | AI2 transporter -!- periplasmic-binding component of ABC superfamily | Periplasmic | COG1879;ABC-type sugar transport system, periplasmic component | | o | tt |
| *lsrC* | CDS | AI2 transporter -!- membrane component of ABC superfamily | Integral Membrane Protein | COG1172;Ribose/xylose/arabinose/galactoside ABC-type transport systems, permease components | | o | tt |
| *lsrD* | CDS | AI2 transporter -!- membrane component of ABC superfamily | Integral Membrane Protein | COG1172;Ribose/xylose/arabinose/galactoside ABC-type transport systems, permease components | | o | tt |
| *lsrF* | CDS | predicted aldolase | Cytoplasmic | COG1830;DhnA-type fructose-1,6-bisphosphate aldolase and related enzymes | | gg | t |
| *lsrG* | CDS | conserved protein | Cytoplasmic | COG1359;Uncharacterized conserved protein | | gg | t |

| | | | | | | | |
|---|---|---|---|---|---|---|---|
| *ltaE* | CDS | L-allo-threonine aldolase, PLP-dependent | Cytoplasmic | COG2008;Threonine aldolase | GO:0006545 glycine biosynthesis | gg | t |
| *luxS* | CDS | S-ribosylhomocysteinase | Cytoplasmic | COG1854;LuxS protein involved in autoinducer AI2 synthesis | | gg | o |
| *lysA* | CDS | diaminopimelate decarboxylase, PLP-binding | Cytoplasmic | COG0019;Diaminopimelate decarboxylase | GO:0009089 lysine biosynthesis via diaminopimelate | o | tt |
| *lysC* | CDS | aspartokinase III | Cytoplasmic | COG0527;Aspartokinases | GO:0009089 lysine biosynthesis via diaminopimelate -!- GO:0009090 homoserine biosynthesis | g | o |
| *lysP* | CDS | lysine transporter | Integral Membrane Protein | COG0833;Amino acid transporters | GO:0009089 lysine biosynthesis via diaminopimelate | o | tt |
| *lysQ* | tRNA | tRNA-Lys(UUU) (Lysine tRNA) | | | | | |
| *lysR* | CDS | DNA-binding transcriptional dual regulator | Cytoplasmic | COG0583;Transcriptional regulator | GO:0009089 lysine biosynthesis via diaminopimelate -!- GO:0006350 transcription | gg | tt |
| *lysS* | CDS | lysine tRNA synthetase, constitutive | Cytoplasmic | COG1190;Lysyl-tRNA synthetase (class II) | GO:0006418 amino acid activation | gg | t |
| *lysT* | tRNA | tRNA-Lys(UUU) (Lysine tRNA) | | | | | |
| *lysU* | CDS | lysine tRNA synthetase, inducible | Cytoplasmic | COG1190;Lysyl-tRNA synthetase (class II) | GO:0006418 amino acid activation | gg | t |
| *lysV* | tRNA | tRNA-Lys(UUU) (Lysine tRNA) | | | | | |
| *lysW* | tRNA | tRNA-Lys(UUU) (Lysine tRNA) | | | | | |
| *lysY* | tRNA | tRNA-Lys(UUU) (Lysine tRNA) | | | | | |
| *lysZ* | tRNA | tRNA-Lys(UUU), (Lysine tRNA) | | | | | |
| *lyxK* | CDS | L-xylulose kinase | Cytoplasmic | COG1070;Sugar (pentulose and hexulose) kinases | GO:0016052 carbohydrate | o | tt |
| *maa* | CDS | maltose O-acetyltransferase | Cytoplasmic | COG0110;Acetyltransferase (isoleucine patch superfamily) | GO:0016052 carbohydrate | gg | o |
| *macA* | CDS | macrolide transporter subunit, membrane fusion protein (MFP) component | Membrane Anchored | | | o | tt |
| *macB* | CDS | fused macrolide transporter subunits -!- ATP-binding component and membrane component of ABC superfamily | Integral Membrane Protein | COG1136;ABC-type antimicrobial peptide transport system, ATPase component -!- COG0577;ABC-type antimicrobial peptide transport system, permease component | | o | tt |
| *maeB* | CDS | fused malic enzyme predicted oxidoreductase -!- predicted phosphotransacetylase | Cytoplasmic | COG0281;Malic enzyme -!- COG0280;Phosphotransacetylase | | g | t |
| *mak* | CDS | manno(fructo)kinase | Cytoplasmic | COG1940;Transcriptional regulator/sugar kinase | GO:0016052 carbohydrate | gg | t |
| *malE* | CDS | maltose transporter subunit -!- periplasmic-binding component of ABC superfamily | Periplasmic | COG2182;Maltose-binding periplasmic proteins/domains | GO:0016052 carbohydrate catabolism -!- GO:0006457 protein | o | tt |
| *malF* | CDS | maltose transporter subunit -!- membrane component of ABC superfamily | Integral Membrane Protein | COG1175;ABC-type sugar transport systems, permease components | GO:0016052 carbohydrate catabolism | o | tt |
| *malG* | CDS | maltose transporter subunit -!- membrane component of ABC superfamily | Integral Membrane Protein | COG3833;ABC-type maltose transport systems, permease component | GO:0016052 carbohydrate catabolism | o | tt |
| *malI* | CDS | DNA-binding transcriptional repressor | Cytoplasmic | COG1609;Transcriptional regulators | GO:0016052 carbohydrate catabolism -!- GO:0006350 | o | tt |
| *malK* | CDS | fused maltose transport subunit, ATP-binding component of ABC superfamily -!- regulatory protein | Cytoplasmic | COG3839;ABC-type sugar transport systems, ATPase components | GO:0016052 carbohydrate catabolism | gg | t |
| *malM* | CDS | maltose regulon periplasmic protein | Periplasmic | | GO:0016052 carbohydrate | o | tt |
| *malP* | CDS | maltodextrin phosphorylase | Cytoplasmic | | GO:0016052 carbohydrate catabolism -!- GO:0000272 polysaccharide catabolism | | |
| *malQ* | CDS | 4-alpha-glucanotransferase (amylomaltase) | Cytoplasmic | COG1640;4-alpha-glucanotransferase | GO:0016052 carbohydrate catabolism -!- GO:0000272 polysaccharide catabolism | o | tt |
| *malS* | CDS | alpha-amylase | Periplasmic | COG0366;Glycosidases | GO:0016052 carbohydrate catabolism -!- GO:0000272 polysaccharide catabolism -!- GO:0006508 proteolysis and | o | tt |
| *malT* | CDS | DNA-binding transcriptional activator, maltotriose-ATP-binding | Cytoplasmic | COG2909;ATP-dependent transcriptional regulator | GO:0016052 carbohydrate catabolism -!- GO:0006350 | o | tt |
| *malX* | CDS | fused maltose and glucose-specific PTS enzyme IIBC components | Integral Membrane Protein | COG1263;Phosphotransferase system IIC components, glucose/maltose/N-acetylglucosamine-specific -!- COG1264;Phosphotransferase system IIB components | GO:0016052 carbohydrate catabolism | o | tt |
| *malY* | CDS | bifunctional beta-cystathionase, PLP-dependent -!- regulator of maltose regulon | Cytoplasmic | COG1168;Bifunctional PLP-dependent enzyme with beta-cystathionase and maltose regulon repressor activities | GO:0016052 carbohydrate catabolism | gg | o |
| *malZ* | CDS | maltodextrin glucosidase | Cytoplasmic | COG0366;Glycosidases | GO:0016052 carbohydrate catabolism -!- GO:0000272 polysaccharide catabolism | o | tt |
| *manA* | CDS | mannose-6-phosphate isomerase | Cytoplasmic | | GO:0016052 carbohydrate catabolism -!- GO:0009226 nucleotide-sugar biosynthesis -!- GO:0009242 colanic acid biosynthesis -!- GO:0009269 | gg | o |
| *manX* | CDS | fused mannose-specific PTS enzyme IIAB components | Cytoplasmic | COG2893;Phosphotransferase system, mannose/fructose-specific component IIA -!- COG3444;Phosphotransferase system, mannose/fructose/N-acetylgalactosamine-specific component IIB | GO:0016052 carbohydrate catabolism | gg | tt |
| *manY* | CDS | mannose-specific enzyme IIC component of PTS | Integral Membrane Protein | COG3715;Phosphotransferase system, mannose/fructose/N-acetylgalactosamine-specific component IIC | GO:0016052 carbohydrate catabolism | o | tt |
| *manZ* | CDS | mannose-specific enzyme IID component of PTS | Integral Membrane Protein | | GO:0016052 carbohydrate | o | tt |
| *maoC* | CDS | fused aldehyde dehydrogenase -!- enoyl-CoA hydratase | Cytoplasmic | COG1012;NAD-dependent aldehyde dehydrogenases -!- COG2030;Acyl dehydratase | GO:0016052 carbohydrate catabolism | g | tt |
| *map* | CDS | methionine aminopeptidase | Cytoplasmic | COG0024;Methionine aminopeptidase | GO:0006457 protein folding | gg | o |
| *marA* | CDS | DNA-binding transcriptional dual regulator | Cytoplasmic | COG2207;AraC-type DNA-binding domain-containing proteins | GO:0006350 transcription -!- GO:0042493 response to drug -!- GO:0006805 xenobiotic metabolism | | |
| *marB* | CDS | predicted protein | Periplasmic | | GO:0042493 response to drug -!- GO:0006805 xenobiotic metabolism | g | tt |
| *marC* | CDS | predicted transporter | Integral Membrane Protein | COG2095;Multiple antibiotic transporter | | g | tt |
| *marR* | CDS | DNA-binding transcriptional repressor | Cytoplasmic | | GO:0006350 transcription -!- GO:0042493 response to drug | | |
| *mazG* | CDS | nucleoside triphosphate pyrophosphohydrolase | Cytoplasmic | COG1694;Predicted pyrophosphatase | | gg | tt |
| *mbhA* | CDS | flagellar system protein, promoterless fragment (pseudogene) | Cytoplasmic | COG1360;Flagellar motor protein | GO:0042330 taxis | | |
| *mcrA* | CDS | e14 prophage; 5-methylcytosine-specific restriction endonuclease B | Cytoplasmic | COG1403;Restriction endonuclease | | gg | t |
| *mcrB* | CDS | 5-methylcytosine-specific restriction enzyme McrBC, subunit McrB | Cytoplasmic | | GO:0006308 DNA catabolism -!- GO:0006308 DNA catabolism | | |
| *mcrC* | CDS | 5-methylcytosine-specific restriction enzyme McrBC, subunit McrC | Cytoplasmic | COG4268;McrBC 5-methylcytosine restriction system component | GO:0006308 DNA catabolism -!- GO:0006308 DNA catabolism | | |
| *mdaB* | CDS | NADPH quinone reductase | Cytoplasmic | COG2249;Putative NADPH-quinone reductase (modulator of drug activity | GO:0042493 response to drug | gg | tt |
| *mdh* | CDS | malate dehydrogenase, NAD(P)-binding | Cytoplasmic | COG0039;Malate/lactate dehydrogenases | GO:0016052 carbohydrate catabolism -!- GO:0006099 tricarboxylic acid cycle -!- GO:0006113 fermentation -!- GO:0009061 anaerobic respiration -!- GO:0006094 gluconeogenesis -!- | gg | t |
| *mdlA* | CDS | fused predicted multidrug transporter subunits -!- ATP-binding components of ABC superfamily | Integral Membrane Protein | COG1132;ABC-type multidrug transport system, ATPase and permease components | GO:0042493 response to drug | | |
| *mdlB* | CDS | fused predicted multidrug transporter subunits -!- ATP binding components of ABC superfamily | Integral Membrane Protein | COG1132;ABC-type multidrug transport system, ATPase and permease components | | | |
| *mdoB* | CDS | phosphoglycerol transferase I | Integral Membrane Protein | COG1368;Phosphoglycerol transferase and related proteins, alkaline phosphatase superfamily | GO:0008654 phospholipid biosynthesis -!- GO:0009250 glucan biosynthesis -!- GO:0006970 response to osmotic stress | | |
| *mdoC* | CDS | membrane protein required for modification of periplasmic glucan | Integral Membrane Protein | | GO:0009250 glucan biosynthesis -!- GO:0006970 response to osmotic stress | o | tt |
| *mdoD* | CDS | glucan biosynthesis protein, periplasmic | Periplasmic | | GO:0006970 response to osmotic stress | gg | t |
| *mdoG* | CDS | glucan biosynthesis protein, periplasmic | Periplasmic | COG3131;Periplasmic glucans biosynthesis protein | GO:0009250 glucan biosynthesis -!- GO:0006970 response to osmotic stress | o | tt |
| *mdoH* | CDS | glucan biosynthesis: glycosyl transferase | Integral Membrane Protein | COG2943;Membrane glycosyltransferase | GO:0006970 response to osmotic stress | o | tt |
| *mdtA* | CDS | multidrug efflux system, subunit A | Membrane Anchored | COG0845;Membrane-fusion protein | | | |
| *mdtB* | CDS | multidrug efflux system, subunit B | Integral Membrane Protein | COG0841;Cation/multidrug efflux pump | | o | tt |
| *mdtC* | CDS | multidrug efflux system, subunit C | Integral Membrane Protein | COG0841;Cation/multidrug efflux pump | | o | tt |
| *mdtD* | CDS | multidrug efflux system protein | Integral Membrane Protein | COG0477;Permeases of the major facilitator superfamily | | o | tt |
| *mdtE* | CDS | multidrug resistance efflux transporter | Inner Membrane Lipoprotein | COG0845;Membrane-fusion protein | | o | tt |
| *mdtF* | CDS | multidrug transporter, RpoS-dependent | Integral Membrane Protein | COG0841;Cation/multidrug efflux pump | | o | tt |
| *mdtG* | CDS | predicted drug efflux system | Integral Membrane Protein | COG0477;Permeases of the major facilitator superfamily | | o | tt |
| *mdtH* | CDS | predicted drug efflux system | Integral Membrane Protein | | | o | tt |
| *mdtI* | CDS | multidrug efflux system transporter | Integral Membrane Protein | COG2076;Membrane transporters of cations and cationic drugs | | | |
| *mdtJ* | CDS | multidrug efflux system transporter | Integral Membrane Protein | COG2076;Membrane transporters of cations and cationic drugs | | o | tt |
| *mdtK* | CDS | multidrug efflux system transporter | Integral Membrane Protein | | GO:0042493 response to drug | gg | tt |
| *mdtL* | CDS | multidrug efflux system protein | Integral Membrane Protein | COG0477;Permeases of the major facilitator superfamily | | o | tt |
| *melA* | CDS | alpha-galactosidase, NAD(P)-binding | Cytoplasmic | COG1486;Alpha-galactosidases/6-phospho-beta-glucos idases, family 4 of glycosyl hydrolases | GO:0016052 carbohydrate catabolism -!- GO:0009435 nicotinamide adenine dinucleotide | gg | t |
| *melB* | CDS | melibiose:sodium symporter | Integral Membrane Protein | COG2211;Na+/melibiose symporter and related transporters | GO:0016052 carbohydrate | o | tt |
| *melR* | CDS | DNA-binding transcriptional dual regulator | Cytoplasmic | COG2207;AraC-type DNA-binding domain-containing proteins | GO:0016052 carbohydrate catabolism -!- GO:0006350 | o | tt |
| *menA* | CDS | 1,4-dihydroxy-2-naphthoate octaprenyltransferase | Integral Membrane Protein | COG1575;1,4-dihydroxy-2-naphthoate octaprenyltransferase | GO:0009061 anaerobic respiration | o | tt |
| *menB* | CDS | dihydroxynaphthoic acid synthetase | Cytoplasmic | COG0447;Dihydroxynaphthoic acid synthase | GO:0009061 anaerobic respiration | gg | o |
| *menC* | CDS | o-succinylbenzoyl-CoA synthase | Cytoplasmic | COG1441;O-succinylbenzoate synthase | GO:0009061 anaerobic respiration | gg | tt |
| *menD* | CDS | bifunctional 2-oxoglutarate decarboxylase -!- SHCHC synthase | Cytoplasmic | COG1165;2-succinyl-6-hydroxy-2,4-cyclohexadiene-1- carboxylate synthase | GO:0009061 anaerobic respiration | | |

| | | | | | | | |
|---|---|---|---|---|---|---|---|
| *menE* | CDS | o-succinylbenzoate-CoA ligase | Cytoplasmic | COG0318;Acyl-CoA synthetases (AMP-forming)/AMP-acid ligases II | GO:0009061 anaerobic respiration | gg | t |
| *menF* | CDS | isochorismate synthase 2 | Cytoplasmic | | GO:0009061 anaerobic respiration | gg | t |
| *mepA* | CDS | murein DD-endopeptidase | Periplasmic | COG3770;Murein endopeptidase | GO:0009252 peptidoglycan biosynthesis -!- GO:0042493 response to drug | o | tt |
| *metA* | CDS | homoserine transsuccinylase | Cytoplasmic | COG1897;Homoserine trans-succinylase | GO:0009086 methionine biosynthesis | gg | t |
| *metB* | CDS | cystathionine gamma-synthase, PLP-dependent | Cytoplasmic | COG0626;Cystathionine beta-lyases/cystathionine gamma-synthases | GO:0009063 amino acid catabolism -!- GO:0009086 methionine biosynthesis | gg | t |
| *metC* | CDS | cystathionine beta-lyase, PLP-dependent | Cytoplasmic | COG0626;Cystathionine beta-lyases/cystathionine gamma-synthases | GO:0009086 methionine biosynthesis | g | t |
| *metE* | CDS | 5-methyltetrahydropteroyltriglutamate-homocysteine S-methyltransferase | Cytoplasmic | COG0620;Methionine synthase II (cobalamin-independent) | GO:0009086 methionine biosynthesis -!- GO:0006556 S-adenosylmethionine biosynthesis | o | tt |
| *metF* | CDS | 5,10-methylenetetrahydrofolate reductase | Cytoplasmic | COG0685;5,10-methylenetetrahydrofolate reductase | GO:0009257 10-formyltetrahydrofolate biosynthesis | gg | t |
| *metG* | CDS | methionyl-tRNA synthetase | Cytoplasmic | COG0143;Methionyl-tRNA synthetase -!- COG0073;EMAP domain | GO:0006418 amino acid activation | g | t |
| *metH* | CDS | homocysteine-N5-methyltetrahydrofolate transmethylase, B12-dependent | Cytoplasmic | COG0646;Methionine synthase I (cobalamin-dependent), methyltransferase domain -!- COG1410;Methionine synthase I, cobalamin-binding domain | GO:0009086 methionine biosynthesis -!- GO:0006556 S-adenosylmethionine biosynthesis -!- GO:0009257 10-formyltetrahydrofolate biosynthesis -!- GO:0009236 vitamin B12 | o | tt |
| *metI* | CDS | DL-methionine transporter subunit -!- membrane component protein of ABC superfamily | Integral Membrane Protein | COG2011;ABC-type metal ion transport system, permease component | | o | o |
| *metJ* | CDS | DNA-binding transcriptional repressor, S-adenosylmethionine-binding | Cytoplasmic | COG3060;Transcriptional regulator of met regulon | GO:0009086 methionine biosynthesis -!- GO:0006350 transcription | o | tt |
| *metK* | CDS | methionine adenosyltransferase 1 | Cytoplasmic | COG0192;S-adenosylmethionine synthetase | GO:0006556 S-adenosylmethionine biosynthesis | gg | t |
| *metL* | CDS | fused aspartokinase II -!- homoserine dehydrogenase II | Cytoplasmic | COG0527;Aspartokinases -!- COG0460;Homoserine dehydrogenase | GO:0009086 methionine biosynthesis -!- GO:0009090 homoserine biosynthesis | o | tt |
| *metN* | CDS | DL-methionine transporter subunit -!- ATP-binding component of ABC superfamily | Cytoplasmic | COG1135;ABC-type metal ion transport system, ATPase component | | g | t |
| *metQ* | CDS | DL-methionine transporter subunit -!- periplasmic-binding component of ABC superfamily | Membrane Lipoprotein | COG1464;ABC-type metal ion transport system, periplasmic component/surface antigen | | o | tt |
| *metR* | CDS | DNA-binding transcriptional activator, homocysteine-binding | Cytoplasmic | COG0583;Transcriptional regulator | GO:0009086 methionine biosynthesis -!- GO:0006350 transcription | o | tt |
| *metT* | tRNA | tRNA-Met(CAU) (Methionine tRNAm) | | | | | |
| *metU* | tRNA | tRNA-Met(CAU) (Methionine tRNAm) | | | | | |
| *metV* | tRNA | tRNA-initiator Met(CAU) (Initiator methionine tRNAf1) | | | | | |
| *metW* | tRNA | tRNA-initiator Met(CAU) (Initiator methionine tRNAf1) | | | | | |
| *metY* | tRNA | tRNA-initiator Met(CAU) (Initiator methionine tRNAf2) | | | | | |
| *metZ* | tRNA | tRNA-initiator Met(CAU) (Initiator methionine tRNAf1) | | | | | |
| *mfd* | CDS | transcription-repair coupling factor | Cytoplasmic | COG1197;Transcription-repair coupling factor (superfamily II helicase) | GO:0006350 transcription | o | tt |
| *mglA* | CDS | fused methyl-galactoside transporter subunitsl -!- ATP-binding components of ABC superfamily | Cytoplasmic | COG1129;ABC-type sugar transport system, ATPase component | GO:0016052 carbohydrate catabolism | o | tt |
| *mglB* | CDS | methyl-galactoside transporter subunit -!- periplasmic-binding component of ABC superfamily | Periplasmic | COG1879;ABC-type sugar transport system, periplasmic component | GO:0016052 carbohydrate catabolism -!- GO:0006457 protein | g | tt |
| *mglC* | CDS | methyl-galactoside transporter subunit -!- membrane component of ABC superfamily | Integral Membrane Protein | COG4211;ABC-type glucose/galactose transport system, permease component | GO:0016052 carbohydrate catabolism | o | tt |
| *mgsA* | CDS | methylglyoxal synthase | Cytoplasmic | COG1803;Methylglyoxal synthase | GO:0009438 methylglyoxal metabolism | | |
| *mgtA* | CDS | magnesium transporter | Integral Membrane Protein | COG0474;Cation transport ATPase | GO:0015986 ATP synthesis coupled proton transport | o | tt |
| *mhpA* | CDS | 3-(3-hydroxyphenyl)propionate hydroxylase | Cytoplasmic | COG0654;2-polyprenyl-6-methoxyphenol hydroxylase and related FAD-dependent oxidoreductases | GO:0019395 fatty acid oxidation | gg | t |
| *mhpB* | CDS | 2,3-dihydroxyphenylpropionate 1,2-dioxygenase | Cytoplasmic | | GO:0019395 fatty acid oxidation | gg | t |
| *mhpC* | CDS | 2-hydroxy-6-ketonona-2,4-dienedioic acid hydrolase | Cytoplasmic | | GO:0019395 fatty acid oxidation | gg | o |
| *mhpD* | CDS | 2-keto-4-pentenoate hydratase | Cytoplasmic | COG3971;2-keto-4-pentenoate hydratase | GO:0019395 fatty acid oxidation | gg | t |
| *mhpE* | CDS | 4-hyroxy-2-oxovalerate/4-hydroxy-2-oxopentanoic acid aldolase, class I | Cytoplasmic | COG4569;Acetaldehyde dehydrogenase (acetylating) | GO:0019395 fatty acid oxidation | gg | tt |
| *mhpF* | CDS | acetaldehyde-CoA dehydrogenase II, NAD-binding | Cytoplasmic | COG0119;Isopropylmalate/homocitrate/citramalate synthases | GO:0019395 fatty acid oxidation | gg | t |
| *mhpR* | CDS | DNA-binding transcriptional activator, 3HPP-binding | Cytoplasmic | COG1414;Transcriptional regulator | GO:0019395 fatty acid oxidation -!- GO:0006350 transcription | o | tt |
| *mhpT* | CDS | predicted 3-hydroxyphenylpropionic transporter | Integral Membrane Protein | COG0477;Permeases of the major facilitator superfamily | GO:0019395 fatty acid oxidation | | |
| *miaA* | CDS | delta(2)-isopentenylpyrophosphate tRNA-adenosine transferase | Cytoplasmic | COG0324;tRNA delta(2)-isopentenylpyrophosphate transferase | GO:0009451 RNA modification | o | tt |
| *miaB* | CDS | isopentenyl-adenosine A37 tRNA methylthiolase | Cytoplasmic | COG0621;2-methylthioadenine synthetase | GO:0009451 RNA modification | | |
| *micC* | misc_RNA | regulatory sRNA | | | | | |
| *micF* | misc_RNA | regulatory sRNA | | | | | |
| *minC* | CDS | cell division inhibitor | Cytoplasmic | COG0850;Septum formation inhibitor | | gg | tt |
| *minD* | CDS | membrane ATPase of the MinC-MinD-MinE system | Cytoplasmic | COG2894;Septum formation inhibitor-activating ATPase | | gg | tt |
| *minE* | CDS | cell division topological specificity factor | Cytoplasmic | COG0851;Septum formation topological specificity factor | | gg | t |
| *mioC* | CDS | FMN-binding protein MioC | Cytoplasmic | COG0716;Flavodoxins | GO:0006261 DNA dependent DNA replication | gg | t |
| *mipA* | CDS | scaffolding protein for murein synthesizing machinery | Outer Membrane B-barrel protein | COG3713;Outer membrane protein V | GO:0009252 peptidoglycan biosynthesis | o | tt |
| *mlrA* | CDS | DNA-binding transcriptional regulator | Cytoplasmic | COG0789;Predicted transcriptional regulators | GO:0006350 transcription | gg | o |
| *mltA* | CDS | membrane-bound lytic murein transglycosylase A | Outer Membrane Lipoprotein | COG2821;Membrane-bound lytic murein transglycosylase | GO:0009252 peptidoglycan biosynthesis | g | tt |
| *mltB* | CDS | membrane-bound lytic murein transglycosylase B | Outer Membrane Lipoprotein | COG2951;Membrane-bound lytic murein transglycosylase B | GO:0009252 peptidoglycan biosynthesis | o | tt |
| *mltC* | CDS | membrane-bound lytic murein transglycosylase C | Outer Membrane Lipoprotein | COG0741;Soluble lytic murein transglycosylase and related regulatory proteins (some contain LysM/invasin domains) | GO:0009252 peptidoglycan biosynthesis | | |
| *mltD* | CDS | predicted membrane-bound lytic murein transglycosylase D | Outer Membrane Lipoprotein | COG0741;Soluble lytic murein transglycosylase and related regulatory proteins (some contain LysM/invasin domains) | GO:0009061 anaerobic respiration -!- GO:0017004 cytochrome biogenesis -!- GO:0006350 transcription | | |
| *mmuM* | CDS | CP4-6 prophage; S-methylmethionine:homocysteine methyltransferase | Cytoplasmic | COG2040;Homocysteine/selenocysteine methylase (S-methylmethionine-dependent) | GO:0006071 glycerol metabolism -!- GO:0009086 methionine biosynthesis | o | tt |
| *mmuP* | CDS | CP4-6 prophage; predicted S-methylmethionine transporter | Integral Membrane Protein | | GO:0009086 methionine biosynthesis | | |
| *mngA* | CDS | fused 2-O-a-mannosyl-D-glycerate specific PTS enzyme IIABC components | Integral Membrane Protein | COG1762;Phosphotransferase system mannitol/fructose-specific IIA domain (Ntr-type) -!- COG1445;Phosphotransferase system fructose-specific component IIB -!- COG1299;Phosphotransferase system, fructose-specific IIC component | GO:0016052 carbohydrate catabolism -!- GO:0006464 protein modification -!- GO:0009266 response to temperature | o | tt |
| *mngB* | CDS | alpha-mannosidase | Cytoplasmic | COG0383;Alpha-mannosidase | GO:0016052 carbohydrate | o | tt |
| *mngR* | CDS | DNA-binding transcriptional dual regulator, fatty-acyl-binding | Cytoplasmic | COG2188;Transcriptional regulators | GO:0006099 tricarboxylic acid cycle -!- GO:0006350 transcription -!- GO:0006355 regulation of transcription, DNA-dependent | o | tt |
| *mntH* | CDS | manganese/divalent cation transporter | Integral Membrane Protein | COG1914;Mn2+ and Fe2+ transporters of the NRAMP family | GO:0006805 xenobiotic metabolism | gg | tt |
| *mntR* | CDS | transcriptional regulator of mntH | Cytoplasmic | COG1321;Mn-dependent transcriptional regulator | | o | tt |
| *moaA* | CDS | molybdopterin biosynthesis protein A | Cytoplasmic | COG2896;Molybdenum cofactor biosynthesis enzyme | GO:0006777 Mo-molybdopterin cofactor biosynthesis | gg | t |
| *moaB* | CDS | molybdopterin biosynthesis protein B | Cytoplasmic | COG0521;Molybdopterin biosynthesis enzymes | GO:0006777 Mo-molybdopterin cofactor biosynthesis | gg | t |
| *moaC* | CDS | molybdopterin biosynthesis, protein C | Cytoplasmic | COG0315;Molybdenum cofactor biosynthesis enzyme | GO:0006777 Mo-molybdopterin cofactor biosynthesis | gg | t |
| *moaD* | CDS | molybdopterin synthase, small subunit | Cytoplasmic | COG1977;Molybdopterin converting factor, small subunit | GO:0006777 Mo-molybdopterin cofactor biosynthesis | gg | t |
| *moaE* | CDS | molybdopterin synthase, large subunit | Cytoplasmic | COG0314;Molybdopterin converting factor, large subunit | GO:0006777 Mo-molybdopterin cofactor biosynthesis | gg | o |
| *mobA* | CDS | molybdopterin-guanine dinucleotide synthase | Cytoplasmic | COG0746;Molybdopterin-guanine dinucleotide biosynthesis protein A | GO:0006777 Mo-molybdopterin cofactor biosynthesis | gg | o |
| *mobB* | CDS | molybdopterin-guanine dinucleotide biosynthesis protein B | Cytoplasmic | COG1763;Molybdopterin-guanine dinucleotide biosynthesis protein | GO:0006777 Mo-molybdopterin cofactor biosynthesis | | |
| *modA* | CDS | molybdate transporter subunit -!- periplasmic-binding component of ABC superfamily | Periplasmic | COG0725;ABC-type molybdate transport system, periplasmic component | GO:0006777 Mo-molybdopterin cofactor biosynthesis | o | tt |
| *modB* | CDS | molybdate transporter subunit -!- membrane component of ABC superfamily | Integral Membrane Protein | COG4149;ABC-type molybdate transport system, permease component | | o | tt |
| *modC* | CDS | molybdate transporter subunit -!- ATP-binding component of ABC superfamily | Cytoplasmic | COG4148;ABC-type molybdate transport system, ATPase component | GO:0006777 Mo-molybdopterin cofactor biosynthesis | g | t |
| *modE* | CDS | DNA-binding transcriptional dual regulator | Cytoplasmic | COG2005;N-terminal domain of molybdenum-binding protein | GO:0006777 Mo-molybdopterin cofactor biosynthesis -!- GO:0006350 transcription | gg | o |
| *modF* | CDS | fused subunits of molybdate transporter -!- ATP-binding components of ABC superfamily | Cytoplasmic | COG1119;ABC-type molybdenum transport system, ATPase component/photorepair protein PhrA | GO:0006777 Mo-molybdopterin cofactor biosynthesis | gg | t |
| *moeA* | CDS | molybdopterin biosynthesis protein | Cytoplasmic | COG0303;Molybdopterin biosynthesis enzyme | GO:0006777 Mo-molybdopterin cofactor biosynthesis | gg | t |
| *moeB* | CDS | molybdopterin synthase sulfurylase | Cytoplasmic | COG0476;Dinucleotide-utilizing enzymes involved in molybdopterin and thiamine biosynthesis family 2 | GO:0006777 Mo-molybdopterin cofactor biosynthesis | g | tt |
| *mog* | CDS | predicted molybdochelatase | Cytoplasmic | COG0521;Molybdopterin biosynthesis enzymes | GO:0006777 Mo-molybdopterin cofactor biosynthesis | gg | tt |
| *mokB* | CDS | regulatory peptide | Cytoplasmic | | | | |
| *mokC* | CDS | regulatory protein for HokC, overlaps CDS of hokC | Periplasmic | | | | |
| *molR* | CDS | DNA-binding transcriptional regulator, N-ter fragment (pseudogene) | Cytoplasmic | | | o | o |
| *molR* | ancestral | DNA-binding transcriptional regulator (pseudogene) | Cytoplasmic | | | | |
| *molR* | CDS | DNA-binding transcriptional regulator, middle fragment (pseudogene) | Cytoplasmic | | | | |

| | | | | | | | |
|---|---|---|---|---|---|---|---|
| *molR* | CDS | DNA-binding transcriptional regulator, C-ter fragment (pseudogene) | Cytoplasmic | | | | |
| *motA* | CDS | proton conductor component of flagella motor | Integral Membrane Protein | COG1291;Flagellar motor component | GO:0042330 taxis | o | tt |
| *motB* | CDS | protein that enables flagellar motor rotation | Membrane Anchored | COG1360;Flagellar motor protein | GO:0042330 taxis | o | tt |
| *mpaA* | CDS | murein peptide amidase A | Cytoplasmic | COG2866;Predicted carboxypeptidase | | gg | t |
| *mpl* | CDS | UDP-N-acetylmuramate:L-alanyl-gamma-D-glutamyl-meso-diaminopimelate ligase | Periplasmic | COG0773;UDP-N-acetylmuramate-alanine ligase | GO:0000270 peptidoglycan metabolism | gg | t |
| *mppA* | CDS | murein tripeptide (L-ala-gamma-D-glutamyl-meso-DAP) transporter subunit -!- periplasmic-binding component of ABC superfamily | Periplasmic | COG4166;ABC-type oligopeptide transport system, periplasmic component | GO:0000270 peptidoglycan metabolism | gg | tt |
| *mprA* | CDS | DNA-binding transcriptional regulator | Cytoplasmic | COG1846;Transcriptional regulators | GO:0006350 transcription -!- GO:0042493 response to drug | o | tt |
| *mqo* | CDS | malate dehydrogenase, FAD/NAD(P)-binding domain | Periplasmic | COG0579;Predicted dehydrogenase | GO:0006099 tricarboxylic acid cycle | g | tt |
| *mraW* | CDS | S-adenosyl-dependent methyltransferase activity on membrane-located substrates | Cytoplasmic | COG0275;Predicted S-adenosylmethionine-dependent methyltransferase involved in cell envelope biogenesis | GO:0042158 lipoprotein biosynthesis | gg | tt |
| *mraY* | CDS | phospho-N-acetylmuramoyl-pentapeptide transferase | Integral Membrane Protein | COG0472;UDP-N-acetylmuramyl pentapeptide phosphotransferase/UDP-N-acetylglucosamine-1-phosp hate transferase | GO:0009252 peptidoglycan biosynthesis | o | tt |
| *mraZ* | CDS | conserved protein | Cytoplasmic | COG2001;Uncharacterized protein conserved in bacteria | | o | tt |
| *mrcA* | CDS | fused penicillin-binding protein 1a murein transglycosylase -!- murein transpeptidase | Membrane Anchored | COG5009;Membrane carboxypeptidase/penicillin-binding protein | GO:0009252 peptidoglycan biosynthesis -!- GO:0042493 response to drug | o | tt |
| *mrcB* | CDS | fused glycosyl transferase -!- transpeptidase | Membrane Anchored | COG0744;Membrane carboxypeptidase (penicillin-binding protein) | GO:0046349 amino sugar biosynthesis -!- GO:0009252 peptidoglycan biosynthesis -!- GO:0042493 response to drug | o | tt |
| *mrdA* | CDS | transpeptidase involved in peptidoglycan synthesis (penicillin-binding protein 2) | Membrane Anchored | COG0768;Cell division protein FtsI/penicillin-binding protein 2 | GO:0009252 peptidoglycan biosynthesis -!- GO:0042493 response to drug | o | tt |
| *mrdB* | CDS | cell wall shape-determining protein | Integral Membrane Protein | COG0772;Bacterial cell division membrane protein | | o | tt |
| *mreB* | CDS | cell wall structural complex MreBCD, actin-like component MreB | Cytoplasmic | COG1077;Actin-like ATPase involved in cell morphogenesis | GO:0042493 response to drug | o | tt |
| *mreC* | CDS | cell wall structural complex MreBCD transmembrane component MreC | Membrane Anchored | COG1792;Cell shape-determining protein | GO:0009252 peptidoglycan biosynthesis | o | tt |
| *mreD* | CDS | cell wall structural complex MreBCD transmembrane component MreD | Integral Membrane Protein | COG2891;Cell shape-determining protein | GO:0009252 peptidoglycan biosynthesis | o | tt |
| *mrp* | CDS | antiporter inner membrane protein | Cytoplasmic | | | gg | o |
| *mrr* | CDS | methylated adenine and cytosine restriction protein | Cytoplasmic | COG1715;Restriction endonuclease | GO:0006306 DNA methylation | o | tt |
| *msbA* | CDS | fused lipid transporter subunits -!- membrane component and ATP-binding component of ABC superfamily | Integral Membrane Protein | COG1132;ABC-type multidrug transport system, ATPase and permease components | | o | tt |
| *mscL* | CDS | mechanosensitive channel | Integral Membrane Protein | COG1970;Large-conductance mechanosensitive channel | | o | tt |
| *mscS* | CDS | mechanosensitive channel | Integral Membrane Protein | COG0668;Small-conductance mechanosensitive channel | | o | tt |
| *msrA* | CDS | methionine sulfoxide reductase A | Cytoplasmic | COG0225;Peptide methionine sulfoxide reductase | GO:0006457 protein folding | gg | o |
| *msyB* | CDS | predicted protein | Cytoplasmic | | | o | tt |
| *mtgA* | CDS | biosynthetic peptidoglycan transglycosylase | Membrane Anchored | COG0744;Membrane carboxypeptidase (penicillin-binding protein) | GO:0009252 peptidoglycan biosynthesis | o | tt |
| *mtlA* | CDS | fused mannitol-specific PTS enzyme IIABC components | Integral Membrane Protein | COG2213;Phosphotransferase system, mannitol-specific IIBC component -!- COG4668;Mannitol/fructose-specific phosphotransferase system, IIA | GO:0016052 carbohydrate catabolism | o | tt |
| *mtlD* | CDS | mannitol-1-phosphate dehydrogenase, NAD(P)-binding | Cytoplasmic | COG0246;Mannitol-1-phosphate/altronate dehydrogenases | GO:0016052 carbohydrate catabolism | gg | t |
| *mtlR* | CDS | DNA-binding transcriptional repressor | Cytoplasmic | COG3722;Transcriptional regulator | GO:0016052 carbohydrate catabolism -!- GO:0006350 | gg | o |
| *mtr* | CDS | tryptophan transporter of high affinity | Integral Membrane Protein | COG0814;Amino acid permeases | GO:0000162 tryptophan biosynthesis | o | tt |
| *mukB* | CDS | fused chromosome partitioning proteins | Cytoplasmic | COG3096;Uncharacterized protein involved in chromosome partitioning | | o | tt |
| *mukE* | CDS | protein involved in chromosome partitioning | Cytoplasmic | COG3095;Uncharacterized protein involved in chromosome partitioning | | gg | t |
| *mukF* | CDS | Involved in chromosome partioning, Ca2+ binding protein | Cytoplasmic | COG3006;Uncharacterized protein involved in chromosome partitioning | | gg | tt |
| *murA* | CDS | UDP-N-acetylglucosamine 1-carboxyvinyltransferase | Cytoplasmic | COG0766;UDP-N-acetylglucosamine enolpyruvyl transferase | GO:0009252 peptidoglycan biosynthesis | o | tt |
| *murB* | CDS | UDP-N-acetylenolpyruvoylglucosamine reductase, FAD-binding | Cytoplasmic | COG0812;UDP-N-acetylmuramate dehydrogenase | GO:0009252 peptidoglycan biosynthesis | gg | t |
| *murC* | CDS | UDP-N-acetylmuramate:L-alanine ligase | Cytoplasmic | COG0773;UDP-N-acetylmuramate-alanine ligase | GO:0009252 peptidoglycan biosynthesis | g | o |
| *murD* | CDS | UDP-N-acetylmuramoyl-L-alanine:D-glutamate ligase | Cytoplasmic | COG0771;UDP-N-acetylmuramoylalanine-D-glutamate ligase | GO:0009252 peptidoglycan biosynthesis | gg | o |
| *murE* | CDS | UDP-N-acetylmuramoyl-L-alanyl-D-glutamate:meso-diaminopimelate ligase | Cytoplasmic | COG0769;UDP-N-acetylmuramyl tripeptide synthase | GO:0009252 peptidoglycan biosynthesis | gg | t |
| *murF* | CDS | UDP-N-acetylmuramoyl-tripeptide:D-alanyl-D-alanine ligase | Cytoplasmic | COG0770;UDP-N-acetylmuramyl pentapeptide synthase | GO:0009252 peptidoglycan biosynthesis -!- GO:0000270 peptidoglycan metabolism | gg | t |
| *murG* | CDS | N-acetylglucosaminyl transferase | Membrane Associated | COG0707;UDP-N-acetylglucosamine:LPS N-acetylglucosamine transferase | GO:0009252 peptidoglycan biosynthesis | o | tt |
| *murI* | CDS | glutamate racemase | Cytoplasmic | COG0796;Glutamate racemase | GO:0009252 peptidoglycan biosynthesis | | |
| *murP* | CDS | fused predicted enzyme IIBC components of PTS | Integral Membrane Protein | COG1264;Phosphotransferase system IIB components -!- COG1263;Phosphotransferase system IIC components, glucose/maltose/N-acetylglucosamine-specific | GO:0016052 carbohydrate catabolism | o | tt |
| *mutH* | CDS | methyl-directed mismatch repair protein | Cytoplasmic | COG3066;DNA mismatch repair protein | GO:0006281 DNA repair | o | o |
| *mutL* | CDS | methyl-directed mismatch repair protein | Cytoplasmic | COG0323;DNA mismatch repair enzyme (predicted ATPase) | GO:0006281 DNA repair | o | o |
| *mutM* | CDS | formamidopyrimidine/5-formyluracil/ 5-hydroxymethyluracil DNA glycosylase | Cytoplasmic | COG0266;Formamidopyrimidine-DNA glycosylase | GO:0006281 DNA repair | o | tt |
| *mutS* | CDS | methyl-directed mismatch repair protein | Cytoplasmic | COG0249;Mismatch repair ATPase (MutS family) | GO:0006281 DNA repair | o | tt |
| *mutT* | CDS | nucleoside triphosphate pyrophosphohydrolase, marked preference for dGTP | Cytoplasmic | COG0494;NTP pyrophosphohydrolases including oxidative damage repair enzymes | GO:0009265 2'-deoxyribonucleotide biosynthesis | gg | o |
| *mutY* | CDS | adenine DNA glycosylase | Cytoplasmic | COG1194;A/G-specific DNA glycosylase | GO:0006281 DNA repair | g | tt |
| *mviM* | CDS | predicted oxidoreductase, NAD(P)-binding Rossmann-fold domain | Periplasmic | COG0673;Predicted dehydrogenases and related proteins | | gg | t |
| *mviN* | CDS | predicted inner membrane protein | Integral Membrane Protein | COG0728;Uncharacterized membrane protein, putative virulence factor | | o | tt |
| *nac* | CDS | DNA-binding transcriptional dual regulator | Cytoplasmic | COG0583;Transcriptional regulator | GO:0006807 nitrogen metabolism -!- GO:0006350 transcription | o | tt |
| *nadA* | CDS | quinolinate synthase, subunit A | Cytoplasmic | COG0379;Quinolinate synthase | GO:0009435 nicotinamide adenine dinucleotide biosynthesis | gg | t |
| *nadB* | CDS | quinolinate synthase, L-aspartate oxidase (B protein) subunit | Cytoplasmic | COG0029;Aspartate oxidase | GO:0009435 nicotinamide adenine dinucleotide biosynthesis | gg | t |
| *nadC* | CDS | quinolinate phosphoribosyltransferase | Cytoplasmic | COG0157;Nicotinate-nucleotide pyrophosphorylase | GO:0009435 nicotinamide adenine dinucleotide biosynthesis | gg | o |
| *nadD* | CDS | nicotinic acid mononucleotide adenylyltransferase, NAD(P)-dependent | Cytoplasmic | COG1057;Nicotinic acid mononucleotide adenylyltransferase | GO:0009435 nicotinamide adenine dinucleotide biosynthesis | gg | o |
| *nadE* | CDS | NAD synthetase, NH3/glutamine-dependent | Cytoplasmic | COG0171;NAD synthase | GO:0009435 nicotinamide adenine dinucleotide biosynthesis | gg | o |
| *nadR* | CDS | bifunctional DNA-binding transcriptional repressor -!- NMN adenylyltransferase | Cytoplasmic | COG1396;Predicted transcriptional regulators -!- COG1056;Nicotinamide mononucleotide adenylyltransferase -!- COG3172;Predicted ATPase/kinase involved in NAD metabolism | GO:0009435 nicotinamide adenine dinucleotide biosynthesis -!- GO:0006350 transcription | | |
| *nagA* | CDS | N-acetylglucosamine-6-phosphate deacetylase | Cytoplasmic | COG1820;N-acetylglucosamine-6-phosphate deacetylase | GO:0046349 amino sugar biosynthesis | gg | t |
| *nagB* | CDS | glucosamine-6-phosphate deaminase | Cytoplasmic | COG0363;6-phosphogluconolactonase/Glucosamine-6-ph osphate isomerase/deaminase | GO:0046349 amino sugar biosynthesis | gg | o |
| *nagC* | CDS | DNA-binding transcriptional dual regulator | Cytoplasmic | COG1940;Transcriptional regulator/sugar kinase | GO:0046349 amino sugar biosynthesis -!- GO:0006350 transcription -!- GO:0009386 translational attenuation | gg | tt |
| *nagD* | CDS | UMP phosphatase | Cytoplasmic | COG0647;Predicted sugar phosphatases of the HAD superfamily | GO:0046349 amino sugar biosynthesis | gg | t |
| *nagE* | CDS | fused N-acetyl glucosamine specific PTS enzyme IICBA components | Integral Membrane Protein | COG1263;Phosphotransferase system IIC components, glucose/maltose/N-acetylglucosamine-specific -!- COG1264;Phosphotransferase system IIB components -!- | GO:0009310 amine catabolism -!- GO:0046349 amino sugar biosynthesis | o | tt |
| *nagZ* | CDS | beta N-acetyl-glucosaminidase | Cytoplasmic | COG1472;Beta-glucosidase-related glycosidases | GO:0000270 peptidoglycan metabolism | gg | t |
| *nanA* | CDS | N-acetylneuraminate lyase | Cytoplasmic | COG0329;Dihydrodipicolinate synthase/N-acetylneuraminate lyase | GO:0046349 amino sugar biosynthesis | g | t |
| *nanE* | CDS | predicted N-acetylmannosamine-6-P epimerase | Cytoplasmic | COG3010;Putative N-acetylmannosamine-6-phosphate epimerase | GO:0046349 amino sugar biosynthesis | gg | o |
| *nanK* | CDS | predicted N-acetylmannosamine kinase | Cytoplasmic | | GO:0046349 amino sugar biosynthesis | | |
| *nanR* | CDS | DNA-binding transcriptional dual regulator | Cytoplasmic | | GO:0006350 transcription | o | tt |
| *nanT* | CDS | sialic acid transporter | Integral Membrane Protein | | GO:0046349 amino sugar biosynthesis | o | tt |
| *napA* | CDS | nitrate reductase, periplasmic, large subunit | Periplasmic | COG0243;Anaerobic dehydrogenases, typically selenocysteine-containing | GO:0009061 anaerobic respiration | gg | tt |
| *napB* | CDS | nitrate reductase, small, cytochrome C550 subunit, periplasmic | Periplasmic | | GO:0009061 anaerobic respiration -!- GO:0017004 cytochrome biogenesis | | |
| *napC* | CDS | nitrate reductase, cytochrome c-type,periplasmic | Periplasmic with N-terminal Membrane | COG3005;Nitrate/TMAO reductases, membrane-bound tetraheme cytochrome c subunit | GO:0009061 anaerobic respiration -!- GO:0017004 cytochrome biogenesis | o | tt |
| *napD* | CDS | assembly protein for periplasmic nitrate reductase | Cytoplasmic | COG3062;Uncharacterized protein involved in formation of periplasmic nitrate reductase | GO:0009061 anaerobic respiration -!- GO:0006464 protein modification | gg | t |
| *napF* | CDS | ferredoxin-type protein, predicted role in electron transfer to periplasmic nitrate reductase (NapA) | Cytoplasmic | COG1145;Ferredoxin | GO:0009061 anaerobic respiration | o | o |
| *napG* | CDS | ferredoxin-type protein essential for electron transfer from ubiquinol to periplasmic nitrate reductase (NapAB) | Periplasmic | COG1145;Ferredoxin | GO:0009061 anaerobic respiration | g | t |
| *napH* | CDS | ferredoxin-type protein essential for electron transfer from ubiquinol to periplasmic nitrate reductase (NapAB) | Integral Membrane Protein | COG0348;Polyferredoxin | GO:0009061 anaerobic respiration | o | tt |

| | | | | | | | |
|---|---|---|---|---|---|---|---|
| *narG* | CDS | nitrate reductase 1, alpha subunit | Cytoplasmic | COG5013;Nitrate reductase alpha subunit | GO:0009061 anaerobic respiration | o | tt |
| *narH* | CDS | nitrate reductase 1, beta (Fe-S) subunit | Cytoplasmic | COG1140;Nitrate reductase beta subunit | GO:0009061 anaerobic respiration | gg | t |
| *narI* | CDS | nitrate reductase 1, gamma (cytochrome b(NR)) subunit | Integral Membrane Protein | COG2181;Nitrate reductase gamma subunit | GO:0009061 anaerobic respiration -!- GO:0017004 cytochrome biogenesis | o | tt |
| *narJ* | CDS | molybdenum-cofactor-assembly chaperone subunit (delta subunit) of nitrate reductase 1 | Cytoplasmic | COG2180;Nitrate reductase delta subunit | GO:0009061 anaerobic respiration -!- GO:0006457 protein folding | gg | t |
| *narK* | CDS | nitrate/nitrite transporter | Integral Membrane Protein | COG2223;Nitrate/nitrite transporter | GO:0006807 nitrogen metabolism | o | tt |
| *narL* | CDS | DNA-binding response regulator in two-component regulatory system with NarX (or NarQ) | Cytoplasmic | COG2197;Response regulator containing a CheY-like receiver domain and an HTH DNA-binding domain | GO:0009061 anaerobic respiration -!- GO:0006350 transcription | gg | t |
| *narP* | CDS | DNA-binding response regulator in two-component regulatory system with NarQ or NarX | Cytoplasmic | COG2197;Response regulator containing a CheY-like receiver domain and an HTH DNA-binding domain | GO:0009061 anaerobic respiration -!- GO:0006350 transcription | o | tt |
| *narQ* | CDS | sensory histidine kinase in two-component regulatory system with NarP (NarL) | Integral Membrane Protein | COG3850;Signal transduction histidine kinase, nitrate/nitrite-specific | GO:0009061 anaerobic respiration -!- GO:0006464 protein modification | o | tt |
| *narU* | CDS | nitrate/nitrite transporter | Integral Membrane Protein | COG2223;Nitrate/nitrite transporter | GO:0006807 nitrogen metabolism | o | tt |
| *narV* | CDS | nitrate reductase 2 (NRZ), gamma subunit | Integral Membrane Protein | COG2181;Nitrate reductase gamma subunit | GO:0009061 anaerobic respiration | o | tt |
| *narW* | CDS | nitrate reductase 2 (NRZ), delta subunit (assembly subunit) | Cytoplasmic | COG2180;Nitrate reductase delta subunit | GO:0009061 anaerobic respiration -!- GO:0006457 protein folding | gg | tt |
| *narX* | CDS | sensory histidine kinase in two-component regulatory system with NarL | Integral Membrane Protein | COG3850;Signal transduction histidine kinase, nitrate/nitrite-specific | GO:0009061 anaerobic respiration -!- GO:0006464 protein modification | o | tt |
| *narY* | CDS | nitrate reductase 2 (NRZ), beta subunit | Cytoplasmic | COG1140;Nitrate reductase beta subunit | GO:0009061 anaerobic respiration | gg | t |
| *narZ* | CDS | nitrate reductase 2 (NRZ), alpha subunit | Cytoplasmic | COG5013;Nitrate reductase alpha subunit | GO:0009061 anaerobic respiration | o | tt |
| *ndh* | CDS | respiratory NADH dehydrogenase 2/cupric reductase | Cytoplasmic | COG1252;NADH dehydrogenase, FAD-containing subunit | GO:0009060 aerobic respiration -!- GO:0009061 anaerobic respiration | gg | tt |
| *ndk* | CDS | multifunctional nucleoside diphosphate kinase -!- apyrimidinic endonuclease -!- 3'-phosphodiesterase | Cytoplasmic | COG0105;Nucleoside diphosphate kinase | GO:0015949 nucleobase, nucleoside and nucleotide interconversion -!- GO:0009152 purine ribonucleotide biosynthesis | gg | t |
| *nei* | CDS | endonuclease VIII -!- 5-formyluracil/5-hydroxymethyluracil DNA glycosylase | Cytoplasmic | COG0266;Formamidopyrimidine-DNA glycosylase | GO:0006281 DNA repair | o | tt |
| *nemA* | CDS | N-ethylmaleimide reductase, FMN-linked | Cytoplasmic | COG1902;NADH:flavin oxidoreductases, Old Yellow Enzyme family | GO:0006805 xenobiotic metabolism | gg | t |
| *nfi* | CDS | endonuclease V | Cytoplasmic | COG1515;Deoxyinosine 3'endonuclease (endonuclease V) | GO:0006308 DNA catabolism -!- GO:0006308 DNA catabolism | | |
| *nfnB* | CDS | dihydropteridine reductase, NAD(P)H-dependent, oxygen-insensitive | Cytoplasmic | COG0778;Nitroreductase | GO:0042493 response to drug | gg | t |
| *nfo* | CDS | endonuclease IV with intrinsic 3'-5' exonuclease activity | Periplasmic | COG0648;Endonuclease IV | GO:0006308 DNA catabolism -!- GO:0006308 DNA catabolism -!- GO:0009314 response to radiation | gg | t |
| *nfrA* | CDS | bacteriophage N4 receptor, outer membrane subunit | Outer Membrane B-barrel protein | COG0457;FOG: TPR repeat | | o | tt |
| *nfrB* | CDS | bacteriophage N4 receptor, inner membrane subunit | Integral Membrane Protein | | | o | tt |
| *nfsA* | CDS | nitroreductase A, NADPH-dependent, FMN- | Cytoplasmic | COG0778;Nitroreductase | GO:0009061 anaerobic respiration | gg | t |
| *nhaA* | CDS | sodium-proton antiporter | Integral Membrane Protein | COG3004;Na+/H+ antiporter | GO:0009268 response to pH | o | tt |
| *nhaB* | CDS | sodium:proton antiporter | Integral Membrane Protein | COG3067;Na+/H+ antiporter | GO:0009268 response to pH | o | tt |
| *nhaR* | CDS | DNA-binding transcriptional activator | Cytoplasmic | COG0583;Transcriptional regulator | GO:0006350 transcription -!- GO:0009268 response to pH | o | tt |
| *nhoA* | CDS | N-hydroxyarylamine O-acetyltransferase | Cytoplasmic | COG2162;Arylamine N-acetyltransferase | GO:0008152 metabolism | gg | o |
| *nikA* | CDS | nickel transporter subunit -!- periplasmic-binding component of ABC superfamily | Periplasmic | COG0747;ABC-type dipeptide transport system, periplasmic component | GO:0042330 taxis | o | tt |
| *nikB* | CDS | nickel transporter subunit -!- membrane component of ABC superfamily | Integral Membrane Protein | COG0601;ABC-type dipeptide/oligopeptide/nickel transport systems, permease components | | o | tt |
| *nikC* | CDS | nickel transporter subunit -!- membrane component of ABC superfamily | Integral Membrane Protein | COG1173;ABC-type dipeptide/oligopeptide/nickel transport systems, permease components | | o | tt |
| *nikD* | CDS | nickel transporter subunit -!- ATP-binding component of ABC superfamily | Cytoplasmic | COG0444;ABC-type dipeptide/oligopeptide/nickel transport system, ATPase component | GO:0009060 aerobic respiration -!- GO:0009061 anaerobic respiration | gg | t |
| *nikE* | CDS | nickel transporter subunit -!- ATP-binding component of ABC superfamily | Cytoplasmic | COG1124;ABC-type dipeptide/oligopeptide/nickel transport system, ATPase component | GO:0009060 aerobic respiration -!- GO:0009061 anaerobic respiration | gg | tt |
| *nikR* | CDS | DNA-binding transcriptional regulator, Ni-binding | Cytoplasmic | COG0864;Predicted transcriptional regulators containing the CopG/Arc/MetJ DNA-binding domain and a metal-binding domain | GO:0006350 transcription | g | o |
| *ninE* | CDS | DLP12 prophage; conserved protein | Cytoplasmic | | | gg | tt |
| *nirB* | CDS | nitrite reductase, large subunit, NAD(P)H-binding | Cytoplasmic | COG1251;NAD(P)H-nitrite reductase | GO:0009061 anaerobic respiration | o | tt |
| *nirC* | CDS | nitrite transporter | Integral Membrane Protein | | GO:0006807 nitrogen metabolism | o | tt |
| *nirD* | CDS | nitrite reductase, NAD(P)H-binding, small subunit | Cytoplasmic | COG2146;Ferredoxin subunits of nitrite reductase and ring-hydroxylating dioxygenases | GO:0009061 anaerobic respiration | gg | tt |
| *nlpA* | CDS | cytoplasmic membrane lipoprotein-28 | Inner Membrane Lipoprotein | COG1464;ABC-type metal ion transport system, periplasmic component/surface antigen | | o | tt |
| *nlpB* | CDS | lipoprotein | Outer Membrane Lipoprotein | | | o | tt |
| *nlpC* | CDS | predicted lipoprotein | Outer Membrane Lipoprotein | COG0791;Cell wall-associated hydrolases (invasion-associated proteins) | | o | tt |
| *nlpD* | CDS | predicted outer membrane lipoprotein | Outer Membrane Lipoprotein | COG0739;Membrane proteins related to metalloendopeptidases | | o | tt |
| *nlpE* | CDS | lipoprotein involved with copper homeostasis and adhesion | Outer Membrane Lipoprotein | COG3015;Uncharacterized lipoprotein NlpE involved in copper resistance | GO:0006805 xenobiotic metabolism | o | tt |
| *nlpI* | CDS | conserved protein | Outer Membrane Lipoprotein | COG4785;Lipoprotein NlpI, contains TPR repeats | | o | tt |
| *nmpC* | CDS | DLP12 prophage; truncated outer membrane porin (pseudogene) | Outer Membrane B-barrel protein | COG3203;Outer membrane protein (porin) | | | |
| *nohA* | CDS | Qin prophage; predicted packaging protein | Cytoplasmic | COG4220;Phage DNA packaging protein, Nu1 subunit of terminase | | gg | o |
| *nohB* | CDS | DLP12 prophage; DNA packaging protein | Cytoplasmic | COG4220;Phage DNA packaging protein, Nu1 subunit of terminase | | gg | t |
| *norR* | CDS | DNA-binding transcriptional activator | Cytoplasmic | COG3604;Transcriptional regulator containing GAF, AAA-type ATPase, and DNA binding domains | GO:0006350 transcription | | |
| *norV* | CDS | flavorubredoxin oxidoreductase | Cytoplasmic | COG0426;Uncharacterized flavoproteins -!- COG1773;Rubredoxin | | gg | t |
| *norW* | CDS | NADH:flavorubredoxin oxidoreductase | Cytoplasmic | COG0446;Uncharacterized NAD(FAD)-dependent dehydrogenases | | gg | t |
| *npr* | CDS | phosphohistidinoprotein-hexose phosphotransferase component of N-regulated PTS system (Npr) | Cytoplasmic | COG1925;Phosphotransferase system, HPr-related proteins | GO:0006464 protein modification | g | o |
| *nrdA* | CDS | ribonucleoside diphosphate reductase 1, alpha subunit | Cytoplasmic | COG0209;Ribonucleotide reductase, alpha subunit | GO:0015949 nucleobase, nucleoside and nucleotide interconversion | gg | t |
| *nrdB* | CDS | ribonucleoside diphosphate reductase 1, beta subunit, ferritin-like | Cytoplasmic | COG0208;Ribonucleotide reductase, beta subunit | GO:0015949 nucleobase, nucleoside and nucleotide interconversion | gg | t |
| *nrdD* | CDS | anaerobic ribonucleoside-triphosphate reductase | Cytoplasmic | COG1328;Oxygen-sensitive ribonucleoside-triphosphate reductase | GO:0009265 2'-deoxyribonucleotide biosynthesis | gg | t |
| *nrdE* | CDS | ribonucleoside-diphosphate reductase 2, alpha subunit | Cytoplasmic | COG0209;Ribonucleotide reductase, alpha subunit | GO:0015949 nucleobase, nucleoside and nucleotide interconversion | | |
| *nrdF* | CDS | ribonucleoside-diphosphate reductase 2, beta subunit, ferritin-like | Cytoplasmic | COG0208;Ribonucleotide reductase, beta subunit | GO:0015949 nucleobase, nucleoside and nucleotide interconversion | gg | t |
| *nrdG* | CDS | anaerobic ribonucleotide reductase activating protein | Cytoplasmic | COG0602;Organic radical activating enzymes | GO:0015949 nucleobase, nucleoside and nucleotide interconversion -!- GO:0006464 protein modification | g | o |
| *nrdH* | CDS | glutaredoxin-like protein | Cytoplasmic | COG0695;Glutaredoxin and related proteins | | | |
| *nrdI* | CDS | protein that stimulates ribonucleotide reduction | Cytoplasmic | COG1780;Protein involved in ribonucleotide reduction | | gg | o |
| *nrfA* | CDS | nitrite reductase, formate-dependent, cytochrome | Periplasmic | COG3303;Formate-dependent nitrite reductase, periplasmic cytochrome c552 subunit | GO:0009061 anaerobic respiration -!- GO:0017004 cytochrome biogenesis | o | tt |
| *nrfB* | CDS | nitrite reductase, formate-dependent, penta-heme cytochrome c | Periplasmic | | GO:0009061 anaerobic respiration -!- GO:0017004 cytochrome biogenesis | o | tt |
| *nrfC* | CDS | formate-dependent nitrite reductase, 4Fe4S subunit | Periplasmic | COG0437;Fe-S-cluster-containing hydrogenase components 1 | GO:0009061 anaerobic respiration | gg | t |
| *nrfD* | CDS | formate-dependent nitrite reductase, membrane subunit | Integral Membrane Protein | COG3301;Formate-dependent nitrite reductase, membrane component | GO:0009061 anaerobic respiration | o | tt |
| *nrfE* | CDS | heme lyase (NrfEFG) for insertion of heme into c552, subunit NrfE | Integral Membrane Protein | COG1138;Cytochrome c biogenesis factor | GO:0009061 anaerobic respiration -!- GO:0017004 cytochrome biogenesis -!- GO:0006457 protein folding | o | tt |
| *nrfF* | CDS | heme lyase (NrfEFG) for insertion of heme into c552, subunit NrfF | Membrane Anchored | COG3088;Uncharacterized protein involved in biosynthesis of c-type cytochromes | GO:0009061 anaerobic respiration -!- GO:0017004 cytochrome biogenesis -!- GO:0006457 protein folding | o | tt |
| *nrfG* | CDS | heme lyase (NrfEFG) for insertion of heme into c552, subunit NrfG | Outer Membrane Lipoprotein | COG4235;Cytochrome c biogenesis factor | GO:0009061 anaerobic respiration -!- GO:0017004 cytochrome biogenesis -!- GO:0006457 protein folding | o | tt |
| *nth* | CDS | DNA glycosylase and apyrimidinic (AP) lyase (endonuclease III) | Cytoplasmic | COG0177;Predicted EndoIII-related endonuclease | GO:0006308 DNA catabolism -!- GO:0006308 DNA catabolism -!- GO:0009314 response to radiation | gg | t |
| *nudB* | CDS | dATP pyrophosphohydrolase | Cytoplasmic | COG0494;NTP pyrophosphohydrolases including oxidative damage repair enzymes | GO:0015949 nucleobase, nucleoside and nucleotide interconversion | o | o |
| *nudC* | CDS | NADH pyrophosphatase | Cytoplasmic | | | | |
| *nudD* | CDS | GDP-mannose mannosyl hydrolase | Cytoplasmic | COG0494;NTP pyrophosphohydrolases including oxidative damage repair enzymes | GO:0009226 nucleotide-sugar biosynthesis -!- GO:0009242 colanic acid biosynthesis | | |
| *nudE* | CDS | ADP-ribose diphosphatase | Cytoplasmic | COG0494;NTP pyrophosphohydrolases including oxidative damage repair enzymes | | g | t |
| *nudF* | CDS | ADP-ribose pyrophosphatase | Cytoplasmic | COG0494;NTP pyrophosphohydrolases including oxidative damage repair enzymes | | gg | tt |
| *nudG* | CDS | pyrimidine (deoxy)nucleoside triphosphate pyrophosphohydrolase | Cytoplasmic | COG0494;NTP pyrophosphohydrolases including oxidative damage repair enzymes | | gg | tt |
| *nudH* | CDS | nucleotide hydrolase | Cytoplasmic | COG0494;NTP pyrophosphohydrolases including oxidative damage repair enzymes | | g | tt |
| *nuoA* | CDS | NADH:ubiquinone oxidoreductase, membrane subunit A | Integral Membrane Protein | | GO:0009060 aerobic respiration -!- GO:0009061 anaerobic respiration | o | tt |
| *nuoB* | CDS | NADH:ubiquinone oxidoreductase, chain B | Cytoplasmic | COG0377;NADH:ubiquinone oxidoreductase 20 kD subunit and related Fe-S oxidoreductases | GO:0009060 aerobic respiration -!- GO:0009061 anaerobic respiration | | |
| *nuoC* | CDS | NADH:ubiquinone oxidoreductase, chain C,D | Cytoplasmic | COG0852;NADH:ubiquinone oxidoreductase 27 kD subunit -!- COG0649;NADH:ubiquinone oxidoreductase 49 kD subunit 7 | GO:0009060 aerobic respiration -!- GO:0009061 anaerobic respiration | | |
| *nuoE* | CDS | NADH:ubiquinone oxidoreductase, chain E | Cytoplasmic | COG1905;NADH:ubiquinone oxidoreductase 24 kD subunit | GO:0009060 aerobic respiration -!- GO:0009061 anaerobic respiration | gg | o |

| | | | | | | | |
|---|---|---|---|---|---|---|---|
| *nuoF* | CDS | NADH:ubiquinone oxidoreductase, chain F | Cytoplasmic | COG1894;NADH:ubiquinone oxidoreductase, NADH-binding (51 kD) subunit | GO:0009060 aerobic respiration -!- GO:0009061 anaerobic respiration | gg | t |
| *nuoG* | CDS | NADH:ubiquinone oxidoreductase, chain G | Cytoplasmic | | GO:0009060 aerobic respiration -!- GO:0009061 anaerobic respiration | o | tt |
| *nuoH* | CDS | NADH:ubiquinone oxidoreductase, membrane subunit H | Integral Membrane Protein | COG1005;NADH:ubiquinone oxidoreductase subunit 1 (chain H) | GO:0009060 aerobic respiration -!- GO:0009061 anaerobic respiration | o | tt |
| *nuoI* | CDS | NADH:ubiquinone oxidoreductase, chain I | Cytoplasmic | COG1143;Formate hydrogenlyase subunit 6/NADH:ubiquinone oxidoreductase 23 kD subunit (chain I) | GO:0009060 aerobic respiration -!- GO:0009061 anaerobic respiration | o | t |
| *nuoJ* | CDS | NADH:ubiquinone oxidoreductase, membrane subunit J | Integral Membrane Protein | COG0839;NADH:ubiquinone oxidoreductase subunit 6 (chain J) | GO:0009060 aerobic respiration -!- GO:0009061 anaerobic respiration | gg | tt |
| *nuoK* | CDS | NADH:ubiquinone oxidoreductase, membrane subunit K | Integral Membrane Protein | COG0713;NADH:ubiquinone oxidoreductase subunit 11 or 4L (chain K) | GO:0009060 aerobic respiration -!- GO:0009061 anaerobic respiration | o | tt |
| *nuoL* | CDS | NADH:ubiquinone oxidoreductase, membrane subunit L | Integral Membrane Protein | COG1009;NADH:ubiquinone oxidoreductase subunit 5 (chain L)/Multisubunit Na+/H+ antiporter, MnhA subunit | GO:0009060 aerobic respiration -!- GO:0009061 anaerobic respiration | o | tt |
| *nuoM* | CDS | NADH:ubiquinone oxidoreductase, membrane subunit M | Integral Membrane Protein | COG1008;NADH:ubiquinone oxidoreductase subunit 4 (chain M) | GO:0009060 aerobic respiration -!- GO:0009061 anaerobic respiration | o | tt |
| *nuoN* | CDS | NADH:ubiquinone oxidoreductase, membrane subunit N | Integral Membrane Protein | COG1007;NADH:ubiquinone oxidoreductase subunit 2 (chain N) | GO:0009060 aerobic respiration -!- GO:0009061 anaerobic respiration | o | tt |
| *nupC* | CDS | nucleoside (except guanosine) transporter | Integral Membrane Protein | COG1972;Nucleoside permease | GO:0015949 nucleobase, nucleoside and nucleotide interconversion | g | tt |
| *nupG* | CDS | nucleoside transporter | Integral Membrane Protein | | GO:0015949 nucleobase, nucleoside and nucleotide interconversion | g | tt |
| *nusA* | CDS | transcription termination/antitermination L factor | Cytoplasmic | COG0195;Transcription elongation factor | GO:0006350 transcription -!- GO:0006355 regulation of transcription, DNA-dependent | gg | t |
| *nusB* | CDS | transcription antitermination protein | Cytoplasmic | COG0781;Transcription termination factor | GO:0006350 transcription | gg | o |
| *nusG* | CDS | transcription termination factor | Cytoplasmic | COG0250;Transcription antiterminator | GO:0006350 transcription | o | tt |
| *obgE* | CDS | GTPase involved in cell partioning and DNA repair | Cytoplasmic | COG0536;Predicted GTPase | | gg | tt |
| *ogrK* | CDS | DNA-binding transcriptional regulator -!-prophage P2 remnant | Cytoplasmic | | | gg | o |
| *ogt* | CDS | O-6-alkylguanine-DNA:cysteine-protein methyltransferase | Cytoplasmic | COG0350;Methylated DNA-protein cysteine methyltransferase | GO:0006306 DNA methylation | gg | t |
| *ompA* | CDS | outer membrane protein A (3a;II*;G;d) | Outer Membrane B-barrel protein | COG2885;Outer membrane protein and related peptidoglycan-associated (lipo)proteins | | o | o |
| *ompC* | CDS | outer membrane porin protein C | Outer Membrane B-barrel protein | COG3203;Outer membrane protein (porin) | | o | tt |
| *ompF* | CDS | outer membrane porin 1a (Ia;b;F) | Outer Membrane B-barrel protein | COG3203;Outer membrane protein (porin) | | o | tt |
| *ompG* | CDS | outer membrane porin | Outer Membrane B-barrel protein | | | o | tt |
| *ompL* | CDS | predicted outer membrane porin L | Outer Membrane B-barrel protein | | | g | o |
| *ompN* | CDS | outer membrane pore protein N, non-specific | Outer Membrane B-barrel protein | COG3203;Outer membrane protein (porin) | | o | tt |
| *ompR* | CDS | DNA-binding response regulator in two-component regulatory system with EnvZ | Cytoplasmic | COG0745;Response regulators consisting of a CheY-like receiver domain and a winged-helix DNA-binding domain | GO:0006350 transcription | o | tt |
| *ompT* | CDS | DLP12 prophage; outer membrane protease VII (outer membrane protein 3b) | Outer Membrane B-barrel protein | COG4571;Outer membrane protease | | o | tt |
| *ompW* | CDS | outer membrane protein W | Outer Membrane B-barrel protein | COG3047;Outer membrane protein W | | o | tt |
| *ompX* | CDS | outer membrane protein | Outer Membrane B-barrel protein | COG3637;Opacity protein and related surface antigens | | o | tt |
| *oppA* | CDS | oligopeptide transporter subunit -!- periplasmic-binding component of ABC superfamily | Periplasmic | COG4166;ABC-type oligopeptide transport system, periplasmic component | GO:0000270 peptidoglycan metabolism -!- GO:0006457 protein folding | o | tt |
| *oppB* | CDS | oligopeptide transporter subunit -!- membrane component of ABC superfamily | Integral Membrane Protein | COG0601;ABC-type dipeptide/oligopeptide/nickel transport systems, permease components | GO:0000270 peptidoglycan metabolism | o | tt |
| *oppC* | CDS | oligopeptide transporter subunit -!- membrane component of ABC superfamily | Integral Membrane Protein | COG1173;ABC-type dipeptide/oligopeptide/nickel transport systems, permease components | GO:0000270 peptidoglycan metabolism | o | tt |
| *oppD* | CDS | oligopeptide transporter subunit -!- ATP-binding component of ABC superfamily | Cytoplasmic | | GO:0000270 peptidoglycan metabolism | g | o |
| *oppF* | CDS | oligopeptide transporter subunit -!- ATP-binding component of ABC superfamily | Cytoplasmic | COG4608;ABC-type oligopeptide transport system, ATPase component | GO:0000270 peptidoglycan metabolism | gg | tt |
| *oriC* | rep_origin | origin of replication | | | | | |
| *orn* | CDS | oligoribonuclease | Cytoplasmic | | GO:0006401 RNA catabolism | | |
| *osmB* | CDS | lipoprotein | Outer Membrane Lipoprotein | | GO:0006970 response to osmotic stress | o | tt |
| *osmC* | CDS | osmotically inducible, stress-inducible membrane protein | Cytoplasmic | COG1764;Predicted redox protein, regulator of disulfide bond formation | GO:0006970 response to osmotic stress | gg | t |
| *osmE* | CDS | DNA-binding transcriptional regulator | Outer Membrane Lipoprotein | | GO:0006350 transcription -!- GO:0006970 response to osmotic stress | o | tt |
| *osmY* | CDS | periplasmic protein | Periplasmic | COG2823;Predicted periplasmic or secreted lipoprotein | GO:0006970 response to osmotic stress | o | tt |
| *otsA* | CDS | trehalose-6-phosphate synthase | Cytoplasmic | COG0380;Trehalose-6-phosphate synthase | GO:0006006 glucose metabolism -!- GO:0006970 response to osmotic stress | | |
| *otsB* | CDS | trehalose-6-phosphate phosphatase, biosynthetic | Cytoplasmic | COG1877;Trehalose-6-phosphatase | GO:0006006 glucose metabolism -!- GO:0006970 response to osmotic stress | gg | o |
| *oxc* | CDS | predicted oxalyl-CoA decarboxylase | Cytoplasmic | COG0028;Thiamine pyrophosphate-requiring enzymes [acetolactate synthase, pyruvate dehydrogenase (cytochrome), glyoxylate carboligase, phosphonopyruvate decarboxylase] | | gg | tt |
| *oxyR* | CDS | DNA-binding transcriptional dual regulator | Cytoplasmic | COG0583;Transcriptional regulator | GO:0006350 transcription | gg | o |
| *oxyS* | misc_RNA | regulatory sRNA, global | | | | | |
| *paaA* | CDS | predicted multicomponent oxygenase/reductase subunit for phenylacetic acid degradation | Cytoplasmic | COG3396;Uncharacterized conserved protein | GO:0016052 carbohydrate catabolism | gg | t |
| *paaB* | CDS | predicted multicomponent oxygenase/reductase subunit for phenylacetic acid degradation | Cytoplasmic | COG3460;Uncharacterized enzyme of phenylacetate metabolism | GO:0016052 carbohydrate catabolism | gg | o |
| *paaC* | CDS | predicted multicomponent oxygenase/reductase subunit for phenylacetic acid degradation | Cytoplasmic | COG3396;Uncharacterized conserved protein | GO:0016052 carbohydrate catabolism | gg | t |
| *paaD* | CDS | predicted multicomponent oxygenase/reductase subunit for phenylacetic acid degradation | Cytoplasmic | COG2151;Predicted metal-sulfur cluster biosynthetic enzyme | GO:0016052 carbohydrate catabolism | | |
| *paaE* | CDS | predicted multicomponent oxygenase/reductase subunit for phenylacetic acid degradation | Cytoplasmic | COG1018;Flavodoxin reductases (ferredoxin-NADPH reductases) family 1 | GO:0016052 carbohydrate catabolism | o | tt |
| *paaF* | CDS | enoyl-CoA hydratase-isomerase | Cytoplasmic | COG1024;Enoyl-CoA hydratase/carnithine racemase | GO:0016052 carbohydrate | gg | t |
| *paaG* | CDS | acyl-CoA hydratase | Cytoplasmic | COG1024;Enoyl-CoA hydratase/carnithine racemase | GO:0016052 carbohydrate | gg | o |
| *paaH* | CDS | 3-hydroxybutyryl-CoA dehydrogenase | Cytoplasmic | COG1250;3-hydroxyacyl-CoA dehydrogenase | GO:0016052 carbohydrate | o | t |
| *paaI* | CDS | predicted thioesterase | Cytoplasmic | COG2050;Uncharacterized protein, possibly involved in aromatic compounds catabolism | GO:0016052 carbohydrate catabolism | gg | t |
| *paaJ* | CDS | predicted beta-ketoadipyl CoA thiolase | Cytoplasmic | COG0183;Acetyl-CoA acetyltransferase | GO:0016052 carbohydrate | o | t |
| *paaK* | CDS | phenylacetyl-CoA ligase | Cytoplasmic | COG1541;Coenzyme F390 synthetase | GO:0016052 carbohydrate | | |
| *paaX* | CDS | DNA-binding transcriptional regulator, aryl-CoA responsive | Cytoplasmic | COG3327;Phenylacetic acid-responsive transcriptional repressor | GO:0016052 carbohydrate catabolism | o | o |
| *paaY* | CDS | predicted hexapeptide repeat acetyltransferase | Cytoplasmic | COG0663;Carbonic anhydrases/acetyltransferases, isoleucine patch superfamily | | gg | t |
| *pabA* | CDS | aminodeoxychorismate synthase, subunit II | Cytoplasmic | COG0512;Anthranilate/para-aminobenzoate synthases component II | GO:0046656 folic acid biosynthesis | gg | t |
| *pabB* | CDS | aminodeoxychorismate synthase, subunit I | Cytoplasmic | COG0147;Anthranilate/para-aminobenzoate synthases component I | GO:0046656 folic acid biosynthesis | gg | o |
| *pabC* | CDS | 4-amino-4-deoxychorismate lyase component of para-aminobenzoate synthase multienzyme complex | Cytoplasmic | COG0115;Branched-chain amino acid aminotransferase/4-amino-4-deoxychorismate lyase | GO:0046656 folic acid biosynthesis | g | o |
| *pal* | CDS | peptidoglycan-associated outer membrane lipoprotein | Outer Membrane Lipoprotein | COG2885;Outer membrane protein and related peptidoglycan-associated (lipo)proteins | | o | tt |
| *panB* | CDS | 3-methyl-2-oxobutanoate hydroxymethyltransferase | Cytoplasmic | COG0413;Ketopantoate hydroxymethyltransferase | GO:0015937 coenzyme A biosynthesis | gg | t |
| *panC* | CDS | pantothenate synthetase | Cytoplasmic | COG0414;Panthothenate synthetase | GO:0015937 coenzyme A biosynthesis | gg | o |
| *panD* | CDS | aspartate 1-decarboxylase | Cytoplasmic | COG0853;Aspartate 1-decarboxylase | GO:0015937 coenzyme A biosynthesis | gg | t |
| *panE* | CDS | 2-dehydropantoate reductase, NADPH-specific | Outer Membrane Lipoprotein | COG1893;Ketopantoate reductase | GO:0006221 pyrimidine nucleotide biosynthesis | o | tt |
| *panF* | CDS | pantothenate:sodium symporter | Integral Membrane Protein | | GO:0015937 coenzyme A biosynthesis | o | tt |
| *parC* | CDS | DNA topoisomerase IV, subunit A | Cytoplasmic | COG0188;Type IIA topoisomerase (DNA gyrase/topo II, topoisomerase IV), A subunit | GO:0006261 DNA dependent DNA replication | o | tt |
| *parE* | CDS | DNA topoisomerase IV, subunit B | Cytoplasmic | COG0187;Type IIA topoisomerase (DNA gyrase/topo II, topoisomerase IV), B subunit | GO:0006261 DNA dependent DNA replication | gg | t |
| *pbl* | CDS | predicted peptidoglycan-binding enzyme (pseudogene) | Cytoplasmic | COG0741;Soluble lytic murein transglycosylase and related regulatory proteins (some contain LysM/invasin domains) | | | |
| *pbpC* | CDS | fused transglycosylase -!- transpeptidase | Periplasmic | COG4953;Membrane carboxypeptidase/penicillin-binding protein PbpC | GO:0009252 peptidoglycan biosynthesis | | |
| *pbpG* | CDS | D-alanyl-D-alanine endopeptidase | Periplasmic | COG1686;D-alanyl-D-alanine carboxypeptidase | GO:0009252 peptidoglycan biosynthesis -!- GO:0042493 response to drug | | |
| *pck* | CDS | phosphoenolpyruvate carboxykinase | Cytoplasmic | COG1866;Phosphoenolpyruvate carboxykinase (ATP) | GO:0006094 gluconeogenesis | gg | t |
| *pcm* | CDS | L-isoaspartate protein carboxylmethyltransferase type II | Cytoplasmic | COG2518;Protein-L-isoaspartate carboxylmethyltransferase | GO:0006464 protein modification | gg | t |
| *pcnB* | CDS | poly(A) polymerase I | Cytoplasmic | COG0617;tRNA nucleotidyltransferase/poly(A) polymerase | GO:0009451 RNA modification | | |
| *pdhR* | CDS | DNA-binding transcriptional dual regulator | Cytoplasmic | COG2186;Transcriptional regulators | GO:0006096 glycolysis -!- GO:0006350 transcription | o | tt |

| | | | | | | | |
|---|---|---|---|---|---|---|---|
| *pdxA* | CDS | 4-hydroxy-L-threonine phosphate dehydrogenase, NAD-dependent | Cytoplasmic | COG1995;Pyridoxal phosphate biosynthesis protein | GO:0008615 pyridoxine biosynthesis | gg | o |
| *pdxB* | CDS | erythronate-4-phosphate dehydrogenase | Cytoplasmic | COG0111;Phosphoglycerate dehydrogenase and related dehydrogenases | GO:0008615 pyridoxine biosynthesis | gg | t |
| *pdxH* | CDS | pyridoxine 5'-phosphate oxidase | Cytoplasmic | COG0259;Pyridoxamine-phosphate oxidase | GO:0008615 pyridoxine biosynthesis -!- GO:0009443 pyridoxal 5'-phosphate salvage | gg | t |
| *pdxJ* | CDS | pyridoxine 5'-phosphate synthase | Cytoplasmic | COG0854;Pyridoxal phosphate biosynthesis protein | GO:0008615 pyridoxine biosynthesis | gg | o |
| *pdxK* | CDS | pyridoxal-pyridoxamine kinase/hydroxymethylpyrimidine kinase | Cytoplasmic | COG2240;Pyridoxal/pyridoxine/pyridoxamine kinase | GO:0009443 pyridoxal 5'-phosphate salvage | gg | tt |
| *pdxY* | CDS | pyridoxal kinase 2/pyridoxine kinase | Cytoplasmic | COG2240;Pyridoxal/pyridoxine/pyridoxamine kinase | GO:0009443 pyridoxal 5'-phosphate salvage | gg | t |
| *pepA* | CDS | aminopeptidase A, a cyteinylglycinase | Cytoplasmic | COG0260;Leucyl aminopeptidase | GO:0006310 DNA recombination -!- GO:0006350 transcription | gg | t |
| *pepB* | CDS | aminopeptidase B | Cytoplasmic | | | gg | tt |
| *pepD* | CDS | aminoacyl-histidine dipeptidase (peptidase D) | Cytoplasmic | COG2195;Di- and tripeptidases | | gg | o |
| *pepE* | CDS | (alpha)-aspartyl dipeptidase | Cytoplasmic | COG3340;Peptidase E | | gg | t |
| *pepN* | CDS | aminopeptidase N | Cytoplasmic | COG0308;Aminopeptidase N | | gg | tt |
| *pepP* | CDS | proline aminopeptidase P II | Cytoplasmic | COG0006;Xaa-Pro aminopeptidase | | o | tt |
| *pepQ* | CDS | proline dipeptidase | Cytoplasmic | COG0006;Xaa-Pro aminopeptidase | | gg | t |
| *pepT* | CDS | peptidase T | Cytoplasmic | COG2195;Di- and tripeptidases | | gg | t |
| *perR* | CDS | CP4-6 prophage; predicted DNA-binding transcriptional regulator | Cytoplasmic | COG0583;Transcriptional regulator | | gg | t |
| *pfkA* | CDS | 6-phosphofructokinase I | Cytoplasmic | COG0205;6-phosphofructokinase | GO:0016052 carbohydrate | g | o |
| *pfkB* | CDS | 6-phosphofructokinase II | Cytoplasmic | | GO:0016052 carbohydrate | | |
| *pflA* | CDS | pyruvate formate lyase activating enzyme 1 | Cytoplasmic | COG1180;Pyruvate-formate lyase-activating enzyme | GO:0009061 anaerobic respiration -!- GO:0006464 protein modification | gg | o |
| *pflB* | CDS | pyruvate formate lyase I | Cytoplasmic | COG1882;Pyruvate-formate lyase | GO:0009061 anaerobic respiration -!- GO:0009063 amino acid catabolism | o | tt |
| *pflC* | CDS | pyruvate formate lyase II activase | Cytoplasmic | | GO:0009061 anaerobic respiration -!- GO:0006464 protein modification | gg | t |
| *pflD* | CDS | predicted formate acetyltransferase 2 (pyruvate formate lyase II) | Cytoplasmic | COG1882;Pyruvate-formate lyase | GO:0009061 anaerobic respiration | o | tt |
| *pfs* | CDS | 5'-methylthioadenosine/S-adenosylhomocysteine nucleosidase | Cytoplasmic | COG0775;Nucleoside phosphorylase | GO:0015949 nucleobase, nucleoside and nucleotide interconversion | gg | o |
| *pgi* | CDS | glucosephosphate isomerase | Cytoplasmic | COG0166;Glucose-6-phosphate isomerase | GO:0006096 glycolysis -!- GO:0006094 gluconeogenesis | gg | o |
| *pgk* | CDS | phosphoglycerate kinase | Cytoplasmic | COG0126;3-phosphoglycerate kinase | GO:0006096 glycolysis -!- GO:0006094 gluconeogenesis | gg | o |
| *pgm* | CDS | phosphoglucomutase | Cytoplasmic | COG0033;Phosphoglucomutase | GO:0016052 carbohydrate catabolism -!- GO:0006006 glucose | gg | tt |
| *pgpA* | CDS | phosphatidylglycerophosphatase A | Integral Membrane Protein | COG1267;Phosphatidylglycerophosphatase A and related proteins | GO:0008654 phospholipid biosynthesis | o | tt |
| *pgpB* | CDS | phosphatidylglycerophosphate phosphatase B | Integral Membrane Protein | COG0671;Membrane-associated phospholipid phosphatase | GO:0008654 phospholipid biosynthesis | o | tt |
| *pgsA* | CDS | phosphatidylglycerophosphate synthetase | Integral Membrane Protein | COG0558;Phosphatidylglycerophosphate synthase | GO:0008654 phospholipid biosynthesis | o | tt |
| *pheA* | CDS | fused chorismate mutase P -!- prephenate dehydratase | Cytoplasmic | COG1605;Chorismate mutase -!- COG0077;Prephenate dehydratase | GO:0009094 L-phenylalanine biosynthesis -!- GO:0006571 tyrosine biosynthesis | gg | o |
| *pheL* | CDS | pheA gene leader peptide | Cytoplasmic | | GO:0009094 L-phenylalanine biosynthesis | gg | o |
| *pheM* | CDS | phenylalanyl-tRNA synthetase operon leader peptide | Cytoplasmic | | GO:0006418 amino acid activation | o | o |
| *pheP* | CDS | phenylalanine transporter | Integral Membrane Protein | COG1113;Gamma-aminobutyrate permease and related permeases | GO:0009094 L-phenylalanine biosynthesis -!- GO:0009063 amino acid catabolism | o | tt |
| *pheS* | CDS | phenylalanine tRNA synthetase, alpha subunit | Cytoplasmic | COG0016;Phenylalanyl-tRNA synthetase alpha subunit | GO:0006418 amino acid activation | | |
| *pheT* | CDS | phenylalanine tRNA synthetase, beta subunit | Cytoplasmic | COG0073;EMAP domain -!- COG0072;Phenylalanyl-tRNA synthetase beta subunit | GO:0006418 amino acid activation | o | tt |
| *pheU* | tRNA | tRNA-Phe(GAA) (Phenylalanine tRNA) | | | | | |
| *pheV* | tRNA | tRNA-Phe(GAA) (Phenylalanine tRNA) | | | | | |
| *phnA* | CDS | predicted phosphonate metabolizing protein | Cytoplasmic | COG2824;Uncharacterized Zn-ribbon-containing protein involved in phosphonate metabolism | | gg | o |
| *phnB* | CDS | conserved protein | Cytoplasmic | COG2764;Uncharacterized protein conserved in bacteria | | gg | o |
| *phnC* | CDS | phosphonate/organophosphate ester transporter subunit -!- ATP-binding component of ABC superfamily | Cytoplasmic | COG3638;ABC-type phosphate/phosphonate transport system, ATPase component | GO:0006793 phosphorus metabolism | o | tt |
| *phnD* | CDS | phosphonate/organophosphate ester transporter subunit -!- periplasmic binding component of ABC superfamily | Periplasmic | COG3221;ABC-type phosphate/phosphonate transport system, periplasmic component | GO:0006793 phosphorus metabolism | o | tt |
| *phnE* | CDS | phosphonate/organophosphate ester transporter (pseudogene) | Integral Membrane Protein | COG3639;ABC-type phosphate/phosphonate transport system, permease component | | o | tt |
| *phnE* | CDS | phosphonate/organophosphate ester transporter subunit -!- membrane component of ABC superfamily | Integral Membrane Protein | COG3639;ABC-type phosphate/phosphonate transport system, permease component | GO:0006793 phosphorus metabolism | o | tt |
| *phnF* | CDS | transcriptional regulator of phosphonate uptake and biodegradation | Cytoplasmic | COG2188;Transcriptional regulators | GO:0006793 phosphorus metabolism -!- GO:0006350 transcription | gg | o |
| *phnG* | CDS | carbon-phosphorus lyase complex subunit | Cytoplasmic | COG3624;Uncharacterized enzyme of phosphonate metabolism | GO:0006793 phosphorus metabolism | gg | o |
| *phnH* | CDS | carbon-phosphorus lyase complex subunit | Cytoplasmic | COG3625;Uncharacterized enzyme of phosphonate metabolism | GO:0006793 phosphorus metabolism | gg | t |
| *phnI* | CDS | carbon-phosphorus lyase complex subunit | Cytoplasmic | COG3626;Uncharacterized enzyme of phosphonate metabolism | GO:0006793 phosphorus metabolism | gg | tt |
| *phnJ* | CDS | carbon-phosphorus lyase complex subunit | Cytoplasmic | COG3627;Uncharacterized enzyme of phosphonate metabolism | | gg | t |
| *phnK* | CDS | carbon-phosphorus lyase complex subunit | Cytoplasmic | | GO:0006793 phosphorus metabolism | | |
| *phnL* | CDS | carbon-phosphorus lyase complex subunit | Cytoplasmic | COG4778;ABC-type phosphonate transport system, ATPase component | GO:0006793 phosphorus metabolism | gg | t |
| *phnM* | CDS | carbon-phosphorus lyase complex subunit | Cytoplasmic | COG3454;Metal-dependent hydrolase involved in phosphonate metabolism | GO:0006793 phosphorus metabolism | gg | t |
| *phnN* | CDS | ribose 1,5-bisphosphokinase | Cytoplasmic | COG3709;Uncharacterized component of phosphonate metabolism | GO:0006793 phosphorus metabolism | gg | tt |
| *phnO* | CDS | predicted acyltransferase with acyl-CoA N-acyltransferase domain | Cytoplasmic | COG0454;Histone acetyltransferase HPA2 and related acetyltransferases | | gg | o |
| *phnP* | CDS | carbon-phosphorus lyase complex accessory protein | Periplasmic | COG1235;Metal-dependent hydrolases of the beta-lactamase superfamily I | GO:0006793 phosphorus metabolism | gg | o |
| *phoA* | CDS | alkaline phosphatase | Periplasmic | | GO:0006793 phosphorus metabolism | o | tt |
| *phoB* | CDS | DNA-binding response regulator in two-component regulatory system with PhoR (or CreC) | Cytoplasmic | COG0745;Response regulators consisting of a CheY-like receiver domain and a winged-helix DNA-binding domain | GO:0006793 phosphorus metabolism -!- GO:0006350 transcription | gg | o |
| *phoE* | CDS | outer membrane phosphoporin protein E | Outer Membrane B-barrel protein | COG3203;Outer membrane protein (porin) | | o | tt |
| *phoH* | CDS | conserved protein with nucleoside triphosphate hydrolase domain | Cytoplasmic | COG1702;Phosphate starvation-inducible protein PhoH, predicted ATPase | GO:0006793 phosphorus metabolism -!- GO:0042594 response to | gg | t |
| *phoP* | CDS | DNA-binding response regulator in two-component regulatory system with PhoQ | Cytoplasmic | COG0745;Response regulators consisting of a CheY-like receiver domain and a winged-helix DNA-binding domain | GO:0006350 transcription | gg | o |
| *phoQ* | CDS | sensory histidine kinase in two-compoent regulatory system with PhoP | Integral Membrane Protein | COG0642;Signal transduction histidine kinase | GO:0006464 protein modification | o | tt |
| *phoR* | CDS | sensory histidine kinase in two-component regulatory system with PhoB | Membrane Anchored | COG0642;Signal transduction histidine kinase | GO:0006793 phosphorus metabolism -!- GO:0006464 protein modification | o | t |
| *phoU* | CDS | DNA-binding transcriptional regulator | Cytoplasmic | COG0704;Phosphate uptake regulator | GO:0006793 phosphorus metabolism | gg | o |
| *php* | CDS | predicted hydrolase | Cytoplasmic | COG1735;Predicted metal-dependent hydrolase with the TIM-barrel fold | | gg | t |
| *phr* | CDS | deoxyribodipyrimidine photolyase, FAD-binding | Cytoplasmic | COG0415;Deoxyribodipyrimidine photolyase | GO:0006281 DNA repair | o | tt |
| *pin* | CDS | e14 prophage; site-specific DNA recombinase | Cytoplasmic | COG1961;Site-specific recombinases, DNA invertase Pin homologs | | o | tt |
| *pinH* | CDS | predicted invertase fragment (pseudogene) | Cytoplasmic | | | | |
| *pinQ* | CDS | Qin prophage; predicted site-specific recombinase | Cytoplasmic | COG1961;Site-specific recombinases, DNA invertase Pin homologs | | o | tt |
| *pinR* | CDS | Rac prophage; predicted site-specific recombinase | Cytoplasmic | COG1961;Site-specific recombinases, DNA invertase Pin homologs | | o | tt |
| *pioO* | CDS | part of gsp divergon involved in type II protein secretion | Membrane Anchored | | GO:0006261 DNA dependent DNA replication | o | tt |
| *pitA* | CDS | phosphate transporter, low-affinity | Integral Membrane Protein | COG0306;Phosphate/sulphate permeases | GO:0006793 phosphorus metabolism | o | tt |
| *pitB* | CDS | phosphate transporter | Integral Membrane Protein | COG0306;Phosphate/sulphate permeases | GO:0006793 phosphorus metabolism | o | tt |
| *pldA* | CDS | outer membrane phospholipase A | Outer Membrane B-barrel protein | COG2829;Outer membrane phospholipase A | GO:0006793 phosphorus metabolism | o | tt |
| *pldB* | CDS | lysophospholipase L(2) | Cytoplasmic | | GO:0006793 phosphorus metabolism | | |
| *plsB* | CDS | glycerol-3-phosphate O-acyltransferase | Cytoplasmic | | GO:0008654 phospholipid biosynthesis | o | tt |
| *plsC* | CDS | 1-acyl-sn-glycerol-3-phosphate acyltransferase | Membrane Anchored | COG0204;1-acyl-sn-glycerol-3-phosphate acyltransferase | GO:0008654 phospholipid biosynthesis | o | tt |
| *plsX* | CDS | fatty acid/phospholipid synthesis protein | Cytoplasmic | COG0416;Fatty acid/phospholipid biosynthesis enzyme | GO:0008654 phospholipid biosynthesis | | |
| *pmbA* | CDS | predicted peptidase required for the maturation and secretion of the antibiotic peptide MccB17 | Cytoplasmic | COG0312;Predicted Zn-dependent proteases and their inactivated homologs | GO:0006457 protein folding | gg | tt |
| *pmrD* | CDS | polymyxin resistance protein B | Cytoplasmic | | GO:0042493 response to drug | gg | o |
| *pncA* | CDS | nicotinamidase/pyrazinamidase | Cytoplasmic | | GO:0009435 nicotinamide adenine dinucleotide biosynthesis | gg | o |
| *pncB* | CDS | nicotinate phosphoribosyltransferase | Cytoplasmic | COG1488;Nicotinic acid phosphoribosyltransferase | GO:0009435 nicotinamide adenine dinucleotide biosynthesis | gg | tt |
| *pnp* | CDS | polynucleotide phosphorylase/polyadenylase | Cytoplasmic | | GO:0006401 RNA catabolism -!- GO:0006401 RNA catabolism | | |
| *pntA* | CDS | pyridine nucleotide transhydrogenase, alpha subunit | Integral Membrane Protein | COG3288;NAD/NADP transhydrogenase alpha subunit | | o | tt |
| *pntB* | CDS | pyridine nucleotide transhydrogenase, beta subunit | Integral Membrane Protein | COG1282;NAD/NADP transhydrogenase beta subunit | | o | tt |
| *pnuC* | CDS | predicted nicotinamide mononucleotide transporter | Integral Membrane Protein | COG3201;Nicotinamide mononucleotide transporter | GO:0015949 nucleobase, nucleoside and nucleotide interconversion | o | tt |
| *polA* | CDS | fused DNA polymerase I 5'->3' exonuclease -!- 3'->5' polymerase -!- 3'->5' exonuclease | Cytoplasmic | COG0258;5'-3' exonuclease (including N-terminal domain of PolI) -!- COG0749;DNA polymerase I - 3'-5' exonuclease and polymerase domains | GO:0006261 DNA dependent DNA replication | o | tt |
| *polB* | CDS | DNA polymerase II | Cytoplasmic | COG0417;DNA polymerase elongation subunit (family B) | GO:0006261 DNA dependent DNA replication -!- GO:0006281 DNA repair -!- GO:0009314 response to radiation -!- GO:0009432 SOS | g | tt |
| *potA* | CDS | polyamine transporter subunit -!- ATP-binding component of ABC superfamily | Cytoplasmic | COG3842;ABC-type spermidine/putrescine transport systems, ATPase components | GO:0006596 polyamine biosynthesis -!- GO:0009310 amine catabolism | gg | t |

| | | | | | | | |
|---|---|---|---|---|---|---|---|
| *potB* | CDS | polyamine transporter subunit -!- membrane component of ABC superfamily | Integral Membrane Protein | COG1176;ABC-type spermidine/putrescine transport system, permease component I | GO:0006596 polyamine biosynthesis -!- GO:0009310 amine catabolism | o | tt |
| *potC* | CDS | polyamine transporter subunit -!- membrane component of ABC superfamily | Integral Membrane Protein | COG1177;ABC-type spermidine/putrescine transport system, permease component II | GO:0006596 polyamine biosynthesis -!- GO:0009310 amine catabolism | o | tt |
| *potD* | CDS | polyamine transporter subunit -!- periplasmic-binding component of ABC superfamily | Periplasmic | COG0687;Spermidine/putrescine-binding periplasmic protein | GO:0006596 polyamine biosynthesis -!- GO:0009310 amine catabolism | o | tt |
| *potE* | CDS | putrescine/proton symporter -!- putrescine/ornithine antiporter | Integral Membrane Protein | COG0531;Amino acid transporters | GO:0009310 amine catabolism -!- GO:0006596 polyamine biosynthesis -!- GO:0006970 response to osmotic stress | o | tt |
| *potF* | CDS | putrescine transporter subunit -!- periplasmic-binding component of ABC superfamily | Periplasmic | COG0687;Spermidine/putrescine-binding periplasmic protein | GO:0006596 polyamine biosynthesis -!- GO:0009310 amine catabolism -!- GO:0015847 putrescine transport | o | tt |
| *potG* | CDS | putrescine transporter subunit -!- ATP-binding component of ABC superfamily | Cytoplasmic | | GO:0006596 polyamine biosynthesis -!- GO:0009310 amine catabolism -!- GO:0015847 putrescine transport | | |
| *potH* | CDS | putrescine transporter subunit -!- membrane component of ABC superfamily | Integral Membrane Protein | COG1176;ABC-type spermidine/putrescine transport system, permease component I | GO:0006596 polyamine biosynthesis -!- GO:0009310 amine catabolism -!- GO:0015847 putrescine transport | o | tt |
| *potI* | CDS | putrescine transporter subunit -!- membrane component of ABC superfamily | Integral Membrane Protein | COG1177;ABC-type spermidine/putrescine transport system, permease component II | GO:0006596 polyamine biosynthesis -!- GO:0009310 amine catabolism -!- GO:0015847 putrescine transport | o | tt |
| *poxA* | CDS | predicted lysyl-tRNA synthetase | Cytoplasmic | COG2269;Truncated, possibly inactive, lysyl-tRNA synthetase (class II) | GO:0006418 amino acid activation | o | tt |
| *poxB* | CDS | pyruvate dehydrogenase (pyruvate oxidase), thiamin-dependent, FAD-binding | Cytoplasmic | COG0028;Thiamine pyrophosphate-requiring enzymes [acetolactate synthase, pyruvate dehydrogenase (cytochrome), glyoxylate carboligase, phosphonopyruvate decarboxylase] | GO:0016052 carbohydrate catabolism -!- GO:0042867 pyruvate catabolism | gg | t |
| *ppa* | CDS | inorganic pyrophosphatase | Cytoplasmic | COG0221;Inorganic pyrophosphatase | GO:0006793 phosphorus metabolism | gg | o |
| *ppc* | CDS | phosphoenolpyruvate carboxylase | Cytoplasmic | COG2352;Phosphoenolpyruvate carboxylase | GO:0006113 fermentation -!- GO:0006099 tricarboxylic acid cycle | gg | tt |
| *ppdA* | CDS | conserved protein | Periplasmic | COG2165;Type II secretory pathway, pseudopilin PulG | GO:0009101 glycoprotein biosynthesis | o | tt |
| *ppdB* | CDS | conserved protein | Membrane Anchored | COG4795;Type II secretory pathway, component PulJ | GO:0009101 glycoprotein biosynthesis | | |
| *ppdC* | CDS | predicted protein | Membrane Anchored | COG4967;Tfp pilus assembly protein PilV | GO:0009101 glycoprotein biosynthesis | o | tt |
| *ppdD* | CDS | predicted major pilin subunit | Membrane Anchored | COG4969;Tfp pilus assembly protein, major pilin PilA | | o | tt |
| *pphA* | CDS | serine/threonine-specific protein phosphatase 1 | Cytoplasmic | | GO:0006464 protein modification -!- GO:0009266 response to | g | t |
| *pphB* | CDS | serine/threonine-specific protein phosphatase 2 | Cytoplasmic | COG0639;Diadenosine tetraphosphatase and related serine/threonine protein phosphatases | GO:0006464 protein modification | g | o |
| *ppiA* | CDS | peptidyl-prolyl cis-trans isomerase A (rotamase A) | Periplasmic | COG0652;Peptidyl-prolyl cis-trans isomerase (rotamase) - cyclophilin family | GO:0006457 protein folding | o | tt |
| *ppiB* | CDS | peptidyl-prolyl cis-trans isomerase B (rotamase B) | Cytoplasmic | COG0652;Peptidyl-prolyl cis-trans isomerase (rotamase) - cyclophilin family | GO:0006457 protein folding | gg | t |
| *ppiC* | CDS | peptidyl-prolyl cis-trans isomerase C (rotamase C) | Cytoplasmic | COG0760;Parvulin-like peptidyl-prolyl isomerase | GO:0006457 protein folding | gg | t |
| *ppiD* | CDS | peptidyl-prolyl cis-trans isomerase (rotamase D) | Membrane Anchored | COG0760;Parvulin-like peptidyl-prolyl isomerase | GO:0006457 protein folding | o | tt |
| *ppk* | CDS | polyphosphate kinase, component of RNA degradosome | Cytoplasmic | COG0855;Polyphosphate kinase | GO:0006793 phosphorus metabolism | o | tt |
| *pppA* | CDS | bifunctional prepilin leader peptidase -!- methylase | Integral Membrane Protein | | | g | tt |
| *pps* | CDS | phosphoenolpyruvate synthase | Cytoplasmic | COG0574;Phosphoenolpyruvate synthase/pyruvate phosphate dikinase | GO:0006094 gluconeogenesis | o | tt |
| *ppx* | CDS | exopolyphosphatase | Cytoplasmic | COG0248;Exopolyphosphatase | GO:0006793 phosphorus metabolism | gg | tt |
| *pqiA* | CDS | paraquat-inducible membrane protein A | Integral Membrane Protein | COG2995;Uncharacterized paraquat-inducible protein A | | o | tt |
| *pqiB* | CDS | paraquat-inducible protein B | Membrane Anchored | COG3008;Paraquat-inducible protein B | | o | tt |
| *pqqL* | CDS | predicted peptidase | Cytoplasmic | COG0612;Predicted Zn-dependent peptidases | | gg | t |
| *prc* | CDS | carboxy-terminal protease for penicillin-binding protein 3 | Periplasmic | COG0793;Periplasmic protease | GO:0009252 peptidoglycan biosynthesis -!- GO:0006457 protein folding -!- GO:0042493 response to drug | o | o |
| *prfA* | CDS | peptide chain release factor RF-1 | Cytoplasmic | COG0216;Protein chain release factor A | GO:0006412 protein biosynthesis | o | tt |
| *prfB* | CDS | peptide chain release factor RF-2 | Cytoplasmic | COG1186;Protein chain release factor B | GO:0006412 protein biosynthesis | | |
| *prfC* | CDS | peptide chain release factor RF-3 | Cytoplasmic | COG4108;Peptide chain release factor RF-3 | GO:0006412 protein biosynthesis | | |
| *prfH* | CDS | predicted peptide chain release factor | Cytoplasmic | COG1186;Protein chain release factor B | GO:0006412 protein biosynthesis | g | o |
| *priA* | CDS | Primosome factor n' (replication factor Y) | Cytoplasmic | COG1198;Primosomal protein N' (replication factor Y) - superfamily II helicase | GO:0006261 DNA dependent DNA replication | o | tt |
| *priB* | CDS | primosomal protein N | Cytoplasmic | COG2965;Primosomal replication protein N | GO:0006261 DNA dependent DNA replication | o | tt |
| *priC* | CDS | primosomal replication protein N'' | Cytoplasmic | COG3923;Primosomal replication protein N'' | GO:0006261 DNA dependent DNA replication | gg | tt |
| *prkB* | CDS | predicted phosphoribulokinase | Periplasmic | COG3954;Phosphoribulokinase | GO:0006006 glucose metabolism | gg | t |
| *prlC* | CDS | oligopeptidase A | Cytoplasmic | COG0339;Zn-dependent oligopeptidases | | gg | o |
| *prmA* | CDS | methylase for 50S ribosomal subunit protein L11 | Cytoplasmic | COG2264;Ribosomal protein L11 methylase | GO:0009451 RNA modification | gg | tt |
| *prmB* | CDS | N5-glutamine methyltransferase | Cytoplasmic | | | | |
| *prmC* | CDS | N5-glutamine methyltransferase, modifies release factors RF-1 and RF-2 | Cytoplasmic | COG2890;Methylase of polypeptide chain release factors | | gg | t |
| *proA* | CDS | gamma-glutamylphosphate reductase | Cytoplasmic | COG0014;Gamma-glutamyl phosphate reductase | GO:0006561 proline biosynthesis | g | t |
| *proB* | CDS | gamma-glutamate kinase | Cytoplasmic | COG0263;Glutamate 5-kinase | GO:0006561 proline biosynthesis | g | t |
| *proC* | CDS | pyrroline-5-carboxylate reductase, NAD(P)-binding | Cytoplasmic | COG0345;Pyrroline-5-carboxylate reductase | GO:0006561 proline biosynthesis | gg | o |
| *proK* | tRNA | tRNA-Pro(CGG) (Proline tRNA1) | | | | | |
| *proL* | tRNA | tRNA-Pro(GGG) (Proline tRNA2) | | | | | |
| *proM* | tRNA | tRNA-Pro(UGG) (Proline tRNA3) | | | | | |
| *proP* | CDS | proline/glycine betaine transporter | Integral Membrane Protein | COG0477;Permeases of the major facilitator superfamily | GO:0006561 proline biosynthesis -!- GO:0006970 response to osmotic stress | o | tt |
| *proQ* | CDS | predicted structural transport element | Cytoplasmic | | GO:0006970 response to osmotic stress | | |
| *proS* | CDS | prolyl-tRNA synthetase | Cytoplasmic | COG0442;Prolyl-tRNA synthetase | GO:0006418 amino acid activation | o | tt |
| *proV* | CDS | glycine betaine transporter subunit -!- ATP-binding compoent of ABC superfamily | Cytoplasmic | COG4175;ABC-type proline/glycine betaine transport system, ATPase component | GO:0006561 proline biosynthesis | gg | o |
| *proW* | CDS | glycine betaine transporter subunit -!- membrane component of ABC superfamily | Integral Membrane Protein | COG4176;ABC-type proline/glycine betaine transport system, permease component | GO:0006561 proline biosynthesis | o | tt |
| *proX* | CDS | glycine betaine transporter subunit -!- periplasmic-binding component of ABC superfamily | Periplasmic | COG2113;ABC-type proline/glycine betaine transport systems, periplasmic components | GO:0016052 carbohydrate catabolism -!- GO:0006561 proline biosynthesis -!- GO:0006545 glycine biosynthesis -!- GO:0006578 betaine biosynthesis -!- GO:0006970 response to osmotic stress | o | tt |
| *proY* | CDS | predicted cryptic proline transporter | Integral Membrane Protein | COG1113;Gamma-aminobutyrate permease and related permeases | GO:0009063 amino acid catabolism -!- GO:0006526 arginine biosynthesis | | |
| *prpB* | CDS | 2-methylisocitrate lyase | Cytoplasmic | COG2513;PEP phosphonomutase and related enzymes | GO:0019395 fatty acid oxidation | gg | t |
| *prpC* | CDS | 2-methylcitrate synthase | Cytoplasmic | COG0372;Citrate synthase | GO:0019395 fatty acid oxidation | gg | t |
| *prpD* | CDS | 2-methylcitrate dehydratase | Cytoplasmic | COG2079;Uncharacterized protein involved in propionate catabolism | GO:0016052 carbohydrate | g | o |
| *prpE* | CDS | predicted propionyl-CoA synthetase with ATPase domain | Cytoplasmic | COG0365;Acyl-coenzyme A synthetases/AMP-(fatty) acid ligases | GO:0019395 fatty acid oxidation | gg | t |
| *prpR* | CDS | DNA-binding transcriptional regulator | Cytoplasmic | COG1221;Transcriptional regulators containing an AAA-type ATPase domain and a DNA-binding domain | GO:0019395 fatty acid oxidation -!- GO:0006350 transcription | g | tt |
| *prsA* | CDS | phosphoribosylpyrophosphate synthase | Cytoplasmic | COG0462;Phosphoribosylpyrophosphate synthetase | GO:0009152 purine ribonucleotide biosynthesis | gg | t |
| *psd* | CDS | phosphatidylserine decarboxylase | Cytoplasmic | COG0688;Phosphatidylserine decarboxylase | GO:0008654 phospholipid biosynthesis | o | tt |
| *psiF* | CDS | conserved protein | Periplasmic | | GO:0006793 phosphorus metabolism -!- GO:0042594 response to | | |
| *pspA* | CDS | regulatory protein for phage-shock-protein operon | Cytoplasmic | COG1842;Phage shock protein A (IM30), suppresses sigma54-dependent transcription | GO:0006350 transcription | gg | o |
| *pspB* | CDS | transcriptional regulator of psp operon | Membrane Anchored | | GO:0006350 transcription | gg | tt |
| *pspC* | CDS | transcriptional activator | Membrane Anchored | COG1983;Putative stress-responsive transcriptional regulator | GO:0006350 transcription | gg | tt |
| *pspD* | CDS | peripheral inner membrane phage-shock protein | Cytoplasmic | | | g | tt |
| *pspE* | CDS | thiosulfate:cyanide sulfurtransferase (rhodanese) | Periplasmic | COG0607;Rhodanese-related sulfurtransferase | | g | tt |
| *pspF* | CDS | DNA-binding transcriptional activator | Cytoplasmic | | GO:0006350 transcription | gg | tt |
| *pssA* | CDS | phosphatidylserine synthase (CDP-diacylglycerol-serine O-phosphatidyltransferase) | Cytoplasmic | | GO:0008654 phospholipid biosynthesis | gg | tt |
| *pstA* | CDS | phosphate transporter subunit -!- membrane component of ABC superfamily | Integral Membrane Protein | COG0581;ABC-type phosphate transport system, permease component | GO:0006793 phosphorus metabolism | o | tt |
| *pstB* | CDS | phosphate transporter subunit -!- ATP-binding component of ABC superfamily | Cytoplasmic | COG1117;ABC-type phosphate transport system, ATPase component | GO:0006793 phosphorus metabolism | g | t |
| *pstC* | CDS | phosphate transporter subunit -!- membrane component of ABC superfamily | Integral Membrane Protein | COG0573;ABC-type phosphate transport system, permease component | GO:0006793 phosphorus metabolism | o | tt |
| *pstS* | CDS | phosphate transporter subunit -!- periplasmic-binding component of ABC superfamily | Periplasmic | COG0226;ABC-type phosphate transport system, periplasmic component | GO:0006793 phosphorus metabolism | o | tt |
| *pta* | CDS | phosphate acetyltransferase | Cytoplasmic | COG0857;BioD-like N-terminal domain of phosphotransacetylase -!- COG0280;Phosphotransacetylase | GO:0016052 carbohydrate catabolism -!- GO:0009063 amino acid catabolism -!- GO:0042867 pyruvate catabolism -!- GO:0045733 acetate catabolism | gg | t |
| *pth* | CDS | peptidyl-tRNA hydrolase | Cytoplasmic | COG0193;Peptidyl-tRNA hydrolase | GO:0006418 amino acid activation | gg | t |
| *ptr* | CDS | protease III | Periplasmic | COG1025;Secreted/periplasmic Zn-dependent peptidases, insulinase-like | | o | tt |
| *ptrB* | CDS | protease II | Cytoplasmic | COG1770;Protease II | | o | tt |
| *ptsA* | CDS | fused predicted PTS enzymes Hpr component -!- enzyme I component -!- enzyme IIA component | Cytoplasmic | COG1080;Phosphoenolpyruvate-protein kinase (PTS system EI component in bacteria) -!- COG1762;Phosphotransferase system mannitol/fructose-specific IIA domain (Ntr-type) | GO:0016052 carbohydrate catabolism | | |
| *ptsG* | CDS | fused glucose-specific PTS enzyme IIBC components | Integral Membrane Protein | COG1263;Phosphotransferase system IIC components, glucose/maltose/N-acetylglucosamine-specific -!- COG1264;Phosphotransferase system IIB components | GO:0016052 carbohydrate catabolism -!- GO:0006464 protein modification | o | tt |

| | | | | | | | |
|---|---|---|---|---|---|---|---|
| *ptsH* | CDS | phosphohistidinoprotein-hexose phosphotransferase component of PTS system (Hpr) | Cytoplasmic | COG1925;Phosphotransferase system, HPr-related proteins | GO:0016052 carbohydrate catabolism | gg | o |
| *ptsI* | CDS | PEP-protein phosphotransferase of PTS system (enzyme I) | Cytoplasmic | COG1080;Phosphoenolpyruvate-protein kinase (PTS system EI component in bacteria) | GO:0016052 carbohydrate catabolism | gg | t |
| *ptsN* | CDS | sugar-specific enzyme IIA component of PTS | Cytoplasmic | COG1762;Phosphotransferase system mannitol/fructose-specific IIA domain (Ntr-type) | | gg | o |
| *ptsP* | CDS | fused PEP-protein phosphotransferase (enzyme I) of PTS system | Cytoplasmic | COG3605;Signal transduction protein containing GAF and PtsI domains | GO:0016052 carbohydrate catabolism | o | t |
| *purA* | CDS | adenylosuccinate synthetase | Cytoplasmic | COG0104;Adenylosuccinate synthase | GO:0006164 purine nucleotide biosynthesis -!- GO:0015949 nucleobase, nucleoside and nucleotide interconversion | gg | o |
| *purB* | CDS | adenylosuccinate lyase | Cytoplasmic | COG0015;Adenylosuccinate lyase | GO:0006164 purine nucleotide biosynthesis | gg | t |
| *purC* | CDS | phosphoribosylaminoimidazole-succinocarboxamide synthetase | Cytoplasmic | COG0152;Phosphoribosylaminoimidazolesuccinocarboxa mide (SAICAR) synthase | GO:0006164 purine nucleotide biosynthesis | gg | o |
| *purD* | CDS | phosphoribosylglycinamide synthetase phosphoribosylamine-glycine ligase | Cytoplasmic | COG0151;Phosphoribosylamine-glycine ligase | GO:0006164 purine nucleotide biosynthesis | gg | tt |
| *purE* | CDS | N5-carboxyaminoimidazole ribonucleotide mutase | Cytoplasmic | COG0041;Phosphoribosylcarboxyaminoimidazole (NCAIR) mutase | GO:0006164 purine nucleotide biosynthesis | gg | o |
| *purF* | CDS | amidophosphoribosyltransferase | Cytoplasmic | COG0034;Glutamine phosphoribosylpyrophosphate amidotransferase | GO:0006164 purine nucleotide biosynthesis | gg | t |
| *purH* | CDS | fused IMP cyclohydrolase -!- phosphoribosylaminoimidazolecarboxamide formyltransferase | Cytoplasmic | COG0138;AICAR transformylase/IMP cyclohydrolase PurH (only IMP cyclohydrolase domain in Aful) | GO:0006164 purine nucleotide biosynthesis -!- GO:0015949 nucleobase, nucleoside and nucleotide interconversion | gg | t |
| *purK* | CDS | N5-carboxyaminoimidazole ribonucleotide synthase | Cytoplasmic | COG0026;Phosphoribosylaminoimidazole carboxylase (NCAIR synthetase) | GO:0006164 purine nucleotide biosynthesis | gg | t |
| *purL* | CDS | phosphoribosylformyl-glycineamide synthetase | Cytoplasmic | COG0046;Phosphoribosylformylglycinamidine (FGAM) synthase, synthetase domain -!- COG0047;Phosphoribosylformylglycinamidine (FGAM) synthase, glutamine amidotransferase domain | GO:0006164 purine nucleotide biosynthesis | o | tt |
| *purM* | CDS | phosphoribosylaminoimidazole synthetase | Cytoplasmic | COG0150;Phosphoribosylaminoimidazole (AIR) synthetase | GO:0006164 purine nucleotide biosynthesis | gg | t |
| *purN* | CDS | phosphoribosylglycinamide formyltransferase 1 | Cytoplasmic | COG0299;Folate-dependent phosphoribosylglycinamide formyltransferase PurN | GO:0006164 purine nucleotide biosynthesis | gg | t |
| *purR* | CDS | DNA-binding transcriptional repressor, hypoxanthine-binding | Cytoplasmic | COG1609;Transcriptional regulators | GO:0006164 purine nucleotide biosynthesis -!- GO:0006596 polyamine biosynthesis -!- GO:0000105 histidine biosynthesis -!- GO:0009063 amino acid catabolism -!- GO:0006545 glycine biosynthesis -!- GO:0006807 nitrogen metabolism -!- GO:0009257 10-formyltetr | gg | t |
| *purT* | CDS | phosphoribosylglycinamide formyltransferase 2 | Cytoplasmic | COG0027;Formate-dependent phosphoribosylglycinamide formyltransferase (GAR transformylase) | GO:0006164 purine nucleotide biosynthesis | gg | t |
| *purU* | CDS | formyltetrahydrofolate hydrolase | Cytoplasmic | COG0788;Formyltetrahydrofolate hydrolase | GO:0009257 10-formyltetrahydrofolate biosynthesis -!- GO:0009152 purine ribonucleotide | g | o |
| *putA* | CDS | fused DNA-binding transcriptional regulator -!- proline dehydrogenase -!- pyrroline-5-carboxylate dehydrogenase | Cytoplasmic | COG0506;Proline dehydrogenase -!- COG4230;Delta 1-pyrroline-5-carboxylate dehydrogenase | GO:0009063 amino acid catabolism -!- GO:0006350 transcription | g | tt |
| *putP* | CDS | proline:sodium symporter | Integral Membrane Protein | COG0591;Na+/proline symporter | GO:0006561 proline biosynthesis | o | tt |
| *puuA* | CDS | gamma-Glu-putrescine synthase | Cytoplasmic | | GO:0006542 glutamine biosynthesis -!- GO:0006807 nitrogen metabolism | | |
| *puuB* | CDS | gamma-Glu-putrescine oxidase, FAD/NAD(P)-binding | Cytoplasmic | COG0665;Glycine/D-amino acid oxidases (deaminating) | | o | tt |
| *puuC* | CDS | gamma-Glu-gamma-aminobutyraldehyde dehydrogenase, NAD(P)H-dependent | Cytoplasmic | COG1012;NAD-dependent aldehyde dehydrogenases | GO:0006113 fermentation | o | tt |
| *puuD* | CDS | gamma-Glu-GABA hydrolase | Cytoplasmic | | GO:0009310 amine catabolism | gg | o |
| *puuE* | CDS | GABA aminotransferase, PLP-dependent | Periplasmic | COG0160;4-aminobutyrate aminotransferase and related aminotransferases | GO:0009310 amine catabolism -!- GO:0008615 pyridoxine biosynthesis | gg | tt |
| *puuP* | CDS | putrescine importer | Integral Membrane Protein | | | gg | tt |
| *puuR* | CDS | DNA-binding transcriptional repressor | Cytoplasmic | COG1396;Predicted transcriptional regulators | | g | o |
| *pykA* | CDS | pyruvate kinase II | Cytoplasmic | COG0469;Pyruvate kinase | GO:0006096 glycolysis -!- GO:0006113 fermentation -!- GO:0009061 anaerobic respiration | gg | t |
| *pykF* | CDS | pyruvate kinase I | Cytoplasmic | COG0469;Pyruvate kinase | GO:0006096 glycolysis -!- GO:0006113 fermentation -!- GO:0009061 anaerobic respiration | o | t |
| *pyrB* | CDS | aspartate carbamoyltransferase, catalytic subunit | Cytoplasmic | COG0540;Aspartate carbamoyltransferase, catalytic chain | | o | t |
| *pyrC* | CDS | dihydro-orotase | Cytoplasmic | COG0418;Dihydroorotase | GO:0006221 pyrimidine nucleotide biosynthesis | gg | t |
| *pyrD* | CDS | dihydro-orotate oxidase, FMN-linked | Cytoplasmic | COG0167;Dihydroorotate dehydrogenase | GO:0006221 pyrimidine nucleotide biosynthesis | gg | o |
| *pyrE* | CDS | orotate phosphoribosyltransferase | Cytoplasmic | COG0461;Orotate phosphoribosyltransferase | GO:0006221 pyrimidine nucleotide biosynthesis | g | t |
| *pyrF* | CDS | orotidine-5'-phosphate decarboxylase | Cytoplasmic | COG0284;Orotidine-5'-phosphate decarboxylase | GO:0006221 pyrimidine nucleotide biosynthesis -!- GO:0015949 nucleobase, nucleoside and nucleotide interconversion | gg | o |
| *pyrG* | CDS | CTP synthetase | Cytoplasmic | COG0504;CTP synthase (UTP-ammonia lyase) | GO:0015949 nucleobase, nucleoside and nucleotide interconversion | gg | t |
| *pyrH* | CDS | uridylate kinase | Cytoplasmic | COG0528;Uridylate kinase | GO:0015949 nucleobase, nucleoside and nucleotide interconversion | gg | o |
| *pyrI* | CDS | aspartate carbamoyltransferase, regulatory subunit | Cytoplasmic | COG1781;Aspartate carbamoyltransferase, regulatory subunit | GO:0006221 pyrimidine nucleotide biosynthesis | gg | o |
| *pyrL* | CDS | pyrBI operon leader peptide | Cytoplasmic | | GO:0006221 pyrimidine nucleotide biosynthesis | o | o |
| *qor* | CDS | quinone oxidoreductase, NADPH-dependent | Cytoplasmic | COG0604;NADPH:quinone reductase and related Zn-dependent oxidoreductases | | gg | t |
| *qseB* | CDS | DNA-binding response regulator in two-component regulatory system with QseC | Cytoplasmic | COG0745;Response regulators consisting of a CheY-like receiver domain and a winged-helix DNA-binding domain | GO:0006355 regulation of transcription, DNA-dependent | gg | t |
| *qseC* | CDS | sensory histidine kinase in two-component regulatory system with QseB | Integral Membrane Protein | COG0642;Signal transduction histidine kinase | GO:0006464 protein modification | o | tt |
| *queA* | CDS | S-adenosylmethionine:tRNA ribosyltransferase-isomerase | Cytoplasmic | COG0809;S-adenosylmethionine:tRNA-ribosyltransfera se-isomerase (queuine synthetase) | GO:0009451 RNA modification | gg | o |
| *racC* | CDS | Rac prophage; predicted protein | Cytoplasmic | | | g | o |
| *racR* | CDS | Rac prophage; predicted DNA-binding transcriptional regulator | Cytoplasmic | | | g | tt |
| *radA* | CDS | predicted repair protein | Cytoplasmic | COG1066;Predicted ATP-dependent serine protease | | o | t |
| *rarA* | CDS | predicted hydrolase | Cytoplasmic | COG0596;Predicted hydrolases or acyltransferases (alpha/beta hydrolase superfamily) | | gg | o |
| *rarD* | CDS | predicted chloramphenical resistance permease | Integral Membrane Protein | | GO:0042493 response to drug -!- GO:0042892 chloramphenicol transport | | |
| *rbbA* | CDS | fused ribosome-associated ATPases | Cytoplasmic | COG1131;ABC-type multidrug transport system, ATPase component -!- COG0842;ABC-type multidrug transport system, permease component | GO:0006412 protein biosynthesis | | |
| *rbfA* | CDS | 30s ribosome binding factor | Cytoplasmic | COG0858;Ribosome-binding factor A | GO:0009451 RNA modification | gg | o |
| *rbn* | CDS | tRNA processing exoribonuclease BN | Cytoplasmic | COG1295;Predicted membrane protein | GO:0009451 RNA modification | o | tt |
| *rbsA* | CDS | fused D-ribose transporter subunits -!- ATP-binding components ABC superfamily | Cytoplasmic | COG1129;ABC-type sugar transport system, ATPase component | GO:0016052 carbohydrate catabolism | gg | o |
| *rbsB* | CDS | D-ribose transporter subunit -!- periplasmic-binding compoent of ABC superfamily | Periplasmic | COG1879;ABC-type sugar transport system, periplasmic component | GO:0016052 carbohydrate catabolism | o | tt |
| *rbsC* | CDS | D-ribose transporter subunit -!- membrane component of ABC superfamily | Integral Membrane Protein | COG1172;Ribose/xylose/arabinose/galactoside ABC-type transport systems, permease components | GO:0016052 carbohydrate catabolism | o | tt |
| *rbsD* | CDS | predicted cytoplasmic sugar-binding protein | Cytoplasmic | COG1869;ABC-type ribose transport system, auxiliary component | GO:0016052 carbohydrate | | |
| *rbsK* | CDS | ribokinase | Cytoplasmic | COG0524;Sugar kinases, ribokinase family | GO:0016052 carbohydrate | gg | t |
| *rbsR* | CDS | DNA-binding transcriptional repressor | Cytoplasmic | COG1609;Transcriptional regulators | GO:0016052 carbohydrate catabolism -!- GO:0006350 | g | o |
| *rcsA* | CDS | DNA-binding transcriptional co-regulator with RcsB | Cytoplasmic | COG2771;DNA-binding HTH domain-containing proteins | GO:0009242 colanic acid biosynthesis -!- GO:0009248 K antigen biosynthesis -!- GO:0006350 transcription | gg | t |
| *rcsB* | CDS | DNA-binding response regulator in two-component regulatory system with RcsC and YojN | Cytoplasmic | COG2197;Response regulator containing a CheY-like receiver domain and an HTH DNA-binding domain | GO:0009242 colanic acid biosynthesis -!- GO:0006350 | o | tt |
| *rcsC* | CDS | hybrid sensory kinase in two-component regulatory system with RcsB and YojN | Integral Membrane Protein | COG0642;Signal transduction histidine kinase -!- COG0784;FOG: CheY-like receiver | GO:0009242 colanic acid biosynthesis -!- GO:0006464 protein | | |
| *rcsC* | CDS | hybrid sensory kinase in two-component regulatory system with RcsB and YojN | Integral Membrane Protein | | | | |
| *rcsC* | CDS | hybrid sensory kinase in two-component regulatory system with RcsB and YojN | Integral Membrane Protein | | | | |
| *rcsD* | CDS | phosphotransfer intermediate protein in two-component regulatory system with RcsBC | Membrane Anchored | COG0642;Signal transduction histidine kinase -!- COG2198;FOG: HPt domain | GO:0006464 protein modification | o | tt |
| *rcsF* | CDS | predicted outer membrane protein, signal | Outer Membrane Lipoprotein | | GO:0009242 colanic acid biosynthesis | gg | o |
| *rdgC* | CDS | DNA-binding protein, non-specific | Cytoplasmic | COG2974;DNA recombination-dependent growth factor C | GO:0006261 DNA dependent DNA replication | o | tt |
| *rdlA* | misc_RNA | regulatory, antisense RNA | | | | | |
| *rdlB* | misc_RNA | regulatory, antisense sRNA | | | | | |
| *rdlC* | misc_RNA | regulatory, antisense sRNA | | | | | |

| | | | | | | | |
|---|---|---|---|---|---|---|---|
| *rdlD* | misc_RNA | regulatory, antisense sRNA | | | | | |
| *recA* | CDS | DNA strand exchange and recombination protein with protease and nuclease activity | Cytoplasmic | COG0468;RecA/RadA recombinase | GO:0006310 DNA recombination -!- GO:0006281 DNA repair -!- GO:0006508 proteolysis and peptidolysis -!- GO:0009432 SOS response | gg | t |
| *recB* | CDS | exonuclease V (RecBCD complex), beta subunit | Cytoplasmic | COG1074;ATP-dependent exoDNAse (exonuclease V) beta subunit (contains helicase and exonuclease domains) | GO:0006308 DNA catabolism -!- GO:0006308 DNA catabolism -!- GO:0006310 DNA recombination | o | tt |
| *recC* | CDS | exonuclease V (RecBCD complex), gamma chain | Cytoplasmic | COG1330;Exonuclease V gamma subunit | GO:0006308 DNA catabolism -!- GO:0006308 DNA catabolism -!- GO:0006310 DNA recombination | o | tt |
| *recD* | CDS | exonuclease V (RecBCD complex), alpha chain | Cytoplasmic | COG0507;ATP-dependent exoDNAse (exonuclease V), alpha subunit - helicase superfamily I member | GO:0006308 DNA catabolism -!- GO:0006308 DNA catabolism -!- GO:0006310 DNA recombination | gg | t |
| *recE* | CDS | Rac prophage; exonuclease VIII, 5' -> 3' specific dsDNA exonuclease | Cytoplasmic | | GO:0006308 DNA catabolism -!- GO:0006281 DNA repair -!- GO:0006308 DNA catabolism | gg | tt |
| *recF* | CDS | gap repair protein | Cytoplasmic | COG1195;Recombinational DNA repair ATPase (RecF pathway) | GO:0006261 DNA dependent DNA replication -!- GO:0006281 DNA repair -!- GO:0006310 DNA | o | tt |
| *recG* | CDS | ATP-dependent DNA helicase | Cytoplasmic | COG1200;RecG-like helicase | GO:0006261 DNA dependent DNA replication | o | tt |
| *recJ* | CDS | ssDNA exonuclease, 5' --> 3'-specific | Cytoplasmic | COG0608;Single-stranded DNA-specific exonuclease | GO:0006308 DNA catabolism -!- GO:0006308 DNA catabolism | g | t |
| *recN* | CDS | recombination and repair protein | Cytoplasmic | | GO:0006310 DNA recombination -!- GO:0006281 DNA repair -!- GO:0009432 SOS response | | |
| *recO* | CDS | gap repair protein | Cytoplasmic | COG1381;Recombinational DNA repair protein (RecF pathway) | GO:0006281 DNA repair -!- GO:0006310 DNA recombination | gg | tt |
| *recQ* | CDS | ATP-dependent DNA helicase | Cytoplasmic | | GO:0006261 DNA dependent DNA replication -!- GO:0006281 DNA repair -!- GO:0006310 DNA | | |
| *recR* | CDS | gap repair protein | Cytoplasmic | COG0353;Recombinational DNA repair protein (RecF pathway) | GO:0006281 DNA repair -!- GO:0006310 DNA recombination | o | tt |
| *recT* | CDS | Rac prophage; recombination and repair protein | Cytoplasmic | COG3723;Recombinational DNA repair protein (RecE pathway) | | gg | tt |
| *recX* | CDS | regulatory protein for RecA | Cytoplasmic | COG2137;Uncharacterized protein conserved in bacteria | | o | tt |
| *relA* | CDS | (p)ppGpp synthetase I/GTP pyrophosphokinase | Cytoplasmic | COG0317;Guanosine polyphosphate pyrophosphohydrolases/synthetases | GO:0015949 nucleobase, nucleoside and nucleotide interconversion | o | t |
| *relB* | CDS | Qin prophage; bifunctional antitoxin of the RelE-RelB toxin-antitoxin system -!- transcriptional repressor | Cytoplasmic | COG3077;DNA-damage-inducible protein J | GO:0006350 transcription | gg | o |
| *relE* | CDS | Qin prophage; toxin of the RelE-RelB toxin-antitoxin system | Cytoplasmic | COG2026;Cytotoxic translational repressor of toxin-antitoxin stability system | GO:0006350 transcription | o | tt |
| *rem* | CDS | Qin prophage; predicted protein | Cytoplasmic | | | o | tt |
| *renD* | CDS | DLP12 prophage; predicted protein | Cytoplasmic | | | gg | o |
| *rep* | CDS | DNA helicase and single-stranded DNA-dependent ATPase | Cytoplasmic | | GO:0006261 DNA dependent DNA replication | | |
| *rfaB* | CDS | UDP-D-galactose:(glucosyl)lipopolysaccharide-1,6-D-galactosyltransferase | Cytoplasmic | COG0438;Glycosyltransferase | | o | tt |
| *rfaC* | CDS | ADP-heptose:LPS heptosyl transferase I | Cytoplasmic | COG0859;ADP-heptose:LPS heptosyltransferase | GO:0009244 lipopolysaccharide core region biosynthesis | o | tt |
| *rfaD* | CDS | ADP-L-glycero-D-mannoheptose-6-epimerase, NAD(P)-binding | Cytoplasmic | COG0451;Nucleoside-diphosphate-sugar epimerases | GO:0009435 nicotinamide adenine dinucleotide biosynthesis | o | o |
| *rfaE* | CDS | fused heptose 7-phosphate kinase -!- heptose 1-phosphate adenyltransferase | Cytoplasmic | COG2870;ADP-heptose synthase, bifunctional sugar kinase/adenylyltransferase | GO:0016052 carbohydrate catabolism -!- GO:0009244 lipopolysaccharide core region | gg | tt |
| *rfaF* | CDS | ADP-heptose:LPS heptosyltransferase II | Cytoplasmic | COG0859;ADP-heptose:LPS heptosyltransferase | GO:0009244 lipopolysaccharide core region biosynthesis | g | t |
| *rfaG* | CDS | glucosyltransferase I | Cytoplasmic | COG0438;Glycosyltransferase | GO:0009244 lipopolysaccharide core region biosynthesis -!- GO:0009432 SOS response | o | tt |
| *rfaH* | CDS | DNA-binding transcriptional antiterminator | Cytoplasmic | COG0250;Transcription antiterminator | GO:0009244 lipopolysaccharide core region biosynthesis -!- GO:0009297 fimbrial biogenesis -!- GO:0006350 transcription | gg | t |
| *rfaI* | CDS | UDP-D-galactose:(glucosyl)lipopolysaccharide-alpha-1,3-D-galactosyltransferase | Cytoplasmic | COG1442;Lipopolysaccharide biosynthesis proteins, LPS:glycosyltransferases | | o | tt |
| *rfaJ* | CDS | UDP-D-glucose:(galactosyl)lipopolysaccharide glucosyltransferase | Cytoplasmic | COG1442;Lipopolysaccharide biosynthesis proteins, LPS:glycosyltransferases | | o | o |
| *rfaK* | CDS | lipopolysaccharide core biosynthesis | Cytoplasmic | COG0859;ADP-heptose:LPS heptosyltransferase | GO:0009244 lipopolysaccharide core region biosynthesis | o | tt |
| *rfaL* | CDS | O-antigen ligase | Integral Membrane Protein | COG3307;Lipid A core - O-antigen ligase and related enzymes | GO:0009244 lipopolysaccharide core region biosynthesis | o | tt |
| *rfaP* | CDS | kinase that phosphorylates core heptose of lipopolysaccharide | Cytoplasmic | | GO:0009244 lipopolysaccharide core region biosynthesis | o | tt |
| *rfaQ* | CDS | lipopolysaccharide core biosynthesis protein | Cytoplasmic | COG0859;ADP-heptose:LPS heptosyltransferase | GO:0009244 lipopolysaccharide core region biosynthesis | gg | t |
| *rfaS* | CDS | lipopolysaccharide core biosynthesis protein | Cytoplasmic | | GO:0009244 lipopolysaccharide core region biosynthesis | g | o |
| *rfaY* | CDS | lipopolysaccharide core biosynthesis protein | Cytoplasmic | | GO:0009244 lipopolysaccharide core region biosynthesis | o | tt |
| *rfaZ* | CDS | lipopolysaccharide core biosynthesis protein | Cytoplasmic | | GO:0009244 lipopolysaccharide core region biosynthesis | g | t |
| *rfbA* | CDS | glucose-1-phosphate thymidylyltransferase | Cytoplasmic | COG1209;dTDP-glucose pyrophosphorylase | GO:0009243 O antigen biosynthesis -!- GO:0019305 dTDP-rhamnose biosynthesis -!- GO:0009103 lipopolysaccharide biosynthesis | gg | t |
| *rfbB* | CDS | dTDP-glucose 4,6 dehydratase, NAD(P)-binding | Cytoplasmic | COG1088;dTDP-D-glucose 4,6-dehydratase | GO:0009243 O antigen biosynthesis -!- GO:0009226 nucleotide-sugar biosynthesis -!- GO:0009435 nicotinamide adenine dinucleotide biosynthesis | gg | t |
| *rfbC* | CDS | dTDP-4-deoxyrhamnose-3,5-epimerase | Cytoplasmic | COG1898;dTDP-4-dehydrorhamnose 3,5-epimerase and related enzymes | GO:0009226 nucleotide-sugar biosynthesis -!- GO:0009243 O antigen biosynthesis -!- GO:0009243 O antigen biosynthesis -!- GO:0009103 lipopolysaccharide biosynthesis -!- GO:0019305 dTDP-rhamnose biosynthesis | g | o |
| *rfbD* | CDS | dTDP-4-dehydrorhamnose reductase subunit, NAD(P)-binding, of dTDP-L-rhamnose synthase | Cytoplasmic | COG1091;dTDP-4-dehydrorhamnose reductase | GO:0009243 O antigen biosynthesis -!- GO:0009226 nucleotide-sugar biosynthesis | gg | o |
| *rfbX* | CDS | predicted polisoprenol-linked O-antigen transporter | Integral Membrane Protein | COG2244;Membrane protein involved in the export of O-antigen and teichoic acid | GO:0009243 O antigen biosynthesis | o | tt |
| *rfe* | CDS | UDP-GlcNAc:undecaprenylphosphate GlcNAc-1-phosphate transferase | Integral Membrane Protein | COG0472;UDP-N-acetylmuramyl pentapeptide phosphotransferase/UDP-N-acetylglucosamine-1-phosp hate transferase | GO:0009246 enterobacterial common antigen biosynthesis | o | tt |
| *rffA* | CDS | TDP-4-oxo-6-deoxy-D-glucose transaminase | Cytoplasmic | COG0399;Predicted pyridoxal phosphate-dependent enzyme apparently involved in regulation of cell wall biogenesis | GO:0009246 enterobacterial common antigen biosynthesis | gg | o |
| *rffC* | CDS | TDP-fucosamine acetyltransferase | Cytoplasmic | | GO:0009244 lipopolysaccharide core region biosynthesis | | |
| *rffD* | CDS | UDP-N-acetyl-D-mannosaminuronic acid dehydrogenase | Periplasmic | | GO:0009246 enterobacterial common antigen biosynthesis | | |
| *rffE* | CDS | UDP-N-acetyl glucosamine-2-epimerase | Cytoplasmic | | GO:0009246 enterobacterial common antigen biosynthesis | | |
| *rffG* | CDS | dTDP-glucose 4,6-dehydratase | Cytoplasmic | | GO:0009243 O antigen biosynthesis -!- GO:0009246 enterobacterial common antigen biosynthesis -!- GO:0009435 nicotinamide adenine dinucleotide biosynthesis | | |
| *rffH* | CDS | glucose-1-phosphate thymidylyltransferase | Cytoplasmic | COG1209;dTDP-glucose pyrophosphorylase | GO:0009243 O antigen biosynthesis -!- GO:0009246 enterobacterial common antigen biosynthesis | gg | t |
| *rffM* | CDS | UDP-N-acetyl-D-mannosaminuronic acid transferase | Cytoplasmic | COG1922;Teichoic acid biosynthesis proteins | GO:0009246 enterobacterial common antigen biosynthesis | gg | tt |
| *rffT* | CDS | TDP-Fuc4NAc:lipidII transferase | Cytoplasmic | | | | |
| *rhaA* | CDS | L-rhamnose isomerase | Cytoplasmic | | GO:0016052 carbohydrate | | |
| *rhaB* | CDS | rhamnulokinase | Cytoplasmic | COG1070;Sugar (pentulose and hexulose) kinases | GO:0016052 carbohydrate | o | tt |
| *rhaD* | CDS | rhamnulose-1-phosphate aldolase | Cytoplasmic | COG0235;Ribulose-5-phosphate 4-epimerase and related epimerases and aldolases | GO:0016052 carbohydrate catabolism | gg | o |
| *rhaR* | CDS | DNA-binding transcriptional activator, L-rhamnose-binding | Cytoplasmic | COG2207;AraC-type DNA-binding domain-containing proteins | GO:0016052 carbohydrate catabolism -!- GO:0006350 | g | t |
| *rhaS* | CDS | DNA-binding transcriptional activator, L-rhamnose-binding | Cytoplasmic | COG2207;AraC-type DNA-binding domain-containing proteins | GO:0016052 carbohydrate catabolism -!- GO:0006350 | gg | o |
| *rhaT* | CDS | L-rhamnose:proton symporter | Integral Membrane Protein | COG0697;Permeases of the drug/metabolite transporter (DMT) superfamily | GO:0016052 carbohydrate | o | tt |
| *rhlB* | CDS | ATP-dependent RNA helicase | Cytoplasmic | COG0513;Superfamily II DNA and RNA helicases | | g | t |
| *rhlE* | CDS | RNA helicase | Cytoplasmic | COG0513;Superfamily II DNA and RNA helicases | GO:0009451 RNA modification | o | tt |
| *rho* | CDS | transcription termination factor | Cytoplasmic | COG1158;Transcription termination factor | GO:0006350 transcription | o | tt |
| *rhoL* | CDS | rho operon leader peptide | Cytoplasmic | | GO:0006350 transcription | gg | o |
| *rhsA* | CDS | rhsA element core protein RshA | Cytoplasmic | COG3209;Rhs family protein | | o | tt |
| *rhsB* | CDS | rhsB element core protein RshB | Cytoplasmic | | | | |
| *rhsC* | CDS | rhsC element core protein RshC | Cytoplasmic | COG3209;Rhs family protein | | o | t |

| | | | | | | | |
|---|---|---|---|---|---|---|---|
| *rhsD* | CDS | rhsD element protein | Cytoplasmic | COG3209;Rhs family protein | | o | t |
| *rhsE* | CDS | rhsE element core protein RshE | Cytoplasmic | COG3209;Rhs family protein | | o | tt |
| *rhtA* | CDS | threonine and homoserine efflux system | Integral Membrane Protein | COG5006;Predicted permease, DMT superfamily | | o | tt |
| *rhtB* | CDS | neutral amino-acid efflux system | Integral Membrane Protein | | GO:0006564 L-serine biosynthesis | | |
| *rhtC* | CDS | threonine efflux system | Integral Membrane Protein | | GO:0009088 threonine biosynthesis | | |
| *ribA* | CDS | GTP cyclohydrolase II | Cytoplasmic | COG0807;GTP cyclohydrolase II | | gg | t |
| *ribB* | CDS | 3,4 dihydroxy-2-butanone-4-phosphate synthase | Cytoplasmic | COG0108;3,4-dihydroxy-2-butanone 4-phosphate synthase | | gg | t |
| *ribC* | CDS | riboflavin synthase, alpha subunit | Cytoplasmic | COG0307;Riboflavin synthase alpha chain | | gg | t |
| *ribD* | CDS | fused diaminohydroxyphosphoribosylaminopyrimidine deaminase -!- 5-amino-6-(5-phosphoribosylamino) uracil reductase | Cytoplasmic | COG0117;Pyrimidine deaminase -!- COG1985;Pyrimidine reductase, riboflavin biosynthesis | | gg | t |
| *ribE* | CDS | riboflavin synthase beta chain | Cytoplasmic | COG0054;Riboflavin synthase beta-chain | | o | o |
| *ribF* | CDS | bifunctional riboflavin kinase -!- FAD synthetase | Cytoplasmic | COG0196;FAD synthase | | o | tt |
| *rihA* | CDS | ribonucleoside hydrolase 1 | Cytoplasmic | COG1957;Inosine-uridine nucleoside N-ribohydrolase | GO:0009226 nucleotide-sugar biosynthesis | gg | t |
| *rihB* | CDS | ribonucleoside hydrolase 2 | Cytoplasmic | COG1957;Inosine-uridine nucleoside N-ribohydrolase | | g | t |
| *rihC* | CDS | ribonucleoside hydrolase 3 | Cytoplasmic | COG1957;Inosine-uridine nucleoside N-ribohydrolase | GO:0009226 nucleotide-sugar biosynthesis | g | o |
| *rimI* | CDS | acetylase for 30S ribosomal subunit protein S18 | Cytoplasmic | COG0456;Acetyltransferases | GO:0006464 protein modification | gg | o |
| *rimJ* | CDS | ribosomal-protein-S5-alanine N-acetyltransferase | Cytoplasmic | COG1670;Acetyltransferases, including N-acetylases of ribosomal proteins | GO:0006464 protein modification | gg | o |
| *rimK* | CDS | ribosomal protein S6 modification protein | Cytoplasmic | COG0189;Glutathione synthase/Ribosomal protein S6 modification enzyme (glutaminyl transferase) | GO:0006412 protein biosynthesis | gg | t |
| *rimL* | CDS | ribosomal-protein-L7/L12-serine acetyltransferase | Cytoplasmic | COG1670;Acetyltransferases, including N-acetylases of ribosomal proteins | GO:0006464 protein modification | gg | o |
| *rimM* | CDS | 16S rRNA processing protein | Cytoplasmic | | GO:0009451 RNA modification | | |
| *rlmB* | CDS | 23S rRNA (Gm2251)-methyltransferase | Cytoplasmic | COG0566;rRNA methylases | GO:0009451 RNA modification | gg | t |
| *rlpA* | CDS | minor lipoprotein | Outer Membrane Lipoprotein | COG0797;Lipoproteins | | o | tt |
| *rlpB* | CDS | minor lipoprotein | Membrane Lipoprotein | COG2980;Rare lipoprotein B | GO:0042158 lipoprotein biosynthesis | o | tt |
| *rluA* | CDS | pseudouridine synthase for 23S rRNA (position 746) and tRNAphe(position 32) | Cytoplasmic | COG0564;Pseudouridylate synthases, 23S RNA-specific | GO:0009451 RNA modification | gg | t |
| *rluB* | CDS | 23S rRNA pseudouridylate synthase | Cytoplasmic | COG1187;16S rRNA uridine-516 pseudouridylate synthase and related pseudouridylate synthases | GO:0009451 RNA modification | gg | t |
| *rluC* | CDS | 23S rRNA pseudouridylate synthase | Cytoplasmic | COG0564;Pseudouridylate synthases, 23S RNA-specific | GO:0009451 RNA modification | g | tt |
| *rluD* | CDS | 23S rRNA pseudouridine synthase | Cytoplasmic | COG0564;Pseudouridylate synthases, 23S RNA-specific | GO:0009451 RNA modification | | |
| *rmf* | CDS | ribosome modulation factor | Cytoplasmic | COG3130;Ribosome modulation factor | GO:0006457 protein folding | gg | t |
| *rmuC* | CDS | predicted recombination limiting protein | Cytoplasmic | COG1322;Uncharacterized protein conserved in bacteria | | o | tt |
| *rna* | CDS | ribonuclease I | Periplasmic | COG3719;Ribonuclease I | GO:0006401 RNA catabolism -!- GO:0006401 RNA catabolism | o | tt |
| *rnb* | CDS | ribonuclease II | Cytoplasmic | COG4776;Exoribonuclease II | GO:0006401 RNA catabolism -!- GO:0006401 RNA catabolism | gg | o |
| *rnc* | CDS | RNase III | Cytoplasmic | COG0571;dsRNA-specific ribonuclease | GO:0006401 RNA catabolism -!- GO:0006401 RNA catabolism | gg | tt |
| *rnd* | CDS | ribonuclease D | Cytoplasmic | COG0349;Ribonuclease D | GO:0006401 RNA catabolism -!- GO:0009451 RNA modification | gg | tt |
| *rne* | CDS | fused ribonucleaseE endoribonuclease -!- scaffold for formation of degradosome | Cytoplasmic | COG1530;Ribonucleases G and E | GO:0006401 RNA catabolism -!- GO:0006401 RNA catabolism | o | tt |
| *rng* | CDS | ribonuclease G | Cytoplasmic | COG1530;Ribonucleases G and E | GO:0009451 RNA modification | g | o |
| *rnhA* | CDS | ribonuclease HI, degrades RNA of DNA-RNA hybrids | Cytoplasmic | COG0328;Ribonuclease HI | GO:0006401 RNA catabolism -!- GO:0006261 DNA dependent DNA replication -!- GO:0006508 proteolysis and peptidolysis | o | tt |
| *rnhB* | CDS | ribonuclease HII, degrades RNA of DNA-RNA hybrids | Cytoplasmic | COG0164;Ribonuclease HII | GO:0006401 RNA catabolism -!- GO:0006401 RNA catabolism | gg | o |
| *rnk* | CDS | regulator of nucleoside diphosphate kinase | Cytoplasmic | COG0782;Transcription elongation factor | GO:0015949 nucleobase, nucleoside and nucleotide interconversion | gg | t |
| *rnpA* | CDS | protein C5 component of RNase P | Cytoplasmic | COG0594;RNase P protein component | GO:0006401 RNA catabolism -!- GO:0009451 RNA modification | g | tt |
| *rnpB* | misc_RNA | RNase P, RNA component precursor RnpB | | | | | |
| *rnr* | CDS | exoribonuclease R, RNase R | Cytoplasmic | | GO:0006401 RNA catabolism -!- GO:0006401 RNA catabolism | | |
| *rnt* | CDS | ribonuclease T (RNase T) | Cytoplasmic | COG0847;DNA polymerase III, epsilon subunit and related 3'-5' exonucleases | GO:0006401 RNA catabolism -!- GO:0006308 DNA catabolism -!- GO:0006401 RNA catabolism -!- GO:0006308 DNA catabolism | o | tt |
| *rob* | CDS | DNA-binding transcriptional activator | Cytoplasmic | COG2207;AraC-type DNA-binding domain-containing proteins | GO:0006261 DNA dependent DNA replication -!- GO:0006350 transcription | o | tt |
| *rof* | CDS | modulator of Rho-dependent transcription termination | Cytoplasmic | COG4568;Transcriptional antiterminator | GO:0006350 transcription | o | tt |
| *rpe* | CDS | D-ribulose-5-phosphate 3-epimerase | Cytoplasmic | COG0036;Pentose-5-phosphate-3-epimerase | GO:0016052 carbohydrate catabolism -!- GO:0009052 pentose-phosphate shunt, non-oxidative | gg | o |
| *rph* | CDS | ribonuclease PH (pseudogene) | Cytoplasmic | COG0689;RNase PH | GO:0006401 RNA catabolism -!- GO:0006401 RNA catabolism | g | o |
| *rpiA* | CDS | ribosephosphate isomerase, constitutive | Cytoplasmic | COG0120;Ribose 5-phosphate isomerase | GO:0016052 carbohydrate catabolism -!- GO:0009052 pentose-phosphate shunt, non-oxidative | | |
| *rpiB* | CDS | ribose 5-phosphate isomerase B/allose 6-phosphate isomerase | Cytoplasmic | COG0698;Ribose 5-phosphate isomerase RpiB | GO:0016052 carbohydrate catabolism -!- GO:0009052 pentose-phosphate shunt, non-oxidative | g | o |
| *rpiR* | CDS | DNA-binding transcriptional repressor | Cytoplasmic | | GO:0016052 carbohydrate catabolism -!- GO:0009052 pentose-phosphate shunt, non-oxidative branch -!- GO:0006350 transcription | gg | t |
| *rplA* | CDS | 50S ribosomal subunit protein L1 | Cytoplasmic | COG0081;Ribosomal protein L1 | GO:0006412 protein biosynthesis -!- GO:0009386 translational attenuation | o | tt |
| *rplB* | CDS | 50S ribosomal subunit protein L2 | Cytoplasmic | COG0090;Ribosomal protein L2 | GO:0006412 protein biosynthesis | o | tt |
| *rplC* | CDS | 50S ribosomal subunit protein L3 | Cytoplasmic | COG0087;Ribosomal protein L3 | GO:0006412 protein biosynthesis | gg | tt |
| *rplD* | CDS | 50S ribosomal subunit protein L4 | Cytoplasmic | COG0088;Ribosomal protein L4 | GO:0006412 protein biosynthesis -!- GO:0006355 regulation of transcription, DNA-dependent -!- GO:0009386 translational attenuation | o | tt |
| *rplE* | CDS | 50S ribosomal subunit protein L5 | Cytoplasmic | COG0094;Ribosomal protein L5 | GO:0006412 protein biosynthesis | gg | t |
| *rplF* | CDS | 50S ribosomal subunit protein L6 | Cytoplasmic | COG0097;Ribosomal protein L6P/L9E | GO:0006412 protein biosynthesis | gg | tt |
| *rplI* | CDS | 50S ribosomal subunit protein L9 | Cytoplasmic | COG0359;Ribosomal protein L9 | GO:0006412 protein biosynthesis | o | o |
| *rplJ* | CDS | 50S ribosomal subunit protein L10 | Cytoplasmic | COG0244;Ribosomal protein L10 | GO:0006412 protein biosynthesis | o | tt |
| *rplK* | CDS | 50S ribosomal subunit protein L11 | Cytoplasmic | COG0080;Ribosomal protein L11 | GO:0006412 protein biosynthesis | gg | t |
| *rplL* | CDS | 50S ribosomal subunit protein L7/L12 | Cytoplasmic | COG0222;Ribosomal protein L7/L12 | GO:0006412 protein biosynthesis | o | o |
| *rplM* | CDS | 50S ribosomal subunit protein L13 | Cytoplasmic | COG0102;Ribosomal protein L13 | GO:0006412 protein biosynthesis | gg | tt |
| *rplN* | CDS | 50S ribosomal subunit protein L14 | Cytoplasmic | COG0093;Ribosomal protein L14 | GO:0006412 protein biosynthesis | o | tt |
| *rplO* | CDS | 50S ribosomal subunit protein L15 | Cytoplasmic | COG0200;Ribosomal protein L15 | GO:0006412 protein biosynthesis | g | tt |
| *rplP* | CDS | 50S ribosomal subunit protein L16 | Cytoplasmic | COG0197;Ribosomal protein L16/L10E | GO:0006412 protein biosynthesis | o | tt |
| *rplQ* | CDS | 50S ribosomal subunit protein L17 | Cytoplasmic | COG0203;Ribosomal protein L17 | GO:0006412 protein biosynthesis | o | tt |
| *rplR* | CDS | 50S ribosomal subunit protein L18 | Cytoplasmic | COG0256;Ribosomal protein L18 | GO:0006412 protein biosynthesis | gg | o |
| *rplS* | CDS | 50S ribosomal subunit protein L19 | Cytoplasmic | COG0335;Ribosomal protein L19 | GO:0006412 protein biosynthesis | o | tt |
| *rplT* | CDS | 50S ribosomal subunit protein L20 | Cytoplasmic | COG0292;Ribosomal protein L20 | GO:0006412 protein biosynthesis | gg | t |
| *rplU* | CDS | 50S ribosomal subunit protein L21 | Cytoplasmic | COG0261;Ribosomal protein L21 | GO:0006412 protein biosynthesis | g | o |
| *rplV* | CDS | 50S ribosomal subunit protein L22 | Cytoplasmic | COG0091;Ribosomal protein L22 | GO:0006412 protein biosynthesis | g | tt |
| *rplW* | CDS | 50S ribosomal subunit protein L23 | Cytoplasmic | COG0089;Ribosomal protein L23 | GO:0006412 protein biosynthesis | gg | tt |
| *rplX* | CDS | 50S ribosomal subunit protein L24 | Cytoplasmic | COG0198;Ribosomal protein L24 | GO:0006412 protein biosynthesis | gg | t |
| *rplY* | CDS | 50S ribosomal subunit protein L25 | Cytoplasmic | COG1825;Ribosomal protein L25 (general stress protein Ctc) | GO:0006412 protein biosynthesis | gg | o |
| *rpmA* | CDS | 50S ribosomal subunit protein L27 | Cytoplasmic | COG0211;Ribosomal protein L27 | GO:0006412 protein biosynthesis | gg | tt |
| *rpmB* | CDS | 50S ribosomal subunit protein L28 | Cytoplasmic | COG0227;Ribosomal protein L28 | GO:0006412 protein biosynthesis | gg | tt |
| *rpmC* | CDS | 50S ribosomal subunit protein L29 | Cytoplasmic | COG0255;Ribosomal protein L29 | GO:0006412 protein biosynthesis | gg | o |
| *rpmD* | CDS | 50S ribosomal subunit protein L30 | Cytoplasmic | COG1841;Ribosomal protein L30/L7E | GO:0006412 protein biosynthesis | gg | t |
| *rpmE* | CDS | 50S ribosomal subunit protein L31 | Cytoplasmic | COG0254;Ribosomal protein L31 | GO:0006412 protein biosynthesis | gg | o |
| *rpmF* | CDS | 50S ribosomal subunit protein L32 | Cytoplasmic | COG0333;Ribosomal protein L32 | GO:0006412 protein biosynthesis | gg | tt |
| *rpmG* | CDS | 50S ribosomal subunit protein L33 | Cytoplasmic | COG0267;Ribosomal protein L33 | GO:0006412 protein biosynthesis | gg | tt |
| *rpmH* | CDS | 50S ribosomal subunit protein L34 | Cytoplasmic | COG0230;Ribosomal protein L34 | GO:0006412 protein biosynthesis | gg | tt |
| *rpmI* | CDS | 50S ribosomal subunit protein L35 | Cytoplasmic | | GO:0006412 protein biosynthesis | o | t |
| *rpmJ* | CDS | 50S ribosomal subunit protein L36 | Cytoplasmic | COG0257;Ribosomal protein L36 | GO:0006412 protein biosynthesis | gg | o |
| *rpoA* | CDS | RNA polymerase, alpha subunit | Cytoplasmic | COG0202;DNA-directed RNA polymerase, alpha subunit/40 kD subunit | GO:0006350 transcription | gg | t |
| *rpoB* | CDS | RNA polymerase, beta subunit | Cytoplasmic | COG0085;DNA-directed RNA polymerase, beta subunit/140 kD subunit | GO:0006350 transcription | g | tt |
| *rpoC* | CDS | RNA polymerase, beta prime subunit | Cytoplasmic | COG0086;DNA-directed RNA polymerase, beta' subunit/160 kD subunit | GO:0006350 transcription | o | tt |
| *rpoD* | CDS | RNA polymerase, sigma 70 (sigma D) factor | Cytoplasmic | COG0568;DNA-directed RNA polymerase, sigma subunit (sigma70/sigma32) | GO:0006350 transcription | o | tt |
| *rpoE* | CDS | RNA polymerase, sigma 24 (sigma E) factor | Cytoplasmic | COG1595;DNA-directed RNA polymerase specialized sigma subunit, sigma24 homolog | GO:0006350 transcription -!- GO:0009266 response to | gg | t |
| *rpoH* | CDS | RNA polymerase, sigma 32 (sigma H) factor | Cytoplasmic | COG0568;DNA-directed RNA polymerase, sigma subunit (sigma70/sigma32) | GO:0006350 transcription -!- GO:0009266 response to | o | tt |
| *rpoN* | CDS | RNA polymerase, sigma 54 (sigma N) factor | Cytoplasmic | COG1508;DNA-directed RNA polymerase specialized sigma subunit, sigma54 homolog | GO:0006807 nitrogen metabolism -!- GO:0006350 transcription | o | tt |
| *rpoS* | CDS | RNA polymerase, sigma S (sigma 38) factor | Cytoplasmic | COG0568;DNA-directed RNA polymerase, sigma subunit (sigma70/sigma32) | GO:0006350 transcription -!- GO:0006970 response to osmotic stress | | |
| *rpoZ* | CDS | RNA polymerase, omega subunit | Cytoplasmic | COG1758;DNA-directed RNA polymerase, subunit K/omega | GO:0006350 transcription | gg | o |
| *rprA* | misc_RNA | regulatory sRNA | | | | | |
| *rpsA* | CDS | 30S ribosomal subunit protein S1 | Cytoplasmic | COG0539;Ribosomal protein S1 | GO:0006412 protein biosynthesis | o | tt |
| *rpsB* | CDS | 30S ribosomal subunit protein S2 | Cytoplasmic | COG0052;Ribosomal protein S2 | GO:0006412 protein biosynthesis | o | tt |
| *rpsC* | CDS | 30S ribosomal subunit protein S3 | Cytoplasmic | COG0092;Ribosomal protein S3 | GO:0006412 protein biosynthesis | o | tt |

| | | | | | | | |
|---|---|---|---|---|---|---|---|
| *rpsD* | CDS | 30S ribosomal subunit protein S4 | Cytoplasmic | COG0522;Ribosomal protein S4 and related proteins | GO:0006412 protein biosynthesis | o | tt |
| *rpsE* | CDS | 30S ribosomal subunit protein S5 | Cytoplasmic | COG0098;Ribosomal protein S5 | GO:0006412 protein biosynthesis | gg | o |
| *rpsF* | CDS | 30S ribosomal subunit protein S6 | Cytoplasmic | COG0360;Ribosomal protein S6 | GO:0006412 protein biosynthesis | o | tt |
| *rpsG* | CDS | 30S ribosomal subunit protein S7 | Cytoplasmic | COG0049;Ribosomal protein S7 | GO:0006412 protein biosynthesis | o | tt |
| *rpsH* | CDS | 30S ribosomal subunit protein S8 | Cytoplasmic | COG0096;Ribosomal protein S8 | GO:0006412 protein biosynthesis -!- GO:0009386 translational attenuation | o | tt |
| *rpsI* | CDS | 30S ribosomal subunit protein S9 | Cytoplasmic | COG0103;Ribosomal protein S9 | GO:0006412 protein biosynthesis | gg | tt |
| *rpsJ* | CDS | 30S ribosomal subunit protein S10 | Cytoplasmic | COG0051;Ribosomal protein S10 | GO:0006412 protein biosynthesis | o | o |
| *rpsK* | CDS | 30S ribosomal subunit protein S11 | Cytoplasmic | COG0100;Ribosomal protein S11 | GO:0006412 protein biosynthesis | gg | t |
| *rpsL* | CDS | 30S ribosomal subunit protein S12 | Cytoplasmic | COG0048;Ribosomal protein S12 | GO:0006412 protein biosynthesis | gg | tt |
| *rpsM* | CDS | 30S ribosomal subunit protein S13 | Cytoplasmic | COG0099;Ribosomal protein S13 | GO:0006412 protein biosynthesis | o | tt |
| *rpsN* | CDS | 30S ribosomal subunit protein S14 | Cytoplasmic | COG0199;Ribosomal protein S14 | GO:0006412 protein biosynthesis | gg | tt |
| *rpsO* | CDS | 30S ribosomal subunit protein S15 | Cytoplasmic | COG0184;Ribosomal protein S15P/S13E | GO:0006412 protein biosynthesis | gg | t |
| *rpsP* | CDS | 30S ribosomal subunit protein S16 | Cytoplasmic | COG0228;Ribosomal protein S16 | GO:0006412 protein biosynthesis | gg | t |
| *rpsQ* | CDS | 30S ribosomal subunit protein S17 | Cytoplasmic | COG0186;Ribosomal protein S17 | GO:0006412 protein biosynthesis | gg | t |
| *rpsR* | CDS | 30S ribosomal subunit protein S18 | Cytoplasmic | COG0238;Ribosomal protein S18 | GO:0006412 protein biosynthesis | o | tt |
| *rpsS* | CDS | 30S ribosomal subunit protein S19 | Cytoplasmic | COG0185;Ribosomal protein S19 | GO:0006412 protein biosynthesis | o | tt |
| *rpsT* | CDS | 30S ribosomal subunit protein S20 | Cytoplasmic | COG0268;Ribosomal protein S20 | GO:0006412 protein biosynthesis | gg | t |
| *rpsU* | CDS | 30S ribosomal subunit protein S21 | Cytoplasmic | COG0828;Ribosomal protein S21 | GO:0006412 protein biosynthesis | gg | t |
| *rraA* | CDS | ribonuclease E (RNase E) inhibitor protein | Cytoplasmic | COG0684;Demethylmenaquinone methyltransferase | | o | tt |
| *rrfA* | rRNA | 5S rRNA (rrfA) | | | | | |
| *rrfB* | rRNA | 5S rRNA (rrfB) | | | | | |
| *rrfC* | rRNA | 5S rRNA (rrfC) | | | | | |
| *rrfD* | rRNA | 5S rRNA (rrfD) | | | | | |
| *rrfE* | rRNA | 5S rRNA (rrfE) | | | | | |
| *rrfF* | rRNA | 5S rRNA (rrfF) | | | | | |
| *rrfG* | rRNA | 5S rRNA (rrfG) | | | | | |
| *rrfH* | rRNA | 5S rRNA (rrfH) | | | | | |
| *rrlA* | rRNA | 23S rRNA (rrlA) | | | | | |
| *rrlB* | rRNA | 23S rRNA (rrlB) | | | | | |
| *rrlC* | rRNA | 23S rRNA (rrlC) | | | | | |
| *rrlD* | rRNA | 23S rRNA (rrlD) | | | | | |
| *rrlE* | rRNA | 23S rRNA (rrlE) | | | | | |
| *rrlG* | rRNA | 23S rRNA (rrlG) | | | | | |
| *rrlH* | rRNA | 23S rRNA (rrlH) | | | | | |
| *rrmA* | CDS | 23S rRNA m1G745 methyltransferase | Cytoplasmic | COG0500;SAM-dependent methyltransferases | GO:0009451 RNA modification | gg | t |
| *rrmJ* | CDS | 23S rRNA methyltransferase | Cytoplasmic | COG0293;23S rRNA methylase | | gg | tt |
| *rrsA* | rRNA | 16S rRNA (rrnA) | | | | | |
| *rrsB* | rRNA | 16S rRNA (rrnB) | | | | | |
| *rrsC* | rRNA | 16S rRNA (rrnC) | | | | | |
| *rrsD* | rRNA | 16S rRNA (rrnD) | | | | | |
| *rrsE* | rRNA | 16S rRNA (rrnE) | | | | | |
| *rrsG* | rRNA | 16S rRNA (rrnG) | | | | | |
| *rrsH* | rRNA | 16S rRNA (rrsH) | | | | | |
| *rsd* | CDS | stationary phase protein, binds sigma 70 RNA polymerase subunit | Cytoplasmic | COG3160;Regulator of sigma D | GO:0006350 transcription | gg | tt |
| *rseA* | CDS | anti-sigma factor | Integral Membrane Protein | COG3073;Negative regulator of sigma E activity | GO:0006350 transcription -!- GO:0009266 response to | o | tt |
| *rseB* | CDS | anti-sigma factor | Periplasmic | COG3026;Negative regulator of sigma E activity | GO:0006350 transcription -!- GO:0042594 response to starvation | o | tt |
| *rseC* | CDS | RseC protein involved in reduction of the SoxR iron-sulfur cluster | Integral Membrane Protein | COG3086;Positive regulator of sigma E activity | GO:0006350 transcription -!- GO:0042594 response to starvation | o | tt |
| *rsgA* | CDS | ribosome small subunit-dependent GTPase A | Cytoplasmic | | | gg | t |
| *rsmB* | CDS | 16S rRNA m5C967 methyltransferase, S-adenosyl-L-methionine-dependent | Cytoplasmic | COG0144;tRNA and rRNA cytosine-C5-methylases | GO:0009451 RNA modification | o | tt |
| *rsmC* | CDS | 16S RNA m2G1207 methylase | Cytoplasmic | COG2813;16S RNA G1207 methylase RsmC | GO:0009451 RNA modification | gg | t |
| *rspA* | CDS | predicted dehydratase | Cytoplasmic | COG4948;L-alanine-DL-glutamate epimerase and related enzymes of enolase superfamily | GO:0042594 response to starvation | gg | t |
| *rspB* | CDS | predicted oxidoreductase, Zn-dependent and NAD(P)-binding | Cytoplasmic | COG1063;Threonine dehydrogenase and related Zn-dependent dehydrogenases | GO:0042594 response to starvation | gg | o |
| *rssA* | CDS | conserved protein | Cytoplasmic | COG1752;Predicted esterase of the alpha-beta hydrolase superfamily | | gg | t |
| *rssB* | CDS | response regulator of RpoS | Cytoplasmic | COG0784;FOG: CheY-like receiver | GO:0006350 transcription | gg | t |
| *rstA* | CDS | DNA-bindng response regulator in two-component regulatory system with RstB | Cytoplasmic | COG0745;Response regulators consisting of a CheY-like receiver domain and a winged-helix DNA-binding domain | GO:0006350 transcription | gg | t |
| *rstB* | CDS | sensory histidine kinase in two-component regulatory system with RstA | Integral Membrane Protein | COG0642;Signal transduction histidine kinase | GO:0006464 protein modification | o | tt |
| *rsuA* | CDS | 16S rRNA pseudouridylate 516 synthase | Cytoplasmic | COG1187;16S rRNA uridine-516 pseudouridylate synthase and related pseudouridylate synthases | GO:0009451 RNA modification | g | o |
| *rsxA* | CDS | predicted inner membrane subunit | Integral Membrane Protein | COG4657;Predicted NADH:ubiquinone oxidoreductase, subunit RnfA | | o | tt |
| *rsxB* | CDS | predicted iron-sulfur protein | Periplasmic | COG2878;Predicted NADH:ubiquinone oxidoreductase, subunit RnfB | | o | tt |
| *rsxC* | CDS | fused predicted 4Fe-4S ferredoxin-type protein | Cytoplasmic | COG4656;Predicted NADH:ubiquinone oxidoreductase, subunit RnfC | | o | tt |
| *rsxD* | CDS | predicted inner membrane oxidoreductase | Integral Membrane Protein | COG4658;Predicted NADH:ubiquinone oxidoreductase, subunit RnfD | | o | tt |
| *rsxE* | CDS | predicted inner membrane NADH-quinone reductase | Integral Membrane Protein | COG4660;Predicted NADH:ubiquinone oxidoreductase, subunit RnfE | | o | tt |
| *rsxG* | CDS | predicted oxidoreductase | Periplasmic | COG4659;Predicted NADH:ubiquinone oxidoreductase, subunit RnfG | | o | tt |
| *rtcA* | CDS | RNA 3'-terminal phosphate cyclase | Cytoplasmic | | GO:0016070 RNA metabolism | | |
| *rtcB* | CDS | conserved protein | Cytoplasmic | COG1690;Uncharacterized conserved protein | | gg | t |
| *rtcR* | CDS | sigma 54-dependent transcriptional regulator of rtcBA expression | Cytoplasmic | COG4650;Sigma54-dependent transcription regulator containing an AAA-type ATPase domain and a DNA-binding domain | GO:0006350 transcription -!- GO:0006355 regulation of transcription, DNA-dependent | g | t |
| *rtn* | CDS | conserved protein | Membrane Anchored | COG2200;FOG: EAL domain | | g | t |
| *rttR* | misc_RNA | regulatory sRNA | | | | | |
| *rumA* | CDS | 23S rRNA (uracil-5)-methyltransferase | Cytoplasmic | COG2265;SAM-dependent methyltransferases related to tRNA (uracil-5-)-methyltransferase | GO:0009451 RNA modification | o | o |
| *rumB* | CDS | 23S rRNA m(5)U747 methyltransferase | Cytoplasmic | COG2265;SAM-dependent methyltransferases related to tRNA (uracil-5-)-methyltransferase | GO:0009451 RNA modification | gg | t |
| *rusA* | CDS | DLP12 prophage; endonuclease RUS | Cytoplasmic | COG4570;Holliday junction resolvase | GO:0006308 DNA catabolism -!- GO:0006281 DNA repair -!- GO:0006308 DNA catabolism | o | tt |
| *ruvA* | CDS | component of RuvABC resolvasome, regulatory subunit | Cytoplasmic | COG0632;Holliday junction resolvasome, DNA-binding subunit | GO:0006281 DNA repair -!- GO:0006310 DNA recombination -!- GO:0009432 SOS response | g | tt |
| *ruvB* | CDS | ATP-dependent DNA helicase, component of RuvABC resolvasome | Cytoplasmic | COG2255;Holliday junction resolvasome, helicase subunit | GO:0006281 DNA repair -!- GO:0006310 DNA recombination -!- GO:0009432 SOS response | gg | t |
| *ruvC* | CDS | component of RuvABC resolvasome, endonuclease | Cytoplasmic | COG0817;Holliday junction resolvasome, endonuclease subunit | GO:0006281 DNA repair -!- GO:0006310 DNA recombination | g | tt |
| *rybA* | misc_RNA | small RNA | | | | | |
| *rybB* | misc_RNA | small RNA | | | | | |
| *rydB* | misc_RNA | regulatory sRNA | | | | | |
| *ryeA* | misc_RNA | small RNA | | | | | |
| *ryeB* | misc_RNA | small RNA | | | | | |
| *ryeC* | misc_RNA | small RNA | | | | | |
| *ryeD* | misc_RNA | small RNA | | | | | |
| *ryeE* | misc_RNA | small RNA | | | | | |
| *ryfA* | misc_RNA | small RNA | | | | | |
| *rygA* | misc_RNA | small RNA | | | | | |
| *rygB* | misc_RNA | small RNA | | | | | |
| *rygC* | misc_RNA | small RNA | | | | | |
| *rygD* | misc_RNA | small RNA | | | | | |
| *ryhA* | misc_RNA | small RNA | | | | | |
| *ryhB* | misc_RNA | regulatory sRNA | | | | | |
| *ryiA* | misc_RNA | regulatory sRNA | | | | | |
| *ryjA* | misc_RNA | small RNA | | | | | |
| *rzoD* | CDS | DLP12 prophage; predicted lipoprotein | Outer Membrane Lipoprotein | | | | |
| *rzoR* | CDS | Rac prophage; predicted lipoprotein | Outer Membrane Lipoprotein | | | | |
| *rzpD* | CDS | DLP12 prophage; predicted murein endopeptidase | Periplasmic | | | | |
| *rzpR* | CDS | Rac prophage; predicted defective peptidase | Cytoplasmic | | | | |
| *sanA* | CDS | predicted protein | Periplasmic | COG2949;Uncharacterized membrane protein | GO:0042493 response to drug | o | tt |
| *sapA* | CDS | predicted antimicrobial peptide transporter subunit -!- periplasmic-binding component of ABC superfamily | Periplasmic | COG4166;ABC-type oligopeptide transport system, periplasmic component | GO:0009063 amino acid catabolism | o | tt |

| | | | | | | | |
|---|---|---|---|---|---|---|---|
| *sapB* | CDS | predicted antimicrobial peptide transporter subunit -!- membrane component of ABC superfamily | Integral Membrane Protein | COG4168;ABC-type antimicrobial peptide transport system, permease component | | gg | tt |
| *sapC* | CDS | predicted antimicrobial peptide transporter subunit -!- membrane component of ABC superfamily | Integral Membrane Protein | COG4171;ABC-type antimicrobial peptide transport system, permease component | | o | tt |
| *sapD* | CDS | predicted antimicrobial peptide transporter subunit -!- ATP-binding component of ABC superfamily | Cytoplasmic | COG4170;ABC-type antimicrobial peptide transport system, ATPase component | GO:0009063 amino acid catabolism | o | tt |
| *sapF* | CDS | predicted antimicrobial peptide transporter subunit -!- ATP-binding component of ABC superfamily | Cytoplasmic | COG4167;ABC-type antimicrobial peptide transport system, ATPase component | GO:0009063 amino acid catabolism | gg | t |
| *sbcB* | CDS | exonuclease I | Cytoplasmic | COG2925;Exonuclease I | GO:0006308 DNA catabolism -!- GO:0006308 DNA catabolism | gg | tt |
| *sbcC* | CDS | exonuclease, dsDNA, ATP-dependent | Cytoplasmic | COG0419;ATPase involved in DNA repair | GO:0006308 DNA catabolism -!- GO:0006308 DNA catabolism | o | tt |
| *sbcD* | CDS | exonuclease, dsDNA, ATP-dependent | Cytoplasmic | COG0420;DNA repair exonuclease | GO:0006308 DNA catabolism -!- GO:0006308 DNA catabolism | gg | t |
| *sbmA* | CDS | predicted transporter | Integral Membrane Protein | COG1133;ABC-type long-chain fatty acid transport system, fused permease and ATPase components | GO:0042493 response to drug | o | tt |
| *sbmC* | CDS | DNA gyrase inhibitor | Cytoplasmic | COG3449;DNA gyrase inhibitor | GO:0006261 DNA dependent DNA replication -!- GO:0006281 DNA repair -!- GO:0009432 SOS response | gg | t |
| *sbp* | CDS | sulfate transporter subunit -!- periplasmic-binding component of ABC superfamily | Periplasmic | COG1613;ABC-type sulfate transport system, periplasmic component | GO:0006790 sulfur metabolism | o | tt |
| *sdaA* | CDS | L-serine deaminase I | Cytoplasmic | COG1760;L-serine deaminase | GO:0009063 amino acid catabolism | gg | t |
| *sdaB* | CDS | L-serine deaminase II | Cytoplasmic | COG1760;L-serine deaminase | GO:0019395 fatty acid oxidation -!- GO:0009063 amino acid catabolism | gg | t |
| *sdaC* | CDS | predicted serine transporter | Integral Membrane Protein | COG0814;Amino acid permeases | GO:0006564 L-serine biosynthesis | g | tt |
| *sdhA* | CDS | succinate dehydrogenase, flavoprotein subunit | Cytoplasmic | COG1053;Succinate dehydrogenase/fumarate reductase, flavoprotein subunit | GO:0006099 tricarboxylic acid cycle -!- GO:0009060 aerobic respiration | gg | t |
| *sdhB* | CDS | succinate dehydrogenase, FeS subunit | Cytoplasmic | COG0479;Succinate dehydrogenase/fumarate reductase, Fe-S protein subunit | GO:0006099 tricarboxylic acid cycle -!- GO:0009060 aerobic respiration | gg | t |
| *sdhC* | CDS | succinate dehydrogenase, membrane subunit, binds cytochrome b556 | Integral Membrane Protein | COG2009;Succinate dehydrogenase/fumarate reductase, cytochrome b subunit | GO:0006099 tricarboxylic acid cycle -!- GO:0009060 aerobic respiration -!- GO:0017004 cytochrome biogenesis | o | tt |
| *sdhD* | CDS | succinate dehydrogenase, membrane subunit, binds cytochrome b556 | Integral Membrane Protein | COG2142;Succinate dehydrogenase, hydrophobic anchor subunit | GO:0006099 tricarboxylic acid cycle -!- GO:0009060 aerobic respiration | o | tt |
| *sdiA* | CDS | DNA-binding transcriptional activator | Cytoplasmic | COG2771;DNA-binding HTH domain-containing proteins | GO:0006350 transcription | gg | t |
| *secA* | CDS | preprotein translocase subunit, ATPase that targets protein precursors to the SecYE core translocon | Cytoplasmic | COG0653;Preprotein translocase subunit SecA (ATPase, RNA helicase) | | o | tt |
| *secB* | CDS | protein export chaperone | Cytoplasmic | COG1952;Preprotein translocase subunit SecB | GO:0006457 protein folding | o | tt |
| *secD* | CDS | SecYEG protein translocase auxillary subunit | Integral Membrane Protein | COG0342;Preprotein translocase subunit SecD | | o | tt |
| *secE* | CDS | preprotein translocase membrane subunit | Integral Membrane Protein | COG0690;Preprotein translocase subunit SecE | | o | tt |
| *secF* | CDS | SecYEG protein translocase auxillary subunit | Integral Membrane Protein | COG0341;Preprotein translocase subunit SecF | | o | tt |
| *secG* | CDS | preprotein translocase membrane subunit | Integral Membrane Protein | COG1314;Preprotein translocase subunit SecG | | o | tt |
| *secM* | CDS | regulator of secA translation | Periplasmic | | | | |
| *secY* | CDS | preprotein translocase membrane subunit | Integral Membrane Protein | COG0201;Preprotein translocase subunit SecY | | o | tt |
| *selA* | CDS | selenocysteine synthase | Cytoplasmic | COG1921;Selenocysteine synthase [seryl-tRNASer selenium transferase] | GO:0008615 pyridoxine biosynthesis -!- GO:0009451 RNA modification | g | t |
| *selB* | CDS | selenocysteinyl-tRNA-specific translation factor | Cytoplasmic | COG3276;Selenocysteine-specific translation elongation factor | GO:0006412 protein biosynthesis | o | t |
| *selC* | tRNA | tRNA-Sec(UCA) (Selenocysteyl tRNA-UCA, converted from serine tRNA) | | | | | |
| *selD* | CDS | selenophosphate synthase | Cytoplasmic | COG0709;Selenophosphate synthase | GO:0006418 amino acid activation | gg | t |
| *seqA* | CDS | regulatory protein for replication initiation | Cytoplasmic | COG3057;Negative regulator of replication initiationR | GO:0006261 DNA dependent DNA replication | o | tt |
| *serA* | CDS | D-3-phosphoglycerate dehydrogenase | Cytoplasmic | COG0111;Phosphoglycerate dehydrogenase and related dehydrogenases | GO:0006564 L-serine biosynthesis | gg | t |
| *serB* | CDS | 3-phosphoserine phosphatase | Cytoplasmic | COG0560;Phosphoserine phosphatase | GO:0006545 glycine biosynthesis -!- GO:0006564 L-serine biosynthesis | gg | t |
| *serC* | CDS | 3-phosphoserine/phosphohydroxythreonine aminotransferase | Cytoplasmic | COG1932;Phosphoserine aminotransferase | GO:0006564 L-serine biosynthesis | gg | t |
| *serS* | CDS | seryl-tRNA synthetase, also charges selenocysteinyl-tRNA with serine | Cytoplasmic | COG0172;Seryl-tRNA synthetase | GO:0006418 amino acid activation | gg | tt |
| *serT* | tRNA | tRNA-Ser(UGA) (Serine tRNA1) | | | | | |
| *serU* | tRNA | tRNA-Ser(CGA) (Serine tRNA2) | | | | | |
| *serV* | tRNA | tRNA-Ser(GCU) (Serine tRNA3) | | | | | |
| *serW* | tRNA | tRNA-Ser(GGA) (Serine tRNA5) | | | | | |
| *serX* | tRNA | tRNA-Ser(GGA) (Serine tRNA5) | | | | | |
| *setA* | CDS | broad specificity sugar efflux system | Integral Membrane Protein | | GO:0016052 carbohydrate | o | tt |
| *setB* | CDS | lactose/glucose efflux system | Integral Membrane Protein | COG0477;Permeases of the major facilitator superfamily | GO:0016052 carbohydrate | o | tt |
| *setC* | CDS | predicted sugar efflux system | Integral Membrane Protein | COG0477;Permeases of the major facilitator superfamily | | o | tt |
| *sfcA* | CDS | malate dehydrogenase, NAD-requiring (malic | Cytoplasmic | COG0281;Malic enzyme | GO:0006094 gluconeogenesis | | |
| *sfmA* | CDS | predicted fimbrial-like adhesin protein | Periplasmic | | | o | tt |
| *sfmC* | CDS | pilin chaperone, periplasmic | Periplasmic | COG3121;P pilus assembly protein, chaperone PapD | GO:0006457 protein folding | o | tt |
| *sfmD* | CDS | predicted outer membrane export usher protein | Outer Membrane B-barrel protein | COG3188;P pilus assembly protein, porin PapC | | o | tt |
| *sfmF* | CDS | predicted fimbrial-like adhesin protein | Periplasmic | COG3539;P pilus assembly protein, pilin FimA | | | |
| *sfmH* | CDS | predicted fimbrial-like adhesin protein | Cytoplasmic | COG3539;P pilus assembly protein, pilin FimA | GO:0009296 flagella biogenesis -!- GO:0042330 taxis | | |
| *sfsA* | CDS | predicted DNA-binding transcriptional regulator | Cytoplasmic | COG1489;DNA-binding protein, stimulates sugar fermentation | GO:0016052 carbohydrate | g | o |
| *sfsB* | CDS | DNA-binding transcriptional regulator | Cytoplasmic | COG3423;Predicted transcriptional regulator | GO:0016052 carbohydrate catabolism -!- GO:0006350 | o | tt |
| *sgbE* | CDS | L-ribulose-5-phosphate 4-epimerase | Cytoplasmic | COG0235;Ribulose-5-phosphate 4-epimerase and related epimerases and aldolases | GO:0016052 carbohydrate catabolism | gg | o |
| *sgbH* | CDS | 3-keto-L-gulonate 6-phosphate decarboxylase | Cytoplasmic | COG0269;3-hexulose-6-phosphate synthase and related proteins | | gg | t |
| *sgbU* | CDS | predicted L-xylulose 5-phosphate 3-epimerase | Cytoplasmic | COG3623;Putative L-xylulose-5-phosphate 3-epimerase | | | |
| *sgcA* | CDS | KpLE2 phage-like element; predicted phosphotransferase enzyme IIA component | Cytoplasmic | COG1762;Phosphotransferase system mannitol/fructose-specific IIA domain (Ntr-type) | | o | t |
| *sgcB* | CDS | predicted enzyme IIB component of PTS | Cytoplasmic | | | | |
| *sgcC* | CDS | KpLE2 phage-like element; predicted phosphotransferase enzyme IIC component | Integral Membrane Protein | COG3775;Phosphotransferase system, galactitol-specific IIC component | GO:0009401 phosphoenolpyruvate-dependent sugar phosphotransferase system | o | tt |
| *sgcE* | CDS | KpLE2 phage-like element; predicted epimerase | Cytoplasmic | COG0036;Pentose-5-phosphate-3-epimerase | | o | t |
| *sgcQ* | CDS | KpLE2 phage-like element; predicted nucleoside triphosphatase | Cytoplasmic | COG0434;Predicted TIM-barrel enzyme | | o | o |
| *sgcR* | CDS | KpLE2 phage-like element; predicted DNA-binding transcriptional regulator | Cytoplasmic | COG1349;Transcriptional regulators of sugar metabolism | GO:0006350 transcription | g | t |
| *sgcX* | CDS | KpLE2 phage-like element; predicted endoglucanase with Zn-dependent exopeptidase domain | Cytoplasmic | | | | |
| *sgrR* | CDS | DNA-binding transcriptional regulator | Cytoplasmic | COG4533;ABC-type uncharacterized transport system, periplasmic component | | o | tt |
| *sgrS* | misc_RNA | small antisense RNA | | | | | |
| *shiA* | CDS | shikimate transporter | Integral Membrane Protein | COG0477;Permeases of the major facilitator superfamily | GO:0009423 chorismate biosynthesis | o | tt |
| *sieB* | CDS | Rac prophage; phage superinfection exclusion | Integral Membrane Protein | | | | |
| *sixA* | CDS | phosphohistidine phosphatase | Cytoplasmic | COG2062;Phosphohistidine phosphatase SixA | GO:0006464 protein modification | g | t |
| *slp* | CDS | outer membrane lipoprotein | Outer Membrane Lipoprotein | | GO:0042594 response to starvation | o | o |
| *slt* | CDS | lytic murein transglycosylase, soluble | Periplasmic | | GO:0009252 peptidoglycan biosynthesis | o | tt |
| *slyA* | CDS | DNA-binding transcriptional activator | Cytoplasmic | COG1846;Transcriptional regulators | GO:0006350 transcription | | |
| *slyB* | CDS | outer membrane lipoprotein | Outer Membrane Lipoprotein | | | o | tt |
| *slyD* | CDS | FKBP-type peptidyl prolyl cis-trans isomerase (rotamase) | Cytoplasmic | COG1047;FKBP-type peptidyl-prolyl cis-trans isomerases 2 | GO:0006457 protein folding | gg | tt |
| *slyX* | CDS | conserved protein | Cytoplasmic | COG2900;Uncharacterized protein conserved in bacteria | | gg | o |
| *smf* | CDS | conserved protein | Cytoplasmic | | | | |
| *smg* | CDS | conserved protein | Cytoplasmic | COG2922;Uncharacterized protein conserved in bacteria | | gg | tt |
| *smpA* | CDS | small membrane lipoprotein | Outer Membrane Lipoprotein | COG2913;Small protein A (tmRNA-binding) | | o | tt |
| *smpB* | CDS | trans-translation protein | Cytoplasmic | COG0691;tmRNA-binding protein | | gg | tt |
| *smtA* | CDS | S-adenosylmethionine-dependent methyltransferase | Cytoplasmic | COG0500;SAM-dependent methyltransferases | GO:0006071 glycerol metabolism | gg | t |
| *sodA* | CDS | superoxide dismutase, Mn | Cytoplasmic | | GO:0006805 xenobiotic metabolism | gg | o |
| *sodB* | CDS | superoxide dismutase, Fe | Cytoplasmic | COG0605;Superoxide dismutase | GO:0006805 xenobiotic metabolism | gg | o |
| *sodC* | CDS | superoxide dismutase, Cu, Zn | Periplasmic | COG2032;Cu/Zn superoxide dismutase | GO:0006805 xenobiotic metabolism | o | tt |
| *sohA* | CDS | predicted regulator | Cytoplasmic | COG2002;Regulators of stationary/sporulation gene expression | | gg | tt |
| *sohB* | CDS | predicted inner membrane peptidase | Integral Membrane Protein | COG0616;Periplasmic serine proteases (ClpP class) | | o | tt |
| *sokB* | misc_RNA | regulatory, antisense sRNA | | | | | |
| *sokC* | misc_RNA | regulatory, antisense RNA | | | | | |
| *solA* | CDS | N-methyltryptophan oxidase, FAD-binding | Cytoplasmic | COG0665;Glycine/D-amino acid oxidases (deaminating) | | gg | t |
| *soxR* | CDS | DNA-binding transcriptional dual regulator, Fe-S center for redox-sensing | Cytoplasmic | COG0789;Predicted transcriptional regulators | GO:0006350 transcription -!- GO:0006805 xenobiotic metabolism | o | tt |
| *soxS* | CDS | DNA-binding transcriptional dual regulator | Cytoplasmic | COG2207;AraC-type DNA-binding domain-containing proteins | GO:0006350 transcription -!- GO:0006805 xenobiotic metabolism | o | tt |
| *speA* | CDS | biosynthetic arginine decarboxylase, PLP-binding | Cytoplasmic | COG1166;Arginine decarboxylase (spermidine biosynthesis) | GO:0006596 polyamine biosynthesis | o | tt |
| *speB* | CDS | agmatinase | Cytoplasmic | COG0010;Arginase/agmatinase/formimionoglutamate hydrolase, arginase family | GO:0006596 polyamine biosynthesis | gg | t |
| *speC* | CDS | ornithine decarboxylase, constitutive | Cytoplasmic | | GO:0006596 polyamine biosynthesis | | |
| *speD* | CDS | S-adenosylmethionine decarboxylase | Cytoplasmic | COG1586;S-adenosylmethionine decarboxylase | GO:0006596 polyamine biosynthesis | gg | o |
| *speE* | CDS | spermidine synthase (putrescine aminopropyltransferase) | Cytoplasmic | COG0421;Spermidine synthase | GO:0006596 polyamine biosynthesis | gg | o |
| *speF* | CDS | ornithine decarboxylase isozyme, inducible | Cytoplasmic | COG1982;Arginine/lysine/ornithine decarboxylases | GO:0006596 polyamine biosynthesis | gg | tt |
| *speG* | CDS | spermidine N1-acetyltransferase | Cytoplasmic | COG1670;Acetyltransferases, including N-acetylases of ribosomal proteins | GO:0006596 polyamine biosynthesis | gg | t |

| | | | | | | | |
|---|---|---|---|---|---|---|---|
| *spf* | misc_RNA | Spot 42 RNA | | | | | |
| *spoT* | CDS | bifunctional (p)ppGpp synthetase II -!- guanosine-3',5'-bis pyrophosphate 3'-pyrophosphohydrolase | Cytoplasmic | COG0317;Guanosine polyphosphate pyrophosphohydrolases/synthetases | GO:0015949 nucleobase, nucleoside and nucleotide interconversion -!- GO:0042594 response to starvation | o | tt |
| *sppA* | CDS | protease IV (signal peptide peptidase) | Membrane Anchored | COG0616;Periplasmic serine proteases (ClpP class) | | o | tt |
| *spr* | CDS | predicted peptidase, outer membrane lipoprotein | Outer Membrane Lipoprotein | COG0791;Cell wall-associated hydrolases (invasion-associated proteins) | | gg | t |
| *sprT* | CDS | conserved protein | Cytoplasmic | COG3091;Uncharacterized protein conserved in bacteria | | o | tt |
| *spy* | CDS | envelope stress induced periplasmic protein | Periplasmic | COG3678;P pilus assembly/Cpx signaling pathway, periplasmic inhibitor/zinc-resistance associated protein | | o | tt |
| *sra* | CDS | 30S ribosomal subunit protein S22 | Cytoplasmic | | GO:0006412 protein biosynthesis | | |
| *sraB* | misc_RNA | small RNA | | | | | |
| *sraD* | misc_RNA | small RNA | | | | | |
| *sraF* | misc_RNA | small RNA | | | | | |
| *sraG* | misc_RNA | small RNA | | | | | |
| *srlA* | CDS | glucitol/sorbitol-specific enzyme IIC component of | Integral Membrane Protein | | GO:0016052 carbohydrate | | |
| *srlB* | CDS | glucitol/sorbitol-specific enzyme IIA component of | Cytoplasmic | COG3731;Phosphotransferase system sorbitol-specific component IIA | GO:0016052 carbohydrate | gg | t |
| *srlD* | CDS | sorbitol-6-phosphate dehydrogenase | Outer Membrane Lipoprotein | COG1028;Dehydrogenases with different specificities (related to short-chain alcohol dehydrogenases) | GO:0016052 carbohydrate catabolism | gg | t |
| *srlE* | CDS | glucitol/sorbitol-specific enzyme IIB component of | Integral Membrane Protein | | GO:0016052 carbohydrate | | |
| *srlR* | CDS | DNA-bindng transcriptional repressor | Cytoplasmic | COG1349;Transcriptional regulators of sugar metabolism | GO:0016052 carbohydrate catabolism -!- GO:0006350 transcription -!- GO:0006355 regulation of transcription, DNA- | o | tt |
| *srmB* | CDS | ATP-dependent RNA helicase | Cytoplasmic | COG0513;Superfamily II DNA and RNA helicases | GO:0009451 RNA modification | gg | tt |
| *sroF* | misc_RNA | small RNA | | | | | |
| *ssb* | CDS | Single-stranded DNA-binding protein | Cytoplasmic | COG0629;Single-stranded DNA-binding protein | GO:0006310 DNA recombination -!- GO:0009432 SOS response | o | tt |
| *sseA* | CDS | 3-mercaptopyruvate sulfurtransferase | Cytoplasmic | | GO:0006790 sulfur metabolism | o | o |
| *sseB* | CDS | rhodanase-like enzyme, sulfur transfer from | Cytoplasmic | | GO:0006790 sulfur metabolism | | |
| *ssnA* | CDS | predicted chlorohydrolase/aminohydrolase | Cytoplasmic | | GO:0007049 cell cycle | o | tt |
| *sspA* | CDS | stringent starvation protein A | Cytoplasmic | COG0625;Glutathione S-transferase | GO:0006350 transcription -!- GO:0006355 regulation of transcription, DNA-dependent -!- GO:0042594 response to starvation | gg | t |
| *sspB* | CDS | ClpXP protease specificity-enhancing factor | Cytoplasmic | COG2969;Stringent starvation protein B | GO:0042594 response to starvation | gg | t |
| *ssrA* | misc_RNA | 10Sa RNA SsrA (tmRNA) | | | | | |
| *ssrS* | misc_RNA | 6S regulatory RNA | | | | | |
| *sstT* | CDS | sodium:serine/threonine symporter | Integral Membrane Protein | COG3633;Na+/serine symporter | GO:0006564 L-serine biosynthesis -!- GO:0009088 threonine biosynthesis | o | tt |
| *ssuA* | CDS | alkanesulfonate transporter subunit -!- periplasmic-binding component of ABC superfamily | Periplasmic | | GO:0006793 phosphorus metabolism | o | tt |
| *ssuB* | CDS | alkanesulfonate transporter subunit -!- ATP-binding component of ABC superfamily | Cytoplasmic | COG1116;ABC-type nitrate/sulfonate/bicarbonate transport system, ATPase component | GO:0006790 sulfur metabolism | gg | t |
| *ssuC* | CDS | alkanesulfonate transporter subunit -!- membrane component of ABC superfamily | Integral Membrane Protein | | GO:0006793 phosphorus metabolism | | |
| *ssuD* | CDS | alkanesulfonate monooxygenase, FMNH(2)-dependent | Cytoplasmic | COG2141;Coenzyme F420-dependent N5,N10-methylene tetrahydromethanopterin reductase and related flavin-dependent oxidoreductases | GO:0006790 sulfur metabolism | gg | t |
| *ssuE* | CDS | NAD(P)H-dependent FMN reductase | Cytoplasmic | COG0431;Predicted flavoprotein | GO:0006790 sulfur metabolism -!- GO:0042594 response to starvation | gg | t |
| *stfE* | CDS | e14 prophage; predicted side tail fiber protein fragment (pseudogene) | Cytoplasmic | | | | |
| *stfQ* | CDS | Qin prophage; predicted side tail fibre assembly protein | Cytoplasmic | | | gg | o |
| *stfR* | CDS | Rac prophage; predicted tail fiber protein | Cytoplasmic | | | o | tt |
| *sthA* | CDS | pyridine nucleotide transhydrogenase, soluble | Cytoplasmic | COG1249;Pyruvate/2-oxoglutarate dehydrogenase complex, dihydrolipoamide dehydrogenase (E3) component, and related enzymes | GO:0009226 nucleotide-sugar biosynthesis | | |
| *stpA* | CDS | DNA binding protein, nucleoid-associated | Cytoplasmic | COG2916;DNA-binding protein H-NS | GO:0006457 protein folding | o | tt |
| *sucA* | CDS | 2-oxoglutarate decarboxylase, thiamin-requiring | Cytoplasmic | COG0567;2-oxoglutarate dehydrogenase complex, dehydrogenase (E1) component, and related enzymes | GO:0006099 tricarboxylic acid cycle | gg | t |
| *sucB* | CDS | dihydrolipoyltranssuccinase | Cytoplasmic | COG0508;Pyruvate/2-oxoglutarate dehydrogenase complex, dihydrolipoamide acyltransferase (E2) component, and related enzymes | GO:0006099 tricarboxylic acid cycle | g | t |
| *sucC* | CDS | succinyl-CoA synthetase, beta subunit | Cytoplasmic | COG0045;Succinyl-CoA synthetase, beta subunit | GO:0006099 tricarboxylic acid cycle | gg | t |
| *sucD* | CDS | succinyl-CoA synthetase, NAD(P)-binding, alpha subunit | Cytoplasmic | COG0074;Succinyl-CoA synthetase, alpha subunit | GO:0006099 tricarboxylic acid cycle | g | t |
| *sufA* | CDS | Fe-S cluster assembly protein | Cytoplasmic | COG0316;Uncharacterized conserved protein | | gg | o |
| *sufB* | CDS | component of SufBCD complex | Cytoplasmic | COG0719;ABC-type transport system involved in Fe-S cluster assembly, permease component | GO:0006457 protein folding | | |
| *sufC* | CDS | component of SufBCD complex, ATP-binding component of ABC superfamily | Cytoplasmic | COG0396;ABC-type transport system involved in Fe-S cluster assembly, ATPase component | | gg | tt |
| *sufD* | CDS | component of SufBCD complex | Cytoplasmic | COG0719;ABC-type transport system involved in Fe-S cluster assembly, permease component | | gg | t |
| *sufE* | CDS | sulfur acceptor protein | Cytoplasmic | COG2166;SufE protein probably involved in Fe-S center assembly | | gg | o |
| *sufI* | CDS | repressor protein for FtsI | Periplasmic | COG2132;Putative multicopper oxidases | | o | tt |
| *sufS* | CDS | selenocysteine lyase, PLP-dependent | Cytoplasmic | COG0520;Selenocysteine lyase | | gg | t |
| *sugE* | CDS | multidrug efflux system protein | Integral Membrane Protein | | | | |
| *suhB* | CDS | inositol monophosphatase | Cytoplasmic | COG0483;Archaeal fructose-1,6-bisphosphatase and related enzymes of inositol monophosphatase family | GO:0006350 transcription | gg | t |
| *sulA* | CDS | SOS cell division inhibitor | Cytoplasmic | COG5404;SOS-response cell division inhibitor, blocks FtsZ ring formation | GO:0006281 DNA repair -!- GO:0009432 SOS response | o | tt |
| *surA* | CDS | peptidyl-prolyl cis-trans isomerase (PPIase) | Periplasmic | COG0760;Parvulin-like peptidyl-prolyl isomerase | GO:0006457 protein folding | o | tt |
| *surE* | CDS | broad specificity 5'(3')-nucleotidase and polyphosphatase | Cytoplasmic | COG0496;Predicted acid phosphatase | GO:0006950 response to stress | gg | t |
| *syd* | CDS | predicted protein | Cytoplasmic | | | gg | t |
| *tadA* | CDS | tRNA-specific adenosine deaminase | Cytoplasmic | COG0590;Cytosine/adenosine deaminases | GO:0009451 RNA modification | o | t |
| *tag* | CDS | 3-methyl-adenine DNA glycosylase I, constitutive | Cytoplasmic | COG2818;3-methyladenine DNA glycosylase | GO:0006281 DNA repair | g | t |
| *talA* | CDS | transaldolase A | Cytoplasmic | COG0176;Transaldolase | GO:0009052 pentose-phosphate shunt, non-oxidative branch | gg | t |
| *talB* | CDS | transaldolase B | Cytoplasmic | COG0176;Transaldolase | GO:0009052 pentose-phosphate shunt, non-oxidative branch | gg | o |
| *tam* | CDS | trans-aconitate methyltransferase | Cytoplasmic | COG4106;Trans-aconitate methyltransferase | | gg | o |
| *tap* | CDS | methyl-accepting protein IV | Integral Membrane Protein | COG0840;Methyl-accepting chemotaxis protein | GO:0042330 taxis | o | tt |
| *tar* | CDS | methyl-accepting chemotaxis protein II | Integral Membrane Protein | COG0840;Methyl-accepting chemotaxis protein | GO:0042330 taxis | o | tt |
| *tas* | CDS | predicted oxidoreductase, NADP(H)-dependent aldo-keto reductase | Cytoplasmic | COG0667;Predicted oxidoreductases (related to aryl-alcohol dehydrogenases) | | g | t |
| *tatA* | CDS | TatABCE protein translocation system subunit | Membrane Anchored | | | o | tt |
| *tatB* | CDS | TatABCE protein translocation system subunit | Membrane Anchored | | | | |
| *tatC* | CDS | TatABCE protein translocation system subunit | Integral Membrane Protein | COG0805;Sec-independent protein secretion pathway component TatC | | o | tt |
| *tatD* | CDS | DNase, magnesium-dependent | Cytoplasmic | | | | |
| *tatE* | CDS | TatABCE protein translocation system subunit | Membrane Anchored | COG1826;Sec-independent protein secretion pathway components | | o | t |
| *tauA* | CDS | taurine transporter subunit -!- periplasmic-binding component of ABC superfamily | Periplasmic | COG4521;ABC-type taurine transport system, periplasmic component | GO:0006790 sulfur metabolism -!- GO:0009310 amine catabolism | o | tt |
| *tauB* | CDS | taurine transporter subunit -!- ATP-binding component of ABC superfamily | Cytoplasmic | COG4525;ABC-type taurine transport system, ATPase component | GO:0006790 sulfur metabolism -!- GO:0009310 amine catabolism | gg | t |
| *tauC* | CDS | taurine transporter subunit -!- membrane component of ABC superfamily | Integral Membrane Protein | COG0600;ABC-type nitrate/sulfonate/bicarbonate transport system, permease component | GO:0006790 sulfur metabolism -!- GO:0009310 amine catabolism | o | tt |
| *tauD* | CDS | taurine dioxygenase, 2-oxoglutarate-dependent | Cytoplasmic | COG2175;Probable taurine catabolism dioxygenase | GO:0009310 amine catabolism -!- GO:0006790 sulfur metabolism | gg | t |
| *tbpA* | CDS | thiamin transporter subunit -!- periplasmic-binding component of ABC superfamily | Periplasmic | COG4143;ABC-type thiamine transport system, periplasmic component | GO:0009228 thiamin biosynthesis | o | tt |
| *tdcA* | CDS | DNA-binding transcriptional activator | Cytoplasmic | COG0583;Transcriptional regulator | GO:0009063 amino acid catabolism -!- GO:0006350 transcription | g | o |
| *tdcB* | CDS | catabolic threonine dehydratase, PLP-dependent | Cytoplasmic | COG1171;Threonine dehydratase | GO:0009063 amino acid catabolism | gg | t |
| *tdcC* | CDS | L-threonine/L-serine transporter | Integral Membrane Protein | COG0814;Amino acid permeases | GO:0009088 threonine biosynthesis | o | tt |
| *tdcD* | CDS | propionate kinase/acetate kinase C, anaerobic | Cytoplasmic | COG0282;Acetate kinase | GO:0009063 amino acid catabolism | | |
| *tdcE* | CDS | pyruvate formate-lyase 4/2-ketobutyrate formate-lyase | Cytoplasmic | | GO:0009063 amino acid catabolism -!- GO:0006113 fermentation -!- GO:0009061 anaerobic respiration | | |
| *tdcF* | CDS | predicted L-PSP (mRNA) endoribonuclease | Cytoplasmic | | | | |
| *tdcG* | CDS | L-serine dehydratase 3 | Cytoplasmic | | GO:0009063 amino acid catabolism | | |
| *tdcR* | CDS | DNA-binding transcriptional activator | Cytoplasmic | | GO:0009063 amino acid catabolism -!- GO:0006350 transcription | | |
| *tdh* | CDS | threonine 3-dehydrogenase, NAD(P)-binding | Cytoplasmic | COG1063;Threonine dehydrogenase and related Zn-dependent dehydrogenases | GO:0009063 amino acid catabolism | g | t |
| *tdk* | CDS | thymidine kinase/deoxyuridine kinase | Cytoplasmic | COG1435;Thymidine kinase | GO:0015949 nucleobase, nucleoside and nucleotide interconversion | gg | t |
| *tehA* | CDS | potassium-tellurite ethidium and proflavin transporter | Integral Membrane Protein | COG1275;Tellurite resistance protein and related permeases | GO:0042493 response to drug | o | tt |
| *tehB* | CDS | predicted S-adenosyl-L-methionine-dependent methyltransferase | Cytoplasmic | COG0500;SAM-dependent methyltransferases | GO:0042493 response to drug | gg | t |
| *tesA* | CDS | multifunctional acyl-CoA thioesterase I -!- protease I -!- lysophospholipase L1 | Periplasmic | COG2755;Lysophospholipase L1 and related esterases | | o | tt |
| *tesB* | CDS | acyl-CoA thioesterase II | Cytoplasmic | COG1946;Acyl-CoA thioesterase | | gg | o |

| | | | | | | | |
|---|---|---|---|---|---|---|---|
| *tfaD* | CDS | DLP12 prophage; predicted tail fiber assembly protein (pseudogene) | Cytoplasmic | | | | |
| *tfaE* | CDS | e14 prophage; predicted tail fiber assembly protein | Cytoplasmic | | | gg | t |
| *tfaQ* | CDS | Qin prophage; predicted tail fibre assembly protein | Cytoplasmic | | | gg | o |
| *tfaR* | CDS | Rac prophage; predicted tail fiber assembly protein | Cytoplasmic | | | gg | o |
| *tfaS* | CDS | CPS-53 (KpLE1) prophage; tail fiber assembly protein fragment (pseudogene) | Cytoplasmic | | | | |
| *tff* | misc_RNA | small RNA | | | | | |
| *tgt* | CDS | tRNA-guanine transglycosylase | Cytoplasmic | COG0343;Queuine/archaeosine tRNA-ribosyltransferase | GO:0009451 RNA modification | gg | t |
| *thiC* | CDS | thiamin (pyrimidine moiety) biosynthesis protein | Cytoplasmic | COG0422;Thiamine biosynthesis protein ThiC | GO:0009228 thiamin biosynthesis | gg | t |
| *thiD* | CDS | bifunctional hydroxy-methylpyrimidine kinase -!- hydroxy-phosphomethylpyrimidine kinase | Cytoplasmic | COG0351;Hydroxymethylpyrimidine/phosphomethylpyrim idine kinase | GO:0009228 thiamin biosynthesis | g | t |
| *thiE* | CDS | thiamin phosphate synthase (thiamin phosphate pyrophosphorylase) | Cytoplasmic | COG0352;Thiamine monophosphate synthase | GO:0009228 thiamin biosynthesis | gg | t |
| *thiF* | CDS | thiamin (thiazole moiety) biosynthesis protein | Membrane Anchored | | GO:0009228 thiamin biosynthesis | gg | o |
| *thiG* | CDS | thiamin biosynthesis ThiGH complex subunit | Cytoplasmic | | GO:0009228 thiamin biosynthesis | | |
| *thiH* | CDS | thiamin biosynthesis ThiGH complex subunit | Cytoplasmic | COG1060;Thiamine biosynthesis enzyme ThiH and related uncharacterized enzymes | GO:0009228 thiamin biosynthesis | gg | t |
| *thiI* | CDS | sulfurtransferase required for thiamine and 4-thiouridine biosynthesis | Cytoplasmic | COG0301;Thiamine biosynthesis ATP pyrophosphatase -!- COG0607;Rhodanese-related sulfurtransferase | GO:0009228 thiamin biosynthesis -!- GO:0009451 RNA modification | gg | tt |
| *thiL* | CDS | thiamin-monophosphate kinase | Cytoplasmic | COG0611;Thiamine monophosphate kinase | GO:0009228 thiamin biosynthesis | gg | o |
| *thiM* | CDS | hydoxyethylthiazole kinase | Cytoplasmic | COG2145;Hydroxyethylthiazole kinase, sugar kinase family | GO:0009228 thiamin biosynthesis | gg | o |
| *thiP* | CDS | fused subunits of thiamin transporter -!- membrane components of ABC superfamily | Integral Membrane Protein | COG1178;ABC-type Fe3+ transport system, permease component | GO:0009228 thiamin biosynthesis | o | tt |
| *thiQ* | CDS | thiamin transporter subunit -!- ATP-binding component of ABC superfamily | Cytoplasmic | COG3840;ABC-type thiamine transport system, ATPase component | GO:0009228 thiamin biosynthesis | o | tt |
| *thiS* | CDS | sulphur carrier protein | Cytoplasmic | | | gg | t |
| *thrA* | CDS | fused aspartokinase I -!- homoserine dehydrogenase I | Cytoplasmic | COG0527;Aspartokinases -!- COG0460;Homoserine dehydrogenase | GO:0009088 threonine biosynthesis -!- GO:0009086 methionine biosynthesis -!- GO:0009090 homoserine biosynthesis | | t |
| *thrB* | CDS | homoserine kinase | Cytoplasmic | COG0083;Homoserine kinase | GO:0009088 threonine biosynthesis | g | o |
| *thrC* | CDS | threonine synthase | Cytoplasmic | COG0498;Threonine synthase | GO:0009088 threonine biosynthesis | gg | t |
| *thrL* | CDS | thr operon leader peptide | Cytoplasmic | | GO:0009088 threonine biosynthesis | | |
| *thrS* | CDS | threonyl-tRNA synthetase | Cytoplasmic | COG0441;Threonyl-tRNA synthetase | GO:0006418 amino acid activation | gg | t |
| *thrT* | tRNA | tRNA-Thr(GGU) (Threonine tRNA3) | | | | | |
| *thrU* | tRNA | tRNA-Thr(UGU) (Threonine tRNA4) | | | | | |
| *thrV* | tRNA | tRNA-Thr(GGU) (Threonine tRNA1) | | | | | |
| *thrW* | tRNA | tRNA-Thr(CGU) (Threonine tRNA2) | | | | | |
| *thyA* | CDS | thymidylate synthetase | Cytoplasmic | COG0207;Thymidylate synthase | GO:0015949 nucleobase, nucleoside and nucleotide interconversion -!- GO:0009257 10-formyltetrahydrofolate biosynthesis | gg | t |
| *tiaE* | CDS | 2-keto-D-gluconate reductase (glyoxalate reductase) (2-ketoaldonate reductase) | Cytoplasmic | COG1052;Lactate dehydrogenase and related dehydrogenases | GO:0016052 carbohydrate catabolism | | |
| *tig* | CDS | peptidyl-prolyl cis/trans isomerase (trigger factor) | Cytoplasmic | COG0544;FKBP-type peptidyl-prolyl cis-trans isomerase (trigger factor) | GO:0006457 protein folding | o | tt |
| *tilS* | CDS | tRNA(Ile)-lysidine synthetase | Cytoplasmic | COG0037;Predicted ATPase of the PP-loop superfamily implicated in cell cycle control | | o | tt |
| *tktA* | CDS | transketolase 1, thiamin-binding | Cytoplasmic | | GO:0016052 carbohydrate catabolism -!- GO:0009052 pentose-phosphate shunt, non-oxidative branch -!- GO:0015949 nucleobase, nucleoside and nucleotide | | |
| *tktB* | CDS | transketolase 2, thiamin-binding | Cytoplasmic | COG0021;Transketolase | GO:0016052 carbohydrate catabolism -!- GO:0009052 pentose-phosphate shunt, non-oxidative branch -!- GO:0015949 nucleobase, nucleoside and nucleotide | gg | t |
| *tldD* | CDS | predicted peptidase | Cytoplasmic | COG0312;Predicted Zn-dependent proteases and their inactivated | | o | tt |
| *tmk* | CDS | thymidylate kinase | Cytoplasmic | COG0125;Thymidylate kinase | GO:0015949 nucleobase, nucleoside and nucleotide interconversion | gg | o |
| *tnaA* | CDS | tryptophanase/L-cysteine desulfhydrase, PLP-dependent | Cytoplasmic | | GO:0009063 amino acid catabolism | gg | o |
| *tnaB* | CDS | tryptophan transporter of low affinity | Integral Membrane Protein | COG0814;Amino acid permeases | GO:0000162 tryptophan biosynthesis | | |
| *tnaB* | CDS | tryptophan transporter of low affinity | Integral Membrane Protein | | | | |
| *tnaB* | CDS | tryptophan transporter of low affinity | Integral Membrane Protein | | | | |
| *tnaC* | CDS | tryptophanase leader peptide | Cytoplasmic | | GO:0009063 amino acid catabolism | gg | o |
| *tolA* | CDS | membrane anchored protein in TolA-TolQ-TolR complex | Membrane Anchored | COG3064;Membrane protein involved in colicin uptake | | o | tt |
| *tolB* | CDS | periplasmic protein | Periplasmic | COG0823;Periplasmic component of the Tol biopolymer transport system | | | |
| *tolC* | CDS | transport channel | Outer Membrane B-barrel protein | COG1538;Outer membrane protein | | | |
| *tolQ* | CDS | membrane spanning protein in TolA-TolQ-TolR complex | Integral Membrane Protein | COG0811;Biopolymer transport proteins | | o | tt |
| *tolR* | CDS | membrane spanning protein in TolA-TolQ-TolR complex | Membrane Anchored | COG0848;Biopolymer transport protein | | o | tt |
| *tonB* | CDS | membrane spanning protein in TonB-ExbB-ExbD complex | Membrane Anchored | COG0810;Periplasmic protein TonB, links inner and outer membranes | | | |
| *topA* | CDS | DNA topoisomerase I, omega subunit | Cytoplasmic | COG0550;Topoisomerase IA -!- COG0551;Zn-finger domain associated with topoisomerase type I | GO:0006261 DNA dependent DNA replication -!- GO:0006350 transcription | g | tt |
| *topB* | CDS | DNA topoisomerase III | Cytoplasmic | COG0550;Topoisomerase IA | GO:0006261 DNA dependent DNA replication -!- GO:0006350 transcription | o | tt |
| *torA* | CDS | trimethylamine N-oxide (TMAO) reductase I, catalytic subunit | Periplasmic | COG0243;Anaerobic dehydrogenases, typically selenocysteine-containing | GO:0009061 anaerobic respiration | gg | tt |
| *torC* | CDS | trimethylamine N-oxide (TMAO) reductase I, cytochrome c-type subunit | Periplasmic with N-terminal Membrane Anchor | COG3005;Nitrate/TMAO reductases, membrane-bound tetraheme cytochrome c subunit | GO:0009061 anaerobic respiration -!- GO:0017004 cytochrome biogenesis -!- GO:0006350 transcription | o | tt |
| *torD* | CDS | chaperone involved in maturation of TorA subunit of trimethylamine N-oxide reductase system I | Cytoplasmic | COG3381;Uncharacterized component of anaerobic dehydrogenases | GO:0006457 protein folding | gg | t |
| *torI* | CDS | response regulator inhibitor for tor operon | | | | | |
| *torR* | CDS | DNA-binding response regulator in two-component regulatory system with TorS | Cytoplasmic | COG0745;Response regulators consisting of a CheY-like receiver domain and a winged-helix DNA-binding domain | GO:0009061 anaerobic respiration -!- GO:0006350 transcription | gg | o |
| *torS* | CDS | hybrid sensory histidine kinase in two-component regulatory system with TorR | Integral Membrane Protein | COG0642;Signal transduction histidine kinase -!- COG2198;FOG: HPt domain | GO:0009061 anaerobic respiration -!- GO:0006464 protein modification | | |
| *torT* | CDS | periplasmic sensory protein associated with the TorRS two-component regulatory system | Periplasmic | COG1879;ABC-type sugar transport system, periplasmic component | GO:0009061 anaerobic respiration -!- GO:0006464 protein modification | o | tt |
| *torY* | CDS | TMAO reductase III (TorYZ), cytochrome c-type subunit | Periplasmic with N-terminal Membrane | COG3005;Nitrate/TMAO reductases, membrane-bound tetraheme cytochrome c subunit | | o | tt |
| *torZ* | CDS | trimethylamine N-oxide reductase system III, catalytic subunit | Periplasmic | | | gg | tt |
| *tpiA* | CDS | triosephosphate isomerase | Cytoplasmic | COG0149;Triosephosphate isomerase | GO:0006096 glycolysis | gg | t |
| *tpr* | CDS | predicted protamine-like protein | Cytoplasmic | | | gg | t |
| *tpx* | CDS | lipid hydroperoxide peroxidase | Periplasmic | COG2077;Peroxiredoxin | GO:0006805 xenobiotic metabolism | gg | o |
| *treA* | CDS | periplasmic trehalase | Periplasmic | COG1626;Neutral trehalase | GO:0006006 glucose metabolism -!- GO:0006970 response to osmotic stress | o | tt |
| *treB* | CDS | fused trehalose(maltose)-specific enzyme IIBC component of PTS | Integral Membrane Protein | COG1264;Phosphotransferase system IIB components -!- COG1263;Phosphotransferase system IIC components, glucose/maltose/N-acetylglucosamine-specific | GO:0006006 glucose metabolism | o | tt |
| *treC* | CDS | trehalose-6-P hydrolase | Cytoplasmic | COG0366;Glycosidases | GO:0016052 carbohydrate | o | tt |
| *treF* | CDS | cytoplasmic trehalase | Cytoplasmic | COG1626;Neutral trehalase | GO:0016052 carbohydrate | o | tt |
| *treR* | CDS | DNA-binding transcriptional repressor | Cytoplasmic | COG1609;Transcriptional regulators | GO:0006006 glucose metabolism -!- GO:0006350 transcription -!- GO:0006970 response to osmotic stress | o | tt |
| *trg* | CDS | methyl-accepting chemotaxis protein III, ribose and galactose sensor receptor | Integral Membrane Protein | COG0840;Methyl-accepting chemotaxis protein | GO:0042330 taxis | o | tt |
| *trkA* | CDS | NAD-binding component of TrK potassium | Cytoplasmic | COG0569;K+ transport systems, NAD-binding component | | g | t |
| *trkD* | CDS | potassium transporter | Integral Membrane Protein | | GO:0006970 response to osmotic stress | | |
| *trkG* | CDS | Rac prophage; potassium transporter subunit | Integral Membrane Protein | COG0168;Trk-type K+ transport systems, membrane components | | o | tt |
| *trkH* | CDS | potassium transporter | Integral Membrane Protein | | | | |
| *trmA* | CDS | tRNA (uracil-5-)-methyltransferase | Cytoplasmic | COG2265;SAM-dependent methyltransferases related to tRNA (uracil-5-)-methyltransferase | GO:0009451 RNA modification | gg | tt |
| *trmC* | CDS | fused 5-methylaminomethyl-2-thiouridine forming enzyme methyltransferase -!- FAD-dependent demodification enzyme | Cytoplasmic | | | | |
| *trmD* | CDS | tRNA (guanine-1-)-methyltransferase | Cytoplasmic | COG0336;tRNA-(guanine-N1)-methyltransferase | GO:0009451 RNA modification | gg | o |
| *trmE* | CDS | GTPase | Cytoplasmic | COG0486;Predicted GTPase | GO:0009451 RNA modification -!- GO:0006805 xenobiotic metabolism | gg | tt |
| *trmH* | CDS | tRNA (Guanosine-2'-O-)-methyltransferase | Cytoplasmic | COG0566;rRNA methylases | GO:0009451 RNA modification | gg | t |
| *trmU* | CDS | tRNA (5-methylaminomethyl-2-thiouridylate)-methyltransferase | Cytoplasmic | | GO:0009451 RNA modification | o | tt |
| *trpA* | CDS | tryptophan synthase, alpha subunit | Cytoplasmic | COG0159;Tryptophan synthase alpha chain | GO:0000162 tryptophan biosynthesis | gg | o |
| *trpB* | CDS | tryptophan synthase, beta subunit | Cytoplasmic | COG0133;Tryptophan synthase beta chain | GO:0000162 tryptophan biosynthesis | gg | tt |
| *trpC* | CDS | fused indole-3-glycerolphosphate synthetase -!- N-(5-phosphoribosyl)anthranilate isomerase | Cytoplasmic | COG0134;Indole-3-glycerol phosphate synthase -!- COG0135;Phosphoribosylanthranilate isomerase | GO:0000162 tryptophan biosynthesis | o | tt |

| | | | | | | | |
|---|---|---|---|---|---|---|---|
| *trpD* | CDS | fused glutamine amidotransferase (component II) of anthranilate synthase -!- anthranilate phosphoribosyl transferase | Cytoplasmic | COG0512;Anthranilate/para-aminobenzoate synthases component II -!- COG0547;Anthranilate phosphoribosyltransferase | GO:0000162 tryptophan biosynthesis | gg | t |
| *trpE* | CDS | component I of anthranilate synthase | Cytoplasmic | COG0147;Anthranilate/para-aminobenzoate synthases component I | GO:0000162 tryptophan biosynthesis | o | tt |
| *trpL* | CDS | trp operon leader peptide | Cytoplasmic | | GO:0000162 tryptophan biosynthesis | gg | o |
| *trpR* | CDS | DNA-binding transcriptional repressor, tryptophan-binding | Cytoplasmic | COG2973;Trp operon repressor | GO:0000162 tryptophan biosynthesis -!- GO:0006350 transcription | o | tt |
| *trpS* | CDS | tryptophanyl-tRNA synthetase | Cytoplasmic | COG0180;Tryptophanyl-tRNA synthetase | GO:0006418 amino acid activation | gg | tt |
| *trpT* | tRNA | tRNA-Trp(CCA) (Tryptophan tRNA) | | | | | |
| *truA* | CDS | pseudouridylate synthase I | Cytoplasmic | COG0101;Pseudouridylate synthase | GO:0006418 amino acid activation | o | tt |
| *truB* | CDS | tRNA pseudouridine synthase | Cytoplasmic | COG0130;Pseudouridine synthase | GO:0009451 RNA modification | gg | tt |
| *truD* | CDS | pseudoruidine synthase | Cytoplasmic | COG0585;Uncharacterized conserved protein | | gg | o |
| *trxA* | CDS | thioredoxin 1 | Cytoplasmic | COG0526;Thiol-disulfide isomerase and thioredoxins | | | |
| *trxB* | CDS | thioredoxin reductase, FAD/NAD(P)-binding | Cytoplasmic | COG0492;Thioredoxin reductase | | gg | t |
| *trxC* | CDS | thioredoxin 2 | Cytoplasmic | COG0526;Thiol-disulfide isomerase and thioredoxins | | gg | tt |
| *tsf* | CDS | protein chain elongation factor EF-Ts | Cytoplasmic | COG0264;Translation elongation factor Ts | GO:0006412 protein biosynthesis | gg | o |
| *tsgA* | CDS | predicted transporter | Integral Membrane Protein | COG0477;Permeases of the major facilitator superfamily | | o | tt |
| *tsr* | CDS | methyl-accepting chemotaxis protein I, serine sensor receptor | Integral Membrane Protein | COG0840;Methyl-accepting chemotaxis protein | GO:0042330 taxis | o | tt |
| *tsx* | CDS | nucleoside channel, receptor of phage T6 and colicin K | Outer Membrane B-barrel protein | COG3248;Nucleoside-binding outer membrane protein | GO:0015949 nucleobase, nucleoside and nucleotide interconversion | o | tt |
| *ttdA* | CDS | L-tartrate dehydratase, alpha subunit | Cytoplasmic | COG1951;Tartrate dehydratase alpha subunit/Fumarate hydratase class I, N-terminal domain | GO:0006113 fermentation | g | t |
| *ttdB* | CDS | L-tartrate dehydratase, beta subunit | Cytoplasmic | COG1838;Tartrate dehydratase beta subunit/Fumarate hydratase class I, C-terminal domain | GO:0006113 fermentation | o | t |
| *ttk* | CDS | division inhibitor | Cytoplasmic | | GO:0006350 transcription | | |
| *tufA* | CDS | protein chain elongation factor EF-Tu (duplicate of tufB) | Cytoplasmic | COG0050;GTPases - translation elongation factors | GO:0006412 protein biosynthesis -!- GO:0006970 response to osmotic stress | o | t |
| *tufB* | CDS | protein chain elongation factor EF-Tu (duplicate of tufA) | Cytoplasmic | COG0050;GTPases - translation elongation factors | GO:0006412 protein biosynthesis -!- GO:0006970 response to osmotic stress | o | tt |
| *tus* | CDS | inhibitor of replication at Ter, DNA-binding protein | Cytoplasmic | | GO:0006261 DNA dependent DNA replication | gg | t |
| *tynA* | CDS | tyramine oxidase, copper-requiring | Periplasmic | COG3733;Cu2+-containing amine oxidase | GO:0009063 amino acid catabolism -!- GO:0009310 amine catabolism | o | tt |
| *tyrA* | CDS | fused chorismate mutase T -!- prephenate dehydrogenase | Cytoplasmic | COG1605;Chorismate mutase -!- COG0287;Prephenate dehydrogenase | GO:0009094 L-phenylalanine biosynthesis -!- GO:0006571 tyrosine biosynthesis | gg | o |
| *tyrB* | CDS | tyrosine aminotransferase, tyrosine-repressible, PLP-dependent | Cytoplasmic | COG1448;Aspartate/tyrosine/aromatic aminotransferase | GO:0009094 L-phenylalanine biosynthesis -!- GO:0006571 tyrosine biosynthesis -!- GO:0009098 leucine biosynthesis | gg | t |
| *tyrP* | CDS | tyrosine transporter | Integral Membrane Protein | COG0814;Amino acid permeases | GO:0006571 tyrosine biosynthesis | o | tt |
| *tyrR* | CDS | DNA-binding transcriptional dual regulator, tyrosine-binding | Cytoplasmic | COG3283;Transcriptional regulator of aromatic amino acids metabolism | GO:0006350 transcription | gg | tt |
| *tyrS* | CDS | tyrosyl-tRNA synthetase | Cytoplasmic | COG0162;Tyrosyl-tRNA synthetase | GO:0006418 amino acid activation | o | tt |
| *tyrT* | tRNA | tRNA-Tyr(GUA) (Tyrosine tRNA1) | | | | | |
| *tyrU* | tRNA | tRNA-Tyr(GUA) (Tyrosine tRNA2) | | | | | |
| *tyrV* | tRNA | tRNA-Tyr(GUA) (Tyrosine tRNA1) | | | | | |
| *ubiA* | CDS | p-hydroxybenzoate octaprenyltransferase | Integral Membrane Protein | COG0382;4-hydroxybenzoate polyprenyltransferase and related prenyltransferases | GO:0009060 aerobic respiration | o | tt |
| *ubiB* | CDS | 2-octaprenylphenol hydroxylase | Integral Membrane Protein | COG0661;Predicted unusual protein kinase | | o | tt |
| *ubiC* | CDS | chorismate pyruvate lyase | Cytoplasmic | | GO:0009060 aerobic respiration | | |
| *ubiD* | CDS | 3-octaprenyl-4-hydroxybenzoate decarboxylase | Cytoplasmic | COG0043;3-polyprenyl-4-hydroxybenzoate decarboxylase and related decarboxylases | | g | t |
| *ubiE* | CDS | bifunctional 2-octaprenyl-6-methoxy-1,4-benzoquinone methylase -!- S-adenosylmethionine:2-DMK methyltransferase | Cytoplasmic | | GO:0009060 aerobic respiration -!- GO:0009061 anaerobic respiration | | |
| *ubiF* | CDS | 2-octaprenyl-3-methyl-6-methoxy-1,4-benzoquinol oxygenase | Cytoplasmic | COG0654;2-polyprenyl-6-methoxyphenol hydroxylase and related FAD-dependent oxidoreductases | | o | tt |
| *ubiG* | CDS | bifunctional 3-demethylubiquinone-9 3-methyltransferase -!- 2-octaprenyl-6-hydroxy phenol methylase | Cytoplasmic | COG2227;2-polyprenyl-3-methyl-5-hydroxy-6-metoxy-1 ,4-benzoquinol methylase | GO:0009060 aerobic respiration | gg | o |
| *ubiH* | CDS | 2-octaprenyl-6-methoxyphenol hydroxylase, FAD/NAD(P)-binding | Cytoplasmic | COG0654;2-polyprenyl-6-methoxyphenol hydroxylase and related FAD-dependent oxidoreductases | GO:0009060 aerobic respiration | gg | t |
| *ubiX* | CDS | 3-octaprenyl-4-hydroxybenzoate carboxy-lyase | Cytoplasmic | COG0163;3-polyprenyl-4-hydroxybenzoate decarboxylase | GO:0009060 aerobic respiration | g | o |
| *ucpA* | CDS | predicted oxidoredutase, sulfate metabolism protein | Cytoplasmic | | | | |
| *udk* | CDS | uridine/cytidine kinase | Cytoplasmic | COG0572;Uridine kinase | GO:0015949 nucleobase, nucleoside and nucleotide interconversion | gg | t |
| *udp* | CDS | uridine phosphorylase | Cytoplasmic | COG2820;Uridine phosphorylase | GO:0015949 nucleobase, nucleoside and nucleotide interconversion | gg | o |
| *ugd* | CDS | UDP-glucose 6-dehydrogenase | Cytoplasmic | COG1004;Predicted UDP-glucose 6-dehydrogenase | GO:0009242 colanic acid biosynthesis -!- GO:0009269 response to dessication | gg | t |
| *ugpA* | CDS | glycerol-3-phosphate transporter subunit -!- membrane component of ABC superfamily | Integral Membrane Protein | COG1175;ABC-type sugar transport systems, permease components | GO:0009060 aerobic respiration -!- GO:0009061 anaerobic respiration -!- GO:0006071 glycerol metabolism -!- GO:0008654 phospholipid biosynthesis | o | tt |
| *ugpB* | CDS | glycerol-3-phosphate transporter subunit -!- periplasmic-binding component of ABC superfamily | Periplasmic | COG1653;ABC-type sugar transport system, periplasmic component | GO:0009060 aerobic respiration -!- GO:0009061 anaerobic respiration -!- GO:0006071 glycerol metabolism -!- GO:0008654 phospholipid biosynthesis | o | tt |
| *ugpC* | CDS | glycerol-3-phosphate transporter subunit -!- ATP-binding component of ABC superfamily | Cytoplasmic | COG3839;ABC-type sugar transport systems, ATPase components | GO:0009060 aerobic respiration -!- GO:0009061 anaerobic respiration -!- GO:0006071 glycerol metabolism -!- GO:0008654 phospholipid biosynthesis | g | o |
| *ugpE* | CDS | glycerol-3-phosphate transporter subunit -!- membrane component of ABC superfamily | Integral Membrane Protein | COG0395;ABC-type sugar transport system, permease component | GO:0009060 aerobic respiration -!- GO:0009061 anaerobic respiration -!- GO:0006071 glycerol metabolism -!- GO:0008654 phospholipid biosynthesis | o | tt |
| *ugpQ* | CDS | glycerophosphodiester phosphodiesterase, cytosolic | Cytoplasmic | COG0584;Glycerophosphoryl diester phosphodiesterase | GO:0006071 glycerol metabolism | gg | t |
| *uhpA* | CDS | DNA-binding response regulator in two-component regulatory system wtih UhpB | Cytoplasmic | COG2197;Response regulator containing a CheY-like receiver domain and an HTH DNA-binding domain | GO:0016052 carbohydrate catabolism -!- GO:0006350 | o | tt |
| *uhpB* | CDS | sensory histidine kinase in two-component regulatory sytem with UhpA | Integral Membrane Protein | | GO:0016052 carbohydrate catabolism -!- GO:0006464 protein | o | tt |
| *uhpC* | CDS | membrane protein regulates uhpT expression | Integral Membrane Protein | COG2271;Sugar phosphate permease | GO:0016052 carbohydrate catabolism -!- GO:0006350 | o | tt |
| *uhpT* | CDS | hexose phosphate transporter | Integral Membrane Protein | COG2271;Sugar phosphate permease | GO:0016052 carbohydrate | o | tt |
| *uidA* | CDS | beta-D-glucuronidase | Cytoplasmic | COG3250;Beta-galactosidase/beta-glucuronidase | GO:0016052 carbohydrate | gg | tt |
| *uidB* | CDS | glucuronide transporter | Integral Membrane Protein | COG2211;Na+/melibiose symporter and related transporters | GO:0016052 carbohydrate | o | tt |
| *uidC* | CDS | predicted outer membrane porin protein | Outer Membrane B-barrel protein | | | o | tt |
| *uidR* | CDS | DNA-binding transcriptional repressor | Cytoplasmic | COG1309;Transcriptional regulator | GO:0016052 carbohydrate catabolism -!- GO:0006350 | o | tt |
| *ulaA* | CDS | L-ascorbate-specific enzyme IIC component of PTS | Integral Membrane Protein | | GO:0009401 phosphoenolpyruvate-dependent sugar phosphotransferase system | | |
| *ulaB* | CDS | L-ascorbate-specific enzyme IIB component of PTS | Cytoplasmic | COG3414;Phosphotransferase system, galactitol-specific IIB component | GO:0009401 phosphoenolpyruvate-dependent sugar phosphotransferase system | o | t |
| *ulaC* | CDS | L-ascorbate-specific enzyme IIA component of PTS | Cytoplasmic | COG1762;Phosphotransferase system mannitol/fructose-specific IIA domain (Ntr-type) | | gg | o |
| *ulaD* | CDS | 3-keto-L-gulonate 6-phosphate decarboxylase | Cytoplasmic | COG0269;3-hexulose-6-phosphate synthase and related proteins | | g | t |
| *ulaE* | CDS | L-xylulose 5-phosphate 3-epimerase | Cytoplasmic | COG3623;Putative L-xylulose-5-phosphate 3-epimerase | | o | tt |
| *ulaF* | CDS | L-ribulose 5-phosphate 4-epimerase | Cytoplasmic | COG0235;Ribulose-5-phosphate 4-epimerase and related epimerases and aldolases | | gg | t |
| *ulaG* | CDS | L-ascorbate 6-phosphate lactonase | Cytoplasmic | COG2220;Predicted Zn-dependent hydrolases of the beta-lactamase fold | | | |
| *ulaR* | CDS | DNA-binding transriptional dual regulator | Cytoplasmic | COG1349;Transcriptional regulators of sugar metabolism | GO:0006350 transcription | o | tt |
| *umuC* | CDS | DNA polymerase V, subunit C | Cytoplasmic | COG0389;Nucleotidyltransferase/DNA polymerase involved in DNA repair | GO:0006261 DNA dependent DNA replication -!- GO:0006281 DNA repair -!- GO:0009432 SOS response | o | tt |
| *umuD* | CDS | DNA polymerase V, subunit D | Cytoplasmic | COG1974;SOS-response transcriptional repressors (RecA-mediated autopeptidases) | GO:0006261 DNA dependent DNA replication -!- GO:0006281 DNA repair -!- GO:0006350 transcription -!- GO:0009432 SOS response | g | o |
| *ung* | CDS | uracil-DNA-glycosylase | Cytoplasmic | COG0692;Uracil DNA glycosylase | GO:0006281 DNA repair | gg | t |
| *upp* | CDS | uracil phosphoribosyltransferase | Cytoplasmic | COG0035;Uracil phosphoribosyltransferase | GO:0015949 nucleobase, nucleoside and nucleotide interconversion | gg | t |
| *uppS* | CDS | NAD kinase | Cytoplasmic | COG0061;Predicted sugar kinase | | g | t |
| *uraA* | CDS | uracil transporter | Integral Membrane Protein | COG2233;Xanthine/uracil permeases | GO:0015949 nucleobase, nucleoside and nucleotide interconversion | o | tt |
| *usg* | CDS | predicted semialdehyde dehydrogenase | Cytoplasmic | COG0136;Aspartate-semialdehyde dehydrogenase | | gg | tt |
| *ushA* | CDS | bifunctional UDP-sugar hydrolase -!- 5'-nucleotidase | Periplasmic | COG0737;5'-nucleotidase/2',3'-cyclic phosphodiesterase and related esterases | GO:0009226 nucleotide-sugar biosynthesis | o | tt |
| *uspA* | CDS | universal stress global response regulator | Cytoplasmic | COG0589;Universal stress protein UspA and related nucleotide-binding proteins | GO:0006950 response to stress | gg | o |

| | | | | | | | |
|---|---|---|---|---|---|---|---|
| *uspE* | CDS | stress-induced protein | Cytoplasmic | COG0589;Universal stress protein UspA and related nucleotide-binding proteins | | gg | tt |
| *uspF* | CDS | stress-induced protein, ATP-binding protein | Cytoplasmic | | | gg | t |
| *uspG* | CDS | universal stress protein UP12 | Cytoplasmic | COG0589;Universal stress protein UspA and related nucleotide-binding proteins | | gg | o |
| *uup* | CDS | fused predicted transporter subunits -!- ATP-binding components of ABC superfamily | Cytoplasmic | COG0488;ATPase components of ABC transporters with duplicated ATPase domains | | gg | tt |
| *uvrA* | CDS | ATPase and DNA damage recognition protein of nucleotide excision repair excinuclease UvrABC | Cytoplasmic | COG0178;Excinuclease ATPase subunit | GO:0006281 DNA repair -!- GO:0009314 response to radiation -!- GO:0009432 SOS response | o | tt |
| *uvrB* | CDS | excinulease of nucleotide excision repair, DNA damage recognition component | Cytoplasmic | COG0556;Helicase subunit of the DNA excision repair complex | GO:0006308 DNA catabolism -!- GO:0006281 DNA repair -!- GO:0006308 DNA catabolism -!- GO:0009314 response to radiation -!- GO:0009432 SOS response | gg | t |
| *uvrC* | CDS | excinuclease UvrABC, endonuclease subunit | Cytoplasmic | | GO:0006308 DNA catabolism -!- GO:0006281 DNA repair -!- GO:0006308 DNA catabolism -!- GO:0009314 response to radiation | o | tt |
| *uvrD* | CDS | DNA-dependent ATPase I and helicase II | Cytoplasmic | COG0210;Superfamily I DNA and RNA helicases | GO:0006261 DNA dependent DNA replication -!- GO:0006281 DNA repair -!- GO:0009432 SOS response | o | tt |
| *uvrY* | CDS | DNA-binding response regulator in two-component regulatory system with BarA | Cytoplasmic | COG2197;Response regulator containing a CheY-like receiver domain and an HTH DNA-binding domain | GO:0006350 transcription | gg | t |
| *uxaA* | CDS | altronate hydrolase | Cytoplasmic | COG2721;Altronate dehydratase | GO:0016052 carbohydrate | gg | t |
| *uxaB* | CDS | altronate oxidoreductase, NAD-dependent | Cytoplasmic | | GO:0016052 carbohydrate | gg | t |
| *uxaC* | CDS | uronate isomerase | Cytoplasmic | COG1904;Glucuronate isomerase | GO:0016052 carbohydrate | o | o |
| *uxuA* | CDS | mannonate hydrolase | Cytoplasmic | COG1312;D-mannonate dehydratase | GO:0016052 carbohydrate | gg | o |
| *uxuB* | CDS | D-mannonate oxidoreductase, NAD-binding | Cytoplasmic | COG0246;Mannitol-1-phosphate/altronate dehydrogenases | GO:0016052 carbohydrate | gg | t |
| *uxuR* | CDS | DNA-binding transcriptional repressor | Cytoplasmic | COG2186;Transcriptional regulators | GO:0016052 carbohydrate catabolism -!- GO:0006350 | o | tt |
| *vacJ* | CDS | predicted lipoprotein | Outer Membrane Lipoprotein | COG2853;Surface lipoprotein | | gg | t |
| *valS* | CDS | valyl-tRNA synthetase | Cytoplasmic | COG0525;Valyl-tRNA synthetase | GO:0006418 amino acid activation | gg | tt |
| *valT* | tRNA | tRNA-Val(UAC) (Valine tRNA1) | | | | | |
| *valU* | tRNA | tRNA-Val(UAC) (Valine tRNA1) | | | | | |
| *valV* | tRNA | tRNA-Val(GAC) (Valine tRNA2B) | | | | | |
| *valW* | tRNA | tRNA-Val(GAC) (Valine tRNA2A) | | | | | |
| *valX* | tRNA | tRNA-Val(UAC), (Valine tRNA1) | | | | | |
| *valY* | tRNA | tRNA-Val(UAC) (Valine tRNA1) | | | | | |
| *valZ* | tRNA | tRNA-Val(UAC) (Valine tRNA1) | | | | | |
| *visC* | CDS | predicted oxidoreductase, FAD/NAD(P)-binding domain | Membrane Lipoprotein | COG0654;2-polyprenyl-6-methoxyphenol hydroxylase and related FAD-dependent oxidoreductases | | gg | t |
| *vsr* | CDS | DNA mismatch endonuclease of very short patch repair | Cytoplasmic | COG3727;DNA G:T-mismatch repair endonuclease | GO:0006308 DNA catabolism -!- GO:0006281 DNA repair -!- GO:0006308 DNA catabolism | gg | o |
| *wbbH* | CDS | O-antigen polymerase | Integral Membrane Protein | | GO:0009243 O antigen biosynthesis -!- GO:0009243 O antigen biosynthesis -!- GO:0009103 lipopolysaccharide biosynthesis | o | tt |
| *wbbI* | CDS | conserved protein | Cytoplasmic | | | o | tt |
| *wbbJ* | CDS | predicted acyl transferase | Cytoplasmic | COG0110;Acetyltransferase (isoleucine patch superfamily) | GO:0009103 lipopolysaccharide biosynthesis | g | tt |
| *wbbK* | CDS | lipopolysaccharide biosynthesis protein | Cytoplasmic | COG0438;Glycosyltransferase | | o | tt |
| *wbbL* | CDS | lipopolysaccharide biosynthesis protein, C-ter fragment (pseudogene) | Cytoplasmic | | | o | o |
| *wbbL* | ancestral | lipopolysaccharide biosynthesis protein | Cytoplasmic | | | | |
| *wbbL* | CDS | lipopolysaccharide biosynthesis protein, N-ter fragment (pseudogene) | Cytoplasmic | | | g | o |
| *wcaA* | CDS | predicted glycosyl transferase | Cytoplasmic | COG0463;Glycosyltransferases involved in cell wall biogenesis | GO:0009242 colanic acid | gg | tt |
| *wcaB* | CDS | predicted acyl transferase | Cytoplasmic | COG1045;Serine acetyltransferase | GO:0009242 colanic acid | o | tt |
| *wcaC* | CDS | predicted glycosyl transferase | Cytoplasmic | COG0438;Glycosyltransferase | GO:0009242 colanic acid | gg | t |
| *wcaD* | CDS | predicted colanic acid polymerase | Integral Membrane Protein | | | o | tt |
| *wcaE* | CDS | predicted glycosyl transferase | Cytoplasmic | COG0463;Glycosyltransferases involved in cell wall biogenesis | GO:0009242 colanic acid | gg | tt |
| *wcaF* | CDS | predicted acyl transferase | Cytoplasmic | COG0110;Acetyltransferase (isoleucine patch superfamily) | GO:0009242 colanic acid | o | tt |
| *wcaI* | CDS | predicted glycosyl transferase | Cytoplasmic | COG0438;Glycosyltransferase | GO:0009242 colanic acid | gg | tt |
| *wcaJ* | CDS | predicted UDP-glucose lipid carrier transferase | Integral Membrane Protein | COG2148;Sugar transferases involved in lipopolysaccharide synthesis | GO:0009242 colanic acid | o | tt |
| *wcaK* | CDS | predicted pyruvyl transferase | Cytoplasmic | COG2327;Uncharacterized conserved protein | GO:0009242 colanic acid | gg | tt |
| *wcaL* | CDS | predicted glycosyl transferase | Cytoplasmic | COG0438;Glycosyltransferase | GO:0009242 colanic acid | gg | t |
| *wcaM* | CDS | predicted colanic acid biosynthesis protein | Periplasmic | | GO:0009242 colanic acid | gg | t |
| *wrbA* | CDS | predicted flavoprotein in Trp regulation | Cytoplasmic | COG0655;Multimeric flavodoxin WrbA | GO:0000162 tryptophan biosynthesis | gg | t |
| *wza* | CDS | lipoprotein required for capsular polysaccharide translocation through the outer membrane | Outer Membrane Lipoprotein | COG1596;Periplasmic protein involved in polysaccharide export | GO:0009242 colanic acid biosynthesis | o | tt |
| *wzb* | CDS | protein-tyrosine phosphatase | Cytoplasmic | COG0394;Protein-tyrosine-phosphatase | GO:0009242 colanic acid | gg | o |
| *wzc* | CDS | protein-tyrosine kinase | Integral Membrane Protein | COG3206;Uncharacterized protein involved in exopolysaccharide biosynthesis -!- COG0489;ATPases involved in chromosome partitioning | GO:0009242 colanic acid biosynthesis -!- GO:0006464 protein | o | tt |
| *wzxC* | CDS | colanic acid exporter | Integral Membrane Protein | COG2244;Membrane protein involved in the export of O-antigen and teichoic acid | | g | tt |
| *wzxE* | CDS | O-antigen translocase | Integral Membrane Protein | COG2244;Membrane protein involved in the export of O-antigen and teichoic acid | GO:0017004 cytochrome biogenesis | o | tt |
| *wzyE* | CDS | predicted Wzy protein involved in ECA polysaccharide chain elongation | Integral Membrane Protein | | GO:0009246 enterobacterial common antigen biosynthesis | o | tt |
| *wzzE* | CDS | Entobacterial Common Antigen (ECA) polysaccharide chain length modulation protein | Integral Membrane Protein | COG3765;Chain length determinant protein | GO:0009246 enterobacterial common antigen biosynthesis | | |
| *xapA* | CDS | purine nucleoside phosphorylase II | Cytoplasmic | COG0005;Purine nucleoside phosphorylase | GO:0015949 nucleobase, nucleoside and nucleotide interconversion | g | o |
| *xapB* | CDS | xanthosine transporter | Integral Membrane Protein | COG0477;Permeases of the major facilitator superfamily | GO:0016052 carbohydrate | o | tt |
| *xapR* | CDS | DNA-binding transcriptional activator | Cytoplasmic | COG0583;Transcriptional regulator | GO:0015949 nucleobase, nucleoside and nucleotide interconversion -!- GO:0006350 transcription | o | t |
| *xdhA* | CDS | xanthine dehydrogenase, molybdenum binding subunit | Cytoplasmic | COG1529;Aerobic-type carbon monoxide dehydrogenase, large subunit CoxL/CutL homologs | | | |
| *xdhB* | CDS | xanthine dehydrogenase, FAD-binding subunit | Cytoplasmic | COG1319;Aerobic-type carbon monoxide dehydrogenase, middle subunit CoxM/CutM homologs | | gg | t |
| *xdhC* | CDS | xanthine dehydrogenase, Fe-S binding subunit | Cytoplasmic | COG2080;Aerobic-type carbon monoxide dehydrogenase, small subunit CoxS/CutS homologs | | gg | t |
| *xdhD* | CDS | fused predicted xanthine/hypoxanthine oxidase, molybdopterin-binding subunit -!- Fe-S binding subunit | Cytoplasmic | COG2080;Aerobic-type carbon monoxide dehydrogenase, small subunit CoxS/CutS homologs -!- COG1529;Aerobic-type carbon monoxide dehydrogenase, large subunit CoxL/CutL homologs | | o | tt |
| *xerC* | CDS | site-specific tyrosine recombinase | Cytoplasmic | COG4973;Site-specific recombinase XerC | GO:0006310 DNA recombination | o | tt |
| *xerD* | CDS | site-specific tyrosine recombinase | Cytoplasmic | COG4974;Site-specific recombinase XerD | GO:0006310 DNA recombination | o | tt |
| *xseA* | CDS | exonuclease VII, large subunit | Cytoplasmic | COG1570;Exonuclease VII, large subunit | GO:0006308 DNA catabolism -!- GO:0006308 DNA catabolism | gg | tt |
| *xseB* | CDS | exonuclease VII small subunit | Cytoplasmic | COG1722;Exonuclease VII small subunit | GO:0006308 DNA catabolism -!- GO:0006281 DNA repair -!- GO:0006308 DNA catabolism | gg | t |
| *xthA* | CDS | exonuclease III | Cytoplasmic | COG0708;Exonuclease III | GO:0006308 DNA catabolism -!- GO:0006308 DNA catabolism -!- GO:0009314 response to radiation | gg | o |
| *xylA* | CDS | D-xylose isomerase | Cytoplasmic | COG2115;Xylose isomerase | GO:0016052 carbohydrate catabolism -!- GO:0006006 glucose | g | t |
| *xylB* | CDS | xylulokinase | Cytoplasmic | COG1070;Sugar (pentulose and hexulose) kinases | GO:0016052 carbohydrate | o | tt |
| *xylE* | CDS | D-xylose transporter | Integral Membrane Protein | COG0477;Permeases of the major facilitator superfamily | GO:0016052 carbohydrate catabolism -!- GO:0015753 D-xylose | o | tt |
| *xylF* | CDS | D-xylose transporter subunit -!- periplasmic-binding component of ABC superfamily | Periplasmic | COG4213;ABC-type xylose transport system, periplasmic component | GO:0016052 carbohydrate catabolism | o | tt |
| *xylG* | CDS | fused subunits of D-xylose transporter -!- ATP-binding components of ABC superfamily | Cytoplasmic | COG1129;ABC-type sugar transport system, ATPase component | GO:0016052 carbohydrate catabolism | o | t |
| *xylH* | CDS | D-xylose transporter subunit -!- membrane component of ABC superfamily | Integral Membrane Protein | COG4214;ABC-type xylose transport system, permease component | GO:0016052 carbohydrate catabolism | o | tt |
| *xylR* | CDS | DNA-binding transcriptional activator, xylose-binding | Cytoplasmic | COG1609;Transcriptional regulators -!- COG4753;Response regulator containing CheY-like receiver domain and AraC-type DNA-binding domain | GO:0016052 carbohydrate catabolism -!- GO:0006350 | gg | t |
| *yaaA* | CDS | conserved protein | Cytoplasmic | COG3022;Uncharacterized protein conserved in bacteria | | g | t |
| *yaaH* | CDS | conserved inner membrane protein associated with acetate transport | Integral Membrane Protein | COG1584;Predicted membrane protein | | o | tt |
| *yaaI* | CDS | predicted protein | Periplasmic | | | o | tt |
| *yaaJ* | CDS | predicted transporter | Integral Membrane Protein | COG1115;Na+/alanine symporter | | o | tt |
| *yaaU* | CDS | predicted transporter | Integral Membrane Protein | COG0477;Permeases of the major facilitator superfamily | | o | tt |
| *yaaW* | CDS | conserved protein | Cytoplasmic | COG4735;Uncharacterized protein conserved in bacteria | | | |
| *yaaX* | CDS | predicted protein | Periplasmic | | | o | tt |
| *yaaY* | CDS | predicted protein | Membrane Anchored | | | | |
| *yabI* | CDS | conserved inner membrane protein | Integral Membrane Protein | COG0586;Uncharacterized membrane-associated protein | | | |
| *yabP* | CDS | predicted protein | Cytoplasmic | | | g | o |
| *yabQ* | CDS | predicted protein | Cytoplasmic | | | gg | o |
| *yacC* | CDS | predicted protein | Periplasmic | | | o | tt |
| *yacF* | CDS | conserved protein | Cytoplasmic | COG4582;Uncharacterized protein conserved in bacteria | | gg | t |
| *yacG* | CDS | conserved protein | Cytoplasmic | COG3024;Uncharacterized protein conserved in bacteria | | | |
| *yacH* | CDS | predicted protein | Periplasmic | | | gg | t |
| *yacL* | CDS | conserved protein | Cytoplasmic | COG3112;Uncharacterized protein conserved in bacteria | | gg | t |
| *yadB* | CDS | glutamyl-Q tRNA(Asp) synthetase | Cytoplasmic | COG0008;Glutamyl- and glutaminyl-tRNA synthetases | GO:0006418 amino acid activation | | |
| *yadC* | CDS | predicted fimbrial-like adhesin protein | Periplasmic | COG3539;P pilus assembly protein, pilin FimA | | o | tt |

| | | | | | | | |
|---|---|---|---|---|---|---|---|
| *yadD* | CDS | predicted transposase | Cytoplasmic | | | | |
| *yadE* | CDS | predicted polysaccharide deacetylase lipoprotein | Periplasmic | COG0726;Predicted xylanase/chitin deacetylase | | o | tt |
| *yadG* | CDS | predicted transporter subunit -!- ATP-binding component of ABC superfamily | Cytoplasmic | COG1131;ABC-type multidrug transport system, ATPase component | | gg | tt |
| *yadH* | CDS | predicted transporter subunit -!- membrane component of ABC superfamily | Integral Membrane Protein | COG0842;ABC-type multidrug transport system, permease component | | o | tt |
| *yadI* | CDS | predicted PTS Enzyme IIA | Cytoplasmic | COG2893;Phosphotransferase system, mannose/fructose-specific component IIA | GO:0009401 phosphoenolpyruvate-dependent sugar phosphotransferase system | gg | o |
| *yadK* | CDS | predicted fimbrial-like adhesin protein | Periplasmic | | | o | tt |
| *yadL* | CDS | predicted fimbrial-like adhesin protein | Periplasmic | | | o | tt |
| *yadM* | CDS | predicted fimbrial-like adhesin protein | Periplasmic | | | o | tt |
| *yadN* | CDS | predicted fimbrial-like adhesin protein | Periplasmic | COG3539;P pilus assembly protein, pilin FimA | | o | tt |
| *yadR* | CDS | conserved protein | Cytoplasmic | COG0316;Uncharacterized conserved protein | | gg | t |
| *yadS* | CDS | conserved inner membrane protein | Integral Membrane Protein | COG2860;Predicted membrane protein | | o | tt |
| *yaeB* | CDS | conserved protein | Cytoplasmic | COG1720;Uncharacterized conserved protein | | gg | o |
| *yaeF* | CDS | predicted lipoprotein | Outer Membrane Lipoprotein | | | | |
| *yaeH* | CDS | conserved protein | Cytoplasmic | | | o | tt |
| *yaeI* | CDS | predicted phosphatase | Membrane Anchored | COG1408;Predicted phosphohydrolases | | | |
| *yaeJ* | CDS | conserved protein | Cytoplasmic | COG1186;Protein chain release factor B | | gg | o |
| *yaeL* | CDS | zinc metallopeptidase | Integral Membrane Protein | COG0750;Predicted membrane-associated Zn-dependent proteases 1 | | o | tt |
| *yaeP* | CDS | conserved protein | Cytoplasmic | | | gg | o |
| *yaeQ* | CDS | conserved protein | Cytoplasmic | COG4681;Uncharacterized protein conserved in bacteria | | o | tt |
| *yaeR* | CDS | predicted lyase | Cytoplasmic | | | gg | t |
| *yaeT* | CDS | conserved protein | Outer Membrane B-barrel protein | COG4775;Outer membrane protein/protective antigen OMA87 | | o | tt |
| *yafC* | CDS | predicted DNA-binding transcriptional regulator | Cytoplasmic | COG0583;Transcriptional regulator | GO:0006350 transcription | o | tt |
| *yafD* | CDS | conserved protein | Cytoplasmic | COG3021;Uncharacterized protein conserved in bacteria | | | |
| *yafE* | CDS | predicted S-adenosyl-L-methionine-dependent methyltransferase | Cytoplasmic | COG0500;SAM-dependent methyltransferases | GO:0009102 biotin biosynthesis | o | o |
| *yafF* | CDS | conserved protein | Cytoplasmic | | | gg | o |
| *yafJ* | CDS | predicted amidotransfease | Cytoplasmic | COG0121;Predicted glutamine amidotransferase | | gg | t |
| *yafK* | CDS | conserved protein | Periplasmic | COG3034;Uncharacterized protein conserved in bacteria | | o | tt |
| *yafL* | CDS | predicted lipoprotein and C40 family peptidase | Periplasmic | COG0791;Cell wall-associated hydrolases (invasion-associated proteins) | | o | t |
| *yafM* | CDS | conserved protein | Cytoplasmic | COG1943;Transposase and inactivated derivatives | | gg | o |
| *yafN* | CDS | predicted antitoxin of the YafO-YafN toxin-antitoxin system | Cytoplasmic | COG2161;Antitoxin of toxin-antitoxin stability system | | | |
| *yafO* | CDS | predicted toxin of the YafO-YafN toxin-antitoxin | Cytoplasmic | | | gg | o |
| *yafP* | CDS | predicted acyltransferase with acyl-CoA N-acyltransferase domain | Cytoplasmic | COG0454;Histone acetyltransferase HPA2 and related acetyltransferases | | gg | tt |
| *yafQ* | CDS | predicted toxin of the YafQ-DinJ toxin-antitoxin | Cytoplasmic | COG3041;Uncharacterized protein conserved in bacteria | | gg | o |
| *yafS* | CDS | predicted S-adenosyl-L-methionine-dependent methyltransferase | Cytoplasmic | COG0500;SAM-dependent methyltransferases | | o | tt |
| *yafT* | CDS | predicted aminopeptidase | Outer Membrane Lipoprotein | | | o | tt |
| *yafU* | CDS | predicted inner membrane protein | Integral Membrane Protein | | | | |
| *yafV* | CDS | predicted C-N hydrolase family amidase, NAD(P)-binding | Cytoplasmic | COG0388;Predicted amidohydrolase | | | |
| *yafW* | CDS | CP4-6 prophage; antitoxin of the YkfI-YafW toxin-antitoxin system | Cytoplasmic | | | o | o |
| *yafX* | CDS | CP4-6 prophage; predicted protein | Cytoplasmic | | | | |
| *yafY* | CDS | CP4-6 prophage; predicted DNA-binding transcriptional regulator | Cytoplasmic | | | o | tt |
| *yafZ* | CDS | CP4-6 prophage; conserved protein | Cytoplasmic | | | g | o |
| *yagA* | CDS | CP4-6 prophage; predicted DNA-binding transcriptional regulator | Cytoplasmic | COG2801;Transposase and inactivated derivatives | | o | tt |
| *yagB* | CDS | CP4-6 prophage; conserved protein | Cytoplasmic | | | o | o |
| *yagE* | CDS | CP4-6 prophage; predicted lyase/synthase | Cytoplasmic | COG0329;Dihydrodipicolinate synthase/N-acetylneuraminate lyase | | gg | tt |
| *yagF* | CDS | CP4-6 prophage; predicted dehydratase | Cytoplasmic | COG0129;Dihydroxyacid dehydratase/phosphogluconate dehydratase | | | |
| *yagG* | CDS | CP4-6 prophage; predicted sugar transporter | Integral Membrane Protein | COG2211;Na+/melibiose symporter and related transporters | | o | tt |
| *yagH* | CDS | CP4-6 prophage; predicted xylosidase/arabinosidase | Cytoplasmic | COG3507;Beta-xylosidase | | gg | tt |
| *yagI* | CDS | CP4-6 prophage; predicted DNA-binding transcriptional regulator | Cytoplasmic | COG1414;Transcriptional regulator | GO:0006350 transcription | o | tt |
| *yagJ* | CDS | CP4-6 prophage; predicted protein | Cytoplasmic | | | gg | o |
| *yagK* | CDS | CP4-6 prophage; conserved protein | Cytoplasmic | | | o | tt |
| *yagL* | CDS | CP4-6 prophage; DNA-binding protein | Cytoplasmic | COG1961;Site-specific recombinases, DNA invertase Pin homologs | | o | tt |
| *yagM* | CDS | CP4-6 prophage; predicted protein | Cytoplasmic | | | o | o |
| *yagN* | CDS | CP4-6 prophage; predicted protein | Cytoplasmic | | | | |
| *yagP* | CDS | predicted transcriptional regulator | Cytoplasmic | COG0583;Transcriptional regulator | GO:0006355 regulation of transcription, DNA-dependent | o | tt |
| *yagQ* | CDS | conserved protein | Cytoplasmic | COG1975;Xanthine and CO dehydrogenases maturation factor, XdhC/CoxF family | | gg | tt |
| *yagR* | CDS | predicted oxidoreductase with molybdenum-binding domain | Periplasmic | COG1529;Aerobic-type carbon monoxide dehydrogenase, large subunit CoxL/CutL homologs | | gg | tt |
| *yagS* | CDS | predicted oxidoreductase with FAD-binding domain | Membrane Anchored | COG1319;Aerobic-type carbon monoxide dehydrogenase, middle subunit CoxM/CutM homologs | | o | tt |
| *yagT* | CDS | predicted xanthine dehydrogenase, 2Fe-2S subunit | Cytoplasmic | COG2080;Aerobic-type carbon monoxide dehydrogenase, small subunit CoxS/CutS homologs | | gg | o |
| *yagU* | CDS | conserved inner membrane protein | Integral Membrane Protein | COG3477;Predicted periplasmic/secreted protein | | o | tt |
| *yagV* | CDS | conserved protein | Periplasmic | | | | |
| *yagW* | CDS | predicted receptor | Periplasmic | | | o | tt |
| *yagX* | CDS | predicted aromatic compound dioxygenase | Periplasmic | | | o | tt |
| *yagY* | CDS | predicted protein | Periplasmic | | | o | tt |
| *yagZ* | CDS | conserved protein | Periplasmic | | | o | tt |
| *yahA* | CDS | predicted DNA-binding transcriptional regulator | Cytoplasmic | COG2771;DNA-binding HTH domain-containing proteins -!- COG2200;FOG: EAL domain | GO:0006355 regulation of transcription, DNA-dependent | o | tt |
| *yahB* | CDS | predicted DNA-bindng transcriptional regulator | Periplasmic | COG0583;Transcriptional regulator | GO:0006350 transcription | gg | t |
| *yahC* | CDS | predicted inner membrane protein | Integral Membrane Protein | | | o | tt |
| *yahD* | CDS | predicted transcriptional regulator with ankyrin | Cytoplasmic | COG0666;FOG: Ankyrin repeat | | gg | o |
| *yahE* | CDS | predicted protein | Cytoplasmic | | | gg | o |
| *yahF* | CDS | predicted acyl-CoA synthetase with NAD(P)-binding domain and succinyl-CoA synthetase domain | Cytoplasmic | COG0074;Succinyl-CoA synthetase, alpha subunit | | o | tt |
| *yahG* | CDS | conserved protein | Cytoplasmic | | | o | tt |
| *yahH* | CDS | predicted protein | Membrane Anchored | | | | |
| *yahI* | CDS | predicted carbamate kinase-like protein | Cytoplasmic | COG0549;Carbamate kinase | GO:0006526 arginine biosynthesis -!- GO:0006221 pyrimidine nucleotide biosynthesis | gg | t |
| *yahJ* | CDS | predicted deaminase with metallo-dependent hydrolase domain | Periplasmic | COG0402;Cytosine deaminase and related metal-dependent hydrolases | GO:0009310 amine catabolism | gg | t |
| *yahK* | CDS | predicted oxidoreductase, Zn-dependent and NAD(P)-binding | Cytoplasmic | COG1064;Zn-dependent alcohol dehydrogenases | | gg | t |
| *yahL* | CDS | predicted protein | Cytoplasmic | | | o | tt |
| *yahM* | CDS | predicted protein | Cytoplasmic | | | | |
| *yahN* | CDS | neutral amino-acid efflux system | Integral Membrane Protein | COG1280;Putative threonine efflux protein | | o | tt |
| *yahO* | CDS | predicted protein | Periplasmic | | | o | tt |
| *yaiA* | CDS | predicted protein | Cytoplasmic | | | gg | o |
| *yaiB* | CDS | predicted protein | Cytoplasmic | | | gg | o |
| *yaiC* | CDS | predicted diguanylate cyclase | Integral Membrane Protein | COG2199;FOG: GGDEF domain | | o | tt |
| *yaiE* | CDS | conserved protein | Cytoplasmic | COG3123;Uncharacterized protein conserved in bacteria | | gg | o |
| *yaiF* | CDS | predicted protein | Cytoplasmic | | | | |
| *yaiI* | CDS | conserved protein | Cytoplasmic | COG1671;Uncharacterized protein conserved in bacteria | | gg | o |
| *yaiL* | CDS | nucleoprotein/polynucleotide-associated enzyme | Cytoplasmic | | | gg | tt |
| *yaiO* | CDS | predicted protein | Periplasmic | | | o | tt |
| *yaiP* | CDS | predicted glucosyltransferase | Integral Membrane Protein | COG1215;Glycosyltransferases, probably involved in cell wall biogenesis | GO:0000271 polysaccharide biosynthesis | o | tt |
| *yaiS* | CDS | conserved protein | Cytoplasmic | | | o | t |
| *yaiT* | CDS | predicted protein | Periplasmic | | | o | tt |
| *yaiU* | CDS | predicted protein | Outer Membrane B-barrel protein | COG3468;Type V secretory pathway, adhesin AidA | GO:0009296 flagella biogenesis -!- GO:0042330 taxis | | |
| *yaiV* | CDS | predicted DNA-binding transcriptional regulator | Cytoplasmic | | | gg | o |
| *yaiW* | CDS | predicted DNA-binding transcriptional regulator | Outer Membrane Lipoprotein | | | o | tt |
| *yaiX* | CDS | predicted acyl transferase (pseudogene) | Cytoplasmic | COG0110;Acetyltransferase (isoleucine patch superfamily) | | gg | t |
| *yaiY* | CDS | predicted inner membrane protein | Integral Membrane Protein | | | o | tt |
| *yaiZ* | CDS | predicted inner membrane protein | Integral Membrane Protein | | | | |
| *yajB* | CDS | conserved protein | Cytoplasmic | COG3124;Uncharacterized protein conserved in bacteria | | gg | t |
| *yajC* | CDS | SecYEG protein translocase auxillary subunit | Membrane Anchored | COG1862;Preprotein translocase subunit YajC | | o | tt |
| *yajD* | CDS | conserved protein | Cytoplasmic | | | gg | o |
| *yajG* | CDS | predicted lipoprotein | Outer Membrane Lipoprotein | | | o | tt |
| *yajI* | CDS | predicted lipoprotein | Outer Membrane Lipoprotein | | | | |
| *yajL* | CDS | conserved protein | Cytoplasmic | | GO:0009102 biotin biosynthesis | | |
| *yajO* | CDS | predicted oxidoreductase, NAD(P)-binding | Cytoplasmic | COG0667;Predicted oxidoreductases (related to aryl-alcohol dehydrogenases) | | gg | t |
| *yajQ* | CDS | predicted nucleotide binding protein | Cytoplasmic | COG1666;Uncharacterized protein conserved in bacteria | | | |
| *yajR* | CDS | predicted transporter | Integral Membrane Protein | | GO:0042493 response to drug | | |
| *ybaA* | CDS | conserved protein | Cytoplasmic | | | gg | o |
| *ybaB* | CDS | conserved protein | Cytoplasmic | COG0718;Uncharacterized protein conserved in bacteria | | gg | t |

| | | | | | | | |
|---|---|---|---|---|---|---|---|
| *ybaD* | CDS | conserved protein | Cytoplasmic | COG1327;Predicted transcriptional regulator, consists of a Zn-ribbon and ATP-cone domains | | gg | o |
| *ybaE* | CDS | predicted transporter subunit -!- periplasmic-binding component of ABC superfamily | Cytoplasmic | COG4533;ABC-type uncharacterized transport system, periplasmic component | | gg | t |
| *ybaJ* | CDS | predicted protein | Cytoplasmic | | | gg | t |
| *ybaK* | CDS | conserved protein | Cytoplasmic | COG2606;Uncharacterized conserved protein | | gg | o |
| *ybaL* | CDS | predicted transporter with NAD(P)-binding Rossmann-fold domain | Integral Membrane Protein | COG4651;Kef-type K+ transport system, predicted NAD-binding component -!- COG1226;Kef-type K+ transport systems, predicted NAD-binding component | | o | tt |
| *ybaM* | CDS | predicted protein | Cytoplasmic | | | gg | o |
| *ybaN* | CDS | conserved inner membrane protein | Integral Membrane Protein | COG2832;Uncharacterized protein conserved in bacteria | | o | tt |
| *ybaO* | CDS | predicted DNA-binding transcriptional regulator | Cytoplasmic | | GO:0006350 transcription | o | o |
| *ybaP* | CDS | conserved protein | Cytoplasmic | COG3735;Uncharacterized protein conserved in bacteria | | gg | tt |
| *ybaQ* | CDS | predicted DNA-binding transcriptional regulator | Cytoplasmic | | | o | tt |
| *ybaS* | CDS | predicted glutaminase | Cytoplasmic | COG2066;Glutaminase | GO:0009063 amino acid catabolism | gg | t |
| *ybaT* | CDS | predicted transporter | Integral Membrane Protein | COG0531;Amino acid transporters | GO:0009063 amino acid catabolism | o | tt |
| *ybaV* | CDS | conserved protein | Periplasmic | COG1555;DNA uptake protein and related DNA-binding proteins | | o | tt |
| *ybaW* | CDS | conserved protein | Cytoplasmic | COG0824;Predicted thioesterase | | o | tt |
| *ybaX* | CDS | predicted aluminum resistance protein | Cytoplasmic | COG0603;Predicted PP-loop superfamily ATPase | GO:0006805 xenobiotic metabolism | gg | t |
| *ybaY* | CDS | predicted outer membrane lipoprotein | Outer Membrane Lipoprotein | | GO:0042158 lipoprotein biosynthesis -!- GO:0009101 glycoprotein biosynthesis | o | tt |
| *ybaZ* | CDS | predicted methyltransferase | Cytoplasmic | COG3695;Predicted methylated DNA-protein cysteine methyltransferase | | o | tt |
| *ybbA* | CDS | predicted transporter subunit -!- ATP-binding component of ABC superfamily | Cytoplasmic | COG4181;Predicted ABC-type transport system involved in lysophospholipase L1 biosynthesis, ATPase component | | gg | tt |
| *ybbB* | CDS | tRNA 2-selenouridine synthase, selenophosphate-dependent | Cytoplasmic | COG2603;Predicted ATPase | | o | t |
| *ybbC* | CDS | predicted protein | Outer Membrane Lipoprotein | | | o | tt |
| *ybbD* | CDS | predicted protein | Membrane Anchored | | | o | tt |
| *ybbJ* | CDS | conserved inner membrane protein | Integral Membrane Protein | COG1585;Membrane protein implicated in regulation of membrane protease activity | | | |
| *ybbK* | CDS | predicted protease, membrane anchored | Membrane Anchored | COG0330;Membrane protease subunits, stomatin/prohibitin homologs | | o | tt |
| *ybbL* | CDS | predicted transporter subunit -!- ATP-binding component of ABC superfamily | Cytoplasmic | COG4619;ABC-type uncharacterized transport system, ATPase component | | gg | o |
| *ybbM* | CDS | predicted inner membrane protein | Integral Membrane Protein | | GO:0042493 response to drug | | |
| *ybbN* | CDS | predicted thioredoxin domain-containing protein | Cytoplasmic | | | | |
| *ybbO* | CDS | predicted oxidoreductase with NAD(P)-binding Rossmann-fold domain | Cytoplasmic | COG1028;Dehydrogenases with different specificities (related to short-chain alcohol dehydrogenases) | | o | tt |
| *ybbP* | CDS | predicted inner membrane protein | Integral Membrane Protein | COG3127;Predicted ABC-type transport system involved in lysophospholipase L1 biosynthesis, permease component | | o | tt |
| *ybbS* | CDS | transcriptional activator of the allD operon | Cytoplasmic | COG0583;Transcriptional regulator | GO:0006350 transcription | gg | t |
| *ybbV* | CDS | predicted protein | Membrane Anchored | | | | |
| *ybbW* | CDS | predicted allantoin transporter | Integral Membrane Protein | | | o | tt |
| *ybbY* | CDS | predicted uracil/xanthine transporter | Integral Membrane Protein | | | o | tt |
| *ybcC* | CDS | DLP12 prophage; predicted exonuclease | Cytoplasmic | | | o | t |
| *ybcD* | CDS | DLP12 prophage; predicted replication protein fragment (pseudogene) | Cytoplasmic | | | gg | t |
| *ybcF* | CDS | predicted carbamate kinase | Cytoplasmic | COG0549;Carbamate kinase | GO:0006526 arginine biosynthesis -!- GO:0006221 pyrimidine nucleotide biosynthesis | gg | t |
| *ybcH* | CDS | predicted protein | Periplasmic | | | o | tt |
| *ybcI* | CDS | conserved inner membrane protein | Integral Membrane Protein | COG1988;Predicted membrane-bound metal-dependent hydrolases | | o | o |
| *ybcJ* | CDS | predicted RNA-binding protein | Cytoplasmic | COG2501;Uncharacterized conserved protein | | | |
| *ybcK* | CDS | DLP12 prophage; predicted recombinase | Cytoplasmic | COG1961;Site-specific recombinases, DNA invertase Pin homologs | | gg | t |
| *ybcL* | CDS | DLP12 prophage; predicted kinase inhibitor | Periplasmic | COG1881;Phospholipid-binding protein | | o | tt |
| *ybcM* | CDS | DLP12 prophage; predicted DNA-binding transcriptional regulator | Cytoplasmic | COG2207;AraC-type DNA-binding domain-containing proteins | | o | tt |
| *ybcN* | CDS | DLP12 prophage; predicted protein | Cytoplasmic | | | gg | o |
| *ybcO* | CDS | DLP12 prophage; predicted protein | Cytoplasmic | | | gg | o |
| *ybcQ* | CDS | DLP12 prophage; predicted antitermination protein | Cytoplasmic | | | o | tt |
| *ybcS* | CDS | DLP12 prophage; predicted lysozyme | Periplasmic | COG3772;Phage-related lysozyme (muraminidase) | | | |
| *ybcV* | CDS | DLP12 prophage; predicted protein | Cytoplasmic | | | | |
| *ybcW* | CDS | DLP12 prophage; predicted protein | Cytoplasmic | | | | |
| *ybcY* | CDS | DLP12 prophage; predicted SAM-dependent methyltransferase | Cytoplasmic | | | | |
| *ybdA* | CDS | predicted transporter | Integral Membrane Protein | COG0477;Permeases of the major facilitator superfamily | | o | tt |
| *ybdB* | CDS | conserved protein | Cytoplasmic | COG2050;Uncharacterized protein, possibly involved in aromatic compounds catabolism | | gg | t |
| *ybdD* | CDS | conserved protein | Cytoplasmic | | | gg | o |
| *ybdF* | CDS | conserved protein | Cytoplasmic | COG2315;Uncharacterized protein conserved in bacteria | | | |
| *ybdG* | CDS | predicted mechanosensitive channel | Integral Membrane Protein | COG0668;Small-conductance mechanosensitive channel | | o | tt |
| *ybdH* | CDS | predicted oxidoreductase | Cytoplasmic | COG0371;Glycerol dehydrogenase and related enzymes | | gg | o |
| *ybdJ* | CDS | predicted inner membrane protein | Integral Membrane Protein | | | o | tt |
| *ybdK* | CDS | gamma-glutamyl:cysteine ligase | Cytoplasmic | COG2170;Uncharacterized conserved protein | | gg | t |
| *ybdL* | CDS | methionine aminotransferase, PLP-dependent | Cytoplasmic | COG0436;Aspartate/tyrosine/aromatic aminotransferase | GO:0008152 metabolism | gg | o |
| *ybdM* | CDS | conserved protein | Cytoplasmic | COG1475;Predicted transcriptional regulators | | gg | t |
| *ybdN* | CDS | conserved protein | Cytoplasmic | COG3969;Predicted phosphoadenosine phosphosulfate sulfotransferase | | g | o |
| *ybdO* | CDS | predicted DNA-binding transcriptional regulator | Cytoplasmic | COG0583;Transcriptional regulator | GO:0006350 transcription -!- GO:0006355 regulation of transcription, DNA-dependent | g | o |
| *ybdR* | CDS | predicted oxidoreductase, Zn-dependent and NAD(P)-binding | Cytoplasmic | COG1063;Threonine dehydrogenase and related Zn-dependent dehydrogenases | | gg | t |
| *ybdZ* | CDS | conserved protein | Cytoplasmic | | | gg | o |
| *ybeA* | CDS | conserved protein | Cytoplasmic | COG1576;Uncharacterized conserved protein | | gg | t |
| *ybeB* | CDS | predicted protein | Cytoplasmic | | | | |
| *ybeD* | CDS | conserved protein | Cytoplasmic | COG2921;Uncharacterized conserved protein | | o | tt |
| *ybeF* | CDS | predicted DNA-binding transcriptional regulator | Cytoplasmic | | GO:0006350 transcription | g | o |
| *ybeH* | CDS | predicted protein | Cytoplasmic | | | gg | t |
| *ybeL* | CDS | conserved protein | Cytoplasmic | | | gg | t |
| *ybeM* | CDS | predicted C-N hydrolase superfamily, NAD(P)-binding amidase/nitrilase | Cytoplasmic | COG0388;Predicted amidohydrolase | | gg | t |
| *ybeQ* | CDS | conserved protein | Cytoplasmic | | | | |
| *ybeR* | CDS | predicted protein | Cytoplasmic | | | gg | t |
| *ybeT* | CDS | conserved outer membrane protein | Outer Membrane Lipoprotein | COG0790;FOG: TPR repeat, SEL1 subfamily | | o | tt |
| *ybeU* | CDS | predicted tRNA ligase | Cytoplasmic | | | o | tt |
| *ybeX* | CDS | predicteed ion transport | Cytoplasmic | COG4535;Putative Mg2+ and Co2+ transporter CorC | | gg | o |
| *ybeY* | CDS | conserved protein | Cytoplasmic | COG0319;Predicted metal-dependent hydrolase | | gg | tt |
| *ybeZ* | CDS | predicted protein with nucleoside triphosphate hydrolase domain | Cytoplasmic | COG1702;Phosphate starvation-inducible protein PhoH, predicted ATPase | | g | t |
| *ybfA* | CDS | predicted protein | Membrane Anchored | | | o | tt |
| *ybfB* | CDS | predicted inner membrane protein | Integral Membrane Protein | | | o | tt |
| *ybfC* | CDS | predicted protein | Periplasmic | | | o | o |
| *ybfD* | CDS | conserved protein | Cytoplasmic | | | g | t |
| *ybfE* | CDS | LexA regulated protein | Cytoplasmic | | GO:0009432 SOS response | | |
| *ybfF* | CDS | conserved protein | Cytoplasmic | COG0596;Predicted hydrolases or acyltransferases (alpha/beta hydrolase superfamily) | | g | o |
| *ybfG* | CDS | predicted protein | Cytoplasmic | | | | |
| *ybfH* | CDS | predicted protein | Cytoplasmic | | | | |
| *ybfL* | CDS | predicted transposase (pseudogene) | Cytoplasmic | | | o | tt |
| *ybfM* | CDS | predicted outer membrane porin | Outer Membrane B-barrel protein | | | o | o |
| *ybfN* | CDS | predicted lipoprotein | Outer Membrane Lipoprotein | | | o | tt |
| *ybfO* | CDS | conserved protein, rhs-like | Cytoplasmic | COG3209;Rhs family protein | | gg | t |
| *ybfP* | CDS | predicted protein | Outer Membrane Lipoprotein | | | | |
| *ybfQ* | CDS | predicted transposase | Cytoplasmic | | | o | t |
| *ybgA* | CDS | conserved protein | Cytoplasmic | COG3272;Uncharacterized conserved protein | | gg | t |
| *ybgC* | CDS | predicted acyl-CoA thioesterase | Cytoplasmic | COG0824;Predicted thioesterase | | gg | o |
| *ybgD* | CDS | predicted fimbrial-like adhesin protein | Periplasmic | COG3539;P pilus assembly protein, pilin FimA | | o | tt |
| *ybgE* | CDS | conserved inner membrane protein | Integral Membrane Protein | COG3790;Predicted membrane protein | | o | tt |
| *ybgF* | CDS | predicted protein | Periplasmic | COG1729;Uncharacterized protein conserved in bacteria | | o | tt |
| *ybgH* | CDS | predicted transporter | Integral Membrane Protein | COG3104;Dipeptide/tripeptide permease | GO:0009063 amino acid catabolism | o | tt |
| *ybgI* | CDS | conserved metal-binding protein | Cytoplasmic | COG0327;Uncharacterized conserved protein | | gg | tt |
| *ybgJ* | CDS | predicted enzyme subunit | Cytoplasmic | COG2049;Allophanate hydrolase subunit 1 | | g | t |
| *ybgK* | CDS | predicted enzyme subunit | Cytoplasmic | COG1984;Allophanate hydrolase subunit 2 | | gg | t |
| *ybgL* | CDS | predicted lactam utilization protein | Cytoplasmic | COG1540;Uncharacterized proteins, homologs of lactam utilization protein B | GO:0016052 carbohydrate catabolism -!- GO:0042493 response | gg | t |
| *ybgO* | CDS | predicted fimbrial-like adhesin protein | Periplasmic | | | | |
| *ybgP* | CDS | predicted assembly protein | Periplasmic | COG3121;P pilus assembly protein, chaperone PapD | GO:0006457 protein folding | o | tt |
| *ybgQ* | CDS | predicted outer membrane protein | Outer Membrane B-barrel protein | COG3188;P pilus assembly protein, porin PapC | | | |
| *ybgS* | CDS | conserved protein | Periplasmic | | | o | tt |
| *ybgT* | CDS | conserved protein | Cytoplasmic | | | o | tt |
| *ybhA* | CDS | predicted hydrolase | Cytoplasmic | COG0561;Predicted hydrolases of the HAD superfamily | | o | t |
| *ybhB* | CDS | predicted kinase inhibitor | Cytoplasmic | COG1881;Phospholipid-binding protein | | o | t |

| | | | | | | | |
|---|---|---|---|---|---|---|---|
| *ybhC* | CDS | predicted pectinesterase | Outer Membrane Lipoprotein | COG4677;Pectin methylesterase | GO:0008152 metabolism -!- GO:0000272 polysaccharide catabolism | o | tt |
| *ybhD* | CDS | predicted DNA-binding transcriptional regulator | Membrane Anchored | | GO:0006350 transcription | | |
| *ybhE* | CDS | 6-phosphogluconolactonase | Cytoplasmic | COG2706;3-carboxymuconate cyclase | | gg | tt |
| *ybhF* | CDS | fused predicted transporter subunits -!- ATP-binding components of ABC superfamily | Cytoplasmic | | | | |
| *ybhG* | CDS | predicted membrane fusion protein (MFP) component of efflux pump, membrane anchor | Membrane Anchored | COG0845;Membrane-fusion protein | | o | tt |
| *ybhH* | CDS | conserved protein | Cytoplasmic | COG2828;Uncharacterized protein conserved in bacteria | | gg | o |
| *ybhI* | CDS | predicted transporter | Integral Membrane Protein | COG0471;Di- and tricarboxylate transporters | | o | tt |
| *ybhJ* | CDS | predicted hydratase | Cytoplasmic | COG1048;Aconitase A | | | |
| *ybhK* | CDS | predicted transferase with NAD(P)-binding Rossmann-fold domain | Cytoplasmic | COG0391;Uncharacterized conserved protein | | gg | tt |
| *ybhL* | CDS | predicted inner membrane protein | Integral Membrane Protein | COG0670;Integral membrane protein, interacts with FtsH | | o | tt |
| *ybhM* | CDS | conserved inner membrane protein | Integral Membrane Protein | COG0670;Integral membrane protein, interacts with FtsH | | o | o |
| *ybhN* | CDS | conserved inner membrane protein | Integral Membrane Protein | COG0392;Predicted integral membrane protein | | o | tt |
| *ybhO* | CDS | cardiolipin synthase 2 | Cytoplasmic | COG1502;Phosphatidylserine/phosphatidylglycerophos phate/cardiolipin synthases and related enzymes | GO:0008654 phospholipid biosynthesis | gg | t |
| *ybhP* | CDS | predicted DNase | Cytoplasmic | COG3568;Metal-dependent hydrolase | GO:0006308 DNA catabolism | gg | t |
| *ybhQ* | CDS | predicted inner membrane protein | Integral Membrane Protein | | | o | tt |
| *ybhR* | CDS | predicted transporter subunit -!- membrane component of ABC superfamily | Integral Membrane Protein | COG0842;ABC-type multidrug transport system, permease component | | | |
| *ybhS* | CDS | predicted transporter subunit -!- membrane component of ABC superfamily | Integral Membrane Protein | COG0842;ABC-type multidrug transport system, permease component | | o | tt |
| *ybhT* | CDS | predicted protein | Membrane Anchored | | | | |
| *ybiA* | CDS | conserved protein | Cytoplasmic | COG3236;Uncharacterized protein conserved in bacteria | | gg | t |
| *ybiB* | CDS | predicted transferase/phosphorylase | Cytoplasmic | COG0547;Anthranilate phosphoribosyltransferase | | o | tt |
| *ybiC* | CDS | predicted dehydrogenase | Cytoplasmic | COG2055;Malate/L-lactate dehydrogenases | | gg | t |
| *ybiH* | CDS | predicted DNA-binding transcriptional regulator | Cytoplasmic | COG1309;Transcriptional regulator | GO:0006350 transcription | o | tt |
| *ybiI* | CDS | conserved protein | Cytoplasmic | COG1734;DnaK suppressor protein | | gg | o |
| *ybiJ* | CDS | predicted protein | Periplasmic | | | o | tt |
| *ybiM* | CDS | predicted protein | Periplasmic | | | | |
| *ybiN* | CDS | predicted SAM-dependent methyltransferase | Cytoplasmic | COG3129;Predicted SAM-dependent methyltransferase | | | |
| *ybiO* | CDS | predicted mechanosensitive channel | Integral Membrane Protein | | | | |
| *ybiP* | CDS | predicted hydrolase, inner membrane | Integral Membrane Protein | COG2194;Predicted membrane-associated, metal-dependent hydrolase | | o | tt |
| *ybiR* | CDS | predicted transporter | Integral Membrane Protein | COG0471;Di- and tricarboxylate transporters | | o | tt |
| *ybiS* | CDS | conserved protein | Periplasmic | COG1376;Uncharacterized protein conserved in bacteria | | o | tt |
| *ybiT* | CDS | fused predicted transporter subunits -!- ATP-binding components of ABC superfamily | Cytoplasmic | COG0488;ATPase components of ABC transporters with duplicated ATPase domains | | gg | t |
| *ybiU* | CDS | predicted protein | Cytoplasmic | | | gg | tt |
| *ybiV* | CDS | predicted hydrolase | Cytoplasmic | COG0561;Predicted hydrolases of the HAD superfamily | | gg | t |
| *ybiW* | CDS | predicted pyruvate formate lyase | Cytoplasmic | COG1882;Pyruvate-formate lyase | GO:0006113 fermentation | gg | t |
| *ybiX* | CDS | conserved protein | Cytoplasmic | | | | |
| *ybiY* | CDS | predicted pyruvate formate lyase activating enzyme | Cytoplasmic | COG1180;Pyruvate-formate lyase-activating enzyme | GO:0009061 anaerobic respiration -!- GO:0006464 protein modification | gg | t |
| *ybjC* | CDS | predicted inner membrane protein | Integral Membrane Protein | | | o | tt |
| *ybjD* | CDS | conserved protein with nucleoside triphosphate hydrolase domain | Cytoplasmic | COG3593;Predicted ATP-dependent endonuclease of the OLD family | | gg | tt |
| *ybjE* | CDS | predicted transporter | Integral Membrane Protein | COG2431;Predicted membrane protein | | o | tt |
| *ybjG* | CDS | undecaprenyl pyrophosphate phosphatase | Integral Membrane Protein | COG0671;Membrane-associated phospholipid phosphatase | | | |
| *ybjH* | CDS | predicted protein | Periplasmic | | | o | tt |
| *ybjI* | CDS | conserved protein | Cytoplasmic | | | | |
| *ybjJ* | CDS | predicted transporter | Integral Membrane Protein | COG0477;Permeases of the major facilitator superfamily | | o | tt |
| *ybjK* | CDS | predicted DNA-binding transcriptional regulator | Cytoplasmic | COG3226;Uncharacterized protein conserved in bacteria | | | |
| *ybjL* | CDS | predicted transporter | Integral Membrane Protein | COG2985;Predicted permease | | o | tt |
| *ybjM* | CDS | predicted inner membrane protein | Integral Membrane Protein | | | | |
| *ybjN* | CDS | predicted oxidoreductase | Cytoplasmic | | | o | tt |
| *ybjO* | CDS | predicted inner membrane protein | Integral Membrane Protein | | | o | tt |
| *ybjP* | CDS | predicted lipoprotein | Outer Membrane Lipoprotein | | | o | tt |
| *ybjQ* | CDS | conserved protein | Cytoplasmic | COG0393;Uncharacterized conserved protein | | gg | t |
| *ybjR* | CDS | predicted amidase and lipoprotein | Outer Membrane Lipoprotein | COG3023;Negative regulator of beta-lactamase expression | | o | tt |
| *ybjS* | CDS | predicted NAD(P)H oxidoreductase with NAD(P)-binding Rossmann-fold domain | Cytoplasmic | COG0451;Nucleoside-diphosphate-sugar epimerases | | | |
| *ybjT* | CDS | conserved protein with NAD(P)-binding Rossmann-fold domain | Membrane Anchored | | | | |
| *ybjX* | CDS | conserved protein | Cytoplasmic | COG2990;Uncharacterized protein conserved in bacteria | | o | tt |
| *ycaC* | CDS | predicted hydrolase | Cytoplasmic | COG1335;Amidases related to nicotinamidase | GO:0009063 amino acid catabolism | gg | o |
| *ycaD* | CDS | predicted transporter | Integral Membrane Protein | COG0477;Permeases of the major facilitator superfamily | | o | tt |
| *ycaI* | CDS | conserved inner membrane protein | Integral Membrane Protein | COG0658;Predicted membrane metal-binding protein -!- COG2333;Predicted hydrolase (metallo-beta-lactamase superfamily) | | | |
| *ycaJ* | CDS | recombination protein | Cytoplasmic | COG2256;ATPase related to the helicase subunit of the Holliday junction resolvase | | o | tt |
| *ycaK* | CDS | conserved protein | Cytoplasmic | COG2249;Putative NADPH-quinone reductase (modulator of drug activity | | gg | o |
| *ycaL* | CDS | predicted peptidase with chaperone function | Outer Membrane Lipoprotein | COG0501;Zn-dependent protease with chaperone function | GO:0009266 response to temperature | o | tt |
| *ycaM* | CDS | predicted transporter | Integral Membrane Protein | COG0531;Amino acid transporters | | | |
| *ycaN* | CDS | predicted DNA-binding transcriptional regulator | Cytoplasmic | COG0583;Transcriptional regulator | GO:0006350 transcription | gg | t |
| *ycaO* | CDS | conserved protein | Cytoplasmic | | | gg | tt |
| *ycaP* | CDS | conserved inner membrane protein | Integral Membrane Protein | COG2323;Predicted membrane protein | | o | tt |
| *ycaQ* | CDS | conserved protein | Cytoplasmic | COG3214;Uncharacterized protein conserved in bacteria | | o | o |
| *ycaR* | CDS | conserved protein | Cytoplasmic | COG2835;Uncharacterized conserved protein | | gg | o |
| *ycbB* | CDS | predicted carboxypeptidase | Periplasmic | COG2989;Uncharacterized protein conserved in bacteria | | o | tt |
| *ycbC* | CDS | conserved inner membrane protein | Integral Membrane Protein | COG1434;Uncharacterized conserved protein | | o | tt |
| *ycbF* | CDS | predicted periplasmic pilini chaperone | Periplasmic | | GO:0006457 protein folding | | |
| *ycbG* | CDS | conserved protein | Cytoplasmic | COG3120;Uncharacterized protein conserved in bacteria | | o | o |
| *ycbJ* | CDS | conserved protein | Cytoplasmic | | | o | t |
| *ycbK* | CDS | conserved protein | Periplasmic | COG3108;Uncharacterized protein conserved in bacteria | | gg | tt |
| *ycbL* | CDS | predicted metal-binding enzyme | Cytoplasmic | COG0491;Zn-dependent hydrolases, including glyoxylases | | gg | o |
| *ycbQ* | CDS | predicted fimbrial-like adhesin protein | Periplasmic | COG3539;P pilus assembly protein, pilin FimA | | | |
| *ycbR* | CDS | predicted periplasmic pilin chaperone | Periplasmic | COG3121;P pilus assembly protein, chaperone PapD | GO:0006457 protein folding | o | tt |
| *ycbS* | CDS | predicted outer membrane usher protein | Outer Membrane B-barrel protein | COG3188;P pilus assembly protein, porin PapC | | o | tt |
| *ycbT* | CDS | predicted fimbrial-like adhesin protein | Periplasmic | COG3539;P pilus assembly protein, pilin FimA | | o | tt |
| *ycbU* | CDS | predicted fimbrial-like adhesin protein | Periplasmic | COG3539;P pilus assembly protein, pilin FimA | | o | tt |
| *ycbV* | CDS | predicted fimbrial-like adhesin protein | Periplasmic | COG3539;P pilus assembly protein, pilin FimA | | | |
| *ycbW* | CDS | predicted protein | Cytoplasmic | | | | |
| *ycbX* | CDS | predicted 2Fe-2S cluster-containing protein | Cytoplasmic | COG3217;Uncharacterized Fe-S protein -!- COG0633;Ferredoxin | | | |
| *ycbY* | CDS | predicted methyltransferase | Cytoplasmic | COG0116;Predicted N6-adenine-specific DNA methylase -!- COG1092;Predicted SAM-dependent methyltransferases | | gg | tt |
| *ycbZ* | CDS | predicted peptidase | Cytoplasmic | COG1067;Predicted ATP-dependent protease | GO:0006508 proteolysis and peptidolysis | o | t |
| *yccA* | CDS | inner membrane protein | Integral Membrane Protein | COG0670;Integral membrane protein, interacts with FtsH | | o | tt |
| *yccC* | CDS | cryptic autophosphorylating protein tyrosine kinase Etk | Integral Membrane Protein | COG3206;Uncharacterized protein involved in exopolysaccharide biosynthesis -!- COG0489;ATPases involved in chromosome partitioning | GO:0006464 protein modification | o | tt |
| *yccE* | CDS | predicted protein | Cytoplasmic | | | gg | tt |
| *yccF* | CDS | conserved inner membrane protein | Integral Membrane Protein | COG3304;Predicted membrane protein | | o | tt |
| *yccJ* | CDS | predicted protein | Cytoplasmic | | | gg | o |
| *yccK* | CDS | predicted sulfite reductase subunit | Cytoplasmic | | | gg | tt |
| *yccM* | CDS | predicted 4Fe-4S membrane protein | Integral Membrane Protein | COG0348;Polyferredoxin | | o | tt |
| *yccR* | CDS | conserved protein | Cytoplasmic | COG3070;Regulator of competence-specific genes | | o | tt |
| *yccS* | CDS | predicted inner membrane protein | Integral Membrane Protein | COG1289;Predicted membrane protein | | | |
| *yccT* | CDS | conserved protein | Periplasmic | COG3110;Uncharacterized protein conserved in bacteria | | o | tt |
| *yccU* | CDS | predicted CoA-binding protein with NAD(P)-binding Rossmann-fold domain | Cytoplasmic | | | | |
| *yccV* | CDS | DNA-binding protein, hemimethylated | Cytoplasmic | | | | |
| *yccW* | CDS | predicted methyltransferase | Cytoplasmic | | | | |
| *yccX* | CDS | predicted acylphosphatase | Cytoplasmic | COG1254;Acylphosphatases | | | |
| *yccZ* | CDS | predicted exopolysaccharide export protein | Outer Membrane Lipoprotein | COG1596;Periplasmic protein involved in polysaccharide export | | o | tt |
| *ycdB* | CDS | conserved protein | Periplasmic | COG2837;Predicted iron-dependent peroxidase | | gg | t |
| *ycdC* | CDS | predicted DNA-binding transcriptional regulator | Cytoplasmic | COG1309;Transcriptional regulator | GO:0006350 transcription | o | tt |
| *ycdG* | CDS | predicted transporter | Integral Membrane Protein | | | | |
| *ycdH* | CDS | predicted oxidoreductase, flavin:NADH component | Cytoplasmic | | | | |
| *ycdI* | CDS | predicted oxidoreductase | Cytoplasmic | COG0778;Nitroreductase | | gg | o |
| *ycdK* | CDS | conserved protein | Periplasmic | COG0251;Putative translation initiation inhibitor, yjgF family | | g | o |
| *ycdL* | CDS | predicted enzyme | Cytoplasmic | COG1335;Amidases related to nicotinamidase | | | |
| *ycdM* | CDS | predicted monooxygenase | Cytoplasmic | | | gg | t |
| *ycdN* | ancestral | predicted membrane protein (pseudogene) | Cytoplasmic | | | | |
| *ycdN* | CDS | predicted protein, N-ter fragment (pseudogene) | Cytoplasmic | | | | |
| *ycdN* | CDS | predicted protein, C-ter fragment (pseudogene) | Cytoplasmic | | | o | tt |
| *ycdO* | CDS | conserved protein | Periplasmic | COG2822;Predicted periplasmic lipoprotein involved in iron transport | | gg | o |
| *ycdP* | CDS | predicted inner membrane protein | Integral Membrane Protein | | | o | tt |
| *ycdQ* | CDS | predicted glycosyl transferase | Integral Membrane Protein | COG1215;Glycosyltransferases, probably involved in cell wall biogenesis | | o | tt |
| *ycdR* | CDS | predicted enzyme associated with biofilm formation | Outer Membrane Lipoprotein | COG0726;Predicted xylanase/chitin deacetylase | | | |
| *ycdS* | CDS | predicted outer membrane protein | Outer Membrane B-barrel protein | | | o | tt |

| | | | | | | | |
|---|---|---|---|---|---|---|---|
| *ycdT* | CDS | predicted diguanylate cyclase | Integral Membrane Protein | COG2199;FOG: GGDEF domain | | | |
| *ycdU* | CDS | predicted inner membrane protein | Integral Membrane Protein | | | o | tt |
| *ycdW* | CDS | 2-ketoacid reductase | Cytoplasmic | COG0111;Phosphoglycerate dehydrogenase and related dehydrogenases | | | |
| *ycdX* | CDS | predicted zinc-binding hydrolase | Cytoplasmic | COG1387;Histidinol phosphatase and related hydrolases of the PHP family | GO:0006261 DNA dependent DNA replication | gg | o |
| *ycdY* | CDS | conserved protein | Cytoplasmic | COG3381;Uncharacterized component of anaerobic dehydrogenases | | gg | o |
| *ycdZ* | CDS | predicted inner membrane protein | Integral Membrane Protein | | | | |
| *yceA* | CDS | conserved protein | Cytoplasmic | COG1054;Predicted sulfurtransferase | GO:0007049 cell cycle | o | tt |
| *yceB* | CDS | predicted lipoprotein | Outer Membrane Lipoprotein | | | o | tt |
| *yceD* | CDS | conserved protein | Cytoplasmic | COG1399;Predicted metal-binding, possibly nucleic acid-binding protein | | gg | tt |
| *yceF* | CDS | predicted protein | Cytoplasmic | COG0424;Nucleotide-binding protein implicated in inhibition of septum formation | | | |
| *yceG* | CDS | predicted aminodeoxychorismate lyase | Membrane Anchored | COG1559;Predicted periplasmic solute-binding protein | | tt | tt |
| *yceH* | CDS | conserved protein | Cytoplasmic | COG3132;Uncharacterized protein conserved in bacteria | | g | tt |
| *yceI* | CDS | predicted protein | Periplasmic | COG2353;Uncharacterized conserved protein | | o | tt |
| *yceJ* | CDS | predicted cytochrome b561 | Integral Membrane Protein | COG3038;Cytochrome B561 | GO:0017004 cytochrome biogenesis | o | tt |
| *yceK* | CDS | predicted lipoprotein | Membrane Lipoprotein | | | | |
| *yceO* | CDS | predicted protein | Membrane Anchored | | | | |
| *yceP* | CDS | predicted protein | Cytoplasmic | | | | |
| *yceQ* | CDS | predicted protein | Cytoplasmic | | | | |
| *ycfD* | CDS | conserved protein | Cytoplasmic | | | gg | tt |
| *ycfH* | CDS | predicted metallodependent hydrolase | Cytoplasmic | COG0084;Mg-dependent DNase | | gg | o |
| *ycfJ* | CDS | predicted protein | Membrane Anchored | COG3134;Predicted outer membrane lipoprotein | | o | tt |
| *ycfK* | CDS | e14 prophage; predicted protein | Cytoplasmic | | | o | t |
| *ycfL* | CDS | predicted protein | Outer Membrane Lipoprotein | | | o | tt |
| *ycfM* | CDS | predicted outer membrane lipoprotein | Outer Membrane Lipoprotein | COG3417;Collagen-binding surface adhesin SpaP (antigen I/II family) | | | |
| *ycfN* | CDS | thiamin kinase | Cytoplasmic | COG0510;Predicted choline kinase involved in LPS biosynthesis | | gg | o |
| *ycfP* | CDS | conserved protein | Cytoplasmic | | | | |
| *ycfQ* | CDS | predicted DNA-binding transcriptional regulator | Cytoplasmic | | GO:0006355 regulation of transcription, DNA-dependent | | |
| *ycfR* | CDS | predicted protein | Periplasmic | | | o | tt |
| *ycfS* | CDS | conserved protein | Periplasmic | COG1376;Uncharacterized protein conserved in bacteria | | | |
| *ycfT* | CDS | predicted inner membrane protein | Integral Membrane Protein | COG4763;Predicted membrane protein | | gg | tt |
| *ycfX* | CDS | predicted DNA-binding transcriptional regulator | Cytoplasmic | COG1940;Transcriptional regulator/sugar kinase | GO:0006355 regulation of transcription, DNA-dependent | gg | t |
| *ycfZ* | CDS | predicted inner membrane protein | Integral Membrane Protein | | | o | tt |
| *ycgB* | CDS | conserved protein | Cytoplasmic | COG2719;Uncharacterized conserved protein | | o | tt |
| *ycgE* | CDS | predicted DNA-binding transcriptional regulator | Cytoplasmic | COG0789;Predicted transcriptional regulators | GO:0006350 transcription | o | tt |
| *ycgF* | CDS | predicted FAD-binding phosphodiesterase | Cytoplasmic | COG2200;FOG: EAL domain | | gg | o |
| *ycgG* | CDS | conserved inner membrane protein | Integral Membrane Protein | COG2200;FOG: EAL domain | | | |
| *ycgH* | ancestral | predicted protein (pseudogene) | Outer Membrane B-barrel protein | | | | |
| *ycgH* | CDS | predicted protein, N-ter fragment (pseudogene) | Outer Membrane B-barrel protein | | | | |
| *ycgH* | CDS | predicted protein, C-ter fragment (pseudogene) | Outer Membrane B-barrel protein | | | | |
| *ycgI* | CDS | predicted protein | Cytoplasmic | | | | |
| *ycgJ* | CDS | predicted protein | Periplasmic | | | o | tt |
| *ycgK* | CDS | predicted protein | Periplasmic | | | o | tt |
| *ycgL* | CDS | conserved protein | Cytoplasmic | | | gg | t |
| *ycgM* | CDS | predicted isomerase/hydrolase | Cytoplasmic | COG0179;2-keto-4-pentenoate hydratase/2-oxohepta-3-ene-1,7-dioic acid hydratase (catechol pathway) | | o | o |
| *ycgN* | CDS | conserved protein | Cytoplasmic | | | | |
| *ycgR* | CDS | protein involved in flagellar function | Cytoplasmic | | GO:0006355 regulation of transcription, DNA-dependent | gg | t |
| *ycgV* | CDS | predicted adhesin | Cytoplasmic | COG3468;Type V secretory pathway, adhesin AidA | | o | tt |
| *ycgX* | CDS | predicted protein | Cytoplasmic | | | gg | o |
| *ycgY* | CDS | predicted protein | Cytoplasmic | | | gg | o |
| *ycgZ* | CDS | predicted protein | Cytoplasmic | | | gg | o |
| *ychA* | CDS | predicted transcriptional regulator | Cytoplasmic | COG2912;Uncharacterized conserved protein | GO:0006355 regulation of transcription, DNA-dependent | gg | tt |
| *ychE* | CDS | predicted inner membrane protein | Integral Membrane Protein | COG2095;Multiple antibiotic transporter | | o | o |
| *ychF* | CDS | predicted GTP-binding protein | Cytoplasmic | COG0012;Predicted GTPase, probable translation factor | | gg | t |
| *ychG* | CDS | predicted protein, C-ter fragment (pseudogene) | Cytoplasmic | | | gg | t |
| *ychG* | ancestral | predicted protein (pseudogene) | Cytoplasmic | | | | |
| *ychG* | CDS | predicted protein, N-ter fragment (pseudogene) | Cytoplasmic | | | | |
| *ychH* | CDS | predicted inner membrane protein | Integral Membrane Protein | | | o | o |
| *ychJ* | CDS | conserved protein | Cytoplasmic | COG3012;Uncharacterized protein conserved in bacteria | | gg | t |
| *ychM* | CDS | predicted transporter | Integral Membrane Protein | COG0659;Sulfate permease and related transporters (MFS superfamily) | | | |
| *ychN* | CDS | conserved protein | Cytoplasmic | COG1553;Uncharacterized conserved protein involved in intracellular sulfur reduction | | g | t |
| *ychP* | CDS | predicted invasin | Cytoplasmic | | | gg | t |
| *ychQ* | CDS | predicted transcriptional regulator | Integral Membrane Protein | COG3094;Uncharacterized protein conserved in bacteria | GO:0006355 regulation of transcription, DNA-dependent | o | tt |
| *ychS* | CDS | predicted protein | Cytoplasmic | | | | |
| *yciA* | CDS | predicted hydrolase | Cytoplasmic | COG1607;Acyl-CoA hydrolase | | gg | o |
| *yciB* | CDS | predicted inner membrane protein | Integral Membrane Protein | COG2917;Intracellular septation protein A | | o | tt |
| *yciC* | CDS | predicted inner membrane protein | Integral Membrane Protein | | | o | tt |
| *yciE* | CDS | conserved protein | Cytoplasmic | COG3685;Uncharacterized protein conserved in bacteria | | gg | o |
| *yciF* | CDS | conserved protein | Cytoplasmic | COG3685;Uncharacterized protein conserved in bacteria | | gg | t |
| *yciG* | CDS | predicted protein | Cytoplasmic | | | gg | o |
| *yciH* | CDS | conserved protein | Cytoplasmic | COG0023;Translation initiation factor 1 (eIF-1/SUI1) and related proteins | | gg | t |
| *yciI* | CDS | predicted enzyme | Cytoplasmic | COG2350;Uncharacterized protein conserved in bacteria | | gg | o |
| *yciK* | CDS | predicted oxoacyl-(acyl carrier protein) reductase, EmrKY-TolC system | Cytoplasmic | COG1028;Dehydrogenases with different specificities (related to short-chain alcohol dehydrogenases) | | gg | t |
| *yciM* | CDS | conserved protein | Periplasmic | COG2956;Predicted N-acetylglucosaminyl transferase | | gg | tt |
| *yciN* | CDS | predicted protein | Cytoplasmic | | | gg | t |
| *yciO* | CDS | conserved protein | Cytoplasmic | | | | |
| *yciQ* | CDS | predicted inner membrane protein | Integral Membrane Protein | | | | |
| *yciS* | CDS | conserved inner membrane protein | Integral Membrane Protein | COG3771;Predicted membrane protein | | o | tt |
| *yciT* | CDS | predicted DNA-binding transcriptional regulator | Cytoplasmic | COG1349;Transcriptional regulators of sugar metabolism | GO:0006350 transcription | | |
| *yciU* | CDS | predicted protein | Cytoplasmic | | | gg | tt |
| *yciV* | CDS | conserved protein | Cytoplasmic | COG0613;Predicted metal-dependent phosphoesterases (PHP family) | | gg | t |
| *yciW* | CDS | predicted oxidoreductase | Cytoplasmic | COG4950;Uncharacterized protein conserved in bacteria -!- COG2128;Uncharacterized conserved protein | | | |
| *yciX* | CDS | predicted protein, N-ter fragment (pseudogene) | Membrane Anchored | | | | |
| *yciX* | ancestral | predicted protein (pseudogene) | Membrane Anchored | | | | |
| *yciX* | CDS | predicted protein, C-ter fragment (pseudogene) | Cytoplasmic | | | | |
| *ycjD* | CDS | conserved protein | Cytoplasmic | COG2852;Uncharacterized protein conserved in bacteria | | gg | t |
| *ycjF* | CDS | conserved inner membrane protein | Integral Membrane Protein | COG3768;Predicted membrane protein | | gg | tt |
| *ycjG* | CDS | L-Ala-D/L-Glu epimerase | Cytoplasmic | | | o | t |
| *ycjM* | CDS | predicted glucosyltransferase | Cytoplasmic | | GO:0000272 polysaccharide catabolism | gg | t |
| *ycjN* | CDS | predicted sugar transporter subunit -!- periplasmic-binding component of ABC superfamily | Outer Membrane Lipoprotein | COG1653;ABC-type sugar transport system, periplasmic component | | o | tt |
| *ycjO* | CDS | predicted sugar transporter subunit -!- membrane component of ABC superfamily | Integral Membrane Protein | COG1175;ABC-type sugar transport systems, permease components | | o | tt |
| *ycjP* | CDS | predicted sugar transporter subunit -!- membrane component of ABC superfamily | Integral Membrane Protein | COG0395;ABC-type sugar transport system, permease component | | o | tt |
| *ycjQ* | CDS | predicted oxidoreductase, Zn-dependent and NAD(P)-binding | Cytoplasmic | COG1063;Threonine dehydrogenase and related Zn-dependent dehydrogenases | | gg | o |
| *ycjR* | CDS | predicted enzyme | Cytoplasmic | COG1082;Sugar phosphate isomerases/epimerases | | | |
| *ycjS* | CDS | predicted oxidoreductase, NADH-binding | Cytoplasmic | COG0673;Predicted dehydrogenases and related proteins | | o | o |
| *ycjT* | CDS | predicted hydrolase | Cytoplasmic | COG1554;Trehalose and maltose hydrolases (possible phosphorylases) | | o | o |
| *ycjU* | CDS | predicted beta-phosphoglucomutase | Cytoplasmic | COG0637;Predicted phosphatase/phosphohexomutase | | g | t |
| *ycjV* | CDS | predicted sugar transporter subunit -!- ATP-binding component of ABC superfamily | Cytoplasmic | COG3839;ABC-type sugar transport systems, ATPase components | | gg | tt |
| *ycjW* | CDS | predicted DNA-binding transcriptional regulator | Cytoplasmic | COG1609;Transcriptional regulators | GO:0006350 transcription | gg | tt |
| *ycjX* | CDS | conserved protein with nucleoside triphosphate hydrolase domain | Cytoplasmic | COG3106;Predicted ATPase | | ttgg | ttgg |
| *ycjY* | CDS | predicted hydrolase | Cytoplasmic | COG1073;Hydrolases of the alpha/beta superfamily | | | |
| *ycjZ* | CDS | predicted DNA-binding transcriptional regulator | Cytoplasmic | COG0583;Transcriptional regulator | GO:0006350 transcription | gg | t |
| *ydaC* | CDS | Rac prophage; predicted protein | Cytoplasmic | | | gg | o |
| *ydaE* | CDS | Rac prophage; conserved protein | Cytoplasmic | | | gg | o |
| *ydaF* | CDS | Rac prophage; predicted protein | Cytoplasmic | | | gg | o |
| *ydaG* | CDS | Rac prophage; predicted protein | Cytoplasmic | | | | |
| *ydaL* | CDS | conserved protein | Cytoplasmic | COG2840;Uncharacterized protein conserved in bacteria | | gg | t |
| *ydaM* | CDS | predicted diguanylate cyclase, GGDEF domain signalling protein | Cytoplasmic | COG2199;FOG: GGDEF domain | | | |
| *ydaN* | CDS | predicted Zn(II) transporter | Integral Membrane Protein | COG0598;Mg2+ and Co2+ transporters | | o | tt |
| *ydaO* | CDS | predicted C32 tRNA thiolase | Cytoplasmic | COG0037;Predicted ATPase of the PP-loop superfamily implicated in cell cycle control | | gg | o |
| *ydaQ* | CDS | Rac prophage; conserved protein | Cytoplasmic | | | | |
| *ydaS* | CDS | Rac prophage; predicted DNA-binding transcriptional regulator | Periplasmic | COG4197;Uncharacterized protein conserved in bacteria, prophage-related | | o | tt |
| *ydaT* | CDS | Rac prophage; predicted protein | Cytoplasmic | | | o | tt |
| *ydaU* | CDS | Rac prophage; conserved protein | Cytoplasmic | COG3756;Uncharacterized protein conserved in bacteria | | g | tt |

| | | | | | | | |
|---|---|---|---|---|---|---|---|
| *ydaV* | CDS | Rac prophage; predicted DNA replication protein | Cytoplasmic | COG1484;DNA replication protein | | gg | tt |
| *ydaW* | CDS | Rac prophage; predicted DNA-binding protein | Cytoplasmic | | | | |
| *ydaY* | CDS | Rac prophage; predicted protein | Cytoplasmic | | | gg | o |
| *ydbA* | CDS | predicted outer membrane protein, N-ter fragment (pseudogene) | Cytoplasmic | | | | |
| *ydbA* | ancestral | predicted outer membrane protein (pseudogene) | Cytoplasmic | | | | |
| *ydbA* | CDS | predicted outer membrane protein, C-ter fragment (pseudogene) | Cytoplasmic | | | o | tt |
| *ydbC* | CDS | predicted oxidoreductase, NAD(P)-binding | Cytoplasmic | COG0667;Predicted oxidoreductases (related to aryl-alcohol dehydrogenases) | | gg | o |
| *ydbD* | CDS | predicted protein | Periplasmic | | | | |
| *ydbH* | CDS | predicted protein | Membrane Anchored | | | o | tt |
| *ydbJ* | CDS | predicted protein | Outer Membrane Lipoprotein | | | | |
| *ydbK* | CDS | fused predicted Fe-S subunit of pyruvate-flavodoxin oxidoreductase | Cytoplasmic | COG0674;Pyruvate:ferredoxin oxidoreductase and related 2-oxoacid:ferredoxin oxidoreductases, alpha subunit -!- COG1014;Pyruvate:ferredoxin oxidoreductase and related 2-oxoacid:ferredoxin oxidoreductases, gamma subunit -!- COG1013;Pyruvate:ferredoxin oxidoreductase and related 2-oxoacid:ferredoxin oxidoreductases, beta subunit | | o | tt |
| *ydbL* | CDS | conserved protein | Periplasmic | COG3784;Uncharacterized protein conserved in bacteria | | | |
| *ydcA* | CDS | predicted protein | Periplasmic | | | o | tt |
| *ydcC* | CDS | conserved protein | Cytoplasmic | | | gg | t |
| *ydcD* | CDS | predicted protein | Membrane Anchored | | | o | t |
| *ydcE* | CDS | 4-oxalocrotonate tautomerase | Cytoplasmic | COG1942;Uncharacterized protein, 4-oxalocrotonate tautomerase homolog | | gg | o |
| *ydcF* | CDS | conserved protein | Cytoplasmic | COG1434;Uncharacterized conserved protein | | gg | o |
| *ydcH* | CDS | predicted protein | Cytoplasmic | | | | |
| *ydcI* | CDS | predicted DNA-binding transcriptional regulator | Cytoplasmic | COG0583;Transcriptional regulator | GO:0006350 transcription | | |
| *ydcJ* | CDS | conserved protein | Cytoplasmic | COG5383;Uncharacterized protein conserved in bacteria | | gg | t |
| *ydcK* | CDS | predicted enzyme | Cytoplasmic | | | o | tt |
| *ydcL* | CDS | predicted lipoprotein | Outer Membrane Lipoprotein | | | o | tt |
| *ydcM* | CDS | predicted transposase | Cytoplasmic | COG0675;Transposase and inactivated derivatives | | | |
| *ydcN* | CDS | predicted DNA-binding transcriptional regulator | Cytoplasmic | COG1396;Predicted transcriptional regulators | | g | t |
| *ydcO* | CDS | predicted benzoate transporter | Integral Membrane Protein | | | | |
| *ydcP* | CDS | predicted peptidase | Cytoplasmic | COG0826;Collagenase and related proteases | | gg | t |
| *ydcQ* | CDS | predicted DNA-binding transcriptional regulator | Cytoplasmic | COG1598;Uncharacterized conserved protein | | o | o |
| *ydcR* | CDS | fused predicted DNA-binding transcriptional regulator -!- predicted amino transferase | Cytoplasmic | COG1167;Transcriptional regulators containing a DNA-binding HTH domain and an aminotransferase domain (MocR family) and their eukaryotic orthologs | GO:0006355 regulation of transcription, DNA-dependent | o | tt |
| *ydcS* | CDS | predicted spermidine/putrescine transporter subunit -!- periplasmic-binding component of ABC superfamily | Periplasmic | COG0687;Spermidine/putrescine-binding periplasmic protein | | gg | tt |
| *ydcT* | CDS | predicted spermidine/putrescine transporter subunit -!- ATP-binding component of ABC superfamily | Cytoplasmic | COG3842;ABC-type spermidine/putrescine transport systems, ATPase components | | gg | o |
| *ydcU* | CDS | predicted spermidine/putrescine transporter subunit -!- membrane component of ABC superfamily | Integral Membrane Protein | COG1176;ABC-type spermidine/putrescine transport system, permease component I | | o | tt |
| *ydcV* | CDS | predicted spermidine/putrescine transporter subunit -!- membrane component of ABC superfamily | Integral Membrane Protein | COG1177;ABC-type spermidine/putrescine transport system, permease component II | | o | tt |
| *ydcW* | CDS | medium chain aldehyde dehydrogenase | Cytoplasmic | COG1012;NAD-dependent aldehyde dehydrogenases | | o | tt |
| *ydcX* | CDS | predicted inner membrane protein | Integral Membrane Protein | | | | |
| *ydcY* | CDS | predicted protein | Cytoplasmic | | | gg | t |
| *ydcZ* | CDS | predicted inner membrane protein | Integral Membrane Protein | COG3238;Uncharacterized protein conserved in bacteria | | o | tt |
| *yddA* | CDS | fused predicted multidrug transporter subunits -!- membrane component and ATP-binding component of ABC superfamily | Integral Membrane Protein | COG4178;ABC-type uncharacterized transport system, permease and ATPase components | | | |
| *yddB* | CDS | predicted porin protein | Periplasmic | | | o | tt |
| *yddE* | CDS | conserved protein | Cytoplasmic | COG0384;Predicted epimerase, PhzC/PhzF homolog | | gg | o |
| *yddG* | CDS | predicted methyl viologen efflux pump | Integral Membrane Protein | | | o | tt |
| *yddH* | CDS | conserved protein | Cytoplasmic | COG1853;Conserved protein/domain typically associated with flavoprotein oxygenases, DIM6/NTAB family | | gg | o |
| *yddJ* | CDS | predicted protein | Cytoplasmic | | | | |
| *yddK* | CDS | predicted protein | Cytoplasmic | COG4886;Leucine-rich repeat (LRR) protein | GO:0009101 glycoprotein biosynthesis | o | o |
| *yddL* | CDS | predicted lipoprotein | Periplasmic | COG3203;Outer membrane protein (porin) | | o | tt |
| *yddM* | CDS | predicted DNA-binding transcriptional regulator | Cytoplasmic | | GO:0006355 regulation of transcription, DNA-dependent | | |
| *yddV* | CDS | predicted diguanylate cyclase | Cytoplasmic | COG2199;FOG: GGDEF domain | | | |
| *yddW* | CDS | predicted liprotein | Outer Membrane Lipoprotein | COG1649;Uncharacterized protein conserved in bacteria | | gg | tt |
| *ydeA* | CDS | predicted arabinose transporter | Integral Membrane Protein | COG2814;Arabinose efflux permease | GO:0016052 carbohydrate | g | t |
| *ydeE* | CDS | predicted transporter | Integral Membrane Protein | COG0477;Permeases of the major facilitator superfamily | | o | tt |
| *ydeH* | CDS | conserved protein | Cytoplasmic | COG2199;FOG: GGDEF domain | | gg | t |
| *ydeI* | CDS | conserved protein | Periplasmic | COG3111;Uncharacterized conserved protein | | g | tt |
| *ydeJ* | CDS | conserved protein | Cytoplasmic | COG1546;Uncharacterized protein (competence- and mitomycin-induced) | | gg | t |
| *ydeK* | CDS | predicted lipoprotein | Outer Membrane Lipoprotein | | | | |
| *ydeM* | CDS | conserved protein | Cytoplasmic | | | gg | o |
| *ydeN* | CDS | conserved protein | Periplasmic | COG3119;Arylsulfatase A and related enzymes | GO:0006790 sulfur metabolism | | |
| *ydeO* | CDS | predicted DNA-binding transcriptional acfivator | Cytoplasmic | COG2207;AraC-type DNA-binding domain-containing proteins | GO:0006350 transcription | gg | tt |
| *ydeP* | CDS | predicted oxidoreductase | Cytoplasmic | COG0243;Anaerobic dehydrogenases, typically selenocysteine-containing | | o | tt |
| *ydeQ* | CDS | predicted fimbrial-like adhesin protein | Periplasmic | | | g | tt |
| *ydeR* | CDS | predicted fimbrial-like adhesin protein | Periplasmic | COG3539;P pilus assembly protein, pilin FimA | | g | tt |
| *ydeS* | CDS | predicted fimbrial-like adhesin protein | Membrane Anchored | COG3539;P pilus assembly protein, pilin FimA | | o | tt |
| *ydeT* | CDS | predicted protein | Cytoplasmic | COG3188;P pilus assembly protein, porin PapC | | gg | t |
| *ydeU* | CDS | conserved protein, predicted pseudogene | Cytoplasmic | COG3468;Type V secretory pathway, adhesin AidA | | gg | t |
| *ydeV* | CDS | predicted sugar kinase | Cytoplasmic | | | gg | tt |
| *ydeW* | CDS | predicted DNA-binding transcriptional regulator | Cytoplasmic | COG2390;Transcriptional regulator, contains sigma factor-related N-terminal domain | | | |
| *ydfA* | CDS | Qin prophage; predicted protein | Cytoplasmic | | | gg | o |
| *ydfB* | CDS | Qin prophage; predicted protein | Cytoplasmic | | | gg | o |
| *ydfC* | CDS | Qin prophage; predicted protein | Cytoplasmic | | | gg | o |
| *ydfD* | CDS | Qin prophage; predicted protein | Cytoplasmic | | | o | tt |
| *ydfE* | CDS | Qin prophage; predicted protein | Cytoplasmic | | | gg | tt |
| *ydfG* | CDS | L-allo-threonine dehydrogenase, NAD(P)-binding | Cytoplasmic | COG4221;Short-chain alcohol dehydrogenase of unknown specificity | | gg | t |
| *ydfH* | CDS | predicted DNA-binding transcriptional regulator | Cytoplasmic | COG1802;Transcriptional regulators | GO:0006350 transcription | o | tt |
| *ydfI* | CDS | predicted mannonate dehydrogenase | Cytoplasmic | COG0246;Mannitol-1-phosphate/altronate dehydrogenases | | g | tt |
| *ydfJ* | CDS | predicted transporter | Integral Membrane Protein | COG0477;Permeases of the major facilitator superfamily | | o | tt |
| *ydfK* | CDS | Qin prophage; predicted DNA-binding transcriptional regulator | Cytoplasmic | | | gg | o |
| *ydfO* | CDS | Qin prophage; predicted protein | Cytoplasmic | | | | |
| *ydfP* | CDS | Qin prophage; conserved protein | Periplasmic | | | o | tt |
| *ydfQ* | CDS | Qin prophage; predicted lysozyme | Membrane Anchored | COG3772;Phage-related lysozyme (muraminidase) | | o | tt |
| *ydfR* | CDS | Qin prophage; predicted protein | Cytoplasmic | | | g | o |
| *ydfT* | CDS | Qin prophage; predicted antitermination protein Q | Cytoplasmic | | | o | tt |
| *ydfU* | CDS | Qin prophage; predicted protein | Cytoplasmic | | | | |
| *ydfV* | CDS | Qin prophage; predicted protein | Cytoplasmic | | | o | t |
| *ydfW* | CDS | Qin prophage; predicted protein | Cytoplasmic | | | gg | o |
| *ydfX* | CDS | Qin prophage; predicted protein | Cytoplasmic | | | gg | o |
| *ydfZ* | CDS | conserved protein | Cytoplasmic | | | gg | o |
| *ydgA* | CDS | conserved protein | Periplasmic | COG5339;Uncharacterized protein conserved in bacteria | | o | tt |
| *ydgC* | CDS | conserved inner membrane protein associated with alginate biosynthesis | Integral Membrane Protein | COG3136;Uncharacterized membrane protein required for alginate biosynthesis | | o | tt |
| *ydgD* | CDS | predicted peptidase | Periplasmic | COG3591;V8-like Glu-specific endopeptidase | | o | tt |
| *ydgG* | CDS | predicted inner membrane protein | Integral Membrane Protein | COG0628;Predicted permease | | o | tt |
| *ydgH* | CDS | predicted protein | Periplasmic | | | o | tt |
| *ydgI* | CDS | predicted arginine/ornithine antiporter transporter | Integral Membrane Protein | COG0531;Amino acid transporters | | o | tt |
| *ydgJ* | CDS | predicted oxidoreductase | Cytoplasmic | | | | |
| *ydgK* | CDS | conserved inner membrane protein | Integral Membrane Protein | | | o | tt |
| *ydgR* | CDS | predicted transporter | Integral Membrane Protein | COG3104;Dipeptide/tripeptide permease | | o | tt |
| *ydgT* | CDS | predicted regulator | Cytoplasmic | | | gg | o |
| *ydhA* | CDS | predicted lipoprotein | Outer Membrane Lipoprotein | | | gg | o |
| *ydhB* | CDS | predicted DNA-binding transcriptional regulator | Cytoplasmic | COG0583;Transcriptional regulator | GO:0006350 transcription | g | t |
| *ydhC* | CDS | predicted transporter | Integral Membrane Protein | | | gg | tt |
| *ydhD* | CDS | conserved protein | Cytoplasmic | COG0278;Glutaredoxin-related protein | | gg | o |
| *ydhF* | CDS | predicted oxidoreductase | Cytoplasmic | | | gg | t |
| *ydhH* | CDS | conserved protein | Cytoplasmic | COG2377;Predicted molecular chaperone distantly related to HSP70-fold metalloproteases | | gg | t |
| *ydhI* | CDS | predicted inner membrane protein | Integral Membrane Protein | | | | |
| *ydhJ* | CDS | undecaprenyl pyrophosphate phosphatase | Membrane Anchored | | | o | tt |
| *ydhK* | CDS | conserved inner membrane protein | Integral Membrane Protein | COG1289;Predicted membrane protein | | o | tt |
| *ydhL* | CDS | conserved protein | Cytoplasmic | COG3313;Predicted Fe-S protein | | | |
| *ydhM* | CDS | predicted DNA-binding transcriptional regulator | Cytoplasmic | COG1309;Transcriptional regulator | GO:0006355 regulation of transcription, DNA-dependent | | |
| *ydhO* | CDS | predicted lipoprotein | Periplasmic | COG0791;Cell wall-associated hydrolases (invasion-associated proteins) | | | |
| *ydhP* | CDS | predicted transporter | Integral Membrane Protein | COG2814;Arabinose efflux permease | | gg | tt |
| *ydhQ* | CDS | conserved protein | Cytoplasmic | COG3468;Type V secretory pathway, adhesin AidA | | gg | t |
| *ydhR* | CDS | predicted protein | Cytoplasmic | | | gg | o |
| *ydhS* | CDS | conserved protein with FAD/NAD(P)-binding domain | Cytoplasmic | COG4529;Uncharacterized protein conserved in bacteria | | gg | t |
| *ydhT* | CDS | conserved protein | Cytoplasmic | | | g | t |

| | | | | | | | |
|---|---|---|---|---|---|---|---|
| *ydhU* | CDS | predicted cytochrome | Integral Membrane Protein | COG4117;Thiosulfate reductase cytochrome B subunit (membrane anchoring protein) | | o | tt |
| *ydhV* | CDS | predicted oxidoreductase | Cytoplasmic | COG2414;Aldehyde:ferredoxin oxidoreductase | | | |
| *ydhW* | CDS | predicted protein | Cytoplasmic | | | gg | t |
| *ydhX* | CDS | predicted 4Fe-4S ferridoxin-type protein | Periplasmic | COG0437;Fe-S-cluster-containing hydrogenase components 1 | | | |
| *ydhY* | CDS | predicted 4Fe-4S ferridoxin-type protein | Cytoplasmic | COG0437;Fe-S-cluster-containing hydrogenase components 1 | | o | t |
| *ydhZ* | CDS | predicted protein | Cytoplasmic | | | gg | o |
| *ydiA* | CDS | conserved protein | Cytoplasmic | COG1806;Uncharacterized protein conserved in bacteria | | gg | o |
| *ydiB* | CDS | quinate/shikimate 5-dehydrogenase, NAD(P)-binding | Cytoplasmic | COG0169;Shikimate 5-dehydrogenase | | gg | o |
| *ydiD* | CDS | short chain acyl-CoA synthetase, anaerobic | Cytoplasmic | | | | |
| *ydiE* | CDS | conserved protein | Cytoplasmic | COG4256;Hemin uptake protein | | gg | o |
| *ydiF* | CDS | fused predicted acetyl-CoA:acetoacetyl-CoA transferase: alpha subunit -!- beta subunit | Cytoplasmic | COG4670;Acyl CoA:acetate/3-ketoacid CoA transferase | GO:0019395 fatty acid oxidation | g | t |
| *ydiH* | CDS | predicted protein | Cytoplasmic | | | o | t |
| *ydiI* | CDS | conserved protein | Cytoplasmic | COG2050;Uncharacterized protein, possibly involved in aromatic compounds catabolism | | g | t |
| *ydiJ* | CDS | predicted FAD-linked oxidoreductase | Cytoplasmic | COG0277;FAD/FMN-containing dehydrogenases -!- COG0247;Fe-S oxidoreductase | | o | tt |
| *ydiK* | CDS | predicted inner membrane protein | Integral Membrane Protein | COG0628;Predicted permease | | o | tt |
| *ydiL* | CDS | conserved protein | Cytoplasmic | | | gg | o |
| *ydiM* | CDS | predicted transporter | Integral Membrane Protein | COG0477;Permeases of the major facilitator superfamily | | o | tt |
| *ydiN* | CDS | predicted transporter | Integral Membrane Protein | | | | |
| *ydiO* | CDS | predicted acyl-CoA dehydrogenase | Cytoplasmic | | | | |
| *ydiP* | CDS | predicted DNA-binding transcriptional regulator | Cytoplasmic | COG2207;AraC-type DNA-binding domain-containing proteins | GO:0006350 transcription -!- GO:0006355 regulation of transcription, DNA-dependent | gg | o |
| *ydiQ* | CDS | conserved protein | Cytoplasmic | COG2086;Electron transfer flavoprotein, beta subunit | | | |
| *ydiR* | CDS | predicted electron transfer flavoprotein, FAD-binding | Cytoplasmic | COG2025;Electron transfer flavoprotein, alpha subunit | | gg | t |
| *ydiS* | CDS | predicted oxidoreductase, FAD/NAD(P)-binding domain | Cytoplasmic | COG0644;Dehydrogenases (flavoproteins) | | gg | t |
| *ydiT* | CDS | predicted 4Fe-4S ferredoxin-type protein | Cytoplasmic | COG2440;Ferredoxin-like protein | | gg | o |
| *ydiU* | CDS | conserved protein | Cytoplasmic | COG0397;Uncharacterized conserved protein | | gg | t |
| *ydiV* | CDS | conserved protein | Cytoplasmic | COG2200;FOG: EAL domain | | g | o |
| *ydiY* | CDS | conserved protein | Periplasmic | COG3137;Putative salt-induced outer membrane protein | | o | tt |
| *ydiZ* | CDS | predicted protein | Cytoplasmic | | | gg | t |
| *ydjA* | CDS | predicted oxidoreductase | Cytoplasmic | COG0778;Nitroreductase | | gg | t |
| *ydjE* | CDS | predicted transporter | Integral Membrane Protein | COG0477;Permeases of the major facilitator superfamily | | o | tt |
| *ydjF* | CDS | predicted DNA-binding transcriptional regulator | Cytoplasmic | COG1349;Transcriptional regulators of sugar metabolism | GO:0006350 transcription -!- GO:0006355 regulation of transcription, DNA-dependent | o | t |
| *ydjG* | CDS | predicted oxidoreductase | Periplasmic | COG0667;Predicted oxidoreductases (related to aryl-alcohol dehydrogenases) | | gg | t |
| *ydjH* | CDS | predicted kinase | Cytoplasmic | | | | |
| *ydjI* | CDS | predicted aldolase | Cytoplasmic | COG0191;Fructose/tagatose bisphosphate aldolase | | gg | t |
| *ydjJ* | CDS | predicted oxidoreductase, Zn-dependent and NAD(P)-binding | Cytoplasmic | COG1063;Threonine dehydrogenase and related Zn-dependent dehydrogenases | | gg | t |
| *ydjK* | CDS | predicted transporter | Integral Membrane Protein | COG0477;Permeases of the major facilitator superfamily | | | |
| *ydjL* | CDS | predicted oxidoreductase, Zn-dependent and NAD(P)-binding | Cytoplasmic | COG1063;Threonine dehydrogenase and related Zn-dependent dehydrogenases | | o | t |
| *ydjM* | CDS | predicted inner membrane protein regulated by LexA | Integral Membrane Protein | | GO:0009432 SOS response | | |
| *ydjN* | CDS | predicted transporter | Integral Membrane Protein | COG1823;Predicted Na+/dicarboxylate symporter | | o | tt |
| *ydjO* | CDS | predicted protein | Cytoplasmic | | | gg | o |
| *ydjQ* | CDS | endonuclease of nucleotide excision repair | Cytoplasmic | COG0322;Nuclease subunit of the excinuclease complex | | gg | t |
| *ydjR* | CDS | conserved protein | Cytoplasmic | COG3758;Uncharacterized protein conserved in bacteria | | g | o |
| *ydjX* | CDS | predicted inner membrane protein | Integral Membrane Protein | | | o | tt |
| *ydjY* | CDS | predicted protein | Inner Membrane Lipoprotein | | | | |
| *ydjZ* | CDS | conserved inner membrane protein | Integral Membrane Protein | COG0398;Uncharacterized conserved protein | | o | tt |
| *yeaA* | CDS | methionine sulfoxide reductase B | Cytoplasmic | COG0229;Conserved domain frequently associated with peptide methionine sulfoxide reductase | | gg | o |
| *yeaB* | CDS | predicted NUDIX hydrolase | Cytoplasmic | COG0494;NTP pyrophosphohydrolases including oxidative damage repair enzymes | | g | o |
| *yeaC* | CDS | conserved protein | Cytoplasmic | | | gg | o |
| *yeaD* | CDS | conserved protein | Cytoplasmic | | | gg | o |
| *yeaE* | CDS | predicted oxidoreductase | Cytoplasmic | COG0656;Aldo/keto reductases, related to diketogulonate reductase | | gg | t |
| *yeaG* | CDS | conserved protein with nucleoside triphosphate hydrolase domain | Cytoplasmic | COG2766;Putative Ser protein kinase | | gg | tt |
| *yeaH* | CDS | conserved protein | Cytoplasmic | COG2718;Uncharacterized conserved protein | | gg | tt |
| *yeaI* | CDS | predicted diguanylate cyclase | Integral Membrane Protein | COG2199;FOG: GGDEF domain | | o | tt |
| *yeaJ* | CDS | predicted diguanylate cyclase | Integral Membrane Protein | COG2199;FOG: GGDEF domain | | | |
| *yeaK* | CDS | conserved protein | Cytoplasmic | COG2606;Uncharacterized conserved protein | | gg | o |
| *yeaL* | CDS | conserved inner membrane protein | Integral Membrane Protein | COG2707;Predicted membrane protein | | o | tt |
| *yeaM* | CDS | predicted DNA-binding transcriptional regulator | Cytoplasmic | COG2207;AraC-type DNA-binding domain-containing proteins | GO:0006350 transcription -!- GO:0006355 regulation of transcription, DNA-dependent | gg | t |
| *yeaN* | CDS | predicted transporter | Integral Membrane Protein | COG2807;Cyanate permease | GO:0042886 amide transport | o | tt |
| *yeaO* | CDS | conserved protein | Cytoplasmic | COG3189;Uncharacterized conserved protein | | gg | t |
| *yeaP* | CDS | predicted diguanylate cyclase | Cytoplasmic | COG2203;FOG: GAF domain -!- COG2199;FOG: GGDEF domain | | | |
| *yeaQ* | CDS | conserved inner membrane protein | Integral Membrane Protein | COG2261;Predicted membrane protein | | o | tt |
| *yeaR* | CDS | conserved protein | Cytoplasmic | COG3615;Uncharacterized protein/domain, possibly involved in tellurite resistance | GO:0006805 xenobiotic metabolism | gg | o |
| *yeaS* | CDS | neutral amino-acid efflux system | Integral Membrane Protein | COG1280;Putative threonine efflux protein | | o | tt |
| *yeaT* | CDS | predicted DNA-binding transcriptional regulator | Cytoplasmic | | GO:0006350 transcription | g | t |
| *yeaU* | CDS | predicted dehydrogenase | Cytoplasmic | COG0473;Isocitrate/isopropylmalate dehydrogenase | GO:0006113 fermentation | gg | t |
| *yeaV* | CDS | predicted transporter | Integral Membrane Protein | COG1292;Choline-glycine betaine transporter | | | |
| *yeaW* | CDS | predicted 2Fe-2S cluster-containing protein | Cytoplasmic | COG4638;Phenylpropionate dioxygenase and related ring-hydroxylating dioxygenases, large terminal subunit | | | |
| *yeaX* | CDS | predicted oxidoreductase | Cytoplasmic | COG1018;Flavodoxin reductases (ferredoxin-NADPH reductases) family 1 | | gg | t |
| *yeaY* | CDS | predicted lipoprotein | Outer Membrane Lipoprotein | COG3065;Starvation-inducible outer membrane lipoprotein | | o | tt |
| *yeaZ* | CDS | predicted peptidase | Cytoplasmic | COG1214;Inactive homolog of metal-dependent proteases, putative molecular chaperone | | gg | o |
| *yebA* | CDS | predicted peptidase | Cytoplasmic | COG0739;Membrane proteins related to metalloendopeptidases | | | |
| *yebB* | CDS | predicted protein | Cytoplasmic | | | | |
| *yebC* | CDS | conserved protein | Cytoplasmic | COG0217;Uncharacterized conserved protein | | gg | o |
| *yebE* | CDS | conserved protein | Membrane Anchored | COG2979;Uncharacterized protein conserved in bacteria | | gg | t |
| *yebF* | CDS | predicted protein | Periplasmic | | | o | tt |
| *yebG* | CDS | conserved protein regulated by LexA | Cytoplasmic | COG3141;Uncharacterized protein conserved in bacteria | GO:0006281 DNA repair -!- GO:0009432 SOS response | gg | t |
| *yebK* | CDS | predicted DNA-binding transcriptional regulator | Cytoplasmic | COG1737;Transcriptional regulators | GO:0006355 regulation of transcription, DNA-dependent | o | tt |
| *yebN* | CDS | conserved inner membrane protein | Integral Membrane Protein | COG1971;Predicted membrane protein | | | |
| *yebO* | CDS | predicted protein | Membrane Anchored | | | o | t |
| *yebQ* | CDS | predicted transporter | Integral Membrane Protein | | | | |
| *yebR* | CDS | conserved protein | Cytoplasmic | | | gg | t |
| *yebS* | CDS | conserved inner membrane protein | Integral Membrane Protein | COG2995;Uncharacterized paraquat-inducible protein A | | o | tt |
| *yebT* | CDS | conserved protein | Membrane Anchored | COG3008;Paraquat-inducible protein B | | o | tt |
| *yebU* | CDS | predicted methyltransferase | Cytoplasmic | COG0144;tRNA and rRNA cytosine-C5-methylases -!- COG3270;Uncharacterized conserved protein | GO:0009451 RNA modification | | |
| *yebV* | CDS | predicted protein | Cytoplasmic | | | | |
| *yebW* | CDS | predicted protein | Cytoplasmic | | | | |
| *yebY* | CDS | predicted protein | Periplasmic | | | o | o |
| *yebZ* | CDS | predicted inner membrane protein | Integral Membrane Protein | COG1276;Putative copper export protein | GO:0042493 response to drug | o | tt |
| *yecA* | CDS | conserved metal-binding protein | Cytoplasmic | COG3318;Predicted metal-binding protein related to the C-terminal domain of SecA | | gg | tt |
| *yecC* | CDS | predicted transporter subunit -!- ATP-binding component of ABC superfamily | Cytoplasmic | COG1126;ABC-type polar amino acid transport system, ATPase component | | gg | tt |
| *yecD* | CDS | predicted hydrolase | Cytoplasmic | COG1335;Amidases related to nicotinamidase | | | |
| *yecE* | CDS | conserved protein | Cytoplasmic | COG1801;Uncharacterized conserved protein | | gg | t |
| *yecF* | CDS | predicted protein | Cytoplasmic | | | gg | o |
| *yecG* | CDS | universal stress protein | Cytoplasmic | COG0589;Universal stress protein UspA and related nucleotide-binding proteins | | gg | o |
| *yecH* | CDS | predicted protein | Cytoplasmic | | | gg | o |
| *yecI* | CDS | predicted ferritin-like protein | Cytoplasmic | COG1528;Ferritin-like protein | GO:0006826 iron ion transport | gg | o |
| *yecJ* | CDS | predicted protein | Cytoplasmic | | | gg | o |
| *yecM* | CDS | predicted metal-binding enzyme | Cytoplasmic | | | | |
| *yecN* | CDS | predicted inner membrane protein | Integral Membrane Protein | | | | |
| *yecO* | CDS | predicted methyltransferase | Cytoplasmic | COG0500;SAM-dependent methyltransferases | | gg | o |
| *yecP* | CDS | predicted methyltransferase | Cytoplasmic | COG0500;SAM-dependent methyltransferases | | gg | t |
| *yecR* | CDS | predicted protein | Outer Membrane Lipoprotein | | | o | tt |
| *yecS* | CDS | predicted transporter subunit -!- membrane component of ABC superfamily | Integral Membrane Protein | COG0765;ABC-type amino acid transport system, permease component | | o | tt |
| *yecT* | CDS | predicted protein | Periplasmic | | | | |
| *yedA* | CDS | predicted inner membrane protein | Integral Membrane Protein | COG0697;Permeases of the drug/metabolite transporter (DMT) superfamily | | gg | tt |
| *yedD* | CDS | predicted protein | Outer Membrane Lipoprotein | | | o | o |
| *yedE* | CDS | predicted inner membrane protein | Integral Membrane Protein | COG2391;Predicted transporter component | | o | tt |
| *yedF* | CDS | conserved protein | Cytoplasmic | COG0425;Predicted redox protein, regulator of disulfide bond formation | | gg | o |
| *yedI* | CDS | conserved inner membrane protein | Integral Membrane Protein | COG2354;Uncharacterized protein conserved in bacteria | | g | tt |

| | | | | | | | |
|---|---|---|---|---|---|---|---|
| *yedJ* | CDS | predicted phosphohydrolase | Cytoplasmic | COG1418;Predicted HD superfamily hydrolase | | gg | t |
| *yedK* | CDS | predicted protein | Cytoplasmic | | | gg | t |
| *yedL* | CDS | predicted acyltransferase | Cytoplasmic | COG0454;Histone acetyltransferase HPA2 and related acetyltransferases | | g | o |
| *yedM* | CDS | conserved protein | Cytoplasmic | COG4886;Leucine-rich repeat (LRR) protein | | gg | o |
| *yedN* | CDS | predicted protein, N-ter fragment (pseudogene) | Cytoplasmic | | | | |
| *yedN* | ancestral | predicted protein (pseudogene) | Cytoplasmic | | | | |
| *yedN* | CDS | predicted protein, C-ter fragment (pseudogene) | Cytoplasmic | | | | |
| *yedO* | CDS | D-cysteine desulfhydrase, PLP-dependent | Cytoplasmic | | | | |
| *yedP* | CDS | conserved protein | Cytoplasmic | COG3769;Predicted hydrolase (HAD superfamily) | | gg | t |
| *yedQ* | CDS | predicted diguanylate cyclase | Integral Membrane Protein | COG2199;FOG: GGDEF domain | | | |
| *yedR* | CDS | predicted inner membrane protein | Integral Membrane Protein | | | gg | o |
| *yedS* | ancestral | predicted protein (pseudogene) | Cytoplasmic | | | | |
| *yedS* | CDS | predicted protein, N-ter fragment (pseudogene) | Cytoplasmic | | | | |
| *yedS* | CDS | predicted protein, middle fragment (pseudogene) | Cytoplasmic | | | o | o |
| *yedS* | CDS | predicted protein, C-ter fragment (pseudogene) | Cytoplasmic | | | | |
| *yedV* | CDS | predicted sensory kinase in two-component regulatory system with YedW | Membrane Anchored | COG0642;Signal transduction histidine kinase | GO:0006464 protein modification | o | tt |
| *yedW* | CDS | predicted DNA-binding response regulator in two-component system withYedV | Cytoplasmic | COG0745;Response regulators consisting of a CheY-like receiver domain and a winged-helix DNA-binding domain | GO:0006350 transcription | | |
| *yedX* | CDS | conserved protein | Periplasmic | COG2351;Transthyretin-like protein | | gg | o |
| *yedY* | CDS | predicted reductase | Periplasmic | COG2041;Sulfite oxidase and related enzymes | | gg | t |
| *yedZ* | CDS | conserved inner membrane protein | Integral Membrane Protein | COG2717;Predicted membrane protein | | o | tt |
| *yeeA* | CDS | conserved inner membrane protein | Integral Membrane Protein | COG1289;Predicted membrane protein | | o | tt |
| *yeeD* | CDS | conserved protein | Cytoplasmic | COG0425;Predicted redox protein, regulator of disulfide bond formation | | gg | o |
| *yeeE* | CDS | predicted inner membrane protein | Integral Membrane Protein | COG2391;Predicted transporter component | | o | o |
| *yeeF* | CDS | predicted amino-acid transporter | Integral Membrane Protein | COG0531;Amino acid transporters | | | |
| *yeeI* | CDS | conserved protein | Cytoplasmic | COG3228;Uncharacterized protein conserved in bacteria | | gg | tt |
| *yeeJ* | CDS | adhesin | Cytoplasmic | | GO:0006355 regulation of transcription, DNA-dependent | | |
| *yeeL* | CDS | predicted protein, C-ter fragment (pseudogene) | Cytoplasmic | | | | |
| *yeeL* | ancestral | predicted protein (pseudogene) | Cytoplasmic | | | | |
| *yeeL* | CDS | predicted protein, N-ter fragment (pseudogene) | Cytoplasmic | | | gg | t |
| *yeeN* | CDS | conserved protein | Cytoplasmic | COG0217;Uncharacterized conserved protein | | gg | t |
| *yeeO* | CDS | predicted multidrug efflux system | Integral Membrane Protein | COG0534;Na+-driven multidrug efflux pump | | o | tt |
| *yeeP* | CDS | CP4-44 prophage; predicted GTP-binding protein (pseudogene) | Cytoplasmic | COG3596;Predicted GTPase | | | |
| *yeeR* | CDS | CP4-44 prophage; predicted membrane protein | Integral Membrane Protein | | | o | tt |
| *yeeS* | CDS | CP4-44 prophage; predicted DNA repair protein | Cytoplasmic | COG2003;DNA repair proteins | | gg | o |
| *yeeT* | CDS | CP4-44 prophage; predicted protein | Cytoplasmic | | | gg | o |
| *yeeU* | CDS | CP4-44 prophage; antitoxin of the YeeV-YeeU toxin-antitoxin system | Cytoplasmic | | | gg | o |
| *yeeV* | CDS | CP4-44 prophage; toxin of the YeeV-YeeU toxin-antitoxin system | Cytoplasmic | | | gg | tt |
| *yeeW* | CDS | CP4-44 prophage; predicted protein | Cytoplasmic | | | gg | o |
| *yeeX* | CDS | conserved protein | Cytoplasmic | | | gg | t |
| *yeeY* | CDS | predicted DNA-binding transcriptional regulator | Cytoplasmic | | GO:0006350 transcription | | |
| *yeeZ* | CDS | predicted epimerase, with NAD(P)-binding Rossmann-fold domain | Periplasmic | COG0451;Nucleoside-diphosphate-sugar epimerases | GO:0016052 carbohydrate catabolism | | |
| *yefM* | CDS | antitoxin of the YoeB-YefM toxin-antitoxin system | Cytoplasmic | COG2161;Antitoxin of toxin-antitoxin stability system | | | |
| *yegD* | CDS | predicted chaperone | Cytoplasmic | | GO:0006457 protein folding | o | tt |
| *yegE* | CDS | predicted diguanylate cyclase, GGDEF domain signalling protein | Integral Membrane Protein | COG3447;Predicted integral membrane sensor domain -!- COG2202;FOG: PAS/PAC domain -!- COG2199;FOG: GGDEF domain -!- COG2200;FOG: EAL domain | GO:0006464 protein modification | o | tt |
| *yegH* | CDS | fused predicted membrane proteins | Integral Membrane Protein | | | | |
| *yegI* | CDS | conserved protein | Cytoplasmic | COG4248;Uncharacterized protein with protein kinase and helix-hairpin-helix DNA-binding domains | | o | tt |
| *yegJ* | CDS | predicted protein | Periplasmic | COG3779;Uncharacterized protein conserved in bacteria | | o | tt |
| *yegK* | CDS | predicted protein | Cytoplasmic | | | o | tt |
| *yegL* | CDS | conserved protein | Cytoplasmic | COG4245;Uncharacterized protein encoded in toxicity protection region of plasmid R478, contains von Willebrand factor (vWF) domain | | gg | t |
| *yegP* | CDS | predicted protein | Cytoplasmic | | | | |
| *yegQ* | CDS | predicted peptidase | Cytoplasmic | COG0826;Collagenase and related proteases | | gg | o |
| *yegR* | CDS | predicted protein | Outer Membrane Lipoprotein | | | | |
| *yegS* | CDS | conserved protein | Cytoplasmic | COG1597;Sphingosine kinase and enzymes related to eukaryotic diacylglycerol kinase | | g | t |
| *yegT* | CDS | predicted nucleoside transporter | Integral Membrane Protein | COG0477;Permeases of the major facilitator superfamily | | o | tt |
| *yegU* | CDS | predicted hydrolase | Cytoplasmic | COG1397;ADP-ribosylglycohydrolase | | gg | o |
| *yegV* | CDS | predicted kinase | Cytoplasmic | COG0524;Sugar kinases, ribokinase family | | | |
| *yegW* | CDS | predicted DNA-binding transcriptional regulator | Cytoplasmic | COG2188;Transcriptional regulators | GO:0006350 transcription | o | tt |
| *yegX* | CDS | predicted hydrolase | Membrane Anchored | COG3757;Lyzozyme M1 (1,4-beta-N-acetylmuramidase) | | | |
| *yegZ* | CDS | predicted protein fragment (pseudogene) | Cytoplasmic | COG3500;Phage protein D | | | |
| *yehA* | CDS | predicted fimbrial-like adhesin protein | Periplasmic | | | o | tt |
| *yehB* | CDS | predicted outer membrane protein | Outer Membrane B-barrel protein | COG3188;P pilus assembly protein, porin PapC | | o | tt |
| *yehC* | CDS | predicted periplasmic pilin chaperone | Periplasmic | COG3121;P pilus assembly protein, chaperone PapD | GO:0006457 protein folding | o | tt |
| *yehD* | CDS | predicted fimbrial-like adhesin protein | Periplasmic | COG3539;P pilus assembly protein, pilin FimA | | o | tt |
| *yehE* | CDS | predicted protein | Periplasmic | | | o | tt |
| *yehI* | CDS | conserved protein | Cytoplasmic | | | o | tt |
| *yehK* | CDS | predicted protein | Cytoplasmic | | | gg | o |
| *yehL* | CDS | predicted transporter subunit -!- ATP-binding compoents of ABC superfamily | Cytoplasmic | COG0714;MoxR-like ATPases | | | |
| *yehM* | CDS | predicted protein | Cytoplasmic | | | gg | tt |
| *yehP* | CDS | conserved protein | Cytoplasmic | | | | |
| *yehQ* | CDS | predicted protein | Cytoplasmic | | | o | tt |
| *yehR* | CDS | conserved protein | Membrane Lipoprotein | COG4808;Uncharacterized protein conserved in bacteria | | | |
| *yehS* | CDS | conserved protein | Cytoplasmic | COG4807;Uncharacterized protein conserved in bacteria | | gg | t |
| *yehT* | CDS | predicted response regulator in two-component system withYehU | Cytoplasmic | | | | |
| *yehU* | CDS | predicted sensory kinase in two-component system with YehT | Integral Membrane Protein | COG3275;Putative regulator of cell autolysis | GO:0006464 protein modification | | |
| *yehW* | CDS | predicted transporter subunit -!- membrane component of ABC superfamily | Integral Membrane Protein | COG1174;ABC-type proline/glycine betaine transport systems, permease component | | o | tt |
| *yehX* | CDS | predicted transporter subunit -!- ATP-binding component of ABC superfamily | Cytoplasmic | COG1125;ABC-type proline/glycine betaine transport systems, ATPase components | | g | t |
| *yehY* | CDS | predicted transporter subunit -!- membrane component of ABC superfamily | Integral Membrane Protein | COG1174;ABC-type proline/glycine betaine transport systems, permease component | | o | tt |
| *yehZ* | CDS | predicted transporter subunit -!- periplasmic-binding component of ABC superfamily | Periplasmic | COG1732;Periplasmic glycine betaine/choline-binding (lipo)protein of an ABC-type transport system (osmoprotectant binding protein) | | gg | o |
| *yeiA* | CDS | predicted oxidoreductase | Cytoplasmic | COG0167;Dihydroorotate dehydrogenase -!- COG1146;Ferredoxin | | g | t |
| *yeiB* | CDS | conserved inner membrane protein | Integral Membrane Protein | COG2311;Predicted membrane protein | | o | tt |
| *yeiC* | CDS | predicted kinase | Cytoplasmic | COG0524;Sugar kinases, ribokinase family | | gg | t |
| *yeiE* | CDS | predicted DNA-binding transcriptional regulator | Cytoplasmic | COG0583;Transcriptional regulator | GO:0006350 transcription | gg | t |
| *yeiG* | CDS | predicted esterase | Cytoplasmic | COG0627;Predicted esterase | | gg | o |
| *yeiH* | CDS | conserved inner membrane protein | Integral Membrane Protein | COG2855;Predicted membrane protein | | o | tt |
| *yeiI* | CDS | predicted kinase | Cytoplasmic | COG2771;DNA-binding HTH domain-containing proteins -!- COG0524;Sugar kinases, ribokinase family | | g | tt |
| *yeiJ* | CDS | predicted nucleoside transporter | Integral Membrane Protein | COG1972;Nucleoside permease | | o | tt |
| *yeiL* | CDS | DNA-binding transcriptional activator | Cytoplasmic | | GO:0006355 regulation of transcription, DNA-dependent | gg | o |
| *yeiM* | CDS | predicted nucleoside transporter | Integral Membrane Protein | COG1972;Nucleoside permease | | g | tt |
| *yeiN* | CDS | conserved protein | Cytoplasmic | COG2313;Uncharacterized enzyme involved in pigment biosynthesis | | gg | o |
| *yeiP* | CDS | predicted elongtion factor | Cytoplasmic | | GO:0006412 protein biosynthesis | | |
| *yeiQ* | CDS | predicted dehydrogenase, NAD-dependent | Cytoplasmic | COG0246;Mannitol-1-phosphate/altronate dehydrogenases | | g | t |
| *yeiR* | CDS | predicted enzyme | Cytoplasmic | COG0523;Putative GTPases (G3E family) | | gg | t |
| *yeiS* | CDS | predicted inner membrane protein | Integral Membrane Protein | | | | |
| *yeiT* | CDS | predicted oxidoreductase | Cytoplasmic | COG0493;NADPH-dependent glutamate synthase beta chain and related oxidoreductases | | gg | t |
| *yeiU* | CDS | undecaprenyl pyrophosphate phosphatase | Integral Membrane Protein | | | o | tt |
| *yeiW* | CDS | conserved protein | | | | | |
| *yejA* | CDS | predicted oligopeptide transporter subunit -!- periplasmic-binding component of ABC superfamily | Periplasmic | | | o | tt |
| *yejB* | CDS | predicted oligopeptide transporter subunit -!- membrane component of ABC superfamily | Integral Membrane Protein | COG4174;ABC-type uncharacterized transport system, permease component | | o | tt |
| *yejE* | CDS | predicted oligopeptide transporter subunit -!- membrane component of ABC superfamily | Integral Membrane Protein | COG4239;ABC-type uncharacterized transport system, permease component | | o | tt |
| *yejF* | CDS | fused predicted oligopeptide transporter subunits -!- ATP-binding components of ABC superfamily | Cytoplasmic | COG4172;ABC-type uncharacterized transport system, duplicated ATPase component | | o | tt |
| *yejG* | CDS | predicted protein | Cytoplasmic | | | gg | o |
| *yejH* | CDS | predicted ATP-dependet helicase | Cytoplasmic | COG1061;DNA or RNA helicases of superfamily II | | g | t |
| *yejK* | CDS | nucleotide associated protein | Cytoplasmic | COG3081;Nucleoid-associated protein | | o | tt |
| *yejL* | CDS | conserved protein | Cytoplasmic | COG3082;Uncharacterized protein conserved in bacteria | | gg | o |
| *yejM* | CDS | predicted hydrolase, inner membrane | Integral Membrane Protein | COG3083;Predicted hydrolase of alkaline phosphatase superfamily | | o | tt |
| *yejO* | CDS | predicted autotransporter outer membrane protein | Cytoplasmic | COG3468;Type V secretory pathway, adhesin AidA | | | |
| *yfaA* | CDS | predicted protein | Periplasmic | | | g | tt |
| *yfaD* | CDS | conserved protein | Cytoplasmic | | | gg | o |
| *yfaE* | CDS | predicted 2Fe-2S cluster-containing protein | Cytoplasmic | COG0633;Ferredoxin | | g | t |
| *yfaH* | CDS | conserved protein | Cytoplasmic | COG0583;Transcriptional regulator | | g | o |
| *yfaL* | CDS | adhesin | Periplasmic | COG3468;Type V secretory pathway, adhesin AidA | | gg | tt |

| | | | | | | | |
|---|---|---|---|---|---|---|---|
| *yfaO* | CDS | predicted NUDIX hydrolase | Cytoplasmic | COG0494;NTP pyrophosphohydrolases including oxidative damage repair enzymes | | g | t |
| *yfaP* | CDS | conserved protein | Periplasmic | COG4676;Uncharacterized protein conserved in bacteria | | o | tt |
| *yfaQ* | CDS | predicted protein | Periplasmic | | | g | tt |
| *yfaS* | CDS | predicted protein, C-ter fragment (pseudogene) | Periplasmic | | | gg | o |
| *yfaS* | ancestral | predicted protein (pseudogene) | Periplasmic | | | | |
| *yfaS* | CDS | predicted protein, N-ter fragment (pseudogene) | Periplasmic | | | o | tt |
| *yfaT* | CDS | predicted protein | Periplasmic | | | o | tt |
| *yfaU* | CDS | predicted 2,4-dihydroxyhept-2-ene-1,7-dioic acid aldolase | Cytoplasmic | COG3836;2,4-dihydroxyhept-2-ene-1,7-dioic acid aldolase | | o | o |
| *yfaV* | CDS | predicted transporter | Integral Membrane Protein | | | o | tt |
| *yfaW* | CDS | predicted enolase | Cytoplasmic | COG4948;L-alanine-DL-glutamate epimerase and related enzymes of enolase superfamily | | gg | t |
| *yfaX* | CDS | predicted DNA-binding transcriptional regulator | Cytoplasmic | COG1414;Transcriptional regulator | GO:0006350 transcription -!- GO:0006355 regulation of transcription, DNA-dependent | gg | t |
| *yfaY* | CDS | conserved protein | Cytoplasmic | COG1058;Predicted nucleotide-utilizing enzyme related to molybdopterin-biosynthesis enzyme MoeA | | g | tt |
| *yfaZ* | CDS | predicted outer membrane porin protein | Outer Membrane B-barrel protein | | | | |
| *yfbB* | CDS | predicted peptidase | Cytoplasmic | COG0596;Predicted hydrolases or acyltransferases (alpha/beta hydrolase superfamily) | | gg | tt |
| *yfbE* | CDS | uridine 5'-(beta-1-threo-pentapyranosyl-4-ulose diphosphate) aminotransferase, PLP-dependent | Cytoplasmic | | | | |
| *yfbF* | CDS | undecaprenyl phosphate-L-Ara4FN transferase | Integral Membrane Protein | COG0463;Glycosyltransferases involved in cell wall biogenesis | | gg | tt |
| *yfbG* | CDS | fused UDP-L-Ara4N formyltransferase -!- UDP-GlcA C-4'-decarboxylase | Cytoplasmic | COG0223;Methionyl-tRNA formyltransferase -!- COG0451;Nucleoside-diphosphate-sugar epimerases | | g | tt |
| *yfbH* | CDS | conserved protein | Cytoplasmic | COG0726;Predicted xylanase/chitin deacetylase | | gg | t |
| *yfbJ* | CDS | predicted inner membrane protein | Integral Membrane Protein | | | | |
| *yfbK* | CDS | conserved protein | Membrane Lipoprotein | COG2304;Uncharacterized protein containing a von Willebrand factor type A (vWA) domain | | g | tt |
| *yfbL* | CDS | predicted peptidase | Membrane Anchored | COG2234;Predicted aminopeptidases | | o | tt |
| *yfbM* | CDS | predicted protein | Cytoplasmic | | | gg | t |
| *yfbN* | CDS | predicted protein | Cytoplasmic | | | gg | t |
| *yfbO* | CDS | predicted protein | Cytoplasmic | | | g | t |
| *yfbP* | CDS | predicted protein | Cytoplasmic | | | gg | o |
| *yfbQ* | CDS | predicted aminotransferase | Cytoplasmic | COG0436;Aspartate/tyrosine/aromatic aminotransferase | | gg | t |
| *yfbR* | CDS | deoxyribonucleoside 5'-monophosphatase | Cytoplasmic | COG1896;Predicted hydrolases of HD superfamily | | gg | o |
| *yfbS* | CDS | predicted transporter | Integral Membrane Protein | COG0471;Di- and tricarboxylate transporters | | o | tt |
| *yfbT* | CDS | predicted hydrolase or phosphatase | Cytoplasmic | COG0637;Predicted phosphatase/phosphohexomutase | | | |
| *yfbU* | CDS | conserved protein | Cytoplasmic | COG3013;Uncharacterized conserved protein | | g | o |
| *yfbV* | CDS | conserved inner membrane protein | Integral Membrane Protein | COG3092;Uncharacterized protein conserved in bacteria | | o | tt |
| *yfbW* | CDS | conserved protein | Integral Membrane Protein | | | o | tt |
| *yfcA* | CDS | conserved inner membrane protein | Integral Membrane Protein | COG0730;Predicted permeases | | o | tt |
| *yfcC* | CDS | predicted inner membrane protein | Integral Membrane Protein | COG1288;Predicted membrane protein | | o | tt |
| *yfcD* | CDS | predicted NUDIX hydrolase | Cytoplasmic | COG0494;NTP pyrophosphohydrolases including oxidative damage repair enzymes | | gg | tt |
| *yfcE* | CDS | predicted phosphatase | Cytoplasmic | | | | |
| *yfcF* | CDS | predicted enzyme | Cytoplasmic | COG0625;Glutathione S-transferase | | g | o |
| *yfcG* | CDS | predicted glutathione S-transferase | Cytoplasmic | COG0625;Glutathione S-transferase | | gg | o |
| *yfcH* | CDS | conserved protein with NAD(P)-binding Rossmann-fold domain | Cytoplasmic | COG1090;Predicted nucleoside-diphosphate sugar epimerase | | gg | tt |
| *yfcI* | CDS | conserved protein | Cytoplasmic | | | gg | t |
| *yfcJ* | CDS | predicted transporter | Integral Membrane Protein | COG0477;Permeases of the major facilitator superfamily | | o | tt |
| *yfcL* | CDS | predicted protein | Cytoplasmic | | | gg | tt |
| *yfcM* | CDS | conserved protein | Cytoplasmic | COG3101;Uncharacterized protein conserved in bacteria | | | |
| *yfcN* | CDS | conserved protein | Cytoplasmic | COG2840;Uncharacterized protein conserved in bacteria | | gg | t |
| *yfcO* | CDS | predicted protein | Periplasmic | | | o | tt |
| *yfcP* | CDS | predicted fimbrial-like adhesin protein | Periplasmic | COG3539;P pilus assembly protein, pilin FimA | | o | tt |
| *yfcQ* | CDS | predicted fimbrial-like adhesin protein | Periplasmic | COG3539;P pilus assembly protein, pilin FimA | | o | tt |
| *yfcR* | CDS | predicted fimbrial-like adhesin protein | Periplasmic | COG3539;P pilus assembly protein, pilin FimA | | o | tt |
| *yfcS* | CDS | predicted periplasmic pilus chaperone | Periplasmic | COG3121;P pilus assembly protein, chaperone PapD | GO:0006457 protein folding | o | tt |
| *yfcT* | CDS | predicted outer membrane export usher protein | Cytoplasmic | | | gg | o |
| *yfcU* | CDS | predicted export usher protein | Periplasmic | | | o | tt |
| *yfcV* | CDS | predicted fimbrial-like adhesin protein | Periplasmic | COG3539;P pilus assembly protein, pilin FimA | | o | tt |
| *yfcX* | CDS | fused enoyl-CoA hydratase-!-epimerase-!-isomerase -!- 3-hydroxyacyl-CoA dehydrogenase, subunit of anaerobic fatty acid oxidation complex | Cytoplasmic | COG1024;Enoyl-CoA hydratase/carnithine racemase -!- COG1250;3-hydroxyacyl-CoA dehydrogenase | GO:0019395 fatty acid oxidation | o | tt |
| *yfcY* | CDS | beta-ketoacyl-CoA thiolase, anaerobic, subunit | Cytoplasmic | COG0183;Acetyl-CoA acetyltransferase | | o | o |
| *yfcZ* | CDS | conserved protein | Cytoplasmic | COG3691;Uncharacterized protein conserved in bacteria | | gg | t |
| *yfdC* | CDS | predicted inner membrane protein | Integral Membrane Protein | COG2116;Formate/nitrite family of transporters | | o | tt |
| *yfdE* | CDS | predicted CoA-transferase, NAD(P)-binding | Cytoplasmic | COG1804;Predicted acyl-CoA transferases/carnitine dehydratase | | gg | t |
| *yfdF* | CDS | predicted protein | Cytoplasmic | | GO:0006259 DNA metabolism | gg | o |
| *yfdG* | CDS | CPS-53 (KpLE1) prophage; bactoprenol-linked glucose translocase (flippase) | Integral Membrane Protein | COG2246;Predicted membrane protein | | o | tt |
| *yfdH* | CDS | CPS-53 (KpLE1) prophage; bactoprenol glucosyl transferase | Integral Membrane Protein | COG0463;Glycosyltransferases involved in cell wall biogenesis | | gg | tt |
| *yfdI* | CDS | CPS-53 (KpLE1) prophage; predicted inner membrane protein | Integral Membrane Protein | | | | |
| *yfdK* | CDS | CPS-53 (KpLE1) prophage; conserved protein | Cytoplasmic | | | gg | t |
| *yfdL* | CDS | CPS-53 (KpLE1) prophage; conserved protein | Cytoplasmic | | | | |
| *yfdM* | CDS | CPS-53 (KpLE1) prophage; predicted methyltransferase | Cytoplasmic | | | gg | t |
| *yfdN* | CDS | CPS-53 (KpLE1) prophage; predicted protein | Cytoplasmic | | | | |
| *yfdO* | CDS | CPS-53 (KpLE1) prophage; predicted defective phage replication protein O | Cytoplasmic | | | gg | t |
| *yfdP* | CDS | CPS-53 (KpLE1) prophage; predicted protein | Cytoplasmic | | | o | tt |
| *yfdQ* | CDS | CPS-53 (KpLE1) prophage; predicted protein | Cytoplasmic | | | g | o |
| *yfdR* | CDS | CPS-53 (KpLE1) prophage; conserved protein | Cytoplasmic | COG1896;Predicted hydrolases of HD superfamily | | gg | t |
| *yfdS* | CDS | CPS-53 (KpLE1) prophage; predicted protein | Cytoplasmic | | | gg | o |
| *yfdT* | CDS | CPS-53 (KpLE1) prophage; predicted protein | Cytoplasmic | | | | |
| *yfdV* | CDS | predicted transporter | Integral Membrane Protein | COG0679;Predicted permeases | | | |
| *yfdX* | CDS | predicted protein | Periplasmic | | | o | tt |
| *yfdY* | CDS | predicted inner membrane protein | Integral Membrane Protein | | | o | tt |
| *yfdZ* | CDS | prediected aminotransferase, PLP-dependent | Cytoplasmic | COG0436;Aspartate/tyrosine/aromatic aminotransferase | | gg | t |
| *yfeA* | CDS | predicted diguanylate cyclase | Integral Membrane Protein | COG2199;FOG: GGDEF domain -!- COG2200;FOG: EAL domain | | | |
| *yfeC* | CDS | predicted DNA-binding transcriptional regulator | Cytoplasmic | | | gg | tt |
| *yfeD* | CDS | predicted DNA-binding transcriptional regulator | Cytoplasmic | | | o | tt |
| *yfeG* | CDS | predicted DNA-binding transcriptional regulator | Cytoplasmic | COG2207;AraC-type DNA-binding domain-containing proteins | GO:0009310 amine catabolism -!- GO:0006350 transcription | gg | t |
| *yfeH* | CDS | predicted inner membrane protein | Integral Membrane Protein | COG0385;Predicted Na+-dependent transporter | | | |
| *yfeK* | CDS | predicted protein | Periplasmic | | | o | tt |
| *yfeN* | CDS | conserved outer membrane protein | Outer Membrane B-barrel protein | | | o | tt |
| *yfeO* | CDS | predicted ion channel protein | Integral Membrane Protein | COG0038;Chloride channel protein EriC | | o | tt |
| *yfeR* | CDS | predicted DNA-binding transcriptional regulator | Cytoplasmic | COG0583;Transcriptional regulator | GO:0006350 transcription | o | tt |
| *yfeS* | CDS | conserved protein | Cytoplasmic | COG3831;Uncharacterized conserved protein -!- COG4884;Uncharacterized protein conserved in bacteria | | g | tt |
| *yfeT* | CDS | predicted DNA-binding transcriptional regulator | Cytoplasmic | COG1737;Transcriptional regulators | | o | tt |
| *yfeU* | CDS | predicted PTS component | Cytoplasmic | COG2103;Predicted sugar phosphate isomerase | | | |
| *yfeW* | CDS | predicted periplasmic esterase | Periplasmic | COG1680;Beta-lactamase class C and other penicillin binding proteins | GO:0016052 carbohydrate | | |
| *yfeX* | CDS | conserved protein | Cytoplasmic | | | gg | o |
| *yfeY* | CDS | predicted protein | Outer Membrane Lipoprotein | | | o | tt |
| *yfeZ* | CDS | predicted inner membrane protein | Integral Membrane Protein | | | o | tt |
| *yffB* | CDS | conserved protein | Cytoplasmic | COG1393;Arsenate reductase and related proteins, glutaredoxin family | | gg | tt |
| *yffH* | CDS | predicted NUDIX hydrolase | Cytoplasmic | COG0494;NTP pyrophosphohydrolases including oxidative damage repair enzymes | | gg | t |
| *yffI* | CDS | predicted carboxysome structural protein with predicted role in ethanolamine utilization | Cytoplasmic | | GO:0009310 amine catabolism | gg | t |
| *yffL* | CDS | CPZ-55 prophage; predicted protein | Cytoplasmic | | | | |
| *yffM* | CDS | CPZ-55 prophage; predicted protein | Cytoplasmic | | | | |
| *yffN* | CDS | CPZ-55 prophage; predicted protein | Cytoplasmic | | | | |
| *yffO* | CDS | CPZ-55 prophage; predicted protein | Cytoplasmic | | | | |
| *yffP* | CDS | CPZ-55 prophage; predicted protein | Cytoplasmic | | | | |
| *yffQ* | CDS | CPZ-55 prophage; predicted protein | Periplasmic | | | | |
| *yffR* | CDS | CPZ-55 prophage; predicted protein | Cytoplasmic | | | | |
| *yffS* | CDS | CPZ-55 prophage; predicted protein | Cytoplasmic | | | | |
| *yfgA* | CDS | conserved protein | Membrane Anchored | COG1426;Uncharacterized protein conserved in bacteria | | o | tt |
| *yfgB* | CDS | predicted enzyme | Cytoplasmic | COG0820;Predicted Fe-S-cluster redox enzyme | | gg | t |
| *yfgC* | CDS | predicted peptidase | Periplasmic | COG4783;Putative Zn-dependent protease, contains TPR repeats | | | |
| *yfgD* | CDS | predicted oxidoreductase | Cytoplasmic | COG1393;Arsenate reductase and related proteins, glutaredoxin family | GO:0006805 xenobiotic metabolism | o | o |
| *yfgF* | CDS | predicted inner membrane protein | Integral Membrane Protein | COG2199;FOG: GGDEF domain -!- COG2200;FOG: EAL domain | | o | tt |
| *yfgG* | CDS | predicted protein | Membrane Anchored | | | | |
| *yfgH* | CDS | predicted outer membrane lipoprotein | Outer Membrane Lipoprotein | | | | |
| *yfgI* | CDS | conserved protein | Periplasmic | | | gg | tt |
| *yfgJ* | CDS | predicted protein | Cytoplasmic | | GO:0042966 biotin carboxyl carrier protein biosynthesis | | |

| | | | | | | | |
|---|---|---|---|---|---|---|---|
| *yfgL* | CDS | protein assembly complex, lipoprotein component | Outer Membrane Lipoprotein | COG1520;FOG: WD40-like repeat | GO:0006464 protein modification | o | tt |
| *yfgM* | CDS | conserved protein | Membrane Anchored | COG2976;Uncharacterized protein conserved in bacteria | | | |
| *yfgO* | CDS | predicted inner membrane protein | Integral Membrane Protein | COG0628;Predicted permease | | o | tt |
| *yfhA* | CDS | predicted DNA-binding response regulator in two-component system | Cytoplasmic | COG2204;Response regulator containing CheY-like receiver, AAA-type ATPase, and DNA-binding domains | GO:0006464 protein modification | gg | t |
| *yfhB* | CDS | conserved protein | Membrane Anchored | | | | |
| *yfhD* | CDS | predicted transglycosylase | Periplasmic | COG4623;Predicted soluble lytic transglycosylase fused to an ABC-type amino acid-binding protein | | gg | t |
| *yfhG* | CDS | conserved protein | Outer Membrane Lipoprotein | | | gg | tt |
| *yfhH* | CDS | predicted DNA-binding transcriptional regulator | Cytoplasmic | COG1737;Transcriptional regulators | | o | tt |
| *yfhJ* | CDS | conserved protein | Cytoplasmic | COG2975;Uncharacterized protein conserved in bacteria | GO:0006457 protein folding | gg | o |
| *yfhK* | CDS | predicted sensory kinase in two-component system | Integral Membrane Protein | COG0642;Signal transduction histidine kinase | GO:0006464 protein modification | | |
| *yfhL* | CDS | predicted 4Fe-4S cluster-containing protein | Cytoplasmic | COG1145;Ferredoxin | | gg | t |
| *yfhM* | CDS | conserved protein | Inner Membrane Lipoprotein | COG2373;Large extracellular alpha-helical protein | | o | tt |
| *yfhQ* | CDS | predicted methyltransferase | Cytoplasmic | COG0565;rRNA methylase | | gg | t |
| *yfhR* | CDS | predicted peptidase | Membrane Anchored | | | o | tt |
| *yfiA* | CDS | cold shock protein associated with 30S ribosomal subunit | Cytoplasmic | COG1544;Ribosome-associated protein Y (PSrp-1) | GO:0006412 protein biosynthesis | gg | t |
| *yfiB* | CDS | predicted outer membrane lipoprotein | Outer Membrane Lipoprotein | COG2885;Outer membrane protein and related peptidoglycan-associated (lipo)proteins | | o | tt |
| *yfiC* | CDS | predicted methyltransferase | Cytoplasmic | COG4123;Predicted O-methyltransferase | | gg | t |
| *yfiD* | CDS | pyruvate formate lyase subunit | Cytoplasmic | COG3445;Acid-induced glycyl radical enzyme | GO:0009061 anaerobic respiration | gg | t |
| *yfiE* | CDS | predicted DNA-binding transcriptional regulator | Cytoplasmic | | GO:0006350 transcription | g | t |
| *yfiF* | CDS | predicted methyltransferase | Cytoplasmic | COG0566;rRNA methylases | GO:0009451 RNA modification | g | tt |
| *yfiH* | CDS | conserved protein | Cytoplasmic | COG1496;Uncharacterized conserved protein | | g | o |
| *yfiK* | CDS | neutral amino-acid efflux system | Integral Membrane Protein | COG1280;Putative threonine efflux protein | | o | tt |
| *yfiL* | CDS | predicted protein | Outer Membrane Lipoprotein | | | | |
| *yfiM* | CDS | predicted protein | Cytoplasmic | | | o | t |
| *yfiN* | CDS | predicted diguanylate cyclase | Integral Membrane Protein | COG2199;FOG: GGDEF domain | | | |
| *yfiO* | CDS | predicted lipoprotein | Outer Membrane Lipoprotein | COG4105;DNA uptake lipoprotein | | o | tt |
| *yfiP* | CDS | conserved protein | Cytoplasmic | | | | |
| *yfiQ* | CDS | fused predicted acyl-CoA synthetase NAD(P)-binding subunit -!- ATP-binding subunit | Cytoplasmic | COG1042;Acyl-CoA synthetase (NDP forming) -!- COG0454;Histone acetyltransferase HPA2 and related acetyltransferases | | gg | tt |
| *yfiR* | CDS | predicted protein | Periplasmic | | | o | tt |
| *yfjD* | CDS | predicted inner membrane protein | Integral Membrane Protein | | | | |
| *yfjF* | CDS | predicted protein | Cytoplasmic | | | gg | o |
| *yfjG* | CDS | conserved protein | Cytoplasmic | COG2867;Oligoketide cyclase/lipid transport protein | | o | tt |
| *yfjH* | CDS | CP4-57 prophage; predicted protein | Cytoplasmic | | | g | tt |
| *yfjI* | CDS | CP4-57 prophage; predicted protein | Cytoplasmic | | | o | tt |
| *yfjJ* | CDS | CP4-57 prophage; predicted protein | Cytoplasmic | | | gg | o |
| *yfjK* | CDS | CP4-57 prophage; conserved protein | Cytoplasmic | COG1204;Superfamily II helicase | | gg | o |
| *yfjL* | CDS | CP4-57 prophage; predicted protein | Cytoplasmic | | | o | t |
| *yfjM* | CDS | CP4-57 prophage; predicted protein | Cytoplasmic | | | gg | o |
| *yfjN* | CDS | CP4-57 prophage; predicted protein | Cytoplasmic | | | g | t |
| *yfjO* | CDS | CP4-57 prophage; predicted protein | Cytoplasmic | | | | |
| *yfjP* | CDS | CP4-57 prophage; predicted GTP-binding protein | Cytoplasmic | | | | |
| *yfjQ* | CDS | CP4-57 prophage; predicted protein | Cytoplasmic | | | g | t |
| *yfjR* | CDS | CP4-57 prophage; predicted DNA-binding transcriptional regulator | Cytoplasmic | COG2378;Predicted transcriptional regulator | | o | tt |
| *yfjS* | CDS | CP4-57 prophage; predicted protein | Inner Membrane Lipoprotein | | | | |
| *yfjT* | CDS | CP4-57 prophage; predicted protein | Periplasmic | | | o | tt |
| *yfjU* | CDS | CP4-57 prophage; conserved protein | Cytoplasmic | COG1393;Arsenate reductase and related proteins, glutaredoxin family | | g | o |
| *yfjV* | CDS | CP4-57 prophage; predicted protein | Cytoplasmic | | | o | tt |
| *yfjW* | CDS | CP4-57 prophage; predicted inner membrane protein | Integral Membrane Protein | | | o | tt |
| *yfjX* | CDS | CP4-57 prophage; predicted antirestriction protein | Cytoplasmic | | | gg | t |
| *yfjY* | CDS | CP4-57 prophage; predicted DNA repair protein | Cytoplasmic | COG2003;DNA repair proteins | | gg | t |
| *yfjZ* | CDS | CP4-57 prophage; antitoxin of the YpjF-YfjZ toxin-antitoxin system | Cytoplasmic | | | gg | t |
| *ygaC* | CDS | predicted protein | Cytoplasmic | | | gg | o |
| *ygaD* | CDS | conserved protein | Cytoplasmic | COG1546;Uncharacterized protein (competence- and mitomycin-induced) | | gg | t |
| *ygaF* | CDS | predicted enzyme | Cytoplasmic | COG0579;Predicted dehydrogenase | | gg | t |
| *ygaH* | CDS | predicted inner membrane protein | Integral Membrane Protein | | | o | tt |
| *ygaM* | CDS | predicted protein | Membrane Anchored | | | g | t |
| *ygaP* | CDS | predicted inner membrane protein with hydrolase activity | Integral Membrane Protein | COG0607;Rhodanese-related sulfurtransferase | | o | tt |
| *ygaQ* | CDS | predicted protein | Cytoplasmic | | | | |
| *ygaR* | CDS | predicted protein | Cytoplasmic | | | o | tt |
| *ygaT* | CDS | predicted protein | Cytoplasmic | | | | |
| *ygaU* | CDS | predicted protein | Cytoplasmic | COG1652;Uncharacterized protein containing LysM domain | | gg | t |
| *ygaV* | CDS | predicted DNA-binding transcriptional regulator | Cytoplasmic | COG0640;Predicted transcriptional regulators | | o | tt |
| *ygaW* | CDS | predicted inner membrane protein | Integral Membrane Protein | | | o | tt |
| *ygaX* | CDS | predicted transporter | Integral Membrane Protein | | GO:0006950 response to stress | o | tt |
| *ygaY* | CDS | predicted transporter (pseudogene) | Integral Membrane Protein | | | | |
| *ygaZ* | CDS | predicted transporter | Integral Membrane Protein | COG1296;Predicted branched-chain amino acid permease (azaleucine resistance) | | o | tt |
| *ygbA* | CDS | predicted protein | Cytoplasmic | | | gg | tt |
| *ygbE* | CDS | conserved inner membrane protein | Integral Membrane Protein | | | o | tt |
| *ygbF* | CDS | predicted protein | Cytoplasmic | | | | |
| *ygbI* | CDS | predicted DNA-binding transcriptional regulator | Cytoplasmic | COG1349;Transcriptional regulators of sugar metabolism | GO:0006350 transcription | o | tt |
| *ygbJ* | CDS | predicted dehydrogenase, with NAD(P)-binding Rossmann-fold domain | Membrane Anchored | COG2084;3-hydroxyisobutyrate dehydrogenase and related beta-hydroxyacid dehydrogenases | | gg | t |
| *ygbK* | CDS | conserved protein | Cytoplasmic | COG3395;Uncharacterized protein conserved in bacteria | | gg | t |
| *ygbL* | CDS | predicted class II aldolase | Cytoplasmic | COG0235;Ribulose-5-phosphate 4-epimerase and related epimerases and aldolases | | gg | t |
| *ygbM* | CDS | conserved protein | Cytoplasmic | COG3622;Hydroxypyruvate isomerase | | g | t |
| *ygbN* | CDS | predicted transporter | Integral Membrane Protein | COG2610;H+/gluconate symporter and related permeases | | o | tt |
| *ygbT* | CDS | conserved protein | Cytoplasmic | COG1518;Uncharacterized protein predicted to be involved in DNA repair | | gg | tt |
| *ygcB* | CDS | conserved protein, member of DEAD box family | Cytoplasmic | COG1203;Predicted helicases | | g | o |
| *ygcE* | CDS | predicted kinase | Cytoplasmic | COG1070;Sugar (pentulose and hexulose) kinases | | | |
| *ygcF* | CDS | conserved protein | Cytoplasmic | COG0602;Organic radical activating enzymes | GO:0046138 coenzymes and prosthetic group biosynthesis | gg | t |
| *ygcG* | CDS | predicted protein | Integral Membrane Protein | | | | |
| *ygcH* | CDS | predicted protein | Cytoplasmic | | | g | t |
| *ygcI* | CDS | predicted protein | Cytoplasmic | | | | |
| *ygcJ* | CDS | predicted protein | Cytoplasmic | | | gg | t |
| *ygcK* | CDS | predicted protein | Cytoplasmic | | | gg | t |
| *ygcL* | CDS | predicted protein | Cytoplasmic | | | gg | o |
| *ygcM* | CDS | 6-pyruvoyl tetrahydrobiopterin synthase (PTPS) | Cytoplasmic | COG0720;6-pyruvoyl-tetrahydropterin synthase | | o | o |
| *ygcN* | CDS | predicted oxidoreductase, FAD/NAD(P)-binding domain | Cytoplasmic | | | g | t |
| *ygcO* | CDS | predicted 4Fe-4S cluster-containing protein | Cytoplasmic | COG2440;Ferredoxin-like protein | | gg | t |
| *ygcP* | CDS | predicted anti-terminator regulatory protein | Cytoplasmic | COG1954;Glycerol-3-phosphate responsive antiterminator (mRNA-binding) | | gg | t |
| *ygcQ* | CDS | predicted flavoprotein | Cytoplasmic | | | | |
| *ygcR* | CDS | predicted flavoprotein | Cytoplasmic | | | | |
| *ygcS* | CDS | predicted transporter | Integral Membrane Protein | COG0477;Permeases of the major facilitator superfamily | | | |
| *ygcU* | CDS | predicted FAD containing dehydrogenase | Cytoplasmic | | | | |
| *ygcW* | CDS | predicted deoxygluconate dehydrogenase | Cytoplasmic | | | | |
| *ygdB* | CDS | predicted protein | Periplasmic | | | | |
| *ygdD* | CDS | conserved inner membrane protein | Integral Membrane Protein | COG2363;Uncharacterized small membrane protein | | o | tt |
| *ygdE* | CDS | predicted methyltransferase | Cytoplasmic | COG2933;Predicted SAM-dependent methyltransferase | | gg | tt |
| *ygdH* | CDS | conserved protein | Cytoplasmic | COG1611;Predicted Rossmann fold nucleotide-binding protein | | gg | t |
| *ygdI* | CDS | predicted protein | Outer Membrane Lipoprotein | | | | |
| *ygdK* | CDS | predicted Fe-S metabolism protein | Cytoplasmic | COG2166;SufE protein probably involved in Fe-S center assembly | | gg | t |
| *ygdL* | CDS | conserved protein | Membrane Anchored | COG1179;Dinucleotide-utilizing enzymes involved in molybdopterin and thiamine biosynthesis family 1 | | gg | tt |
| *ygdQ* | CDS | predicted inner membrane protein | Integral Membrane Protein | COG0861;Membrane protein TerC, possibly involved in tellurium resistance | | o | tt |
| *ygdR* | CDS | predicted protein | Outer Membrane Lipoprotein | | | o | tt |
| *ygeA* | CDS | predicted racemase | Cytoplasmic | COG1794;Aspartate racemase | GO:0042493 response to drug | o | tt |
| *ygeD* | CDS | predicted inner membrane protein | Integral Membrane Protein | COG0477;Permeases of the major facilitator superfamily | GO:0042493 response to drug | o | tt |
| *ygeF* | CDS | predicted protein | Cytoplasmic | | | g | o |
| *ygeG* | CDS | predicted chaperone | Cytoplasmic | COG0457;FOG: TPR repeat | | gg | t |
| *ygeH* | CDS | predictedtranscriptional regulator | Cytoplasmic | COG3710;DNA-binding winged-HTH domains -!- COG0457;FOG: TPR repeat | | g | o |
| *ygeI* | CDS | predicted protein | Cytoplasmic | | | | |
| *ygeK* | CDS | predicted DNA-binding transcriptional regulator | Cytoplasmic | COG2197;Response regulator containing a CheY-like receiver domain and an HTH DNA-binding domain | GO:0006350 transcription | | |
| *ygeL* | CDS | predicted protein | Cytoplasmic | COG2197;Response regulator containing a CheY-like receiver domain and an HTH DNA-binding domain | | gg | o |
| *ygeM* | CDS | predicted protein | Cytoplasmic | | | | |
| *ygeN* | CDS | predicted protein (pseudogene) | Cytoplasmic | | | | |
| *ygeO* | CDS | predicted protein | Cytoplasmic | | | | |
| *ygeP* | CDS | predicted protein | Cytoplasmic | | | g | o |

| | | | | | | | |
|---|---|---|---|---|---|---|---|
| *ygeQ* | CDS | predicted protein | Periplasmic | | | | |
| *ygeR* | CDS | Tetratricopeptide repeat transcriptional regulator | Outer Membrane Lipoprotein | | | o | tt |
| *ygeV* | CDS | predicted DNA-binding transcriptional regulator | Cytoplasmic | COG3829;Transcriptional regulator containing PAS, AAA-type ATPase, and DNA-binding domains | GO:0006350 transcription | gg | t |
| *ygeW* | CDS | conserved protein | Cytoplasmic | | | | |
| *ygeX* | CDS | 2,3-diaminopropionate ammonia-lyase | Cytoplasmic | COG1171;Threonine dehydratase | | gg | o |
| *ygeY* | CDS | predicted peptidase | Cytoplasmic | COG0624;Acetylornithine deacetylase/Succinyl-diaminopimelate desuccinylase and related deacylases | | gg | tt |
| *ygfA* | CDS | predicted ligase | Cytoplasmic | | | gg | o |
| *ygfB* | CDS | predicted protein | Cytoplasmic | | | | |
| *ygfF* | CDS | predicted NAD(P)-binding oxidoreductase with NAD(P)-binding Rossmann-fold domain | Cytoplasmic | COG1028;Dehydrogenases with different specificities (related to short-chain alcohol dehydrogenases) | | gg | t |
| *ygfG* | CDS | methylmalonyl-CoA decarboxylase, biotin- | Cytoplasmic | | | gg | t |
| *ygfH* | CDS | propionyl-CoA:succinate-CoA transferase | Cytoplasmic | COG0427;Acetyl-CoA hydrolase | | gg | t |
| *ygfI* | CDS | predicted DNA-binding transcriptional regulator | Cytoplasmic | | GO:0006350 transcription | | |
| *ygfJ* | CDS | conserved protein | Cytoplasmic | COG2068;Uncharacterized MobA-related protein | | g | o |
| *ygfK* | CDS | predicted oxidoreductase, Fe-S subunit | Cytoplasmic | COG0493;NADPH-dependent glutamate synthase beta chain and related oxidoreductases | | | |
| *ygfM* | CDS | predicted oxidoreductase | Cytoplasmic | COG1319;Aerobic-type carbon monoxide dehydrogenase, middle subunit CoxM/CutM homologs | | o | t |
| *ygfO* | CDS | predicted transporter | Integral Membrane Protein | COG2233;Xanthine/uracil permeases | | o | tt |
| *ygfQ* | CDS | predicted transporter | Integral Membrane Protein | | | | |
| *ygfS* | CDS | predicted oxidoreductase, 4Fe-4S ferredoxin-type subunit | Cytoplasmic | | | | |
| *ygfT* | CDS | fused predicted oxidoreductase Fe-S subunit -!- nucleotide-binding subunit | Cytoplasmic | COG1142;Fe-S-cluster-containing hydrogenase components 2 -!- COG0493;NADPH-dependent glutamate synthase beta chain and related oxidoreductases | | | |
| *ygfU* | CDS | predicted transporter | Integral Membrane Protein | COG2233;Xanthine/uracil permeases | | | |
| *ygfX* | CDS | predicted protein | Membrane Anchored | | | o | tt |
| *ygfY* | CDS | conserved protein | Cytoplasmic | COG2938;Uncharacterized conserved protein | | gg | o |
| *ygfZ* | CDS | predicted folate-dependent regulatory protein | Cytoplasmic | COG0354;Predicted aminomethyltransferase related to GcvT | | gg | o |
| *yggC* | CDS | conserved protein with nucleoside triphosphate hydrolase domain | Cytoplasmic | COG1072;Panthothenate kinase | | gg | tt |
| *yggD* | CDS | predicted DNA-binding transcriptional regulator | Cytoplasmic | COG3722;Transcriptional regulator | | gg | t |
| *yggE* | CDS | conserved protein | Periplasmic | COG2968;Uncharacterized conserved protein | | o | tt |
| *yggF* | CDS | predicted hexoseP phosphatase | Cytoplasmic | COG1494;Fructose-1,6-bisphosphatase/sedoheptulose 1,7-bisphosphatase and related proteins | | g | t |
| *yggG* | CDS | predicted peptidase | Outer Membrane Lipoprotein | COG0501;Zn-dependent protease with chaperone function | | gg | tt |
| *yggH* | CDS | tRNA (m7G46) methyltransferase, SAM-dependent | Cytoplasmic | COG0220;Predicted S-adenosylmethionine-dependent methyltransferase | | gg | t |
| *yggJ* | CDS | predicted protein | Cytoplasmic | | | g | t |
| *yggL* | CDS | predicted protein | Cytoplasmic | | | o | tt |
| *yggM* | CDS | conserved protein | Periplasmic | | | o | tt |
| *yggN* | CDS | predicted protein | Periplasmic | | | o | tt |
| *yggP* | CDS | predicted dehydrogenase | Cytoplasmic | | | | |
| *yggR* | CDS | predicted transporter | Cytoplasmic | COG2805;Tfp pilus assembly protein, pilus retraction ATPase PilT | | o | tt |
| *yggS* | CDS | predicted enzyme | Cytoplasmic | COG0325;Predicted enzyme with a TIM-barrel fold | | o | tt |
| *yggT* | CDS | predicted inner membrane protein | Integral Membrane Protein | COG0762;Predicted integral membrane protein | | o | tt |
| *yggU* | CDS | conserved protein | Cytoplasmic | COG1872;Uncharacterized conserved protein | | | |
| *yggV* | CDS | dITP/XTP pyrophosphatase | Cytoplasmic | COG0127;Xanthosine triphosphate pyrophosphatase | | gg | o |
| *yggW* | CDS | predicted oxidoreductase | Cytoplasmic | COG0635;Coproporphyrinogen III oxidase and related Fe-S | | g | tt |
| *yggX* | CDS | protein that protects iron-sulfur proteins against oxidative damage | Cytoplasmic | COG2924;Fe-S cluster protector protein | | gg | o |
| *yghA* | CDS | predicted glutathionylspermidine synthase, with NAD(P)-binding Rossmann-fold domain | Cytoplasmic | COG1028;Dehydrogenases with different specificities (related to short-chain alcohol dehydrogenases) | | g | o |
| *yghB* | CDS | conserved inner membrane protein | Integral Membrane Protein | COG0586;Uncharacterized membrane-associated protein | | o | tt |
| *yghD* | CDS | predicted secretion pathway M-type protein, membrane anchored | Membrane Anchored | COG3149;Type II secretory pathway, component PulM | | g | tt |
| *yghE* | CDS | predicted secretion pathway protein, L-type protein (pseudogene) | Cytoplasmic | COG3297;Type II secretory pathway, component PulL | | | |
| *yghF* | CDS | predicted secretion pathway protein, C-type protein | Cytoplasmic | COG3031;Type II secretory pathway, component PulC | | | |
| *yghG* | CDS | predicted protein | Outer Membrane Lipoprotein | | | o | tt |
| *yghJ* | CDS | predicted inner membrane lipoprotein | Inner Membrane Lipoprotein | | | | |
| *yghK* | CDS | glycolate transporter | Integral Membrane Protein | COG1620;L-lactate permease | GO:0016052 carbohydrate | o | tt |
| *yghO* | CDS | predicted DNA-binding transcriptional regulator | Cytoplasmic | | | | |
| *yghQ* | CDS | predicted inner membrane protein | Integral Membrane Protein | | | | |
| *yghR* | CDS | predicted protein with nucleoside triphosphate hydrolase domain | Cytoplasmic | | | o | tt |
| *yghS* | CDS | predicted protein with nucleoside triphosphate hydrolase domain | Cytoplasmic | | | | |
| *yghT* | CDS | predicted protein with nucleoside triphosphate hydrolase domain | Cytoplasmic | | | g | t |
| *yghU* | CDS | predicted S-transferase | Membrane Anchored | | | | |
| *yghW* | CDS | predicted protein | Cytoplasmic | | | gg | o |
| *yghX* | CDS | predicted hydrolase (pseudogene) | Cytoplasmic | | | | |
| *yghY* | CDS | predicted dienlactone hydrolase (pseudogene) | Cytoplasmic | | | | |
| *yghZ* | CDS | aldo-keto reductase | Cytoplasmic | COG0667;Predicted oxidoreductases (related to aryl-alcohol dehydrogenases) | | gg | t |
| *ygiA* | CDS | predicted protein | Cytoplasmic | | | | |
| *ygiB* | CDS | conserved outer membrane protein | Cytoplasmic | | | | |
| *ygiC* | CDS | predicted enzyme | Cytoplasmic | COG0754;Glutathionylspermidine synthase | | o | o |
| *ygiE* | CDS | zinc transporter | Integral Membrane Protein | COG0428;Predicted divalent heavy-metal cations transporter | | o | tt |
| *ygiF* | CDS | predicted adenylate cyclase | Cytoplasmic | COG3025;Uncharacterized conserved protein | | g | t |
| *ygiH* | CDS | conserved inner membrane protein | Integral Membrane Protein | COG0344;Predicted membrane protein | | o | tt |
| *ygiL* | CDS | predicted fimbrial-like adhesin protein | Periplasmic | COG3539;P pilus assembly protein, pilin FimA | | gg | tt |
| *ygiN* | CDS | quinol monooxygenase | Cytoplasmic | COG1359;Uncharacterized conserved protein | | g | t |
| *ygiP* | CDS | predicted DNA-binding transcriptional regulator | Cytoplasmic | COG0583;Transcriptional regulator | GO:0006350 transcription | g | tt |
| *ygiQ* | CDS | conserved protein | Cytoplasmic | | | | |
| *ygiS* | CDS | predicted transporter subunit -!- periplasmic-binding component of ABC superfamily | Periplasmic | COG4166;ABC-type oligopeptide transport system, periplasmic component | | o | tt |
| *ygiT* | CDS | predicted DNA-binding transcriptional regulator | Cytoplasmic | COG1396;Predicted transcriptional regulators | | | |
| *ygiU* | CDS | predicted cyanide hydratase | Cytoplasmic | | | | |
| *ygiV* | CDS | predicted transcriptional regulator | Cytoplasmic | | | | |
| *ygiW* | CDS | conserved protein | Periplasmic | COG3111;Uncharacterized conserved protein | | o | tt |
| *ygiZ* | CDS | conserved inner membrane protein | Integral Membrane Protein | | | | |
| *ygjD* | CDS | predicted peptidase | Cytoplasmic | COG0533;Metal-dependent proteases with possible chaperone activity | | gg | t |
| *ygjE* | CDS | predicted tartrate:succinate antiporter | Integral Membrane Protein | COG0471;Di- and tricarboxylate transporters | | o | tt |
| *ygjF* | CDS | G/U mismatch-specific DNA glycosylase | Cytoplasmic | COG3663;G:T/U mismatch-specific DNA glycosylase | GO:0006281 DNA repair | gg | t |
| *ygjG* | CDS | putrescine:2-oxoglutaric acid aminotransferase, PLP-dependent | Periplasmic | | GO:0006526 arginine biosynthesis -!- GO:0008615 pyridoxine biosynthesis | | |
| *ygjH* | CDS | conserved protein | Cytoplasmic | COG0073;EMAP domain | GO:0006418 amino acid activation | gg | o |
| *ygjI* | CDS | predicted transporter | Integral Membrane Protein | | | | |
| *ygjJ* | CDS | conserved protein | Periplasmic | | | o | tt |
| *ygjK* | CDS | predicted glycosyl hydrolase | Periplasmic | | | g | tt |
| *ygjM* | CDS | predicted DNA-binding transcriptional regulator | Cytoplasmic | | GO:0006355 regulation of transcription, DNA-dependent | gg | o |
| *ygjN* | CDS | conserved protein | Cytoplasmic | COG4680;Uncharacterized protein conserved in bacteria | | o | t |
| *ygjO* | CDS | predicted methyltransferase small domain | Cytoplasmic | | | | |
| *ygjP* | CDS | predicted metal dependent hydrolase | Cytoplasmic | COG1451;Predicted metal-dependent hydrolase | | | |
| *ygjQ* | CDS | predicted thioredoxin-like | Periplasmic | COG2949;Uncharacterized membrane protein | | o | tt |
| *ygjR* | CDS | predicted NAD(P)-binding dehydrogenase | Cytoplasmic | COG0673;Predicted dehydrogenases and related proteins | | gg | t |
| *ygjV* | CDS | conserved inner membrane protein | Integral Membrane Protein | | | o | tt |
| *yhaB* | CDS | predicted protein | Cytoplasmic | | | g | o |
| *yhaC* | CDS | predicted protein | Cytoplasmic | | | gg | o |
| *yhaH* | CDS | predicted inner membrane protein | Integral Membrane Protein | COG3152;Predicted membrane protein | | o | tt |
| *yhaI* | CDS | predicted inner membrane protein | Integral Membrane Protein | COG3152;Predicted membrane protein | | o | tt |
| *yhaJ* | CDS | predicted DNA-binding transcriptional regulator | Cytoplasmic | COG0583;Transcriptional regulator | GO:0006350 transcription | o | tt |
| *yhaK* | CDS | predicted pirin-related protein | Cytoplasmic | COG1741;Pirin-related protein | | g | t |
| *yhaL* | CDS | predicted protein | Cytoplasmic | | | | |
| *yhaM* | CDS | conserved protein | Cytoplasmic | | | | |
| *yhaO* | CDS | predicted transporter | Integral Membrane Protein | | | | |
| *yhaV* | CDS | conserved protein | Cytoplasmic | | | o | tt |
| *yhbC* | CDS | conserved protein | Cytoplasmic | | | | |
| *yhbE* | CDS | conserved inner membrane protein | Integral Membrane Protein | COG0697;Permeases of the drug/metabolite transporter (DMT) superfamily | | o | tt |
| *yhbG* | CDS | predicted transporter subunit -!- ATP-binding component of ABC superfamily | Cytoplasmic | COG1137;ABC-type (unclassified) transport system, ATPase component | | g | o |
| *yhbH* | CDS | predicted ribosome-associated, sigma 54 modulation protein | Cytoplasmic | COG1544;Ribosome-associated protein Y (PSrp-1) | GO:0006350 transcription | gg | o |
| *yhbJ* | CDS | predicted protein with nucleoside triphosphate hydrolase domain | Cytoplasmic | COG1660;Predicted P-loop-containing kinase | | gg | o |
| *yhbN* | CDS | predicted transporter subunit -!- periplasmic-binding component of ABC superfamily | Periplasmic | COG1934;Uncharacterized protein conserved in bacteria | | o | tt |
| *yhbO* | CDS | predicted intracellular protease | Cytoplasmic | COG0693;Putative intracellular protease/amidase | | | |
| *yhbP* | CDS | conserved protein | Cytoplasmic | COG3787;Uncharacterized protein conserved in bacteria | | o | tt |
| *yhbQ* | CDS | predicted endonuclease | Cytoplasmic | COG2827;Predicted endonuclease containing a URI domain | | gg | t |
| *yhbS* | CDS | predicted acyltransferase with acyl-CoA N-acyltransferase domain | Cytoplasmic | COG3153;Predicted acetyltransferase | | gg | t |

| | | | | | | | |
|---|---|---|---|---|---|---|---|
| *yhbT* | CDS | predicted lipid carrier protein | Cytoplasmic | COG3154;Putative lipid carrier protein | | o | tt |
| *yhbU* | CDS | predicted peptidase (collagenase-like) | Cytoplasmic | COG0826;Collagenase and related proteases | | gg | t |
| *yhbV* | CDS | predicted protease | Cytoplasmic | COG0826;Collagenase and related proteases | | | |
| *yhbW* | CDS | predicted enzyme | Cytoplasmic | COG2141;Coenzyme F420-dependent N5,N10-methylene tetrahydromethanopterin reductase and related flavin-dependent oxidoreductases | | g | t |
| *yhbX* | CDS | predicted hydrolase, inner membrane | Integral Membrane Protein | | | | |
| *yhbY* | CDS | predicted RNA-binding protein | Cytoplasmic | COG1534;Predicted RNA-binding protein containing KH domain, possibly ribosomal protein | | gg | t |
| *yhcA* | CDS | predicted periplasmic chaperone protein | Periplasmic | COG3121;P pilus assembly protein, chaperone PapD | GO:0006457 protein folding | o | tt |
| *yhcB* | CDS | conserved protein | Membrane Anchored | COG3105;Uncharacterized protein conserved in bacteria | | | |
| *yhcC* | CDS | predicted Fe-S oxidoreductase | Cytoplasmic | COG1242;Predicted Fe-S oxidoreductase | | gg | t |
| *yhcD* | CDS | predicted outer membrane protein | Outer Membrane B-barrel protein | COG3188;P pilus assembly protein, porin PapC | | o | tt |
| *yhcE* | CDS | predicted protein, N-ter fragment (pseudogene) | Periplasmic | | | g | o |
| *yhcE* | ancestral | predicted protein (pseudogene) | Periplasmic | | | | |
| *yhcE* | CDS | predicted protein, C-ter fragment (pseudogene) | Cytoplasmic | | | g | o |
| *yhcF* | CDS | predicted transcriptional regulator | Periplasmic | | GO:0006350 transcription -!- GO:0006355 regulation of transcription, DNA-dependent | o | tt |
| *yhcG* | CDS | conserved protein | Cytoplasmic | COG4804;Uncharacterized conserved protein | | o | t |
| *yhcH* | CDS | conserved protein | Cytoplasmic | COG2731;Beta-galactosidase, beta subunit | | g | o |
| *yhcM* | CDS | conserved protein with nucleoside triphosphate hydrolase domain | Cytoplasmic | COG1485;Predicted ATPase | | o | tt |
| *yhcN* | CDS | conserved protein | Periplasmic | | | | |
| *yhcO* | CDS | predicted barnase inhibitor | Cytoplasmic | COG2732;Barstar, RNAse (barnase) inhibitor | | gg | t |
| *yhdA* | CDS | conserved inner membrane protein | Integral Membrane Protein | COG2199;FOG: GGDEF domain -!- COG2200;FOG: EAL domain | | o | tt |
| *yhdE* | CDS | conserved protein | Cytoplasmic | COG0424;Nucleotide-binding protein implicated in inhibition of septum formation | | gg | t |
| *yhdH* | CDS | predicted oxidoreductase, Zn-dependent and NAD(P)-binding | Cytoplasmic | COG0604;NADPH:quinone reductase and related Zn-dependent oxidoreductases | | gg | t |
| *yhdJ* | CDS | predicted methyltransferase | Cytoplasmic | | | | |
| *yhdL* | CDS | conserved protein | Cytoplasmic | | | o | o |
| *yhdN* | CDS | conserved protein | Cytoplasmic | | | gg | t |
| *yhdP* | CDS | conserved membrane protein, predicted transporter | Periplasmic | | | | |
| *yhdT* | CDS | conserved inner membrane protein | Integral Membrane Protein | COG3924;Predicted membrane protein | | o | tt |
| *yhdU* | CDS | predicted membrane protein | Membrane Anchored | | | o | tt |
| *yhdV* | CDS | predicted outer membrane protein | Outer Membrane Lipoprotein | | | o | tt |
| *yhdW* | CDS | predicted amino-acid transporter subunit -!- periplasmic-binding component of ABC superfamily | Cytoplasmic | COG0834;ABC-type amino acid transport/signal transduction systems, periplasmic component/domain | GO:0009063 amino acid catabolism | g | o |
| *yhdX* | CDS | predicted amino-acid transporter subunit -!- membrane component of ABC superfamily | Integral Membrane Protein | | | | |
| *yhdY* | CDS | predicted amino-acid transporter subunit -!- membrane component of ABC superfamily | Integral Membrane Protein | COG0765;ABC-type amino acid transport system, permease component | | | |
| *yhdZ* | CDS | predicted amino-acid transporter subunit -!- ATP-binding component of ABC superfamily | Cytoplasmic | COG1126;ABC-type polar amino acid transport system, ATPase component | | g | t |
| *yheL* | CDS | predicted intracellular sulfur oxidation protein | Cytoplasmic | COG2168;Uncharacterized conserved protein involved in oxidation of intracellular sulfur | | g | t |
| *yheM* | CDS | predicted intraceullular sulfur oxidation protein | Cytoplasmic | COG2923;Uncharacterized protein involved in the oxidation of intracellular sulfur | | g | o |
| *yheN* | CDS | predicted intracellular sulfur oxidation protein | Cytoplasmic | COG1553;Uncharacterized conserved protein involved in intracellular sulfur reduction | | g | t |
| *yheO* | CDS | predicted DNA-binding transcriptional regulator | Cytoplasmic | | | | |
| *yheS* | CDS | fused predicted transporter subunits -!- ATP-binding components of ABC superfamily | Cytoplasmic | COG0488;ATPase components of ABC transporters with duplicated ATPase domains | | gg | o |
| *yheT* | CDS | predicted hydrolase | Cytoplasmic | COG0429;Predicted hydrolase of the alpha/beta-hydrolase fold | | o | tt |
| *yheU* | CDS | conserved protein | Cytoplasmic | COG3089;Uncharacterized protein conserved in bacteria | | gg | o |
| *yheV* | CDS | predicted protein | Cytoplasmic | | | gg | o |
| *yhfA* | CDS | conserved protein | Cytoplasmic | COG1765;Predicted redox protein, regulator of disulfide bond formation | | g | o |
| *yhfG* | CDS | predicted protein | Cytoplasmic | | | gg | o |
| *yhfK* | CDS | conserved inner membrane protein | Integral Membrane Protein | COG1289;Predicted membrane protein | GO:0008152 metabolism | | |
| *yhfL* | CDS | conserved secreted peptide | Outer Membrane Lipoprotein | | | o | tt |
| *yhfS* | CDS | conserved protein | Cytoplasmic | | | gg | t |
| *yhfT* | CDS | predicted inner membrane protein | Integral Membrane Protein | | | o | tt |
| *yhfU* | CDS | predicted protein | Membrane Anchored | | | | |
| *yhfW* | CDS | predicted mutase | Cytoplasmic | COG1015;Phosphopentomutase | | gg | t |
| *yhfX* | CDS | predicted amino acid racemase | Cytoplasmic | COG3457;Predicted amino acid racemase | | o | tt |
| *yhfY* | CDS | conserved protein | Cytoplasmic | | | | |
| *yhfZ* | CDS | conserved protein | Cytoplasmic | | | | |
| *yhgA* | CDS | predicted transposase | Cytoplasmic | | | gg | t |
| *yhgE* | CDS | predicted inner membrane protein | Integral Membrane Protein | | | o | tt |
| *yhgF* | CDS | predicted transcriptional accessory protein | Cytoplasmic | | | g | tt |
| *yhgG* | CDS | predicted DNA-binding transcriptional regulator | Cytoplasmic | | GO:0006355 regulation of transcription, DNA-dependent | gg | tt |
| *yhgN* | CDS | predicted antibiotic transporter | Integral Membrane Protein | COG2095;Multiple antibiotic transporter | | o | tt |
| *yhhA* | CDS | conserved protein | Periplasmic | | | o | tt |
| *yhhF* | CDS | predicted methyltransferase | Cytoplasmic | COG0742;N6-adenine-specific methylase | | gg | t |
| *yhhH* | CDS | predicted protein | Membrane Anchored | | | o | o |
| *yhhI* | CDS | predicted transposase | Cytoplasmic | | | gg | t |
| *yhhJ* | CDS | predicted transporter subunit -!- membrane component of ABC superfamily | Integral Membrane Protein | | | | |
| *yhhK* | CDS | conserved protein | Cytoplasmic | | | gg | o |
| *yhhL* | CDS | conserved inner membrane protein | Integral Membrane Protein | COG3776;Predicted membrane protein | | | |
| *yhhM* | CDS | conserved protein | Membrane Anchored | | | o | tt |
| *yhhN* | CDS | conserved inner membrane protein | Integral Membrane Protein | COG3714;Predicted membrane protein | | o | tt |
| *yhhP* | CDS | conserved protein required for cell growth | Cytoplasmic | COG0425;Predicted redox protein, regulator of disulfide bond formation | | gg | t |
| *yhhQ* | CDS | conserved inner membrane protein | Integral Membrane Protein | COG1738;Uncharacterized conserved protein | | o | tt |
| *yhhS* | CDS | predicted transporter | Integral Membrane Protein | COG0477;Permeases of the major facilitator superfamily | | | |
| *yhhT* | CDS | predicted inner membrane protein | Integral Membrane Protein | | | | |
| *yhhW* | CDS | predicted protein | Cytoplasmic | COG1741;Pirin-related protein | | g | t |
| *yhhX* | CDS | predicted oxidoreductase with NAD(P)-binding Rossmann-fold domain | Cytoplasmic | COG0673;Predicted dehydrogenases and related proteins | | gg | t |
| *yhhY* | CDS | predicted acetyltransferase | Cytoplasmic | COG0454;Histone acetyltransferase HPA2 and related acetyltransferases | | g | t |
| *yhhZ* | CDS | conserved protein | Cytoplasmic | COG3157;Hemolysin-coregulated protein (uncharacterized) | | g | o |
| *yhiD* | CDS | predicted Mg(2+) transport ATPase inner membrane protein | Integral Membrane Protein | COG1285;Uncharacterized membrane protein | | | |
| *yhiF* | CDS | predicted DNA-binding ranscriptional regulator | Cytoplasmic | COG2771;DNA-binding HTH domain-containing proteins | GO:0006355 regulation of transcription, DNA-dependent | gg | t |
| *yhiI* | CDS | predicted HlyD family secretion protein | Membrane Anchored | COG0845;Membrane-fusion protein | | o | tt |
| *yhiJ* | CDS | predicted protein | Cytoplasmic | | | gg | o |
| *yhiK* | CDS | predicted protein (pseudogene) | Cytoplasmic | | | | |
| *yhiL* | CDS | predicted protein | Cytoplasmic | | | o | o |
| *yhiM* | CDS | conserved inner membrane protein | Integral Membrane Protein | | | | |
| *yhiN* | CDS | predicted oxidoreductase, FAD/NAD(P)-binding domain | Periplasmic | COG2081;Predicted flavoproteins | | g | t |
| *yhiO* | CDS | predicted universal stress (ethanol tolerance) protein B | Integral Membrane Protein | | GO:0006950 response to stress -!- GO:0006805 xenobiotic metabolism | o | tt |
| *yhiP* | CDS | predicted transporter | Integral Membrane Protein | COG3104;Dipeptide/tripeptide permease | | o | tt |
| *yhiQ* | CDS | predicted SAM-dependent methyltransferase | Cytoplasmic | COG0500;SAM-dependent methyltransferases | | | |
| *yhiR* | CDS | predicted DNA (exogenous) processing protein | Cytoplasmic | COG2961;Protein involved in catabolism of external DNA | | gg | o |
| *yhiS* | CDS | predicted protein | Cytoplasmic | | | o | o |
| *yhjA* | CDS | predicted cytochrome C peroxidase | Membrane Anchored | COG1858;Cytochrome c peroxidase | GO:0017004 cytochrome biogenesis | o | tt |
| *yhjB* | CDS | predicted DNA-binding response regulator in two-component regulatory system | Cytoplasmic | COG2197;Response regulator containing a CheY-like receiver domain and an HTH DNA-binding domain | GO:0006350 transcription | gg | t |
| *yhjC* | CDS | predicted DNA-binding transcriptional regulator | Cytoplasmic | COG0583;Transcriptional regulator | GO:0006350 transcription | o | t |
| *yhjD* | CDS | conserved inner membrane protein | Integral Membrane Protein | COG1295;Predicted membrane protein | | o | tt |
| *yhjE* | CDS | predicted transporter | Integral Membrane Protein | COG0477;Permeases of the major facilitator superfamily | | o | tt |
| *yhjG* | CDS | predicted outer membrane biogenesis protein | Membrane Anchored | COG2982;Uncharacterized protein involved in outer membrane biogenesis | | o | tt |
| *yhjH* | CDS | EAL domain containing protein involved in flagellar function | Cytoplasmic | COG2200;FOG: EAL domain | | g | o |
| *yhjJ* | CDS | predicted zinc-dependent peptidase | Periplasmic | COG0612;Predicted Zn-dependent peptidases | | o | tt |
| *yhjK* | CDS | predicted diguanylate cyclase | Integral Membrane Protein | COG3706;Response regulator containing a CheY-like receiver domain and a GGDEF domain -!- COG2200;FOG: EAL domain | | | |
| *yhjQ* | CDS | cell division protein (chromosome partitioning ATPase) (pseudogene) | Cytoplasmic | COG1192;ATPases involved in chromosome partitioning | | | |
| *yhjR* | CDS | conserved protein | Cytoplasmic | | | gg | t |
| *yhjV* | CDS | predicted transporter | Integral Membrane Protein | COG0814;Amino acid permeases | | o | tt |
| *yhjX* | CDS | predicted transporter | Integral Membrane Protein | COG0477;Permeases of the major facilitator superfamily | GO:0042493 response to drug | o | tt |
| *yhjY* | CDS | conserved protein | Periplasmic | | GO:0019395 fatty acid oxidation | | |
| *yiaA* | CDS | conserved inner membrane protein | Integral Membrane Protein | | | o | tt |
| *yiaB* | CDS | conserved inner membrane protein | Integral Membrane Protein | COG4682;Predicted membrane protein | | | |
| *yiaC* | CDS | predicted acyltransferase with acyl-CoA N-acyltransferase domain | Cytoplasmic | COG0454;Histone acetyltransferase HPA2 and related acetyltransferases | | gg | t |
| *yiaD* | CDS | predicted outer membrane lipoprotien | Outer Membrane Lipoprotein | | | | |
| *yiaF* | CDS | conserved protein | Membrane Lipoprotein | | | | |
| *yiaG* | CDS | predicted transcriptional regulator | Cytoplasmic | COG2944;Predicted transcriptional regulator | GO:0006355 regulation of transcription, DNA-dependent | gg | o |

| | | | | | | | |
|---|---|---|---|---|---|---|---|
| *yiaH* | CDS | conserved inner membrane protein | Integral Membrane Protein | COG3274;Uncharacterized protein conserved in bacteria | | o | tt |
| *yiaI* | CDS | predicted hydrogenase, 4Fe-4S ferredoxin-type component | Cytoplasmic | COG1142;Fe-S-cluster-containing hydrogenase components 2 | | gg | t |
| *yiaJ* | CDS | predicted DNA-binding transcriptional repressor | Cytoplasmic | COG1414;Transcriptional regulator | GO:0006350 transcription | gg | tt |
| *yiaK* | CDS | 2,3-diketo-L-gulonate dehydrogenase, NADH-dependent | Cytoplasmic | COG2055;Malate/L-lactate dehydrogenases | | g | t |
| *yiaL* | CDS | conserved protein | Cytoplasmic | COG2731;Beta-galactosidase, beta subunit | | o | t |
| *yiaM* | CDS | predicted transporter | Integral Membrane Protein | COG3090;TRAP-type C4-dicarboxylate transport system, small permease component | | o | tt |
| *yiaN* | CDS | predicted transporter | Integral Membrane Protein | | | | |
| *yiaO* | CDS | predicted transporter | Periplasmic | COG1638;TRAP-type C4-dicarboxylate transport system, periplasmic component | | o | tt |
| *yiaT* | CDS | predicted protein | Outer Membrane B-barrel protein | COG3713;Outer membrane protein V | | o | tt |
| *yiaU* | CDS | predicted DNA-binding transcriptional regulator | Cytoplasmic | COG0583;Transcriptional regulator | GO:0006350 transcription | gg | t |
| *yiaV* | CDS | membrane fusion protein (MFP) component of efflux pump, signal anchor | Integral Membrane Protein | COG1566;Multidrug resistance efflux pump | | o | tt |
| *yiaW* | CDS | conserved inner membrane protein | Integral Membrane Protein | | | o | tt |
| *yiaY* | CDS | predicted Fe-containing alcohol dehydrogenase | Cytoplasmic | | | | |
| *yibA* | CDS | lyase containing HEAT-repeat | Cytoplasmic | COG1413;FOG: HEAT repeat | | o | tt |
| *yibD* | CDS | predicted glycosyl transferase | Cytoplasmic | COG0463;Glycosyltransferases involved in cell wall biogenesis | | o | tt |
| *yibF* | CDS | predicted glutathione S-transferase | Cytoplasmic | COG0625;Glutathione S-transferase | | gg | o |
| *yibG* | CDS | conserved protein | Periplasmic | | | o | tt |
| *yibH* | CDS | predicted protein | Membrane Anchored | COG1566;Multidrug resistance efflux pump | | o | tt |
| *yibI* | CDS | predicted inner membrane protein | Integral Membrane Protein | | | o | tt |
| *yibJ* | CDS | predicted Rhs-family protein | Cytoplasmic | | | | |
| *yibK* | CDS | predicted rRNA methylase | Cytoplasmic | COG0219;Predicted rRNA methylase (SpoU class) | GO:0009451 RNA modification | g | o |
| *yibL* | CDS | conserved protein | Cytoplasmic | | | gg | o |
| *yibN* | CDS | predicted rhodanese-related sulfurtransferase | Membrane Anchored | COG0607;Rhodanese-related sulfurtransferase | | o | tt |
| *yibQ* | CDS | predicted polysaccharide deacetylase | Periplasmic | | | | |
| *yibT* | CDS | predicted protein | Cytoplasmic | | | gg | o |
| *yicC* | CDS | conserved protein | Cytoplasmic | COG1561;Uncharacterized stress-induced protein | | gg | tt |
| *yicE* | CDS | predicted transporter | Integral Membrane Protein | COG2233;Xanthine/uracil permeases | | o | tt |
| *yicG* | CDS | conserved inner membrane protein | Integral Membrane Protein | | | o | tt |
| *yicH* | CDS | conserved protein | Membrane Anchored | | | o | tt |
| *yicI* | CDS | predicted alpha-glucosidase | Cytoplasmic | COG1501;Alpha-glucosidases, family 31 of glycosyl hydrolases | | o | tt |
| *yicJ* | CDS | predicted transporter | Integral Membrane Protein | | | | |
| *yicL* | CDS | predicted inner membrane protein | Integral Membrane Protein | COG0697;Permeases of the drug/metabolite transporter (DMT) superfamily | | o | tt |
| *yicM* | CDS | predicted transporter | Integral Membrane Protein | COG2814;Arabinose efflux permease | | | |
| *yicN* | CDS | conserved protein | Membrane Anchored | | | | |
| *yicO* | CDS | predicted xanthine/uracil permease | Integral Membrane Protein | COG2252;Permeases | | | |
| *yicR* | CDS | protein associated with replication fork, possible DNA repair protein | Cytoplasmic | COG2003;DNA repair proteins | GO:0006261 DNA dependent DNA replication -!- GO:0006281 DNA | | |
| *yicS* | CDS | predicted protein | Periplasmic | | | | |
| *yidA* | CDS | predicted hydrolase | Cytoplasmic | COG0561;Predicted hydrolases of the HAD superfamily | | o | tt |
| *yidB* | CDS | conserved protein | Cytoplasmic | | | gg | t |
| *yidC* | CDS | cytoplasmic insertase into membrane protein, Sec system | Integral Membrane Protein | COG0706;Preprotein translocase subunit YidC | | o | tt |
| *yidD* | CDS | predicted protein | Cytoplasmic | | | g | tt |
| *yidE* | CDS | predicted transporter | Integral Membrane Protein | | | o | tt |
| *yidF* | CDS | predicted DNA-binding transcriptional regulator | Cytoplasmic | COG0641;Arylsulfatase regulator (Fe-S oxidoreductase) | | gg | o |
| *yidG* | CDS | predicted inner membrane protein | Integral Membrane Protein | | | o | tt |
| *yidH* | CDS | conserved inner membrane protein | Integral Membrane Protein | COG2149;Predicted membrane protein | | o | tt |
| *yidI* | CDS | predicted inner membrane protein | Integral Membrane Protein | | | o | tt |
| *yidJ* | CDS | predicted sulfatase/phosphatase | Cytoplasmic | COG3119;Arylsulfatase A and related enzymes | GO:0006790 sulfur metabolism | gg | t |
| *yidK* | CDS | predicted transporter | Integral Membrane Protein | COG4146;Predicted symporter | | o | tt |
| *yidL* | CDS | predicted DNA-binding transcriptional regulator | Cytoplasmic | COG2207;AraC-type DNA-binding domain-containing proteins | GO:0006350 transcription | gg | t |
| *yidP* | CDS | predicted DNA-binding transcriptional regulator | Cytoplasmic | COG2188;Transcriptional regulators | GO:0006350 transcription | g | t |
| *yidQ* | CDS | conserved outer membrane protein | Outer Membrane Lipoprotein | | | | |
| *yidR* | CDS | conserved protein | Cytoplasmic | | | | |
| *yidX* | CDS | predicted lipoproteinC | Outer Membrane Lipoprotein | | | | |
| *yidZ* | CDS | predicted DNA-binding transcriptional regulator | Cytoplasmic | COG0583;Transcriptional regulator | GO:0006350 transcription | g | t |
| *yieE* | CDS | predicted phosphopantetheinyl transferase | Cytoplasmic | COG2091;Phosphopantetheinyl transferase | | gg | o |
| *yieF* | CDS | chromate reductase, Class I, flavoprotein | Cytoplasmic | COG0431;Predicted flavoprotein | | gg | t |
| *yieG* | CDS | predicted inner membrane protein | Integral Membrane Protein | COG2252;Permeases | | o | tt |
| *yieH* | CDS | predicted hydrolase | Cytoplasmic | COG0637;Predicted phosphatase/phosphohexomutase | | gg | tt |
| *yieI* | CDS | predicted inner membrane protein | Integral Membrane Protein | | | o | tt |
| *yieK* | CDS | predicted 6-phosphogluconolactonase | Cytoplasmic | | | | |
| *yieL* | CDS | predicted xylanase | Periplasmic | COG2382;Enterochelin esterase and related enzymes | GO:0016052 carbohydrate | | |
| *yieM* | CDS | predicted von Willibrand factor containing protein | Cytoplasmic | | | | |
| *yieN* | CDS | fused predicted transcriptional regulators | Cytoplasmic | | | g | t |
| *yieP* | CDS | predicted transcriptional regulator | Cytoplasmic | | | | |
| *yifB* | CDS | predicted bifunctional protein, enzyme and transcriptional regulator | Periplasmic | COG0606;Predicted ATPase with chaperone activity | | o | t |
| *yifE* | CDS | conserved protein | Cytoplasmic | COG3085;Uncharacterized protein conserved in bacteria | GO:0006355 regulation of transcription, DNA-dependent | gg | t |
| *yifK* | CDS | predicted transporter | Integral Membrane Protein | | | | |
| *yifL* | CDS | predicted lipoprotein | Membrane Lipoprotein | | | o | tt |
| *yifN* | CDS | conserved protein (pseudogene) | Cytoplasmic | | | gg | o |
| *yifO* | CDS | conserved protein | Cytoplasmic | | | | |
| *yigA* | CDS | conserved protein | Cytoplasmic | COG3159;Uncharacterized protein conserved in bacteria | | gg | t |
| *yigB* | CDS | predicted hydrolase | Cytoplasmic | COG1011;Predicted hydrolase (HAD superfamily) | | gg | o |
| *yigE* | CDS | predicted protein | Periplasmic | | | | |
| *yigF* | CDS | conserved inner membrane protein | Integral Membrane Protein | | | o | tt |
| *yigG* | CDS | predicted inner membrane protein | Integral Membrane Protein | | | | |
| *yigI* | CDS | conserved protein | Cytoplasmic | | | | |
| *yigL* | CDS | predicted hydrolase | Cytoplasmic | | | | |
| *yigM* | CDS | predicted inner membrane protein | Integral Membrane Protein | COG0697;Permeases of the drug/metabolite transporter (DMT) superfamily | | o | tt |
| *yigP* | CDS | conserved protein | Cytoplasmic | COG3165;Uncharacterized protein conserved in bacteria | | gg | t |
| *yigZ* | CDS | predicted elongation factor | Cytoplasmic | | GO:0006412 protein biosynthesis | | |
| *yihA* | CDS | GTP-binding protein | Cytoplasmic | COG0218;Predicted GTPase | GO:0007049 cell cycle | | |
| *yihD* | CDS | conserved protein | Cytoplasmic | COG3084;Uncharacterized protein conserved in bacteria | | gg | o |
| *yihE* | CDS | predicted kinase | Cytoplasmic | COG2334;Putative homoserine kinase type II (protein kinase fold) | | g | tt |
| *yihF* | CDS | conserved protein | Membrane Anchored | | | | |
| *yihG* | CDS | predicted endonuclease | Cytoplasmic | COG0204;1-acyl-sn-glycerol-3-phosphate acyltransferase | GO:0009451 RNA modification -!- GO:0006401 RNA catabolism | o | tt |
| *yihI* | CDS | conserved protein | Cytoplasmic | COG3078;Uncharacterized protein conserved in bacteria | | gg | t |
| *yihL* | CDS | predicted DNA-binding transcriptional regulator | Cytoplasmic | COG2188;Transcriptional regulators | GO:0006350 transcription | o | tt |
| *yihM* | CDS | predicted sugar phosphate isomerase | Cytoplasmic | | | gg | t |
| *yihN* | CDS | predicted transporter | Membrane Lipoprotein | COG0477;Permeases of the major facilitator superfamily | | o | tt |
| *yihO* | CDS | predited transporter | Integral Membrane Protein | | | | |
| *yihP* | CDS | predicted transporter | Integral Membrane Protein | | | o | tt |
| *yihQ* | CDS | alpha-glucosidase | Cytoplasmic | COG1501;Alpha-glucosidases, family 31 of glycosyl hydrolases | | gg | t |
| *yihR* | CDS | predicted aldose-1-epimerase | Cytoplasmic | | | o | t |
| *yihS* | CDS | predicted glucosamine isomerase | Cytoplasmic | | | | |
| *yihT* | CDS | predicted aldolase | Cytoplasmic | COG3684;Tagatose-1,6-bisphosphate aldolase | | gg | t |
| *yihU* | CDS | predicted oxidoreductase with NAD(P)-binding Rossmann-fold domain | Cytoplasmic | COG2084;3-hydroxyisobutyrate dehydrogenase and related beta-hydroxyacid dehydrogenases | | gg | t |
| *yihV* | CDS | predicted sugar kinase | Cytoplasmic | COG0524;Sugar kinases, ribokinase family | | | |
| *yihW* | CDS | predicted DNA-binding transcriptional regulator | Cytoplasmic | COG1349;Transcriptional regulators of sugar metabolism | GO:0006350 transcription | | |
| *yihX* | CDS | predicted hydrolase | Cytoplasmic | | | | |
| *yiiD* | CDS | predicted acetyltransferase | Cytoplasmic | COG0454;Histone acetyltransferase HPA2 and related acetyltransferases | | o | tt |
| *yiiE* | CDS | predicted transcriptional regulator | Cytoplasmic | | | | |
| *yiiF* | CDS | conserved protein | Cytoplasmic | | | | |
| *yiiG* | CDS | conserved protein | Inner Membrane Lipoprotein | | | o | tt |
| *yiiL* | CDS | L-rhamnose mutarotase | Cytoplasmic | COG3254;Uncharacterized conserved protein | | gg | t |
| *yiiM* | CDS | conserved protein | Cytoplasmic | COG2258;Uncharacterized protein conserved in bacteria | | | |
| *yiiQ* | CDS | conserved protein | Periplasmic | | | o | tt |
| *yiiR* | CDS | conserved inner membrane protein | Integral Membrane Protein | COG3152;Predicted membrane protein | | o | tt |
| *yiiS* | CDS | conserved protein | Cytoplasmic | COG3691;Uncharacterized protein conserved in bacteria | | gg | o |
| *yiiT* | CDS | stress-induced protein | Cytoplasmic | COG0589;Universal stress protein UspA and related nucleotide-binding proteins | GO:0006950 response to stress | gg | t |
| *yiiU* | CDS | conserved protein | Cytoplasmic | COG3074;Uncharacterized protein conserved in bacteria | | gg | tt |
| *yiiX* | CDS | predicted peptidoglycan peptidase | Periplasmic | | | o | tt |
| *yijD* | CDS | conserved inner membrane protein | Integral Membrane Protein | | | o | tt |
| *yijE* | CDS | predicted permease | Integral Membrane Protein | | | | |
| *yijF* | CDS | conserved protein | Periplasmic | COG3738;Uncharacterized protein conserved in bacteria | | o | tt |
| *yijO* | CDS | predicted DNA-binding transcriptional regulator | Cytoplasmic | COG2207;AraC-type DNA-binding domain-containing proteins | GO:0006350 transcription | o | tt |
| *yijP* | CDS | conserved inner membrane protein | Integral Membrane Protein | COG2194;Predicted membrane-associated, metal-dependent hydrolase | | o | tt |
| *yjaA* | CDS | conserved protein | Cytoplasmic | | | gg | t |
| *yjaB* | CDS | predicted acetyltransferase | Cytoplasmic | COG0454;Histone acetyltransferase HPA2 and related acetyltransferases | | gg | o |
| *yjaG* | CDS | conserved protein | Cytoplasmic | COG3068;Uncharacterized protein conserved in bacteria | | gg | tt |
| *yjaH* | CDS | conserved protein | Outer Membrane Lipoprotein | | | o | tt |
| *yjbA* | CDS | predicted phosphate starvation inducible protein | Integral Membrane Protein | COG3223;Predicted membrane protein | | o | tt |
| *yjbB* | CDS | predicted transporter | Integral Membrane Protein | COG1283;Na+/phosphate symporter | | o | tt |

| | | | | | | | |
|---|---|---|---|---|---|---|---|
| *yjbC* | CDS | 23S rRNA pseudouridine synthase | Cytoplasmic | COG1187;16S rRNA uridine-516 pseudouridylate synthase and related pseudouridylate synthases | | gg | t |
| *yjbD* | CDS | conserved protein | Cytoplasmic | | | gg | t |
| *yjbE* | CDS | predicted protein | Periplasmic | | | o | tt |
| *yjbF* | CDS | predicted lipoprotein | Outer Membrane Lipoprotein | | | | |
| *yjbG* | CDS | conserved protein | Periplasmic | | | o | tt |
| *yjbH* | CDS | predicted porin | Membrane Lipoprotein | | | o | tt |
| *yjbI* | CDS | conserved protein | Cytoplasmic | COG1357;Uncharacterized low-complexity proteins | | g | o |
| *yjbJ* | CDS | predicted stress response protein | Cytoplasmic | COG3237;Uncharacterized protein conserved in bacteria | | gg | t |
| *yjbL* | CDS | predicted protein | Cytoplasmic | | | gg | o |
| *yjbM* | CDS | predicted protein | Cytoplasmic | | | o | tt |
| *yjbN* | CDS | tRNA-dihydrouridine synthase A | Cytoplasmic | COG0042;tRNA-dihydrouridine synthase | | | |
| *yjbO* | CDS | phage shock protein G | Integral Membrane Protein | | | | |
| *yjbQ* | CDS | conserved protein | Cytoplasmic | COG0432;Uncharacterized conserved protein | | gg | o |
| *yjbR* | CDS | conserved protein | Cytoplasmic | COG2315;Uncharacterized protein conserved in bacteria | | gg | o |
| *yjcB* | CDS | predicted inner membrane protein | Integral Membrane Protein | | | | |
| *yjcC* | CDS | predicted signal transduction protein (EAL domain containing protein) | Integral Membrane Protein | COG4943;Predicted signal transduction protein containing sensor and EAL domains | | o | tt |
| *yjcD* | CDS | predicted permease | Integral Membrane Protein | COG2252;Permeases | | o | tt |
| *yjcE* | CDS | predicted cation/proton antiporter | Integral Membrane Protein | COG0025;NhaP-type Na+/H+ and K+/H+ antiporters | | o | tt |
| *yjcF* | CDS | conserved protein | Cytoplasmic | COG1357;Uncharacterized low-complexity proteins | | o | o |
| *yjcH* | CDS | conserved inner membrane protein involved in acetate transport | Integral Membrane Protein | COG3162;Predicted membrane protein | | o | tt |
| *yjcO* | CDS | conserved protein | Periplasmic | COG0790;FOG: TPR repeat, SEL1 subfamily | | o | tt |
| *yjcP* | CDS | predicted outer membrane factor of efflux pump | Outer Membrane Lipoprotein | COG1538;Outer membrane protein | | o | tt |
| *yjcQ* | CDS | predicted multidrug efflux system component | Integral Membrane Protein | COG1289;Predicted membrane protein | | o | tt |
| *yjcR* | CDS | predicted membrane fusion protein of efflux pump | Membrane Anchored | COG1566;Multidrug resistance efflux pump | | o | tt |
| *yjcS* | CDS | predicted alkyl sulfatase | Periplasmic | COG2015;Alkyl sulfatase and related hydrolases | GO:0006790 sulfur metabolism | | |
| *yjcZ* | CDS | conserved protein | Cytoplasmic | | | | |
| *yjdA* | CDS | conserved protein with nucleoside triphosphate hydrolase domain | Cytoplasmic | COG0699;Predicted GTPases (dynamin-related) | | g | t |
| *yjdC* | CDS | predicted transcriptional regulator | Cytoplasmic | COG1309;Transcriptional regulator | GO:0006355 regulation of transcription, DNA-dependent | | |
| *yjdF* | CDS | conserved inner membrane protein | Integral Membrane Protein | COG3647;Predicted membrane protein | | o | tt |
| *yjdI* | CDS | conserved protein | Cytoplasmic | COG3592;Uncharacterized conserved protein | | gg | o |
| *yjdJ* | CDS | predicted acyltransferase with acyl-CoA N-acyltransferase domain | Cytoplasmic | COG2388;Predicted acetyltransferase | | gg | o |
| *yjdK* | CDS | predicted protein | Cytoplasmic | | | gg | o |
| *yjdL* | CDS | predicted transporter | Integral Membrane Protein | COG3104;Dipeptide/tripeptide permease | | o | tt |
| *yjdO* | CDS | predicted protein | Integral Membrane Protein | | | | |
| *yjdP* | CDS | conserved protein | Periplasmic | | | | |
| *yjeB* | CDS | predicted DNA-binding transcriptional regulator | Cytoplasmic | COG1959;Predicted transcriptional regulator | | o | tt |
| *yjeE* | CDS | ATPase with strong ADP affinity | Cytoplasmic | COG0802;Predicted ATPase or kinase | | gg | o |
| *yjeF* | CDS | predicted carbohydrate kinase | Cytoplasmic | COG0062;Uncharacterized conserved protein -!- COG0063;Predicted sugar kinase | | gg | o |
| *yjeH* | CDS | predicted transporter | Integral Membrane Protein | COG0531;Amino acid transporters | | o | tt |
| *yjeI* | CDS | conserved protein | Outer Membrane Lipoprotein | | | | |
| *yjeJ* | CDS | predicted protein | Cytoplasmic | | | o | t |
| *yjeK* | CDS | predicted lysine aminomutase | Cytoplasmic | COG1509;Lysine 2,3-aminomutase | | gg | t |
| *yjeM* | CDS | predicted transporter | Integral Membrane Protein | COG0531;Amino acid transporters | | | |
| *yjeN* | CDS | predicted protein | Cytoplasmic | | | gg | o |
| *yjeO* | CDS | conserved inner membrane protein | Integral Membrane Protein | | | o | tt |
| *yjeP* | CDS | predicted mechanosensitive channel | Integral Membrane Protein | COG3264;Small-conductance mechanosensitive channel | | o | tt |
| *yjeS* | CDS | predicted Fe-S electron transport protein | Cytoplasmic | COG1600;Uncharacterized Fe-S protein | | gg | t |
| *yjeT* | CDS | conserved inner membrane protein | Integral Membrane Protein | COG3242;Uncharacterized protein conserved in bacteria | | o | tt |
| *yjfC* | CDS | predicted synthetase/amidase | Cytoplasmic | COG0754;Glutathionylspermidine synthase | | g | t |
| *yjfF* | CDS | predicted sugar transporter subunit -!- membrane component of ABC superfamily | Integral Membrane Protein | COG1172;Ribose/xylose/arabinose/galactoside ABC-type transport systems, permease components | | | |
| *yjfI* | CDS | conserved protein | Cytoplasmic | COG3789;Uncharacterized protein conserved in bacteria | | g | t |
| *yjfJ* | CDS | predicted transcriptional regulator effector protein | Cytoplasmic | COG1842;Phage shock protein A (IM30), suppresses sigma54-dependent transcription | | gg | o |
| *yjfK* | CDS | conserved protein | Cytoplasmic | | | gg | t |
| *yjfL* | CDS | conserved inner membrane protein | Integral Membrane Protein | COG3766;Predicted membrane protein | | o | tt |
| *yjfM* | CDS | conserved protein | Cytoplasmic | | | o | tt |
| *yjfN* | CDS | predicted protein | Periplasmic | | | | |
| *yjfO* | CDS | conserved protein | Outer Membrane Lipoprotein | | | | |
| *yjfP* | CDS | predicted hydrolase | Cytoplasmic | COG1073;Hydrolases of the alpha/beta superfamily | | gg | o |
| *yjfY* | CDS | predicted protein | Periplasmic | | | gg | o |
| *yjfZ* | CDS | predicted protein | Cytoplasmic | | | g | o |
| *yjgA* | CDS | conserved protein | Cytoplasmic | COG3028;Uncharacterized protein conserved in bacteria | | gg | t |
| *yjgB* | CDS | predicted alcohol dehydrogenase, Zn-dependent and NAD(P)-binding | Cytoplasmic | | | | |
| *yjgD* | CDS | conserved protein | Cytoplasmic | COG3076;Uncharacterized protein conserved in bacteria | | gg | tt |
| *yjgF* | CDS | ketoacid-binding protein | Cytoplasmic | COG0251;Putative translation initiation inhibitor, yjgF family | | | |
| *yjgH* | CDS | predicted mRNA endoribonuclease | Cytoplasmic | COG0251;Putative translation initiation inhibitor, yjgF family | GO:0006412 protein biosynthesis | g | o |
| *yjgI* | CDS | predicted oxidoreductase with NAD(P)-binding Rossmann-fold domain | Cytoplasmic | COG1028;Dehydrogenases with different specificities (related to short-chain alcohol dehydrogenases) | | gg | o |
| *yjgJ* | CDS | predicted transcriptional regulator | Cytoplasmic | | | o | tt |
| *yjgK* | CDS | conserved protein | Cytoplasmic | | | | |
| *yjgL* | CDS | predicted protein | Cytoplasmic | | | | |
| *yjgM* | CDS | predicted acetyltransferase | Cytoplasmic | | | | |
| *yjgN* | CDS | conserved inner membrane protein | Integral Membrane Protein | | | | |
| *yjgP* | CDS | conserved inner membrane protein | Integral Membrane Protein | COG0795;Predicted permeases | | o | tt |
| *yjgQ* | CDS | conserved inner membrane protein | Integral Membrane Protein | | | | |
| *yjgR* | CDS | predicted ATPase | Cytoplasmic | COG0433;Predicted ATPase | | gg | t |
| *yjgW* | CDS | KpLE2 phage-like element; predicted protein | Cytoplasmic | | | o | t |
| *yjgX* | CDS | KpLE2 phage-like element; predicted protein, C-ter fragment (pseudogene) | Cytoplasmic | | | | |
| *yjgX* | ancestral | KpLE2 phage-like element; predicted protein (pseudogene) | Membrane Anchored | | | | |
| *yjgX* | CDS | KpLE2 phage-like element; predicted protein, middle fragment (pseudogene) | Cytoplasmic | COG2194;Predicted membrane-associated, metal-dependent hydrolase | | gg | o |
| *yjgX* | CDS | KpLE2 phage-like element; predicted protein, N-ter fragment (pseudogene) | Membrane Anchored | | | | |
| *yjgZ* | CDS | KpLE2 phage-like element; predicted protein | Cytoplasmic | | | g | o |
| *yjhA* | CDS | N-acetylnuraminic acid outer membrane channel protein | Periplasmic | | | | |
| *yjhB* | CDS | KpLE2 phage-like element; predicted transporter | Integral Membrane Protein | | | | |
| *yjhC* | CDS | KpLE2 phage-like element; predicted oxidoreductase | Cytoplasmic | COG0673;Predicted dehydrogenases and related proteins | | | |
| *yjhD* | CDS | KpLE2 phage-like element; predicted protein (pseudogene) | Cytoplasmic | | | | |
| *yjhE* | CDS | KpLE2 phage-like element; predicted membrane protein (pseudogene) | Integral Membrane Protein | | | o | tt |
| *yjhF* | CDS | KpLE2 phage-like element; predicted transporter | Integral Membrane Protein | COG2610;H+/gluconate symporter and related permeases | | o | tt |
| *yjhG* | CDS | KpLE2 phage-like element; predicted dehydratase | Cytoplasmic | COG0129;Dihydroxyacid dehydratase/phosphogluconate dehydratase | | o | tt |
| *yjhH* | CDS | KpLE2 phage-like element; predicted lyase/synthase | Cytoplasmic | | | | |
| *yjhI* | CDS | KpLE2 phage-like element; predicted DNA-binding transcriptional regulator | Cytoplasmic | COG1414;Transcriptional regulator | GO:0006350 transcription | gg | o |
| *yjhP* | CDS | KpLE2 phage-like element; predicted methyltransferase | Cytoplasmic | COG0500;SAM-dependent methyltransferases | | o | t |
| *yjhQ* | CDS | KpLE2 phage-like element; predicted | Cytoplasmic | COG3153;Predicted acetyltransferase | GO:0016070 RNA metabolism | o | o |
| *yjhR* | CDS | KpLE2 phage-like element; predicted frameshift suppressor | Cytoplasmic | COG1112;Superfamily I DNA and RNA helicases and helicase subunits | | gg | t |
| *yjhS* | CDS | conserved protein | Cytoplasmic | COG2801;Transposase and inactivated derivatives | | o | o |
| *yjhT* | CDS | conserved protein | Periplasmic | | | | |
| *yjhU* | CDS | KpLE2 phage-like element; predicted DNA-binding transcriptional regulator | Cytoplasmic | COG2390;Transcriptional regulator, contains sigma factor-related N-terminal domain | GO:0006355 regulation of transcription, DNA-dependent | | |
| *yjhV* | CDS | KpLE2 phage-like element; predicted protein | Cytoplasmic | | | gg | o |
| *yjhW* | CDS | KpLE2 phage-like element; predicted transposase fragment (pseudogene) | Cytoplasmic | | | | |
| *yjhX* | CDS | conserved protein | Cytoplasmic | | | | |
| *yjiA* | CDS | predicted GTPase | Cytoplasmic | | GO:0009236 vitamin B12 | | |
| *yjiC* | CDS | predicted protein | Cytoplasmic | | | g | o |
| *yjiD* | CDS | DNA replication/recombination/repair protein | Cytoplasmic | | | | |
| *yjiE* | CDS | predicted DNA-binding transcriptional regulator | Cytoplasmic | COG0583;Transcriptional regulator | GO:0006350 transcription | g | o |
| *yjiG* | CDS | conserved inner membrane protein | Integral Membrane Protein | COG0700;Uncharacterized membrane protein | | o | tt |
| *yjiH* | CDS | conserved inner membrane protein | Integral Membrane Protein | COG3314;Uncharacterized protein conserved in bacteria | | | |
| *yjiJ* | CDS | predicted inner membrane protein | Integral Membrane Protein | COG0477;Permeases of the major facilitator superfamily | | o | tt |
| *yjiK* | CDS | conserved protein | Membrane Anchored | COG3204;Uncharacterized protein conserved in bacteria | | | |
| *yjiL* | CDS | predicted ATPase, activator of (R)-hydroxyglutaryl-CoA dehydratase | Cytoplasmic | COG1924;Activator of 2-hydroxyglutaryl-CoA dehydratase (HSP70-class ATPase domain) | | | |
| *yjiM* | CDS | predicted 2-hydroxyglutaryl-CoA dehydratase | Cytoplasmic | | | | |
| *yjiN* | CDS | conserved inner membrane protein | Integral Membrane Protein | COG2733;Predicted membrane protein | | o | tt |
| *yjiO* | CDS | multidrug efflux system protein | Integral Membrane Protein | COG0477;Permeases of the major facilitator superfamily | | o | tt |
| *yjiP* | CDS | predicted transposase (pseudogene) | Cytoplasmic | | | | |
| *yjiQ* | CDS | predicted transposase | Cytoplasmic | | | gg | o |

| | | | | | | | |
|---|---|---|---|---|---|---|---|
| *yjiR* | CDS | fused predicted DNA-binding transcriptional regulator -!- predicted aminotransferase | Cytoplasmic | COG1167;Transcriptional regulators containing a DNA-binding HTH domain and an aminotransferase domain (MocR family) and their eukaryotic orthologs | GO:0006350 transcription | o | t |
| *yjiS* | CDS | conserved protein | Cytoplasmic | | | gg | tt |
| *yjiT* | CDS | conserved protein | Cytoplasmic | | | | |
| *yjiV* | CDS | conserved protein (pseudogene) | Cytoplasmic | | | | |
| *yjiW* | CDS | conserved protein | Cytoplasmic | | GO:0009432 SOS response | o | tt |
| *yjiX* | CDS | conserved protein | Cytoplasmic | COG2879;Uncharacterized small protein | | gg | o |
| *yjiY* | CDS | predicted inner membrane protein | Integral Membrane Protein | COG1966;Carbon starvation protein, predicted membrane protein | GO:0042594 response to starvation | | |
| *yjiZ* | CDS | predicted transporter | Integral Membrane Protein | COG0477;Permeases of the major facilitator superfamily | | o | tt |
| *yjjA* | CDS | conserved protein | Periplasmic | | | | |
| *yjjB* | CDS | conserved inner membrane protein | Integral Membrane Protein | COG3610;Uncharacterized conserved protein | | o | tt |
| *yjjG* | CDS | predicted hydrolase | Cytoplasmic | COG1011;Predicted hydrolase (HAD superfamily) | | o | tt |
| *yjjI* | CDS | conserved protein | Cytoplasmic | | | gg | t |
| *yjjJ* | CDS | predicted DNA-binding transcriptional regulator | Cytoplasmic | | | o | tt |
| *yjjK* | CDS | fused predicted transporter subunits -!- ATP-binding components of ABC superfamily | Cytoplasmic | COG0488;ATPase components of ABC transporters with duplicated ATPase domains | | gg | t |
| *yjjM* | CDS | predicted DNA-binding transcriptional regulator | Cytoplasmic | | GO:0006355 regulation of transcription, DNA-dependent | | |
| *yjjN* | CDS | predicted oxidoreductase, Zn-dependent and NAD(P)-binding | Cytoplasmic | COG1063;Threonine dehydrogenase and related Zn-dependent dehydrogenases | | | |
| *yjjP* | CDS | predicted inner membrane protein | Integral Membrane Protein | | | | |
| *yjjQ* | CDS | predicted DNA-binding transcriptional regulator | Cytoplasmic | COG2197;Response regulator containing a CheY-like receiver domain and an HTH DNA-binding domain | | o | tt |
| *yjjU* | CDS | predicted esterase | Cytoplasmic | COG4667;Predicted esterase of the alpha-beta hydrolase superfamily | GO:0006355 regulation of transcription, DNA-dependent | gg | tt |
| *yjjV* | CDS | predicted DNase | Cytoplasmic | | | g | o |
| *yjjW* | CDS | predicted pyruvate formate lyase activating enzyme | Cytoplasmic | COG1180;Pyruvate-formate lyase-activating enzyme | GO:0009061 anaerobic respiration -!- GO:0006464 protein modification | g | t |
| *yjjX* | CDS | thiamin metabolism associated protein | Cytoplasmic | COG1986;Uncharacterized conserved protein | | | |
| *yjjY* | CDS | predicted protein | Cytoplasmic | | | o | t |
| *yjjZ* | CDS | predicted protein | Integral Membrane Protein | | | | |
| *yjtD* | CDS | predicted rRNA methyltransferase | Cytoplasmic | COG0565;rRNA methylase | GO:0009451 RNA modification | g | t |
| *ykfA* | CDS | CP4-6 prophage; predicted GTP-binding protein | Cytoplasmic | | | o | t |
| *ykfB* | CDS | CP4-6 prophage; predicted protein | Periplasmic | | | o | tt |
| *ykfC* | CDS | CP4-6 prophage; conserved protein | Cytoplasmic | COG3344;Retron-type reverse transcriptase | | | |
| *ykfF* | CDS | CP4-6 prophage; predicted protein | Cytoplasmic | | | | |
| *ykfG* | CDS | CP4-6 prophage; predicted DNA repair protein | Cytoplasmic | COG2003;DNA repair proteins | | gg | o |
| *ykfH* | CDS | predicted protein | Cytoplasmic | | | | |
| *ykfI* | CDS | CP4-6 prophage; toxin of the YkfI-YafW toxin-antitoxin system | Cytoplasmic | | | gg | tt |
| *ykfJ* | CDS | conserved protein | Cytoplasmic | COG1690;Uncharacterized conserved protein | | | |
| *ykgA* | CDS | predicted DNA-binding transcriptional regulator | Cytoplasmic | COG2207;AraC-type DNA-binding domain-containing proteins | GO:0006350 transcription | | |
| *ykgB* | CDS | conserved inner membrane protein | Integral Membrane Protein | COG3059;Predicted membrane protein | | | |
| *ykgC* | CDS | predicted oxidoreductase with FAD/NAD(P)-binding domain and dimerization domain | Cytoplasmic | COG1249;Pyruvate/2-oxoglutarate dehydrogenase complex, dihydrolipoamide dehydrogenase (E3) component, and related enzymes | | | |
| *ykgD* | CDS | predicted DNA-binding transcriptional regulator | Cytoplasmic | COG2207;AraC-type DNA-binding domain-containing proteins | GO:0006350 transcription | o | tt |
| *ykgE* | CDS | predicted oxidoreductase | Cytoplasmic | COG0247;Fe-S oxidoreductase | | | |
| *ykgF* | CDS | predicted amino acid dehydrogenase with NAD(P)-binding domain and ferridoxin-like domain | Cytoplasmic | COG1139;Uncharacterized conserved protein containing a ferredoxin-like domain | | o | tt |
| *ykgG* | CDS | predicted transporter | Cytoplasmic | | | | |
| *ykgH* | CDS | predicted inner membrane protein | Integral Membrane Protein | | | | |
| *ykgI* | CDS | predicted protein | Periplasmic | | | | |
| *ykgJ* | CDS | predicted ferredoxin | Periplasmic | COG0727;Predicted Fe-S-cluster oxidoreductase | | gg | o |
| *ykgK* | CDS | predicted regulator | Cytoplasmic | COG2771;DNA-binding HTH domain-containing proteins | | | |
| *ykgL* | CDS | predicted protein | Cytoplasmic | | | | |
| *ykgM* | CDS | predicted ribosomal protein | Cytoplasmic | | | | |
| *ykgM* | CDS | predicted ribosomal protein | Cytoplasmic | COG0254;Ribosomal protein L31 | | | |
| *ykgN* | CDS | predicted IS protein | Cytoplasmic | | | o | tt |
| *ykiA* | CDS | predicted protein | Cytoplasmic | | | gg | o |
| *ykiB* | CDS | predicted protein | Cytoplasmic | | | | |
| *ylaB* | CDS | conserved inner membrane protein | Integral Membrane Protein | COG4943;Predicted signal transduction protein containing sensor and EAL domains | | | |
| *ylaC* | CDS | predicted inner membrane protein | Integral Membrane Protein | | | | |
| *ylbA* | CDS | conserved protein | Cytoplasmic | COG3257;Uncharacterized protein, possibly involved in glyoxylate | | gg | t |
| *ylbE* | CDS | predicted protein, N-ter fragment (pseudogene) | Cytoplasmic | | | gg | o |
| *ylbE* | ancestral | predicted protein (pseudogene) | Cytoplasmic | | | | |
| *ylbE* | CDS | predicted protein, C-ter fragment (pseudogene) | Cytoplasmic | | | o | o |
| *ylbF* | CDS | conserved protein | Cytoplasmic | | | o | tt |
| *ylbG* | CDS | predicted DNA-binding transcriptional regulator | Cytoplasmic | | | | |
| *ylbH* | CDS | conserved protein, rhs-like | Cytoplasmic | COG3209;Rhs family protein | | o | o |
| *ylcE* | CDS | DLP12 prophage; predicted protein | Cytoplasmic | | | | |
| *ylcG* | CDS | DLP12 prophage; predicted protein | Cytoplasmic | | | | |
| *yliA* | CDS | fused predicted peptide transport subunits -!- ATP-binding components of ABC superfamily | Cytoplasmic | COG1123;ATPase components of various ABC-type transport systems, contain duplicated ATPase | | | |
| *yliB* | CDS | predicted peptide transporter subunit -!- periplasmic-binding component of ABC superfamily | Periplasmic | COG0747;ABC-type dipeptide transport system, periplasmic component | | | |
| *yliC* | CDS | predicted peptide transporter subunit -!- membrane component of ABC superfamily | Integral Membrane Protein | COG0601;ABC-type dipeptide/oligopeptide/nickel transport systems, permease components | | o | tt |
| *yliD* | CDS | predicted peptide transporter subunit -!- membrane component of ABC superfamily | Integral Membrane Protein | COG1173;ABC-type dipeptide/oligopeptide/nickel transport systems, permease components | | o | tt |
| *yliE* | CDS | conserved inner membrane protein | Integral Membrane Protein | COG2200;FOG: EAL domain | | o | tt |
| *yliF* | CDS | predicted diguanylate cyclase | Integral Membrane Protein | COG2199;FOG: GGDEF domain | | o | tt |
| *yliG* | CDS | predicted SAM-dependent methyltransferase | Cytoplasmic | COG0621;2-methylthioadenine synthetase | | gg | t |
| *yliH* | CDS | conserved protein | Cytoplasmic | | | gg | t |
| *yliI* | CDS | predicted dehydrogenase | Periplasmic | COG2133;Glucose/sorbosone dehydrogenases | | o | tt |
| *yliJ* | CDS | predicted glutathione S-transferase | Cytoplasmic | | | gg | t |
| *yliK* | CDS | methylmalonyl-CoA mutase | Cytoplasmic | COG1884;Methylmalonyl-CoA mutase, N-terminal domain/subunit -!- COG2185;Methylmalonyl-CoA mutase, C-terminal domain/subunit (cobalamin-binding) | GO:0019395 fatty acid oxidation | gg | t |
| *yliL* | CDS | predicted protein | Cytoplasmic | | | | |
| *ymbA* | CDS | conserved protein | Cytoplasmic | COG3009;Uncharacterized protein conserved in bacteria | | | |
| *ymcA* | CDS | conserved protein | Periplasmic | | | gg | tt |
| *ymcB* | CDS | conserved protein | Periplasmic | | | gg | o |
| *ymcC* | CDS | predicted outer membrane lipoprotein | Outer Membrane Lipoprotein | | | o | tt |
| *ymcD* | CDS | predicted protein | Membrane Anchored | | | | |
| *ymcE* | CDS | cold shock gene | Membrane Anchored | | | o | tt |
| *ymdA* | CDS | predicted protein | Periplasmic | | | o | tt |
| *ymdB* | CDS | conserved protein | Cytoplasmic | COG2110;Predicted phosphatase homologous to the C-terminal domain of histone macroH2A1 | | o | o |
| *ymdC* | CDS | predicted hydrolase | Cytoplasmic | COG1502;Phosphatidylserine/phosphatidylglycerophos phate/cardiolipin synthases and related enzymes | | | |
| *ymdE* | CDS | predicted protein (pseudogene) | Cytoplasmic | | | | |
| *ymdF* | CDS | conserved protein | Cytoplasmic | | | | |
| *ymfA* | CDS | predicted inner membrane protein | Integral Membrane Protein | | | | |
| *ymfB* | CDS | bifunctional thiamin pyrimidine pyrophosphate hydrolase -!- thiamin pyrophosphate hydrolase | Cytoplasmic | COG0494;NTP pyrophosphohydrolases including oxidative damage repair enzymes | | gg | t |
| *ymfC* | CDS | 23S rRNA pseudouridine synthase | Cytoplasmic | COG1187;16S rRNA uridine-516 pseudouridylate synthase and related pseudouridylate synthases | GO:0009451 RNA modification | gg | o |
| *ymfD* | CDS | e14 prophage; predicted SAM-dependent methyltransferase | Cytoplasmic | COG0500;SAM-dependent methyltransferases | | o | o |
| *ymfE* | CDS | e14 prophage; predicted inner membrane protein | Integral Membrane Protein | | | | |
| *ymfG* | CDS | e14 prophage; predicted excisionase | Cytoplasmic | | | o | tt |
| *ymfH* | CDS | e14 prophage; predicted protein | Integral Membrane Protein | | | o | o |
| *ymfI* | CDS | e14 prophage; predicted protein | Cytoplasmic | | | | |
| *ymfJ* | CDS | e14 prophage; predicted protein | Cytoplasmic | | | gg | o |
| *ymfK* | CDS | e14 prophage; repressor protein phage e14 | Cytoplasmic | COG1974;SOS-response transcriptional repressors (RecA-mediated autopeptidases) | | o | tt |
| *ymfL* | CDS | e14 prophage; predicted DNA-binding transcriptional regulator | Cytoplasmic | | | gg | o |
| *ymfM* | CDS | e14 prophage; predicted protein | Cytoplasmic | | | g | t |
| *ymfN* | CDS | e14 prophage; predicted DNA-binding transcriptional regulator | Cytoplasmic | COG4626;Phage terminase-like protein, large subunit | | o | tt |
| *ymfO* | CDS | e14 prophage; conserved protein | Cytoplasmic | COG4695;Phage-related protein | | g | o |
| *ymfP* | CDS | e14 prophage; conserved protein | Cytoplasmic | COG3299;Uncharacterized homolog of phage Mu protein gp47 | | | |
| *ymfQ* | CDS | e14 prophage; conserved protein | Cytoplasmic | COG3778;Uncharacterized protein conserved in bacteria | | o | t |
| *ymfR* | CDS | e14 prophage; predicted protein | Integral Membrane Protein | | | o | tt |
| *ymfS* | CDS | e14 prophage; predicted protein | Cytoplasmic | | | | |
| *ymfT* | CDS | e14 prophage; predicted DNA-binding transcriptional regulator | Cytoplasmic | | | | |
| *ymgA* | CDS | predicted protein | Cytoplasmic | | | gg | o |
| *ymgB* | CDS | predicted protein | Cytoplasmic | | | | |
| *ymgC* | CDS | predicted protein | Cytoplasmic | | | g | o |
| *ymgD* | CDS | predicted protein | Periplasmic | | | | |
| *ymgE* | CDS | predicted inner membrane protein | Integral Membrane Protein | COG2261;Predicted membrane protein | | o | tt |
| *ymgF* | CDS | predicted protein | Integral Membrane Protein | | | o | tt |
| *ymgG* | CDS | predicted protein | Membrane Anchored | | | | |
| *ymgH* | CDS | predicted protein | Cytoplasmic | | | | |
| *ymjA* | CDS | predicted protein | Cytoplasmic | | | gg | o |

| | | | | | | | |
|---|---|---|---|---|---|---|---|
| ymjB | CDS | predicted protein | Cytoplasmic | | | | |
| ymjC | CDS | predicted oxidoreductase | Cytoplasmic | | | | |
| ynaA | CDS | Rac prophage; predicted tail protein | Cytoplasmic | | | gg | tt |
| ynaE | CDS | Rac prophage; predicted DNA-binding transcriptional regulator | Cytoplasmic | | | o | tt |
| ynaI | CDS | conserved inner membrane protein | Integral Membrane Protein | COG0668;Small-conductance mechanosensitive channel | | o | tt |
| ynaJ | CDS | predicted inner membrane protein | Integral Membrane Protein | | | g | tt |
| ynaK | CDS | Rac prophage; conserved protein | Cytoplasmic | COG1475;Predicted transcriptional regulators | | gg | o |
| ynbA | CDS | predicted inner membrane protein | Integral Membrane Protein | | | o | tt |
| ynbB | CDS | predicted CDP-diglyceride synthase | Integral Membrane Protein | COG4589;Predicted CDP-diglyceride synthetase/phosphatidate cytidylyltransferase | GO:0015949 nucleobase, nucleoside and nucleotide interconversion | o | tt |
| ynbC | CDS | predicted hydrolase | Cytoplasmic | COG2267;Lysophospholipase -!- COG0500;SAM-dependent methyltransferases | | g | t |
| ynbD | CDS | predicted phosphatase, inner membrane protein | Integral Membrane Protein | COG0671;Membrane-associated phospholipid phosphatase -!- COG2453;Predicted protein-tyrosine phosphatase | | o | tt |
| ynbE | CDS | predicted lipoprotein | Outer Membrane Lipoprotein | | | o | tt |
| yncA | CDS | predicted acyltransferase with acyl-CoA N-acyltransferase domain | Cytoplasmic | COG1247;Sortase and related acyltransferases | GO:0042493 response to drug | | |
| yncB | CDS | predicted oxidoreductase, Zn-dependent and NAD(P)-binding | Cytoplasmic | | | | |
| yncC | CDS | predicted DNA-binding transcriptional regulator | Cytoplasmic | COG1802;Transcriptional regulators | GO:0006350 transcription -!- GO:0006355 regulation of transcription, DNA-dependent | o | tt |
| yncD | CDS | predicted iron outer membrane transporter | Outer Membrane B-barrel protein | COG1629;Outer membrane receptor proteins, mostly Fe transport | | o | tt |
| yncE | CDS | conserved protein | Periplasmic | COG3391;Uncharacterized conserved protein | | o | tt |
| yncG | CDS | predicted enzyme | Cytoplasmic | COG0625;Glutathione S-transferase | | g | t |
| yncH | CDS | predicted protein | Cytoplasmic | | | | |
| yncI | CDS | predicted protein | Cytoplasmic | | | o | tt |
| yncJ | CDS | predicted protein | Periplasmic | | | o | tt |
| yncK | CDS | predicted transposase, N-ter fragment (pseudogene) | Cytoplasmic | | | | |
| yncK | ancestral | predicted transposase (pseudogene) | Cytoplasmic | | | | |
| yncK | CDS | predicted transposase, C-ter fragment (pseudogene) | Cytoplasmic | | | | |
| yncM | CDS | predicted protein | Cytoplasmic | | | | |
| yncN | CDS | predicted protein | Cytoplasmic | | | | |
| yneE | CDS | conserved inner membrane protein | Integral Membrane Protein | COG3781;Predicted membrane protein | | | |
| yneF | CDS | predicted diguanylate cyclase | Integral Membrane Protein | COG2199;FOG: GGDEF domain | | | |
| yneG | CDS | conserved protein | Cytoplasmic | | | gg | tt |
| yneH | CDS | predicted glutaminase | Cytoplasmic | COG2066;Glutaminase | GO:0009063 amino acid catabolism | o | tt |
| yneI | CDS | predicted aldehyde dehydrogenase | Cytoplasmic | COG1012;NAD-dependent aldehyde dehydrogenases | | | |
| yneJ | CDS | predicted DNA-binding transcriptional regulator | Cytoplasmic | COG0583;Transcriptional regulator | GO:0006350 transcription | g | t |
| yneK | CDS | predicted protein | Cytoplasmic | | | g | o |
| yneL | CDS | predicted transcriptional regulator | Cytoplasmic | COG2207;AraC-type DNA-binding domain-containing proteins | | | |
| ynfA | CDS | conserved inner membrane protein | Integral Membrane Protein | COG1742;Uncharacterized conserved protein | | o | tt |
| ynfB | CDS | predicted protein | Periplasmic | | | o | tt |
| ynfC | CDS | predicted protein | Inner Membrane Lipoprotein | | | | |
| ynfD | CDS | predicted protein | Periplasmic | | | | |
| ynfE | CDS | oxidoreductase subunit | Periplasmic | COG0243;Anaerobic dehydrogenases, typically selenocysteine-containing | | gg | tt |
| ynfF | CDS | oxidoreductase subunit | Periplasmic | COG0243;Anaerobic dehydrogenases, typically selenocysteine-containing | | | |
| ynfG | CDS | oxidoreductase, Fe-S subunit | Periplasmic | COG0437;Fe-S-cluster-containing hydrogenase components 1 | | g | t |
| ynfH | CDS | oxidoreductase, membrane subunit | Integral Membrane Protein | COG3302;DMSO reductase anchor subunit | | | |
| ynfK | CDS | predicted dethiobiotin synthetase | Cytoplasmic | | GO:0009102 biotin biosynthesis | | |
| ynfL | CDS | predicted DNA-binding transcriptional regulator | Cytoplasmic | COG0583;Transcriptional regulator | GO:0006355 regulation of transcription, DNA-dependent | g | o |
| ynfM | CDS | predicted transporter | Integral Membrane Protein | COG0477;Permeases of the major facilitator superfamily | | o | tt |
| ynfN | CDS | Qin prophage; predicted protein | Cytoplasmic | | | | |
| ynfO | CDS | Qin prophage; predicted protein | Cytoplasmic | | | | |
| ynfP | CDS | predicted protein (pseudogene) | Cytoplasmic | | | | |
| ynhG | CDS | conserved protein | Periplasmic | COG1376;Uncharacterized protein conserved in bacteria | | o | tt |
| yniA | CDS | predicted phosphotransferase/kinase | Cytoplasmic | COG3001;Fructosamine-3-kinase | | g | t |
| yniB | CDS | predicted inner membrane protein | Integral Membrane Protein | | | o | tt |
| yniC | CDS | predicted hydrolase | Cytoplasmic | COG0637;Predicted phosphatase/phosphohexomutase | | gg | t |
| yniD | CDS | predicted protein | Membrane Anchored | | | | |
| ynjA | CDS | conserved protein | Cytoplasmic | COG2128;Uncharacterized conserved protein | | | |
| ynjB | CDS | conserved protein | Periplasmic | COG4134;ABC-type uncharacterized transport system, periplasmic component | | | |
| ynjC | CDS | fused transporter subunits -!- membrane component of ABC superfamily | Integral Membrane Protein | | | | |
| ynjD | CDS | predicted transporter subunit -!- ATP-binding component of ABC superfamily | Cytoplasmic | COG4136;ABC-type uncharacterized transport system, ATPase component | | | |
| ynjE | CDS | predicted thiosulfate sulfur transferase | Periplasmic | | GO:0006790 sulfur metabolism | | |
| ynjF | CDS | predicted phosphatidyl transferase, inner membrane protein | Integral Membrane Protein | COG0558;Phosphatidylglycerophosphate synthase | | o | tt |
| ynjH | CDS | predicted protein | Periplasmic | | | o | tt |
| ynjI | CDS | predicted inner membrane protein | Integral Membrane Protein | | | | |
| yoaA | CDS | conserved protein with nucleoside triphosphate hydrolase domain | Cytoplasmic | COG1199;Rad3-related DNA helicases | | o | tt |
| yoaB | CDS | conserved protein | Cytoplasmic | | | | |
| yoaC | CDS | predicted protein | Cytoplasmic | | | | |
| yoaD | CDS | predicted phosphodiesterase | Membrane Anchored | | | o | tt |
| yoaE | CDS | fused predicted membrane proteins | Integral Membrane Protein | COG0861;Membrane protein TerC, possibly involved in tellurium resistance -!- COG1253;Hemolysins and related proteins containing CBS domains | | o | tt |
| yoaF | CDS | conserved outer membrane protein | Outer Membrane Lipoprotein | COG3042;Putative hemolysin | | o | tt |
| yoaG | CDS | predicted protein | Cytoplasmic | | | gg | o |
| yoaH | CDS | conserved protein | Cytoplasmic | COG3140;Uncharacterized protein conserved in bacteria | | gg | o |
| yoaI | CDS | predicted protein | Cytoplasmic | | | | |
| yobA | CDS | conserved protein | Periplasmic | COG2372;Uncharacterized protein, homolog of Cu resistance protein | | o | tt |
| yobB | CDS | conserved protein | Cytoplasmic | COG0388;Predicted amidohydrolase | | g | o |
| yobD | CDS | conserved inner membrane protein | Integral Membrane Protein | COG4811;Predicted membrane protein | | o | tt |
| yobF | CDS | predicted protein | Cytoplasmic | | | | |
| yobG | CDS | predicted protein | Periplasmic | | | | |
| yobH | CDS | predicted protein | Membrane Anchored | | | | |
| yodA | CDS | conserved metal-binding protein | Periplasmic | COG3443;Predicted periplasmic or secreted protein | | o | tt |
| yodB | CDS | predicted cytochrome | Integral Membrane Protein | | GO:0017004 cytochrome biogenesis | | |
| yodC | CDS | predicted protein | Cytoplasmic | | | gg | t |
| yodD | CDS | predicted protein | Cytoplasmic | | | | |
| yoeA | CDS | CP4-44 prophage; predicted disrupted hemin or colicin receptor (pseudogene) | Outer Membrane B-barrel protein | COG1629;Outer membrane receptor proteins, mostly Fe transport | | | |
| yoeB | CDS | toxin of the YoeB-YefM toxin-antitoxin system | Cytoplasmic | | | | |
| yoeE | CDS | CP4-44 prophage; predicted disrupted hemin or colicin receptor | Cytoplasmic | COG1629;Outer membrane receptor proteins, mostly Fe transport | | gg | o |
| yoeF | CDS | conserved protein | Cytoplasmic | | | | |
| yohC | CDS | predicted inner membrane protein | Integral Membrane Protein | | | | |
| yohD | CDS | conserved inner membrane protein | Integral Membrane Protein | COG0586;Uncharacterized membrane-associated protein | | o | tt |
| yohF | CDS | predicted oxidoreductase with NAD(P)-binding Rossmann-fold domain | Cytoplasmic | COG1028;Dehydrogenases with different specificities (related to short-chain alcohol dehydrogenases) | | g | t |
| yohG | CDS | predicted outer membrane protein | Cytoplasmic | COG1538;Outer membrane protein | | | |
| yohH | CDS | conserved protein | Cytoplasmic | COG1538;Outer membrane protein | | | |
| yohJ | CDS | conserved inner membrane protein | Integral Membrane Protein | COG1380;Putative effector of murein hydrolase LrgA | | o | tt |
| yohK | CDS | predicted inner membrane protein | Integral Membrane Protein | COG1346;Putative effector of murein hydrolase | | o | tt |
| yohL | CDS | conserved protein | Cytoplasmic | COG1937;Uncharacterized protein conserved in bacteria | | gg | o |
| yohM | CDS | membrane protein conferring nickel and cobalt resistance | Integral Membrane Protein | COG2215;ABC-type uncharacterized transport system, permease component | | o | tt |
| yohN | CDS | predicted protein | Periplasmic | | | | |
| yohO | CDS | predicted protein | Membrane Anchored | | | | |
| yojI | CDS | fused predicted multidrug transport subunits -!- membrane component and ATP-binding component of ABC superfamily | Integral Membrane Protein | COG4615;ABC-type siderophore export system, fused ATPase and permease components | | g | tt |
| yojL | CDS | predicted thiamine biosynthesis lipoprotein | Inner Membrane Lipoprotein | COG1477;Membrane-associated lipoprotein involved in thiamine biosynthesis | GO:0009228 thiamin biosynthesis | | |
| ypaA | CDS | predicted protein (pseudogene) | Cytoplasmic | | | | |
| ypdA | CDS | predicted sensory kinase in two-component system with YpdB | Integral Membrane Protein | COG3275;Putative regulator of cell autolysis | | | |
| ypdB | CDS | predicted response regulator in two-component system withYpdA | Cytoplasmic | COG3279;Response regulator of the LytR/AlgR family | GO:0006350 transcription | gg | t |
| ypdC | CDS | predicted DNA-binding protein | Cytoplasmic | COG2207;AraC-type DNA-binding domain-containing proteins | GO:0006350 transcription | o | tt |
| ypdD | CDS | fused predicted PTS enzymes Hpr component -!- enzyme I component -!- enzyme IIA component | Cytoplasmic | COG1080;Phosphoenolpyruvate-protein kinase (PTS system EI component in bacteria) -!- COG1762;Phosphotransferase system mannitol/fructose-specific IIA domain (Ntr-type) | GO:0009401 phosphoenolpyruvate-dependent sugar phosphotransferase system | gg | tt |
| ypdE | CDS | predicted peptidase | Cytoplasmic | COG1363;Cellulase M and related proteins | | gg | t |
| ypdF | CDS | predicted peptidase | Cytoplasmic | COG0006;Xaa-Pro aminopeptidase | | g | tt |
| ypdG | CDS | predicted enzyme IIC component of PTS | Integral Membrane Protein | COG1299;Phosphotransferase system, fructose-specific IIC component | GO:0016052 carbohydrate | g | tt |
| ypdH | CDS | predicted enzyme IIB component of PTS | Periplasmic | COG1445;Phosphotransferase system fructose-specific component IIB | GO:0009401 phosphoenolpyruvate-dependent sugar phosphotransferase system | | |
| ypdI | CDS | predicted lipoprotein involved in colanic acid biosynthesis | Outer Membrane Lipoprotein | | | | |

| | | | | | | | |
|---|---|---|---|---|---|---|---|
| *ypdJ* | CDS | predicted protein | Cytoplasmic | | | | |
| *ypeA* | CDS | predicted acyltransferase with acyl-CoA N-acyltransferase domain | Cytoplasmic | | | gg | t |
| *ypeB* | CDS | predicted protein | Cytoplasmic | | | | |
| *ypeC* | CDS | conserved protein | Periplasmic | | | o | tt |
| *ypfE* | CDS | predicted carboxysome structural protein with predicted role in ethanol utilization | Cytoplasmic | | GO:0009310 amine catabolism | gg | t |
| *ypfG* | CDS | predicted protein | Periplasmic | | | g | tt |
| *ypfH* | CDS | predicted hydrolase | Cytoplasmic | COG0400;Predicted esterase | | | |
| *ypfI* | CDS | predicted hydrolase | Cytoplasmic | COG1444;Predicted P-loop ATPase fused to an acetyltransferase | | g | t |
| *ypfJ* | CDS | conserved protein | Membrane Anchored | COG2321;Predicted metalloprotease | | o | tt |
| *ypfN* | CDS | predicted protein | Membrane Anchored | | | g | tt |
| *yphA* | CDS | predicted inner membrane protein | Integral Membrane Protein | | | o | tt |
| *yphB* | CDS | conserved protein | Cytoplasmic | COG2017;Galactose mutarotase and related enzymes | | gg | tt |
| *yphC* | CDS | predicted oxidoreductase, Zn-dependent and NAD(P)-binding | Cytoplasmic | COG1063;Threonine dehydrogenase and related Zn-dependent dehydrogenases | | | |
| *yphD* | CDS | predicted sugar transporter subunit -!- membrane component of ABC superfamily | Integral Membrane Protein | COG1172;Ribose/xylose/arabinose/galactoside ABC-type transport systems, permease components | | o | tt |
| *yphE* | CDS | fused predicted sugar transporter subunits -!- ATP-binding components of ABC superfamily | Cytoplasmic | COG1129;ABC-type sugar transport system, ATPase component | | gg | tt |
| *yphF* | CDS | predicted sugar transporter subunit -!- periplasmic-binding component of ABC superfamily | Periplasmic | COG1879;ABC-type sugar transport system, periplasmic component | | o | tt |
| *yphG* | CDS | conserved protein | Cytoplasmic | | | | |
| *yphH* | CDS | predicted DNA-binding transcriptional regulator | Cytoplasmic | | GO:0006355 regulation of transcription, DNA-dependent | | |
| *ypjA* | CDS | adhesin-like autotransporter | Outer Membrane B-barrel protein | COG3468;Type V secretory pathway, adhesin AidA | | | |
| *ypjB* | CDS | predicted protein | Cytoplasmic | | | gg | o |
| *ypjC* | CDS | predicted protein | Cytoplasmic | | | | |
| *ypjD* | CDS | predicted inner membrane protein | Integral Membrane Protein | | | o | tt |
| *ypjF* | CDS | CP4-57 prophage; toxin of the YpjF-YfjZ toxin-antitoxin system | Cytoplasmic | | | gg | tt |
| *ypjJ* | CDS | predicted protein | Cytoplasmic | | | | |
| *ypjK* | CDS | CP4-57 prophage; predicted inner membrane protein | Integral Membrane Protein | | | | |
| *ypjL* | CDS | CP4-57 prophage; predicted inner membrane protein | Integral Membrane Protein | | | | |
| *ypjM* | CDS | CP4-57 prophage; predicted protein (pseudogene) | Cytoplasmic | | | | |
| *yqaA* | CDS | conserved inner membrane protein | Integral Membrane Protein | COG1238;Predicted membrane protein | | o | tt |
| *yqaB* | CDS | predicted hydrolase | Cytoplasmic | COG0637;Predicted phosphatase/phosphohexomutase | | gg | t |
| *yqaC* | CDS | conserved protein (pseudogene) | Cytoplasmic | COG0366;Glycosidases | | | |
| *yqaD* | CDS | conserved protein | Cytoplasmic | COG0366;Glycosidases | | g | o |
| *yqaE* | CDS | predicted membrane protein | Integral Membrane Protein | COG0401;Uncharacterized homolog of Blt101 | | o | tt |
| *yqcA* | CDS | predicted flavoprotein | Cytoplasmic | COG0716;Flavodoxins | | gg | tt |
| *yqcB* | CDS | tRNA pseudouridine synthase | Cytoplasmic | COG0564;Pseudouridylate synthases, 23S RNA-specific | | gg | t |
| *yqcC* | CDS | conserved protein | Cytoplasmic | COG3098;Uncharacterized protein conserved in bacteria | | gg | t |
| *yqcD* | CDS | conserved protein | Cytoplasmic | COG2904;Uncharacterized protein conserved in bacteria -!- COG0780;Enzyme related to GTP cyclohydrolase I | | gg | t |
| *yqcE* | CDS | predicted transporter | Integral Membrane Protein | COG0477;Permeases of the major facilitator superfamily | | o | tt |
| *yqeA* | CDS | predicted amino acid kinase | Cytoplasmic | COG0549;Carbamate kinase | | gg | t |
| *yqeB* | CDS | conserved protein with NAD(P)-binding Rossman fold | Cytoplasmic | COG1975;Xanthine and CO dehydrogenases maturation factor, XdhC/CoxF family | | gg | t |
| *yqeC* | CDS | conserved protein | Cytoplasmic | | | | |
| *yqeF* | CDS | predicted acyltransferase | Cytoplasmic | COG0183;Acetyl-CoA acetyltransferase | | | |
| *yqeG* | CDS | predicted transporter | Integral Membrane Protein | COG0814;Amino acid permeases | | o | tt |
| *yqeH* | CDS | conserved protein with bipartite regulator domain | Cytoplasmic | COG2771;DNA-binding HTH domain-containing proteins | | | |
| *yqeI* | CDS | predicted transcriptional regulator | Cytoplasmic | COG3710;DNA-binding winged-HTH domains | | | |
| *yqeJ* | CDS | predicted protein | Periplasmic | | | | |
| *yqeK* | CDS | predicted protein | Cytoplasmic | | | o | o |
| *yqfA* | CDS | predicted oxidoreductase, inner membrane subunit | Integral Membrane Protein | COG1272;Predicted membrane protein, hemolysin III homolog | | o | tt |
| *yqfB* | CDS | conserved protein | Cytoplasmic | COG3097;Uncharacterized protein conserved in bacteria | | gg | o |
| *yqfE* | CDS | conserved protein | Cytoplasmic | COG0583;Transcriptional regulator | | gg | t |
| *yqgA* | CDS | predicted inner membrane protein | Integral Membrane Protein | COG1811;Uncharacterized membrane protein, possible Na+ channel or pump | | g | tt |
| *yqgB* | CDS | predicted protein | Cytoplasmic | | | g | o |
| *yqgC* | CDS | predicted protein | Cytoplasmic | | | g | t |
| *yqgD* | CDS | predicted inner membrane protein | Integral Membrane Protein | | | o | tt |
| *yqgE* | CDS | predicted protein | Cytoplasmic | | | gg | o |
| *yqgF* | CDS | predicted Holliday junction resolvase | Cytoplasmic | COG0816;Predicted endonuclease involved in recombination (possible Holliday junction resolvase in Mycoplasmas and B. subtilis) | | gg | t |
| *yqhA* | CDS | conserved inner membrane protein | Integral Membrane Protein | COG2862;Predicted membrane protein | | o | tt |
| *yqhC* | CDS | predicted DNA-binding transcriptional regulator | Cytoplasmic | COG2207;AraC-type DNA-binding domain-containing proteins | GO:0006350 transcription | | |
| *yqhD* | CDS | alcohol dehydrogenase, NAD(P)-dependent | Cytoplasmic | COG1979;Uncharacterized oxidoreductases, Fe-dependent alcohol dehydrogenase family | | gg | t |
| *yqhG* | CDS | conserved protein | Periplasmic | | | | |
| *yqhH* | CDS | predicted outer membrane lipoprotein | Outer Membrane Lipoprotein | COG4238;Murein lipoprotein | | o | tt |
| *yqiA* | CDS | predicted esterase | Cytoplasmic | COG3150;Predicted esterase | | o | tt |
| *yqiB* | CDS | predicted dehydrogenase | Cytoplasmic | COG3151;Uncharacterized protein conserved in bacteria | | gg | t |
| *yqiC* | CDS | conserved protein | Cytoplasmic | | | | |
| *yqiG* | CDS | predicted outer membrane usher protein | Outer Membrane B-barrel protein | | | | |
| *yqiH* | CDS | predicted periplasmic pilin chaperone | Outer Membrane B-barrel protein | | GO:0006457 protein folding | | |
| *yqiI* | CDS | conserved protein | Periplasmic | | | | |
| *yqiJ* | CDS | predicted inner membrane protein | Integral Membrane Protein | | | o | tt |
| *yqiK* | CDS | conserved protein | Membrane Anchored | COG2268;Uncharacterized protein conserved in bacteria | | o | tt |
| *yqjA* | CDS | conserved inner membrane protein | Integral Membrane Protein | COG0586;Uncharacterized membrane-associated protein | | gg | tt |
| *yqjB* | CDS | conserved protein | Membrane Anchored | | | o | tt |
| *yqjC* | CDS | conserved protein | Periplasmic | | | | |
| *yqjD* | CDS | conserved protein | Membrane Anchored | COG4575;Uncharacterized conserved protein | | o | o |
| *yqjE* | CDS | conserved inner membrane protein | Integral Membrane Protein | COG5393;Predicted membrane protein | | o | tt |
| *yqjF* | CDS | predicted quinol oxidase subunit | Integral Membrane Protein | COG2259;Predicted membrane protein | | | |
| *yqjG* | CDS | predicted S-transferase | Cytoplasmic | COG0435;Predicted glutathione S-transferase | | gg | t |
| *yqjH* | CDS | predicted siderophore interacting protein | Cytoplasmic | COG2375;Siderophore-interacting protein | | gg | t |
| *yqjI* | CDS | predicted transcriptional regulator | Cytoplasmic | COG1695;Predicted transcriptional regulators | | g | o |
| *yqjK* | CDS | conserved protein | Membrane Anchored | | | o | tt |
| *yraH* | CDS | predicted fimbrial-like adhesin protein | Periplasmic | COG3539;P pilus assembly protein, pilin FimA | | o | tt |
| *yraI* | CDS | predicted periplasmic pilin chaperone | Periplasmic | COG3121;P pilus assembly protein, chaperone PapD | GO:0006457 protein folding | o | tt |
| *yraJ* | CDS | predicted outer membrane protein | Outer Membrane B-barrel protein | COG3188;P pilus assembly protein, porin PapC | | o | tt |
| *yraK* | CDS | predicted fimbrial-like adhesin protein | Periplasmic | COG3539;P pilus assembly protein, pilin FimA | | o | tt |
| *yraL* | CDS | predicted methyltransferase | Cytoplasmic | COG0313;Predicted methyltransferases | | gg | t |
| *yraM* | CDS | conserved protein | Membrane Lipoprotein | COG3107;Putative lipoprotein | | o | tt |
| *yraN* | CDS | conserved protein | Cytoplasmic | COG0792;Predicted endonuclease distantly related to archaeal Holliday junction resolvase | | gg | t |
| *yraO* | CDS | DnaA initiator-associating factor for replication initiation | Cytoplasmic | COG0279;Phosphoheptose isomerase | | gg | t |
| *yraP* | CDS | predicted protein | Outer Membrane Lipoprotein | COG2823;Predicted periplasmic or secreted lipoprotein | | o | tt |
| *yraQ* | CDS | predicted permease | Integral Membrane Protein | COG0701;Predicted permeases | | o | tt |
| *yraR* | CDS | predicted nucleoside-diphosphate-sugar epimerase | Cytoplasmic | COG0702;Predicted nucleoside-diphosphate-sugar epimerases | | | |
| *yrbA* | CDS | predicted DNA-binding transcriptional regulator | Cytoplasmic | COG5007;Predicted transcriptional regulator, BolA superfamily | GO:0006355 regulation of transcription, DNA-dependent | gg | o |
| *yrbB* | CDS | predicted protein | Cytoplasmic | | | | |
| *yrbC* | CDS | predicted ABC-type organic solvent transporter | Periplasmic | COG2854;ABC-type transport system involved in resistance to organic solvents, auxiliary component | | o | tt |
| *yrbD* | CDS | predicted ABC-type organic solvent transporter | Periplasmic | COG1463;ABC-type transport system involved in resistance to organic solvents, periplasmic component | | o | tt |
| *yrbE* | CDS | predicted toluene transporter subunit-!- membrane component of ABC superfamily | Integral Membrane Protein | COG0767;ABC-type transport system involved in resistance to organic solvents, permease component | | o | tt |
| *yrbF* | CDS | predicted toluene transporter subunit -!- ATP-binding component of ABC superfamily | Cytoplasmic | COG1127;ABC-type transport system involved in resistance to organic solvents, ATPase component | | gg | t |
| *yrbG* | CDS | predicted calcium/sodium:proton antiporter | Integral Membrane Protein | COG0530;Ca2+/Na+ antiporter | | o | tt |
| *yrbK* | CDS | conserved protein | Membrane Anchored | COG3117;Uncharacterized protein conserved in bacteria | | o | tt |
| *yrbL* | CDS | predicted protein | Cytoplasmic | | | gg | t |
| *yrdA* | CDS | conserved protein | Cytoplasmic | | | | |
| *yrdB* | CDS | conserved protein | Cytoplasmic | | | gg | o |
| *yrdC* | CDS | dsRNA-binding protein | Cytoplasmic | COG0009;Putative translation factor (SUA5) | | gg | t |
| *yrdD* | CDS | predicted DNA topoisomerase | Cytoplasmic | | | | |
| *yrfA* | CDS | predicted protein | Cytoplasmic | | | | |
| *yrfB* | CDS | conserved membrane protein | Membrane Anchored | | | o | tt |
| *yrfC* | CDS | predicted fimbrial assembly protein | Membrane Anchored | COG3166;Tfp pilus assembly protein PilN | | o | tt |
| *yrfD* | CDS | predicted pilus assembly protein | Cytoplasmic | | | | |
| *yrfF* | CDS | predicted inner membrane protein | Integral Membrane Protein | | | o | tt |
| *yrfG* | CDS | predicted hydrolase | Cytoplasmic | | | | |
| *yrhA* | CDS | conserved protein | Cytoplasmic | | | | |
| *yrhB* | CDS | predicted protein | Cytoplasmic | | | gg | o |
| *yrhC* | CDS | predicted protein fragment (pseudogene) | Cytoplasmic | | | | |
| *ysaB* | CDS | predicted protein | Outer Membrane Lipoprotein | | | o | tt |
| *ysdC* | CDS | predicted protein (pseudogene) | Cytoplasmic | | | | |

| | | | | | | | |
|---|---|---|---|---|---|---|---|
| *ysgA* | CDS | predicted hydrolase | Cytoplasmic | | | | |
| *ytfA* | CDS | predicted transcriptional regulator | Cytoplasmic | COG1309;Transcriptional regulator | | g | o |
| *ytfB* | CDS | predicted cell envelope opacity-associated protein | Cytoplasmic | COG3061;Cell envelope opacity-associated protein A | | | |
| *ytfE* | CDS | predicted regulator of cell morphogenesis and cell wall metabolism | Cytoplasmic | COG2846;Regulator of cell morphogenesis and NO signaling | | gg | t |
| *ytfF* | CDS | predicted inner membrane protein | Integral Membrane Protein | COG0697;Permeases of the drug/metabolite transporter (DMT) superfamily | | o | tt |
| *ytfG* | CDS | NAD(P)H:quinone oxidoreductase | Cytoplasmic | COG0702;Predicted nucleoside-diphosphate-sugar epimerases | | gg | o |
| *ytfH* | CDS | predicted transcriptional regulator | Cytoplasmic | | GO:0006355 regulation of transcription, DNA-dependent | | |
| *ytfI* | CDS | predicted protein | Cytoplasmic | | | | |
| *ytfJ* | CDS | predicted transcriptional regulator | Periplasmic | COG3054;Predicted transcriptional regulator | | o | tt |
| *ytfK* | CDS | conserved protein | Cytoplasmic | | | | |
| *ytfL* | CDS | predicted inner membrane protein | Integral Membrane Protein | COG1253;Hemolysins and related proteins containing CBS domains | | o | tt |
| *ytfM* | CDS | predicted outer membrane protein and surface antigen | Outer Membrane B-barrel protein | COG0729;Outer membrane protein | | o | tt |
| *ytfN* | CDS | conserved protein | Membrane Anchored | COG2911;Uncharacterized protein conserved in bacteria | | o | tt |
| *ytfP* | CDS | conserved protein | Cytoplasmic | COG2105;Uncharacterized conserved protein | | o | tt |
| *ytfQ* | CDS | predicted sugar transporter subunit -!- periplasmic-binding component of ABC superfamily | Periplasmic | COG1879;ABC-type sugar transport system, periplasmic component | | o | tt |
| *ytfR* | CDS | predicted sugar transporter subunit -! ATP-binding component of ABC superfamily | Cytoplasmic | | | | |
| *ytfT* | CDS | predicted sugar transporter subunit -!- membrane component of ABC superfamily | Integral Membrane Protein | COG1172;Ribose/xylose/arabinose/galactoside ABC-type transport systems, permease components | | | |
| *ytjA* | CDS | predicted protein | Integral Membrane Protein | | | | |
| *ytjB* | CDS | conserved protein | Membrane Anchored | COG3726;Uncharacterized membrane protein affecting hemolysin expression | | o | tt |
| *ytjC* | CDS | phosphoglyceromutase 2, co-factor independent | Membrane Anchored | COG0406;Fructose-2,6-bisphosphatase | GO:0006096 glycolysis -!- GO:0006094 gluconeogenesis | gg | t |
| *yzcX* | CDS | predicted protein | Cytoplasmic | | | | |
| *yzfA* | CDS | predicted protein | Cytoplasmic | | | gg | t |
| *yzgL* | CDS | predicted protein | Integral Membrane Protein | | | o | tt |
| *zapA* | CDS | protein that localizes to the cytokinetic ring | Cytoplasmic | COG3027;Uncharacterized protein conserved in bacteria | | gg | tt |
| *zipA* | CDS | cell division protein involved in Z ring assembly | Periplasmic | COG3115;Cell division protein | | o | tt |
| *zitB* | CDS | zinc efflux system | Integral Membrane Protein | COG1230;Co/Zn/Cd efflux system component | | o | tt |
| *zntA* | CDS | zinc, cobalt and lead efflux system | Integral Membrane Protein | COG2217;Cation transport ATPase | GO:0015986 ATP synthesis coupled proton transport -!- GO:0006805 xenobiotic metabolism | o | tt |
| *zntR* | CDS | DNA-binding transcriptional activator | Cytoplasmic | COG0789;Predicted transcriptional regulators | GO:0006350 transcription -!- GO:0006805 xenobiotic metabolism | gg | o |
| *znuA* | CDS | zinc transporter subunit -!- periplasmic-binding component of ABC superfamily | Periplasmic | | | | |
| *znuB* | CDS | zinc transporter subunit -!- membrane component of ABC superfamily | Integral Membrane Protein | COG1108;ABC-type Mn2+/Zn2+ transport systems, permease components | | o | tt |
| *znuC* | CDS | zinc transporter subunit -!- ATP-binding component of ABC superfamily | Cytoplasmic | COG1121;ABC-type Mn/Zn transport systems, ATPase component | | gg | tt |
| *zraP* | CDS | Zn-binding periplasmic protein | Periplasmic | COG3678;P pilus assembly/Cpx signaling pathway, periplasmic inhibitor/zinc-resistance associated protein | | | |
| *zraR* | CDS | fused DNA-binding response regulator in two-component regulatory system with ZraS | Cytoplasmic | COG2204;Response regulator containing CheY-like receiver, AAA-type ATPase, and DNA-binding domains | GO:0006113 fermentation -!- GO:0006350 transcription | o | t |
| *zraS* | CDS | sensory histidine kinase in two-component regulatory system with ZraR | Integral Membrane Protein | COG0642;Signal transduction histidine kinase | GO:0006113 fermentation -!- GO:0006464 protein modification | o | tt |
| *zupT* | CDS | predicted dioxygenase | Cytoplasmic | COG3384;Uncharacterized conserved protein | | g | tt |
| *zur* | CDS | DNA-binding transcriptional activator, Zn(II)-binding | Cytoplasmic | | GO:0006350 transcription | | |
| *zwf* | CDS | glucose-6-phosphate dehydrogenase | Cytoplasmic | COG0364;Glucose-6-phosphate 1-dehydrogenase | GO:0016052 carbohydrate catabolism -!- GO:0009051 pentose-phosphate shunt, oxidative branch | gg | t |

**Table S2. Toxicity of essential genes.** Toxicity severity was assessed by survivor frequency values. We tested the essential genes causing both severe growth inhibition and no GFP production in Table S1. Results are shown in order of toxicity severity.

| Gene | Feature | Product | Location | Function | Process | Survivor frequency (ppm) |
|---|---|---|---|---|---|---|
| *dnaX* | CDS | DNA polymerase III/DNA elongation factor III, tau and gamma subunits | Cytoplasmic | COG2812;DNA polymerase III, gamma/tau subunits | GO:0006261 DNA dependent DNA replication | 0 |
| *ftsK* | CDS | DNA-binding membrane protein required for chromosome resolution and partitioning | Integral Membrane Protein | COG1674;DNA segregation ATPase FtsK/SpoIIIE and related proteins | GO:0009432 SOS response | 0 |
| *mrdA* | CDS | transpeptidase involved in peptidoglycan synthesis (penicillin-binding protein 2) | Membrane Anchored | COG0768;Cell division protein FtsI/penicillin-binding protein 2 | GO:0009252 peptidoglycan biosynthesis -!- GO:0042493 response to drug | 0 |
| *mrdB* | CDS | cell wall shape-determining protein | Integral Membrane Protein | COG0772;Bacterial cell division membrane protein | | 0 |
| *mukB* | CDS | fused chromosome partitioning proteins | Cytoplasmic | COG3096;Uncharacterized protein involved in chromosome partitioning | | 0 |
| *rpsD* | CDS | 30S ribosomal subunit protein S4 | Cytoplasmic | COG0522;Ribosomal protein S4 and related proteins | GO:0006412 protein biosynthesis | 0 |
| *lexA* | CDS | DNA-binding transcriptional repressor | Cytoplasmic | COG1974;SOS-response transcriptional repressors (RecA-mediated autopeptidases) | GO:0006281 DNA repair -!- GO:0006350 transcription -!- GO:0009432 SOS response | 0.59 |
| *pheT* | CDS | phenylalanine tRNA synthetase, beta subunit | Cytoplasmic | COG0073;EMAP domain -!- COG0072;Phenylalanyl-tRNA synthetase beta subunit | GO:0006418 amino acid activation | 1.4 |
| *prfA* | CDS | peptide chain release factor RF-1 | Cytoplasmic | COG0216;Protein chain release factor A | GO:0006412 protein biosynthesis | 1.77 |
| *lepB* | CDS | leader peptidase (signal peptidase I) | Integral Membrane Protein | COG0681;Signal peptidase I | | 1.8 |
| *polA* | CDS | fused DNA polymerase I 5'->3' exonuclease -!- 3'->5' polymerase -!- 3'->5' exonuclease | Cytoplasmic | COG0258;5'-3' exonuclease (including N-terminal domain of PolI) -!- COG0749;DNA polymerase I - 3'-5' exonuclease and polymerase domains | GO:0006261 DNA dependent DNA replication | 3.98 |
| *leuS* | CDS | leucyl-tRNA synthetase | Cytoplasmic | COG0495;Leucyl-tRNA synthetase | GO:0006418 amino acid activation | 5.57 |
| *ftsI* | CDS | transpeptidase involved in septal peptidoglycan synthesis (penicillin-binding protein 3) | Membrane Anchored | COG0768;Cell division protein FtsI/penicillin-binding protein 2 | GO:0009252 peptidoglycan biosynthesis -!- GO:0042493 response to drug | 8.13 |
| *yaeL* | CDS | zinc metallopeptidase | Integral Membrane Protein | COG0750;Predicted membrane-associated Zn-dependent proteases 1 | | 8.26 |
| *secA* | CDS | preprotein translocase subunit, ATPase that targets protein precursors to the SecYE core | Cytoplasmic | COG0653;Preprotein translocase subunit SecA (ATPase, RNA helicase) | | 8.84 |
| *parC* | CDS | DNA topoisomerase IV, subunit A | Cytoplasmic | COG0188;Type IIA topoisomerase (DNA gyrase/topo II, topoisomerase IV), A subunit | GO:0006261 DNA dependent DNA replication | 11.16 |
| *ftsQ* | CDS | membrane anchored protein involved in growth of wall at septum | Membrane Anchored | COG1589;Cell division septal protein | | 11.51 |
| *rplP* | CDS | 50S ribosomal subunit protein L16 | Cytoplasmic | COG0197;Ribosomal protein L16/L10E | GO:0006412 protein biosynthesis | 16.54 |
| *dnaC* | CDS | DNA biosynthesis protein | Cytoplasmic | COG1484;DNA replication protein | GO:0006261 DNA dependent DNA replication | 19.64 |
| *mviN* | CDS | predicted inner membrane protein | Integral Membrane Protein | COG0728;Uncharacterized membrane protein, putative virulence factor | | 20.33 |
| *lolE* | CDS | outer membrane-specific lipoprotein transporter subunit -!- membrane component of ABC superfamily | Integral Membrane Protein | COG4591;ABC-type transport system, involved in lipoprotein release, permease component | | 20.34 |
| *yejM* | CDS | predicted hydrolase, inner membrane | Integral Membrane Protein | COG3083;Predicted hydrolase of alkaline phosphatase superfamily | | 21.01 |
| *tyrS* | CDS | tyrosyl-tRNA synthetase | Cytoplasmic | COG0162;Tyrosyl-tRNA synthetase | GO:0006418 amino acid activation | 22.51 |
| *dnaA* | CDS | chromosomal replication initiator protein DnaA, DNA-binding transcriptional dual regulator | Cytoplasmic | COG0593;ATPase involved in DNA replication initiation | GO:0006261 DNA dependent DNA replication | 23.89 |
| *rplD* | CDS | 50S ribosomal subunit protein L4 | Cytoplasmic | COG0088;Ribosomal protein L4 | GO:0006412 protein biosynthesis -!- GO:0006355 regulation of transcription, DNA-dependent -!- GO:0009386 translational attenuation | 24.27 |
| *rpoD* | CDS | RNA polymerase, sigma 70 (sigma D) factor | Cytoplasmic | COG0568;DNA-directed RNA polymerase, sigma subunit (sigma70/sigma32) | GO:0006350 transcription | 26.09 |
| *ftsH* | CDS | protease, ATP-dependent zinc-metallo | Integral Membrane Protein | COG0465;ATP-dependent Zn proteases | | 27.62 |
| *secY* | CDS | preprotein translocase membrane subunit | Integral Membrane Protein | COG0201;Preprotein translocase subunit SecY | | 28.88 |
| *hemH* | CDS | ferrochelatase | Cytoplasmic | COG0276;Protoheme ferro-lyase (ferrochelatase) | | 30.85 |
| *rpoH* | CDS | RNA polymerase, sigma 32 (sigma H) factor | Cytoplasmic | COG0568;DNA-directed RNA polymerase, sigma subunit (sigma70/sigma32) | GO:0006350 transcription -!- GO:0009266 response to temperature | 34.94 |

| | | | | | | |
|---|---|---|---|---|---|---|
| *lgt* | CDS | phosphatidylglycerol-prolipoprotein diacylglyceryl transferase | Integral Membrane Protein | COG0682;Prolipoprotein diacylglyceryltransferase | GO:0008654 phospholipid biosynthesis -!- GO:0006464 protein modification | 39.68 |
| *ftsL* | CDS | membrane bound cell division protein at septum containing leucine zipper motif | Membrane Anchored | COG3116;Cell division protein | | 40.4 |
| *rplS* | CDS | 50S ribosomal subunit protein L19 | Cytoplasmic | COG0335;Ribosomal protein L19 | GO:0006412 protein biosynthesis | 40.62 |
| *ribF* | CDS | bifunctional riboflavin kinase -!- FAD synthetase | Cytoplasmic | COG0196;FAD synthase | | 43.72 |
| *rpsB* | CDS | 30S ribosomal subunit protein S2 | Cytoplasmic | COG0052;Ribosomal protein S2 | GO:0006412 protein biosynthesis | 51.92 |
| *rpsC* | CDS | 30S ribosomal subunit protein S3 | Cytoplasmic | COG0092;Ribosomal protein S3 | GO:0006412 protein biosynthesis | 55.12 |
| *ftsX* | CDS | predicted transporter subunit -!- membrane component of ABC superfamily | Integral Membrane Protein | COG2177;Cell division protein | | 56.57 |
| *yhbN* | CDS | predicted transporter subunit -!- periplasmic-binding component of ABC superfamily | Periplasmic | COG1934;Uncharacterized protein conserved in bacteria | | 58.4 |
| *ygjE* | CDS | predicted tartrate:succinate antiporter | Integral Membrane Protein | COG0471;Di- and tricarboxylate transporters | | 64.97 |
| *ileS* | CDS | isoleucyl-tRNA synthetase | Cytoplasmic | COG0060;Isoleucyl-tRNA synthetase | GO:0006418 amino acid activation | 67 |
| *rplB* | CDS | 50S ribosomal subunit protein L2 | Cytoplasmic | COG0090;Ribosomal protein L2 | GO:0006412 protein biosynthesis | 71.05 |
| *rpsA* | CDS | 30S ribosomal subunit protein S1 | Cytoplasmic | COG0539;Ribosomal protein S1 | GO:0006412 protein biosynthesis | 75.1 |
| *lpxB* | CDS | tetraacyldisaccharide-1-P synthase | Cytoplasmic | COG0763;Lipid A disaccharide synthetase | GO:0009245 lipid A biosynthesis | 86.56 |
| *pgsA* | CDS | phosphatidylglycerophosphate synthetase | Integral Membrane Protein | COG0558;Phosphatidylglycerophosphate synthase | GO:0008654 phospholipid biosynthesis | 87.01 |
| *tilS* | CDS | tRNA(Ile)-lysidine synthetase | Cytoplasmic | COG0037;Predicted ATPase of the PP-loop superfamily implicated in cell cycle control | | 89.4 |
| *msbA* | CDS | fused lipid transporter subunits -!- membrane component and ATP-binding component of ABC | Integral Membrane Protein | COG1132;ABC-type multidrug transport system, ATPase and permease components | | 94.48 |
| *ftsW* | CDS | integral membrane protein involved in stabilising FstZ ring during cell division | Integral Membrane Protein | COG0772;Bacterial cell division membrane protein | | 114.92 |
| *rpsS* | CDS | 30S ribosomal subunit protein S19 | Cytoplasmic | COG0185;Ribosomal protein S19 | GO:0006412 protein biosynthesis | 118.51 |
| *degS* | CDS | serine endoprotease, periplasmic | Membrane Anchored | COG0265;Trypsin-like serine proteases, typically periplasmic, contain C-terminal PDZ domain | GO:0006508 proteolysis and peptidolysis | 120.19 |
| *infB* | CDS | fused protein chain initiation factor 2, IF2 | Cytoplasmic | COG0532;Translation initiation factor 2 (IF-2; GTPase) | GO:0006412 protein biosynthesis | 130.69 |
| *mreC* | CDS | cell wall structural complex MreBCD transmembrane component MreC | Membrane Anchored | COG1792;Cell shape-determining protein | GO:0009252 peptidoglycan biosynthesis | 135 |
| *trmU* | CDS | tRNA (5-methylaminomethyl-2-thiouridylate)-methyltransferase | Cytoplasmic | | GO:0009451 RNA modification | 158.46 |
| *yfcC* | CDS | predicted inner membrane protein | Integral Membrane Protein | COG1288;Predicted membrane protein | | 159 |
| *dnaB* | CDS | replicative DNA helicase | Cytoplasmic | COG0305;Replicative DNA helicase | GO:0006261 DNA dependent DNA replication | 161.14 |
| *lpxC* | CDS | UDP-3-O-acyl N-acetylglucosamine deacetylase | Cytoplasmic | COG0774;UDP-3-O-acyl-N-acetylglucosamine deacetylase | GO:0009245 lipid A biosynthesis | 176.47 |
| *zipA* | CDS | cell division protein involved in Z ring assembly | Periplasmic | COG3115;Cell division protein | | 183.23 |
| *yjgP* | CDS | conserved inner membrane protein | Integral Membrane Protein | COG0795;Predicted permeases | | 191.73 |
| *yrbK* | CDS | conserved protein | Membrane Anchored | COG3117;Uncharacterized protein conserved in bacteria | | 191.78 |
| *lspA* | CDS | prolipoprotein signal peptidase (signal peptidase II) | Integral Membrane Protein | COG0597;Lipoprotein signal peptidase | | 219 |
| *murG* | CDS | N-acetylglucosaminyl transferase | Membrane Associated | COG0707;UDP-N-acetylglucosamine:LPS N-acetylglucosamine transferase | GO:0009252 peptidoglycan biosynthesis | 219.69 |
| *yrfF* | CDS | predicted inner membrane protein | Integral Membrane Protein | | | 237.18 |
| *proS* | CDS | prolyl-tRNA synthetase | Cytoplasmic | COG0442;Prolyl-tRNA synthetase | GO:0006418 amino acid activation | 266.67 |
| *lolA* | CDS | chaperone for lipoproteins | Periplasmic | COG2834;Outer membrane lipoprotein-sorting protein | | 271.54 |
| *mreD* | CDS | cell wall structural complex MreBCD transmembrane component MreD | Integral Membrane Protein | COG2891;Cell shape-determining protein | GO:0009252 peptidoglycan biosynthesis | 282.95 |
| *plsB* | CDS | glycerol-3-phosphate O-acyltransferase | Cytoplasmic | | GO:0008654 phospholipid biosynthesis | 290.23 |
| *dxs* | CDS | 1-deoxyxylulose-5-phosphate synthase, thiamine-requiring, FAD-requiring | Cytoplasmic | COG1154;Deoxyxylulose-5-phosphate synthase | GO:0008615 pyridoxine biosynthesis -!- GO:0009228 thiamin biosynthesis | 291.14 |

| | | | | | | |
|---|---|---|---|---|---|---|
| *ssb* | CDS | Single-stranded DNA-binding protein | Cytoplasmic | COG0629;Single-stranded DNA-binding protein | GO:0006310 DNA recombination -!- GO:0009432 SOS response | 291.14 |
| *frr* | CDS | ribosome recycling factor | Cytoplasmic | COG0233;Ribosome recycling factor | GO:0006412 protein biosynthesis -!- GO:0009386 translational attenuation | 342.39 |
| *mraY* | CDS | phospho-N-acetylmuramoyl-pentapeptide transferase | Integral Membrane Protein | COG0472;UDP-N-acetylmuramyl pentapeptide phosphotransferase/UDP-N-acetylglucosamine-1-phosp hate transferase | GO:0009252 peptidoglycan biosynthesis | 345.24 |
| *rho* | CDS | transcription termination factor | Cytoplasmic | COG1158;Transcription termination factor | GO:0006350 transcription | 355.88 |
| *accB* | CDS | acetyl CoA carboxylase, BCCP subunit | Cytoplasmic | COG0511;Biotin carboxyl carrier protein | GO:0042966 biotin carboxyl carrier protein biosynthesis | 361.49 |
| *argS* | CDS | arginyl-tRNA synthetase | Cytoplasmic | COG0018;Arginyl-tRNA synthetase | GO:0006418 amino acid activation | 393.73 |
| *ftsE* | CDS | predicted transporter subunit -!- ATP-binding component of ABC | Cytoplasmic | COG2884;Predicted ATPase involved in cell division | | 396.37 |
| *secD* | CDS | SecYEG protein translocase auxillary subunit | Integral Membrane Protein | COG0342;Preprotein translocase subunit SecD | | 406.78 |
| *yidC* | CDS | cytoplasmic insertase into membrane protein, Sec system | Integral Membrane Protein | COG0706;Preprotein translocase subunit YidC | | 465.62 |
| *grpE* | CDS | heat shock protein | Cytoplasmic | COG0576;Molecular chaperone GrpE (heat shock protein) | GO:0006457 protein folding | 471.07 |
| *psd* | CDS | phosphatidylserine decarboxylase | Cytoplasmic | COG0688;Phosphatidylserine decarboxylase | GO:0008654 phospholipid biosynthesis | 552.24 |
| *ftsY* | CDS | fused Signal Recognition Particle (SRP) receptor | Cytoplasmic | COG0552;Signal recognition particle GTPase | | 700.89 |
| *groL* | CDS | Cpn60 chaperonin GroEL, large subunit of GroESL | Cytoplasmic | COG0459;Chaperonin GroEL (HSP60 family) | GO:0006457 protein folding | 715.35 |
| *lnt* | CDS | apolipoprotein N-acyltransferase | Integral Membrane Protein | COG0815;Apolipoprotein N-acyltransferase | GO:0042158 lipoprotein biosynthesis | 783.51 |
| *rplQ* | CDS | 50S ribosomal subunit protein L17 | Cytoplasmic | COG0203;Ribosomal protein L17 | GO:0006412 protein biosynthesis | 828.27 |
| *secE* | CDS | preprotein translocase membrane subunit | Integral Membrane Protein | COG0690;Preprotein translocase subunit SecE | | 852.43 |
| *hemA* | CDS | glutamyl tRNA reductase | Cytoplasmic | COG0373;Glutamyl-tRNA reductase | GO:0042493 response to drug | 863.95 |
| *rpsM* | CDS | 30S ribosomal subunit protein S13 | Cytoplasmic | COG0099;Ribosomal protein S13 | GO:0006412 protein biosynthesis | 906.63 |
| *accC* | CDS | acetyl-CoA carboxylase, biotin carboxylase subunit | Cytoplasmic | COG0439;Biotin carboxylase | | 977.62 |
| *plsC* | CDS | 1-acyl-sn-glycerol-3-phosphate acyltransferase | Membrane Anchored | COG0204;1-acyl-sn-glycerol-3-phosphate acyltransferase | GO:0008654 phospholipid biosynthesis | 983.05 |
| *fmt* | CDS | 10-formyltetrahydrofolate:L-methionyl-tRNA(fMet) N-formyltransferase | Cytoplasmic | COG0223;Methionyl-tRNA formyltransferase | GO:0009451 RNA modification | 1044.25 |
| *ubiA* | CDS | p-hydroxybenzoate octaprenyltransferase | Integral Membrane Protein | COG0382;4-hydroxybenzoate polyprenyltransferase and related prenyltransferases | GO:0009060 aerobic respiration | 1087.98 |
| *ftsB* | CDS | cell division protein | Membrane Anchored | COG2919;Septum formation initiator | | 1234.03 |
| *nusG* | CDS | transcription termination factor | Cytoplasmic | COG0250;Transcription antiterminator | GO:0006350 transcription | 1331.23 |
| *secF* | CDS | SecYEG protein translocase auxillary subunit | Integral Membrane Protein | COG0341;Preprotein translocase subunit SecF | | 1355.12 |
| *rlpB* | CDS | minor lipoprotein | Membrane Lipoprotein | COG2980;Rare lipoprotein B | GO:0042158 lipoprotein biosynthesis | 1561.64 |
| *rpoC* | CDS | RNA polymerase, beta prime subunit | Cytoplasmic | COG0086;DNA-directed RNA polymerase, beta' subunit/160 kD subunit | GO:0006350 transcription | 1804.79 |
| *rpsG* | CDS | 30S ribosomal subunit protein S7 | Cytoplasmic | COG0049;Ribosomal protein S7 | GO:0006412 protein biosynthesis | 2018.29 |
| *glyS* | CDS | glycine tRNA synthetase, beta subunit | Cytoplasmic | COG0751;Glycyl-tRNA synthetase, beta subunit | GO:0006418 amino acid activation | 2035.76 |
| *rpsH* | CDS | 30S ribosomal subunit protein S8 | Cytoplasmic | COG0096;Ribosomal protein S8 | GO:0006412 protein biosynthesis -!- GO:0009386 translational attenuation | 3325.3 |
| *rplN* | CDS | 50S ribosomal subunit protein L14 | Cytoplasmic | COG0093;Ribosomal protein L14 | GO:0006412 protein biosynthesis | 4195.98 |
| *dnaE* | CDS | DNA polymerase III alpha subunit | Cytoplasmic | COG0587;DNA polymerase III, alpha subunit | GO:0006261 DNA dependent DNA replication | 5206.75 |
| *holB* | CDS | DNA polymerase III, delta prime subunit | Cytoplasmic | COG0470;ATPase involved in DNA replication | GO:0006261 DNA dependent DNA replication | 5900.2 |
| *rpsR* | CDS | 30S ribosomal subunit protein S18 | Cytoplasmic | COG0238;Ribosomal protein S18 | GO:0006412 protein biosynthesis | 7792.75 |
| *rne* | CDS | fused ribonucleaseE endoribonuclease -!- scaffold for formation of degradosome | Cytoplasmic | COG1530;Ribonucleases G and E | GO:0006401 RNA catabolism -!- GO:0006401 RNA catabolism | 9038.83 |
| *hemG* | CDS | protoporphyrin oxidase, flavoprotein | Cytoplasmic | COG4635;Flavodoxin | | 9679.35 |

| | | | | | | |
|---|---|---|---|---|---|---|
| *kdtA* | CDS | 3-deoxy-D-manno-octulosonic-acid transferase (KDO transferase) | Cytoplasmic | COG1519;3-deoxy-D-manno-octulosonic-acid transferase | GO:0009245 lipid A biosynthesis -!- GO:0009244 lipopolysaccharide core region biosynthesis -!- GO:0009248 K antigen biosynthesis | 10533.18 |
| *rplJ* | CDS | 50S ribosomal subunit protein L10 | Cytoplasmic | COG0244;Ribosomal protein L10 | GO:0006412 protein biosynthesis | 13807.38 |
| *mreB* | CDS | cell wall structural complex MreBCD, actin-like component MreB | Cytoplasmic | COG1077;Actin-like ATPase involved in cell morphogenesis | GO:0042493 response to drug | 18005.26 |
| *ftsA* | CDS | ATP-binding cell division protein involved in recruitment of FtsK to Z | Cytoplasmic | COG0849;Actin-like ATPase involved in cell division | | 26525.2 |
| *kdsC* | CDS | 3-deoxy-D-manno-octulosonate 8-phosphate phosphatase | Periplasmic | COG1778;Low specificity phosphatase (HAD superfamily) | | |
| *lpxH* | CDS | UDP-2,3-diacylglucosamine pyrophosphatase | Cytoplasmic | COG2908;Uncharacterized protein conserved in bacteria | GO:0009245 lipid A biosynthesis | |
| *spoT* | CDS | bifunctional (p)ppGpp synthetase II -!- guanosine-3',5'-bis pyrophosphate 3'-pyrophosphohydrolase | Cytoplasmic | COG0317;Guanosine polyphosphate pyrophosphohydrolases/synthetases | GO:0015949 nucleobase, nucleoside and nucleotide interconversion -!- GO:0042594 response to starvation | |
| *ftsN* | CDS | essential cell division protein | Membrane Anchored | COG3087;Cell division protein | | |
| *murA* | CDS | UDP-N-acetylglucosamine 1-carboxyvinyltransferase | Cytoplasmic | COG0766;UDP-N-acetylglucosamine enolpyruvyl transferase | GO:0009252 peptidoglycan biosynthesis | |

Table S3. Toxicity of aminoacyl-tRNA synthetase genes. Genes that exhibit the toxic-overexpression phenotype are highlighted. *tyrS* is highlighted in green. Essential genes are indicated as "e" in the "Essentiality" column.

| Gene | Feature | Essentiality | Product | Location | Function | GFP fluorescence | IPTG growth | Survivor frequency (ppm) |
|---|---|---|---|---|---|---|---|---|
| *pheT* | CDS | e | phenylalanine tRNA synthetase, beta subunit | Cytoplasmic | COG0073;EMAP domain -!- COG0072;Phenylalanyl-tRNA synthetase beta subunit | o | tt | 1.4 |
| *leuS* | CDS | e | leucyl-tRNA synthetase | Cytoplasmic | COG0495;Leucyl-tRNA synthetase | o | tt | 5.6 |
| *tyrS* | CDS | e | tyrosyl-tRNA synthetase | Cytoplasmic | COG0162;Tyrosyl-tRNA | o | tt | 22.5 |
| *ileS* | CDS | e | isoleucyl-tRNA synthetase | Cytoplasmic | COG0060;Isoleucyl-tRNA | o | tt | 67 |
| *tilS* | CDS | e | tRNA(Ile)-lysidine synthetase | Cytoplasmic | COG0037;Predicted ATPase of the PP-loop superfamily implicated in cell cycle control | o | tt | 89.4 |
| *proS* | CDS | e | prolyl-tRNA synthetase | Cytoplasmic | COG0442;Prolyl-tRNA synthetase | o | tt | 266.7 |
| *argS* | CDS | e | arginyl-tRNA synthetase | Cytoplasmic | COG0018;Arginyl-tRNA synthetase | o | tt | 393.7 |
| *glyS* | CDS | e | glycine tRNA synthetase, beta subunit | Cytoplasmic | COG0751;Glycyl-tRNA synthetase, beta subunit | o | tt | 2035.8 |
| *poxA* | CDS | | predicted lysyl-tRNA synthetase | Cytoplasmic | COG2269;Truncated, possibly inactive, lysyl-tRNA synthetase (class II) | o | tt | |
| *alaS* | CDS | e | alanyl-tRNA synthetase | Cytoplasmic | COG0013;Alanyl-tRNA synthetase | gg | tt | |
| *cysS* | CDS | e | cysteinyl-tRNA synthetase | Cytoplasmic | COG0215;Cysteinyl-tRNA | gg | tt | |
| *glnS* | CDS | e | glutamyl-tRNA synthetase | Cytoplasmic | COG0008;Glutamyl- and glutaminyl-tRNA synthetases | gg | tt | |
| *gltX* | CDS | e | glutamyl-tRNA synthetase | Cytoplasmic | COG0008;Glutamyl- and glutaminyl-tRNA synthetases | gg | tt | |
| *serS* | CDS | e | seryl-tRNA synthetase, also charges selenocysteinyl-tRNA with | Cytoplasmic | COG0172;Seryl-tRNA synthetase | gg | tt | |
| *trpS* | CDS | e | tryptophanyl-tRNA synthetase | Cytoplasmic | COG0180;Tryptophanyl-tRNA synthetase | gg | tt | |
| *valS* | CDS | e | valyl-tRNA synthetase | Cytoplasmic | COG0525;Valyl-tRNA synthetase | gg | tt | |
| *asnS* | CDS | e | asparaginyl tRNA synthetase | Cytoplasmic | COG0017;Aspartyl/asparaginyl-tRNA synthetases | gg | t | |
| *aspS* | CDS | e | aspartyl-tRNA synthetase | Cytoplasmic | COG0173;Aspartyl-tRNA | gg | t | |
| *thrS* | CDS | e | threonyl-tRNA synthetase | Cytoplasmic | COG0441;Threonyl-tRNA | gg | t | |
| *lysS* | CDS | | lysine tRNA synthetase, constitutive | Cytoplasmic | COG1190;Lysyl-tRNA synthetase (class II) | gg | t | |
| *lysU* | CDS | | lysine tRNA synthetase, inducible | Cytoplasmic | COG1190;Lysyl-tRNA synthetase (class II) | gg | t | |
| *glyQ* | CDS | e | glycine tRNA synthetase, alpha subunit | Cytoplasmic | COG0752;Glycyl-tRNA synthetase, alpha subunit | g | t | |
| *metG* | CDS | e | methionyl-tRNA synthetase | Cytoplasmic | COG0143;Methionyl-tRNA synthetase -!- COG0073;EMAP | g | t | |
| *hisS* | CDS | e | histidyl tRNA synthetase | Cytoplasmic | COG0124;Histidyl-tRNA | gg | o | |
| *pheS* | CDS | e | phenylalanine tRNA synthetase, alpha subunit | Cytoplasmic | COG0016;Phenylalanyl-tRNA synthetase alpha subunit | | | |
| *yadB* | CDS | | glutamyl-Q tRNA(Asp) synthetase | Cytoplasmic | COG0008;Glutamyl- and glutaminyl-tRNA synthetases | | | |

Table S4. Genomic mutations found in IPTG-resistant mutants. Mutations found in P14R and P30R genomes are shown.

| position | P0S | P14R | position | P0S | P30R |
|---|---|---|---|---|---|
| | | | 112795 | GTTTTTTTTT | GTGGTTTTTTTTTTTTT (upstream) |
| 1737060 | AAC | AAAACAC (frame shift) *chbC* | 1737060 | AAC | AAAACAC,AACCAC (frame shift, insertion) *chbC* |
